# Systems Age: A single blood methylation test to quantify aging heterogeneity across 11 physiological systems

**DOI:** 10.1101/2023.07.13.548904

**Authors:** Raghav Sehgal, Yaroslav Markov, Chenxi Qin, Margarita Meer, Courtney Hadley, Aladdin H. Shadyab, Ramon Casanova, JoAnn E. Manson, Parveen Bhatti, Eileen M. Crimmins, Sara Hagg, Themistocles L. Assimes, Eric A. Whitsel, Albert T. Higgins-Chen, Morgan Levine

## Abstract

Individuals, organs, tissues, and cells age in diverse ways throughout the lifespan. Epigenetic clocks attempt to quantify differential aging between individuals, but they typically summarize aging as a single measure, ignoring within-person heterogeneity. Our aim was to develop novel systems-based methylation clocks that, when assessed in blood, capture aging in distinct physiological systems. We combined supervised and unsupervised machine learning methods to link DNA methylation, system-specific clinical chemistry and functional measures, and mortality risk. This yielded a panel of 11 system-specific scores– Heart, Lung, Kidney, Liver, Brain, Immune, Inflammatory, Blood, Musculoskeletal, Hormone, and Metabolic. Each system score predicted a wide variety of outcomes, aging phenotypes, and conditions specific to the respective system. We also combined the system scores into a composite Systems Age clock that is predictive of aging across physiological systems in an unbiased manner. Finally, we showed that the system scores clustered individuals into unique aging subtypes that had different patterns of age-related disease and decline. Overall, our biological systems based epigenetic framework captures aging in multiple physiological systems using a single blood draw and assay and may inform the development of more personalized clinical approaches for improving age-related quality of life.

## Introduction

The geroscience hypothesis states that targeting aging biology can prevent, delay, or treat multiple chronic diseases simultaneously. To test this hypothesis, reliable biomarkers must be developed that reflect valid age-related changes and responses to interventions.^1,2^ Biomarkers have been proposed based on diverse data types, such as clinical laboratory tests, functional phenotypes, or molecular omics data, each of which have their own advantages and are likely complementary.^1,3,4^

“Epigenetic clocks”, based on DNA methylation (DNAm), are among the most studied aging biomarkers.^2,5,6^ Many studies including meta-analyses have found that differences in epigenetic age among individuals of matched chronological age are predictive of morbidity, mortality, and other age-related phenotypes.^7–9^ Epigenetic clocks have been trained to predict chronological age, phenotypic aging, mortality risk, or pace of aging.^5,10,11^

Existing epigenetic clocks report an individual’s overall degree or pace of aging as a single value, capturing the quantitative heterogeneity in aging between individuals. However, there is also within-individual heterogeneity in the aging, at all levels of biological organization from organ systems down to cells (Figure 1).^12–15^ Recently, multi-dimensional aging biomarker panels based on clinical, functional, or plasma proteomic data have been developed to capture aging in different physiological systems.^16–18^ Different systems are complementary for predicting all-cause mortality, and are cross-sectionally associated with different diseases. For example, gait impairment was associated with muscle aging, and heart attacks with heart aging.^18^ Individuals age preferentially in different systems owing to divergent risk factors, and therefore there exists qualitative heterogeneity in the aging process which is not captured by a single uni-dimensional aging metric.

**Figure 1:**
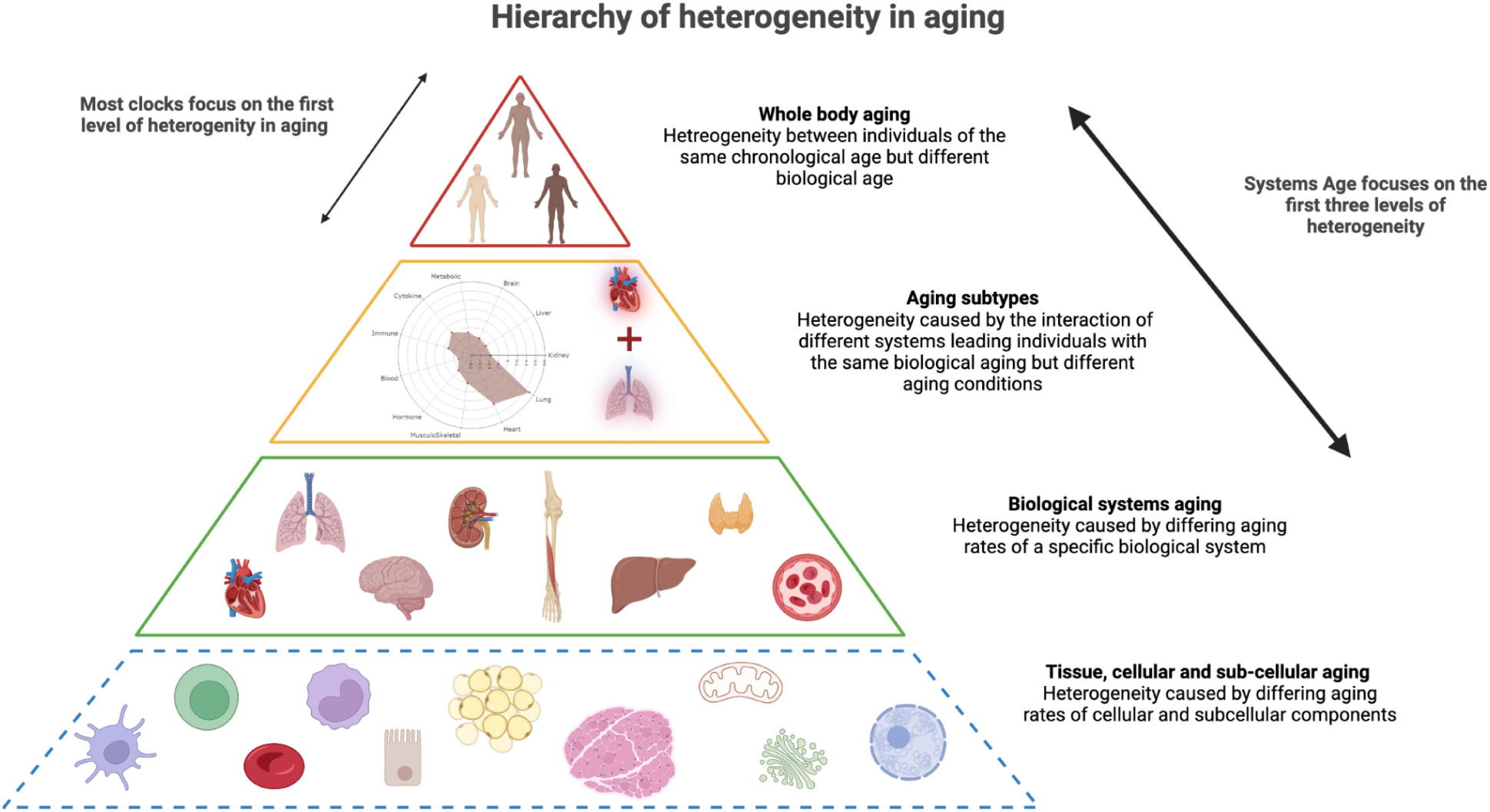
Hierarchy of heterogeneity in aging. Heterogeneity in aging starts at the very cellular and subcellular levels due to genetic and environmental factors. These variations in aging go on to accumulate at the tissue, organ and the biological system level causing differences in the rates of aging of different systems within an individual. These systems do not behave independently of each other and this leads to certain common patterns of deterioration across systems giving rise to aging subtypes. Eventually, all of these variations accumulate at the whole body level to cause variations in overall aging rates across individuals. Most epigenetic aging clocks typically focus on the whole body aging level of heterogeneity, Systems Age attempts to capture both the systems level heterogeneity and aging subtypes other than the whole body aging itself. Image created using Biorender.com.

System-specific epigenetic clocks would extend these findings using the distinct advantages of DNA methylation data. DNAm measurement requires only a single test, and data currently exists in numerous large human aging cohorts and an increasing number of interventional studies.^1,2,6,7^ Most studies utilize either the 450K or EPIC arrays and most epigenetic clocks are compatible, facilitating validation across multiple studies and contexts. In contrast, clinical and functional biomarkers can differ markedly between aging studies and involve many different tests. Other standardized omics types are currently not as widespread as DNA methylation. It is currently unclear whether system-specific predictors predict future aging outcomes other than all-cause mortality, and DNAm availability in longitudinal cohorts will help test this. DNAm data availability also facilitates explicit training to predict mortality risk; existing system-specific biomarkers using other data types^16–18^ were trained as chronological age predictors, which have significant biological and statistical limitations.^2,19–21^

Most existing aging studies measured DNAm in blood due to ease of access. Prior attempts have been made to build blood DNAm biomarkers that capture cardiac or metabolic disease risk,^22–24^ but it is currently unclear how much information about various other organ systems can be gleaned from blood methylation alone.

We aimed to construct a panel of systems-specific aging scores from blood DNA methylation data, leveraging linked clinical, functional, and mortality data to inform their construction. We then tested associations between the aging scores with baseline aging phenotypes and future disease risk in multiple cohorts. Our study demonstrates it is possible to capture heterogeneity across many physiological systems using a single blood DNA methylation test, in turn predicting decline and disease specific to each system.

## Results

### Systems Age Pipeline for modeling systems-specific aging

Systems Age was constructed in a five step process (Figure 2, Methods). First, we mapped clinical chemistry biomarkers, hematology biomarkers, functional assessments, and disease status available in the Health and Retirement study (HRS) to specific biological systems (Supplementary Table 1). Second, we performed principal component analysis (PCA) on measures within each system to identify latent signals captured by system-specific principal components (PCs). Third, we trained proxies of each system PC using methylation PCs. These proxies are analogous to existing methylation predictors of individual traits like BMI, smoking, or CRP, except that now we are capturing broader trends using PCs. Using PCs increases test-retest reliability without sacrificing validity^25^ (Supplementary Figure 1). Fourth, we calculated the predicted DNAm proxies of system PCs (using the models trained in HRS) in the Framingham heart Study (FHS). To integrate these into a single “system score” for each system, we trained a mortality prediction model for each system via elastic net Cox penalized regression. Finally, we trained an elastic net Cox model to combine the system scores into a unified whole body score called ‘Systems Age’. Scores were scaled to the expected age range for interpretability. Given the complexity of the training, we associated each system score with its corresponding biomarkers in HRS to determine which clinical chemistry and functional biomarkers contributed the most to each system score to aid in interpretation (Supplementary Figure 2).

**Figure 2:**
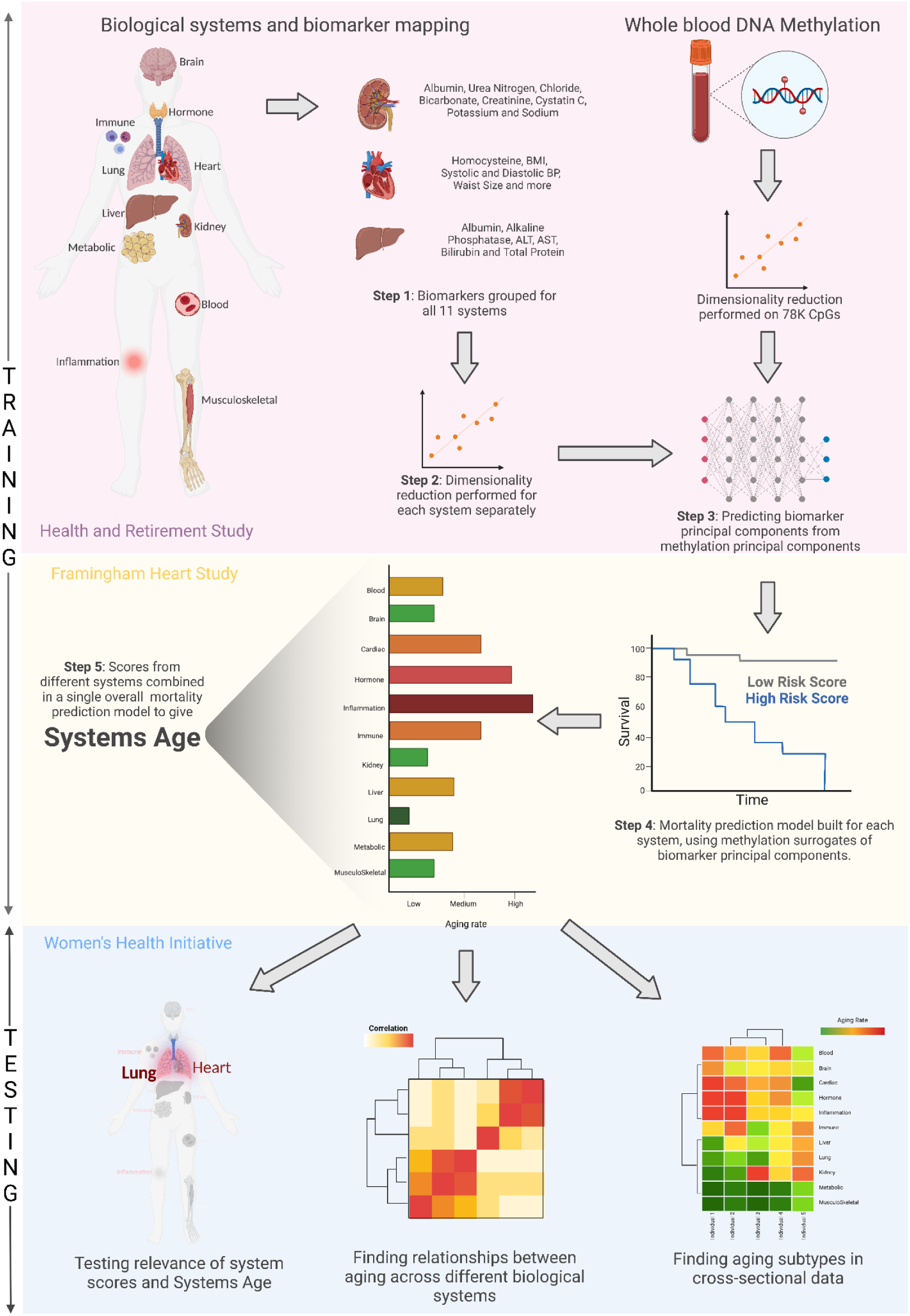
Analysis pipeline: Step 1 - Grouping Biomarkers into systems; Step 2 - Deconvoluting systems into principal components; Step 3 - Building DNAm surrogates of system PCs using ElasticNet regression; Step 4 - Building system scores by combining system PCs using Cox ElasticNet regression; Step 5 - Building Systems Age by combining system scores using Cox ElasticNet regression. Training done in HRS and FHS datasets while testing for specificity and aging subtypes done in WHI. Image created using Biorender.com.

### System scores capture meaningful and specific aging signals

Specificity of system scores was assessed in an independent sample from 3 cohorts of the Women’s Health Initiative (total N = ∼5,600; details in Methods). Cohorts were stratified by race when applicable, totalling 7 groups. Meta-analyses adjusting for chronological age (Figure 3, Supplementary Figure 3, 4A, Supplementary Tables 2-5) were performed testing associations with disease incidence (using Cox proportional hazard models), disease prevalence (logistic regression), and functional parameters of aging (OLS regression). To systematically compare associations between different clocks and different outcomes (Methods), we present z-scores in our main analyses, with standardized effect sizes and p-values provided in Supplement.

**Figure 3:**
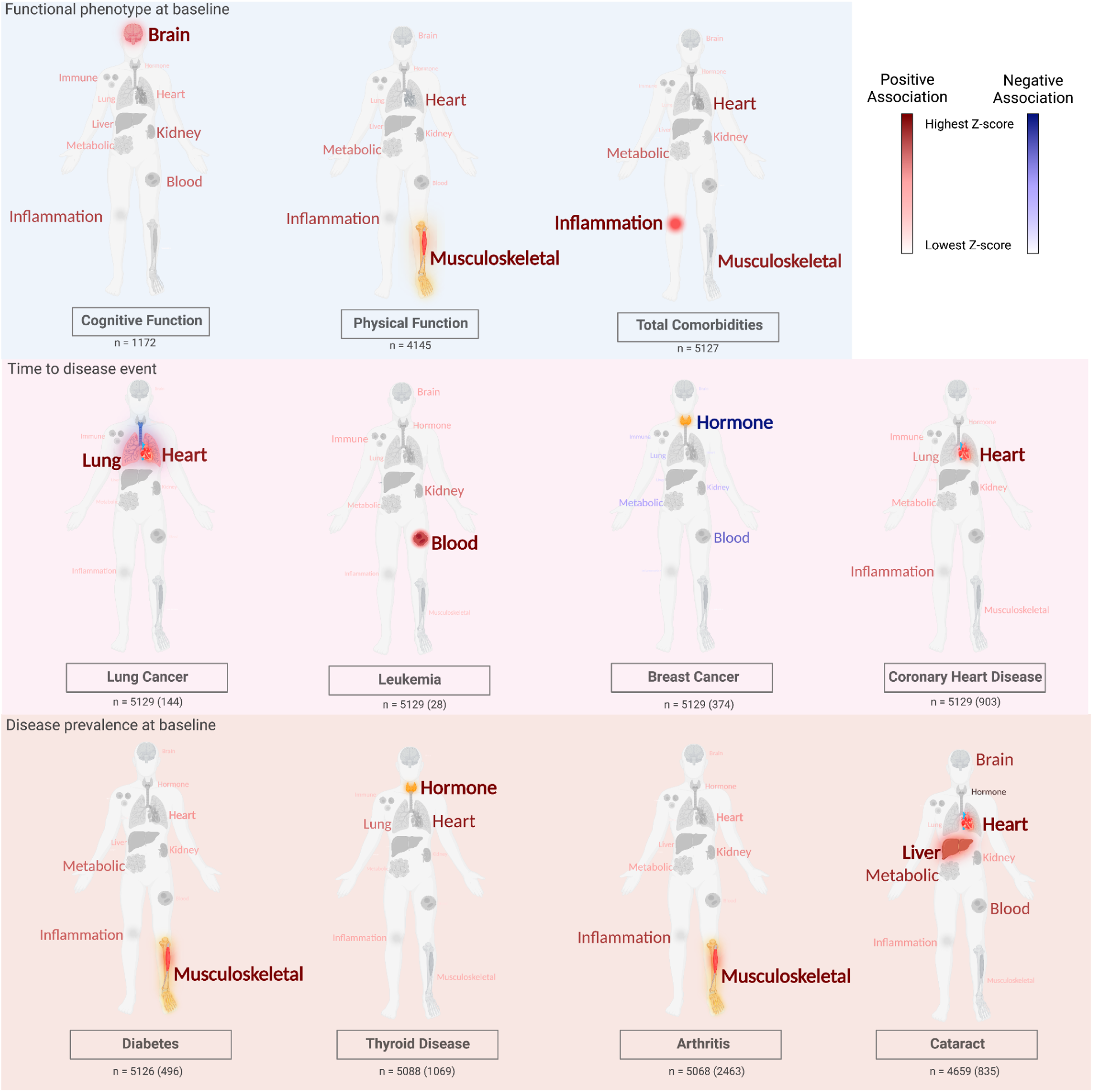
Meta-analysis associations (z-scores calculated using a race stratified analysis of 3 WHI datasets) for specific diseases and aging phenotypes with system score age accelerations depicted with text size and color. Additionally the system(s) with the highest positive association (or lowest in case of negative association) is bolded and the organ colored on the human figure. N for functional phenotypes ranges between 1172 and 5127. For time to disease events and disease prevalence at baseline, total N as well as number of events or individuals with diseases has been provided (in brackets). Total N for the time to disease events and disease prevalence at baseline is typically around 5000. For functional phenotypes at baselines Ordinary Least Squares regression model was used, for time to disease events cox proportional hazard models were used and for disease prevalence at baseline logistic regression models were used. Models built for each racial group separately and then meta-analyzed via a fixed effects model with inverse variance weights. Exact z-scores as well as heterogeneity p-values and other phenotypes are given in supplementary table 2 and 3. Image created using Biorender.com.

Our results suggested specificity of system scores to the expected organ system (Figure 3, Supplementary Figure 3 and 4A, Supplementary Tables 2 and 3), for both baseline diseases and functional phenotypes, as well as future conditions. For baseline function, the Brain score showed the strongest association with cognitive function (meta z-score = 3.51), and Musculoskeletal with physical function (z = 8.53). Heart was most strongly associated with time to CHD events (z = 8.29) and myocardial infarction (z = 6.30). Lung and Heart were most strongly associated with time to lung cancer (z = 9.69 and 9.49 respectively). For time to stroke, Heart (z = 3.40) was most strongly associated, followed by Metabolic (z = 3.35). Blood was most strongly associated with time to leukemia (z = 4.88).

In general, increased age indicators were associated with increased risk as expected. The one exception was time to reproductive organ cancers, which was negatively associated with all systems. Hormone (z = −3.04) and Blood (z = −2.22, not significant after multiple testing correction) were most negatively associated with breast and endometrial cancer, respectively.

For diseases at baseline, Musculoskeletal was most strongly associated with diabetes (z = 10.90) and arthritis (z = 5.15), Hormone was strongest for thyroid disease (z = 2.65), and Inflammation was strongest for total comorbidities (z = 7.65).

### System scores maintain predictive ability of epigenetic clocks while adding specificity

To ensure predictive ability is not compromised to achieve specificity, we repeated our meta-analyses to compare system scores to previously trained epigenetic clocks (Figure 4, Supplementary Figure 4B, 5 and 6, Supplementary Tables 2, 3, 4 and 5). We focused on three prominent epigenetic clocks previously demonstrated to be strongly associated with aging outcomes: PCPhenoAge, DNAmGrimAge and DunedinPACE.

**Figure 4:**
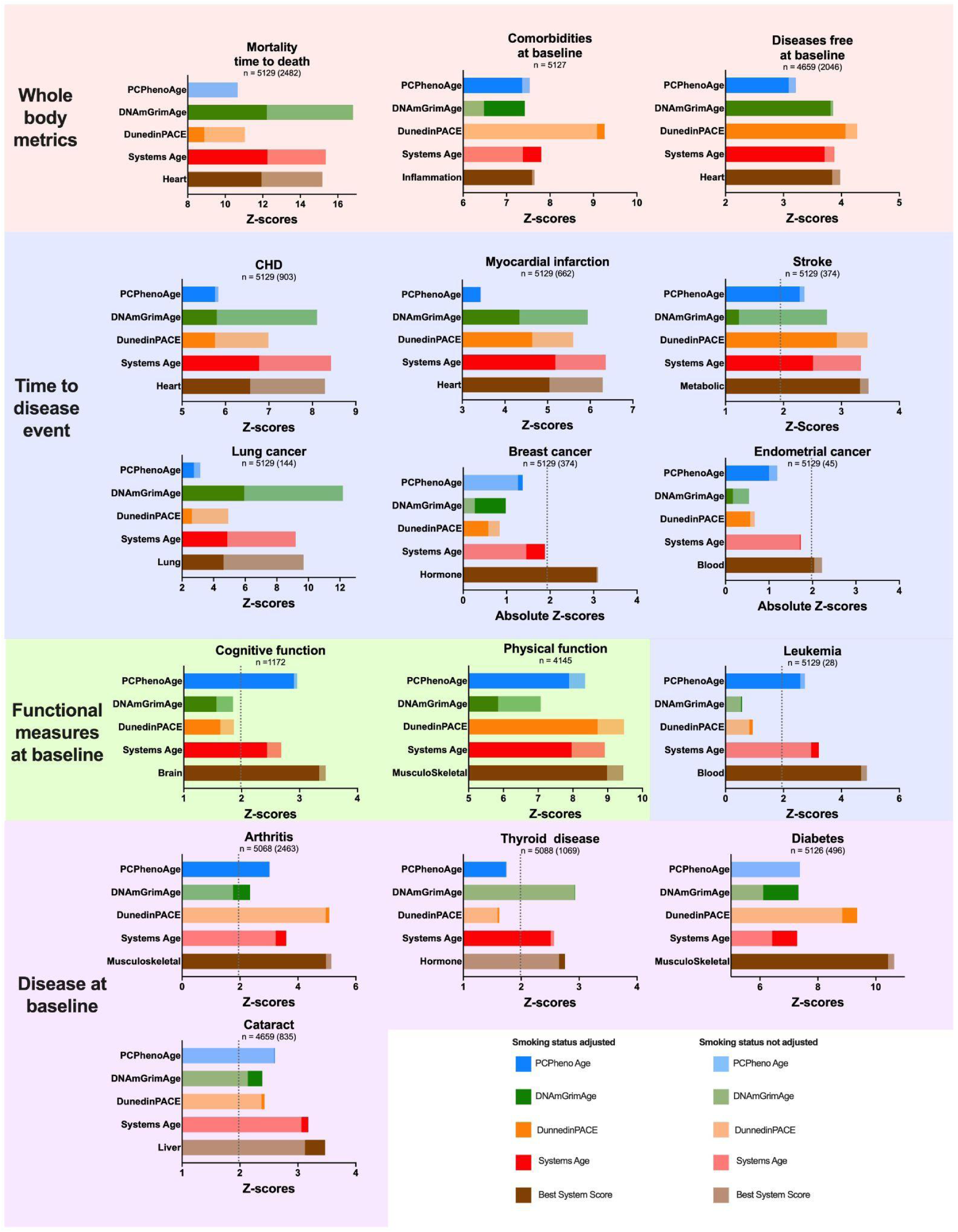
Meta-analysis associations (z-scores calculated using a race stratified analysis of 3 WHI datasets) for specific diseases and aging phenotypes with age accelerations of different clocks, Systems Age and the best system score plotted for smoking status adjusted (in darker shades) and no smoking status adjusted (lighter shades). N for functional measures ranges between 1172 and 4145. For time to disease events and disease prevalence at baseline, total N as well as number of events or individuals with diseases has been provided (in brackets). Total N for the time to disease events and disease prevalence at baseline is typically around 5000. For functional measures at baseline Ordinary Least Squares regression model was used, for time to disease events cox proportional hazard models were used and for disease prevalence at baseline logistic regression models were used. Models built for each racial group separately and then meta-analyzed via a fixed effects model with inverse variance weights. Note, because endpoints differ in the statistical test being used (e.g. linear vs cox vs logistic models), and the sample sizes or number of cases, Z-scores between outcomes may have different magnitudes. Thus, we focus on the differences between the clocks within condition/outcome, which is why the axis have been scaled to focus on the differences between the clocks. Exact z-scores as well as heterogeneity p-values are given in supplementary table 2, 3, 4 and 5. Plots generated using Prism 9.

The most relevant system scores had a higher meta z-score based on effect sizes to existing epigenetic clocks for 10 of the 14 diseases, and was a close second for the remaining 4. To determine whether these differences were statistically significant, we performed receiver operating characteristic curve (ROC) analysis (Figure 5, Supplementary Figure 7, Supplementary Tables 6, 7 and 8) on each cohort comparing the best system score to the different clocks. The system scores had a significantly better area under the curve compared to some clocks when it came to prediction of associations with Mortality (3.55 and 3.84 meta z-score for Heart compared to DunedinPACE and PCPhenoAge ROC, respectively), total comorbidities at baseline (2.39 and 2.17 for Inflammation compared to DNAmGrimAge and PCPhenoAge respectively, not significant after multiple testing correction for DNAmGrimAge), CHD (2.15 for Heart compared to PCPhenoAge), MI (2.5 for Heart compared to PCPhenoAge), leukemia (5.63 and 6.13 for Immune compared to DunedinPACE and DNAmGrimAge respectively), lung cancer (3.13 and 3.92 for Lung compared to DunedinPACE and PCPhenoAge respectively), cognitive function (2.99, 3.87, and 3.23 for Brain compared to DunedinPACE, DNAmGrimAge, and PCPhenoAge respectively), physical function (3.53 and 2.50 for MusculoSkeletal compared to DNAmGrimAge and PCPhenoAge respectively), arthritis (3.62 and 3.43 for MusculoSkeletal compared to DNAmGrimAge and PCPhenoAge respectively), and diabetes (2.66, 5.17, and 5.66 compared to DunedinPACE, DNAmGrimAge, and PCPhenoAge respectively). We did not find statistically significant improvements for cataracts or stroke.

**Figure 5:**
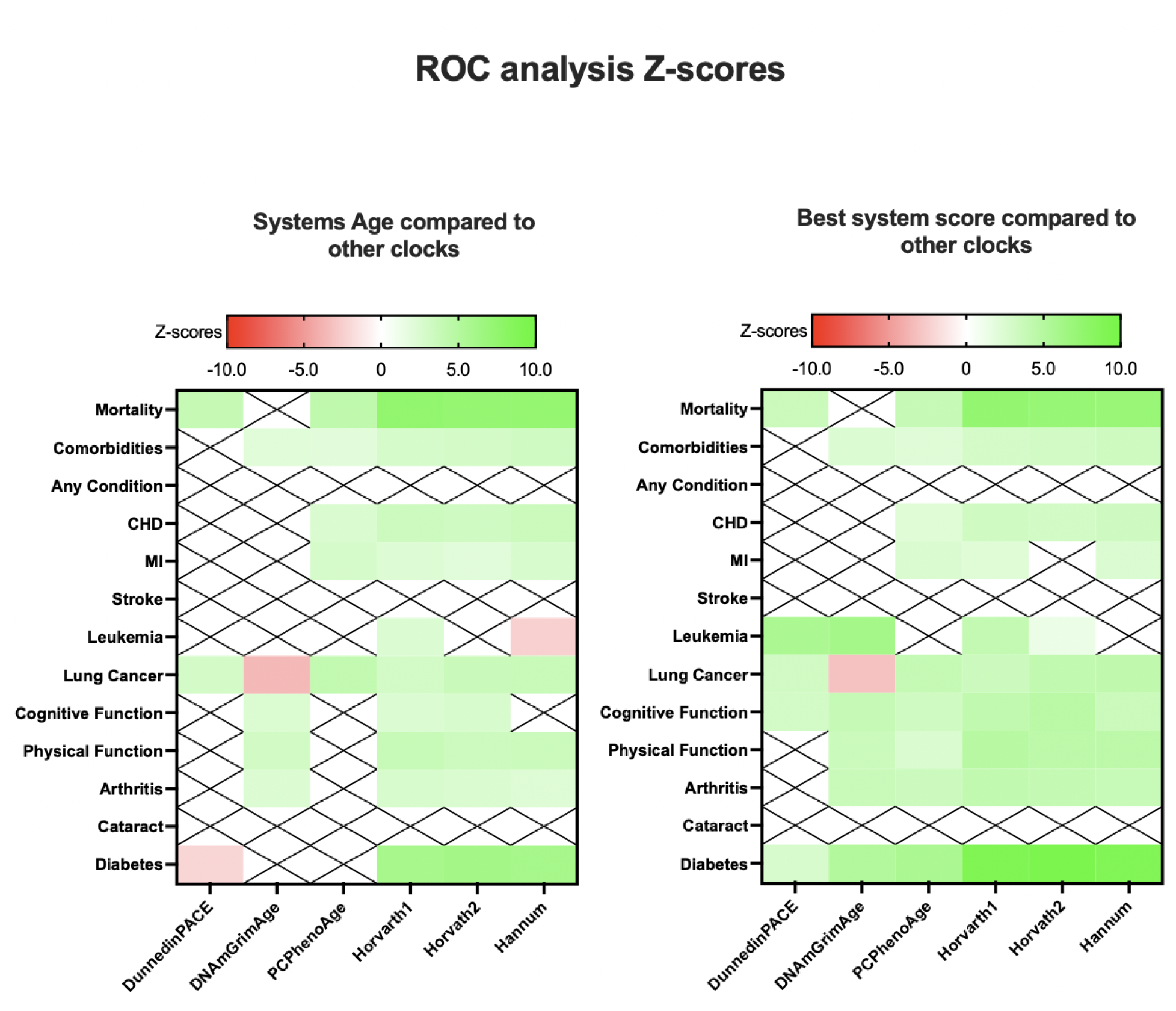
ROC analysis comparing each clock to Systems Age or best system score. Z-scores were calculated using cohort stratified meta analysis using Stouffer’s method on 3 WHI cohorts: EMPC, AS311 and BAA23. Scores with no significant difference have been crossed out. Significance is defined as absolute z-scores with values greater than 1.96. Exact z-scores provided in supplementary tables 6 and 9 with corresponding AUCs in supplementary tables 8. Plots generated using Prism 9.

Overall, these results suggest that predictive ability was not sacrificed to improve specificity, and indeed there may have been some improvements in prediction.

### Systems Age: the golden mean

When training clocks, each may be biased towards specific aspects of aging based on the combination of variables and datasets used for training. Indeed, in our WHI analyses, we noted that DNAmGrimAge was particularly strong in predicting mortality, lung cancer, heart disease, and thyroid disease, but did not predict leukemia, arthritis, or cognitive function (Figure 4, Supplementary Figure 8, Supplementary Tables 2 and 3, Supplementary File 1). Similarly, DunedinPACE was particularly strong in predicting physical function, arthritis, stroke, diabetes, and number of comorbidities, but did not predict leukemia or thyroid disease. PCPhenoAge did predict leukemia and cognitive function, but it was not as strong as DNAmGrimAge or DunedinPACE in other phenotypes. Since each systems score showed superior or equivalent associations with specific diseases and aging phenotypes, we hypothesized that combining them into a single Systems Age score would lead to a more uniform prediction across all diseases and aging phenotypes. Indeed, we found that Systems Age was associated with all of the above conditions.

Among the four clocks, Systems Age had the strongest associations of all clocks for 4 conditions, was the second best for 8 conditions, and third best for 2 conditions. ROC analysis (Figure 5, Supplementary figure 7, Supplementary Tables 8-10) indicated that Systems Age was comparable to the best of the 3 other clocks for each phenotype. In only two cases, Systems Age showed significantly worse association than the best clock (lung cancer for GrimAge, diabetes for DunedinPACE). Thus Systems Age retains unbiased associations with all phenotypes, without significantly sacrificing predictive ability.

### System scores capturing aging signals beyond smoking

Many associations between epigenetic clocks and age-related conditions may be driven by smoking^26^. To determine the degree to which smoking affects the system scores, we repeated the meta-analyses while adjusting for smoking status (Supplementary Figures 5B, 8B, Supplementary Table 4). For example, the Metabolic score’s strong association with time to stroke was changed minimally when adjusting for smoking (3.35 to 3.20 after adjustment). In contrast, GrimAge’s association with stroke decreased >50% and was no longer significant when adjusting for smoking (2.75 to 1.23 after adjustment). Reduced associations with CHD were observed across all clocks after adjusting for smoking (as expected given that smoking is a major risk factor for heart disease), but Systems Age and Heart score retain much of their association after smoking status adjustment (meta z-score in CHD before and after smoking status adjusted for Heart 8.29 vs 6.57, Systems Age 8.43 vs 6.77, DNAmGrimAge 8.11 vs 5.80). Similar expected impacts of smoking were seen for time to myocardial infarction, lung cancer, and death. For other diseases and aging phenotypes, Systems Age and system scores retained their prediction after adjusting for smoking, indicating they captured epigenetic signals beyond just cigarette exposure. We also performed an analysis among never smokers (Supplementary Figure 5C and 8C as well as Supplementary Table 5) finding that system scores retained strong associations.

### System scores predict morbidity and mortality in three additional cohort studies

WHI is composed of multiple diverse cohorts with a variety of disease and functional outcomes, but it is limited to postmenopausal women. To test broader applicability of system scores, we examined three additional cohorts, including the Alzheimer’s Disease Neuroimaging Initiative (ADNI),^27^ Baltimore Longitudinal Study of Aging (BLSA),^28^ and Swedish Adoption Twin Study on Aging (SATSA)^29^.

We again observed specificity of system scores (Figure 6, Supplementary Tables 11 and 12). In ADNI, we tested associations with clinically relevant outcomes among the presumed AD population, after adjustment for age, sex and education status. The Brain score showed the strongest associations with cranial volume decline in AD patients (Figure 6A) (whole brain volume decline 2.98; hippocampal 2.38; fusiform 3.31, hippocampal volume not significant after multiple testing correction) followed by Metabolic score (whole brain 2.86; hippocampal 2.10; fusiform 2.80; hippocampal volume not significant after multiple testing correction). No system scores were associated with cognitive scales such as MMSE, ADAS13 and MOCA in AD patients (Supplementary Tables 11). When considering the full ADNI cohort, including healthy, MCI and AD patients, (Figure 6B) the Inflammation score showed the strongest associations with whole brain volume decline (z-score 3.27) and ADAS13 (z-score 2.20, not significant after multiple testing correction). We tested whether system scores could differentiate NCI, MCI, and AD subjects (Figure 6C). The Inflammation score showed the strongest association with diagnosis (one-way anova p =0.007, not significant after multiple testing correction) and that this is more robust than prior epigenetic clocks (DunedinPACE p = 0.08).

**Figure 6:**
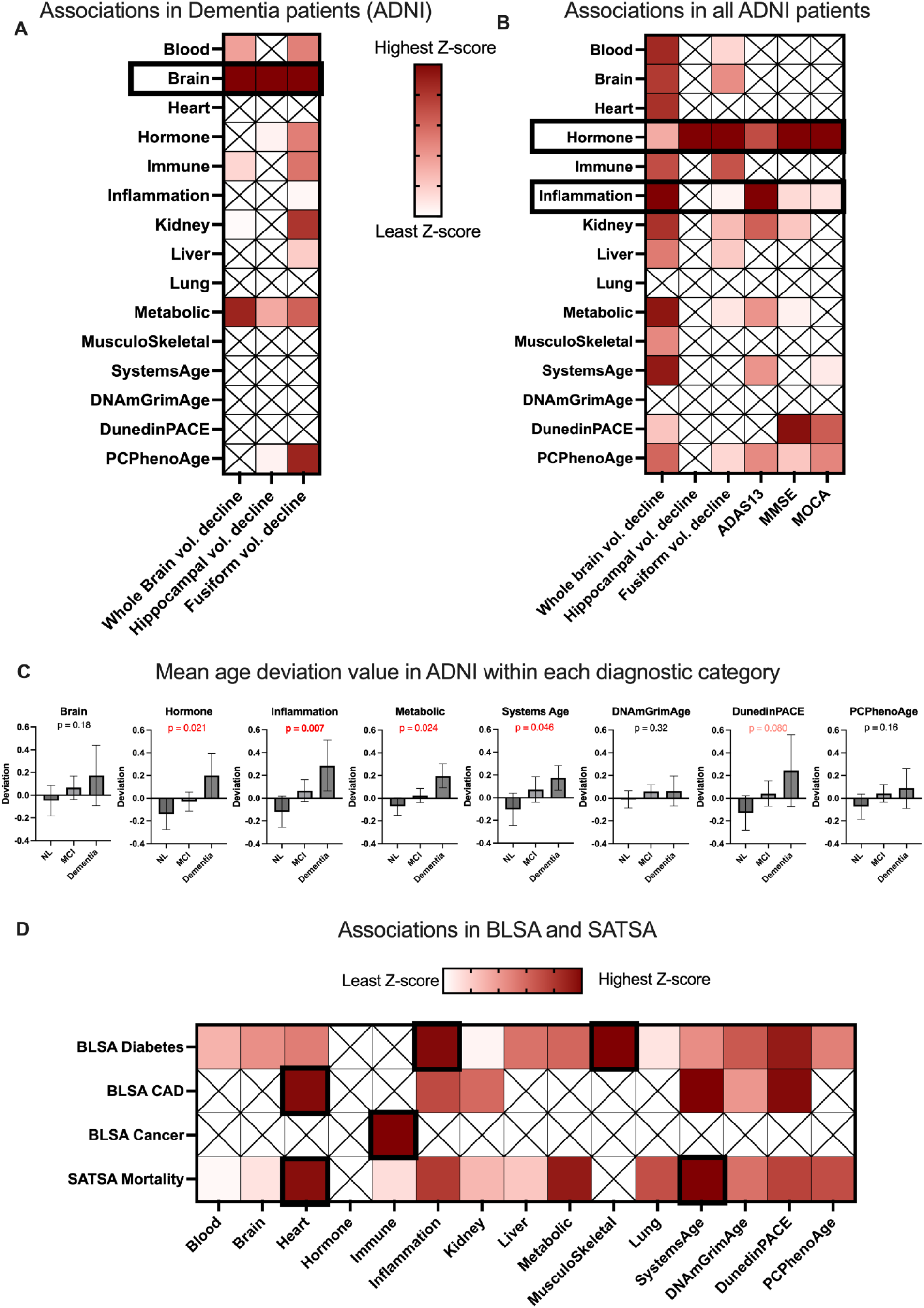
Validation in additional cohorts. A) Association of different brain volume decline metrics with clocks in AD patients in ADNI dataset, a mixed effects cox model was used, details described in methods section, exact associations provided in supplementary tables 11. B) Association of different brain volume decline and cognitive score metrics with clocks in all ADNI samples, including healthy, MCI and AD patients, a mixed effects cox model was used details described in methods section, exact associations provided in supplementary tables 11. C) Mean Age deviation value in ADNI within each diagnostic category for clocks that showed some of the most significant changes in Fig 6A and 6B, p-values calculated using a one way anova test. D) Associations of different aging diseases with clocks in BLSA and SATSA datasets with description of analysis and datasets provided in the methods section. Exact associations are provided in supplementary tables 12. All graphs were generated in Prism 9.

In BLSA (Figure 6D, Supplementary Table 12), baseline Diabetes status was most strongly associated with Inflammation (z = 5.6), MusculoSkeletal (5.66), and Metabolic 4.19). CAD at baseline was most strongly associated with Heart (z = 2.96) and Inflammation (2.69) Notably, the same systems were most strongly associated with diabetes and CAD in WHI. Cancer was associated with the Immune score (z = 2.64, not significant after multiple testing correction). In SATSA (Figure 6D, Supplementary Table 12), Heart (z = 5.38) and Systems Age (5.5) showed strongest associations with mortality, again replicating prior WHI results.

### System scores capture distinct dimensions of aging

To better define relationships between systems, we looked at correlations between age-adjusted system scores across all WHI cohorts (Figure 7A and Supplementary Table 13). Some systems were highly correlated with each other, such as Heart and Lung (r = 0.759), or Inflammation and Musculoskeletal (r = 0.716). Hierarchical clustering revealed that Heart and Lung formed a cluster, Liver, Brain and Blood formed a second cluster, and Metabolic, Inflammation, and Kidney formed a third cluster. The latter two clusters formed a super-cluster that also included Heart and Musculoskeletal.

**Figure 7:**
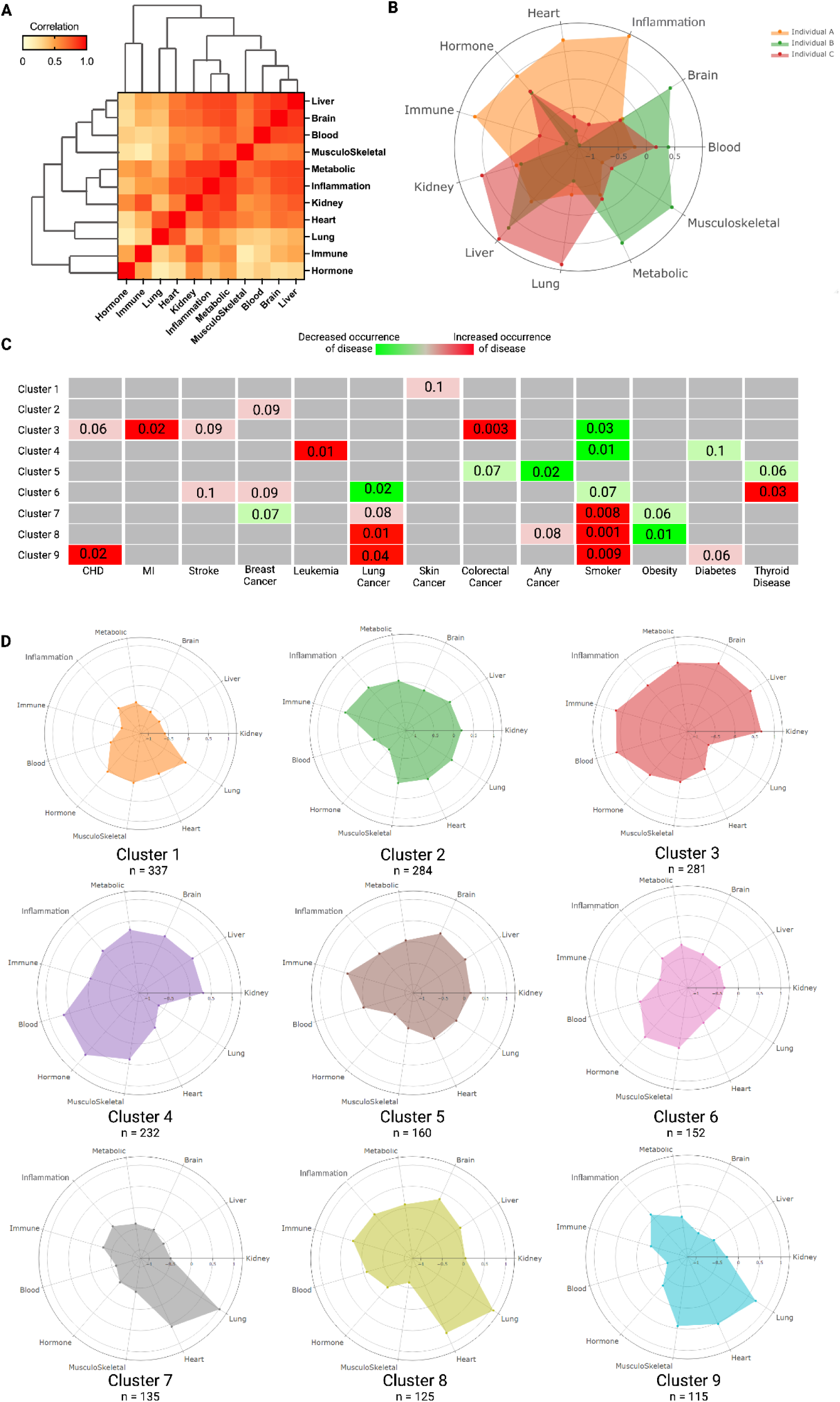
Discovering Aging Subtypes. A) Correlations between system scores across all WHI cohorts corrected for batch effects (N = 5129). Exact correlations provided in supplementary table 8 B) Three chronological age matched individuals with the same race and gender as well as similar age-accelerated Systems Age having very different age-accelerated system scores. C) Overrepresentation analysis of presence or absence of diseases amongst individuals from 9 different clusters. P Values have been calculated using fisher’s exact test and are available in supplementary tables 14. D) Mean age accelerated score has been depicted for each cluster using a spider plot and is also available in supplementary tables 15. Image created using Biorender.com.

We hypothesized individuals can be grouped based on their system scores to generate aging subtypes with distinct predisposition to aging related diseases and conditions. We observed there existed individuals with the same chronological age, overall Systems Age and demographics, yet entirely different patterns of age-accelerated system scores (example of 3 individuals shown in Figure 7B). To test whether these patterns of system scores constituted biologically relevant aging subtypes, we clustered individuals using adaptive hierarchical clustering in the WHI EMPC cohort. This analysis identified 9 unique groups or clusters, with each cluster showing distinct patterns across the different system aging scores, as well as different associations with the prevalence or future occurrence of certain diseases (Figure 7C and 7D, Supplementary Tables 14-15). For example, Cluster 8 and 9 were found to have a high mean Lung aging score (Lung age accelerations 1.06 and 0.71) and a higher risk of future lung cancer (p value: 0.01, 0.04). These groups also had an overrepresentation of smokers (p: 0.001, 0.009), indicating they captured smoking pathophysiology. As expected, their distinct clustering stemmed from other pathophysiology: Group 8 had fewer obese individuals (p = 0.01) while Group 9 was enriched for future CHD events (p = 0.02). At the same time, Cluster 3 demonstrated an increased riskof MI (p = 0.02) yet it showed low Lung aging and decreased prevalence of smokers (p = 0.03). It also demonstrated increased Metabolic (average age acceleration 0.55) and Inflammation (average age acceleration 0.34) aging. Thus, while Cluster 3 and 9 are both at risk for cardiovascular diseases, the risk stems from different sources (smoking vs. metabolic and inflammatory aging). Overall, the system’s scores were capturing relevant aging subtypes that had distinct behavioral and genetic patterns predisposing individuals to certain types of aging phenotypes and diseases.

## Discussion

Commonly used epigenetic clocks predict chronological age, composite measures of biological age, and the pace of aging.^1^ These have been extensively validated due to widespread availability of DNAm data and the compatibility of most epigenetic clocks across many datasets.^2,5,7^ However, these are typically used as uni-dimensional measures, which would not capture the multi-dimensional nature of aging. Defining these dimensions at the level of physiological systems is useful given that modern medicine is often organized at this level, and we expect risk factors, diseases, and aging interventions to affect specific subsets of systems. Prior studies have shown the utility of system-specific aging biomarker panels using clinical biomarkers or omic data types other than DNA methylation.^16–18^ Here we were able to accomplish a similar goal using blood DNA methylation. The standardization offered by DNA methylation data generation allows for both potential wide-spread usage and ease of cross-comparison across experiments. We validate the specificity and predictive ability of our single blood draw test in multiple large aging cohorts, and anticipate application in many more cohort and intervention studies that have measured blood DNAm ^1,2^. We also extend system-specific prediction of future outcomes beyond all-cause mortality to multiple phenotypes, including lung cancer, breast cancer, leukemia, and coronary heart disease. Systems Age is the first predictor to fulfill multiple criteria simultaneously: 1) it is trained to predict mortality rather than chronological age, 2) it captures heterogeneity across different biological systems and 3) it only requires a single assay performed on samples from a single blood draw.

The use of a single assay performed on single blood draw offers multiple advantages: the assay can be easily repeated over time for longitudinal measurements, it greatly reduces missing data, and it does not require optimization of numerous tests each with their own complexities. The same DNAm variables are consistently measured across studies, allowing for validation in many settings. In contrast, different studies usually have very different sets of clinical and functional variables.

Prior to our study, it was unclear to what extent signals in other organ systems could be captured in blood DNAm alone. However, our results demonstrate that this is possible. Systems Age scores were not only predictive of various aging outcomes, but were also specific to the pathophysiology of their intended system. For example, the Heart score was most associated with heart disorders and mortality in both WHI and BLSA,^30^ reflecting that cardiovascular disease is the leading cause of mortality worldwide.^31^ However, Heart demonstrated specificity in that it was only very weakly associated with diseases such as baseline arthritis or time-to-leukemia (associated with Musculoskeletal and Blood respectively).^32^ Likewise, the Inflammation score was strongly associated consistent with known pathophysiology with time-to-CHD^33^, baseline arthritis^34^, baseline physical and cognitive functioning,^35,36^ and total number of comorbidities at baseline.^37^ Interestingly, the Inflammation score showed the most significant difference between AD, MCI and healthy patients in ADNI and was associated with brain volume and cognition, consistent with prior studies highlighting the role of systemic inflammation in Alzheimer’s disease.^38^ The Brain score was associated with baseline cognitive functioning and time-to-stroke in WHI, as well as brain volumes in dementia patients in ADNI, as expected ^39,40^. The Musculoskeletal score was strongly associated with physical function and baseline arthritis as expected (WHI and BLSA). The Musculoskeletal score’s association with diabetes may reflect that sarcopenic obesity is a risk factor for type 2 diabetes through reduced glucose disposal with reduced muscle mass ^41^, and its association with total comorbidities may reflect sarcopenia’s role in frailty as well as the predisposition to injury in frail individuals ^42^. Thus, blood DNA methylation data alone can be used to derive many different specific aging scores for various physiological systems, rather than just a single blood-specific or generalized aging process.

Epigenetic aging was negatively associated with breast cancer risk in women, with the strongest negative association being Hormone. This is consistent with prior literature that showed epigenetic clock acceleration is associated with a younger age at menopause,^43^ which is in turn linked to a lower risk of reproductive organ cancers.^44,45^ TIn moving forward, there may be a further need to develop reproductive system specific scores for different sexes as well as expand the Hormone score.

Given the interconnectedness of aging, it was unclear prior to our study whether a given aging phenotype would be better predicted by epigenetic clocks that detected more global aging signals, or clocks trained on a limited set of clinical biomarkers specifically related to that phenotype. Our results demonstrated the advantages of the latter approach. PCPhenoAge, DNAmGrimAge, and DunedinPACE^8,9,11^ were not intended to capture global aging signals that are not limited to any particular system. For 10 of the 14 system specific phenotypes and diseases we tested, the most relevant system score had higher meta-Z scores than all three of these clocks. ROC analysis demonstrated that many of these improvements were statistically significant. While the relevant system score did not surpass the predictive ability of all existing clocks for other outcomes, the system scores were nearly as predictive, while additionally being more interpretable and granular concerning which systems are related to which phenotypes. The overall Systems Age also showed more uniform prediction across all phenotypes, showing either the strongest or second-strongest associations with 12 of the 14 conditions, compared to the other three clocks. Thus, Systems Age appeared to not be strongly biased by a particular dimension of aging, which is likely the result of first training predictors of mortality in each physiological system independently before combining them. In further support of this idea, system scores and Systems Age remain highly associated with all these phenotypes after correction for smoking.

In addition to heterogeneity at the systems level, we examined the heterogeneity that arose due to interaction of different systems. Interestingly, some system scores were more correlated than others. Heart and Lung were highly related, consistent with their common vulnerability to smoking and shared pulmonary circulation. The Liver and Blood scores showed strong correlations, potentially reflecting the liver’s blood filtration function and production of blood products. Metabolic, Inflammation, and Kidney were highly correlated, reflecting numerous links between metabolism and inflammation^46^ as well as the inclusion of IL-6 and CRP in both systems, and the contributions of both to kidney aging ^47,48^. Of note, while the correlations between systems do likely reflect true physiological interactions, it is also possible that some of the correlation structure can be attributed to similar mechanisms by which they impact the blood methylome and vice versa.

We defined subtypes as groups of individuals with similar systems specific aging scores that have an over or under representation of specific diseases and conditions. We showed the existence of 9 such distinct clusters or aging subtypes that had predisposition to very distinct diseases and conditions. Aging subtypes have already been documented in the literature.^12^ However, we provided evidence that these distinct groups can be observed even when using a single assay–in our case, DNA methylation assessed in blood. In the future, information on longitudinal changes will be critical for further defining age subtypes and their relevant disease risks. This could facilitate sub-classification of aging conditions and eventually inform targeted therapies based on aging subtypes.

Systems Age as a framework for capturing heterogeneity shows a lot of promise, yet limitations remain. Due to insufficient clinical measures paired with DNAm, aging of the reproductive, digestive, peripheral nervous, and other systems are not represented in Systems Age. It is also important to note that one could add more clinical biomarkers and phenotypes to each system, and there are other ways to map biomarkers to systems (there is no gold standard mapping). Another caveat is that it is unknown why DNAm in blood reflects aging in other physiological systems, and what is the molecular relationship between the clinical biomarkers, disease states and blood DNAm. It could reflect shared genetic variation, exposures, age-related patterns between tissues. Alternatively it could involve intercellular signaling influencing DNAm in blood, or blood DNAm reflecting processes by which immune cells affect aging in those systems. Further analysis needs to be performed to understand these relationships better. Finally, Systems Age uses only clinical data to first generate scores that are then predicted from epigenetic data. Other data types, such as proteomics, metabolomics, or imaging, may help capture more diverse dimensions of aging, and the increasing amount of multi-omics data will be invaluable for this effort ^2,49^.

Overall, we highlight the importance of capturing heterogeneity in aging while also building a reusable framework for quantifying multisystem aging. We show that system-specific predictors can be estimated from blood DNA methylation. By providing system-specific scores, Systems Age may better pinpoint which age-related conditions that individuals are at risk for. Given that therapies are often targeted at particular systems, it will be interesting to determine whether the specificity of Systems Age can provide guidance when selecting geroscience-based interventions.

## Acknowledgements and Funding

We would like to thank John Gonzalez, Kyra Thrush, Jenel Armstrong, Jessica Kasamato, Daniel Borrus, Ahana Priyanka, Perry Kuo and other members of the Higgins-Chen lab at Yale University and members of the Levine lab at Altos for their feedback and support.

This work was supported by the National Institute on Aging (NIA:1R01AG065403-03, 1R01AG057912-05 to M.E.L. and A.H.C.). It was also supported by the Impetus Grant (R.S.), the Gruber Science Fellowship at Yale University (R.S.), the Thomas P. Detre Fellowship Award in Translational Neuroscience Research from Yale University (to A.H.C.), and the Medical Informatics Fellowship Program at the West Haven, CT Veterans Healthcare Administration (to A.H.C.). A.H.S. was supported by grant RF1AG074345 from the National Institute on Aging.

The HRS study was supported by NIA R01AG060110-05, R01AG068937-02 and U01AG009740.

The Framingham Heart Study is funded by National Institutes of Health contract N01-HC-25195 and HHSN268201500001I. The laboratory work for this investigation was funded by the Division of Intramural Research, National Heart, Lung, and Blood Institute, National Institutes of Health. The analytical component of this project was funded by the Division of Intramural Research, National Heart, Lung, and Blood Institute, and the Center for Information Technology, National Institutes of Health, Bethesda, MD.

We would like to recognize the support of the WHI Program Office: (National Heart, Lung, and Blood Institute, Bethesda, Maryland) Jacques Rossouw, Shari Ludlam, Joan McGowan, Leslie Ford, and Nancy Geller, the WHI Clinical Coordinating Center: (Fred Hutchinson Cancer Research Center, Seattle, WA) Garnet Anderson, Ross Prentice, Andrea LaCroix, and Charles Kooperberg, the WHI Investigators and Academic Centers: (Brigham and Women’s Hospital, Harvard Medical School, Boston, MA) JoAnn E. Manson; (MedStar Health Research Institute/Howard University, Washington, DC) Barbara V. Howard; (Stanford Prevention Research Center, Stanford, CA) Marcia L. Stefanick; (The Ohio State University, Columbus, OH) Rebecca Jackson; (University of Arizona, Tucson/Phoenix, AZ) Cynthia A. Thomson; (University at Buffalo, Buffalo, NY) Jean Wactawski-Wende; (University of Florida, Gainesville/Jacksonville, FL) Marian Limacher; (University of Iowa, Iowa City/Davenport, IA) Jennifer Robinson; (University of Pittsburgh, Pittsburgh, PA) Lewis Kuller; (Wake Forest University School of Medicine, Winston-Salem, NC) Sally Shumaker; (University of Nevada, Reno, NV) Robert Brunner Women’s Health Initiative Memory Study: (Wake Forest University School of Medicine, Winston-Salem, NC) Mark Espeland. Data from the Epigenetic Mechanisms of PM-Mediated CVD Risk (EMPC) were generated under the National Institute of Environmental Health Sciences grant R01-ES020836.

The BLSA study is conducted by the Intramural Research Program of the National Institute on Aging (NIA), part of the National Institutes of Health at the U.S. Department of Health and Human Services. The authors also wish to thank the staffs and participants of the BLSA.

We thank Ida Karlsson for assistance with data in the SATSA study.

## Author Contributions

M.E.L. conceived the project. R.S. and M.E.L. conceived the study design. R.S. and M.E.L. built the pipeline for training Systems Age. R.S. performed analysis in WHI to validate Systems Age. R.S. performed analysis in BLSA to validate Systems Age. R.S. and Y.M. performed analysis in ADNI to validate Systems Age. R.S. and C.Q. performed analysis in SATSA to validate Systems Age. A.H.C. and R.S. performed reliability analysis. R.S., A.H.C. and M.E.L. interpreted the results. A.H.C. and M.E.L. provided supervision and managed the overall project. Other authors contributed data and analyses related to WHI (A.S., R.C., J.E.M., P.B., T.A., E.A.W.) and HRS (E.M.C.). All authors reviewed and contributed to the manuscript.

## Data Availability Statement

HRS data is available at hrsdata.isr.umich.edu. FHS data is available at dbGaP, accession number: phs000724.v7.p11. Access to WHI data is available upon review and subsequent approval of proposals submitted through the study website (https://www.whi.org/propose-a-paper). The reliability dataset is available on GEO, accession ID GSE55763. Details about the SATSA cohort can be found on the Maelstrom Research platform (https://www.maelstrom-research.org/mica/individual-study/satsa), Phenotypic data are archived in the National Archive of Computerized Data on Aging (https://www.icpsr.umich.edu/icpsrweb/ICPSR/studies/3843) Methylation data is archived in EMBL-EBL (https://www.ebi.ac.uk/biostudies/studies/S-BSST1206). ADNI data is publicly available; access is granted after application approval from the study’s research review committee (adni.loni.usc.edu/data-samples/access-data/). The BLSA data are available upon request. Applications should be made through the website https://www.blsa.nih.gov/.

## Code Availability Statement

Code to Systems Age will be available upon publication.

## Licensing System Age

Systems Age and its components have been licensed out to a commercial partner. For more details please.

Systems Age code was first developed in July 2021. Since the first development, the Systems Age code has been updated with new biomarkers and systems, and has now been validated in independent datasets (WHI, BLSA, ADNI and SATSA).

## Methods

### Datasets used for training Systems Age

Two different longitudinal studies were used for training Systems Age: the Health and Retirement Study (HRS) and Framingham Heart Study (FHS). We previously utilized and described methylation data from these datasets in a separate study ^25^. Briefly, as stated on their website, HRS is a nationally representative sample of Americans over age 50 years. HRS had biomarker information available for 9,933 participants of which Infinium Methylation EPIC BeadChip data was available for 4,018 individuals ^50^. Out of the 4018 individuals only 3,593 had clinical data (age range 51–100 years) which were used for training of Systems Age. All participants provided written informed consent. The study was approved by the Institutional Review Board (IRB) at the University of Michigan (HUM00061128).

FHS includes 2,748 FHS Offspring cohort participants attending the eighth exam cycle (2005–2008) and 1,457 Third Generation cohort participants attending the second exam cycle (2005–2008), who consented to provide their DNA for genomic research ^51,52^. DNA methylation was assayed with the Infinium HumanMethylation450 BeadChip and is available in dbGaP (accession no. phs000724.v7.p11). For the purpose of training Systems Age FHS Offspring data was used but for scaling of system scores and systems age to age range both the Offspring and Third generation data was used. Deaths of FHS participants occurring before 1 January 2014 were ascertained by routine contact with participants, surveillance at the local hospital, local obituaries and queries to the National Death Index dates. Causes of death were reviewed by an endpoint panel of three investigators. The study protocol was approved by the IRB at Boston University Medical Center. All participants provided written informed consent at the time of each examination visit.

**Table 1:**
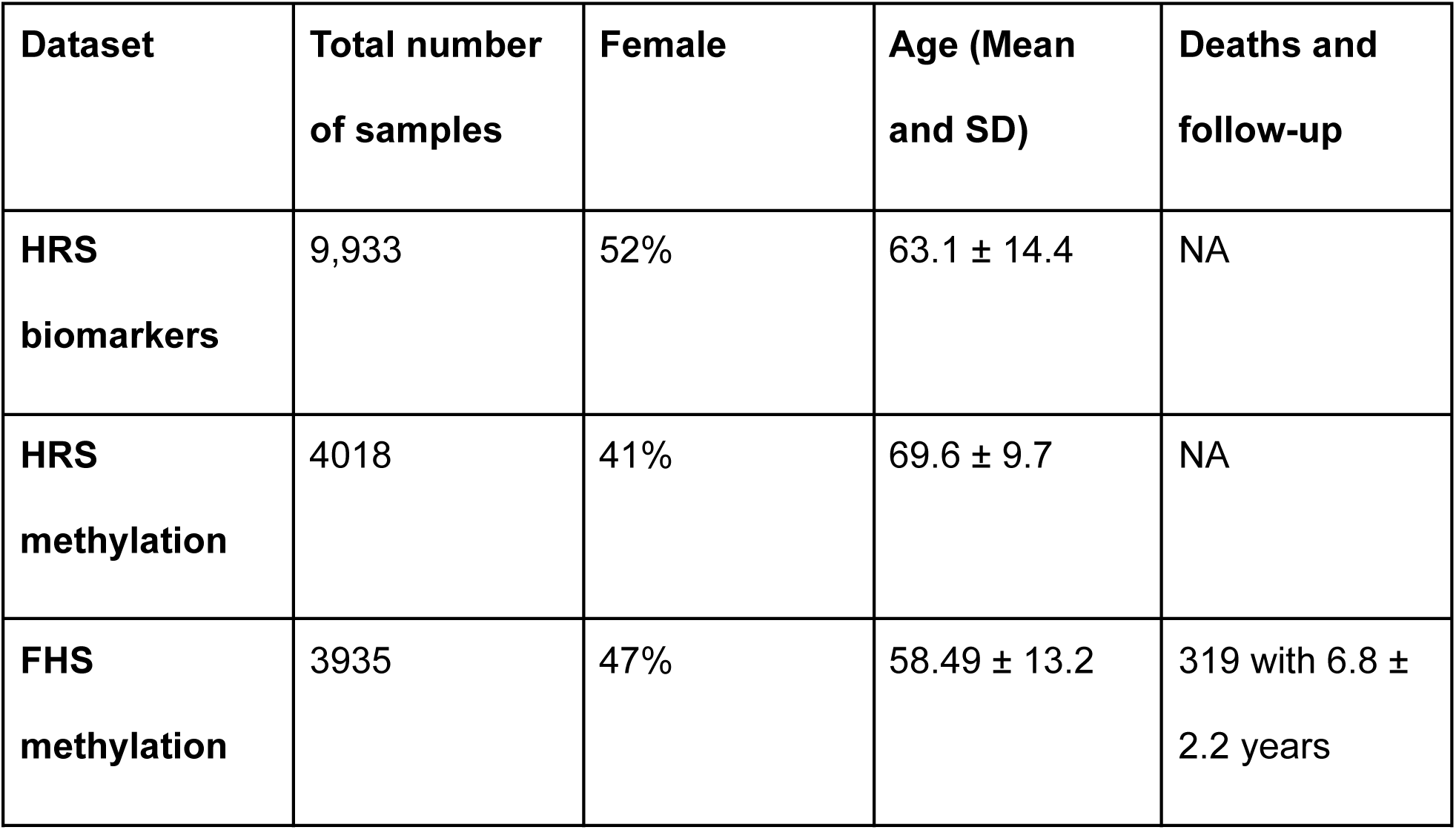
Datasets used for Systems Age training, including information on total number of samples, female percentage, age distribution, deaths and follow-up years.

| **Dataset** | **Total number of samples** | **Female** | **Age (Mean and SD)** | **Deaths and follow-up** |
| --- | --- | --- | --- | --- |
| **HRS biomarkers** | 9,933 | 52% | 63.1 ± 14.4 | NA |
| **HRS methylation** | 4018 | 41% | 69.6 ± 9.7 | NA |
| **FHS methylation** | 3935 | 47% | 58.49 ± 13.2 | 319 with 6.8 ± 2.2 years |

### Systems Age pipeline

**Glossary 1:** terms used to describe Systems Age and intermediate values derived during Systems Age calculation.

| **Term** | **Definition** |
| --- | --- |
| Methylation PCs | Principal components generated by performing PCA on methylation data with 125K CpGs from HRS |
| System biomarker PCs | Principal components generated by performing PCA on groups of Biomarkers in HRS |
| DNAm system PCs | Methylation PC based surrogates of system biomarker PCs calculated in HRS |
| System scores | Cox elastic net regression model scores generated by combining DNAm system PCs for each system calculated in FHS |
| System score PCs | Principal components generated by performing PCA on the 11 system scores plus a chronological age prediction score in FHS |
| Systems Age | Whole body aging score generated by combining system score PCs using a cox elastic net regression model |

#### Step 1: Grouping biomarkers into systems

We utilized molecular and cellular biomarker data from the Health and Retirement Study (HRS) 2016 Venous Blood Study (VBS), for which a subset also has paired DNA methylation data. We assessed the available biomarkers, and manually annotated them as biomarkers for specific physiological systems, totaling 11 systems. To each system we added functional biomarkers (e.g. grip strength) and system-specific disease and condition history (e.g. history of stroke or chronic lung disease).

Our goal was to develop epigenetic aging clocks that are interpretable in terms of physiological systems for clinical and epidemiological applications. Thus, for manual annotation, we required biomarkers to fulfill at least one of two criteria to be assigned to a system: 1) Is there evidence that the biomarkers predict risk of age-related diseases for that physiological system? 2) Would a clinician utilize the biomarker in assessing the status of that physiological system? Annotations were done by multiple team members supported by literature searches to validate disease prediction and clinical interpretations. The team included a gerontologist (M.E.L.) and a physician-scientist (A.H.C.). Most of the biomarkers were transformed and thresholded such that their distribution is more normal. The biomarker-to-system mapping, dataset-specific variable names, and transformations used can be found in Supplementary Table 1.

There is no gold standard list of biomarkers for each physiological system and there is often not a clear delineation between systems because of their biological integration. We do not claim that these are the only 11 systems or the only correct mapping of the Biomarkers to these 11 systems. The Systems Age pipeline can be easily adapted to other biomarker-to-system mappings. Our work here is intended as a proof-of-concept that omics clocks can capture aging in specific physiological systems, and thus the most important validation of our chosen mappings is the high specificity of the System scores in our WHI validation dataset, rather than the exact list of starting biomarkers.

#### Step 2: Principal component analysis of system biomarkers and DNA methylation data

We previously found that performing principal component analysis (PCA) and then using principal components (PCs) as input into supervised machine learning models produces more robust and reliable epigenetic clocks ^25^. PCA removes collinearity, reduces dimensionality of the data, and better separates signal from technical noise. Thus, for each system, we performed PCA on the selected system biomarkers. Before performing PCA, the biomarkers were first transformed to have a normal distribution as described in Supplementary Table 1 as well as scaled before inputting into the prcomp function (stats 4.1.1) in R. In parallel, we performed PCA on DNA methylation data as previously described ^25^, utilizing 125,175 CpGs that 1) are in all of our training and validation data and 2) present on commercially available methylation arrays including the Infinium HumanMethylation450 BeadChip and Infinium Methylation EPIC BeadChip. Practically, this was done using the prcomp function in R. This yielded two sets of PCs (see Glossary 1 for terms): 1) system biomarker PCs (the number of PCs per system is equivalent to the number of biomarkers for each system, since number of samples is greater than number of features) and 2) 4,017 DNA methylation PCs (one less than number of samples, since the number of samples is less than number of features). We did not filter out low-variance PCs (for example using scree plots or random matrix theory methods). Low-variance PCs can still capture relevant variation for prediction (Jolliffe 1982; Yan et al. 2020; Aschard et al. 2014; Tarashansky et al. 2019), while those that are irrelevant are removed or minimized at later supervised machine learning steps. Thus, when predicting system biomarker PCs from DNA methylation PCs (Step 3), we can predict both dominant, shared signals between biomarkers (high-variance PCs) as well as more subtle variations.

#### Step 3: Building DNAm surrogates of system PCs

We utilize elastic net regression to train a model using methylation PCs to predict each system biomarker PC using the glmnet 4.1-4 package in R. We refer to the resulting models as DNAm system PCs. This was done as described previously ^25^. The L1 to L2 regularization ratio was 1 (α = 0.5), the λ tuning parameter was selected via tenfold cross-validation, and the final methylation PC was excluded as it is not meaningful in cases where the number of samples is less than the number of features. Not all system PCs are predicted well using methylation PCs. We retained DNAm system PCs with at least 20 DNAm PCs being used at the minimum mean cross-validated error in the model and at least 5 DNAm PCs at the cross-validated error one standard error from the minimum mean cross-validated error in the model. All selected PCs by the above method had a correlation coefficient greater than 0.4 in HRS.This allows us to take only well predicted DNAm system PCs to the next step.

#### Step 4: Building system scores

To build system scores we first calculate DNAm system PCs in FHS based on parameters previously trained in HRS (first calculating methylation PCs, then predicting system PCs). Then, for each system separately, we predicted mortality using DNAm system PCs in a Cox elastic net mortality prediction model using the glmnet 4.1-4 package. The L1 to L2 regularization ratio was 1 (α = 0.5), the λ tuning parameter was selected via tenfold cross-validation. This yielded 11 separate mortality prediction models that we term system scores, and can serve as a measure of mortality-related deterioration of each system.

#### Step 5: Building Systems Age

To build Systems Age, we first perform PCA on the DNAm system scores and age prediction score using the prcomp(stats 4.1.1) function in R, as the system scores and age prediction score are partially correlated with one another.

The age prediction score is built specifically to predict chronological age and was trained in HRS. The DNAm PCs in HRS were first used to predict chronological age. The scores thus generated were then used to predict chronological age again but instead now using a second degree polynomial function fitted to the 5 year interval averages of the predicted chronological age score (previous step) predicting for the 5 year interval averages of chronological age. The score obtained from the second degree polynomial is referred to as age prediction in our model.

Using all system score PCs, we then predict mortality using another Cox elastic net mortality prediction model using glmnet 4.1-4 package in R. The L1 to L2 regularization ratio was 1 (α = 0.5), the λ tuning parameter was selected via tenfold cross-validation, Again, using PCs as input is intended to reduce redundancy, increase reliability, and allow for more subtle variations in system scores to have an important role in the overall model.

#### Step 6: Scaling scores to age range

The 11 system scores and Systems Age are first standardized to have mean 0 and standard deviation 1. They are then scaled to match the mean and standard deviation of chronological age for the 3935 samples from FHS Offspring and Gen3 cohorts. It should be noted that prescaling all system scores and Systems Age had a correlation between 0.8 and 0.34 in healthy patients of BLSA. (Supplementary Figure 9) All correlations were positive and in the moderate to high correlation range. BLSA has a mean of 69 and SD of 13, age range 24-98. These correlations were comparable to the system-specific predictors reported by Oh 2023, Nature (proteomics; correlations range from 0.34 to 0.73) and Tian 2023, Nature Medicine (clinical phenotypes; correlations range from 0.3 to 0.8).

### Using z-scores as metric for comparison in association analysis

For all models, we list Hazard Ratios, Odds Ratios and Beta Coefficients in our Supplementary Tables. However, we utilize z-scores as our primary metric for comparing the association of different clocks with different outcomes, for the following reasons:

1. Z-scores track with effect sizes: Z-scores even though do not directly measure effect size however, a stronger effect size leads to a more significant z-score. Z-scores measure the significance of the coefficient in our models and the stronger the coefficient the more likely it is to have a higher z-score.
2. Uniformity across models: We use a wide variety of models ranging from linear to cox to logistic and from fixed effects to random effects to meta-analysis. HRs and ORs are on the exponential scale, while beta coefficients are on the linear scale, which makes it challenging to compare across models. Instead, z-scores provide a standardized measure across all these models which allows for comparison.
3. Standardization: Metrics such as beta coefficients are not standardized, meaning they are influenced by the scale of the variables involved in the regression model. Standardizing scores requires an additional step of analysis and it is not always clear how best to scale to compare different clocks and different outcomes. On the other hand, z-scores are already standardized, providing a standardized measure of the magnitude of the effect relative to the variability of the data. This standardization can be helpful for comparing the magnitudes of effects across different predictors or models.

Statistical Inference: z-scores are directly associated with p-values, which indicate the statistical significance of coefficients. This makes them particularly useful for assessing the significance of predictors in regression models. Beta coefficients, HRs and ORs, while informative about the magnitude of the effect, do not directly convey information about statistical significance.

### Association meta-analysis in Women’s Health Initiative cohorts

The Women’s Health Initiative (WHI) is a long-term national health study ^53^. WHI is funded by the National Heart, Lung, and Blood Institute, or NHLB and ran from the early 1990s to 2005. Post 2005, there have been Extension Studies, which continue to collect data on health outcomes annually. We used 3 WHI cohorts which had methylation data available. In each WHI cohort (Table 2) we calculated system scores then regressed all epigenetic aging clocks on chronological age using a linear regression model and defined clock age acceleration as the corresponding residual. We then calculated associations between these clock accelerations and different diseases and aging phenotypes in all WHI cohorts. We stratified the cohorts by race (except WHI AS311 where analysis of the Black and Hispanic populations would be underpowered), for a total of 7 groups. Depending on condition and disease we either built linear regression models (cognitive function, physical function, comorbidities and more), cox prediction models (Lung Cancer, Breast Cancer, Leukemia, CHD, MI and more) or logistic regression models (Thyroid disease, Diabetes, Arthritis, Cataract and more) to look at associations with the age accelerated scores. An example of the formula used is as follows: *Cognitive function ∼ AgeAccel + Age*. In certain cases, additional factors such as Education level (for Cognitive function) were also added to the models. Sex was not a covariate as all WHI participants are female. We combined the associations from the different cohorts and racial groups in a fixed effects model meta-analysis with inverse variance weights, obtaining meta-analysis z-scores for the associations (package metafor 2.4-0, function rma and forest). Forest plots, z-scores, heterogeneity p-values and other meta-analysis results are provided in Supplementary Figure 3 and 4 and Supplementary Tables 2-5.

**Table 2:**
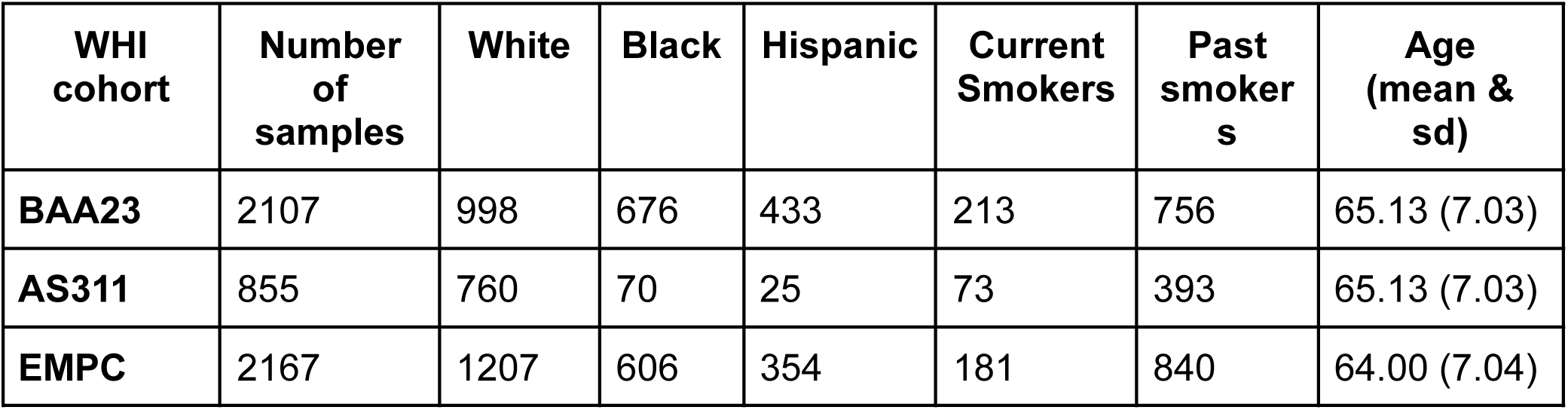
WHI cohorts used for testing with distribution of race, ethnicity, sex, smokers and age.

| WHI cohort | Number of samples | White | Black | Hispanic | Current Smokers | Past smokers | Age (mean & sd) |
| --- | --- | --- | --- | --- | --- | --- | --- |
| **BAA23** | 2107 | 998 | 676 | 433 | 213 | 756 | 65.13 (7.03) |
| **AS311** | 855 | 760 | 70 | 25 | 73 | 393 | 65.13 (7.03) |
| **EMPC** | 2167 | 1207 | 606 | 354 | 181 | 840 | 64.00 (7.04) |

We performed additional analyses adjusting for smoking status by adding smoking status (present smoker, ex-smoker or never smoked) into the linear (package stats 4.0.2, function lm), Cox (package survival 3.2-11, function coxph) and logistic models (package stats 4.0.2, function glm(family=”binomial”)). We also examined non-smokers separately. A list of the variables used from WHI are shown in Table 3. It is important to note that even though multiple disease variables were available we could not test for a majority of the variables because they were underpowered. Rather, for many of the variables we calculated z-scores and showed them in our supplementary data. Variables which had insufficient N have not been listed below. (Table 3)

**Table 3:**
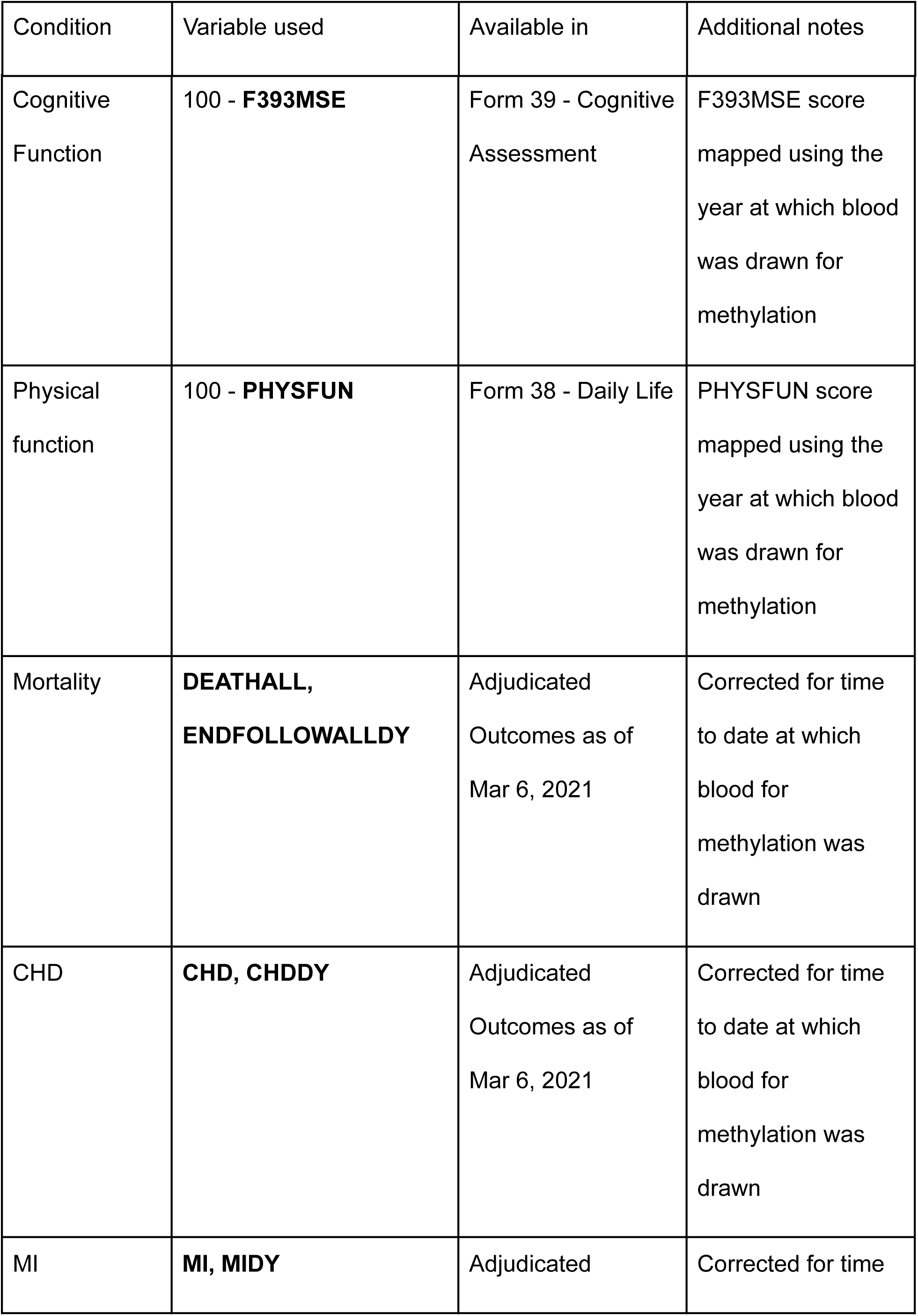
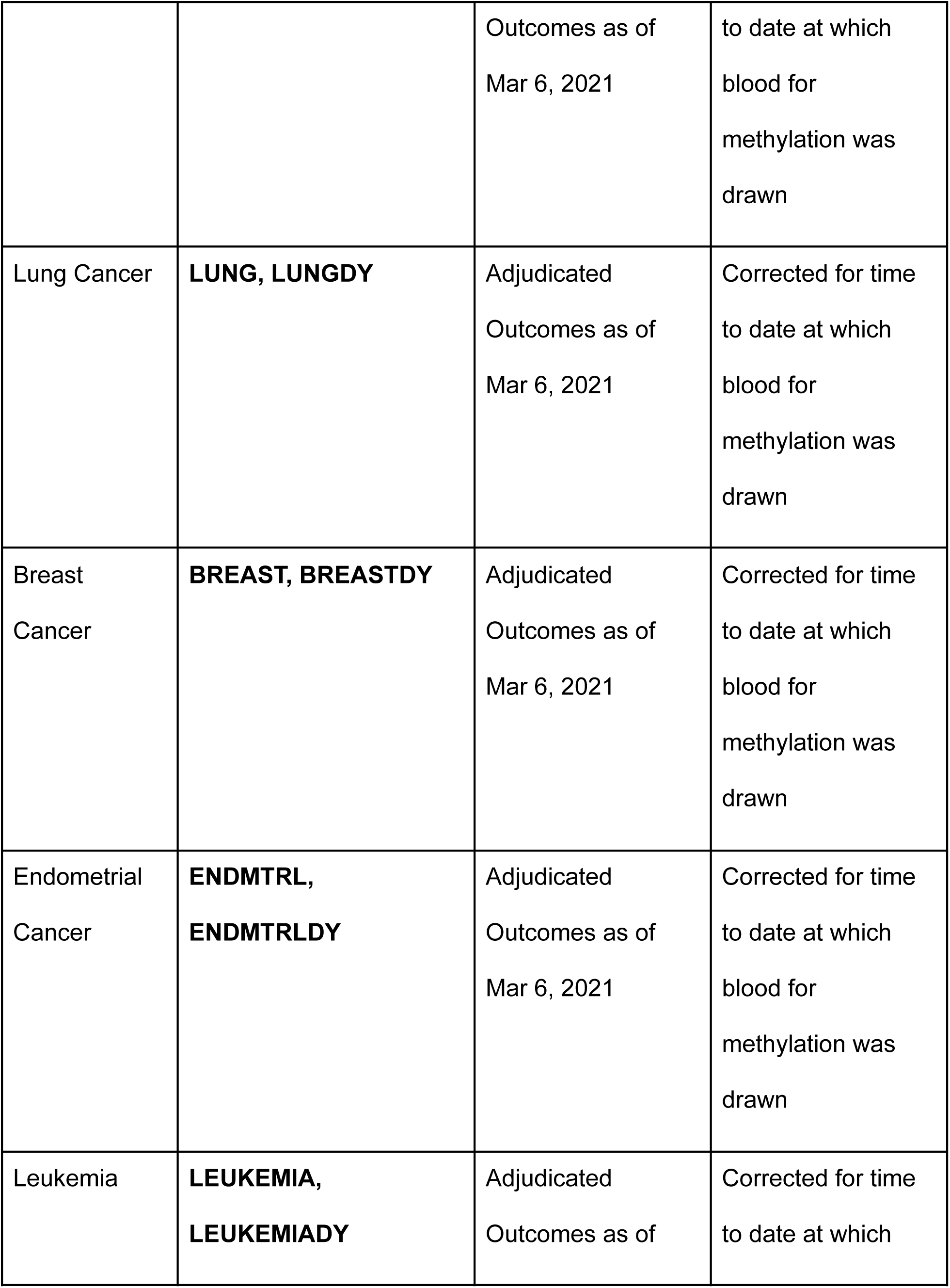
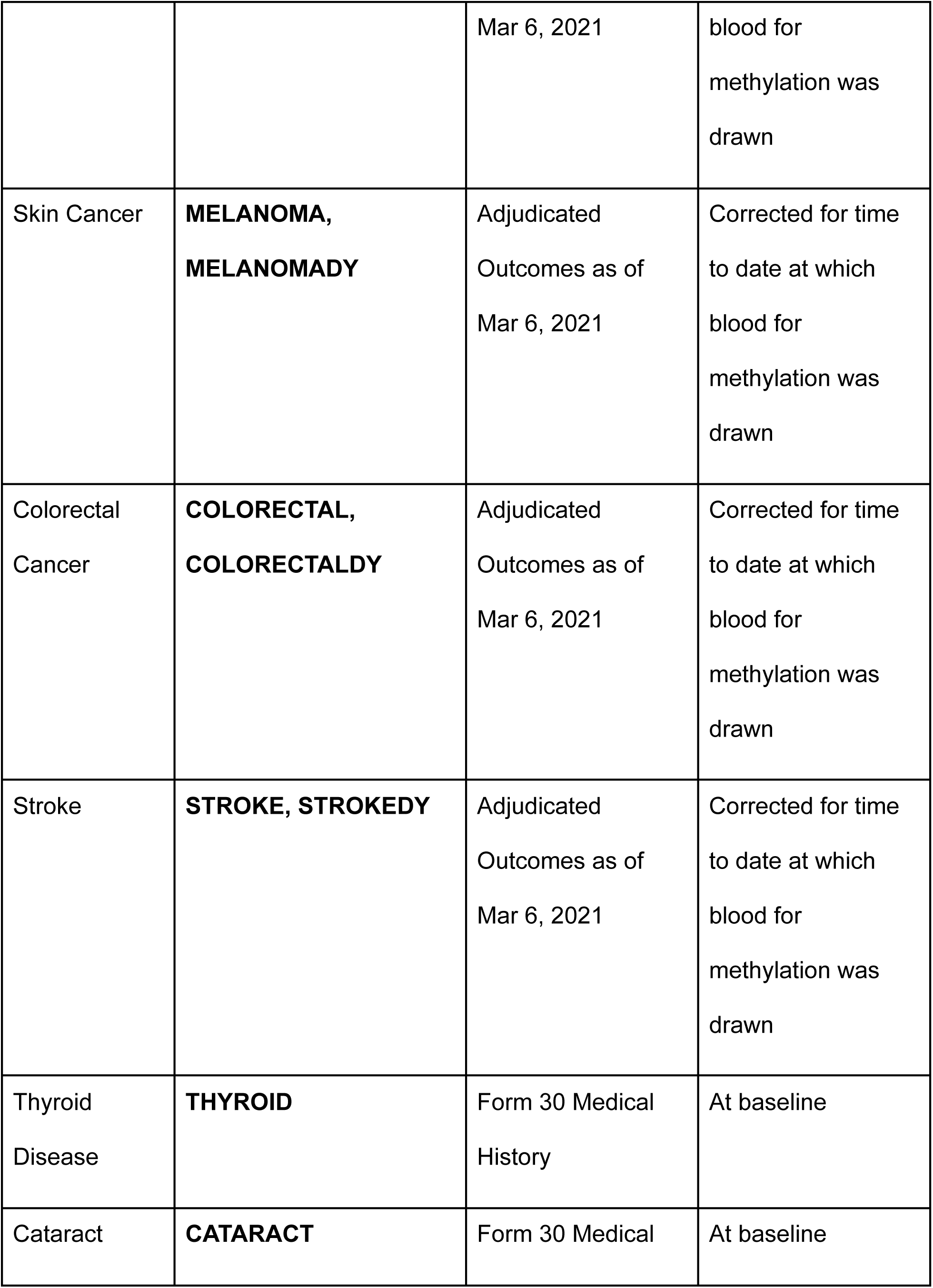
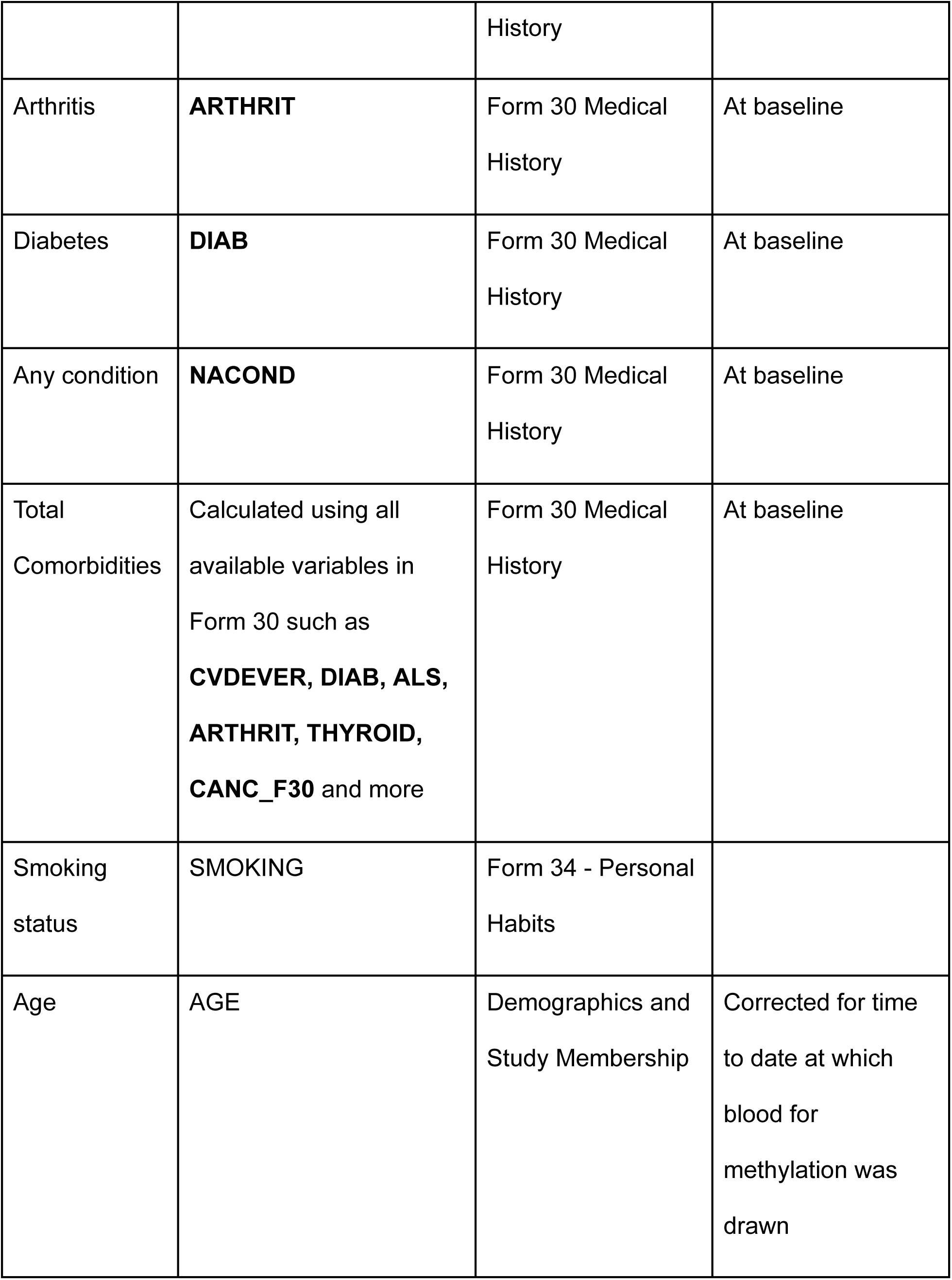
WHI variables used for testing associations of scores.

### ROC meta-analysis in Women’s Health Initiative cohorts

For performing ROC meta analysis, we first converted all our variables to binary format. Continuous variables such as cognitive function and physical function were converted to binary variables by marking the lowest quintile as diseased and the rest as healthy. For total comorbidities we marked greater than 2 comorbidities as diseased. For time to event variables such as mortality, CHD, MI and more, we censored at t=6,000 days. The censoring time was determined manually by looking at the longest follow-up times for event-free individuals. Models for ROC analysis were built including the score along with age. Each ROC curve for a clock was either compared with Systems Age or best system score using roc.test() function in the pROC package in R. Meta-analysis was then performed by analyzing z-scores comparing each clock to Systems Age or best system score for all cohorts using Stouffer’s method.

### Association analysis in BLSA

The Baltimore Longitudinal Study of Aging (BLSA) is America’s longest-running scientific study of human aging. The study began in 1958. We built logistic regression models to look at associations between the age accelerated scores and disease status at baseline for Cancer, Diabetes and CAD. We adjusted for age, sex and smoking status, with an additional sensitivity analysis adjusting for race. An example of the formula used is as follows: *CAD ∼ AgeAccel + Age + Sex + Smoking_Status*. All z-scores and ORs are provided in Supplementary Table 12. Demographics are provided in Table 4.

**Table 4:** Additional cohorts used for testing with distribution of race, sex, smokers and age.

| **Cohort Study** | **Number of samples** | **Female** | **Black** | **Current Smokers** | **Past smokers** | **Age (mean & sd)** | **Age Range (years)** |
| --- | --- | --- | --- | --- | --- | --- | --- |
| **BLSA** | 820 | 403 | 216 | 17 | 338 | 69.1 (13.6) | 24 - 98 |
| **SATSA** | 535 | 312 | 0 | 95 | 20 | 68.2 (9.5) | 48 - 94 |
| **ADNI** | 1696 | 758 | 9 | - | - | 75.4 (7.64) | 55 - 91.4 |

### Association analysis in SATSA

The Swedish Adoption Twin Study of Aging was started in 1984 and has previously been described.^29^ For the purpose of our analysis we analyzed methylation data from Infinium HumanMethylation450 BeadChips and EPIC array, available for 535 individuals. All participants in SATSA provided written informed consents and the study was approved by the ethics committee at Karolinska Institutet with Dnr 2015/1729-31/5.

We built a Cox mixed effects model with twin pair ID as a random effect, and smoking status, sex, age, and stratified birth cohort as the fixed effects. All z-scores and HRs are provided in Supplementary Table 12. Demographics are provided in Table 4.

### Association analysis in ADNI

The Alzheimer’s Disease and Neuroimaging Initiative began in 2004 as a private-public partnership with 20 companies along with the National Institutes of Health and the National Institute on Aging. The initial five-year study (ADNI-1) was extended by two years in 2009 by a Grand Opportunities grant (ADNI-GO), and in 2011 and 2016 by further competitive renewals of the ADNI-1 grant (ADNI-2, and ADNI-3, respectively). For our analysis we use the complete ADNI dataset with DNA methylation data publicly available up to and including ADNI-2 cohort. We have performed 3 different types of analysis in the ADNI cohort. DNA methylation was profiled in blood or buffy coat samples using Illumina EPIC chips according to the Illumina protocols. A detailed protocol describing DNA measurement has been described previously.^54^

First, association only in AD patients where we associate brain volume measures such as whole brain volume, hippocampal volume and fusiform volume along with cognitive measures such as ADAS13, MMSE and MOCA. Since we have longitudinal data for the same individuals we used linear mixed effects models where age, sex and education status are the fixed effects and ADNI ID is the random effect. We used the lme4 package in R for this purpose.

In our second analysis, we associated all the above mentioned variables but this across all ADNI subjects which included healthy, MCI and AD categories. We again used a linear mixed effects model as described above to perform this analysis.

In the last analysis, we compared scaled age deviation values of clocks across each diagnostic category. For this purpose we only used baseline samples for each individual and their diagnosis category. P-values were determined using anova test performed in Prism 9.

### Multiple Testing

Given that we were testing for multiple system scores as well as whole body clocks (15 in total), we decided to perform multiple testing corrections across all the different tests we performed for a given disease or condition related analysis. We chose to use Benjamin-Hochberg correction instead of using Bonferroni correction. Our choice was dictated by the fact that Bonferroni correction assumes the tests are independent of each other. If the tests are correlated, Bonferroni correction may be overly conservative. We know given our correlation plot that some scores are indeed correlated and which is why we decided to go with a relatively less conservative multiple testing method that is the benjamin-hochberg correction. We first converted the Z-scores to P-values and then used the p.adjust function from the stats package in R for this purpose.

Almost all results have significant p-values after multiple testing corrections and the results which did not have been specifically mentioned in the results.

### Calculating different clocks

In addition to Systems Age, we calculated a large number of additional existing clocks for comparison. We used the following packages or sources to do so (Table 4).

**Table 4:**
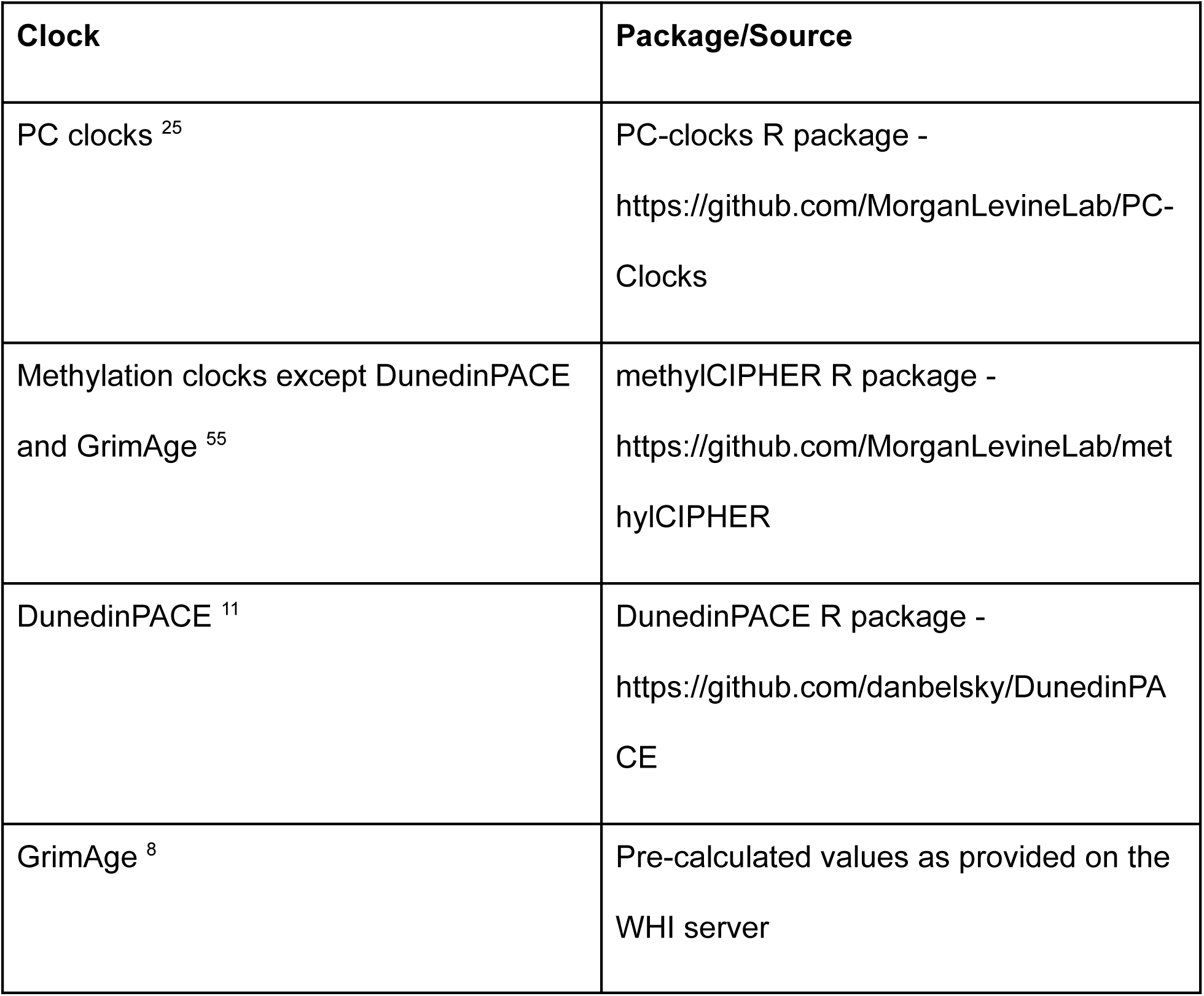
Packages used for calculating epigenetic clocks.

### Aging Subtypes and overrepresentation of diseases in subtypes

Age-adjusted system scores were used to perform adaptive hierarchical clustering using the Dynamic Tree Cut library (dynamicTreeCut 1.63-1, function cutreeDynamicTree) in R. Parameters used other than default settings included minModuleSize which was set at 100. Based on the most stable node distance, 9 clusters were identified. Average score for each system for each cluster was plotted on polar spider plots. An over representation analysis comparing occurrence of disease in the cluster compared to the whole population was performed using Fisher’s exact test. Binary disease status variables were used without transformation, continuous variables such as cognitive function and physical function were converted into binary variables by marking values lesser than 1 standard deviation from mean as disease states. For time-to-event variables, the model was built only for individuals who were alive until the 7 year follow-up or died because of the condition.

### Association of Biomarkers with System specific scores in HRS

We wanted to show how the different Biomarkers used in training were associated with the methylation based system scores. For this purpose we built linear regression models associating the system score with each biomarker. The z-score of association was then plotted on the y axis for each score in supplementary figure 1.

### Test-retest reliability analysis

Reliability was calculated as described before ^25^. Briefly, reliability was calculated in GSE55763 which consisted of 36 whole-blood samples measured in duplicate (age range 37.3 to 74.6). We used the icc function in the irr R package version 0.84.1, using a single-rater, absolute-agreement, two-way random-effects model ^56^.

