## Supplementary Figures for "Systems Age: A single blood methylation test to quantify aging heterogeneity across 11 physiological systems"

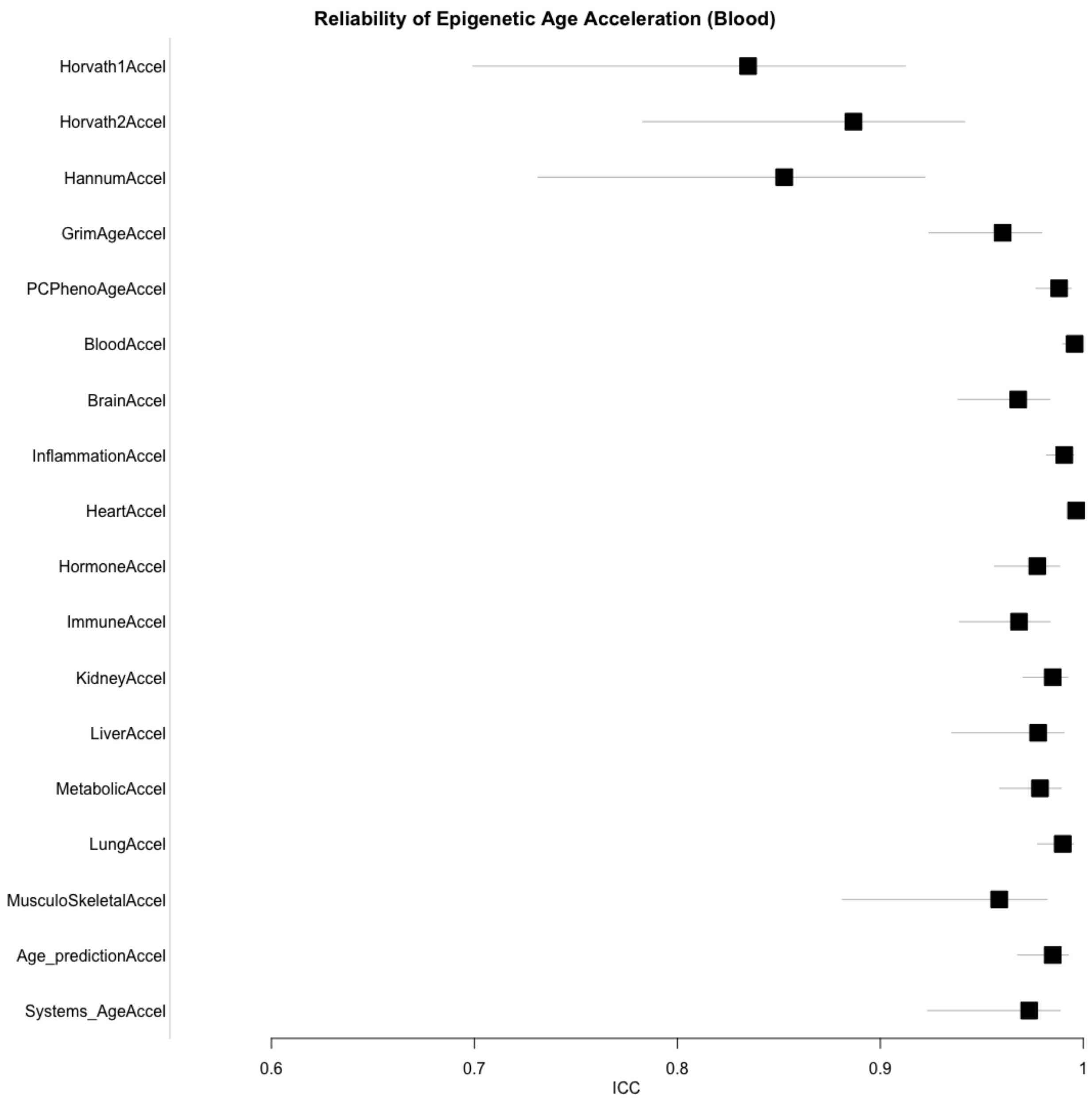
Reliability of Epigenetic Age Acceleration (Blood)
Horvath1Accel
Horvath2Accel
HannumAccel
GrimAgeAccel
PCPhenoAgeAccel
BloodAccel
BrainAccel
InflammationAccel
HeartAccel
HormoneAccel
ImmuneAccel
KidneyAccel
LiverAccel
MetabolicAccel
LungAccel
MusculoSkeletalAccel
Age_predictionAccel
Systems_AgeAccel
0.6
0.7
0.8
0.9
1
ICC

*Supplementary figure 1: Reliability of different age accelerated system scores and Systems Age as compared to other clocks. Reliability was calculated as described before (Higgins-Chen et al. 2022). Briefly, reliability was calculated in GSE55763 which consisted of 36 whole-blood samples measured in duplicate (age range 37.3 to 74.6). We used the icc function in the irr R package version 0.84.1, using a single-rater, absolute-agreement, two-way random-effects model.*

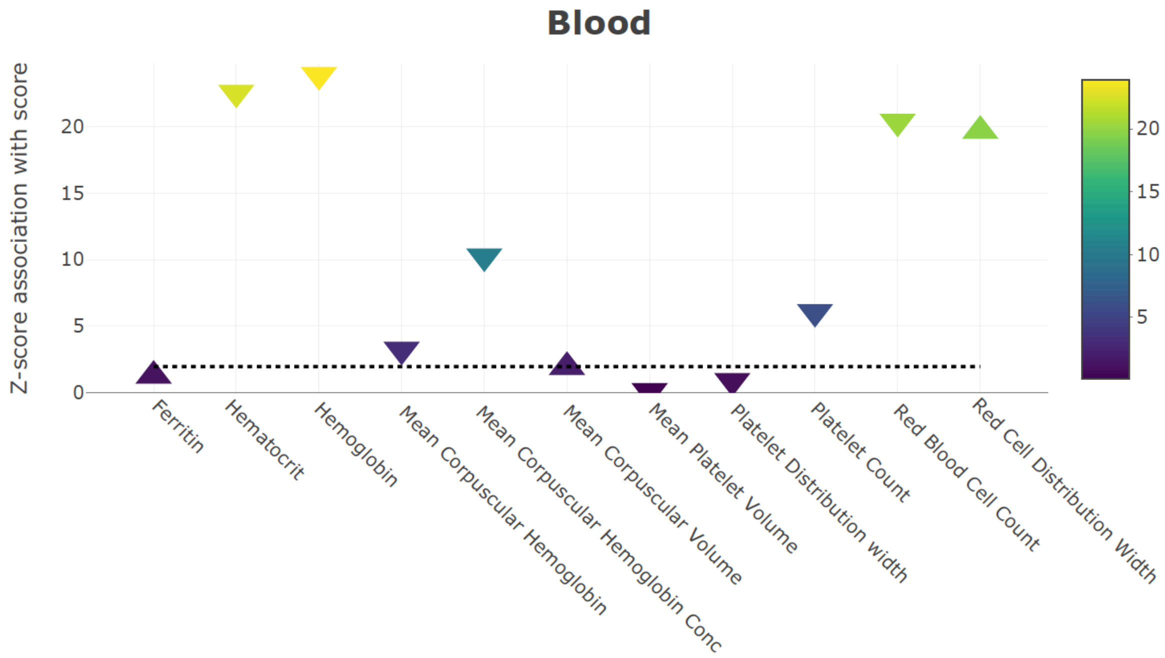
Blood
Z-score association with score
0
5
10
15
20
Ferritin
Hematocrit
Hemoglobin
Mean Corpuscular Hemoglobin
Mean Corpuscular Hemoglobin Conc
Mean Corpuscular Volume
Mean Platelet Volume
Platelet Distribution width
Platelet Count
Red Blood Cell Count
Red Cell Distribution Width

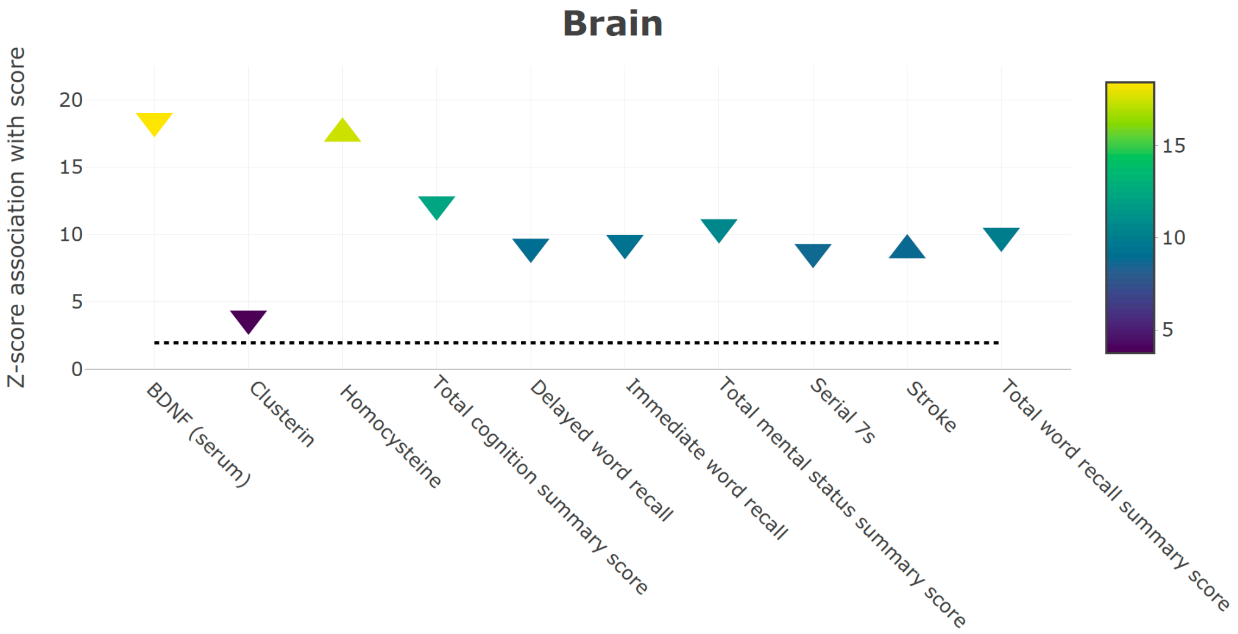
Brain
Z-score association with score
0
5
10
15
20
BDNF (serum)
Clusterin
Homocysteine
Total cognition summary score
Delayed word recall
Immediate word recall
Total mental status summary score
Serial 7s
Stroke
Total word recall summary score

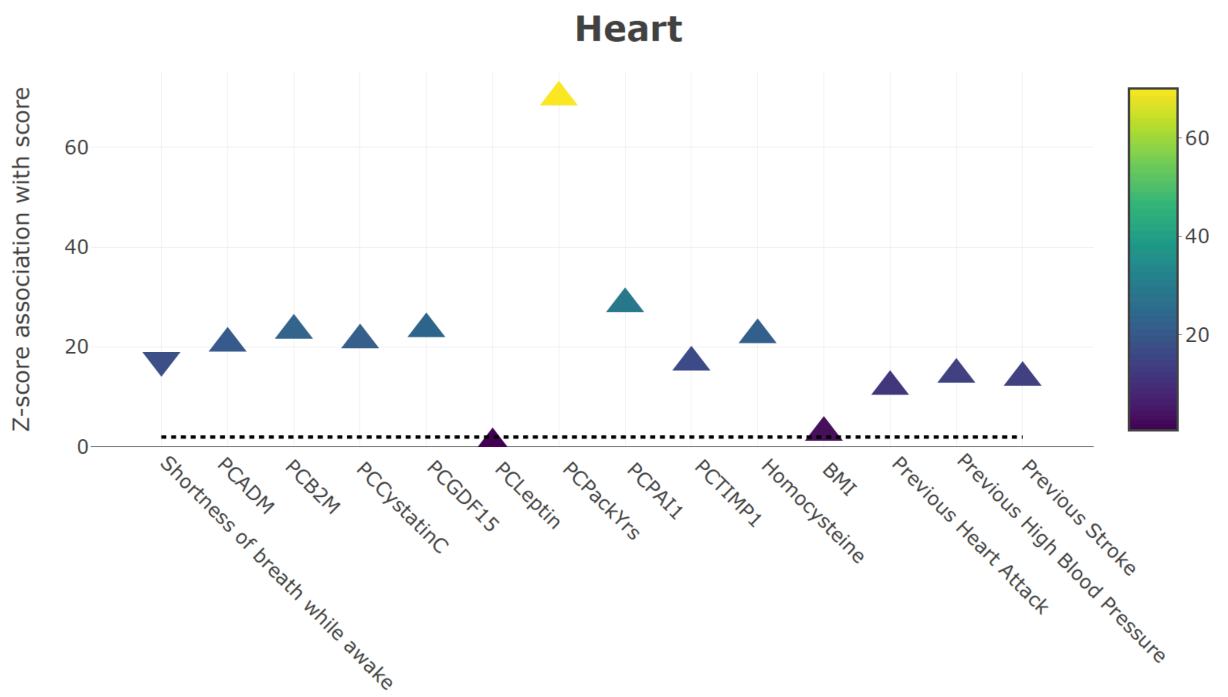
Heart
Z-score association with score
0
20
40
60
Shortness of breath while awake
PCADM
PCB2M
PCCystatinC
PCGDF15
PCLeptin
PCPackYrs
PCPAI1
PCTIMP1
Homocysteine
BMI
Previous Heart Attack
Previous High Blood Pressure
Previous Stroke
20
40
60

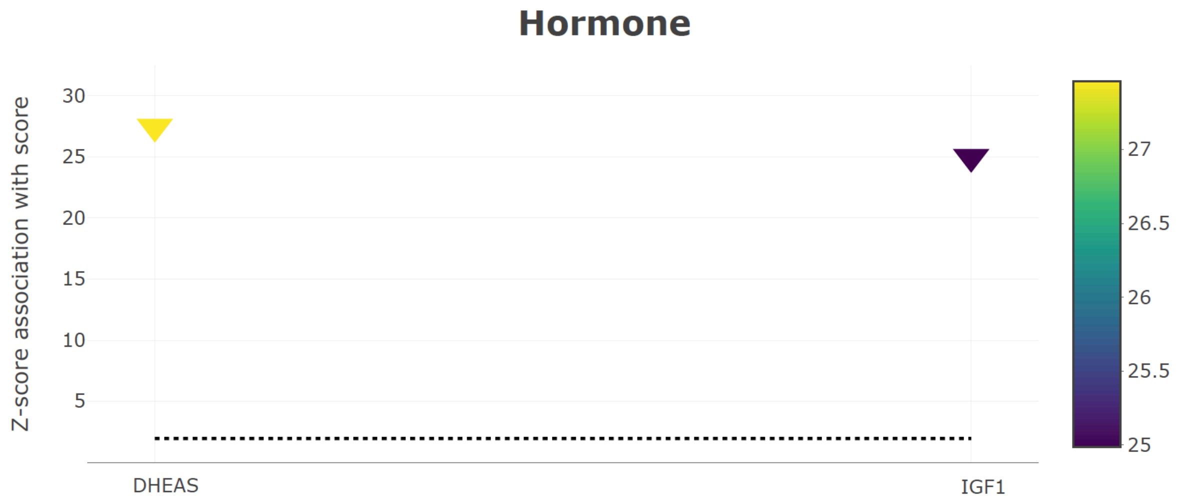
Hormone
Z-score association with score
5
10
15
20
25
30
DHEAS
IGF1
25
25.5
26
26.5
27

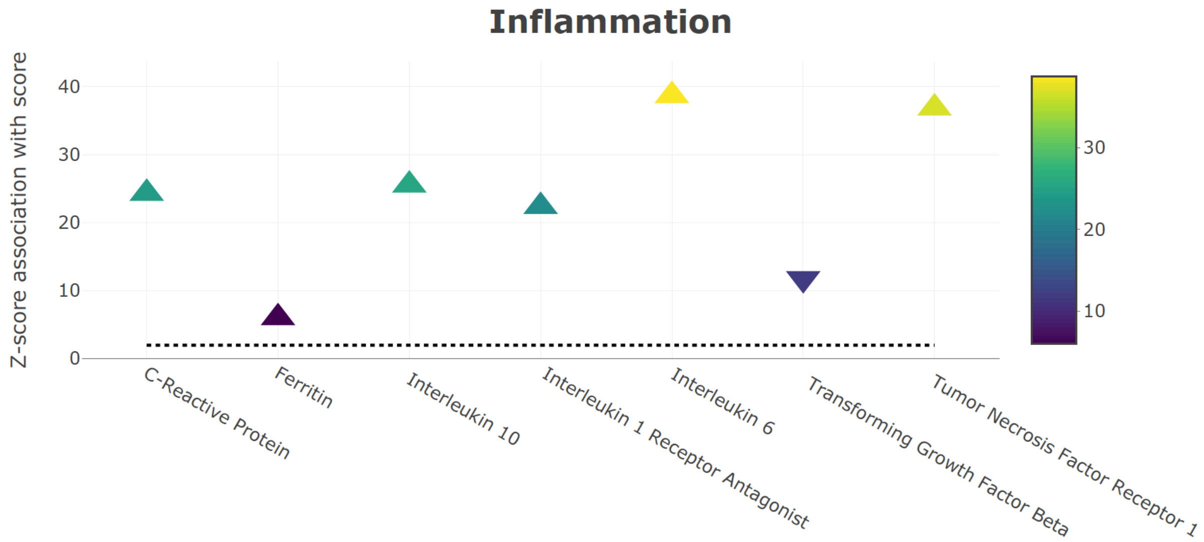
Inflammation
Z-score association with score
40
30
20
10
0
C-Reactive Protein
Ferritin
Interleukin 10
Interleukin 1 Receptor Antagonist
Interleukin 6
Transforming Growth Factor Beta
Tumor Necrosis Factor Receptor 1
30
20
10

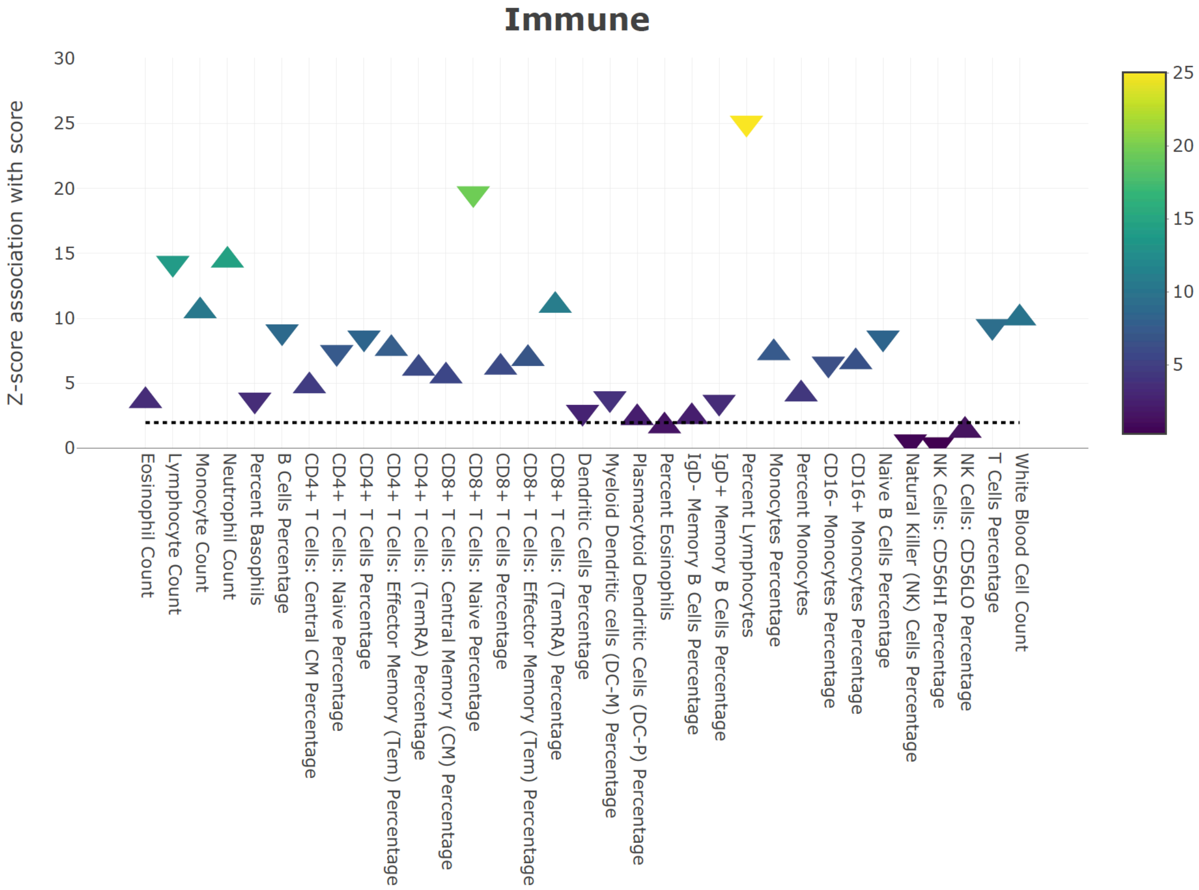
Immune
Z-score association with score
30
25
20
15
10
5
0
Eosinophil Count
Lymphocyte Count
Monocyte Count
Neutrophil Count
Percent Basophils
B Cells Percentage
CD4+ T Cells: Central CM Percentage
CD4+ T Cells: Naive Percentage
CD4+ T Cells Percentage
CD4+ T Cells: Effector Memory (Tem) Percentage
CD4+ T Cells: (TemRA) Percentage
CD8+ T Cells: Central Memory (CM) Percentage
CD8+ T Cells: Naive Percentage
CD8+ T Cells Percentage
CD8+ T Cells: Effector Memory (Tem) Percentage
CD8+ T Cells: (TemRA) Percentage
Dendritic Cells Percentage
Myeloid Dendritic cells (DC-M) Percentage
Plasmacytoid Dendritic Cells (DC-P) Percentage
Percent Eosinophils
IgD- Memory B Cells Percentage
IgD+ Memory B Cells Percentage
Percent Lymphocytes
Monocytes Percentage
Percent Monocytes
CD16- Monocytes Percentage
CD16+ Monocytes Percentage
Naive B Cells Percentage
Natural Killer (NK) Cells Percentage
NK Cells: CD56HI Percentage
NK Cells: CD56LO Percentage
T Cells Percentage
White Blood Cell Count
25
20
15
10
5

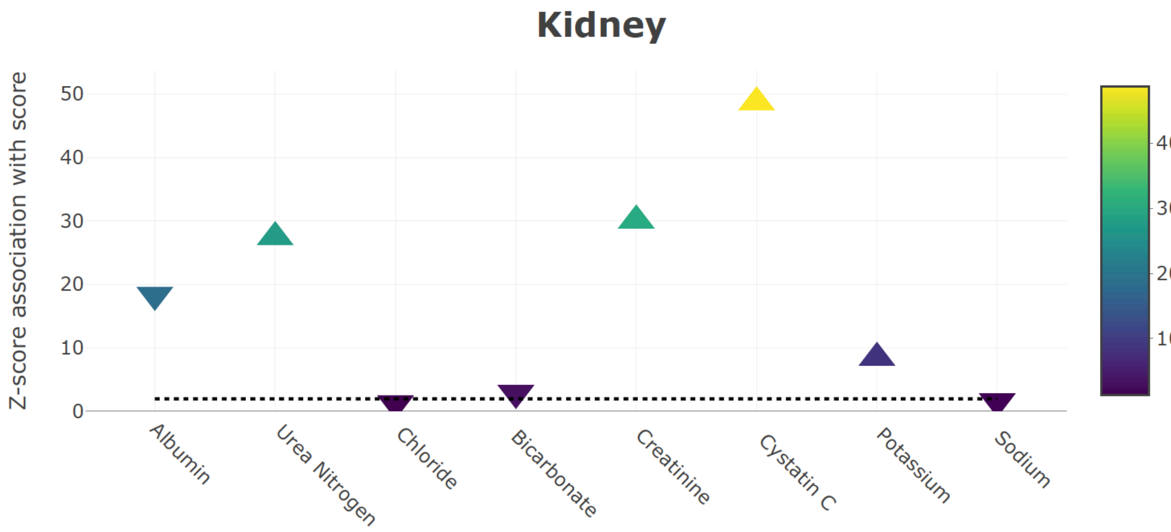
Kidney
Z-score association with score
50
40
30
20
10
0
Albumin
Urea Nitrogen
Chloride
Bicarbonate
Creatinine
Cystatin C
Potassium
Sodium
40
30
20
10

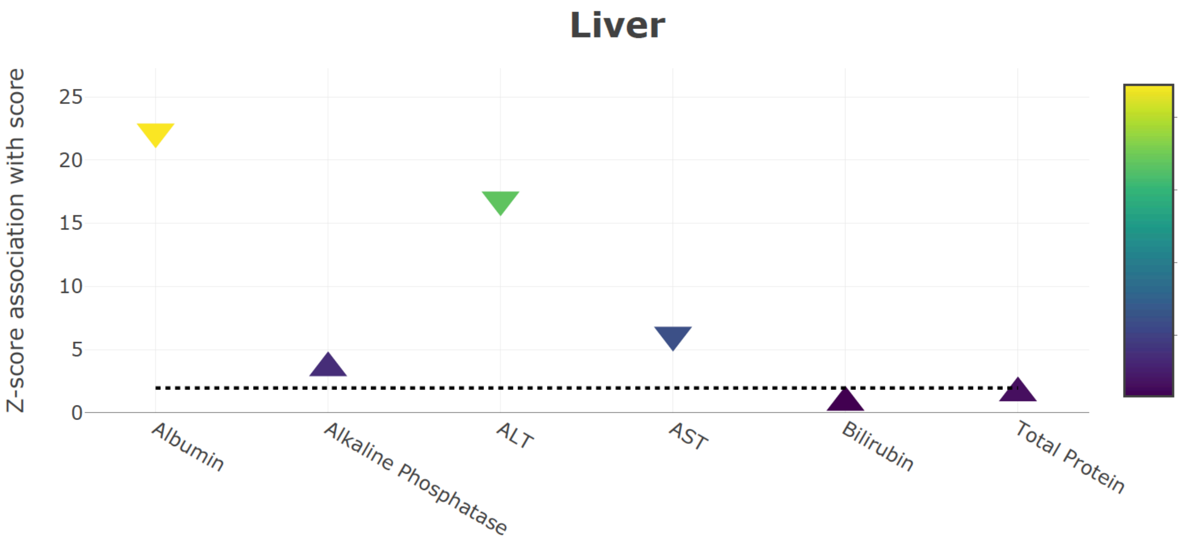
Liver
Z-score association with score
25
20
15
10
5
0
Albumin
Alkaline Phosphatase
ALT
AST
Bilirubin
Total Protein
20
15
10
5

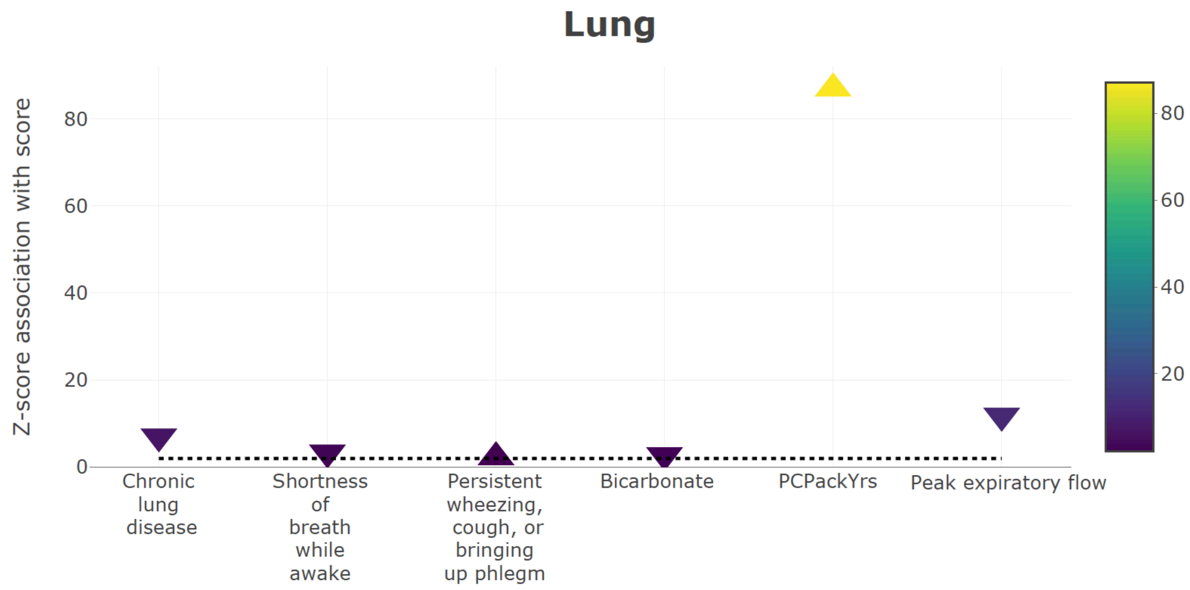
Lung
Z-score association with score
80
60
40
20
0
Chronic lung disease
Shortness of breath while awake
Persistent wheezing, cough, or bringing up phlegm
Bicarbonate
PCPackYrs
Peak expiratory flow
80
60
40
20

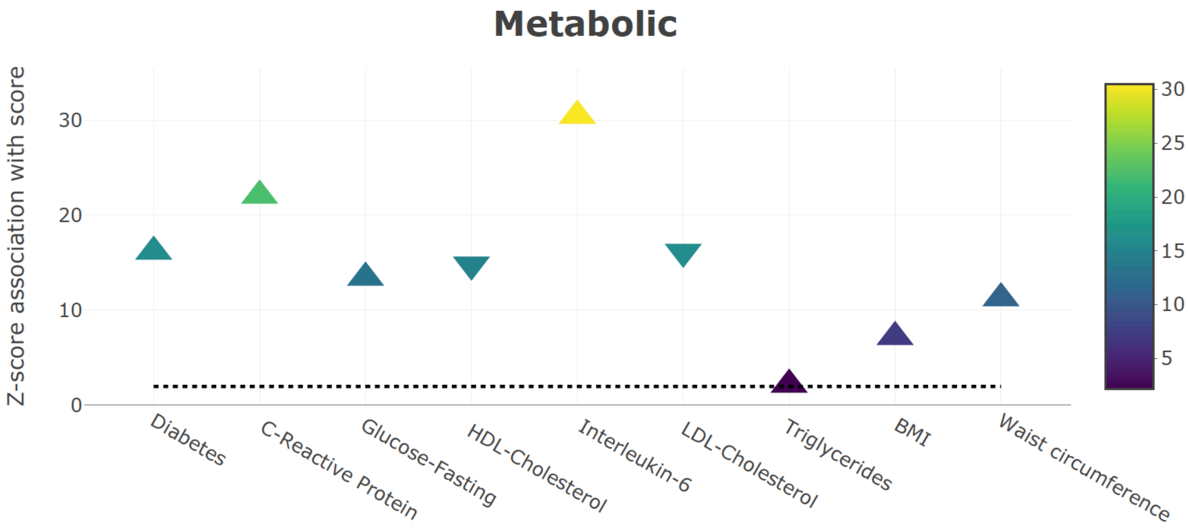
Metabolic
Z-score association with score
30
20
10
0
Diabetes
C-Reactive Protein
Glucose-Fasting
HDL-Cholesterol
Interleukin-6
LDL-Cholesterol
Triglycerides
BMI
Waist circumference
30
25
20
15
10
5

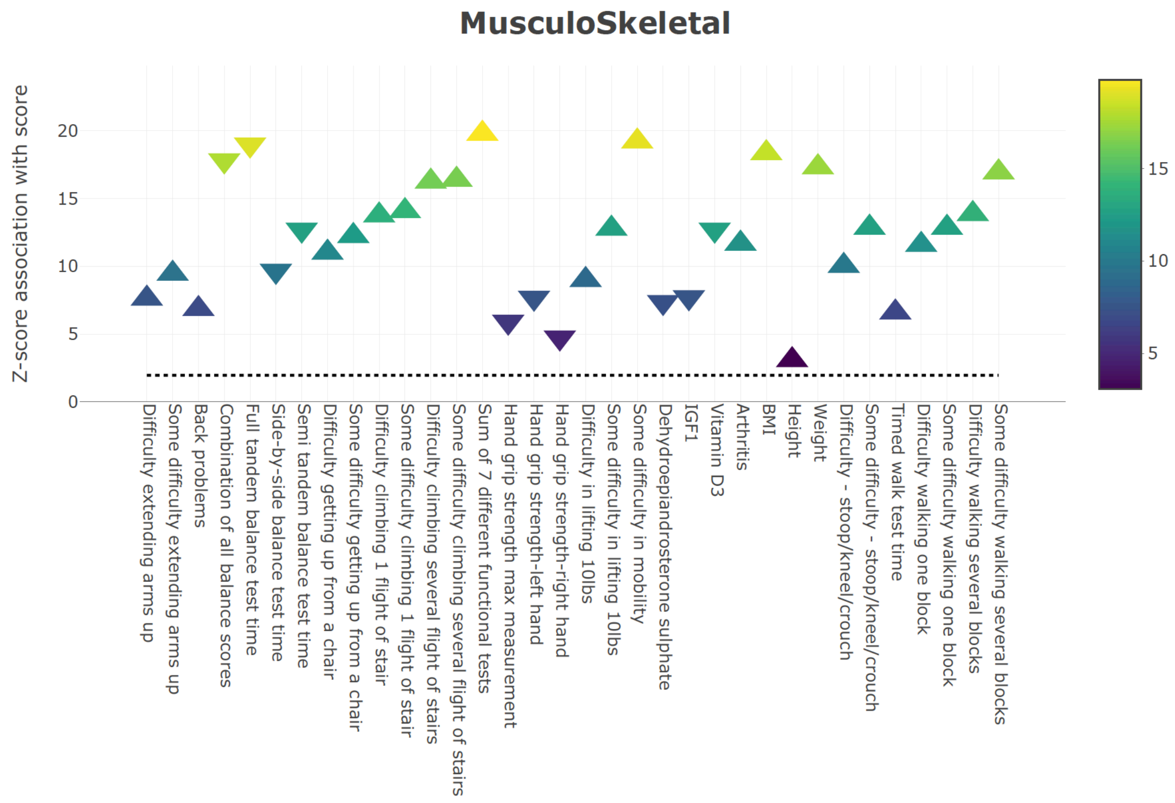


*Supplementary Figures 2 : Associations of Biomarkers with System specific scores in HRS. Each system score was calculated in 3593 samples in HRS. For each of the system scores the respective biomarkers pertaining to each system were associated using a linear model to observe the z-score of association which were then plotted on the above graphs.*

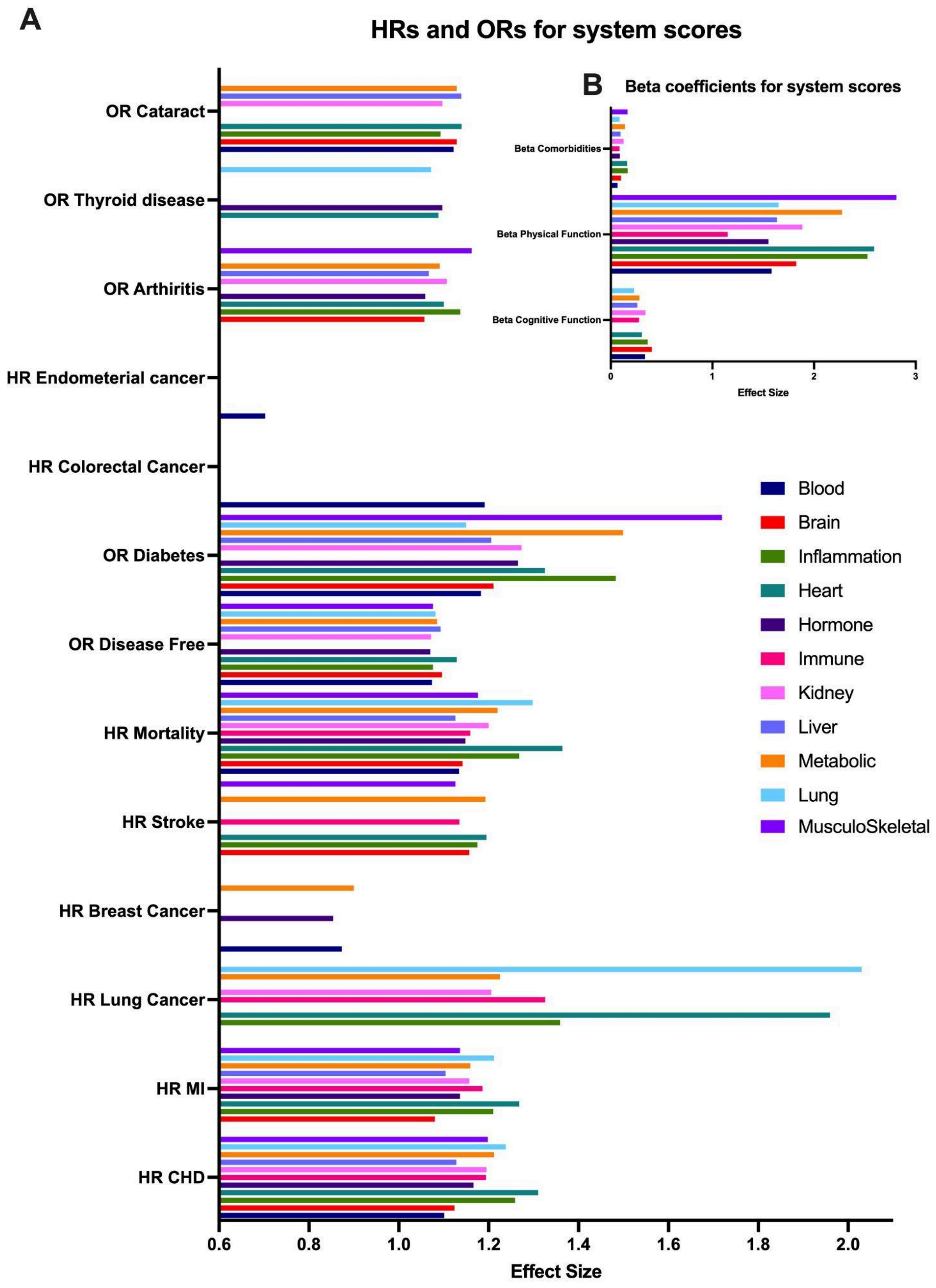
A
HRs and ORs for system scores
OR Cataract
OR Thyroid disease
OR Arthiritis
HR Endometerial cancer
HR Colorectal Cancer
OR Diabetes
OR Disease Free
HR Mortality
HR Stroke
HR Breast Cancer
HR Lung Cancer
HR MI
HR CHD
0.6
0.8
1.0
1.2
1.4
1.6
1.8
2.0
Effect Size
B
Beta coefficients for system scores
Beta Comorbidities
Beta Physical Function
Beta Cognitive Function
0
1
2
3
Effect Size
Blood
Brain
Inflammation
Heart
Hormone
Immune
Kidney
Liver
Metabolic
Lung
MusculoSkeletal

*Supplementary figure 3: Comparing effect sizes for different system scores. Hazard Ratios and Odds Ratios for different models (A) and beta coefficients for different models (B). Only significant effect sizes have been plotted as measured by Z-score. Effect sizes were calculated using a race stratified meta analysis on 3 WHI cohorts: EMPC, AS311 and BAA23. The values for effect sizes are available in supplementary tables 3. Plots generated using Prism 9.*

A

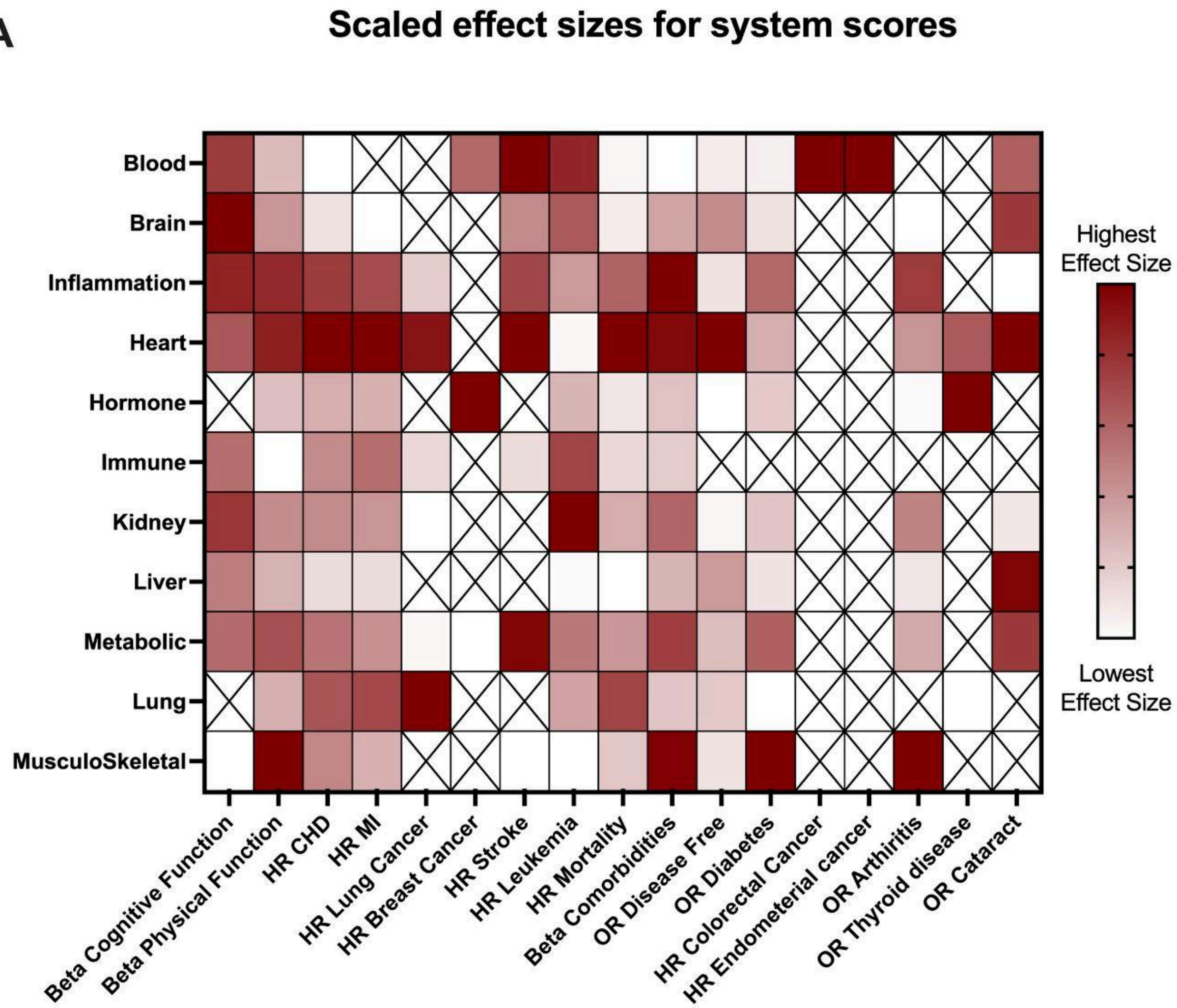

Scaled effect sizes for system scores
Blood
Brain
Inflammation
Heart
Hormone
Immune
Kidney
Liver
Metabolic
Lung
MusculoSkeletal
Beta Cognitive Function
Beta Physical Function
HR CHD
HR MI
HR Lung Cancer
HR Breast Cancer
HR Stroke
HR Leukemia
HR Mortality
Beta Comorbidities
OR Disease Free
OR Diabetes
HR Colorectal Cancer
HR Endometerial cancer
OR Arthiritis
OR Thyroid disease
OR Cataract
Highest Effect Size
Lowest Effect Size


B

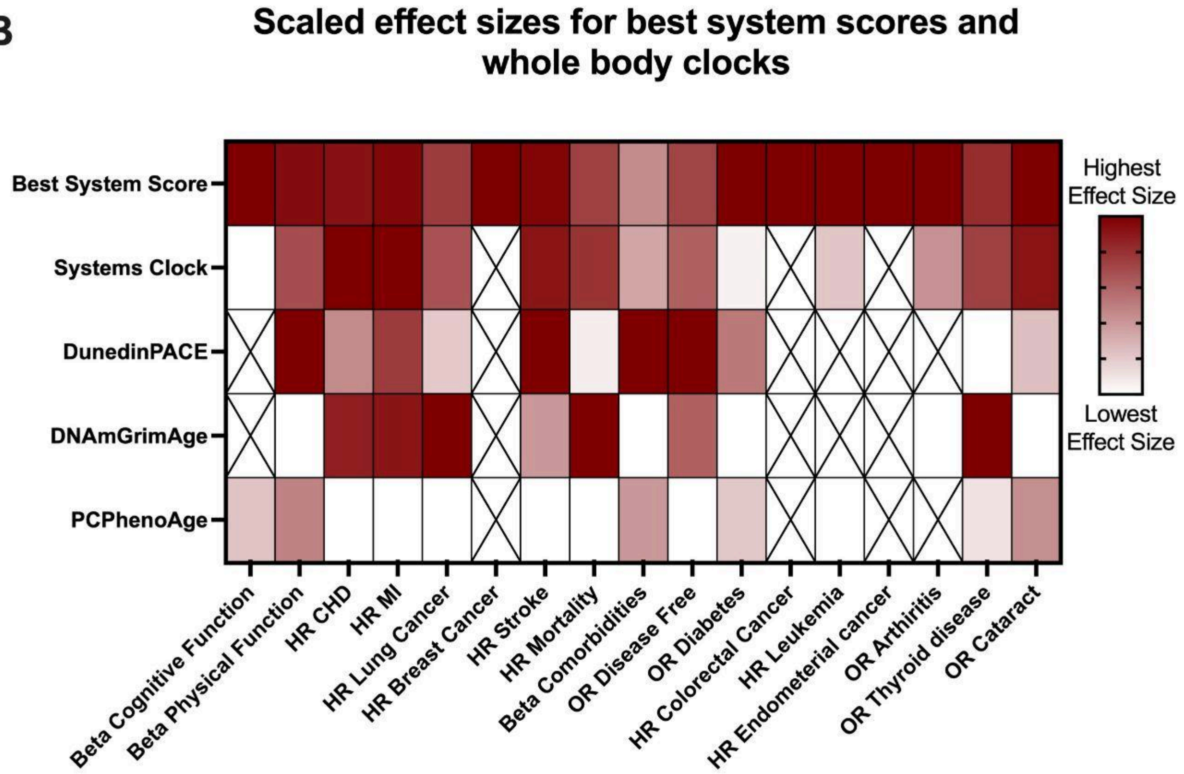

Scaled effect sizes for best system scores and whole body clocks
Best System Score
Systems Clock
DunedinPACE
DNAmGrimAge
PCPhenoAge
Beta Cognitive Function
Beta Physical Function
HR CHD
HR MI
HR Lung Cancer
HR Breast Cancer
HR Stroke
HR Mortality
Beta Comorbidities
OR Disease Free
OR Diabetes
HR Colorectal Cancer
HR Leukemia
HR Endometerial cancer
OR Arthiritis
OR Thyroid disease
OR Cataract
Highest Effect Size
Lowest Effect Size

*Supplementary figure 4: Scaled Effect sizes for (A) system scores (B) best system score and whole body clocks. A cross represents the effect size was not significant as measured by the z-score. Effect sizes were calculated using a race stratified meta analysis on 3 WHI cohorts: EMPC, AS311 and BAA23. Note, effect sizes for Endometrial cancer and Breast cancer have been inverted because of negative associations. The values for effect sizes are available in supplementary tables 3. Plots generated using Prism 9.*

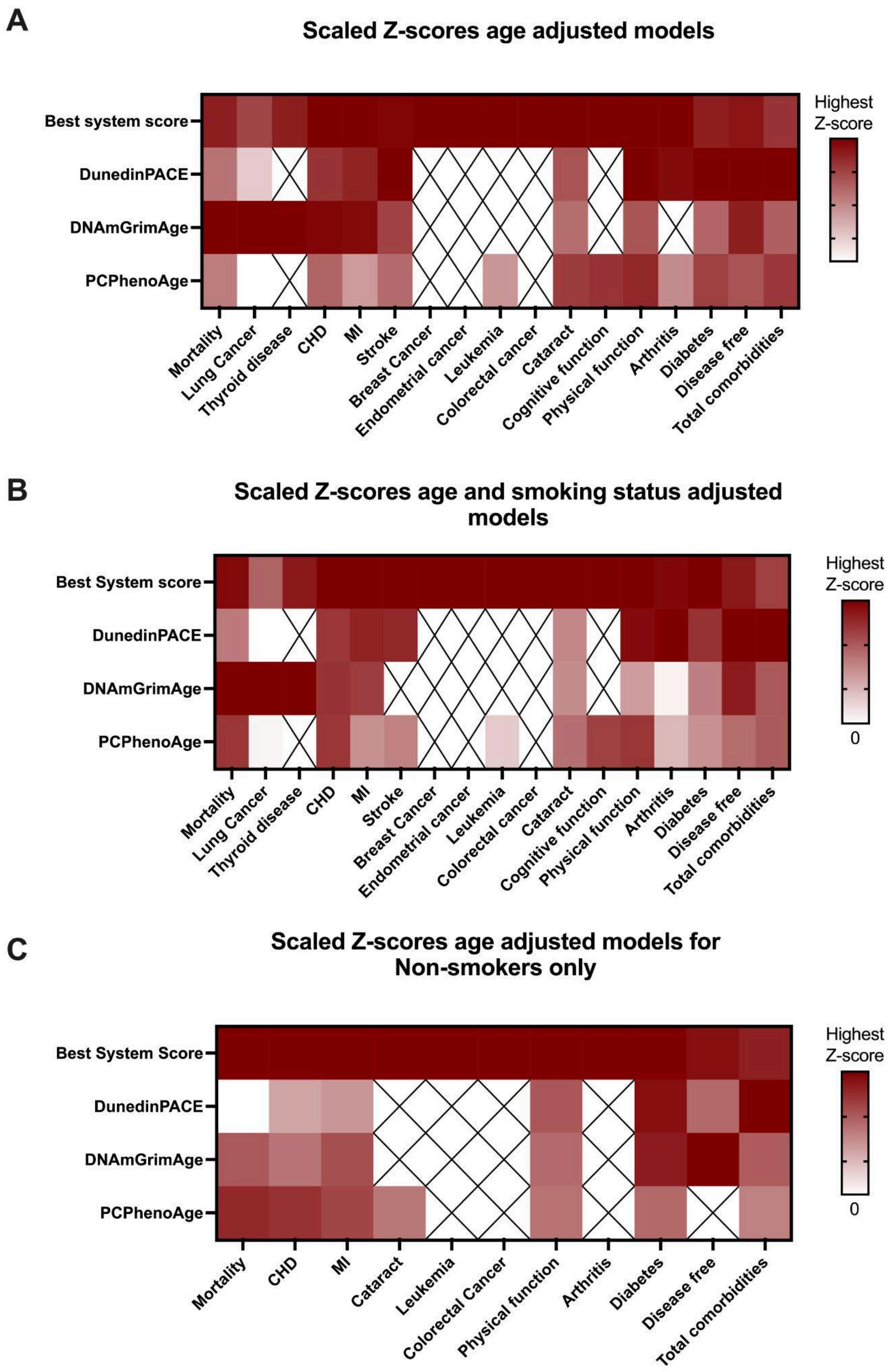
A
Scaled Z-scores age adjusted models
Best system score
DunedinPACE
DNAmGrimAge
PCPhenoAge
Mortality
Lung Cancer
Thyroid disease
CHD
MI
Stroke
Breast Cancer
Endometrial cancer
Leukemia
Colorectal cancer
Cataract
Cognitive function
Physical function
Arthritis
Diabetes
Disease free
Total comorbidities
Highest Z-score
B
Scaled Z-scores age and smoking status adjusted models
Best System score
DunedinPACE
DNAmGrimAge
PCPhenoAge
Mortality
Lung Cancer
Thyroid disease
CHD
MI
Stroke
Breast Cancer
Endometrial cancer
Leukemia
Colorectal cancer
Cataract
Cognitive function
Physical function
Arthritis
Diabetes
Disease free
Total comorbidities
Highest Z-score
0
C
Scaled Z-scores age adjusted models for Non-smokers only
Best System Score
DunedinPACE
DNAmGrimAge
PCPhenoAge
Mortality
CHD
MI
Cataract
Leukemia
Colorectal Cancer
Physical function
Arthritis
Diabetes
Disease free
Total comorbidities
Highest Z-score
0

*Supplementary figure 5: Comparing the best system score to other whole body clocks. Z-scores are scaled by condition with age adjusted (A), age and smoking status adjusted (B) and age adjusted only for non-smokers (C). A cross represents a not significant Z-score. Z-scores were calculated using race stratified meta analysis on 3 WHI cohorts: EMPC, AS311 and BAA23. Z-scores are shown for only those variables which had at least 1 significant z-score. Plots generated using Prism 9.*

A

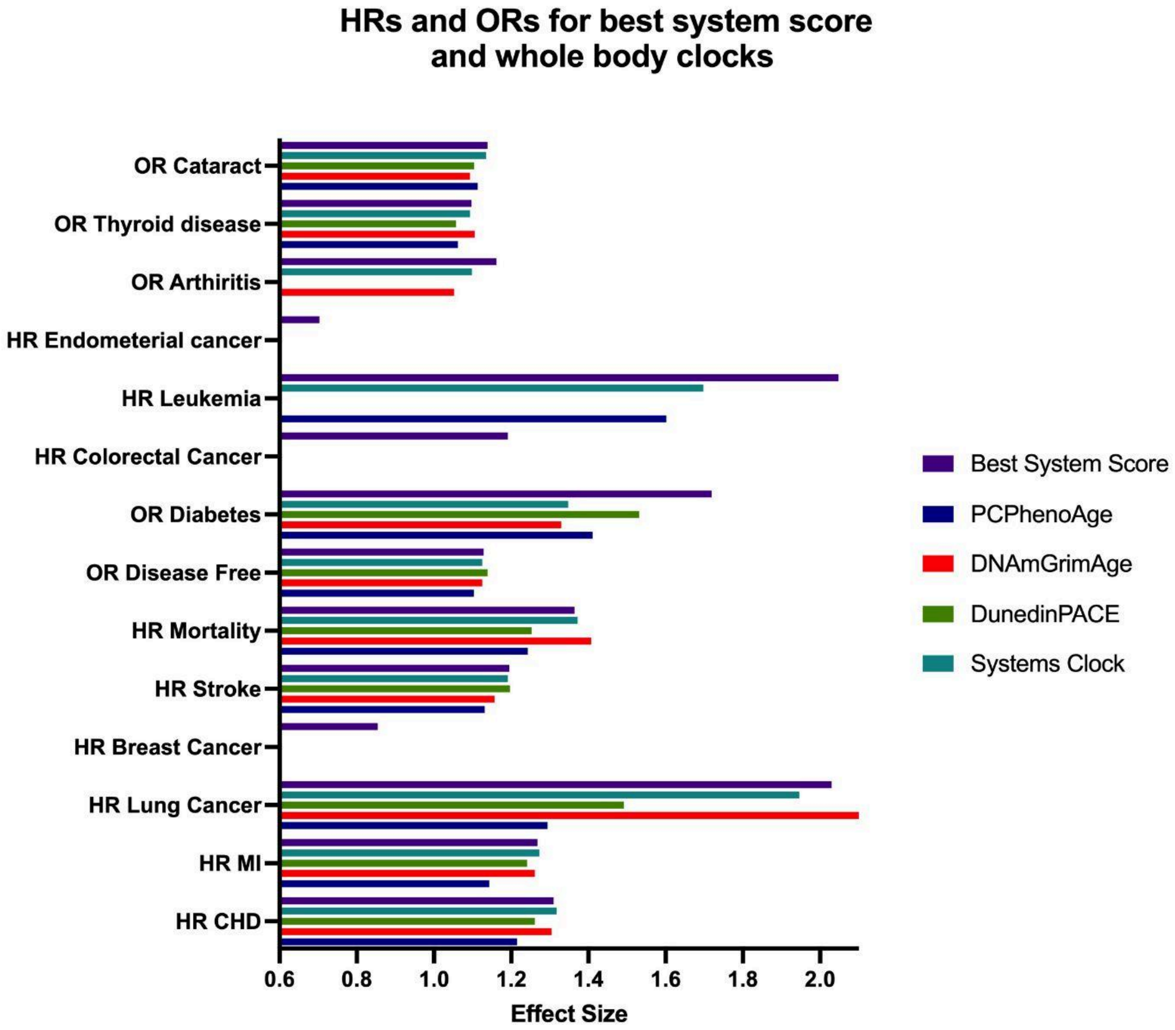
HRs and ORs for best system score and whole body clocks
OR Cataract
OR Thyroid disease
OR Arthiritis
HR Endometerial cancer
HR Leukemia
HR Colorectal Cancer
OR Diabetes
OR Disease Free
HR Mortality
HR Stroke
HR Breast Cancer
HR Lung Cancer
HR MI
HR CHD
0.6
0.8
1.0
1.2
1.4
1.6
1.8
2.0
Effect Size
Best System Score
PCPhenoAge
DNAmGrimAge
DunedinPACE
Systems Clock

B

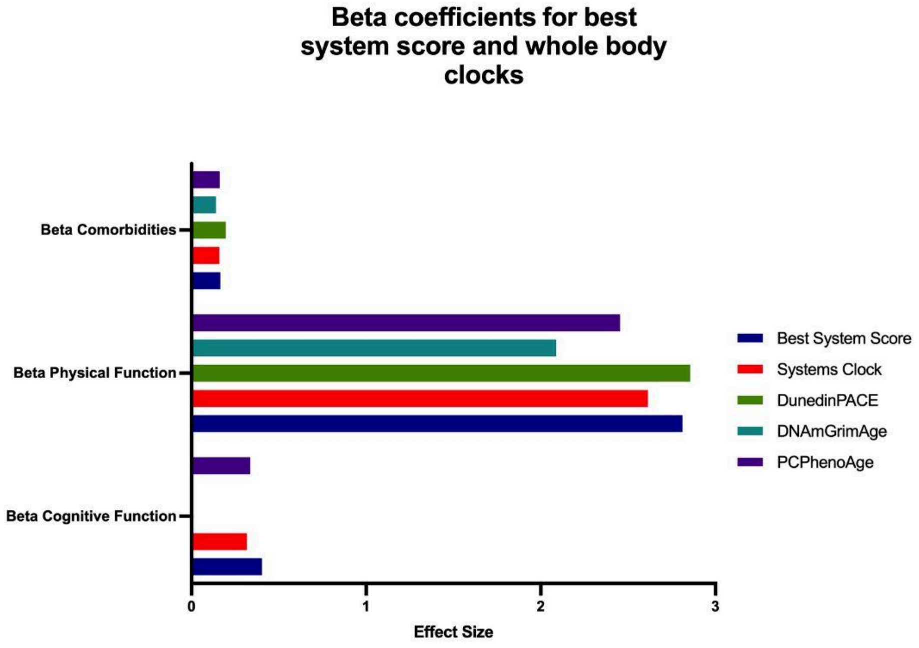
Beta coefficients for best system score and whole body clocks
Beta Comorbidities
Beta Physical Function
Beta Cognitive Function
0
1
2
3
Effect Size
Best System Score
Systems Clock
DunedinPACE
DNAmGrimAge
PCPhenoAge

*Supplementary figure 6: Comparing effect sizes for best system score to whole body clocks. Hazard Ratios and Odds Ratios for different models (A) and beta coefficients for different models (B). Only significant effect sizes have been plotted as measured by Z-score. Effect sizes were calculated using a race stratified meta analysis on 3 WHI cohorts: EMPC, AS311 and BAA23. The values for effect sizes are available in supplementary tables 3. Plots generated using Prism 9.*

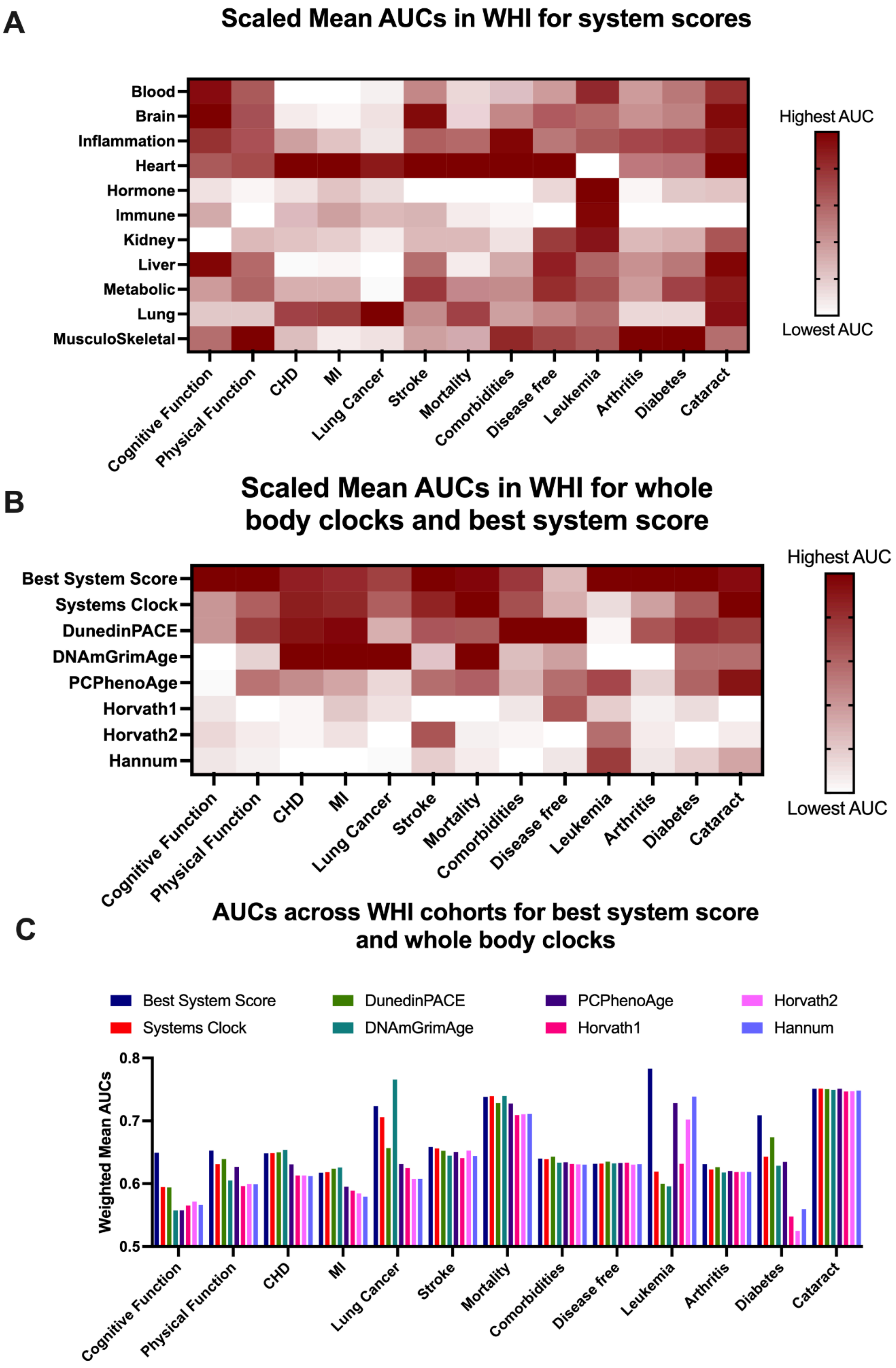
A
Scaled Mean AUCs in WHI for system scores
Blood
Brain
Inflammation
Heart
Hormone
Immune
Kidney
Liver
Metabolic
Lung
MusculoSkeletal
Cognitive Function
Physical Function
CHD
MI
Lung Cancer
Stroke
Mortality
Comorbidities
Disease free
Leukemia
Arthritis
Diabetes
Cataract
Highest AUC
Lowest AUC
B
Scaled Mean AUCs in WHI for whole body clocks and best system score
Best System Score
Systems Clock
DunedinPACE
DNAmGrimAge
PCPhenoAge
Horvath1
Horvath2
Hannum
Highest AUC
Lowest AUC
C
AUCs across WHI cohorts for best system score and whole body clocks
Best System Score
Systems Clock
DunedinPACE
DNAmGrimAge
PCPhenoAge
Horvath1
Horvath2
Hannum
Weighted Mean AUCs
0.8
0.7
0.6
0.5

*Supplementary figure 7: Mean AUCs A) scaled by condition for system scores, B) scaled by condition for best system score and whole body clocks, C) unscaled for all best system score and whole body clocks. AUCs were calculated using mean weighted AUCs from 3 WHI cohorts: EMPC, AS311 and BAA23. The values for AUCs are available in supplementary tables 8. Plots generated using Prism 9.*

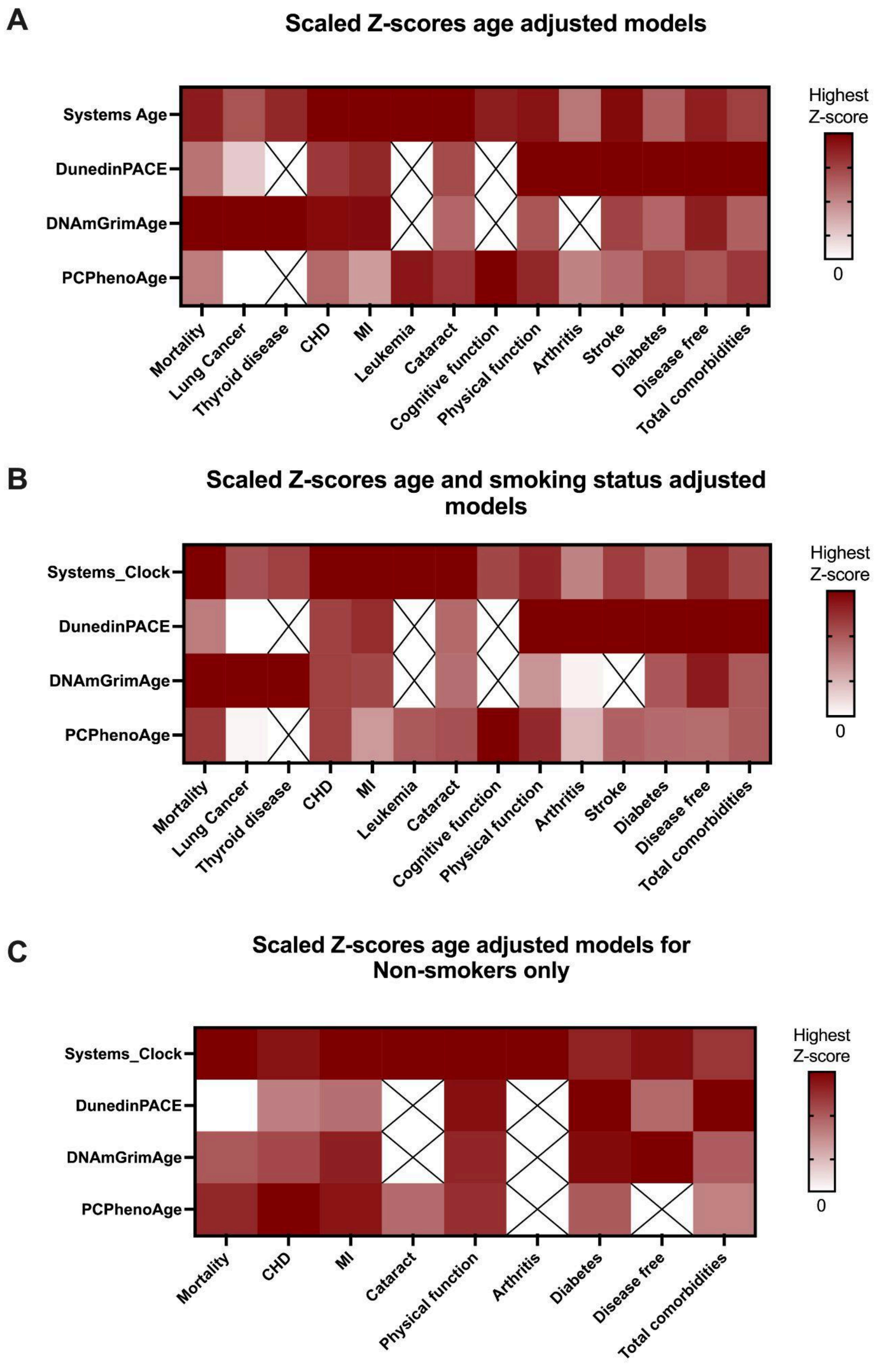
A
Scaled Z-scores age adjusted models
Systems Age
DunedinPACE
DNAmGrimAge
PCPhenoAge
Mortality
Lung Cancer
Thyroid disease
CHD
MI
Leukemia
Cataract
Cognitive function
Physical function
Arthritis
Stroke
Diabetes
Disease free
Total comorbidities
Highest Z-score
0
B
Scaled Z-scores age and smoking status adjusted models
Systems_Clock
DunedinPACE
DNAmGrimAge
PCPhenoAge
Mortality
Lung Cancer
Thyroid disease
CHD
MI
Leukemia
Cataract
Cognitive function
Physical function
Arthritis
Stroke
Diabetes
Disease free
Total comorbidities
Highest Z-score
0
C
Scaled Z-scores age adjusted models for Non-smokers only
Systems_Clock
DunedinPACE
DNAmGrimAge
PCPhenoAge
Mortality
CHD
MI
Cataract
Physical function
Arthritis
Diabetes
Disease free
Total comorbidities
Highest Z-score
0

*Supplementary figure 8: Comparing Systems Age to other whole body clocks. Z-scores are scaled by condition with age adjusted (A), age and smoking status adjusted (B) and age adjusted only for non-smokers (C). A cross represents a not significant Z-score. Z-scores were calculated using race stratified meta analysis on 3 WHI cohorts: EMPC, AS311 and BAA23. Z-scores are shown for only those variables which had at least 1 significant z-score. Plots generated using Prism 9.*

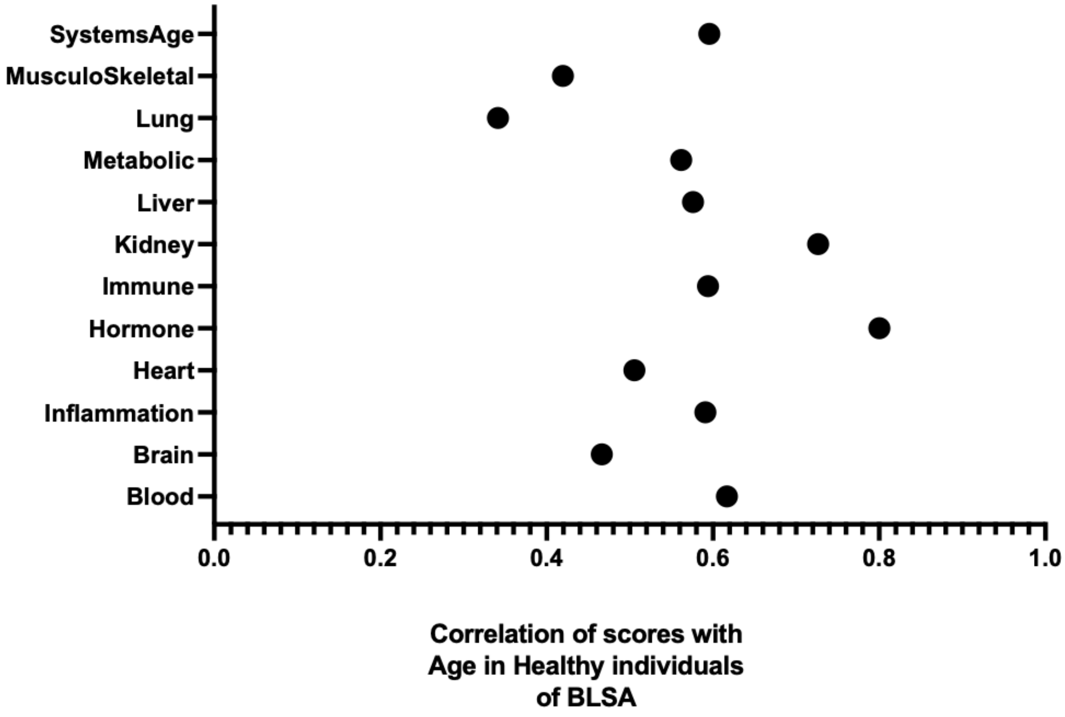


*Supplementary figure 9: Correlations of Systems Age and system scores with chronological age in healthy individuals (no disease) of BLSA. Plots generated using Prism 9.*
