## Supplementary figures and images for "Systems Age: A single blood methylation test to quantify aging heterogeneity across 11 physiological systems"

### Age_prediction_Disease free.pdf

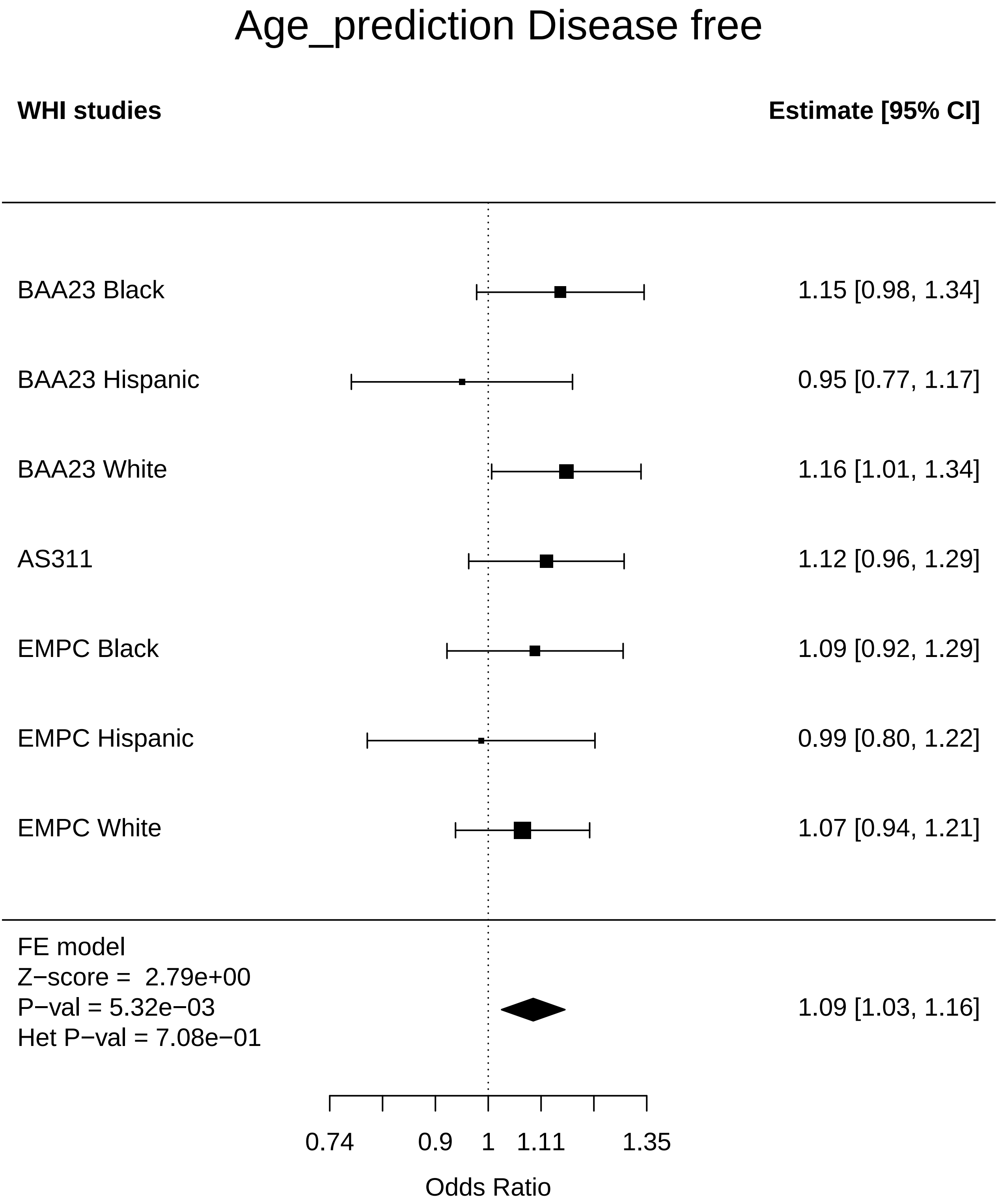

### Blood_Arthritis.pdf

#### Blood Arthritis

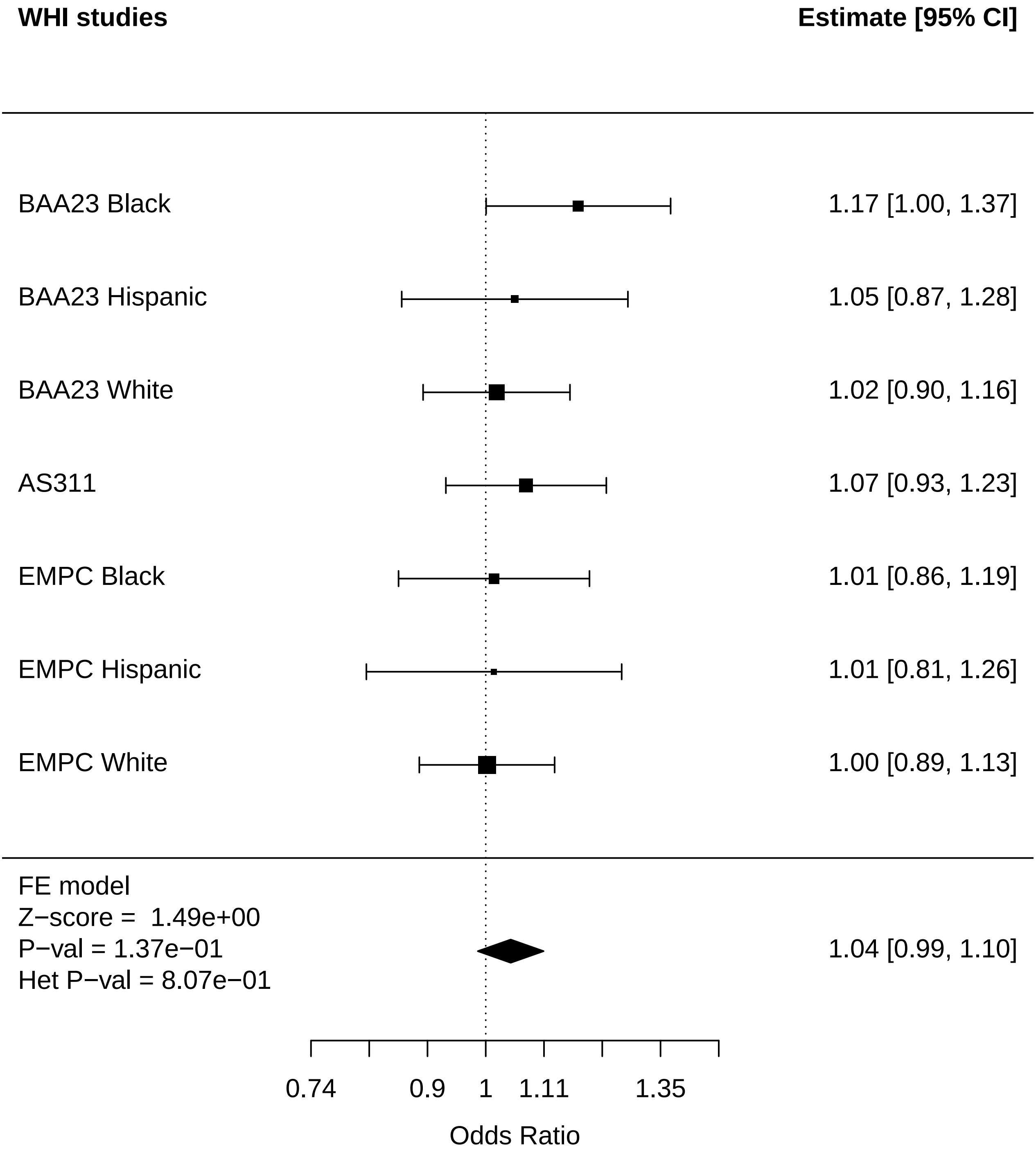

### Heart_Arthritis.pdf

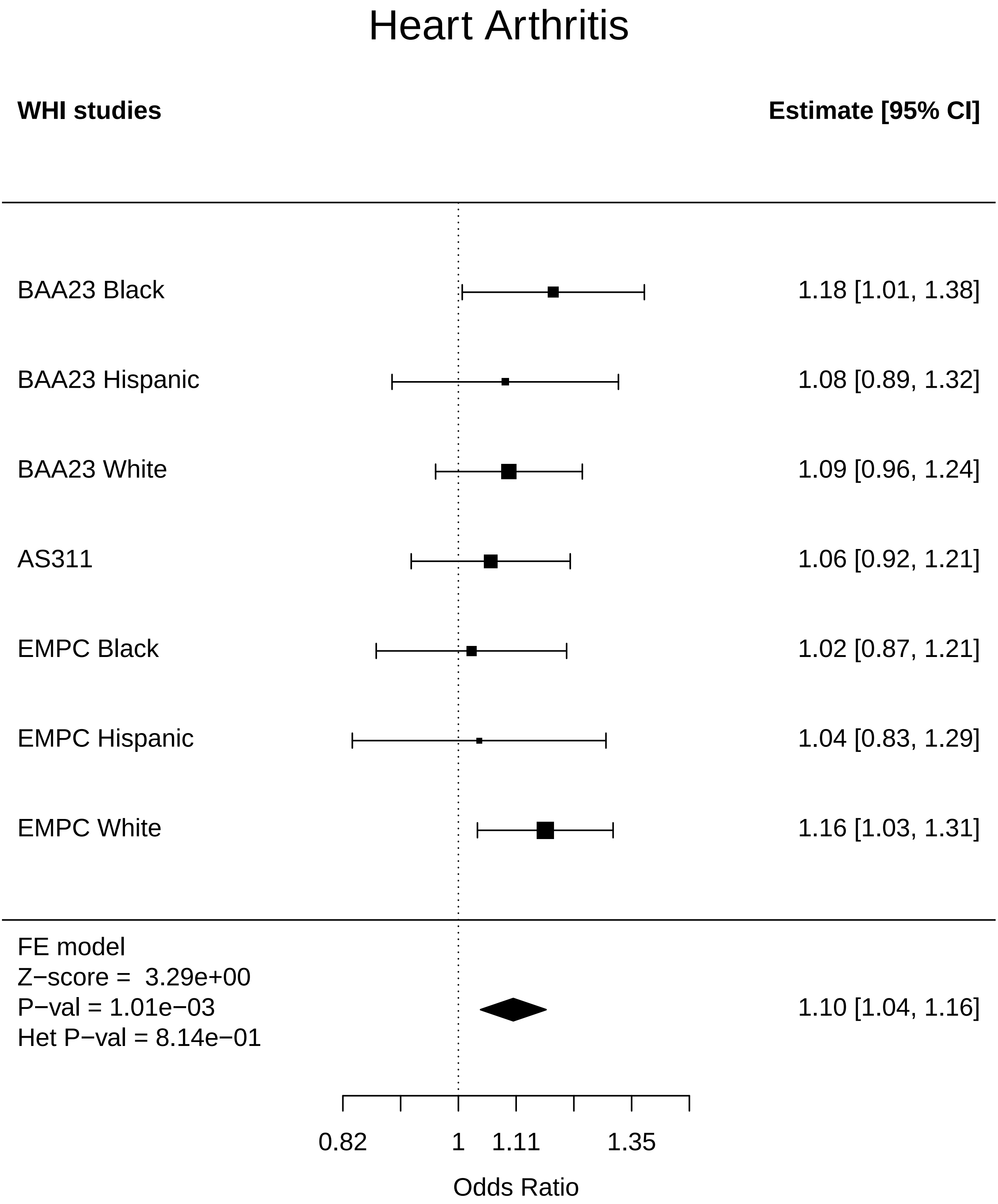

### Heart_Breast Cancer.pdf

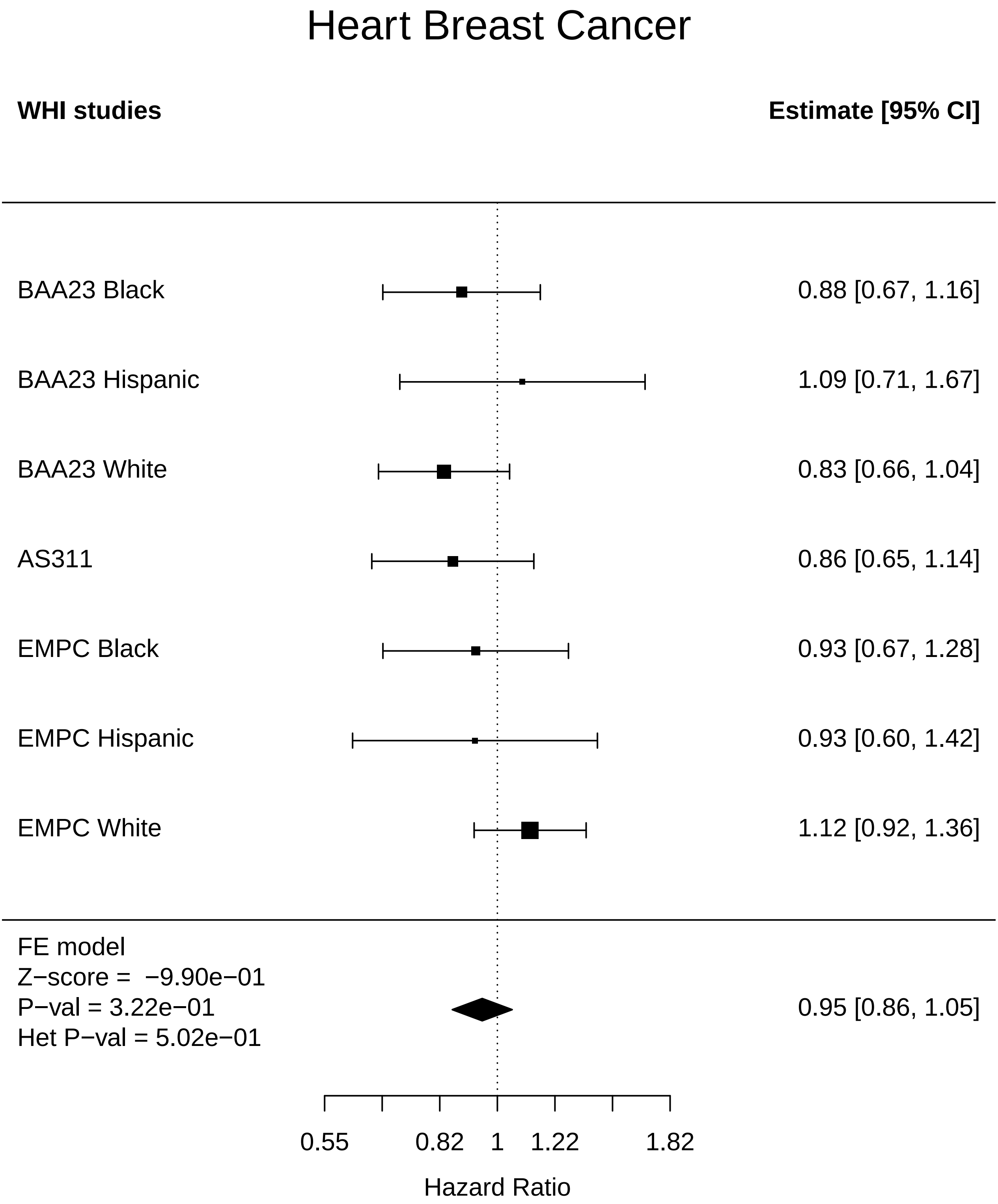

### Heart_Cataract.pdf

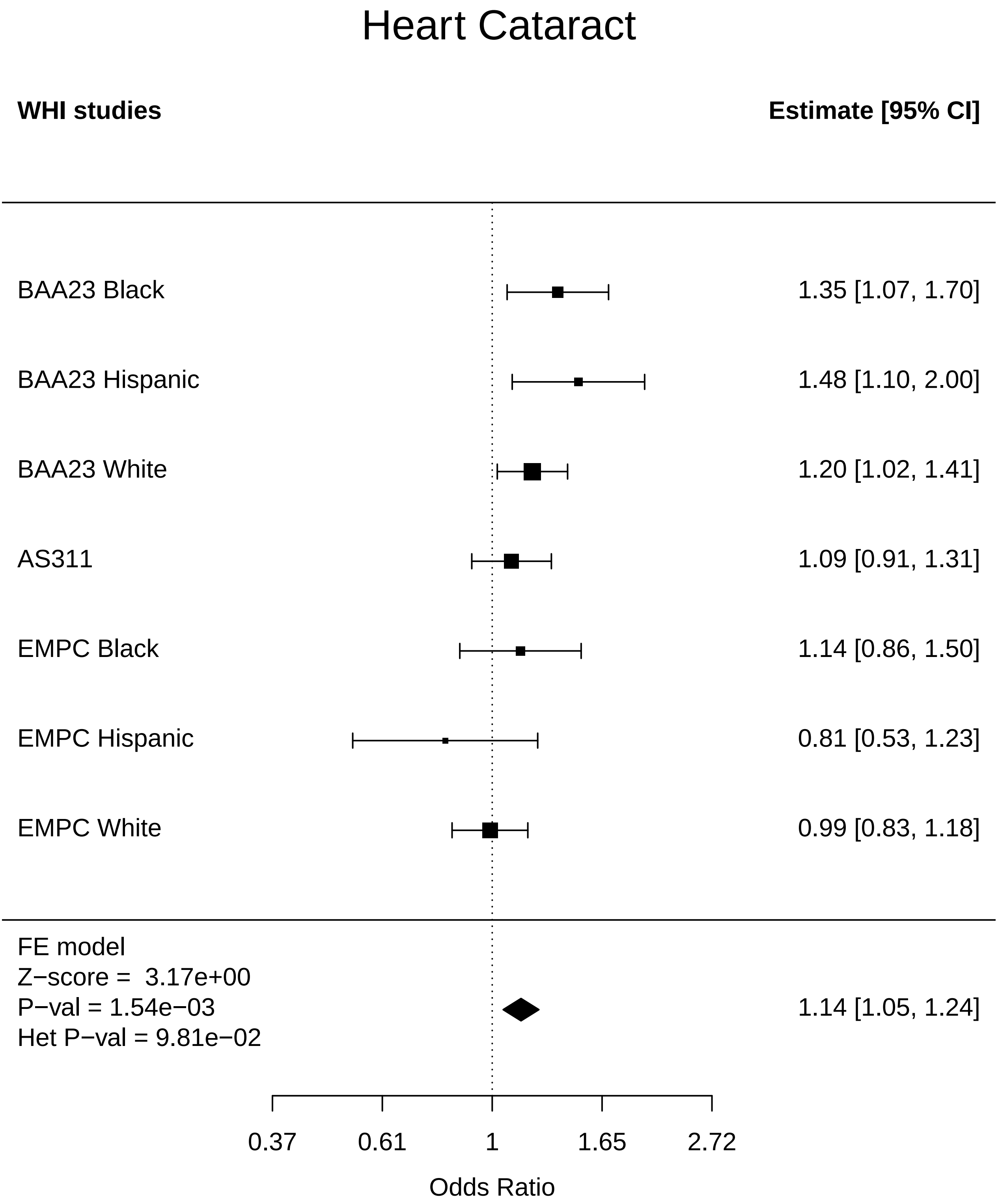

### Heart_CHD.pdf

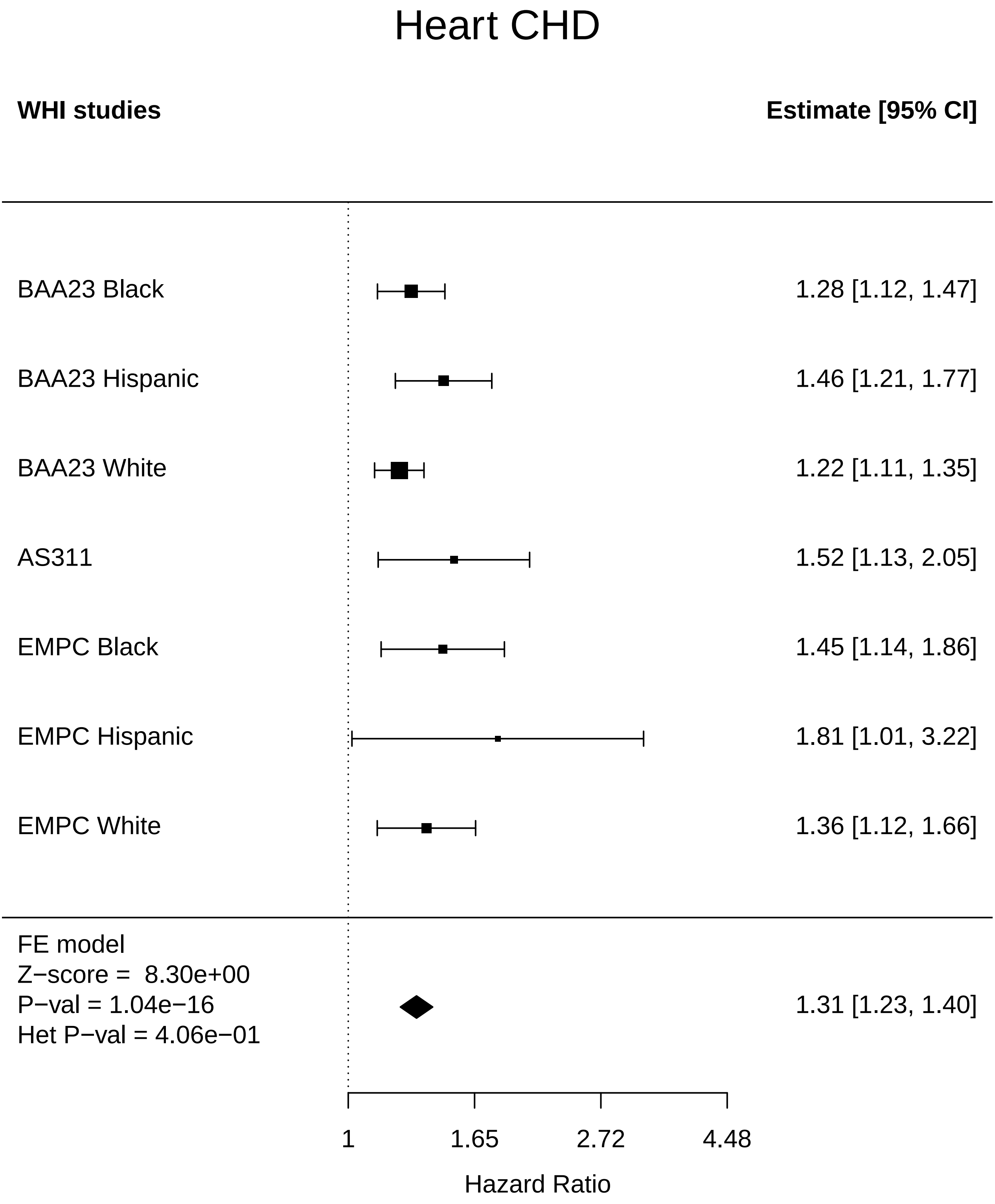

### Heart_Cognitive function.pdf

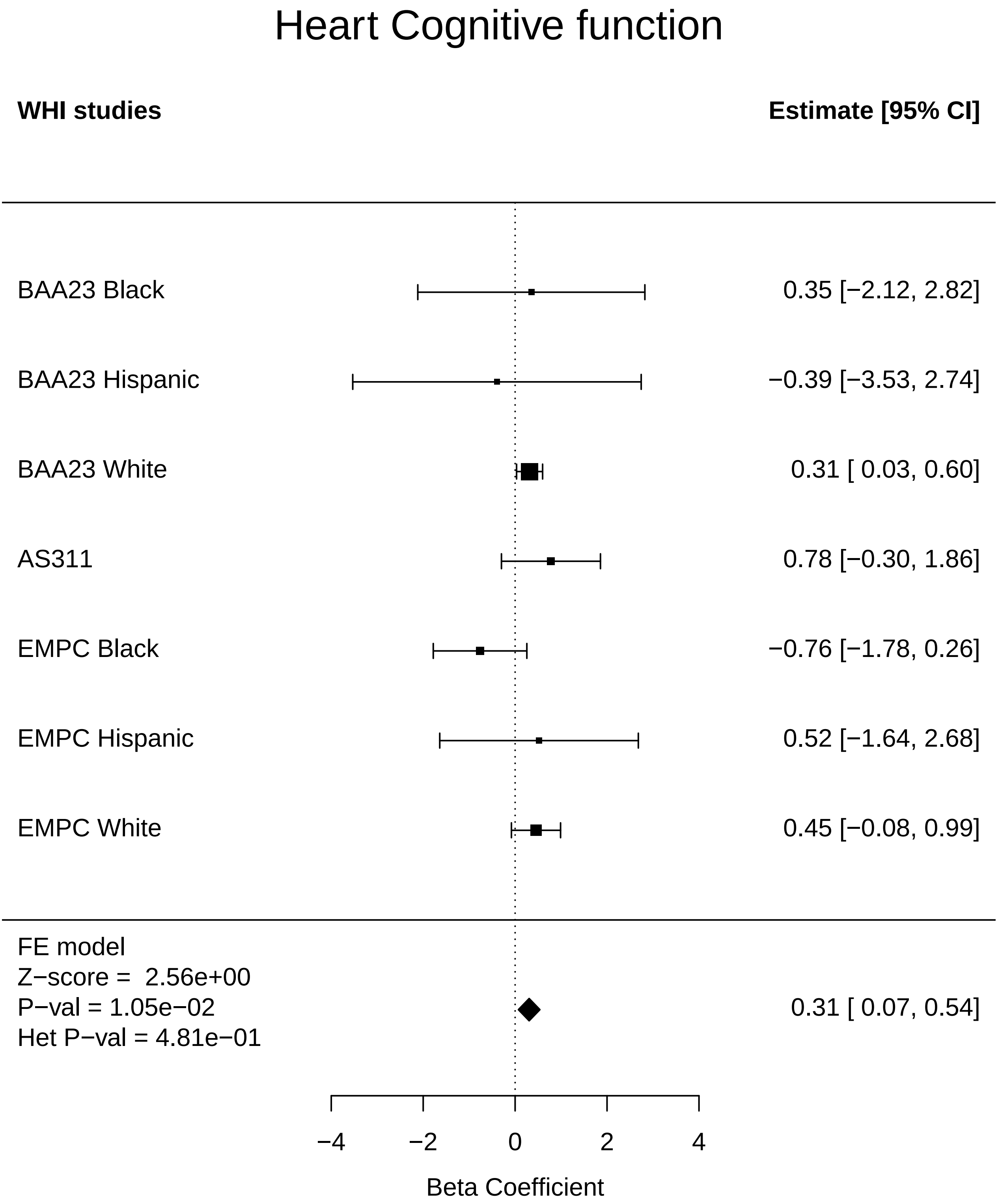

### Heart_Colorectal cancer.pdf

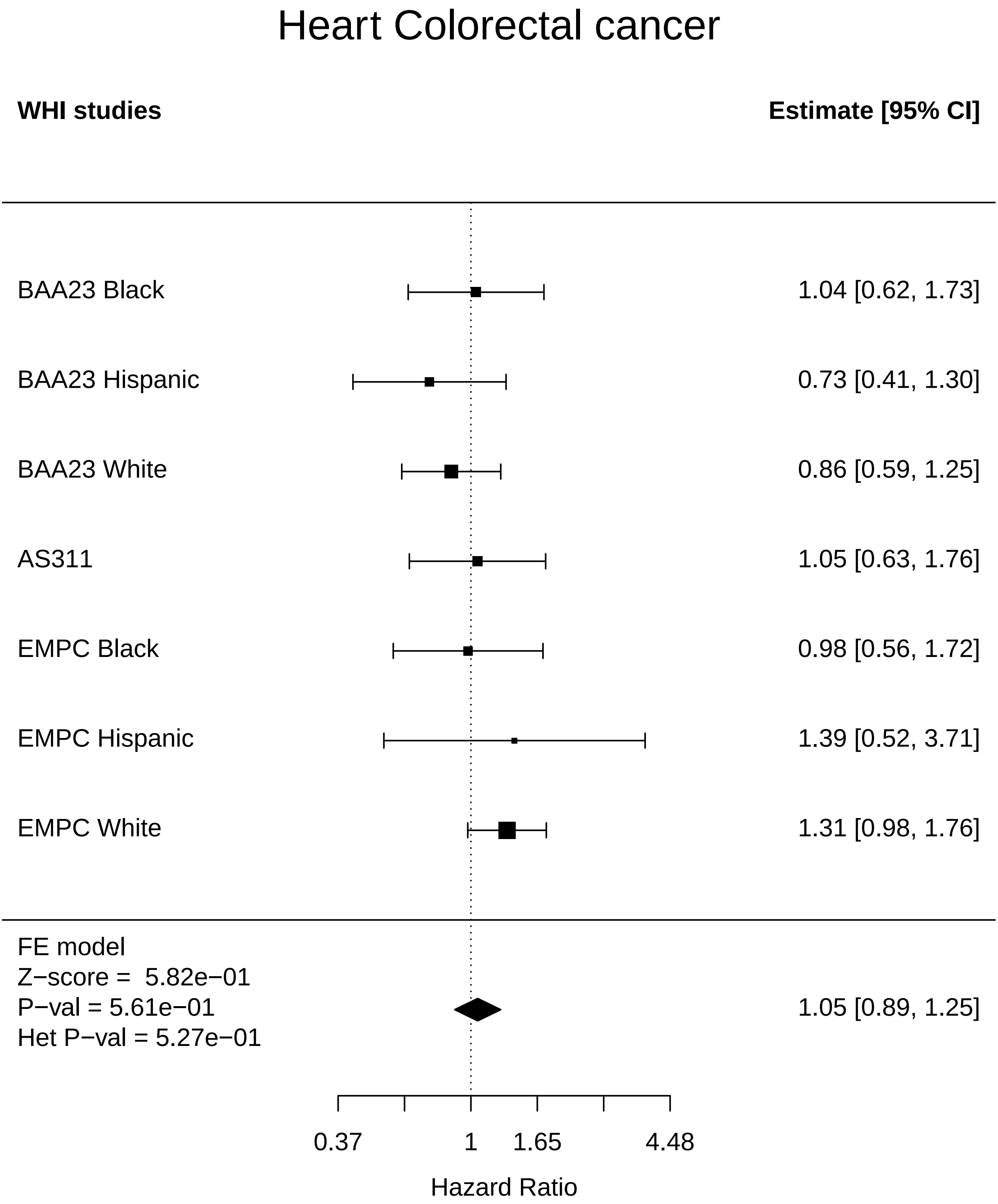

### Heart_Diabetes.pdf

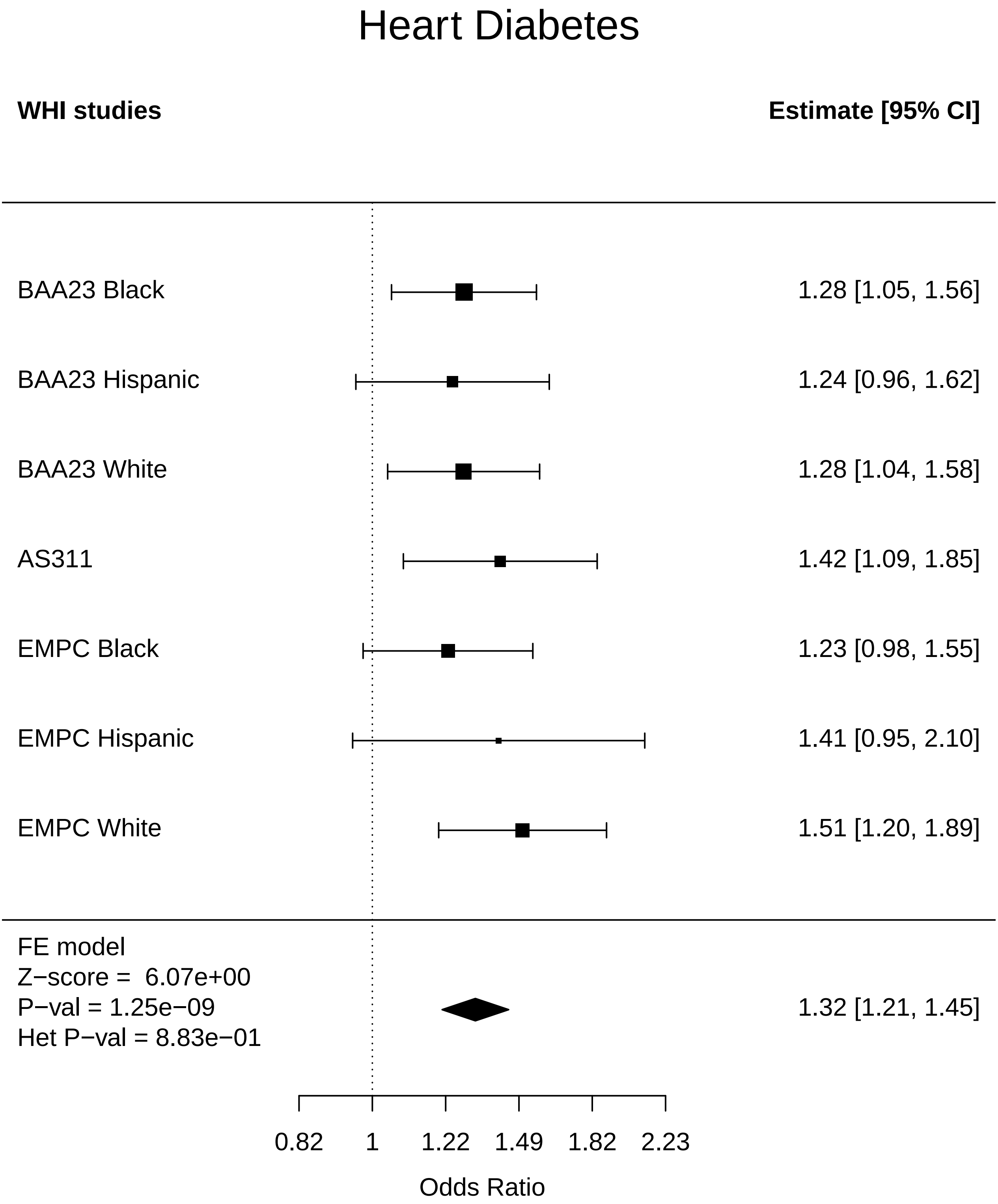

### Heart_Disease free.pdf

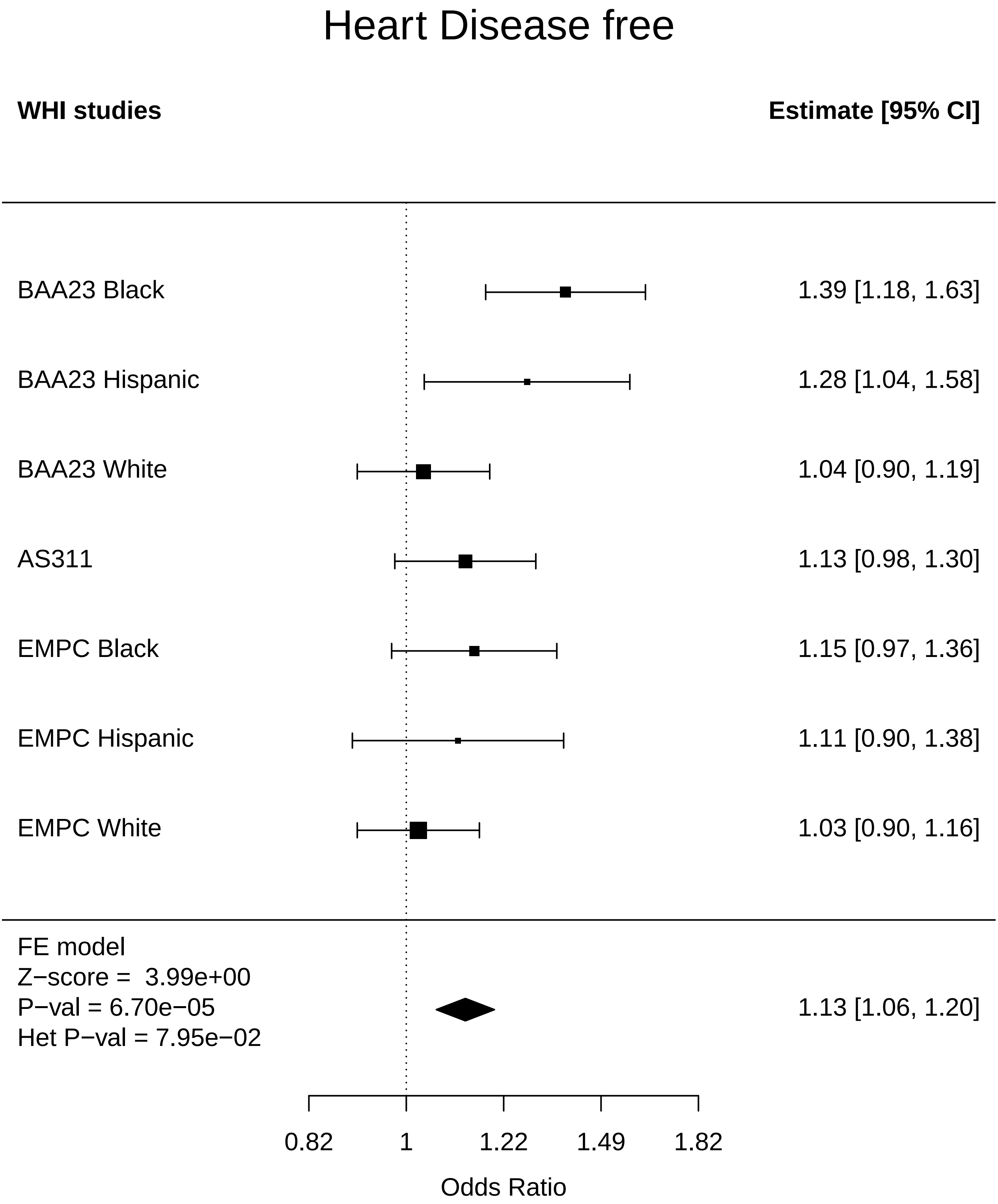

### Heart_Endometrial cancer.pdf

#### Heart Endometrial cancer

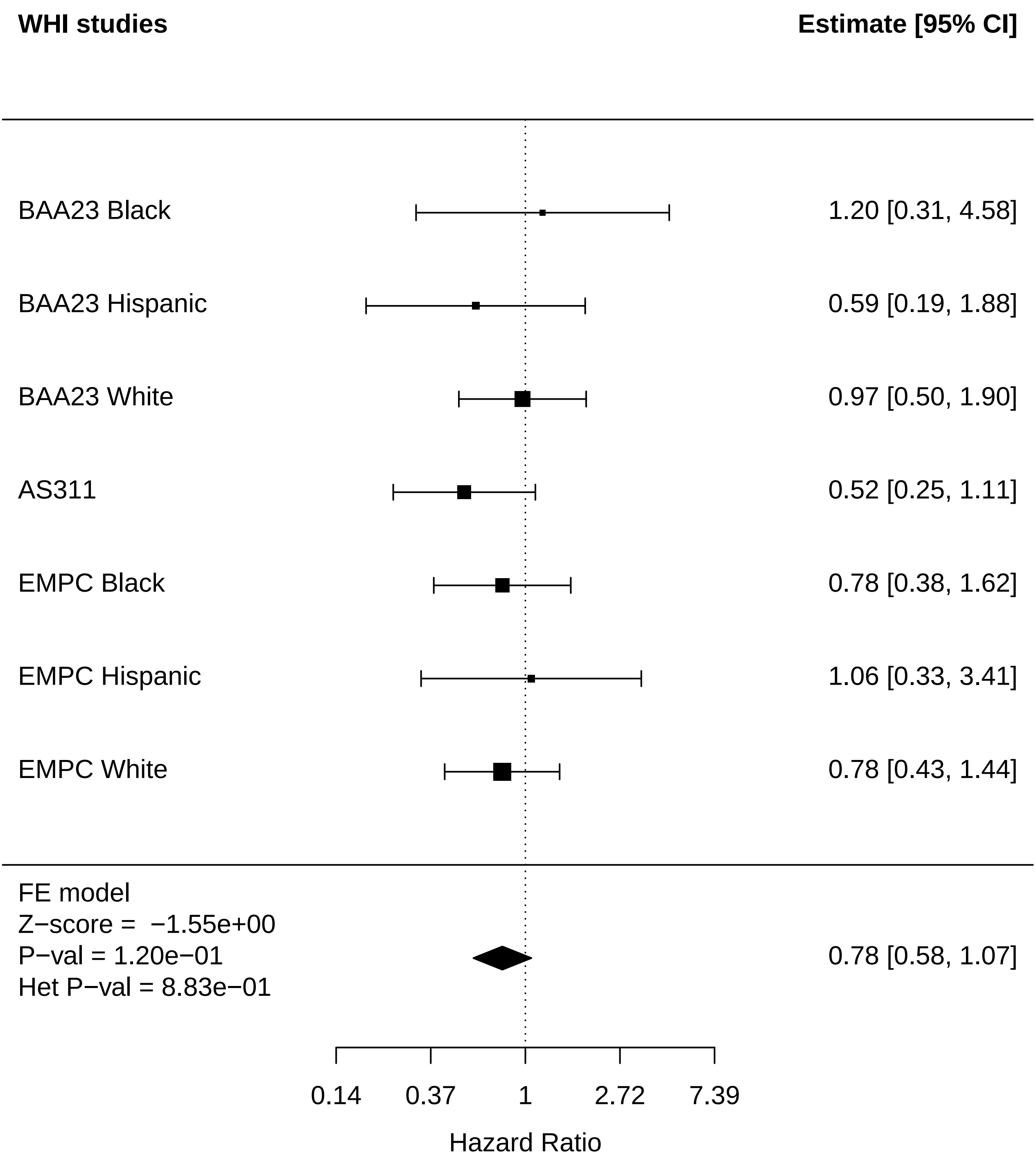

### Heart_Lung Cancer.pdf

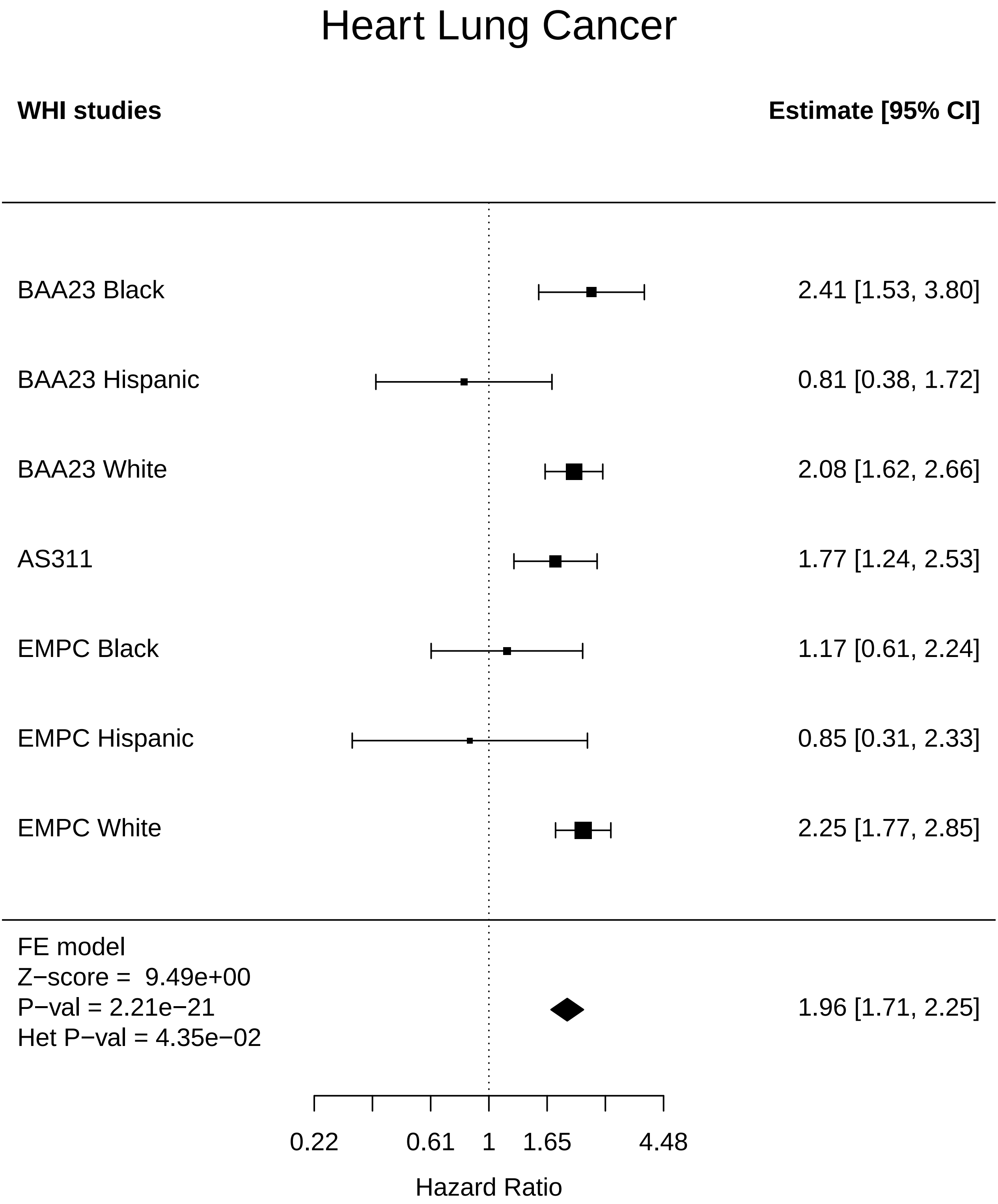

### Heart_MI.pdf

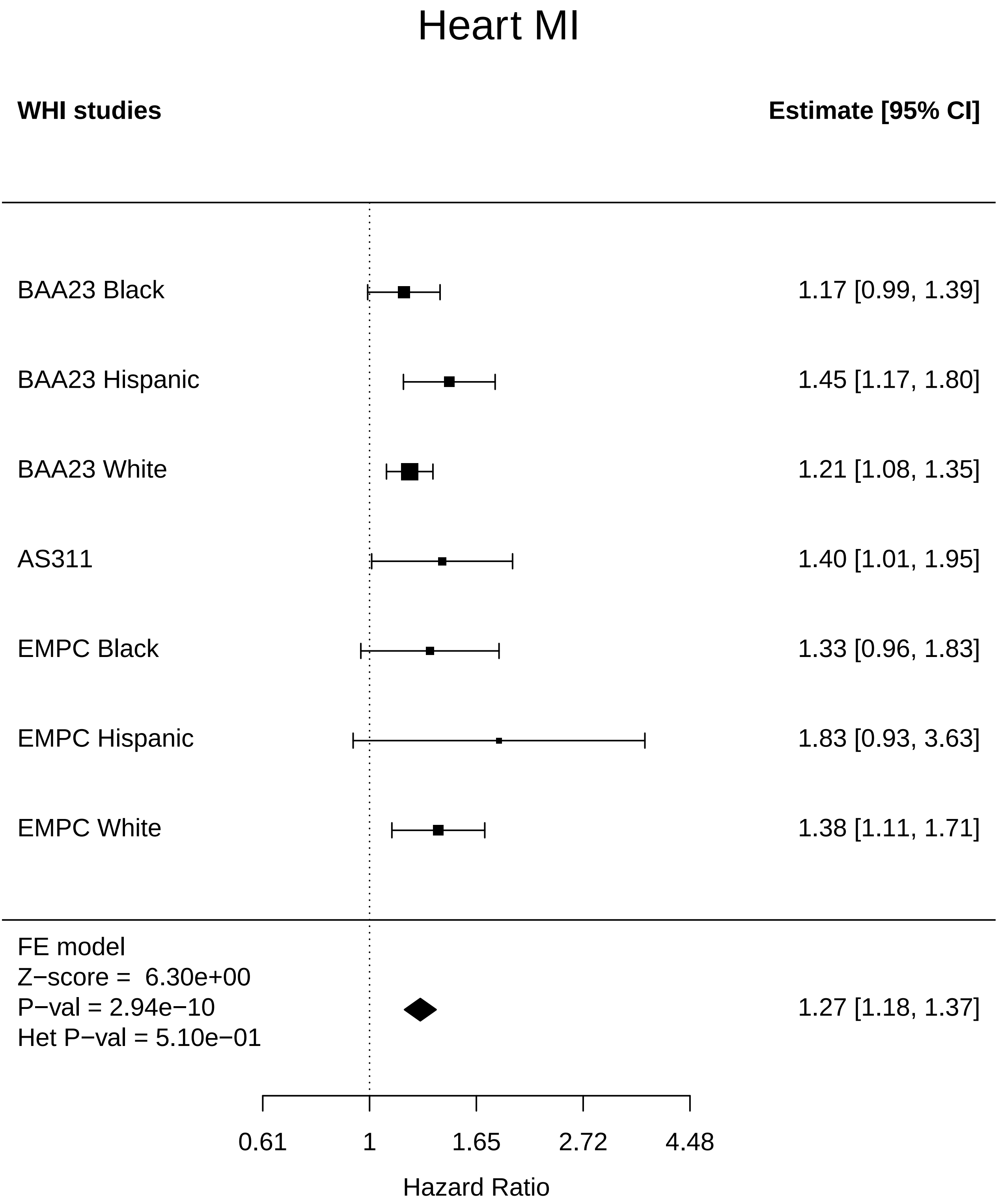

### Heart_Physical function.pdf

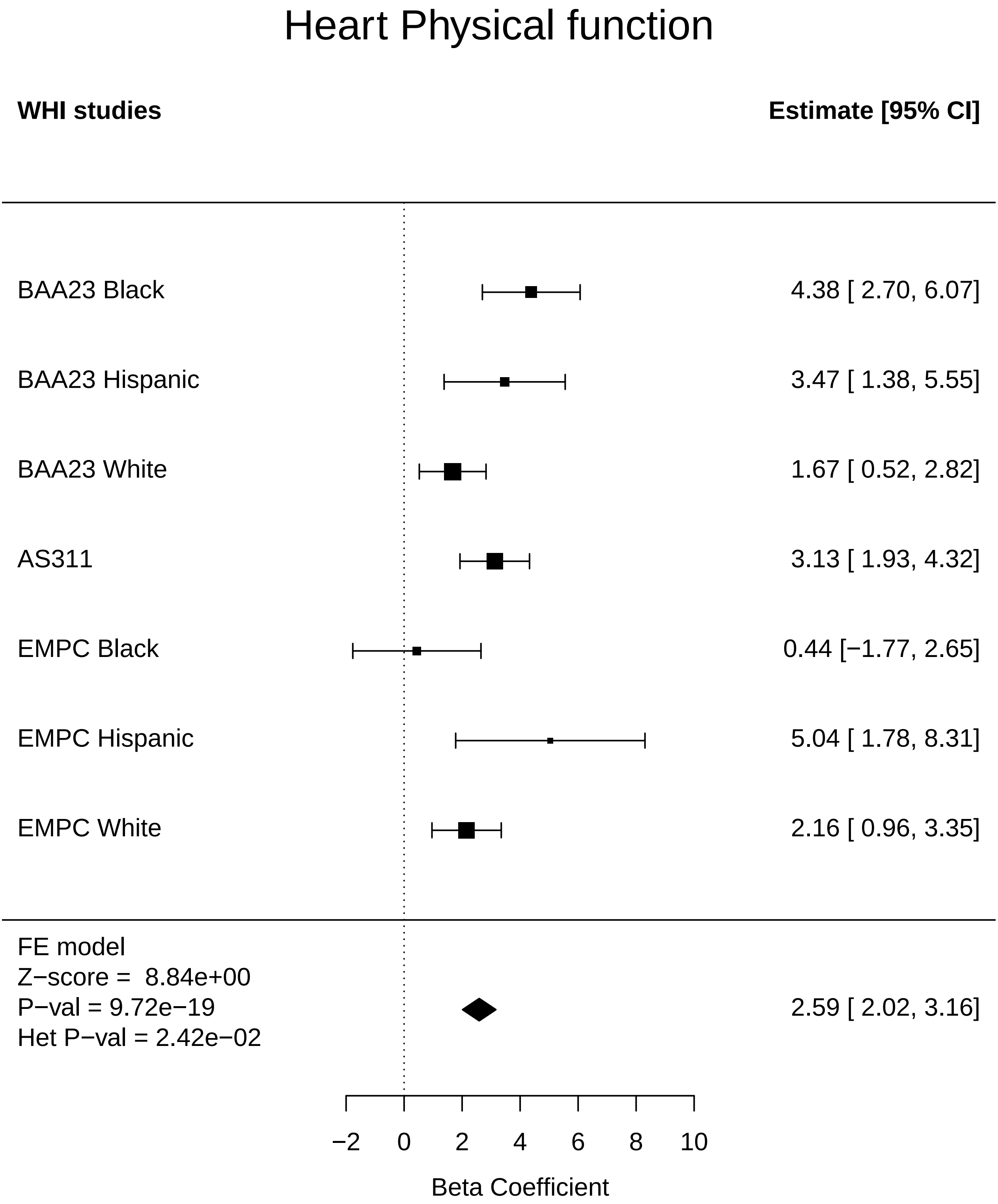

### Heart_Stroke.pdf

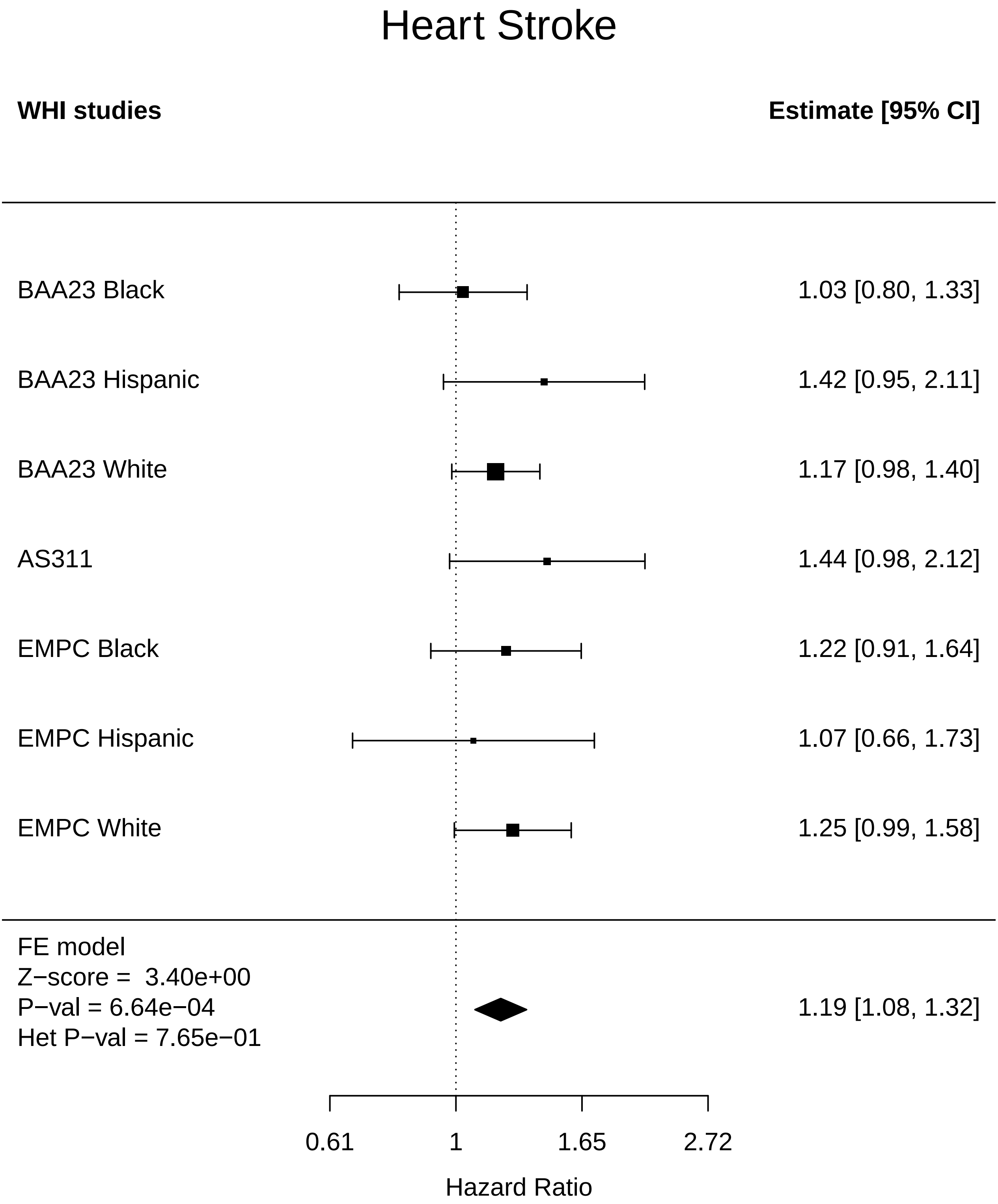

### Heart_Thyroid disease.pdf

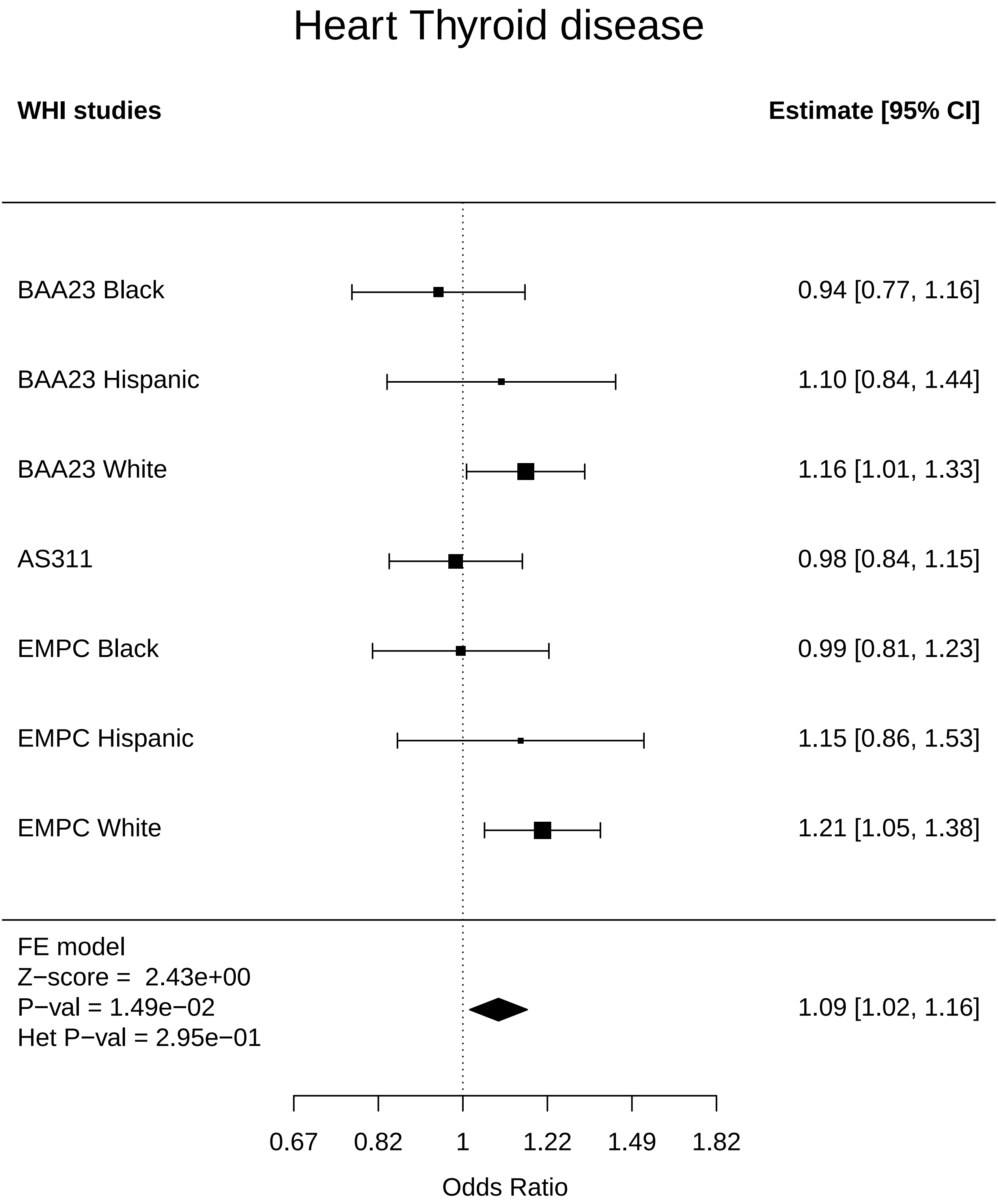

### Heart_Total comorbidities.pdf

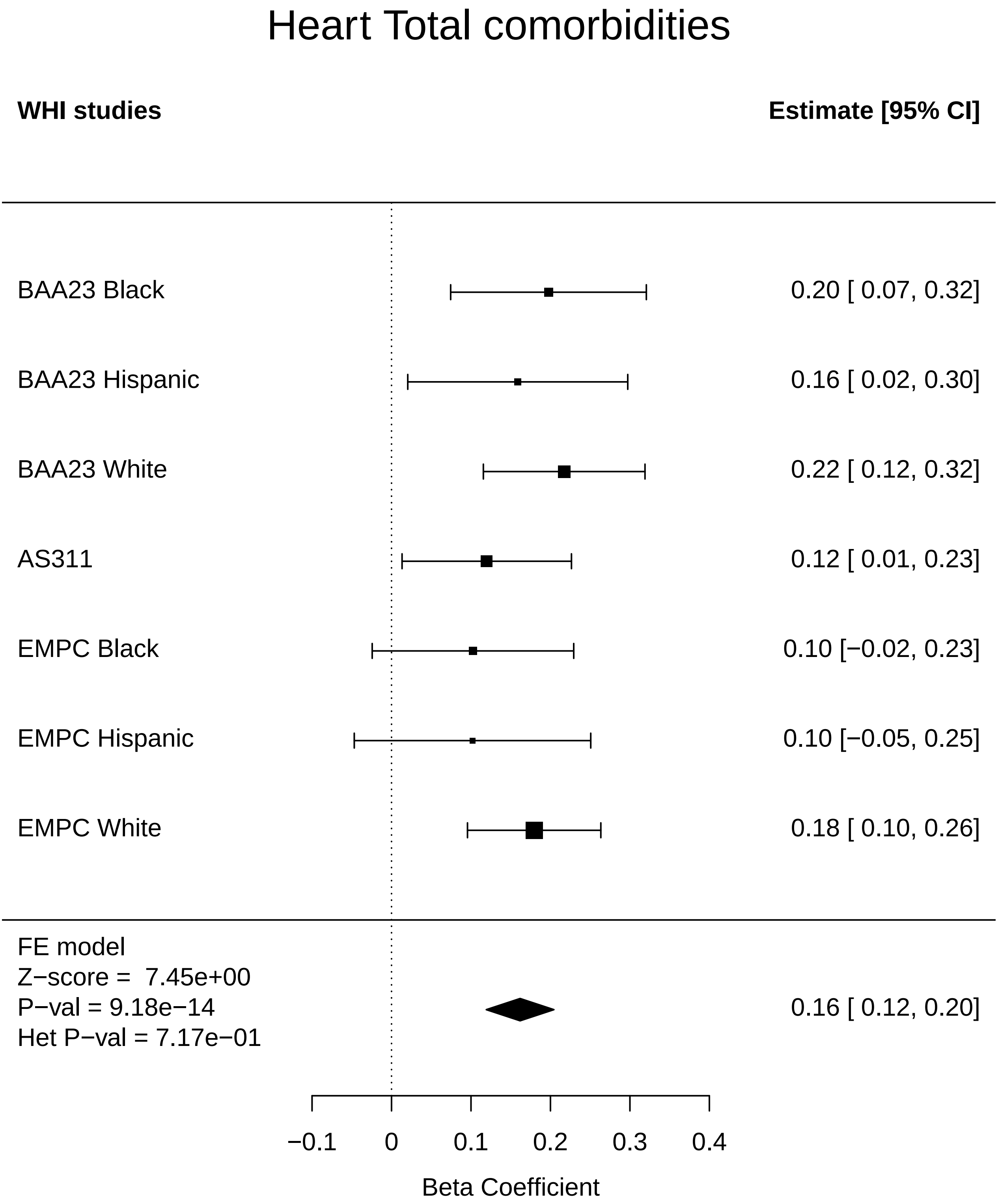

### Hormone_Arthritis.pdf

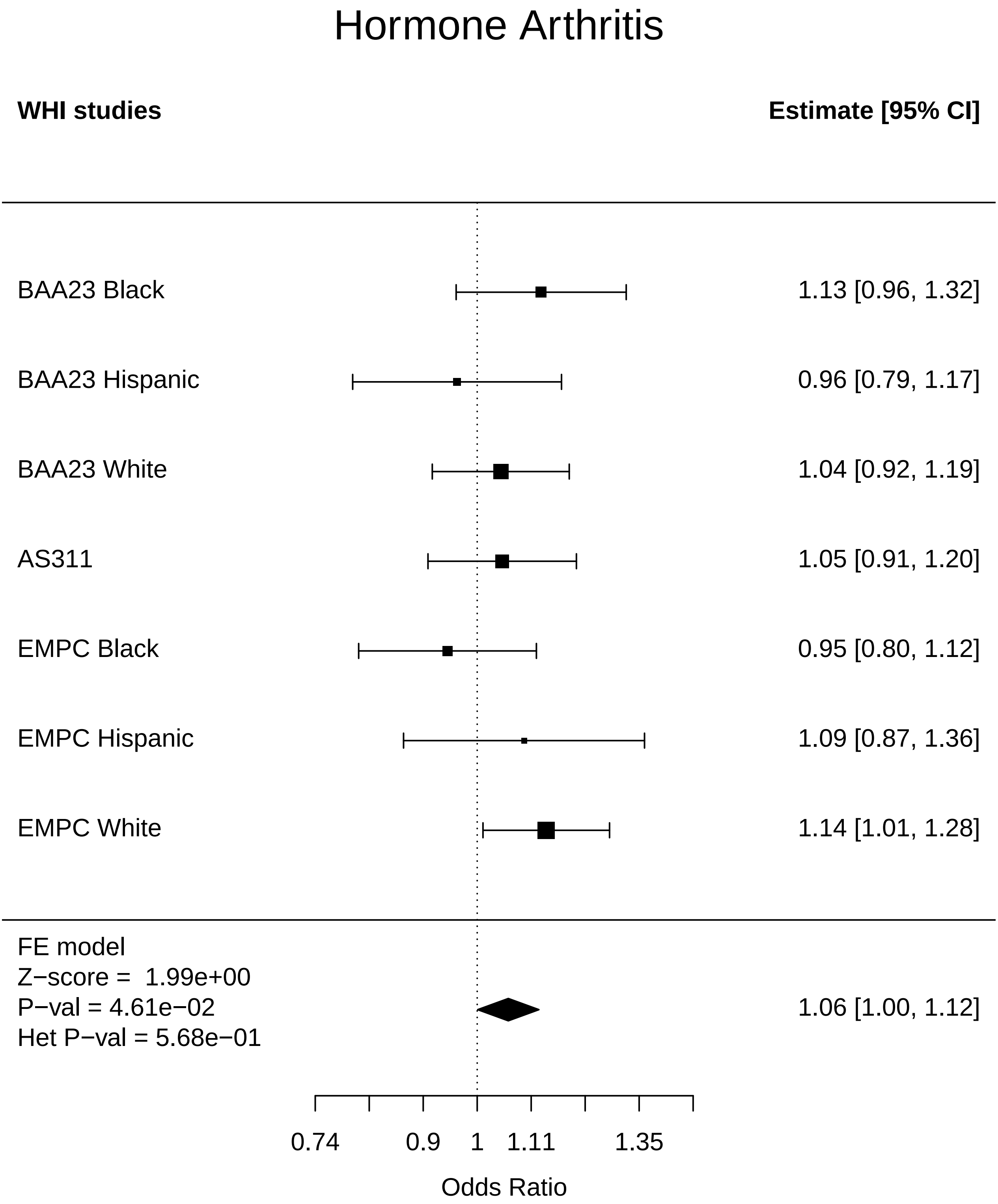

### Hormone_Breast Cancer.pdf

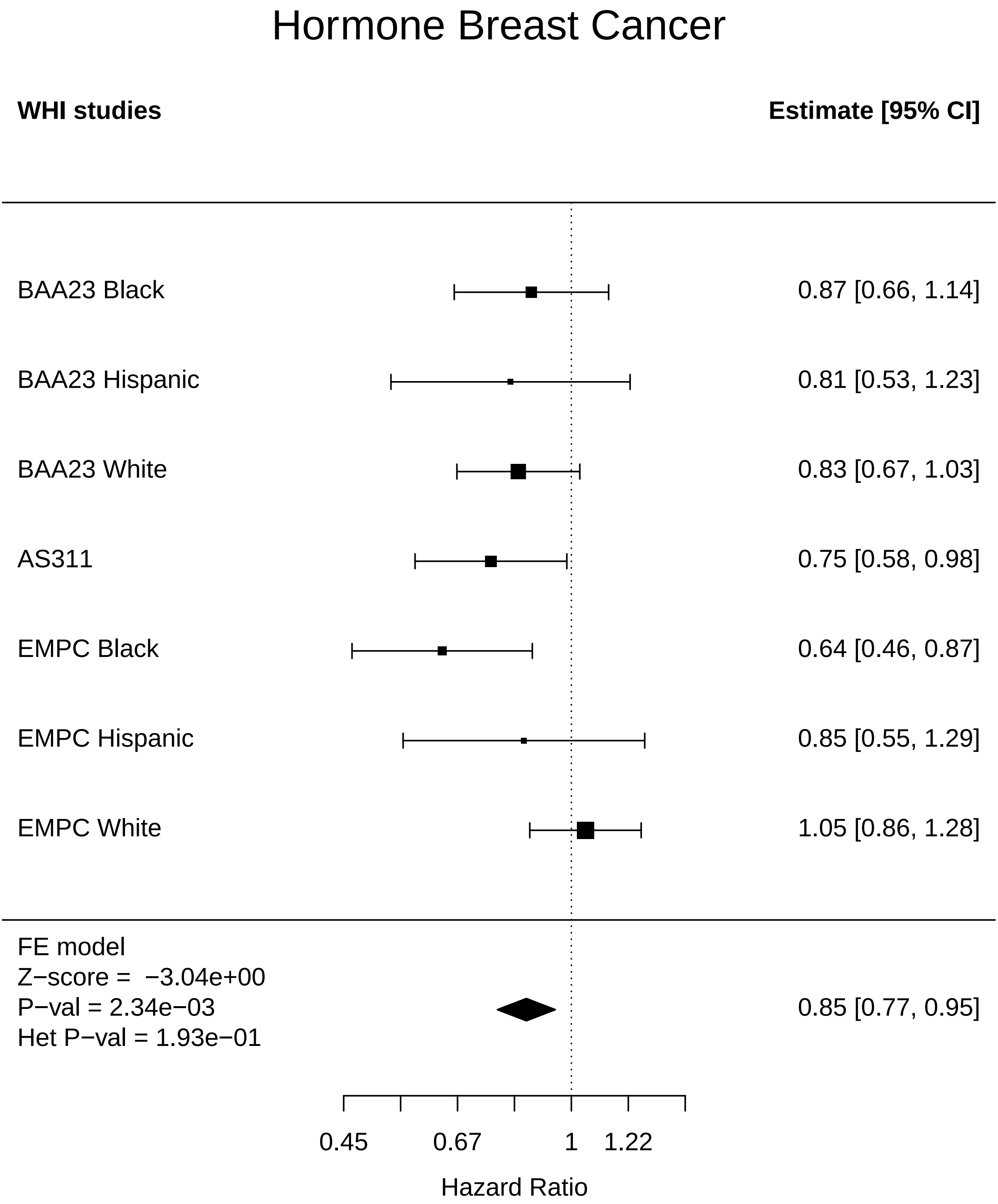

### Hormone_Cataract.pdf

#### Hormone Cataract

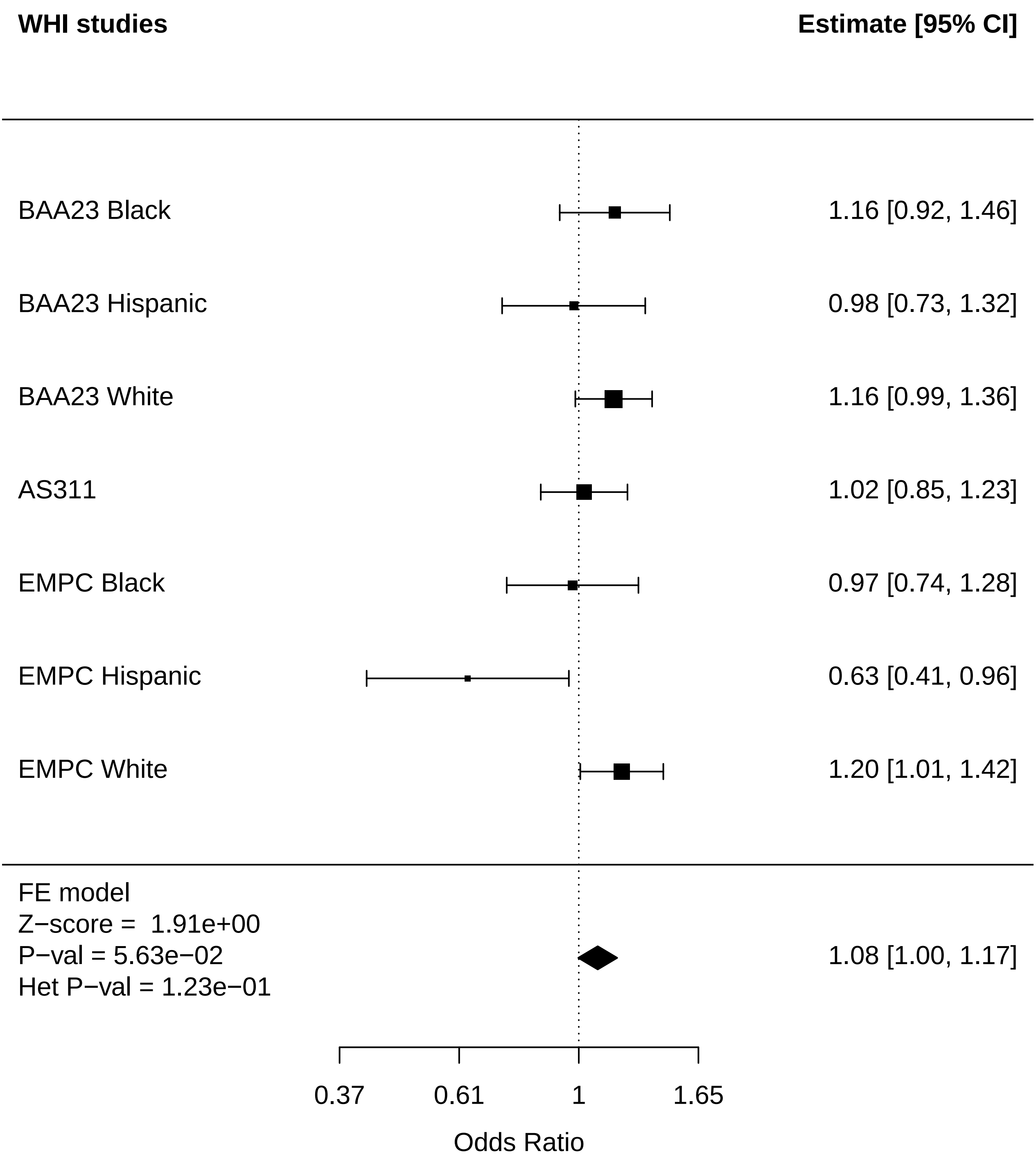

### Hormone_CHD.pdf

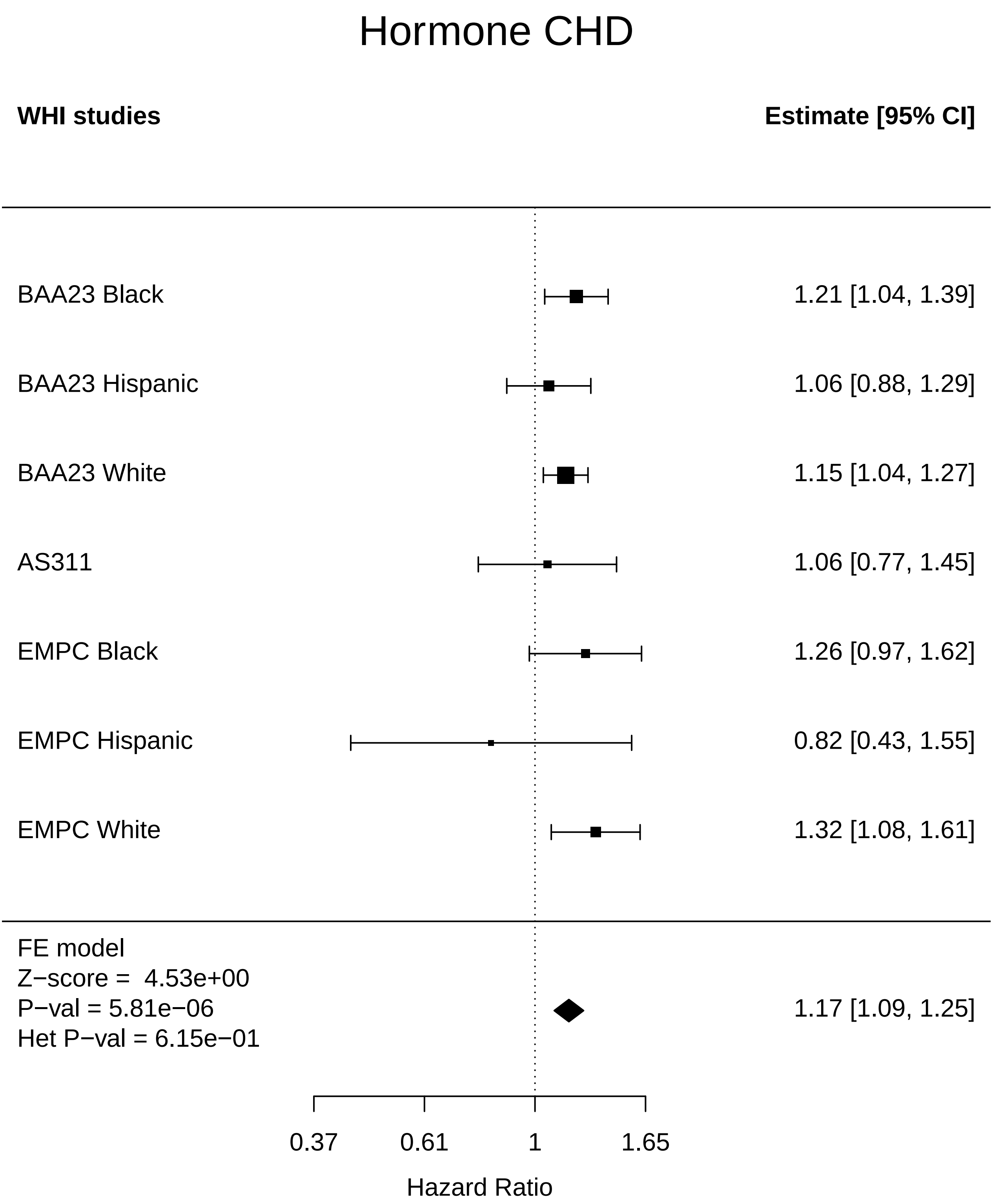

### Hormone_Cognitive function.pdf

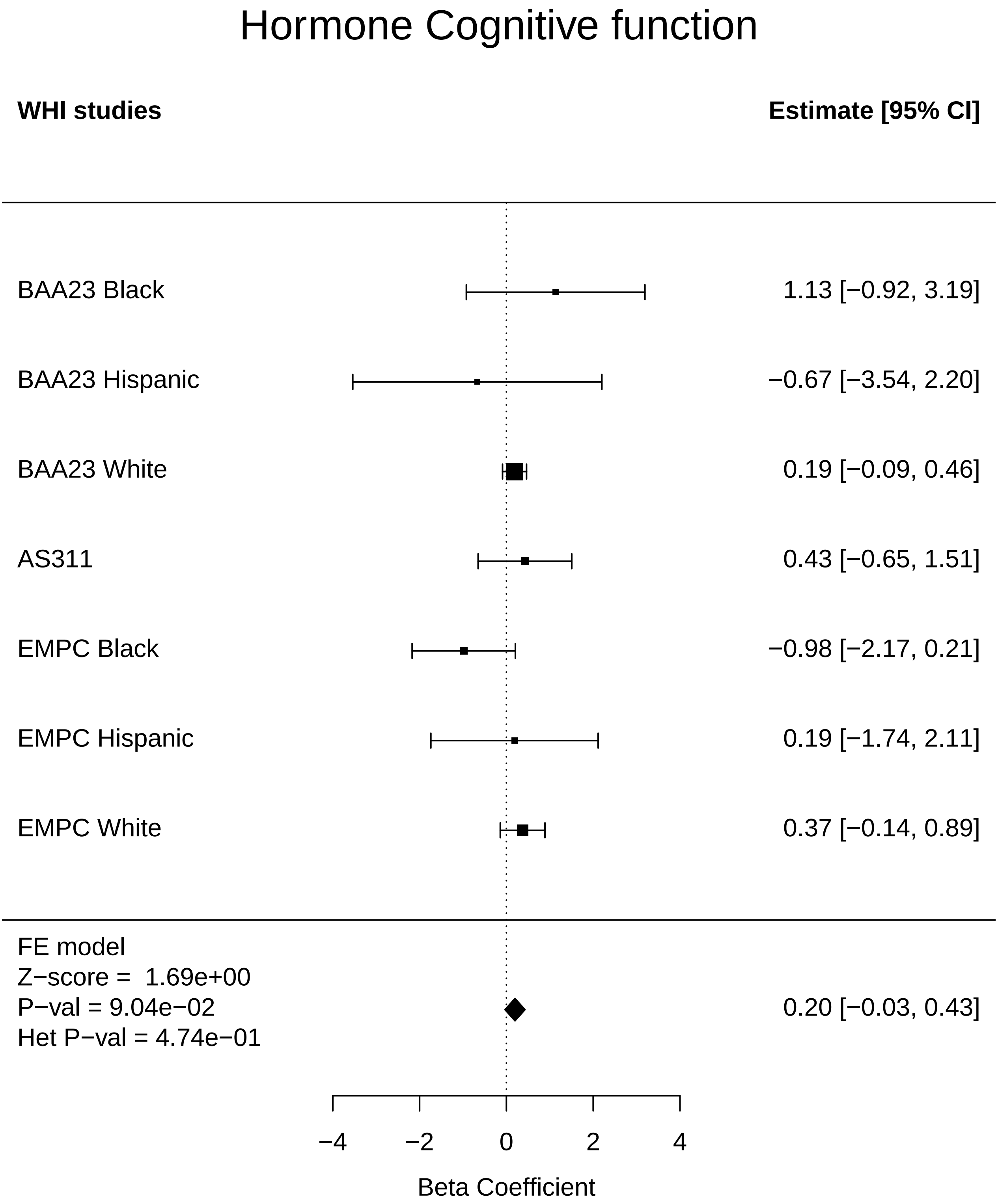

### Hormone_Colorectal cancer.pdf

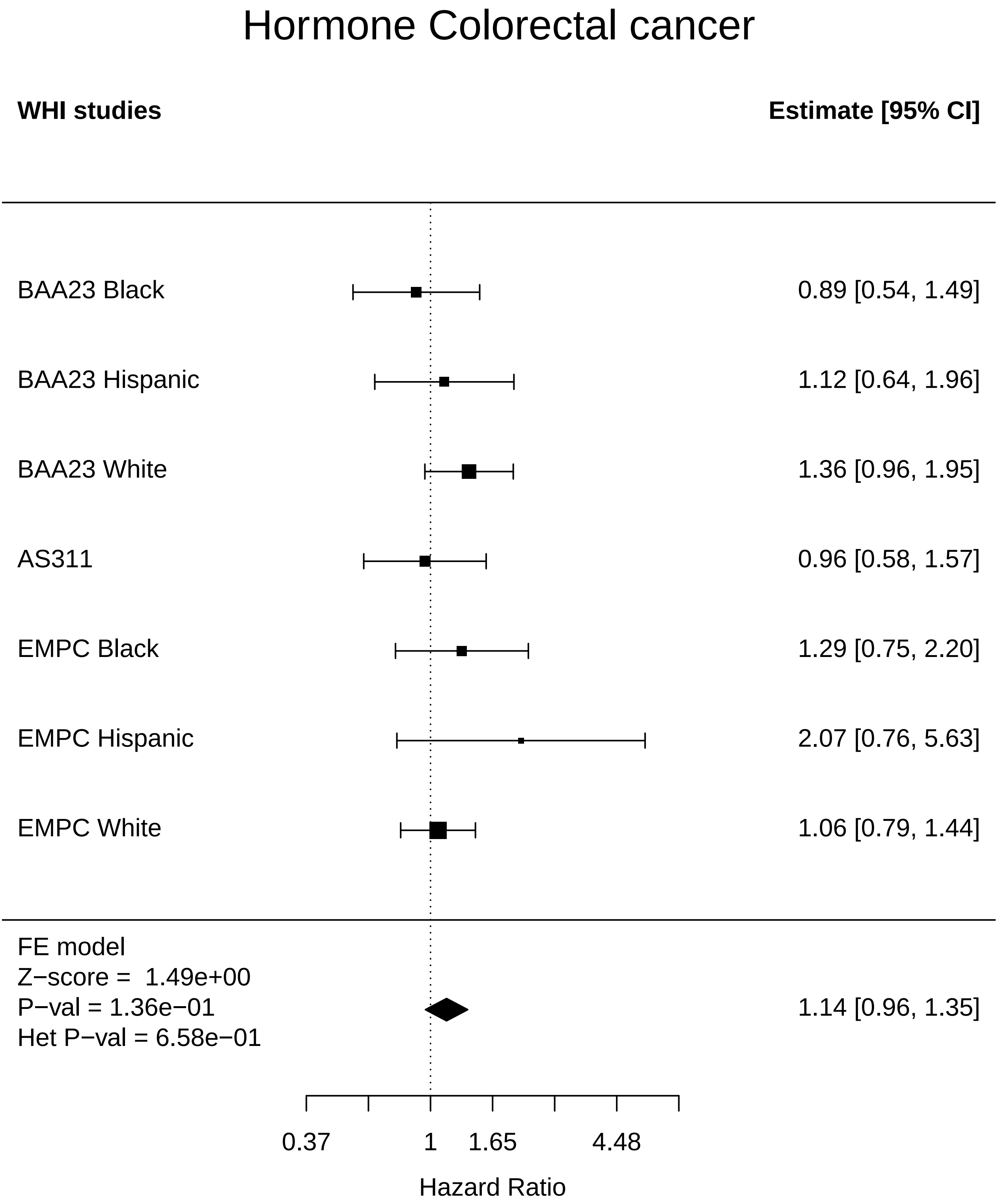

### Hormone_Diabetes.pdf

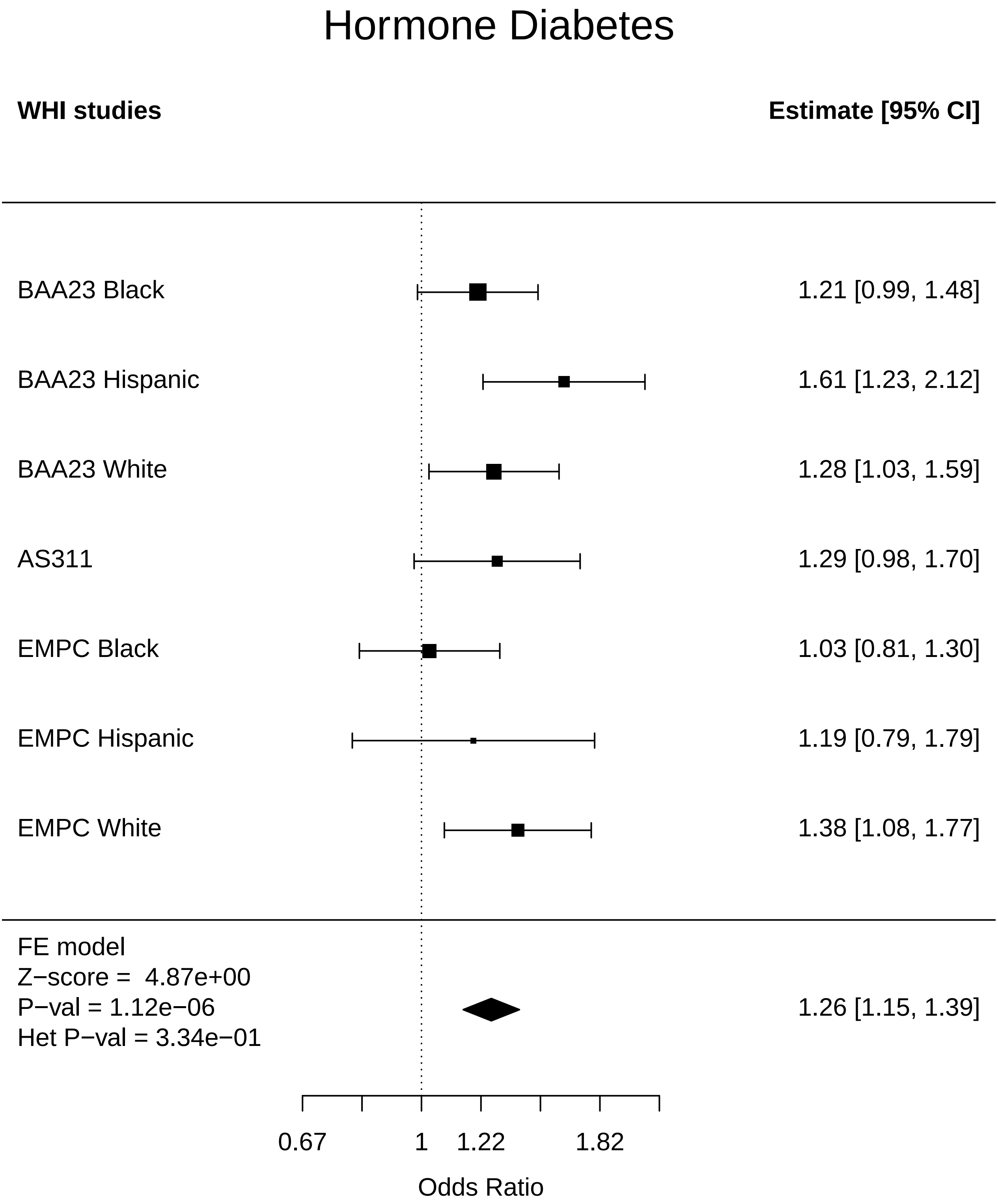

### Hormone_Disease free.pdf

#### Hormone Disease free

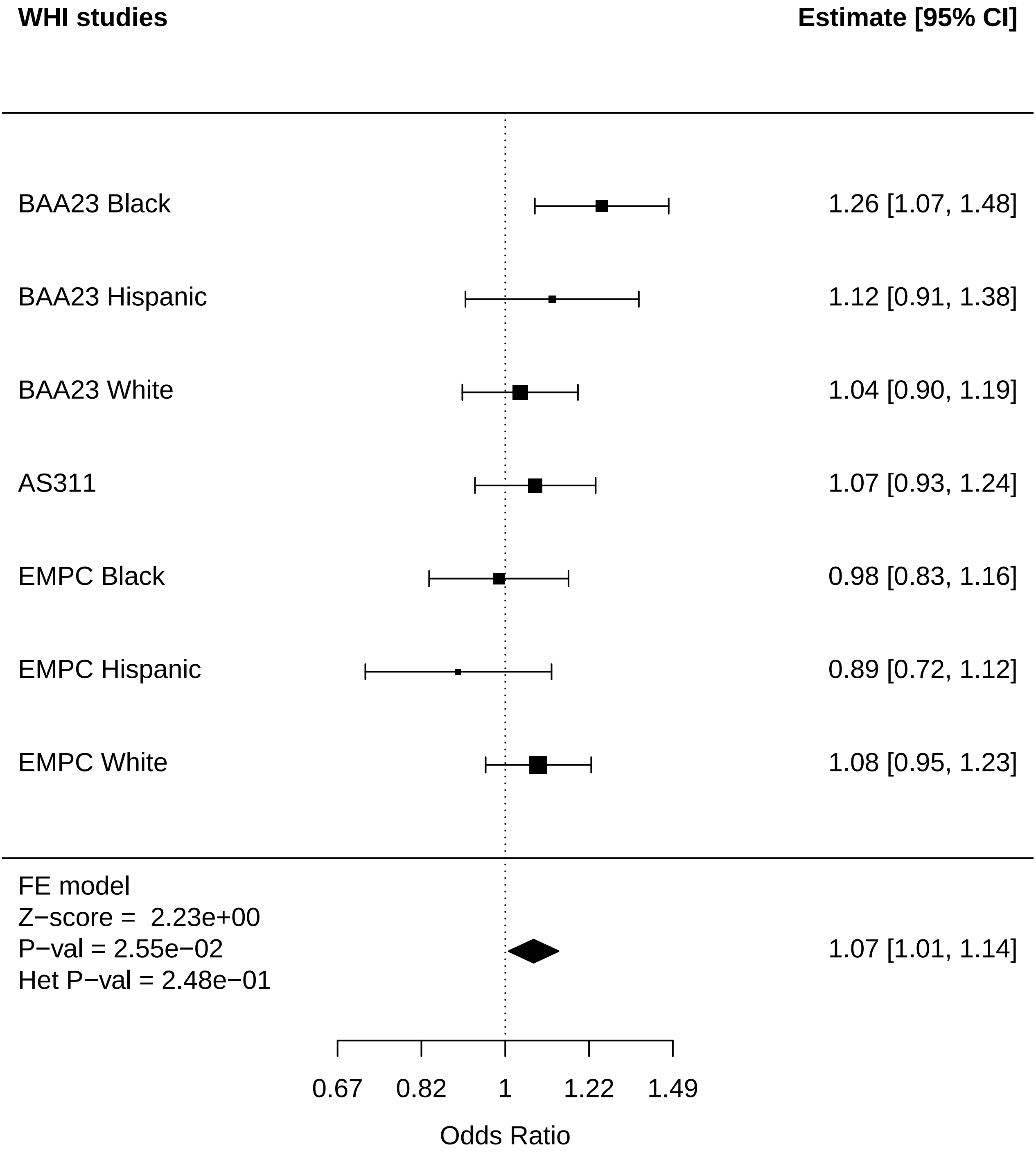

### Hormone_Endometrial cancer.pdf

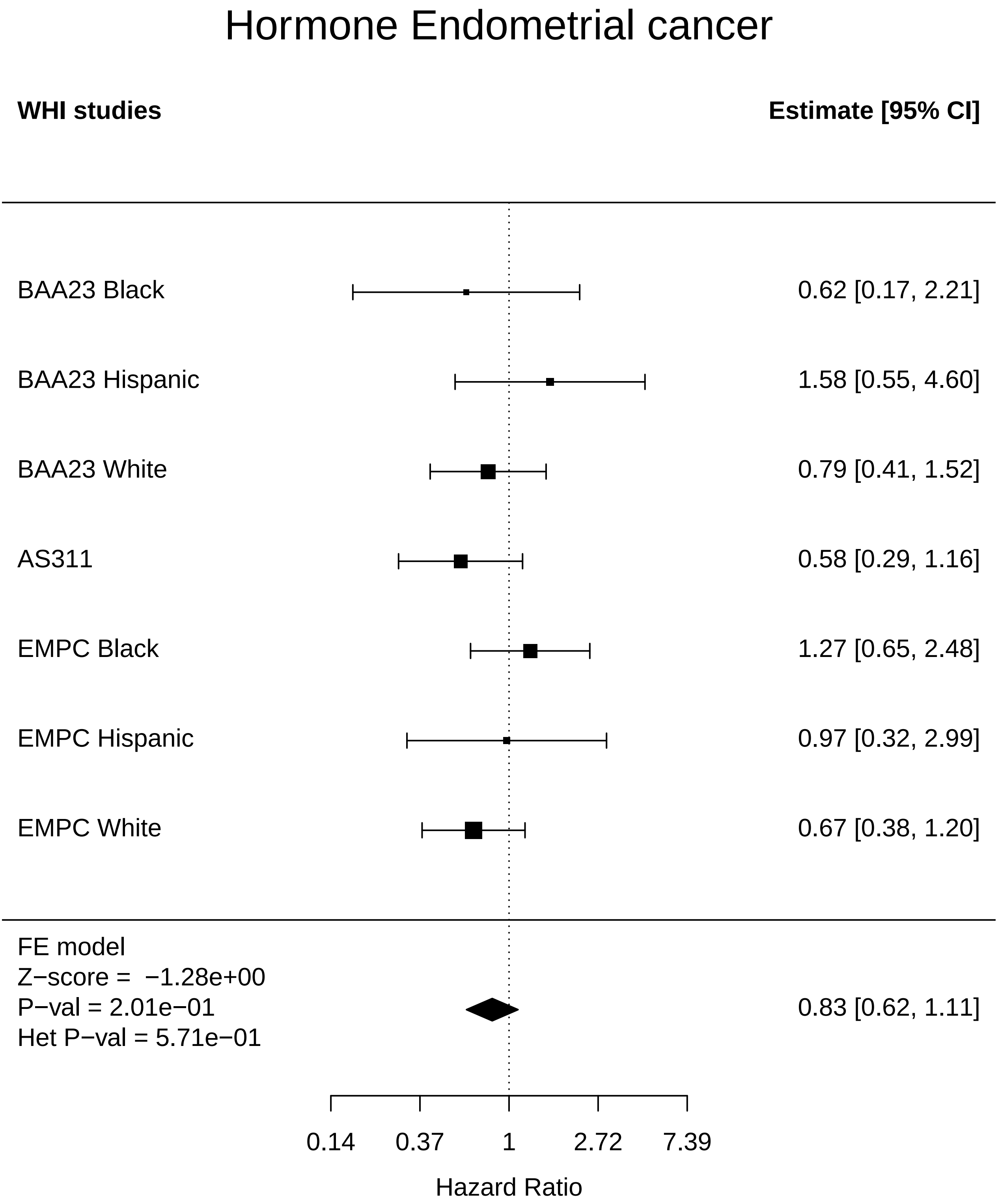

### Hormone_Lung Cancer.pdf

#### Hormone Lung Cancer

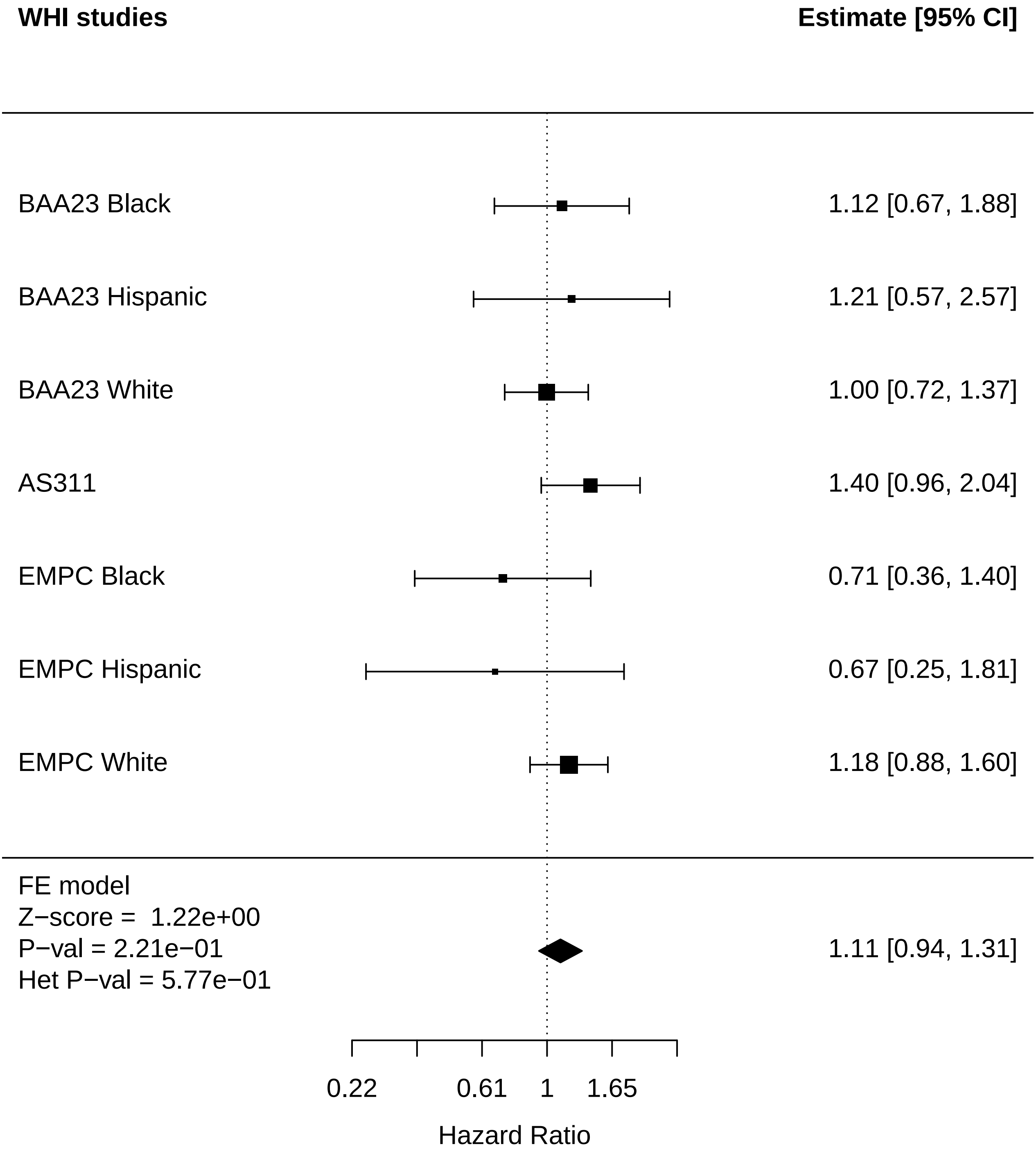

### Hormone_MI.pdf

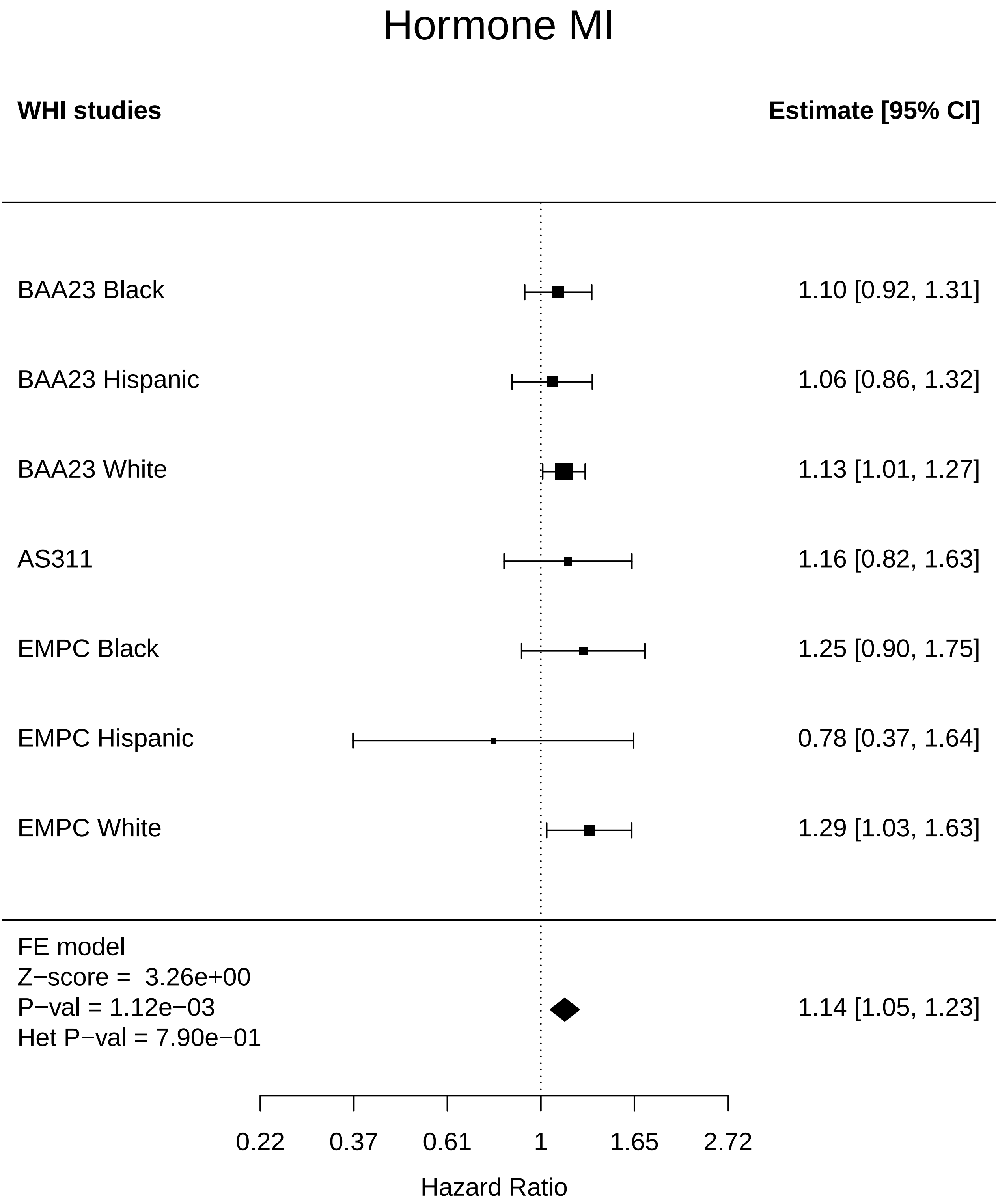

### Hormone_Mortality.pdf

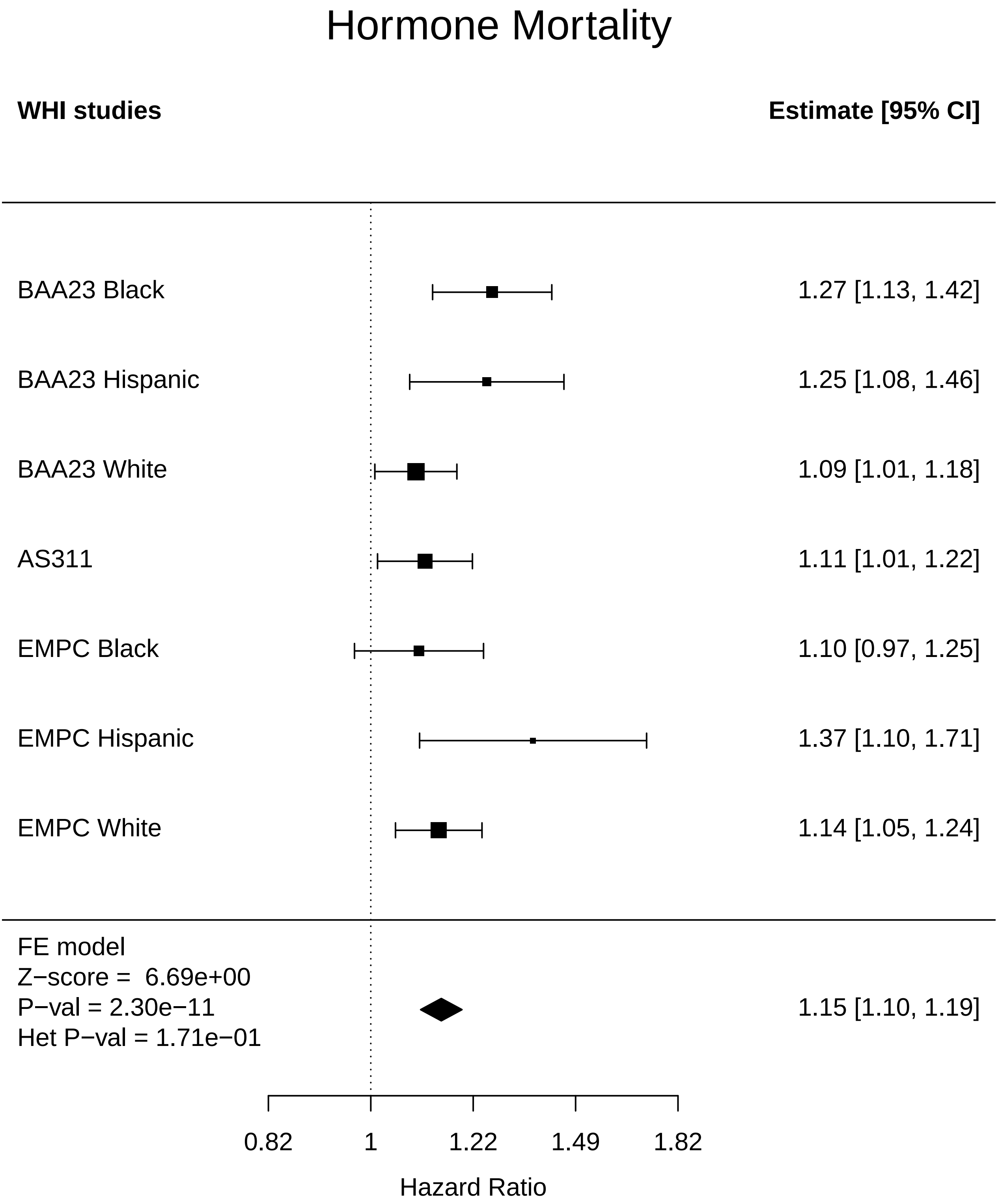

### Hormone_Physical function.pdf

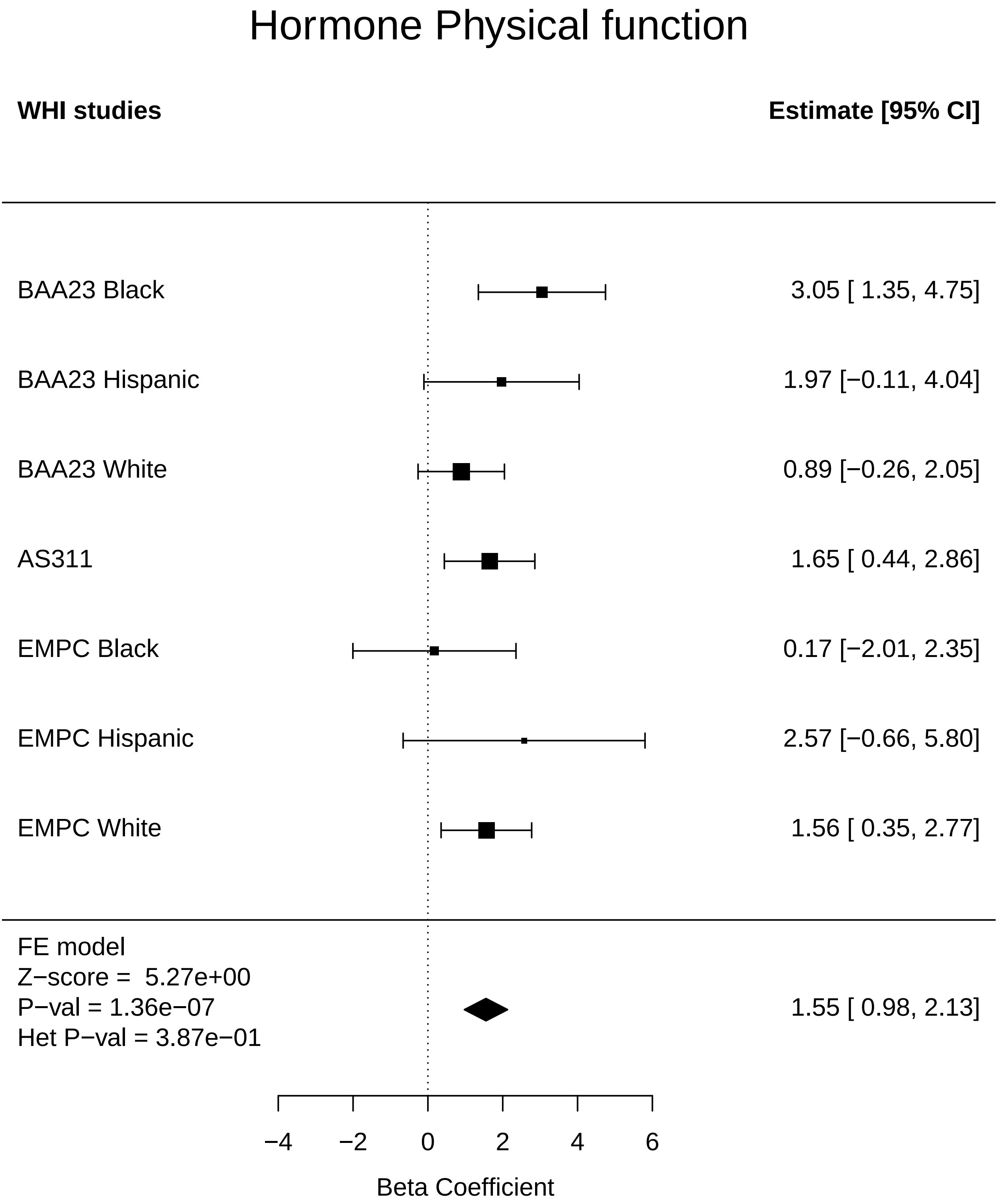

### Hormone_Stroke.pdf

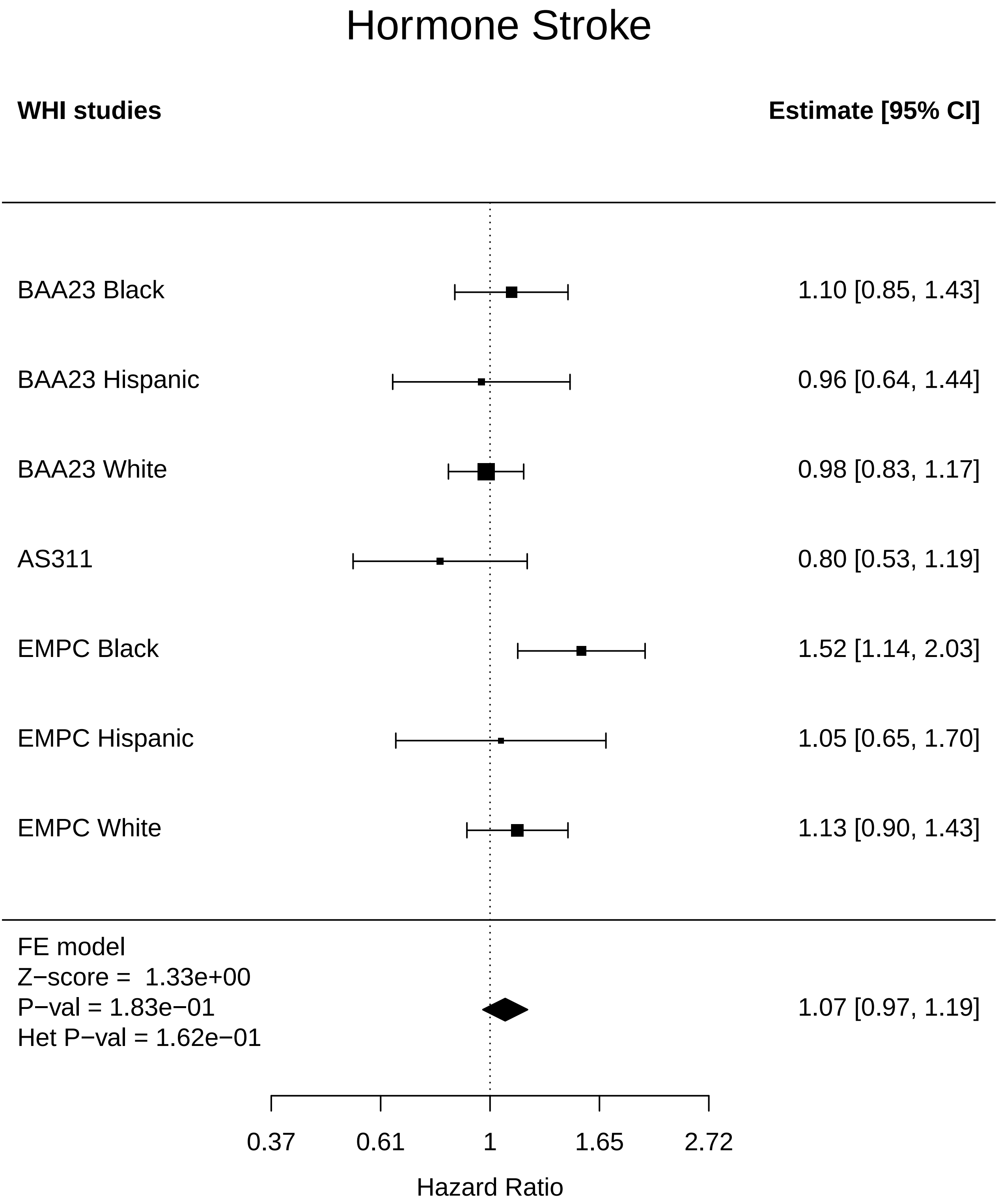

### Hormone_Thyroid disease.pdf

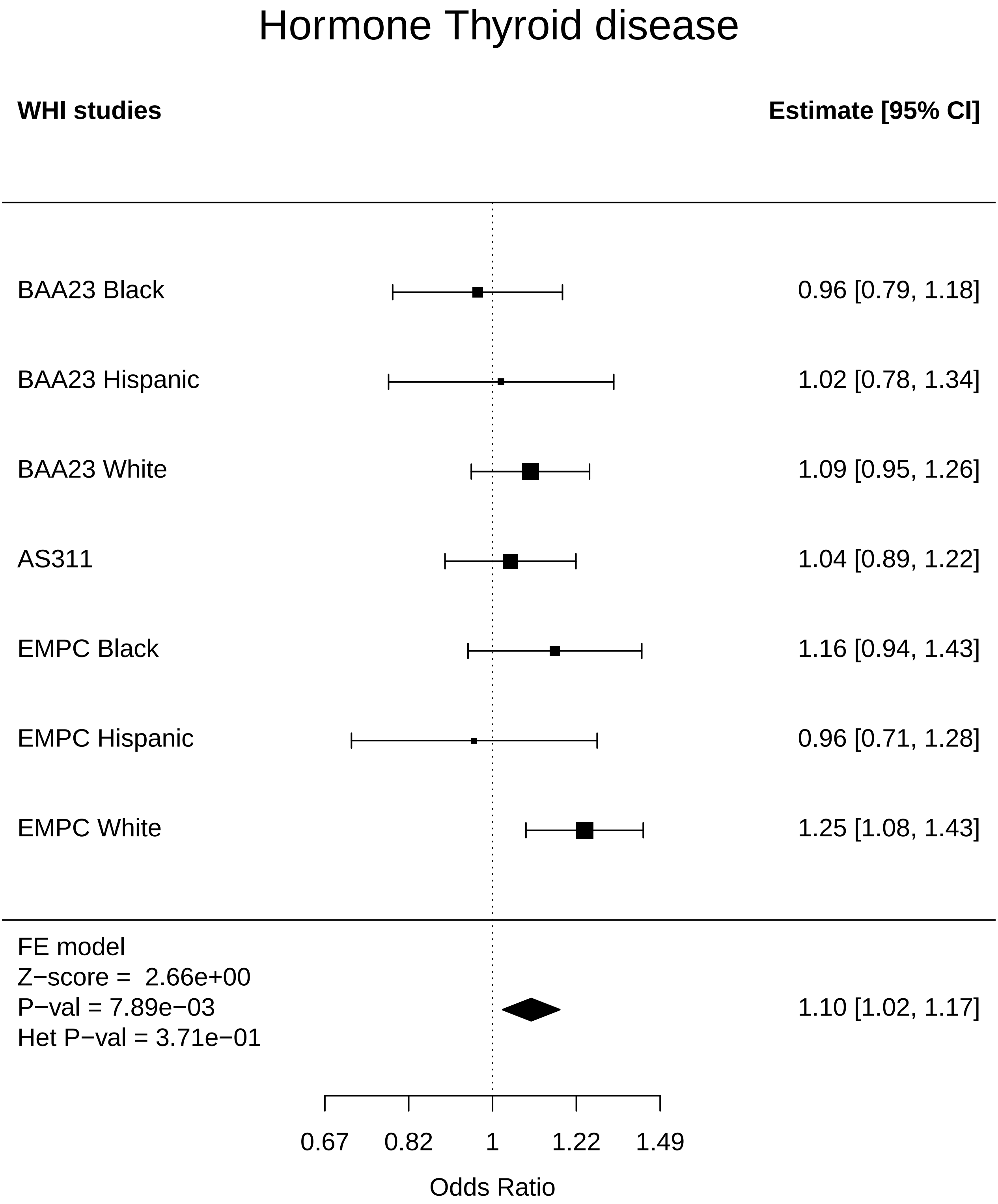

### Hormone_Total comorbidities.pdf

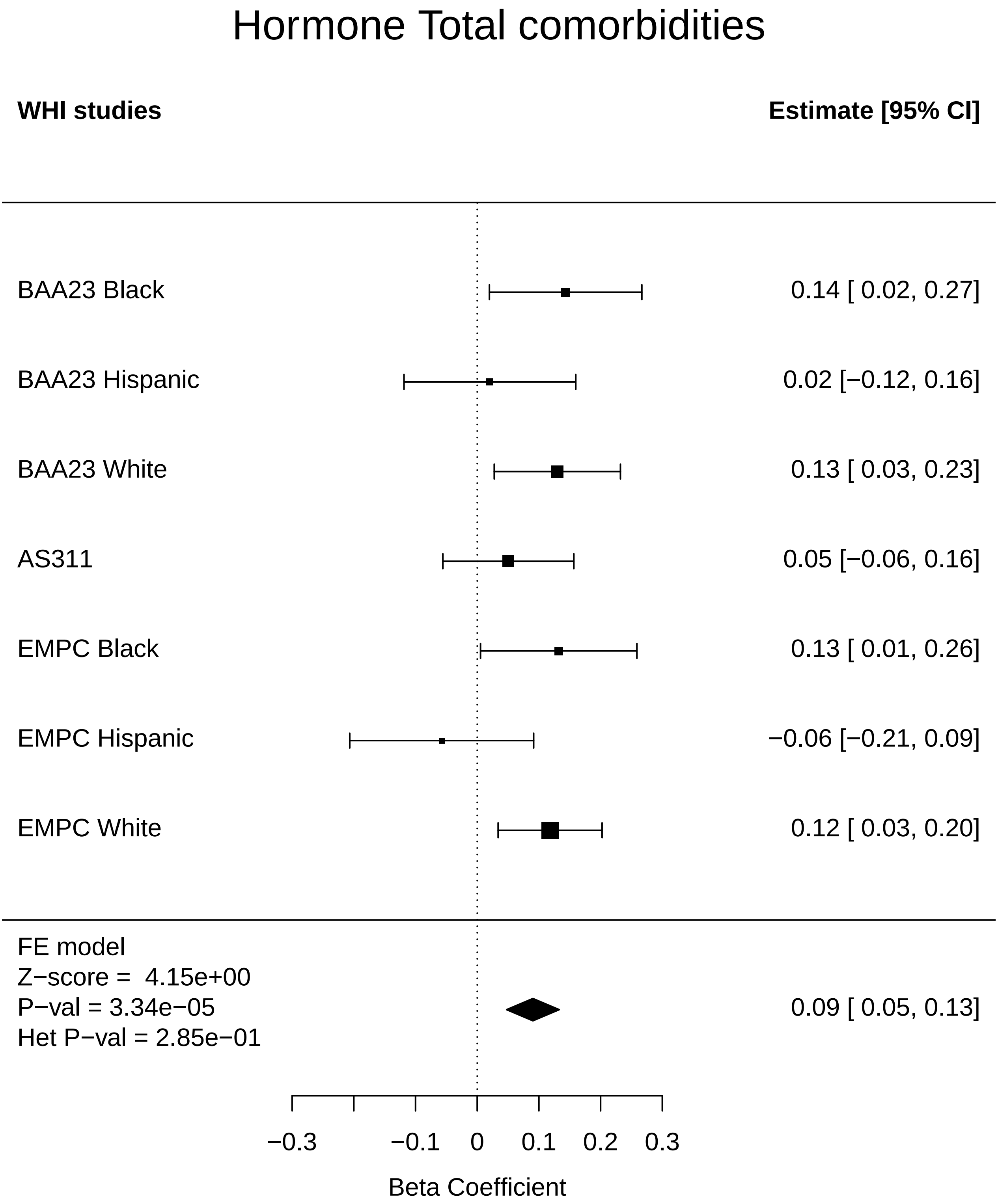

### Immune_Arthritis.pdf

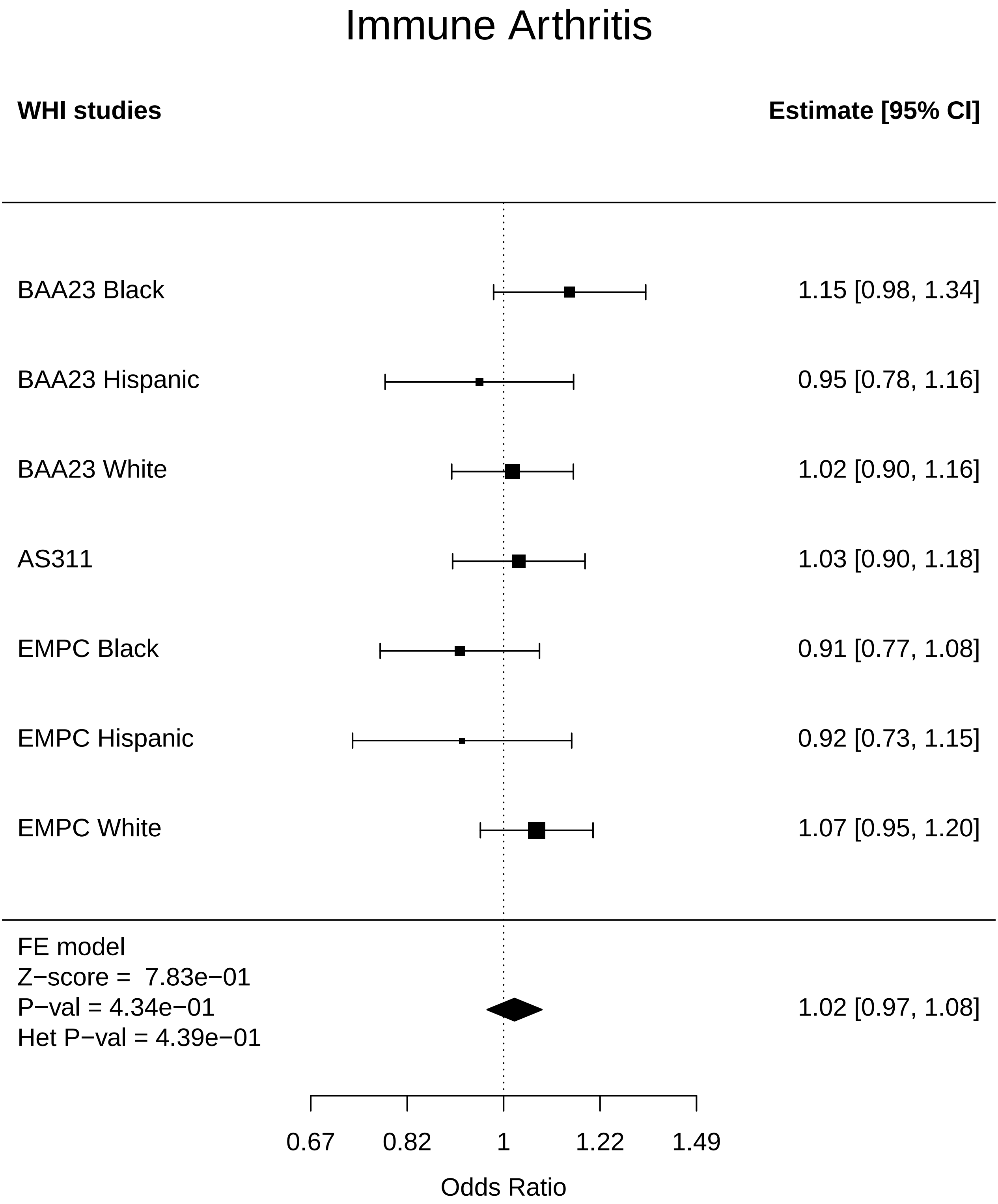

### Immune_CHD.pdf

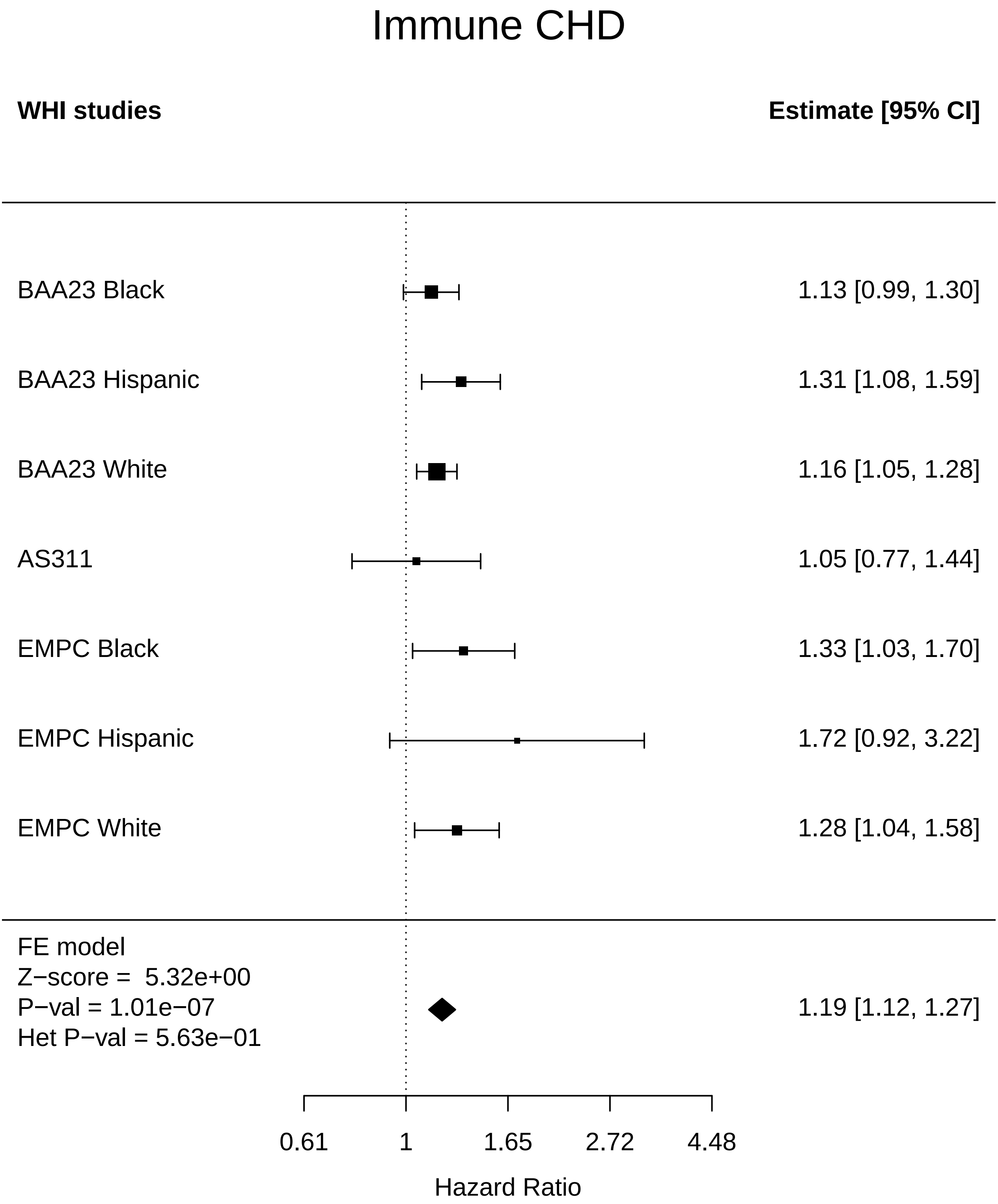

### Immune_Colorectal cancer.pdf

#### Immune Colorectal cancer

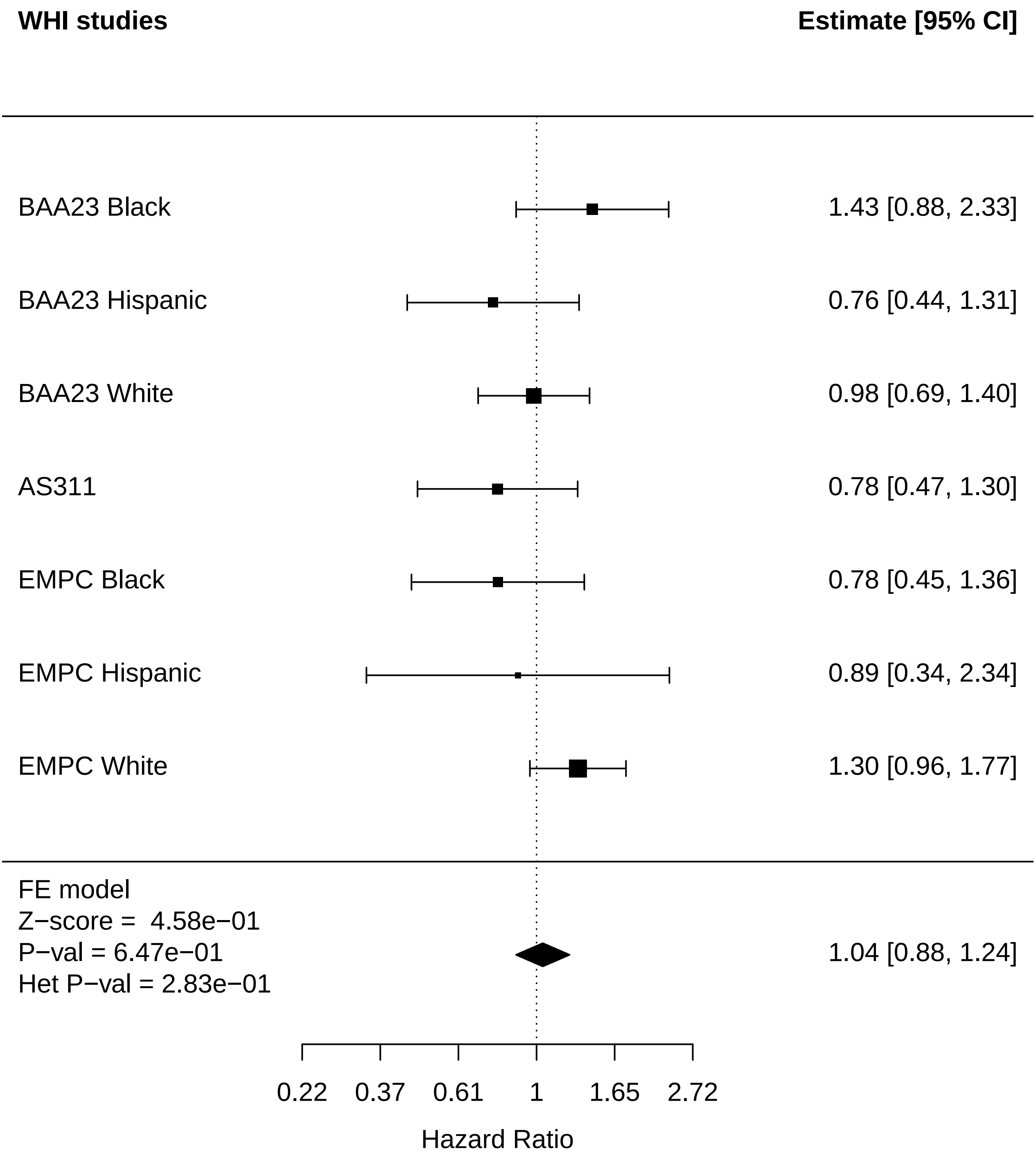

### Immune_Diabetes.pdf

#### Immune Diabetes

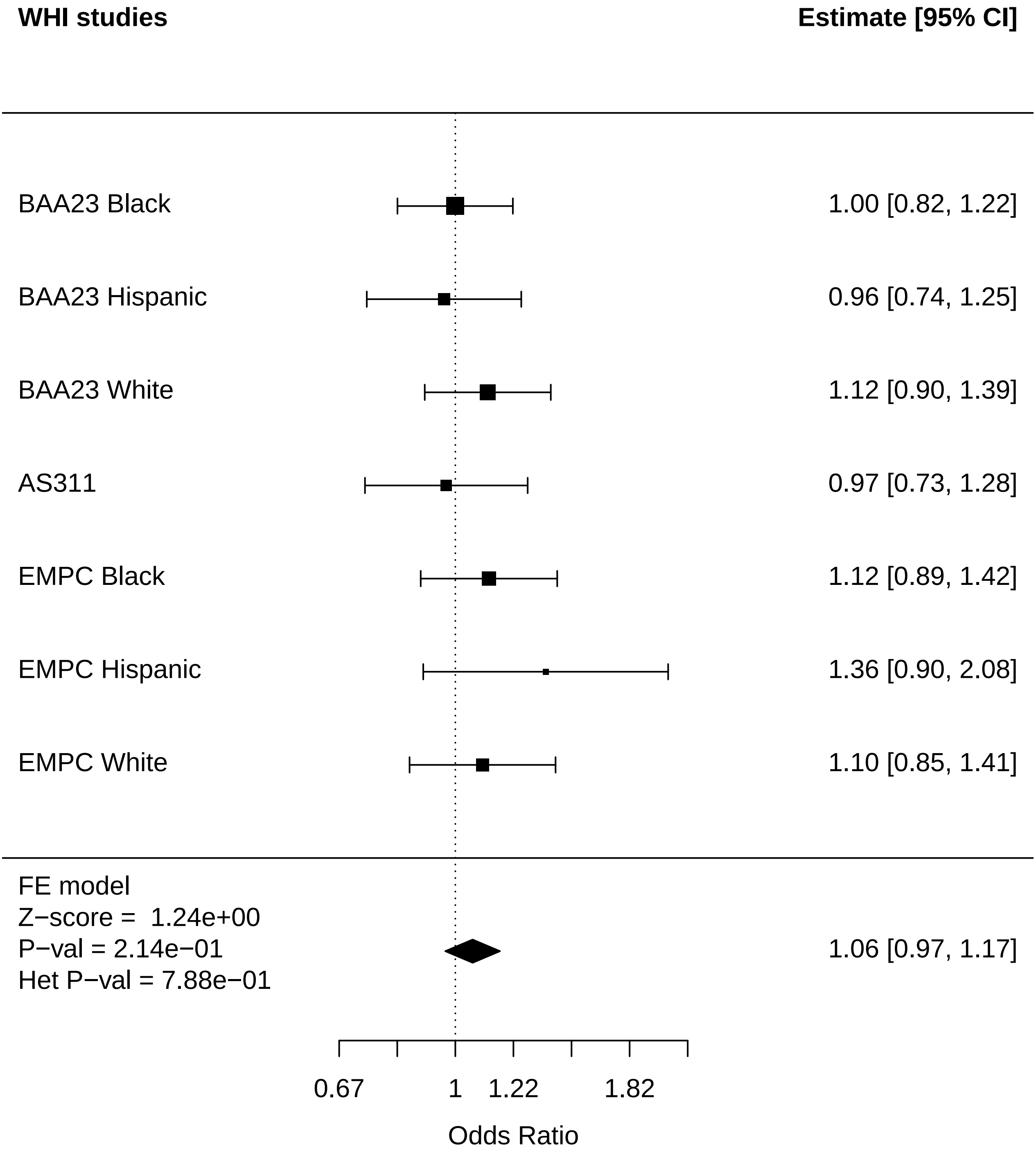

### Immune_Disease free.pdf

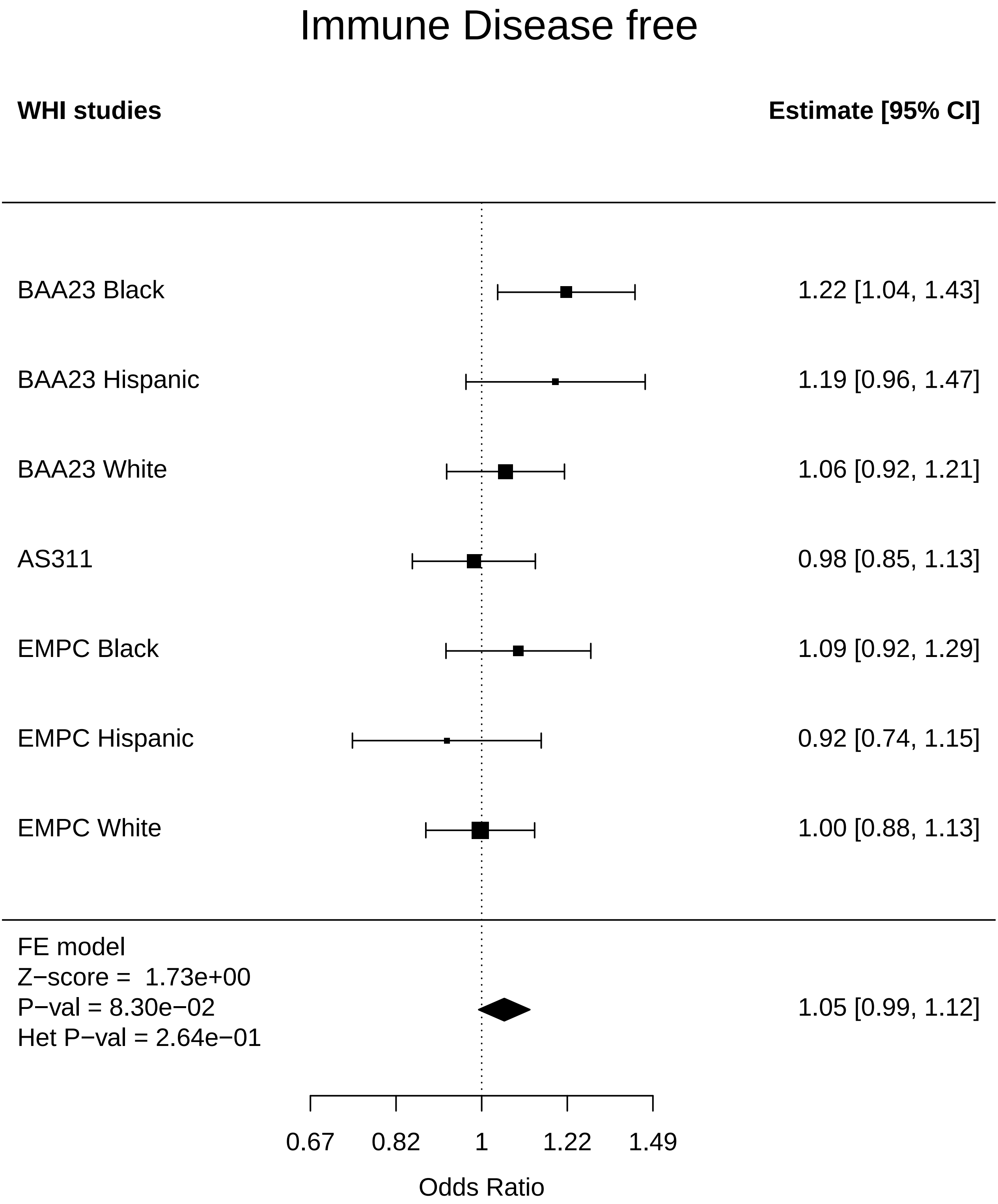

### Immune_Endometrial cancer.pdf

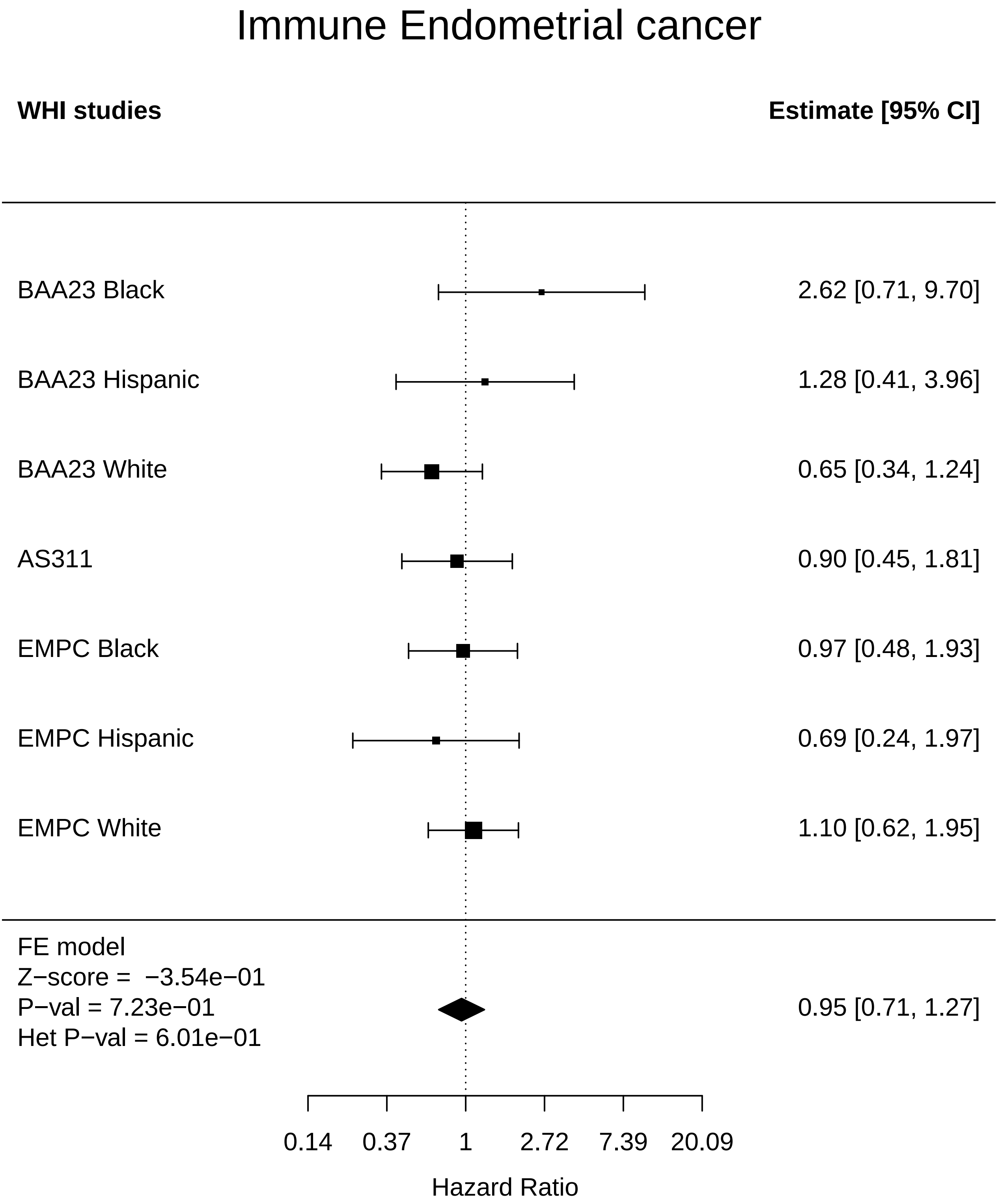

### Immune_Leukemia.pdf

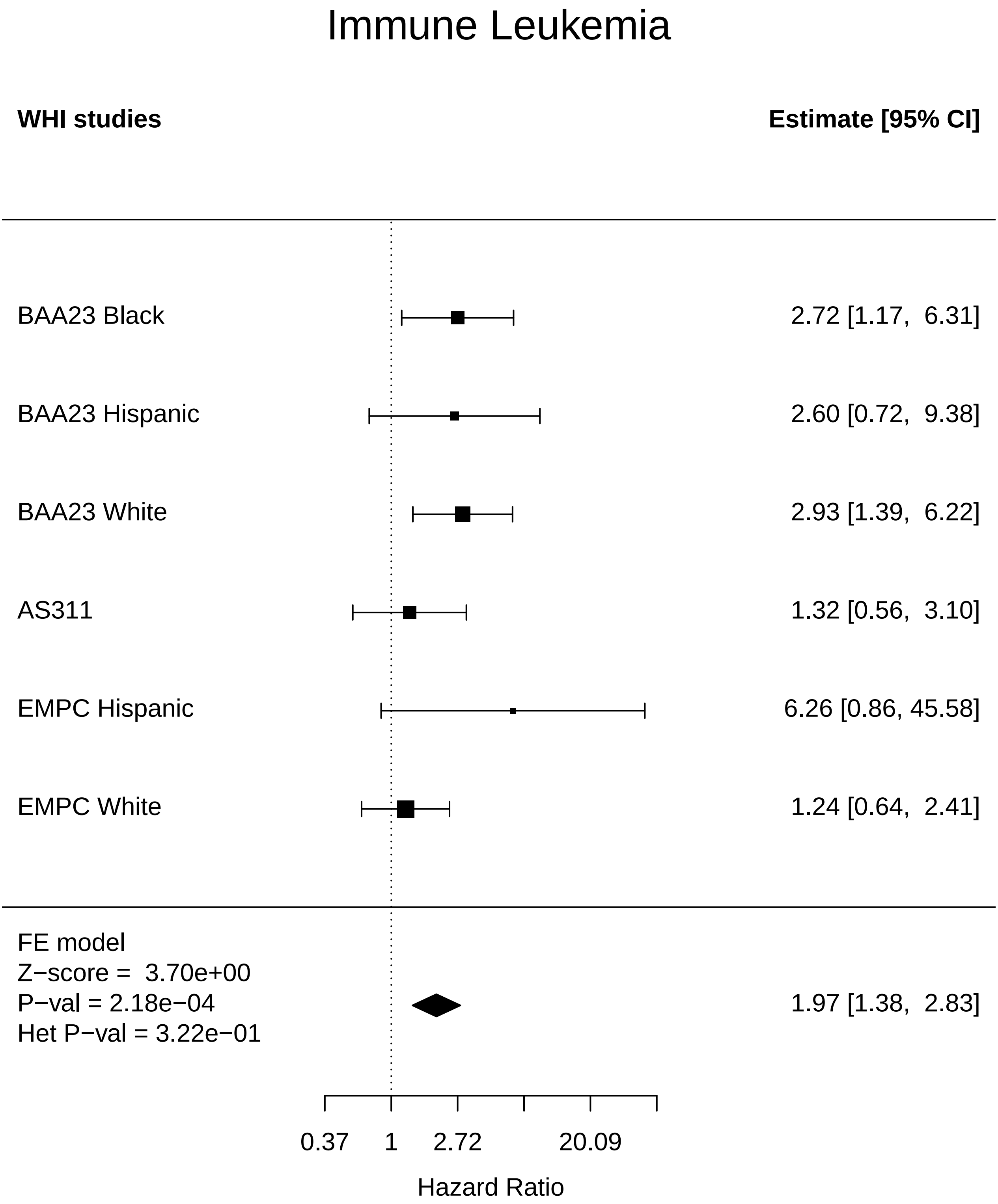

### Immune_MI.pdf

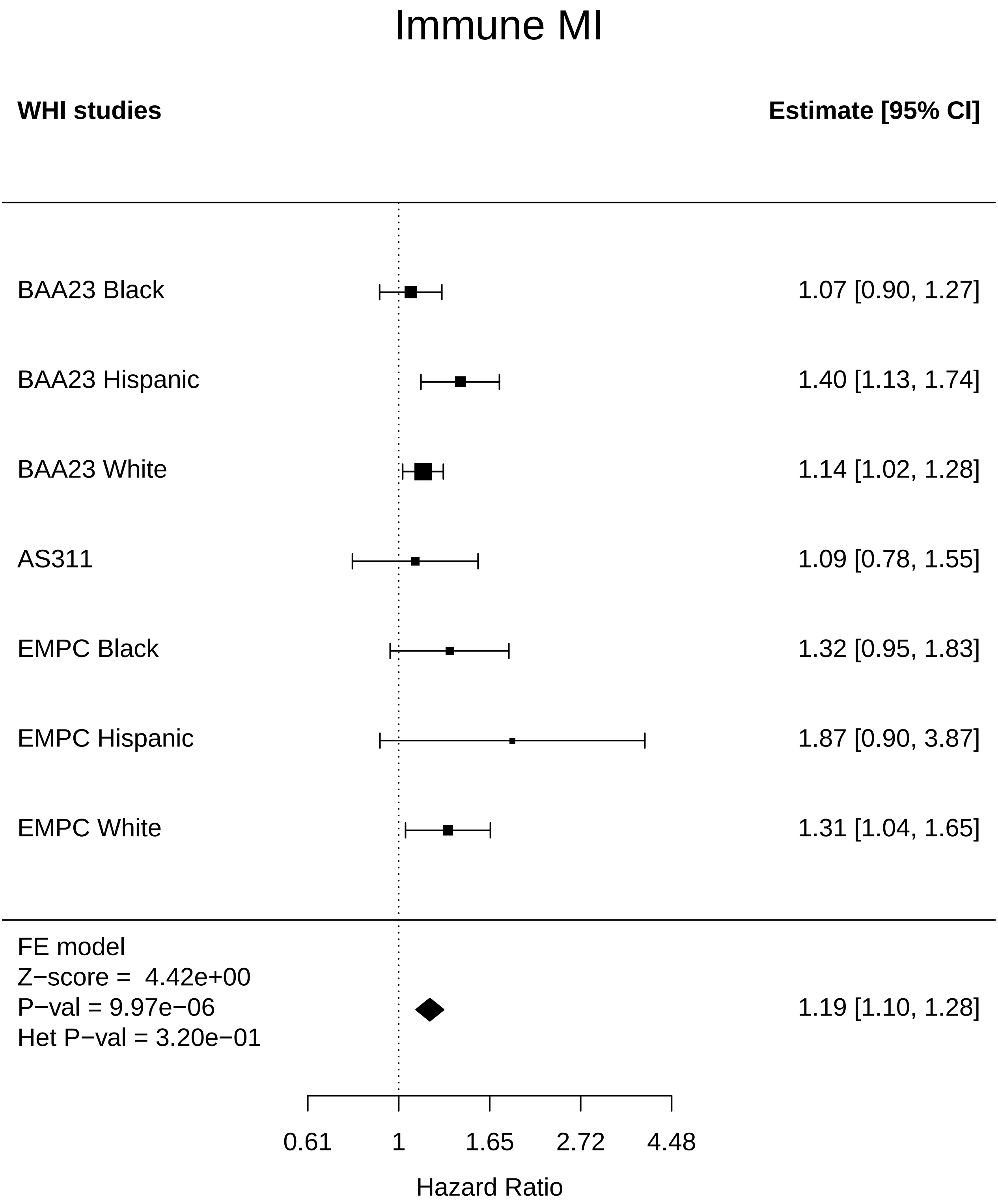

### Immune_Physical function.pdf

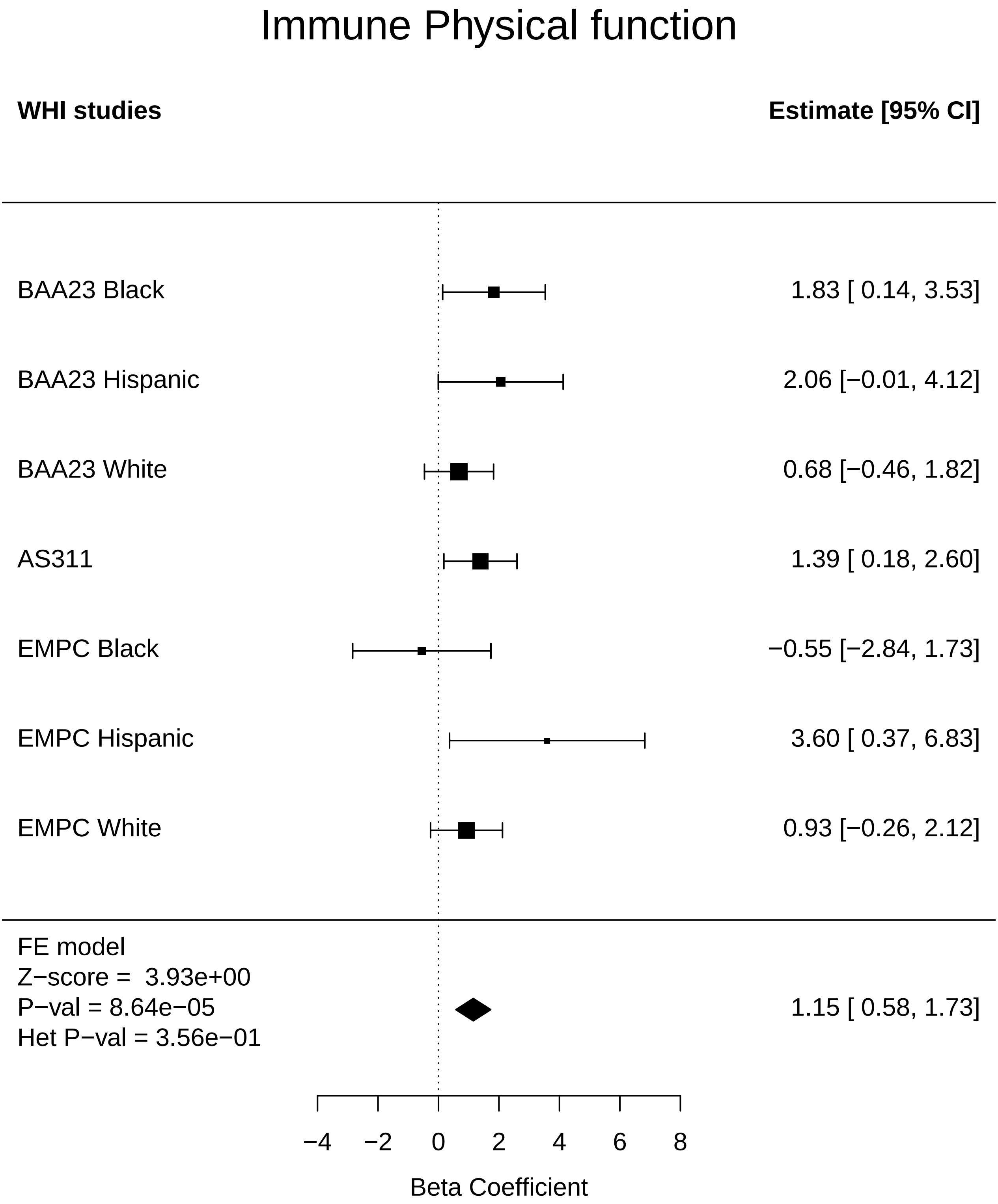

### Immune_Stroke.pdf

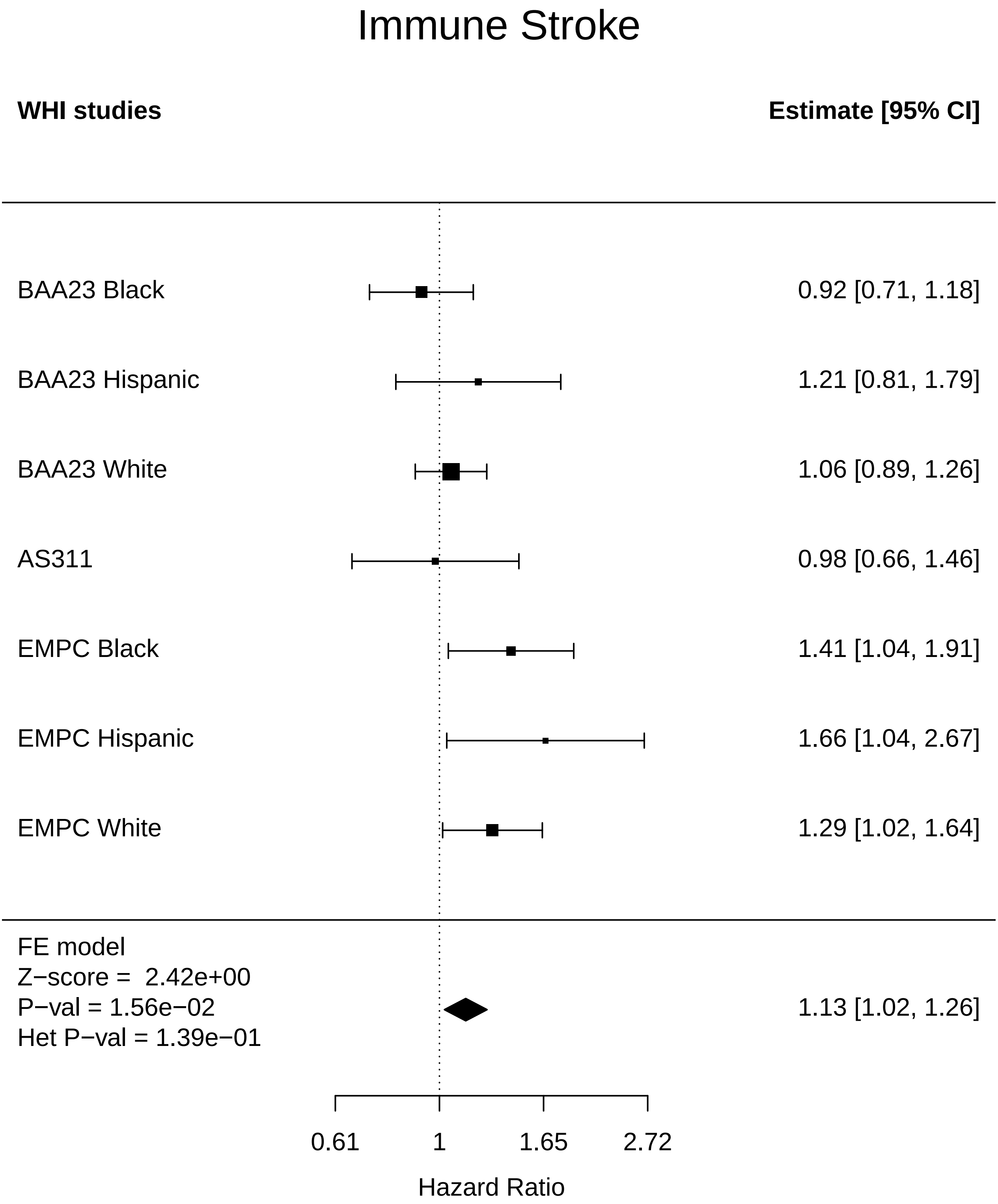

### Immune_Total comorbidities.pdf

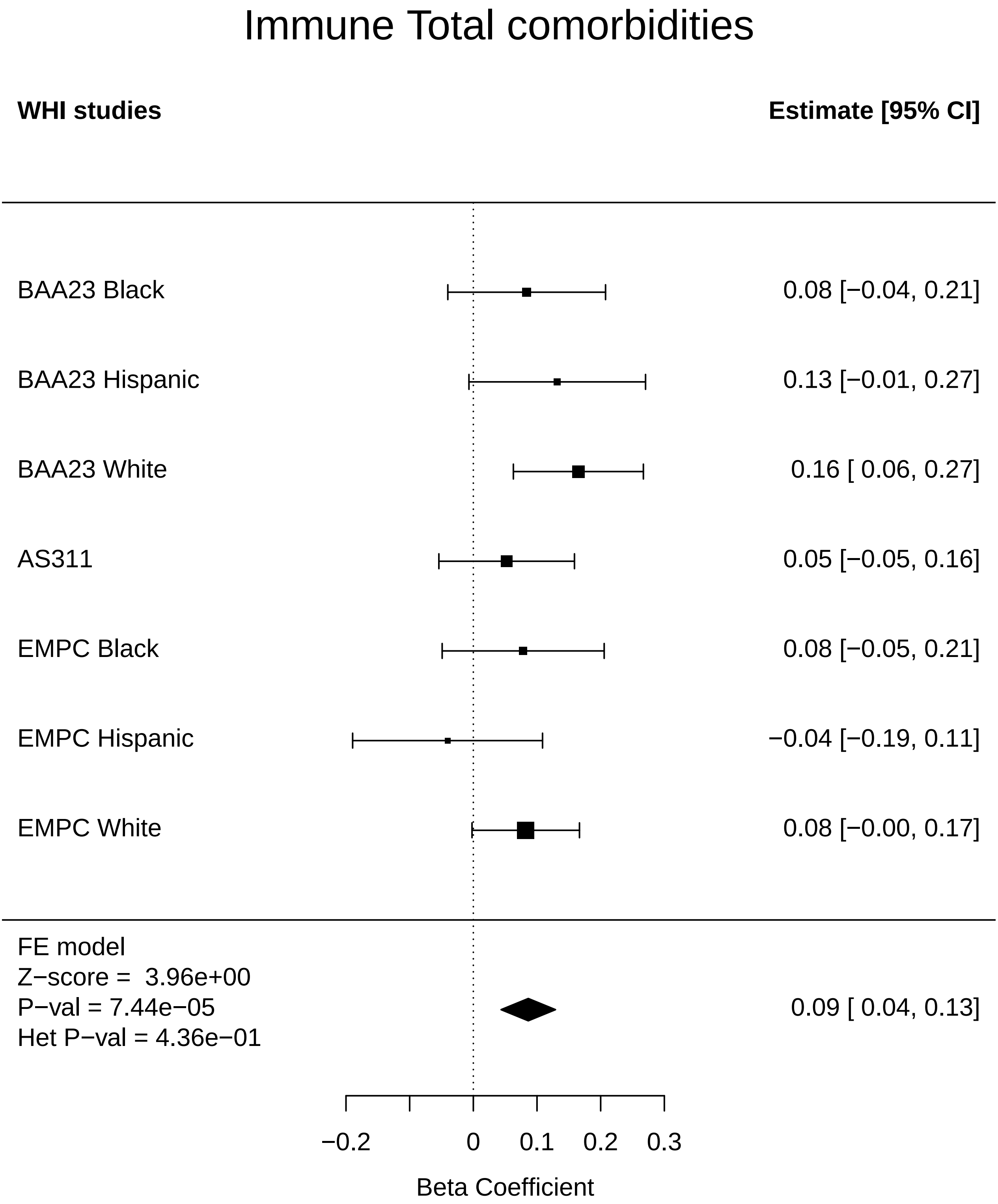

### Inflammation_Any Cancer.pdf

#### Inflammation Any Cancer

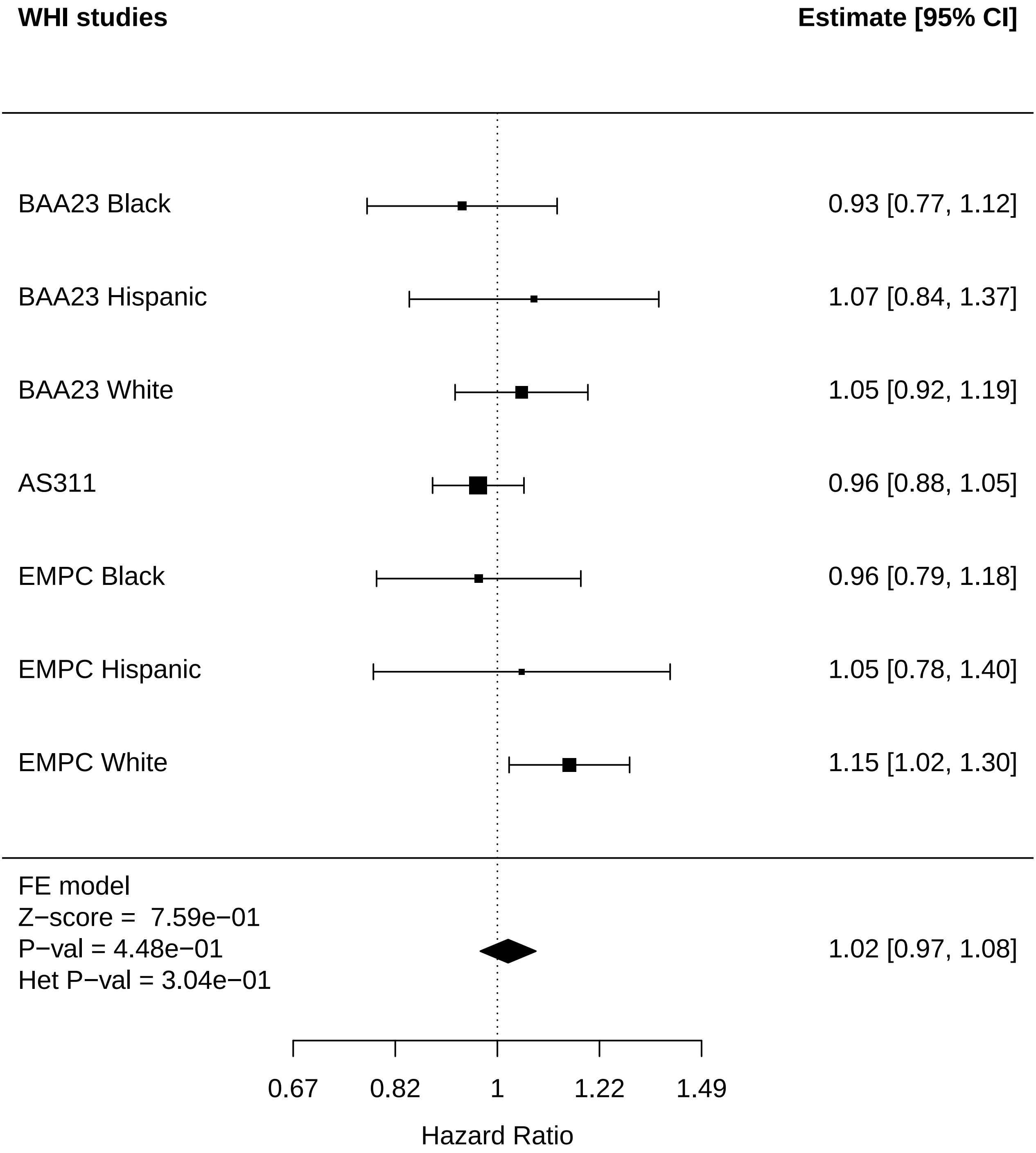

### Inflammation_Arthritis.pdf

#### Inflammation Arthritis

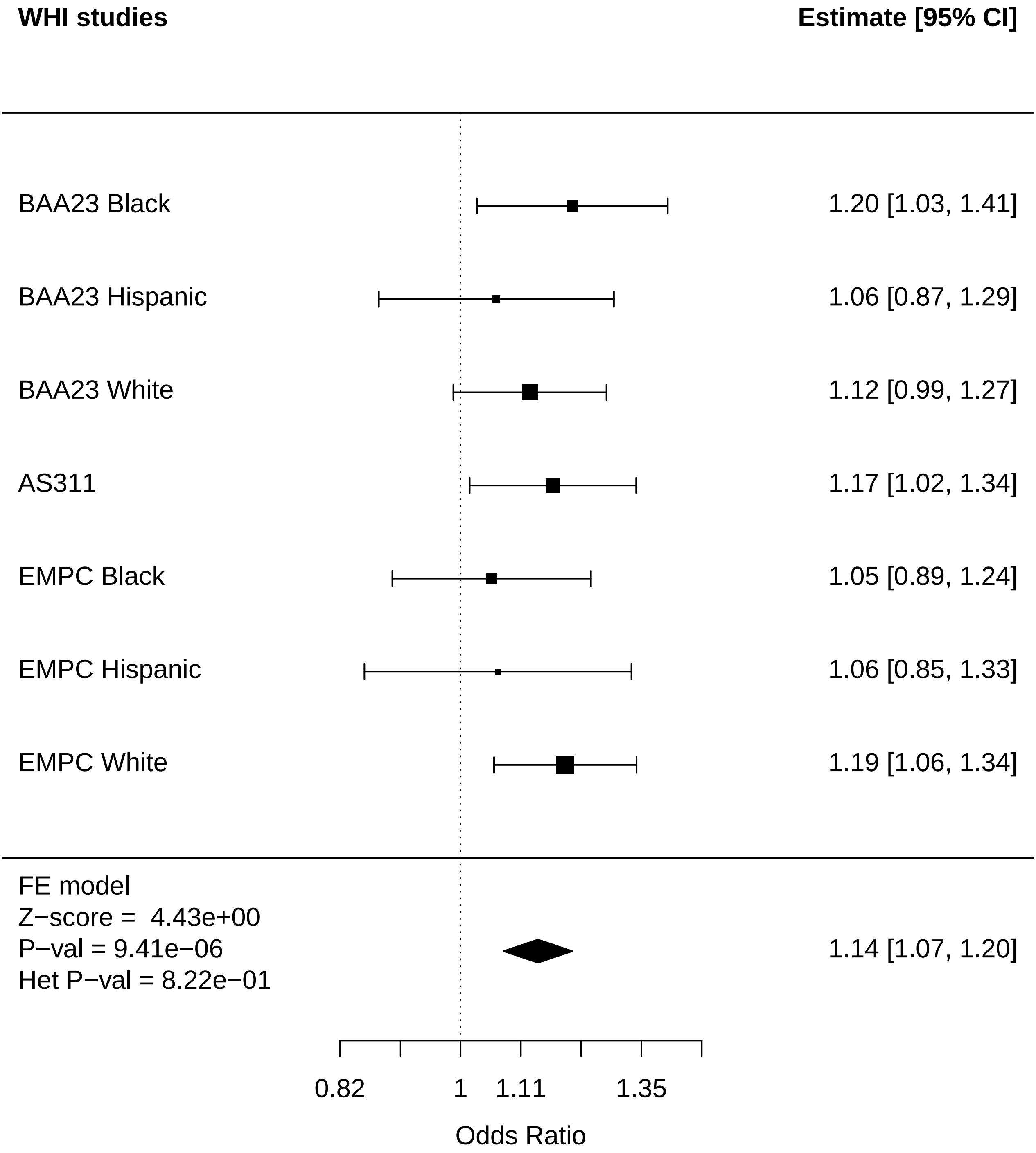

### Inflammation_Breast Cancer.pdf

#### Inflammation Breast Cancer

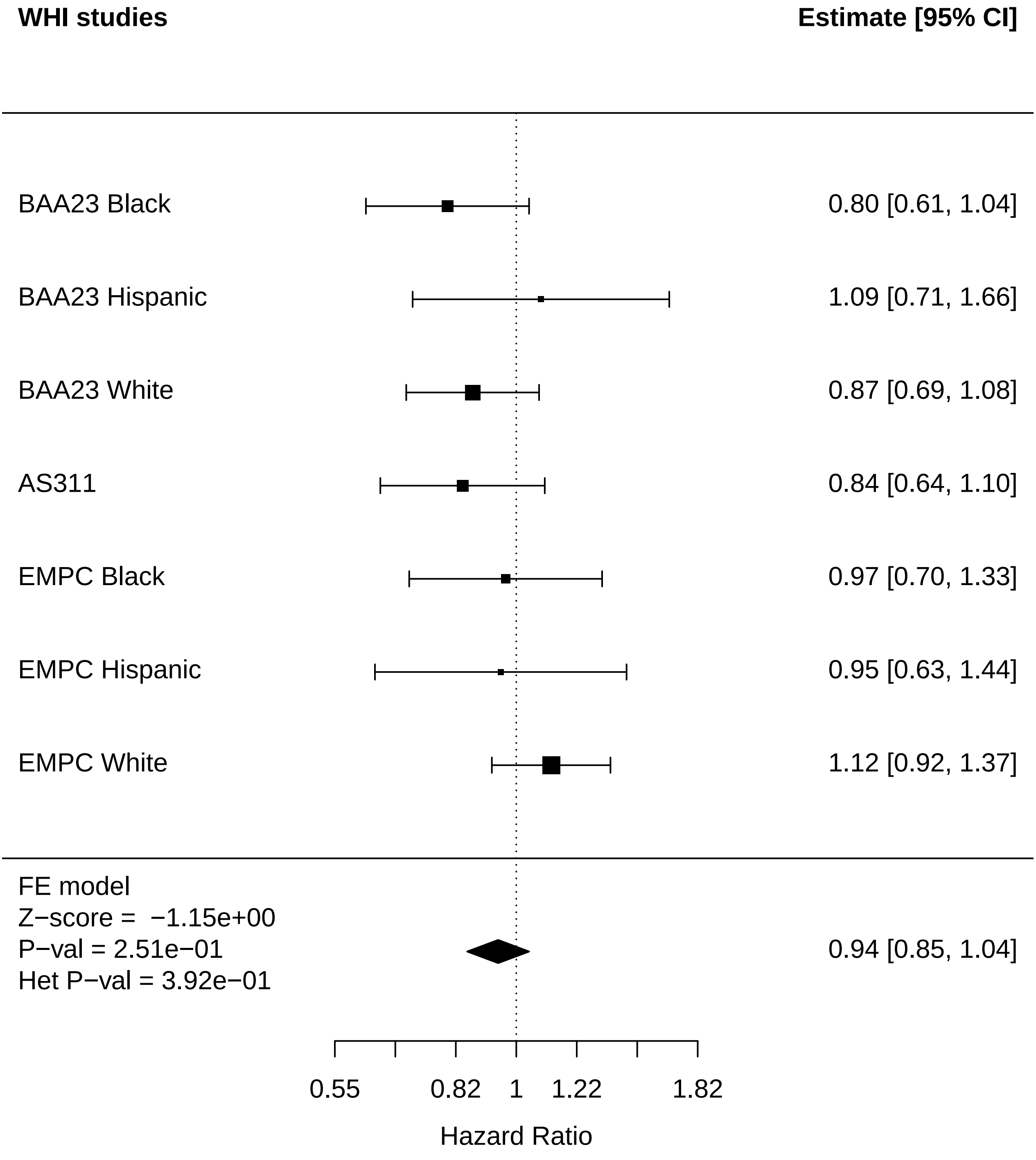

### Inflammation_Cataract.pdf

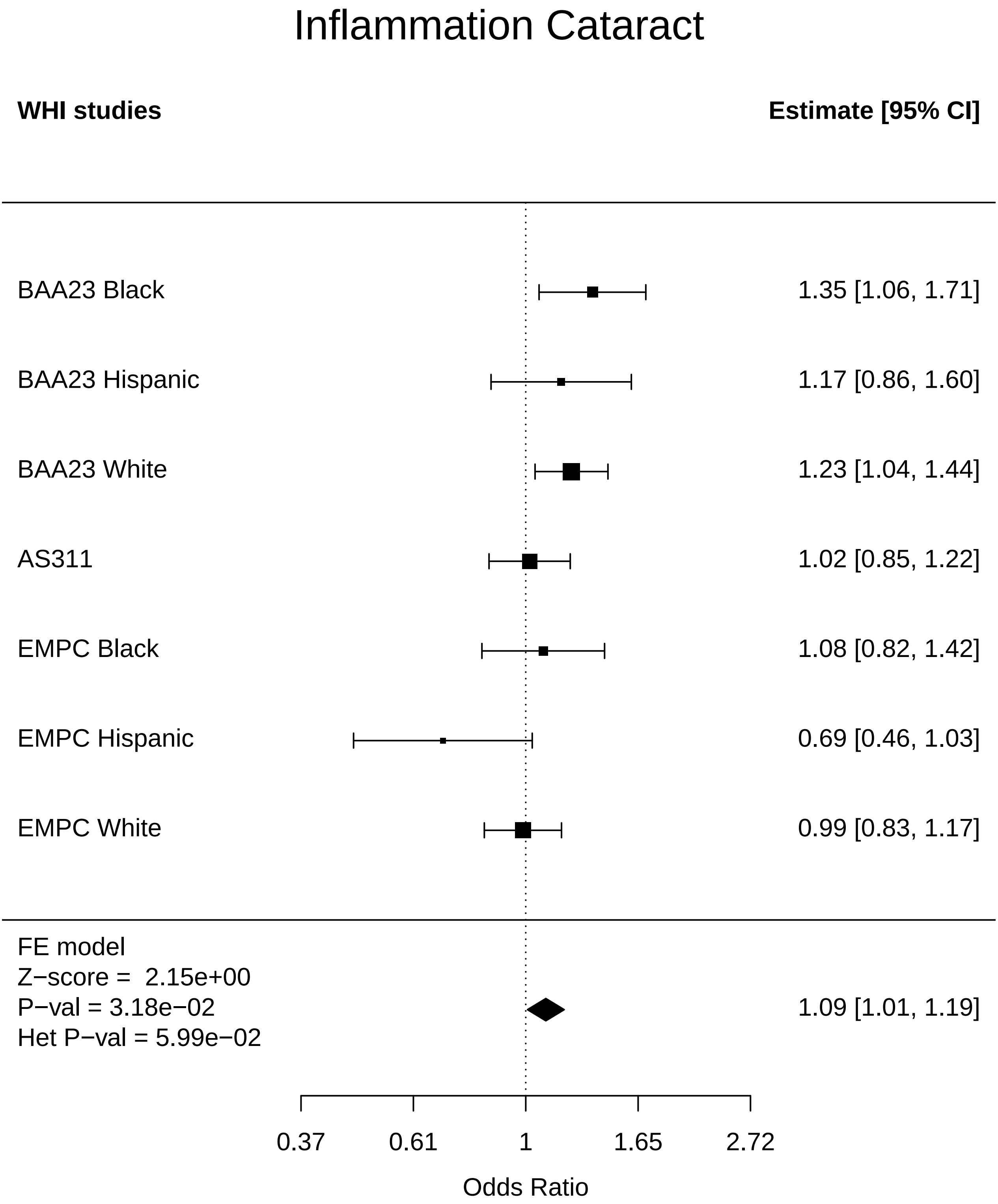

### Inflammation_Colorectal cancer.pdf

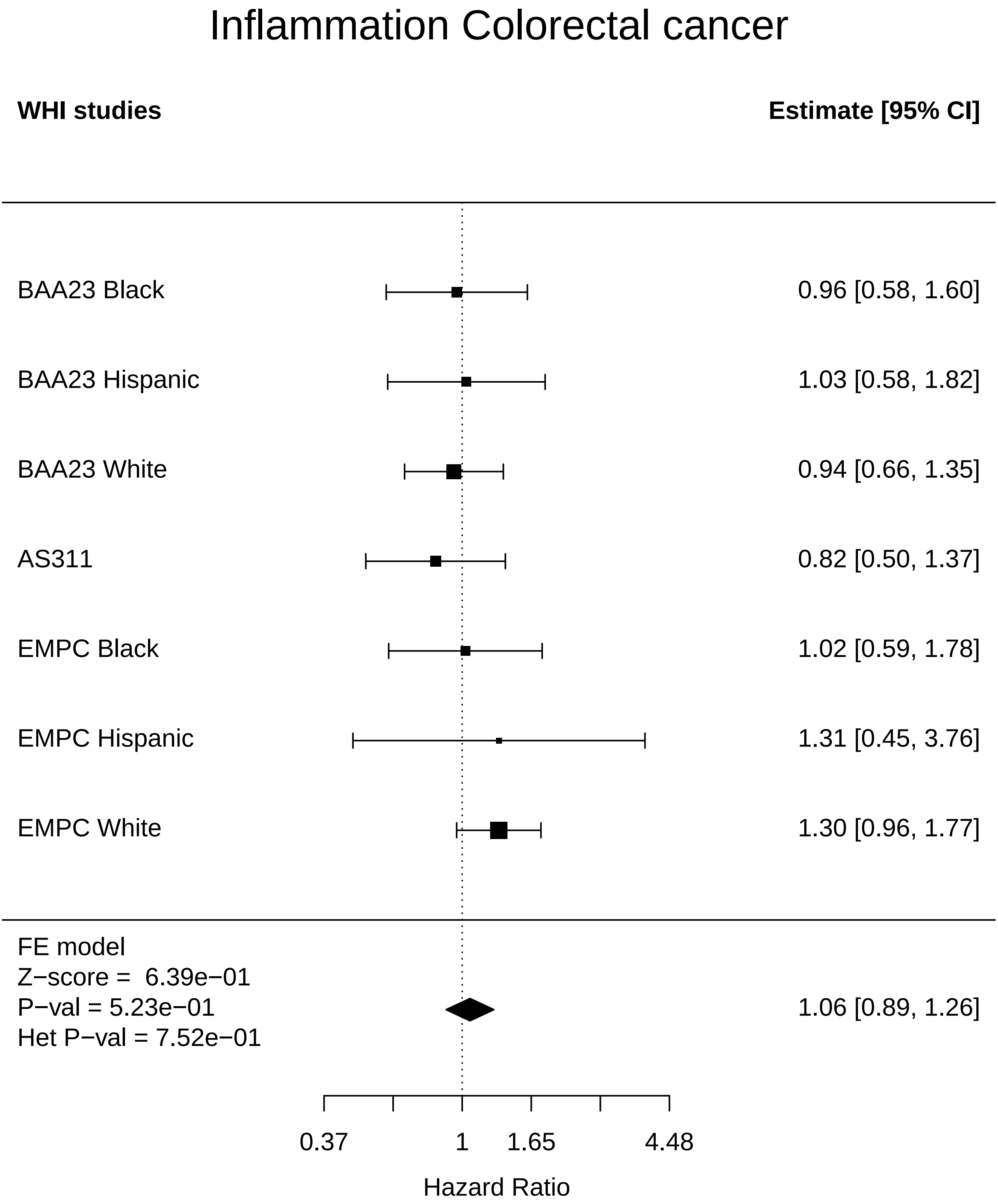

### Inflammation_Diabetes.pdf

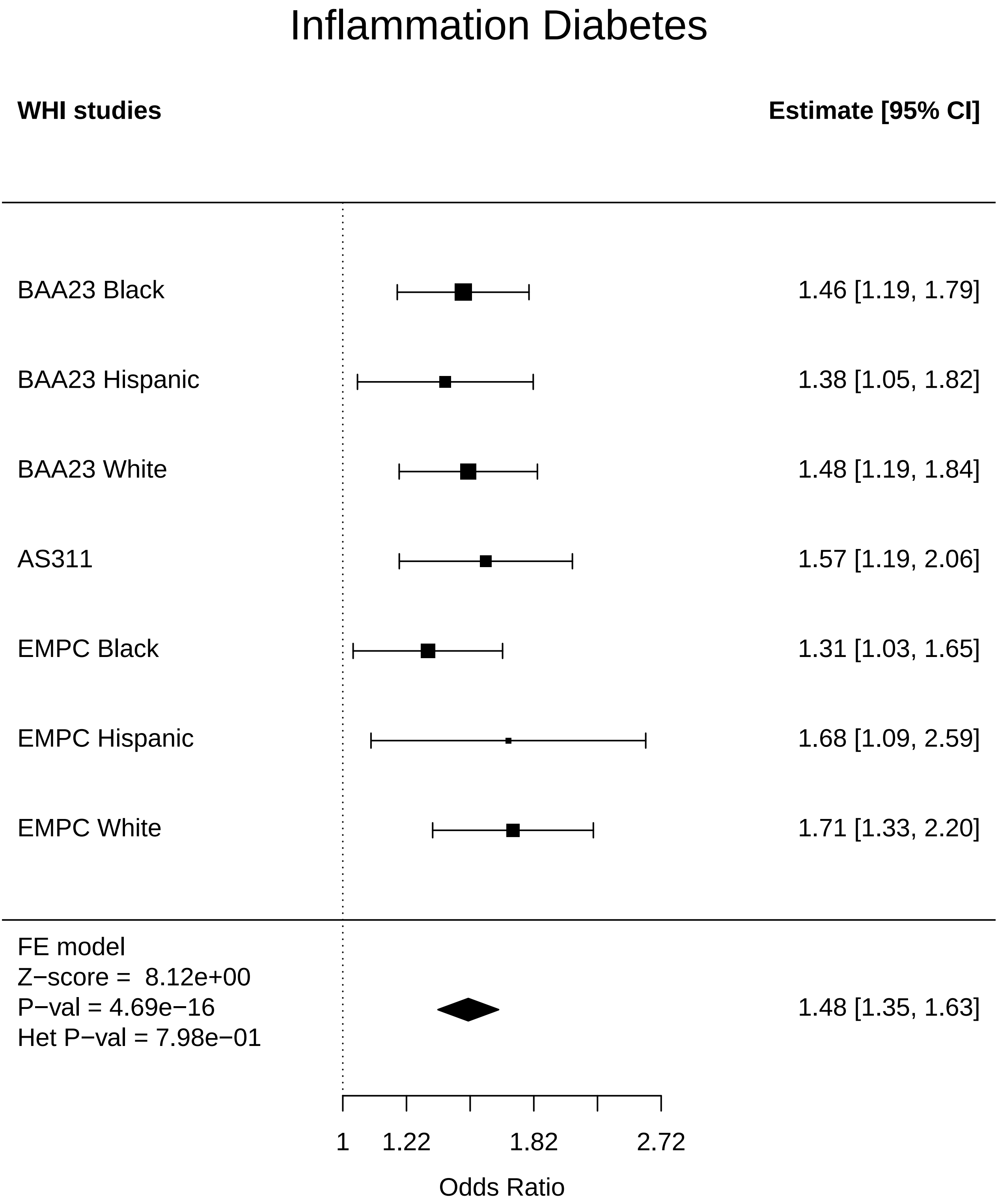

### Inflammation_Disease free.pdf

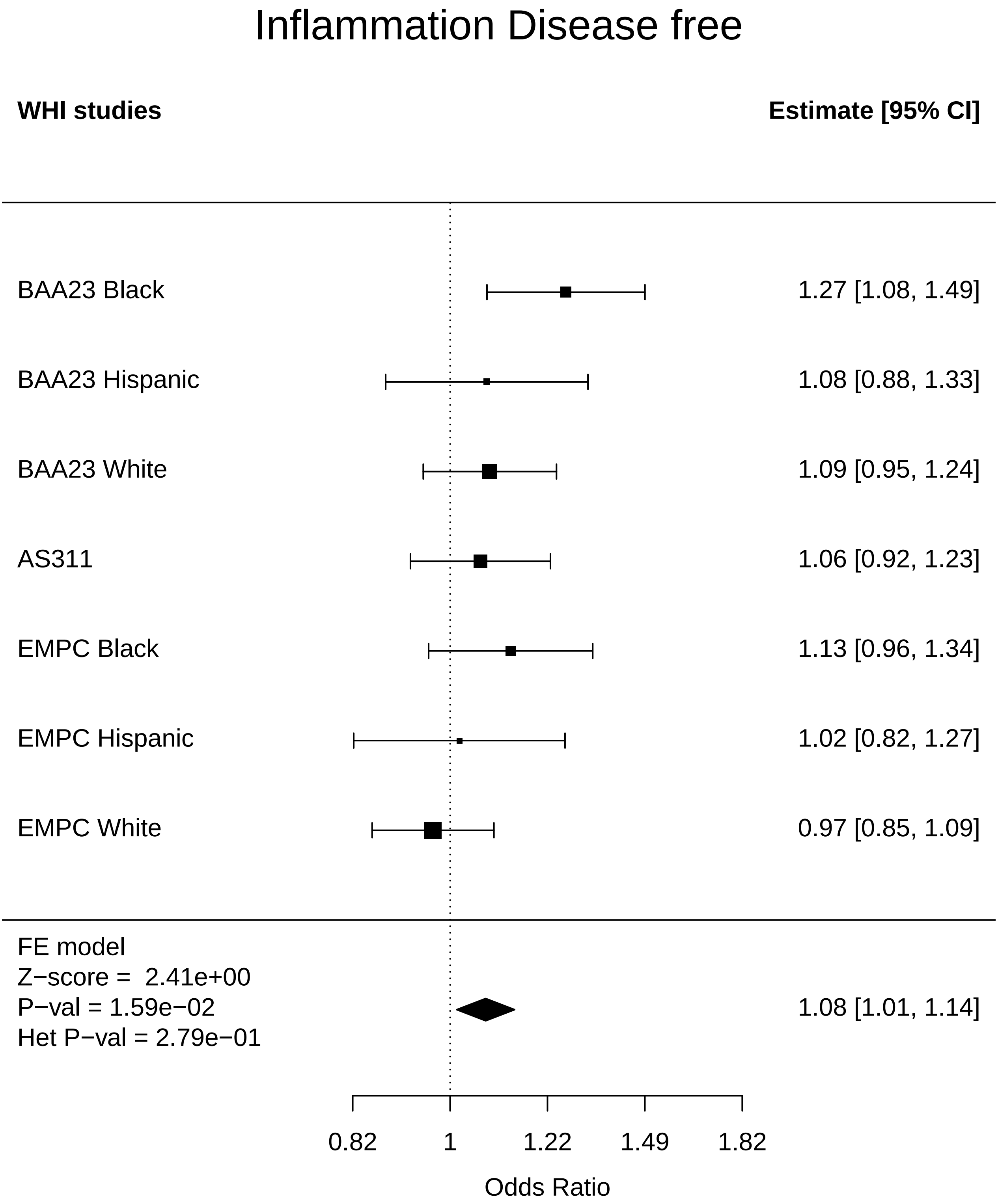

### Inflammation_Endometrial cancer.pdf

#### Inflammation Endometrial cancer

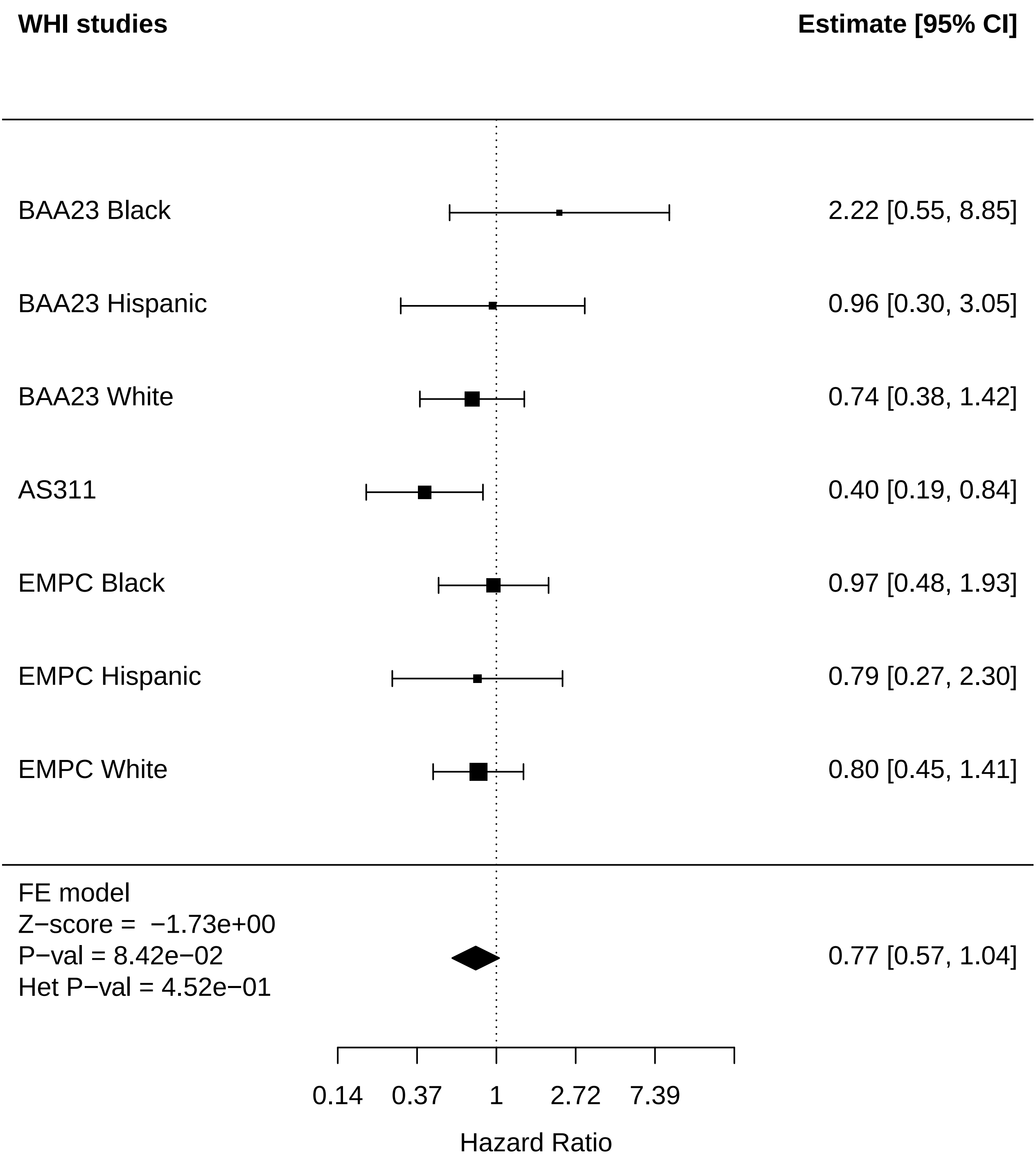

### Inflammation_Leukemia.pdf

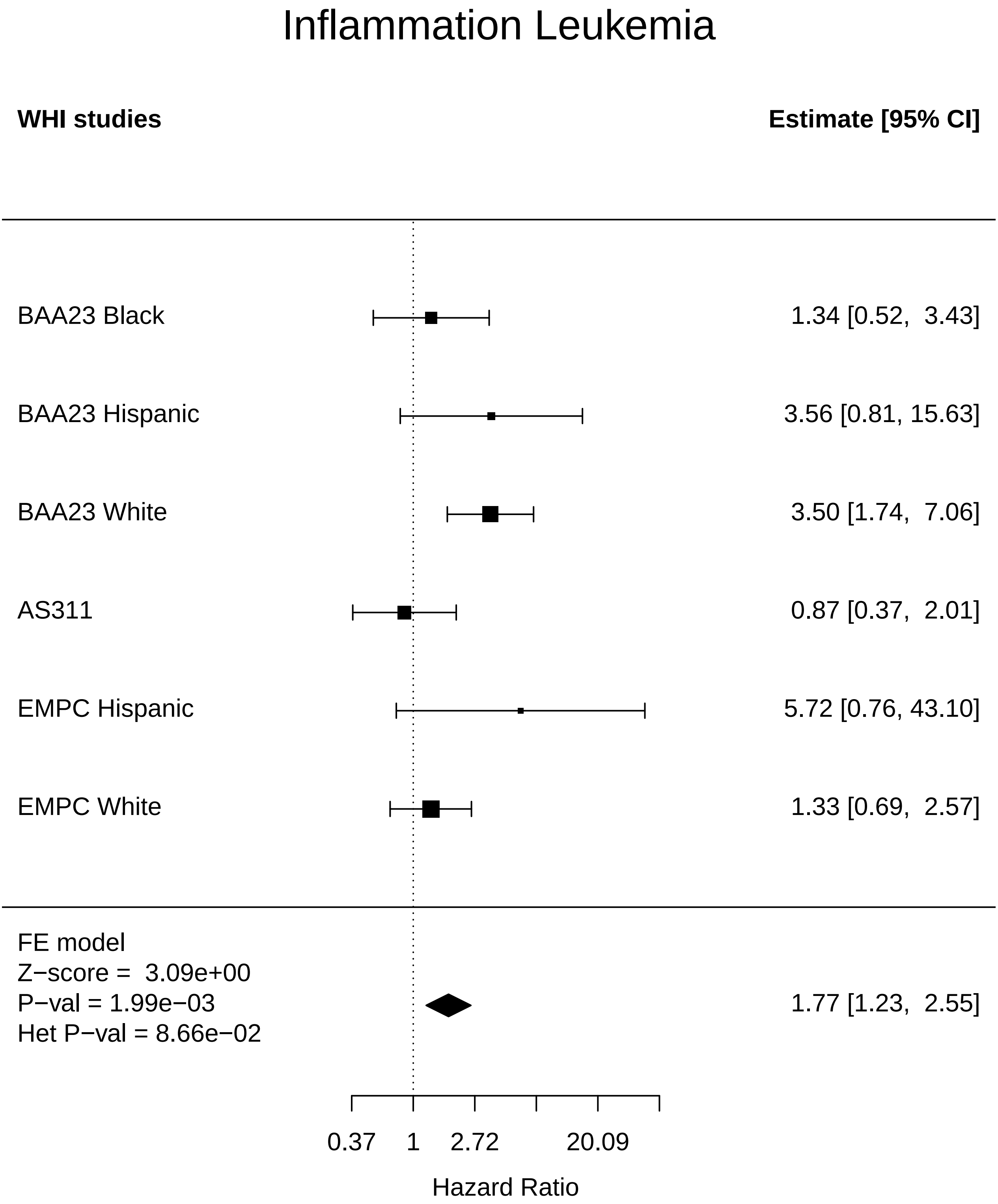

### Inflammation_Lung Cancer.pdf

#### Inflammation Lung Cancer

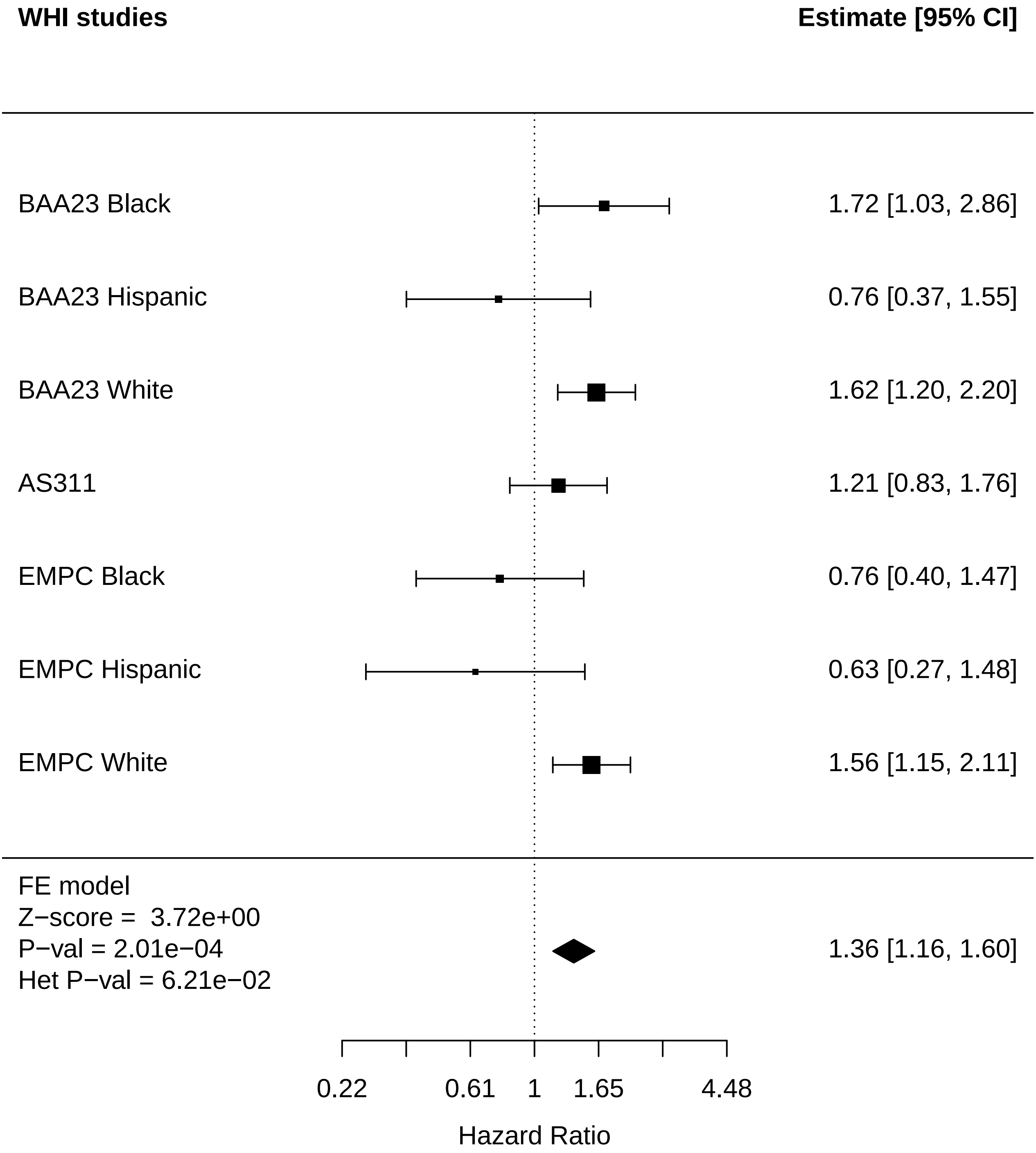

### Inflammation_MI.pdf

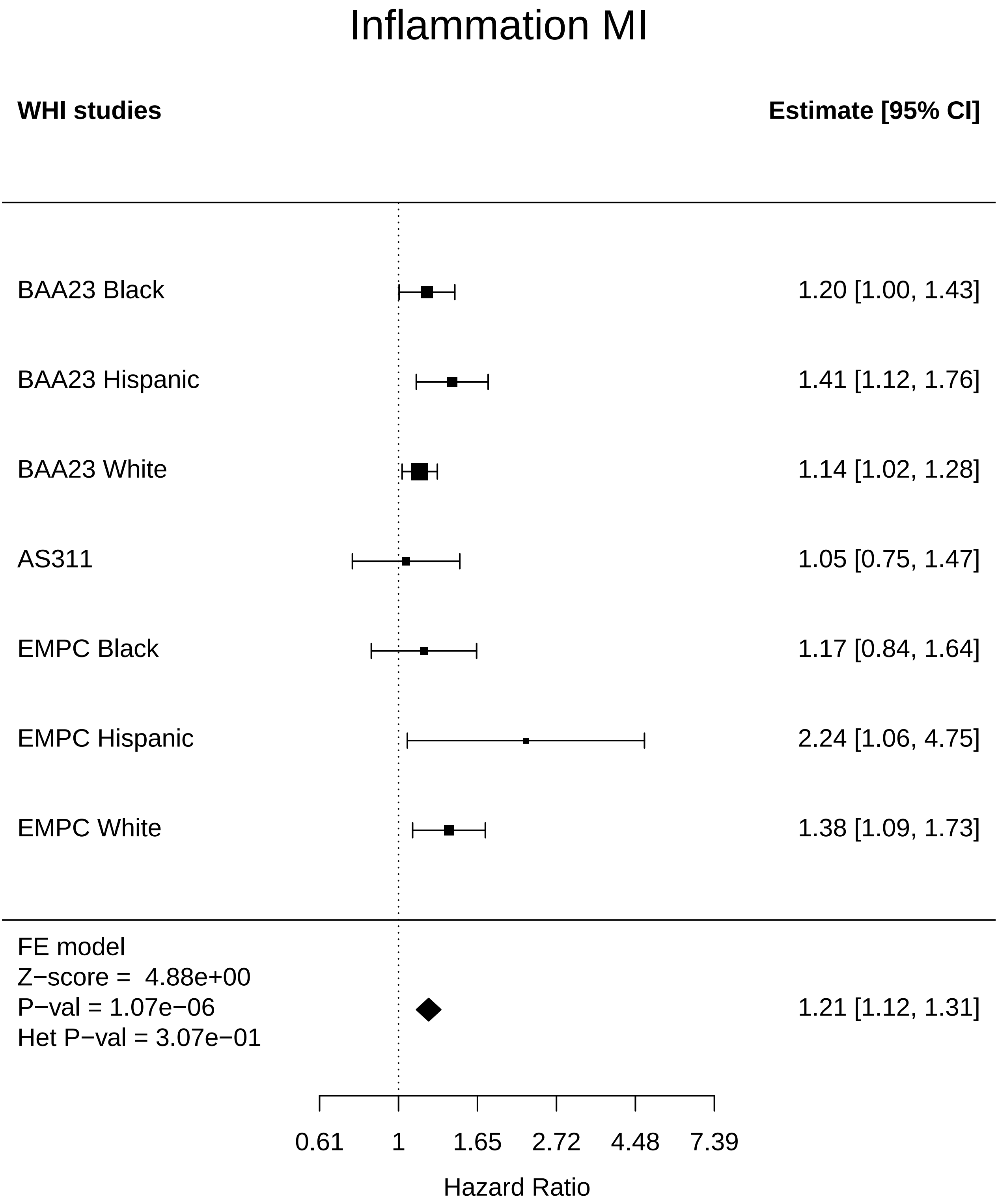

### Inflammation_Mortality.pdf

#### Inflammation Mortality

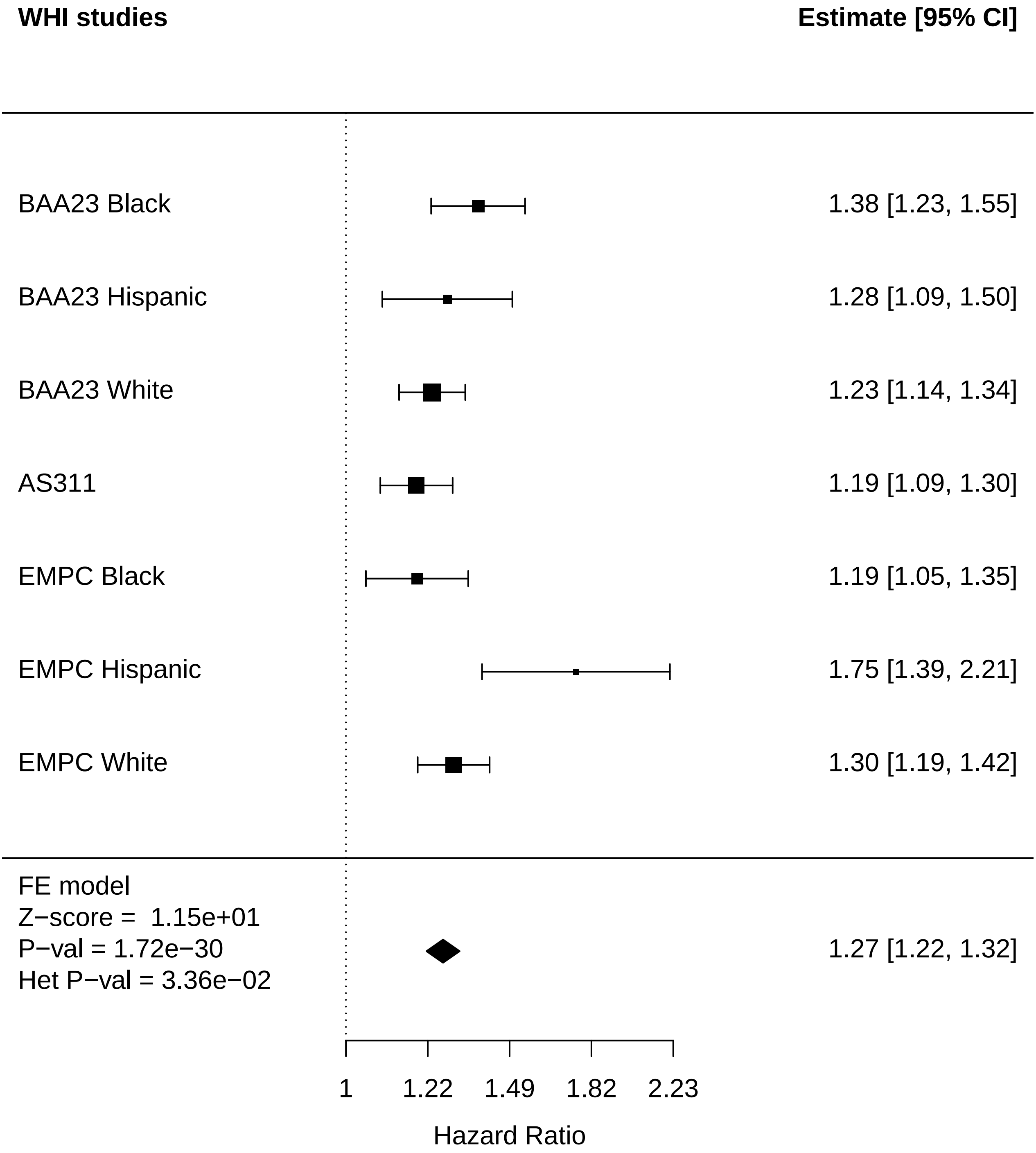

### Inflammation_Stroke.pdf

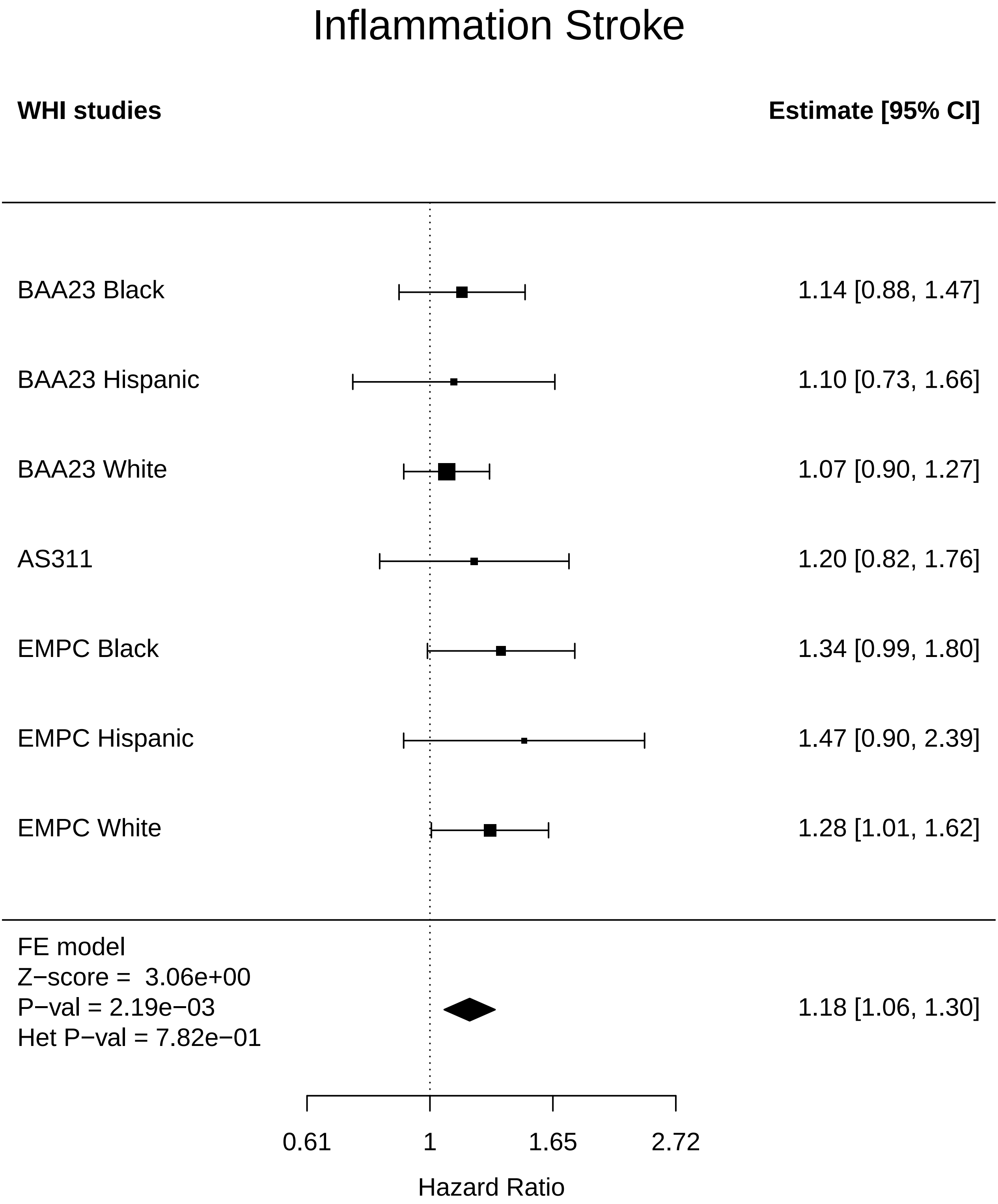

### Inflammation_Thyroid disease.pdf

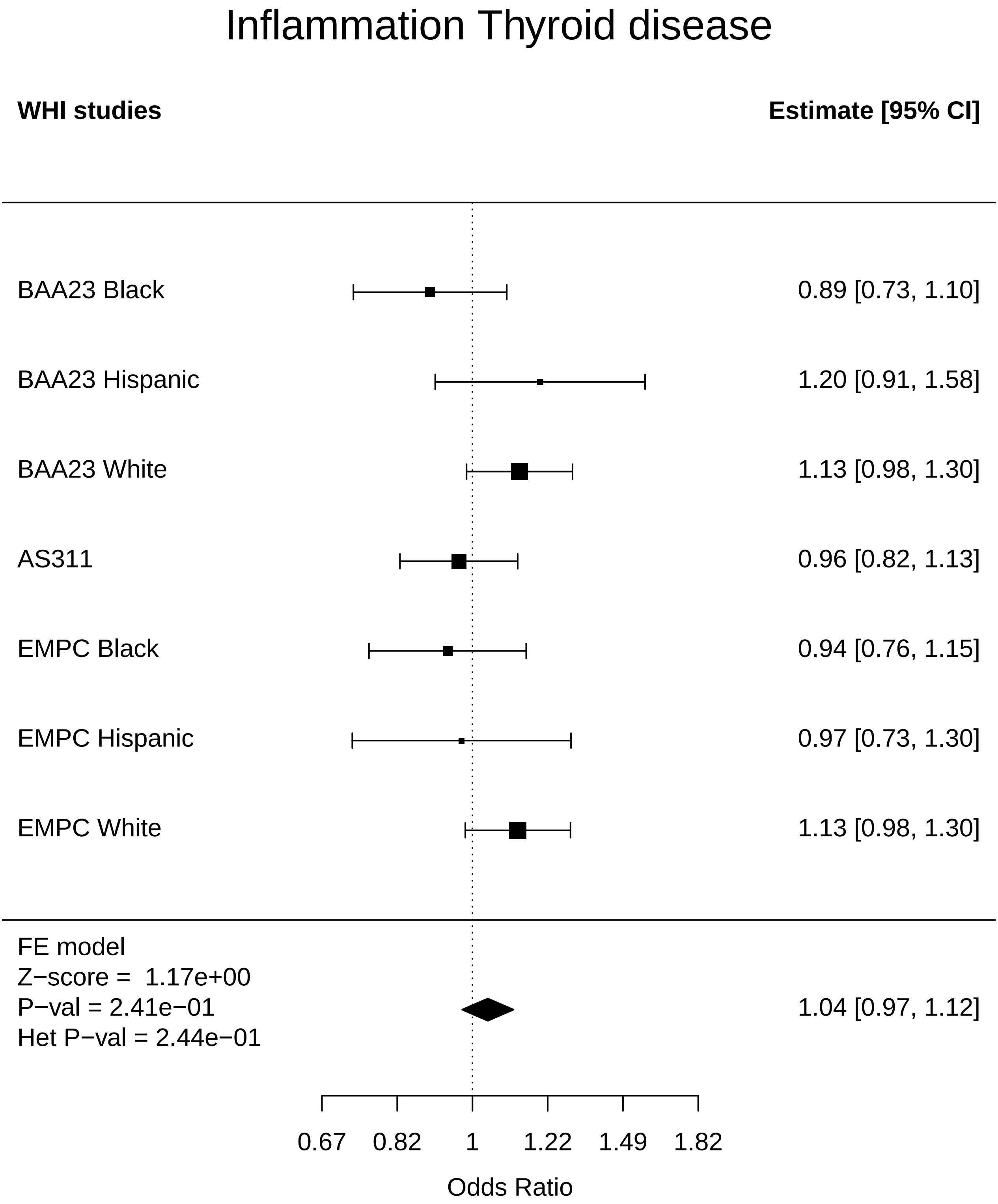

### Inflammation_Total comorbidities.pdf

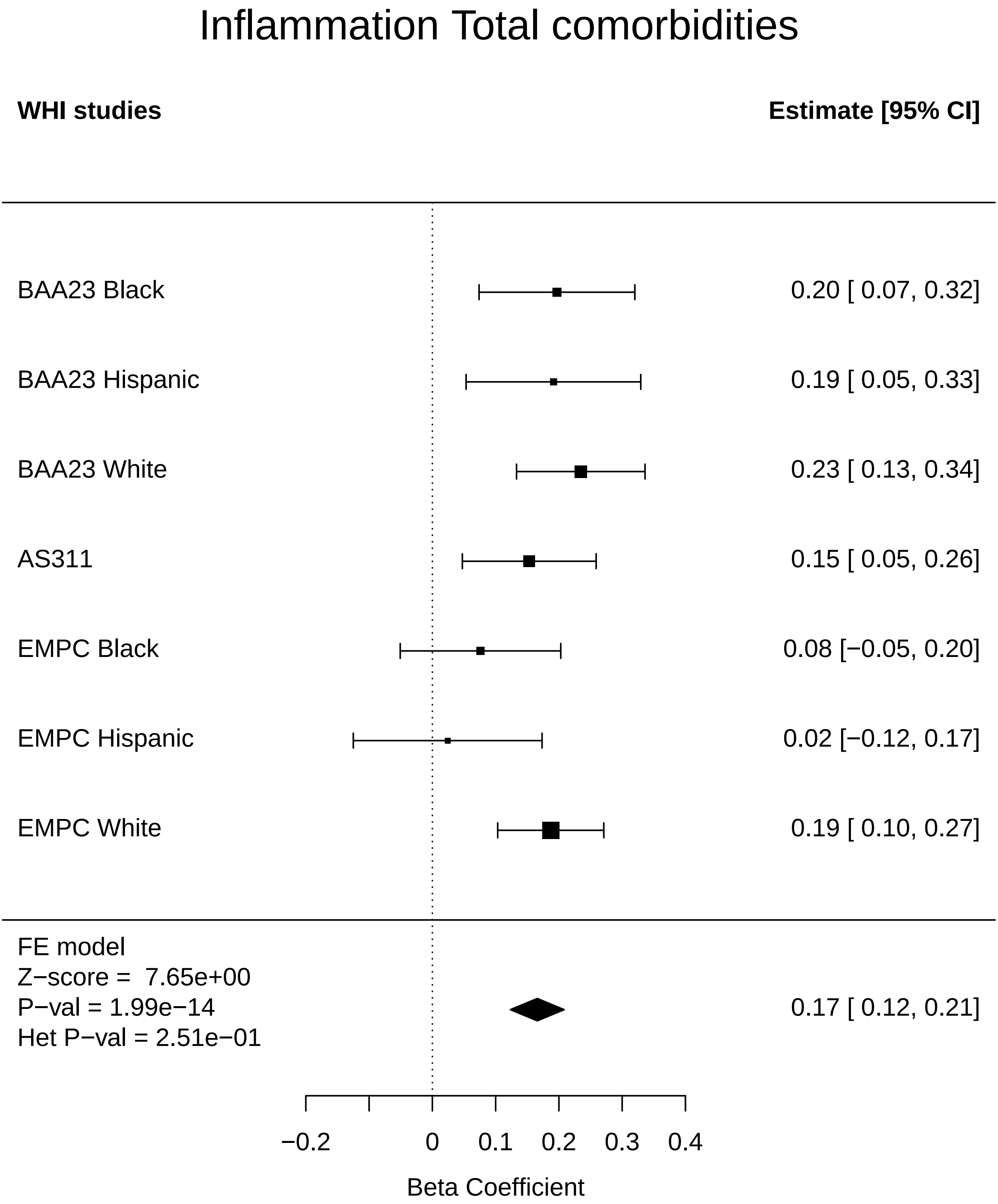

### Kidney_Arthritis.pdf

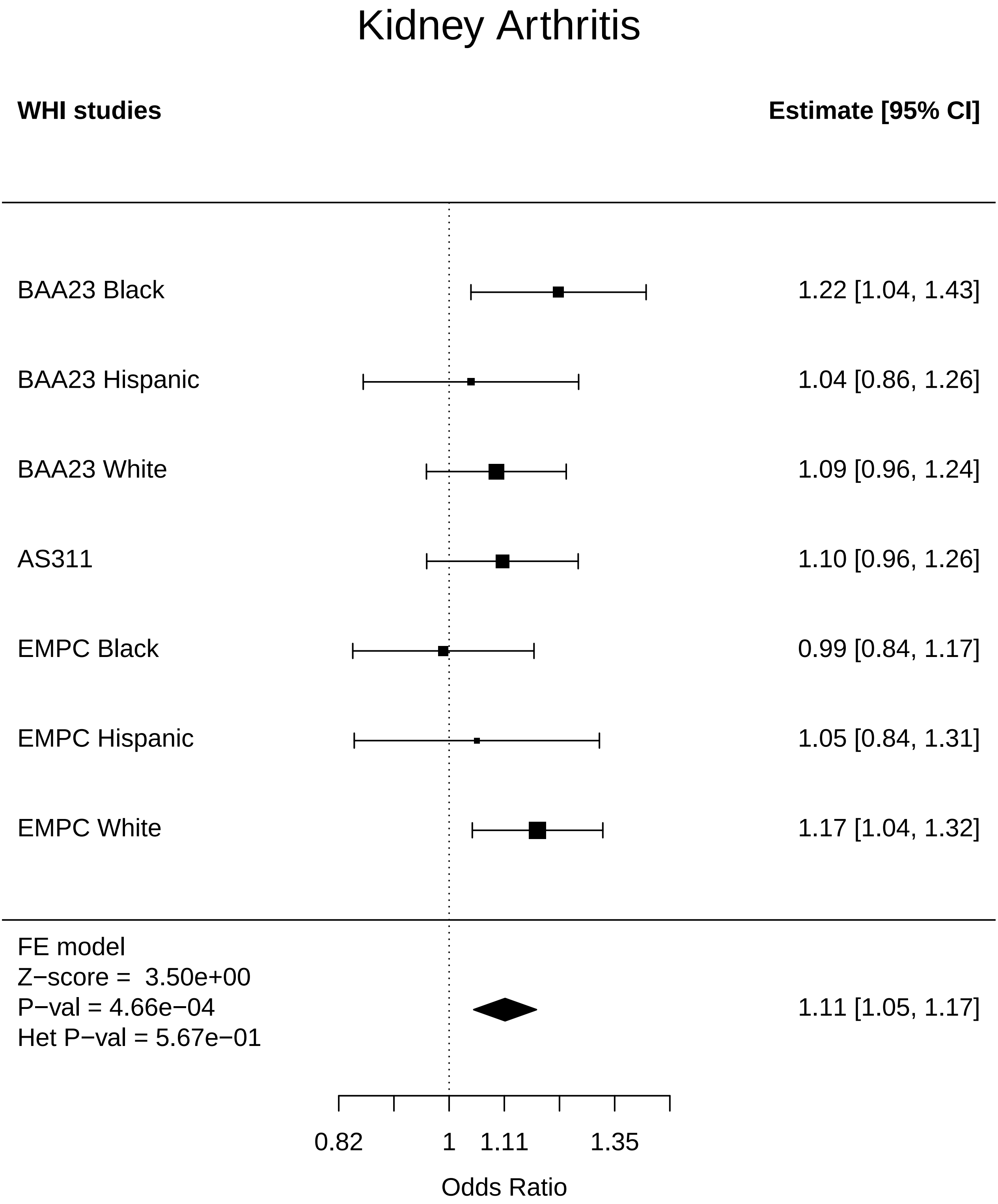

### Kidney_CHD.pdf

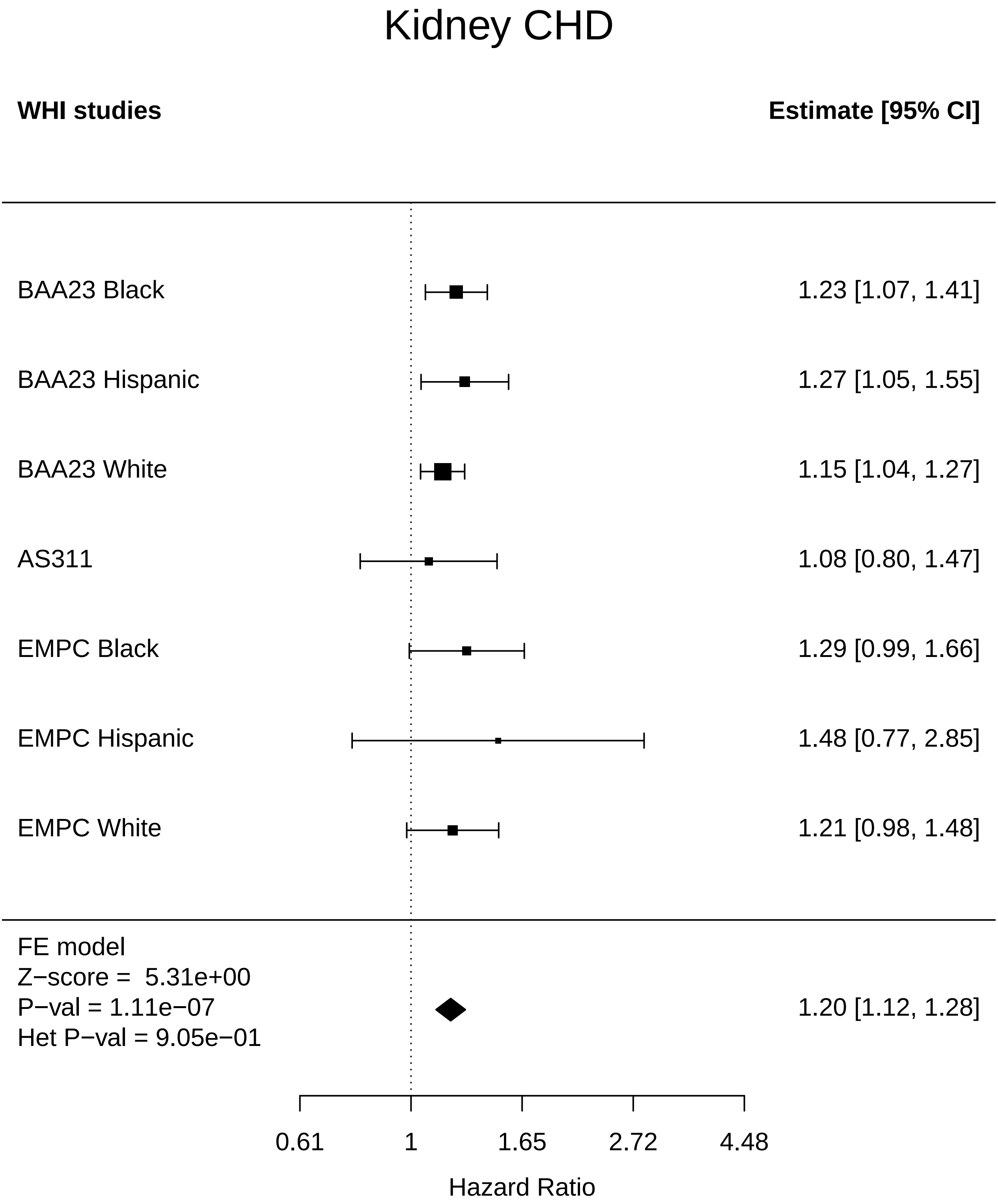

### Kidney_Diabetes.pdf

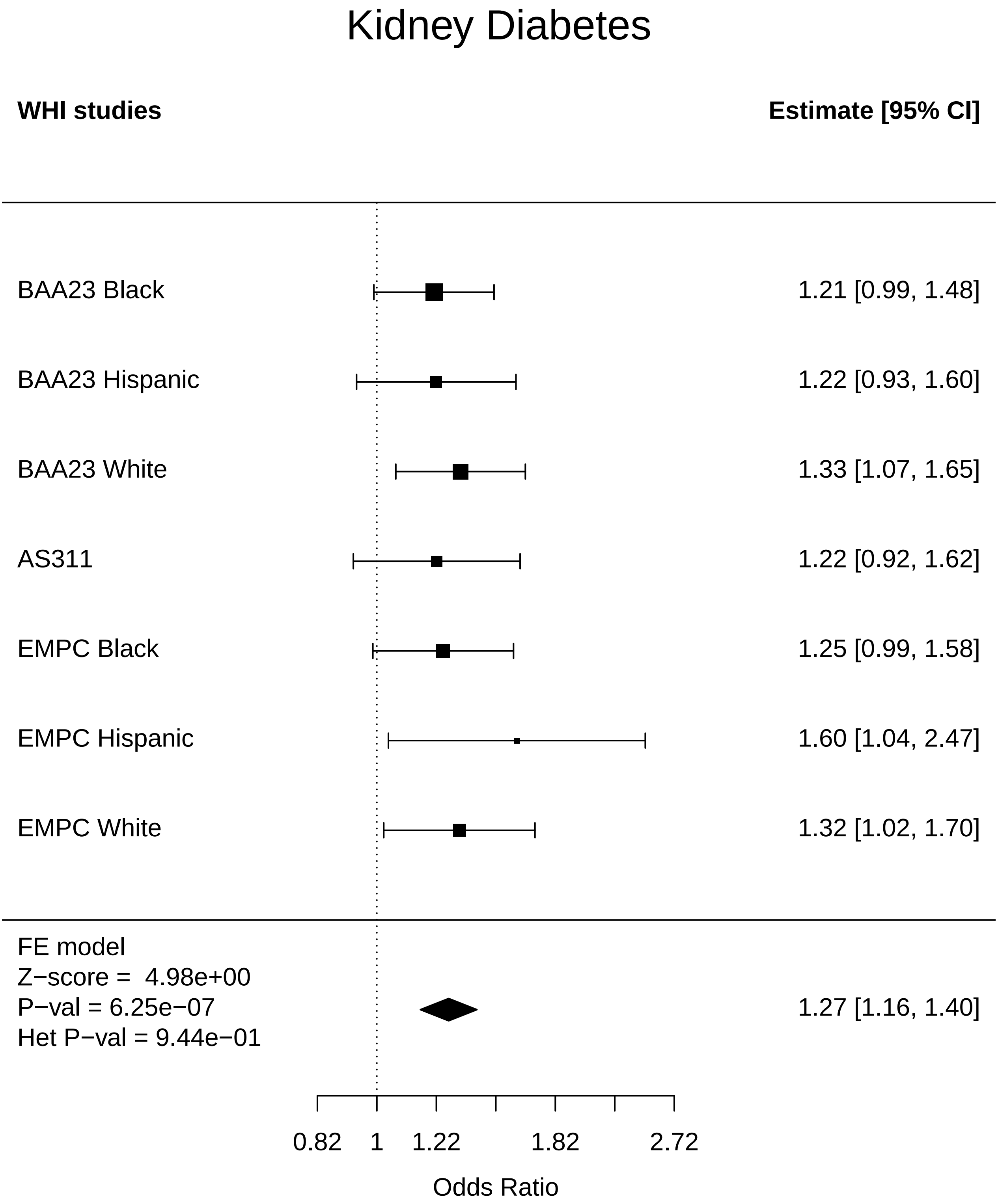

### Kidney_Disease free.pdf

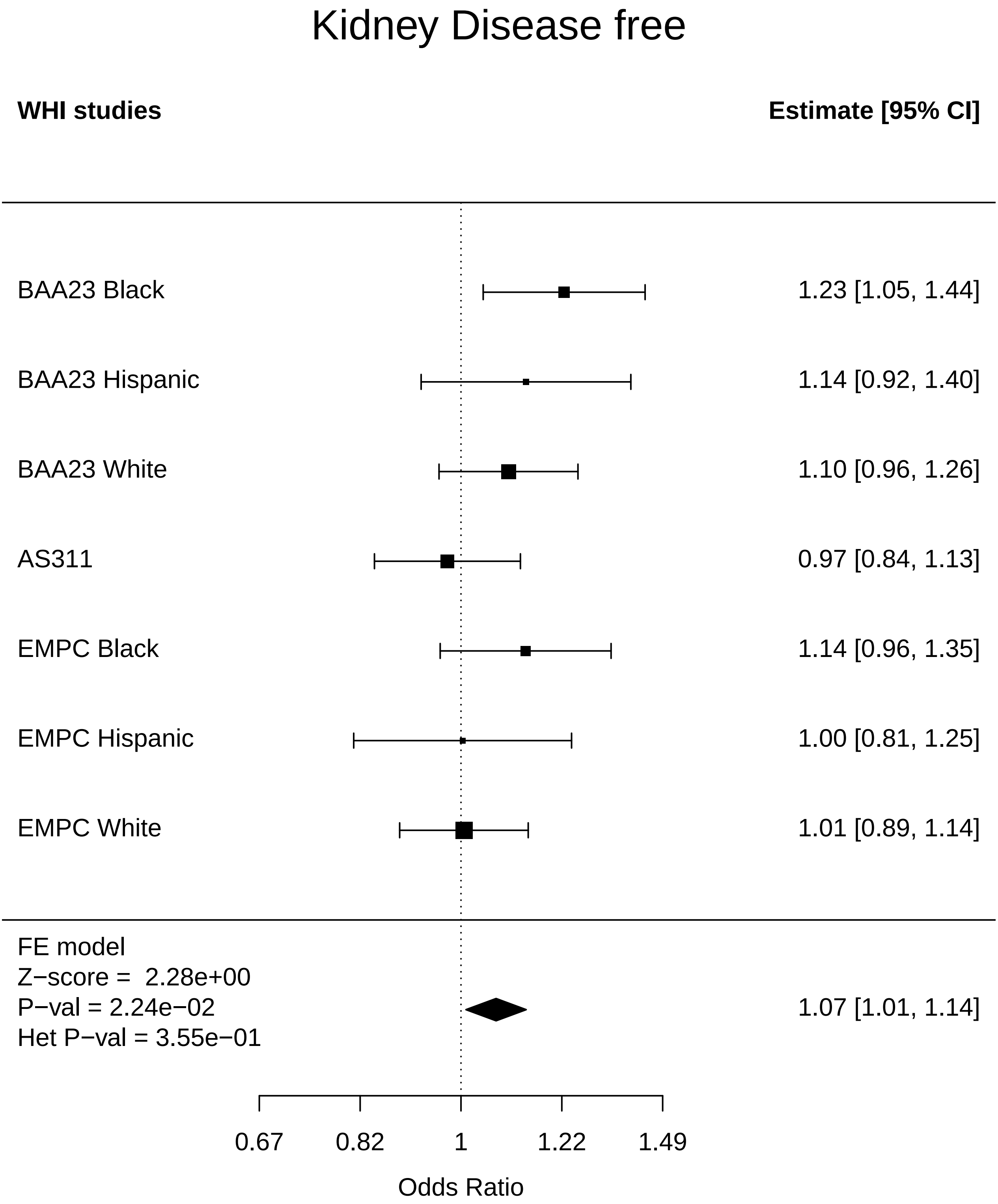

### Kidney_Endometrial cancer.pdf

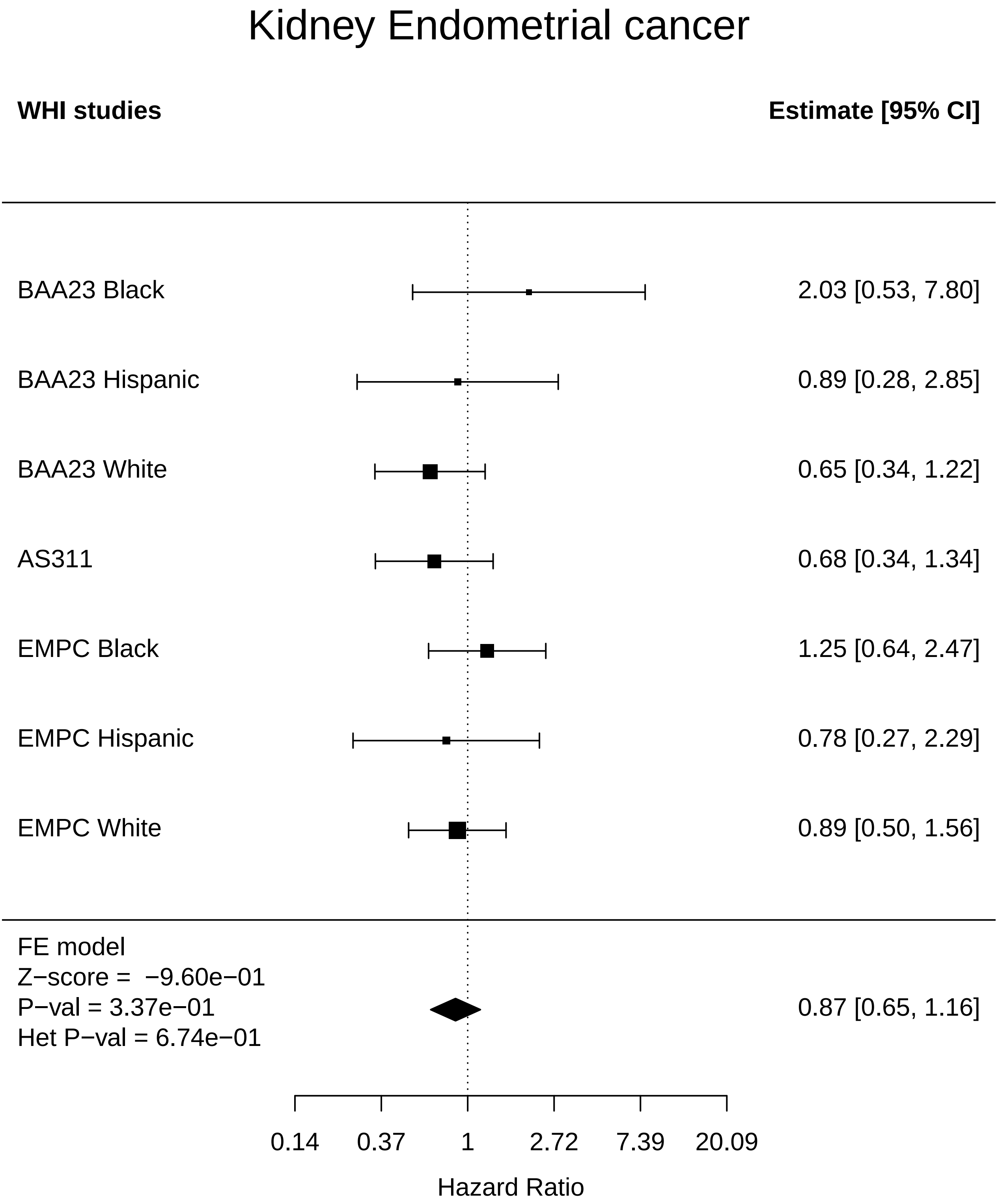

### Kidney_Lung Cancer.pdf

#### Kidney Lung Cancer

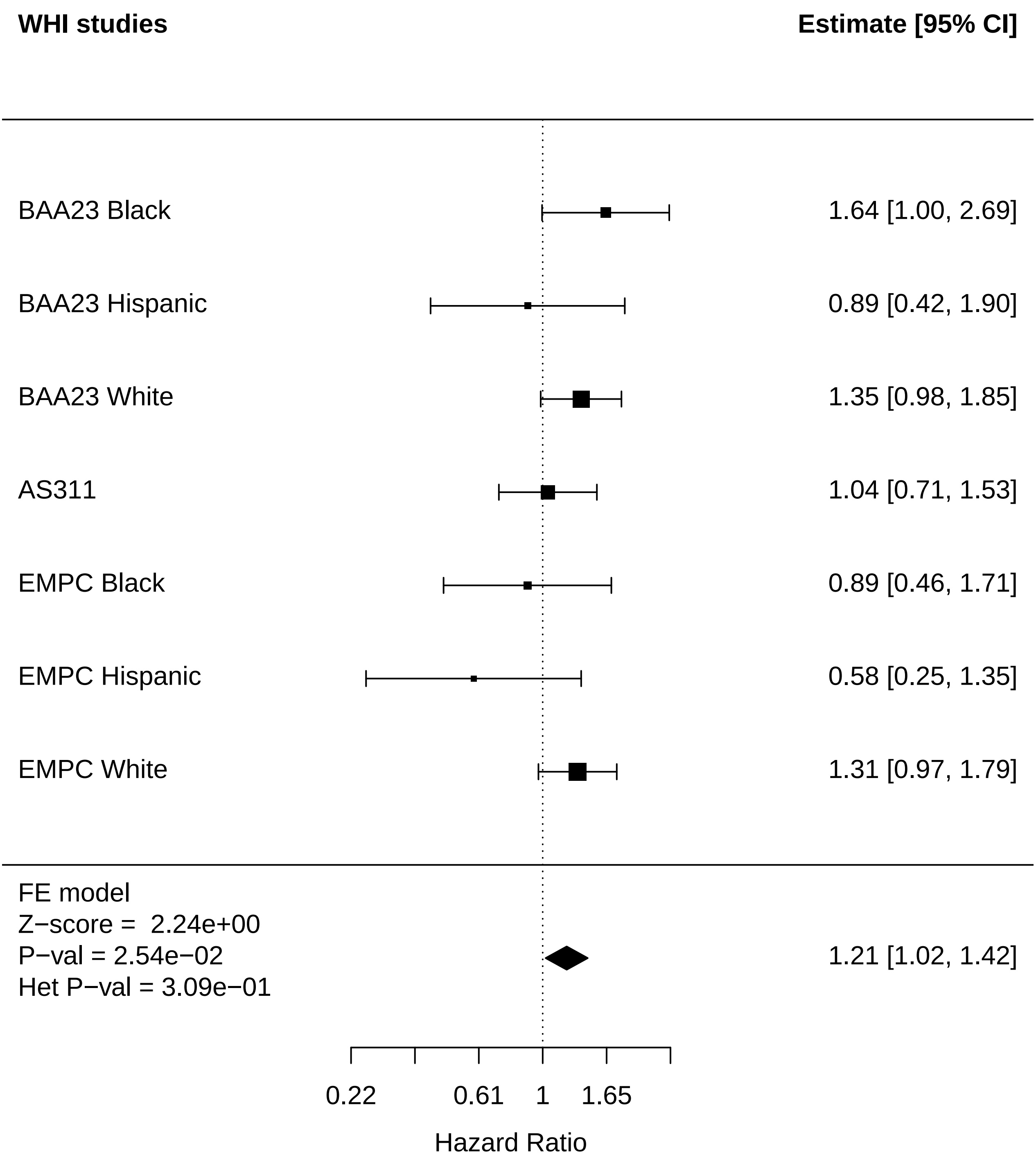

### Kidney_MI.pdf

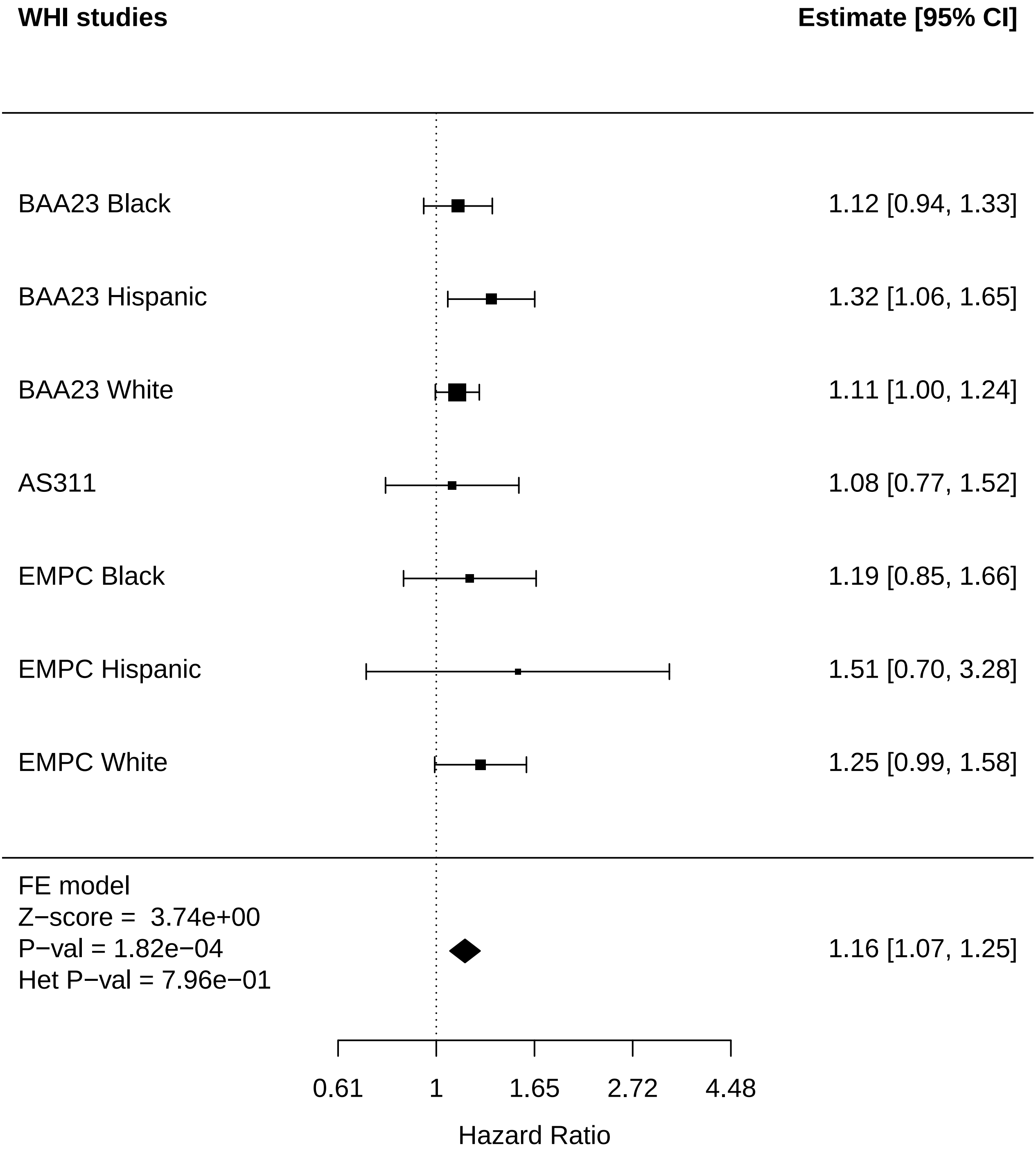

### Kidney_Physical function.pdf

#### Kidney Physical function

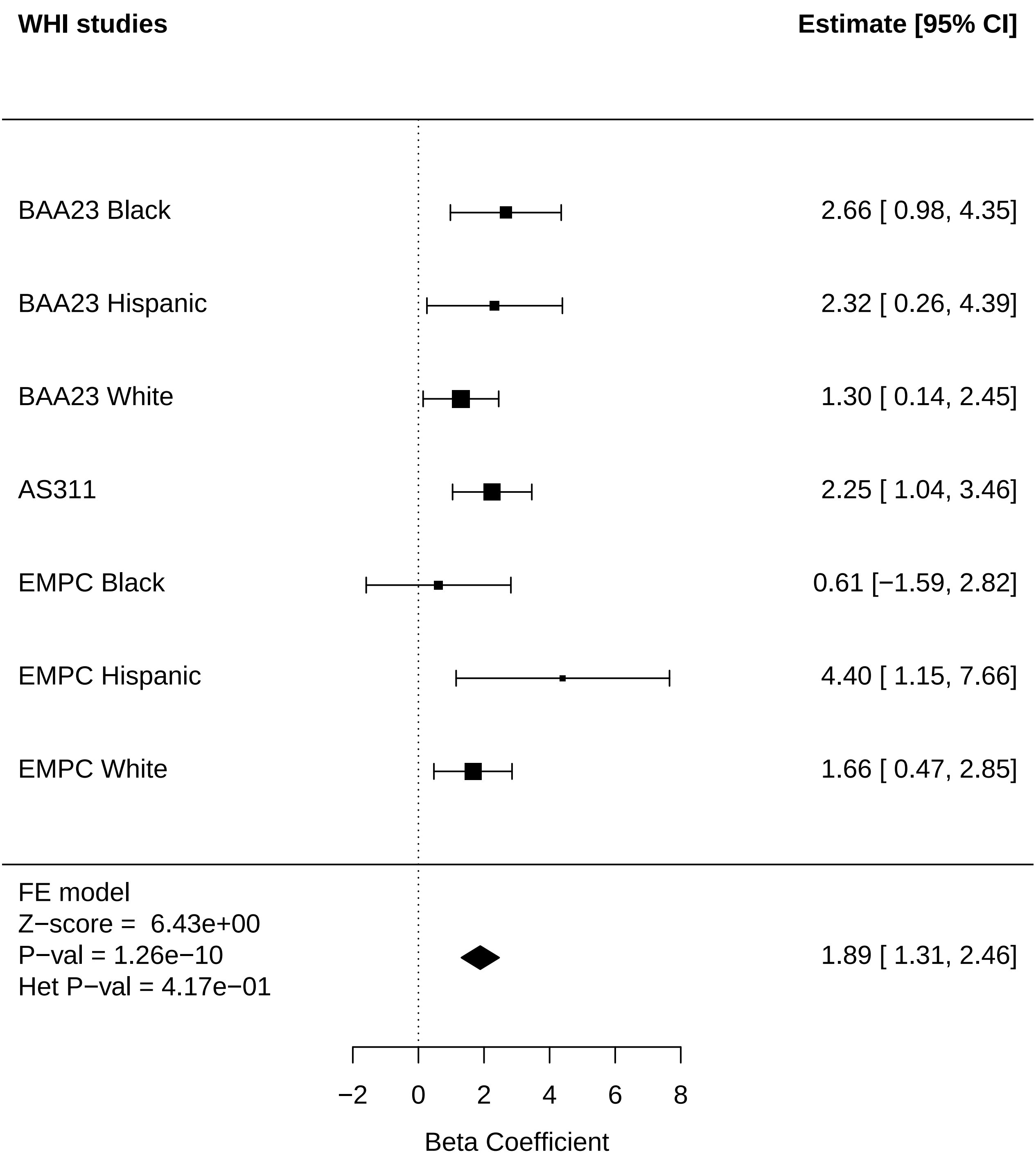

### Kidney_Stroke.pdf

#### Kidney Stroke

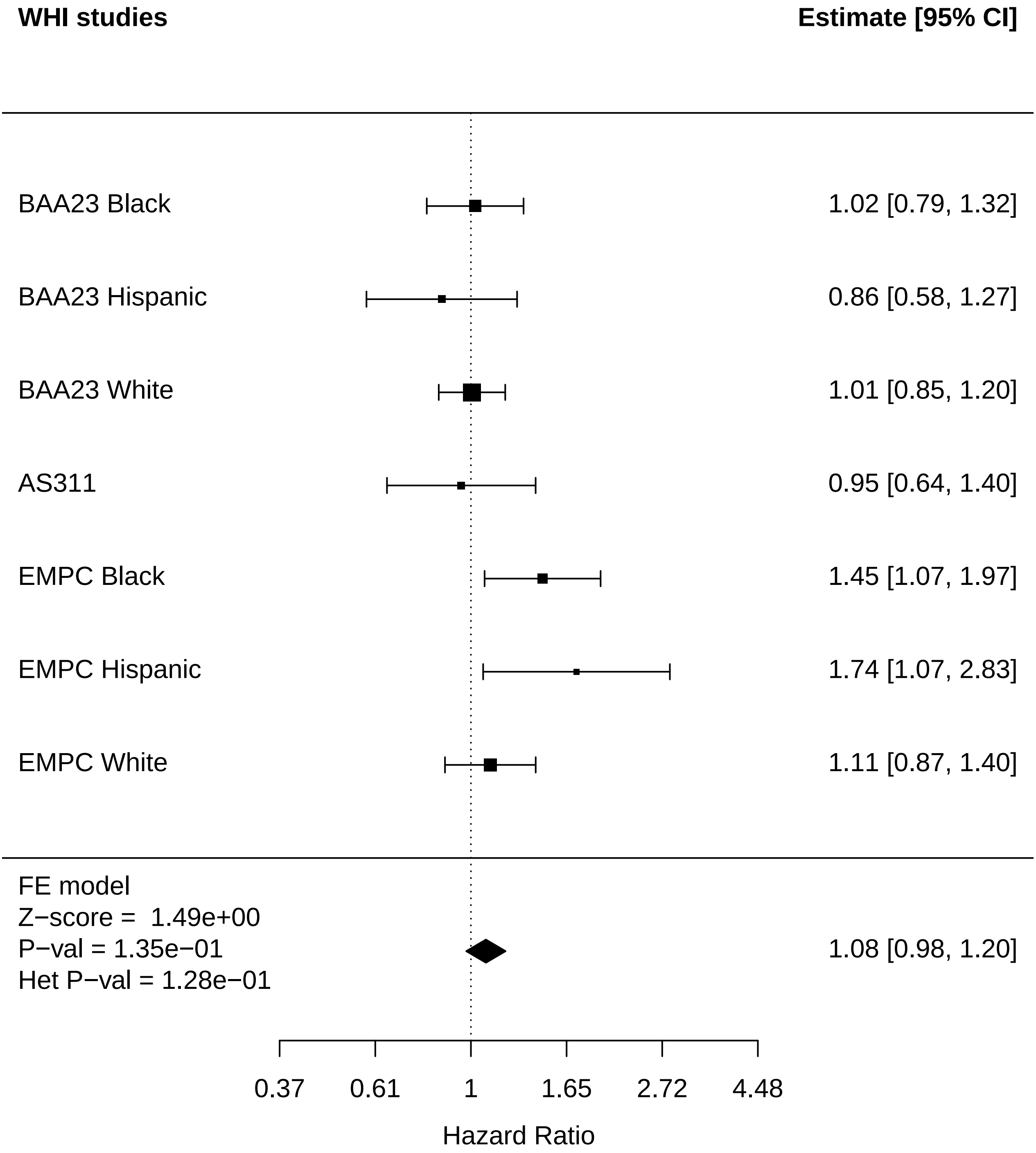

### Kidney_Thyroid disease.pdf

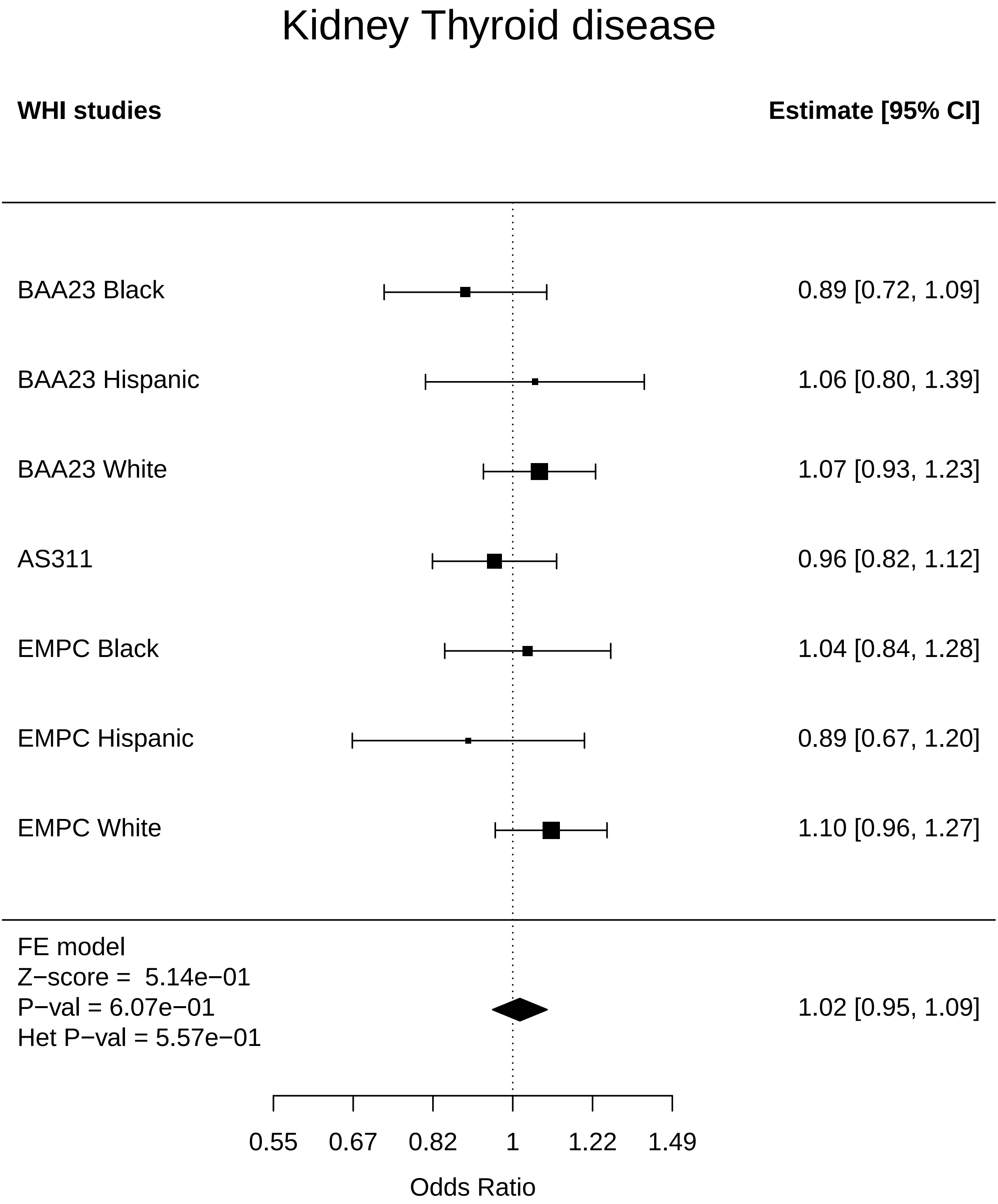

### Kidney_Total comorbidities.pdf

#### Kidney Total comorbidities

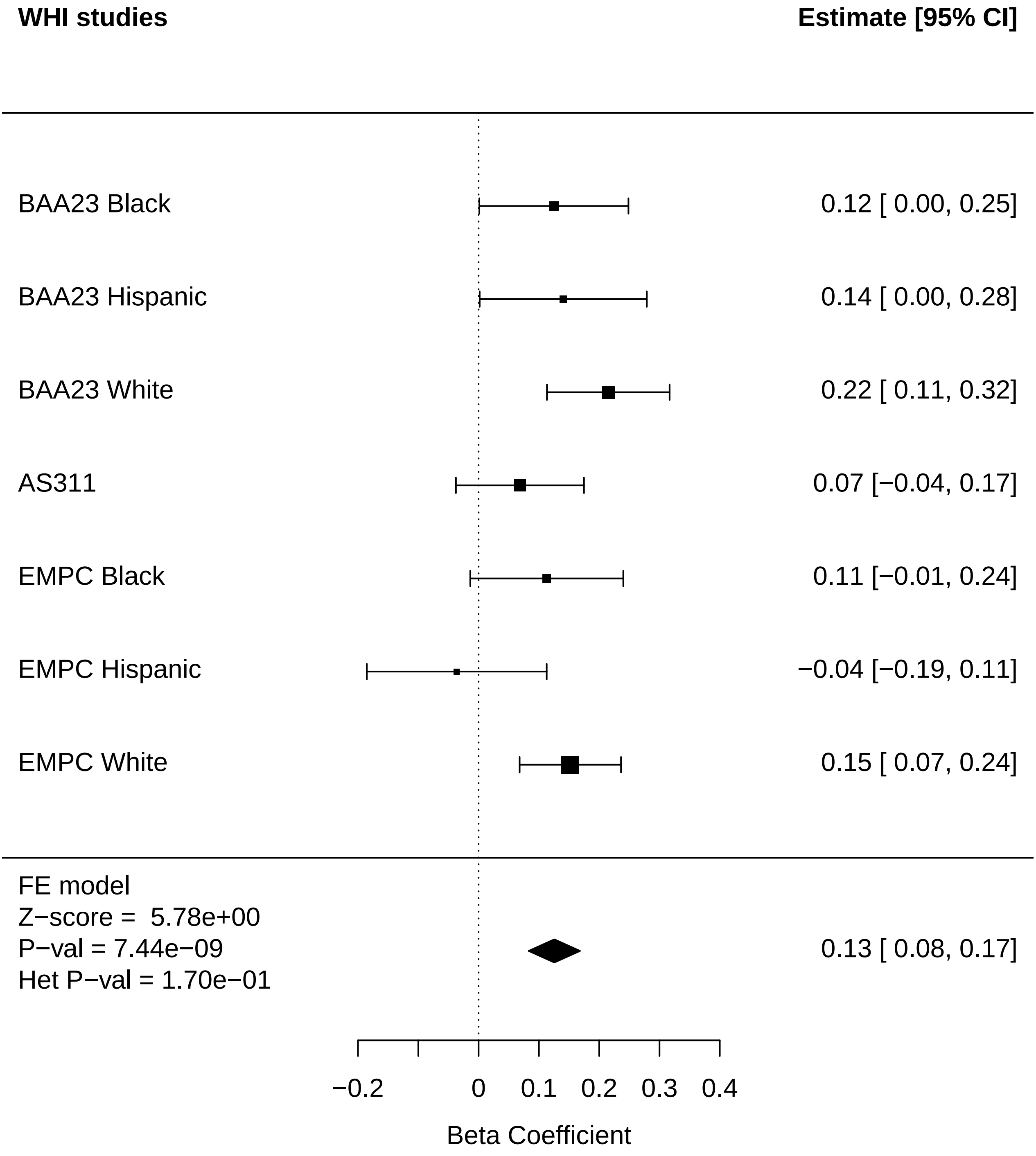

### Liver_Arthritis.pdf

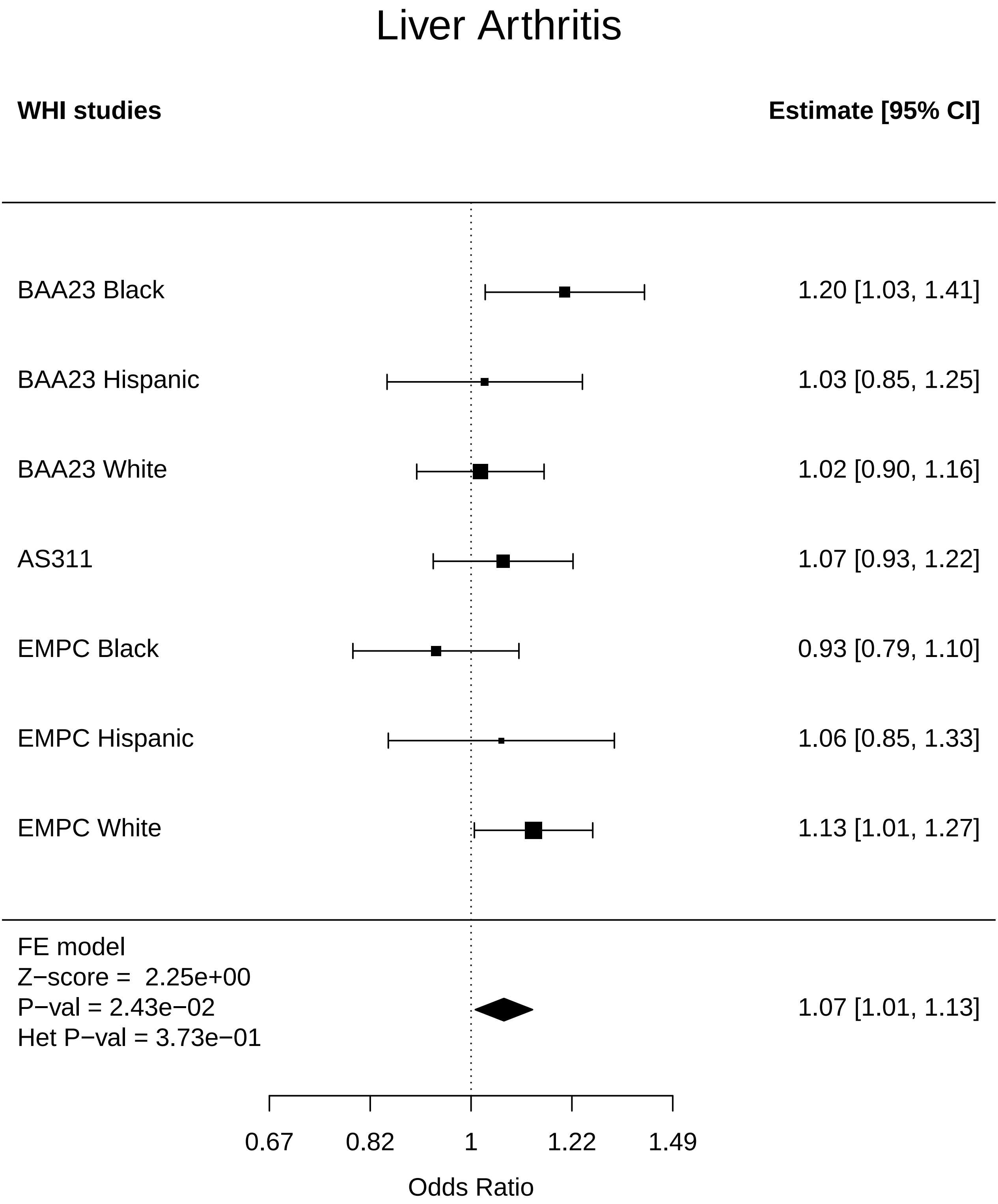

### Liver_Breast Cancer.pdf

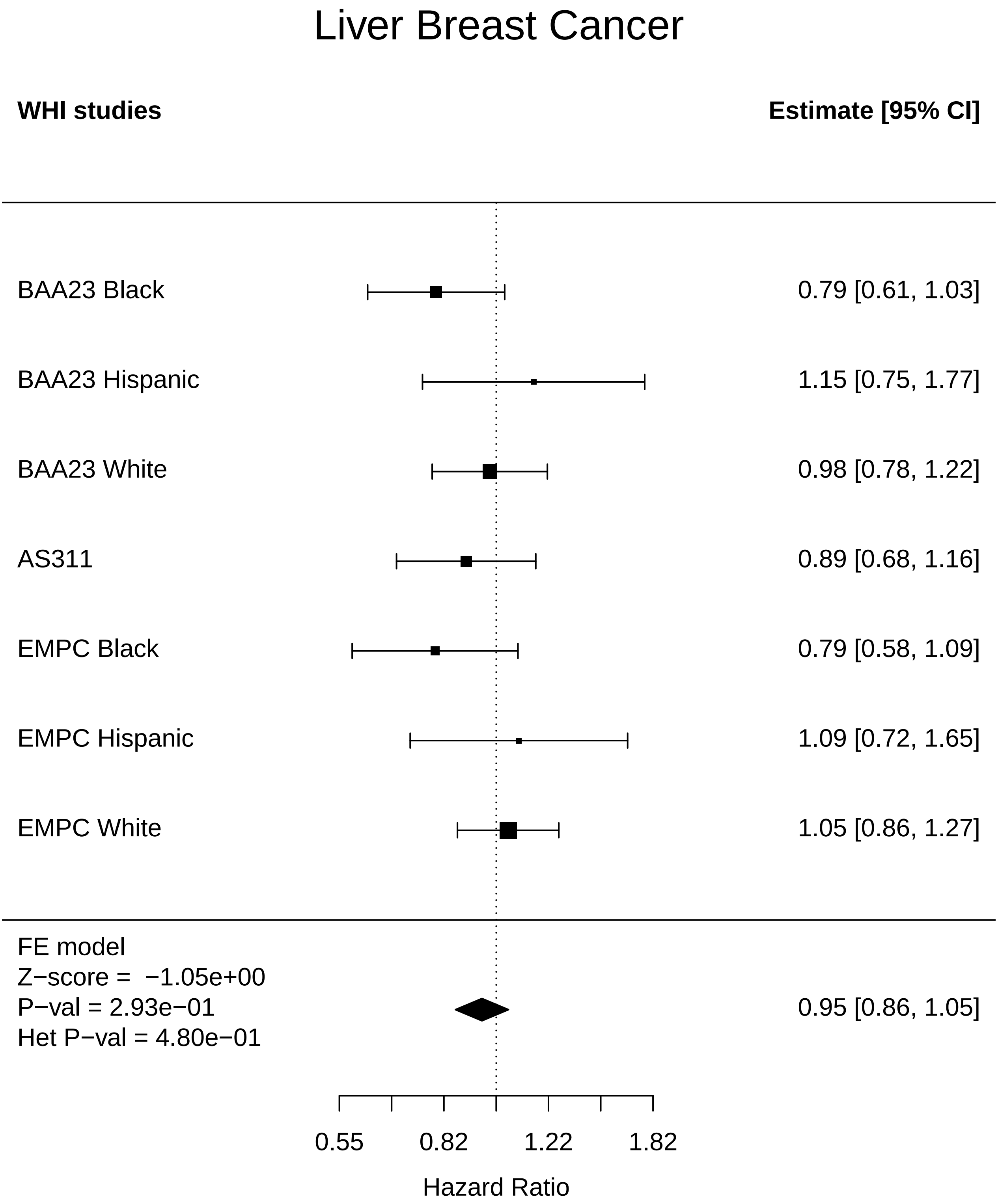

### Liver_Diabetes.pdf

#### Liver Diabetes

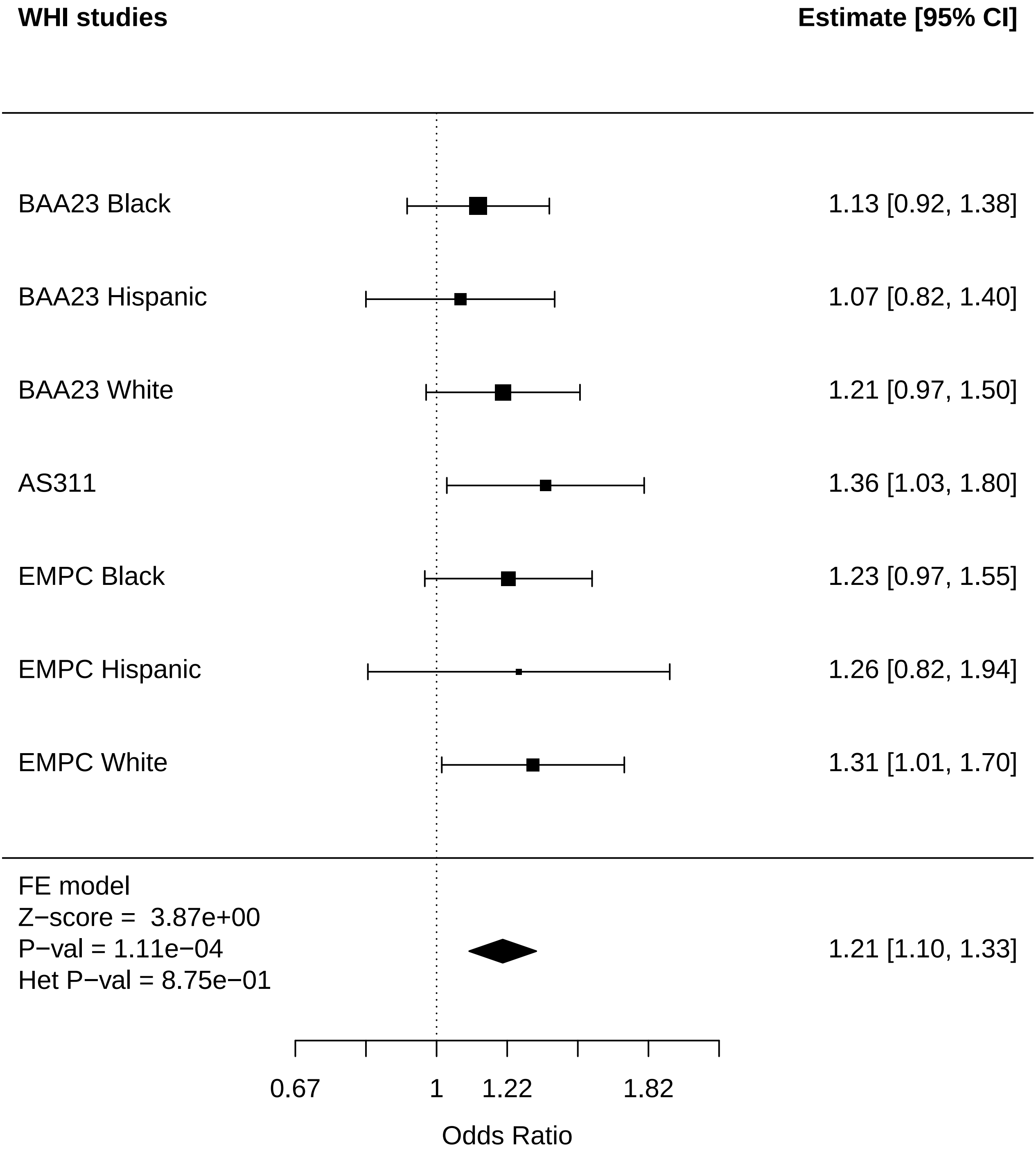

### Liver_Disease free.pdf

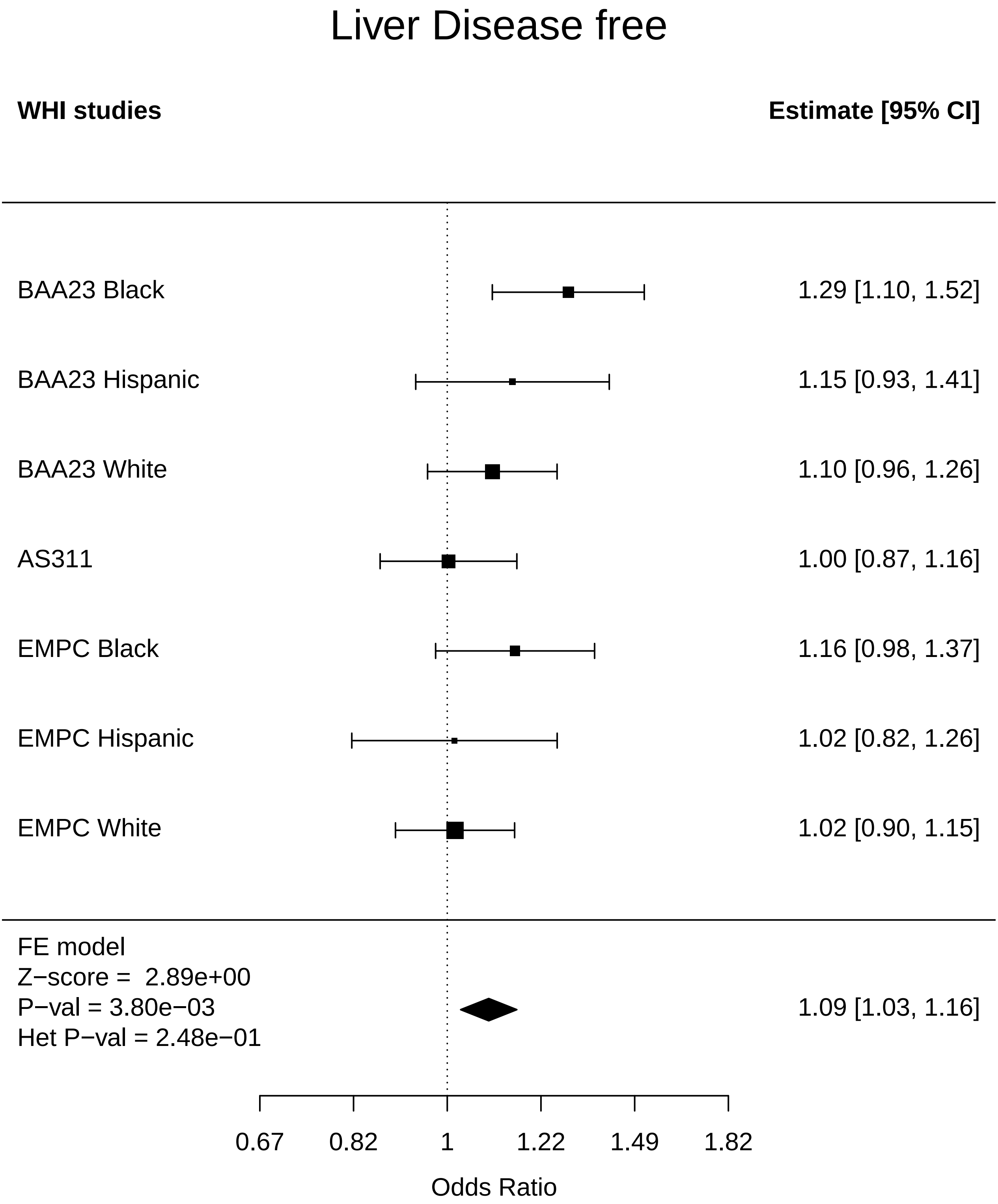

### Liver_Endometrial cancer.pdf

#### Liver Endometrial cancer

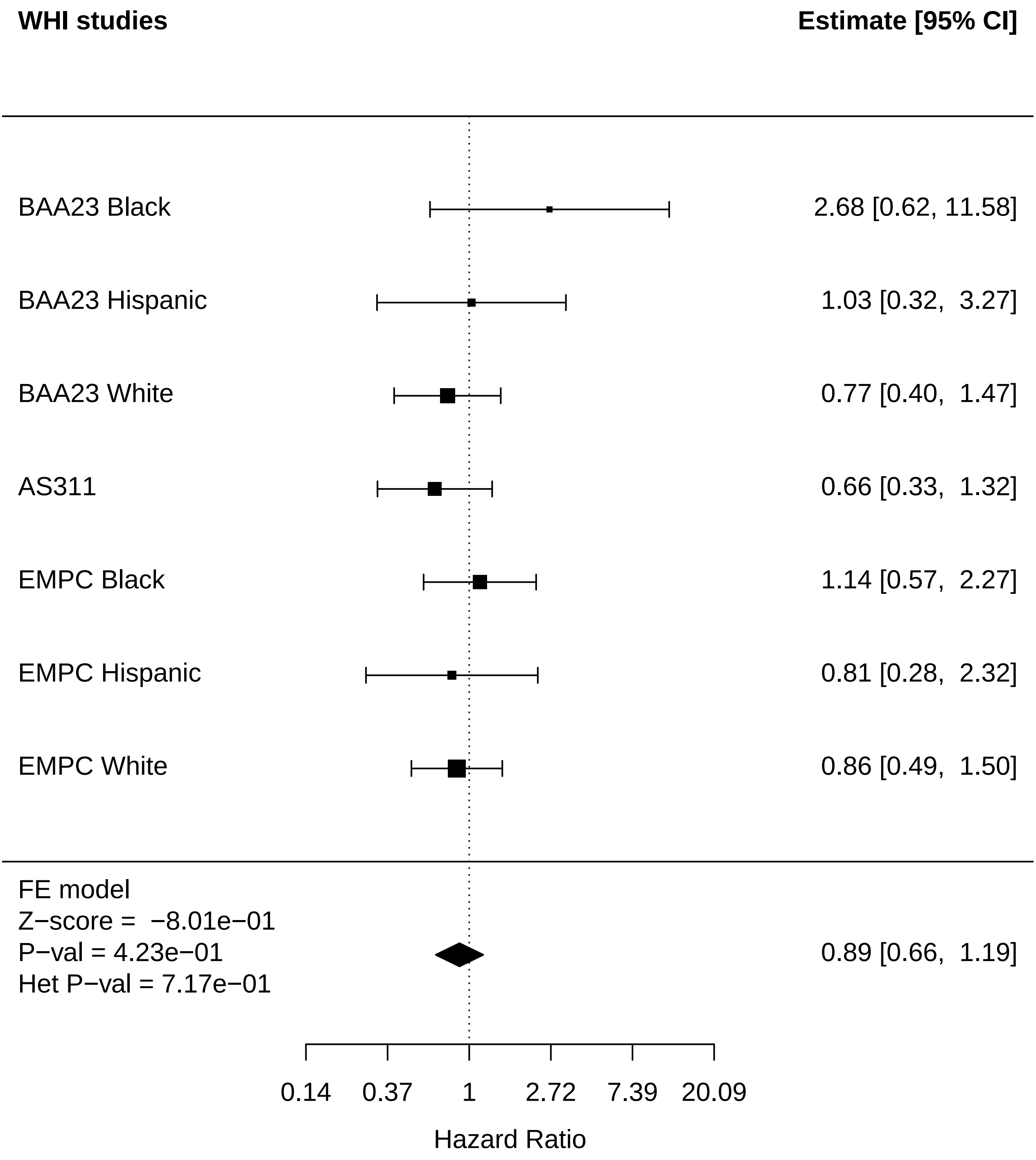

### Liver_MI.pdf

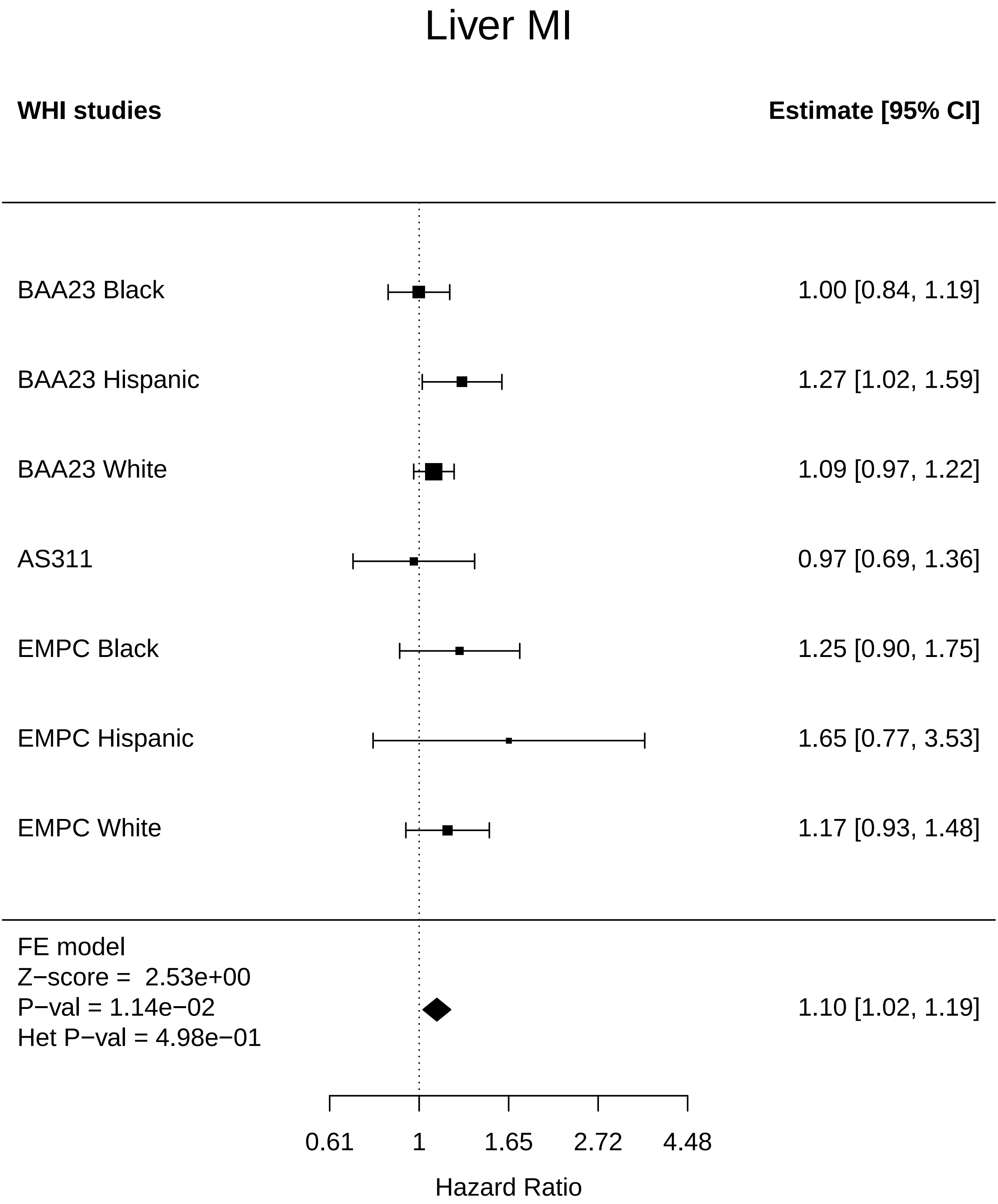

### Liver_Physical function.pdf

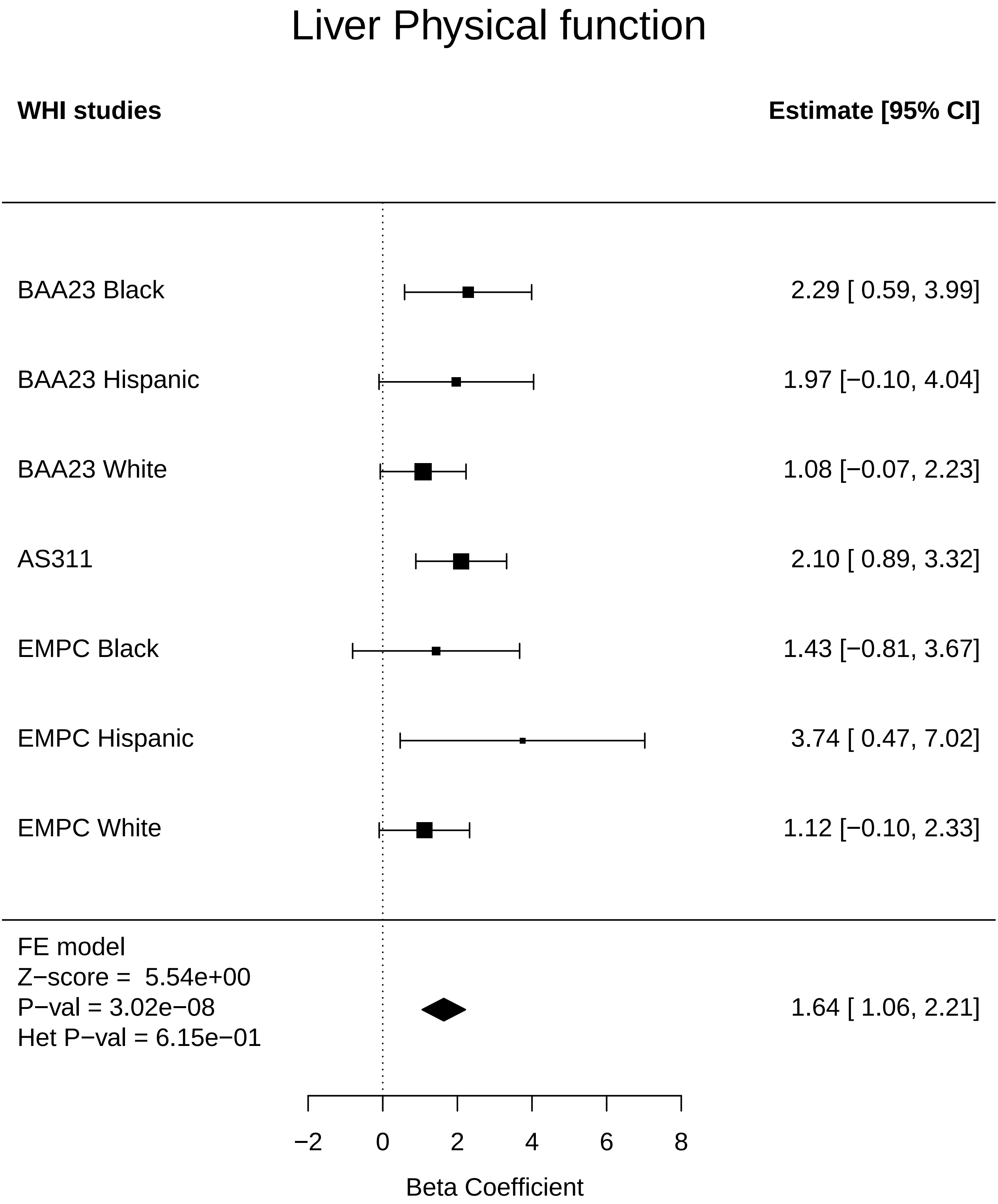

### Liver_Stroke.pdf

#### Liver Stroke

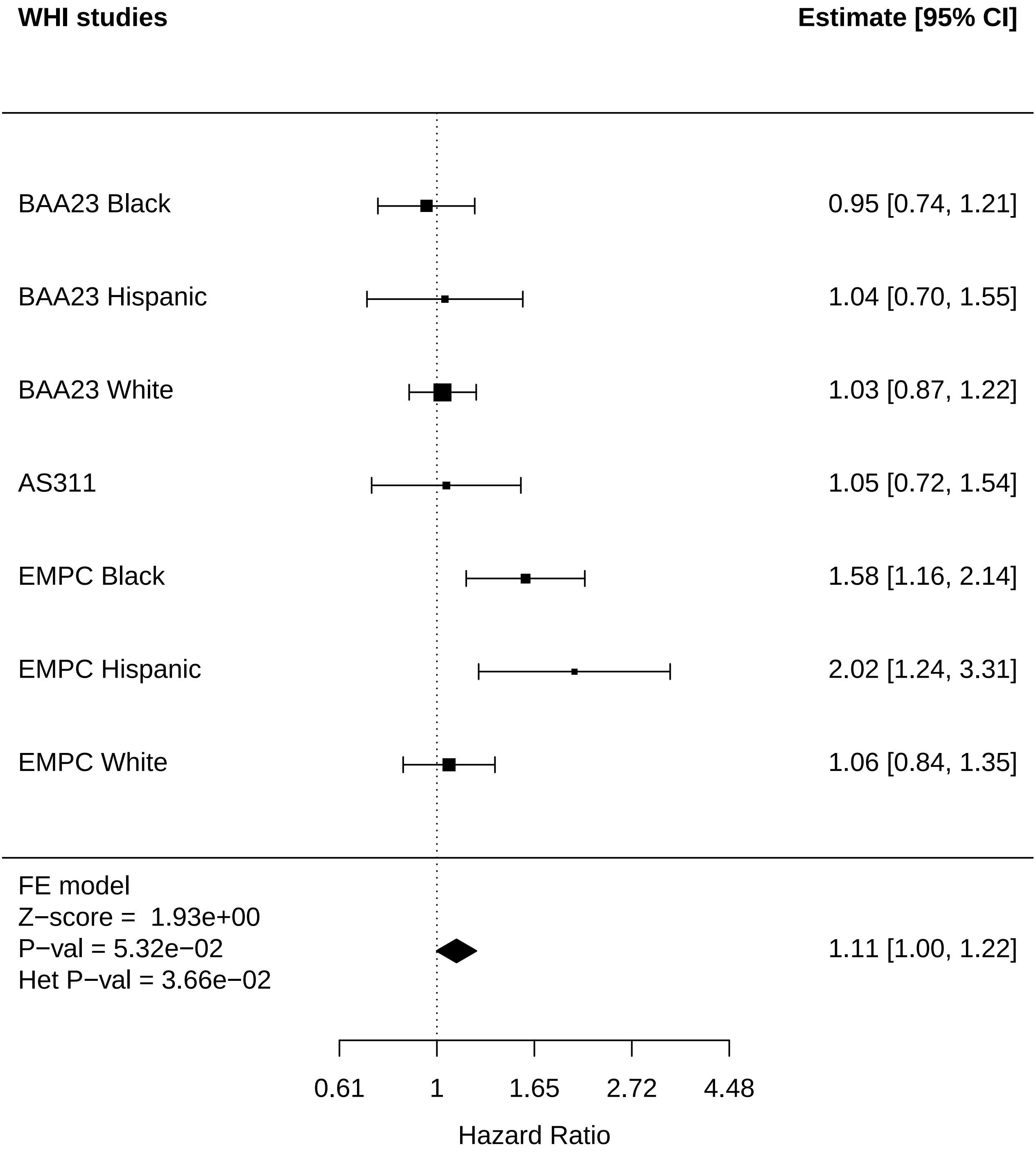

### Lung_Cataract.pdf

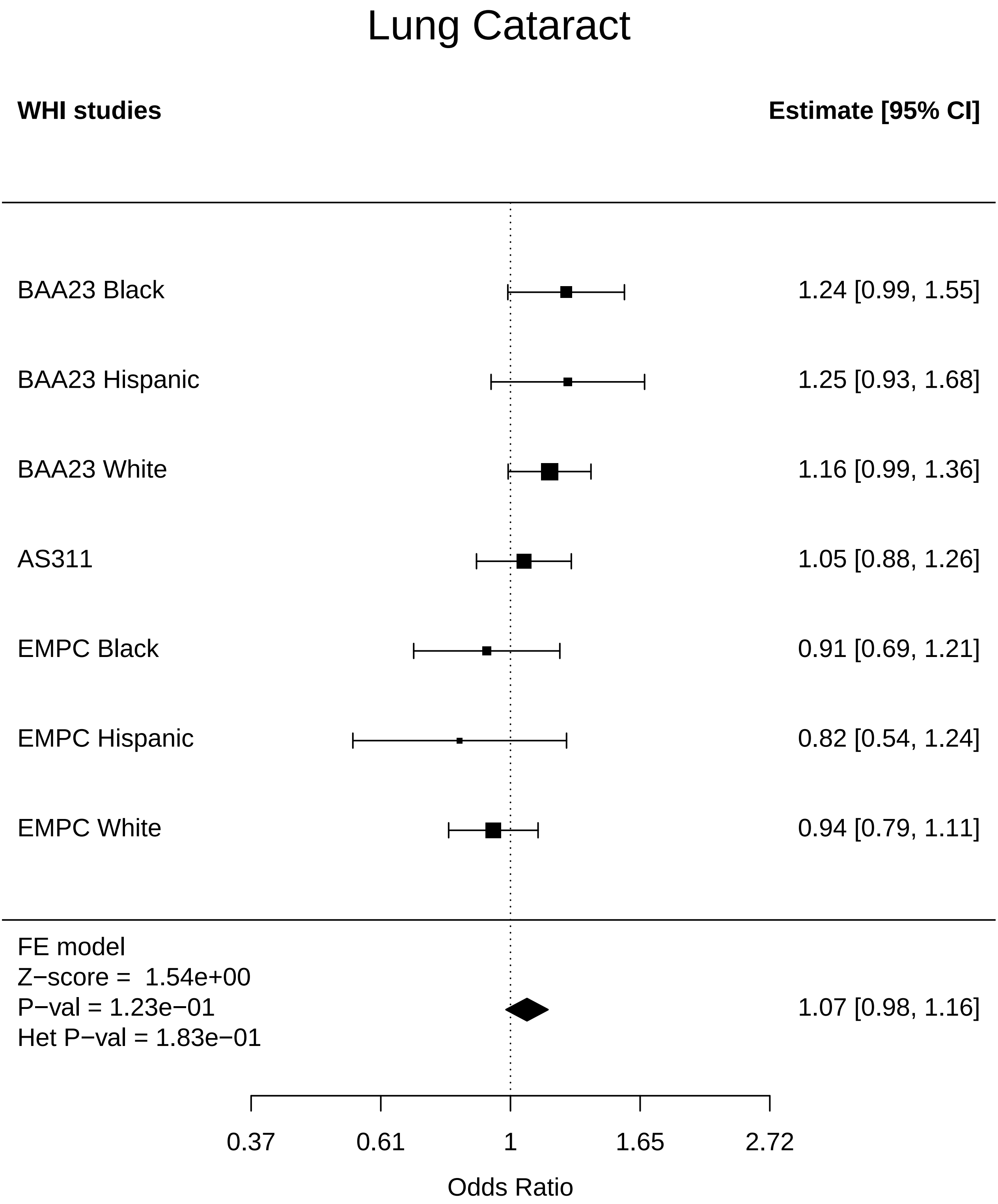

### Lung_CHD.pdf

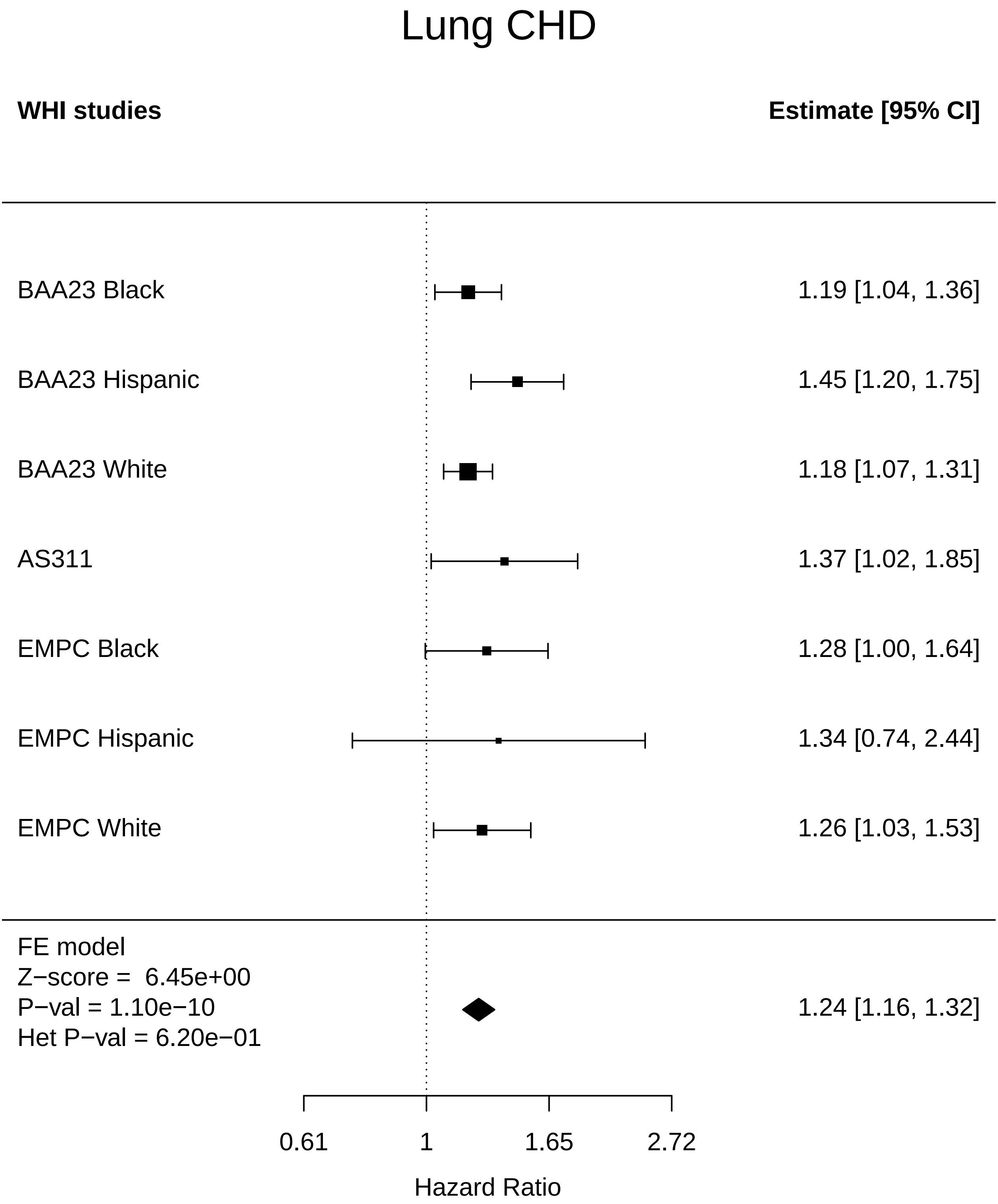

### Lung_Cognitive function.pdf

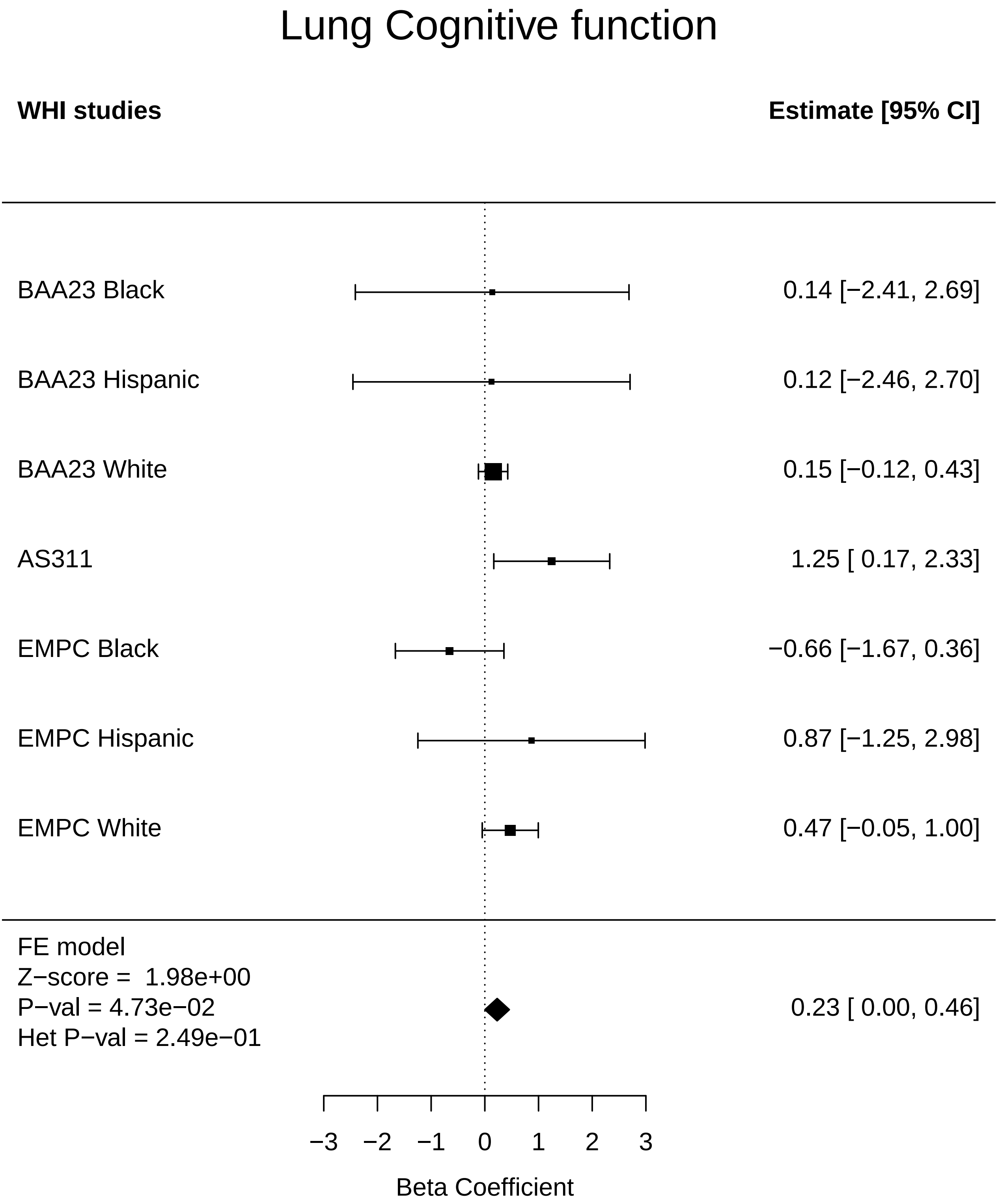

### Lung_Diabetes.pdf

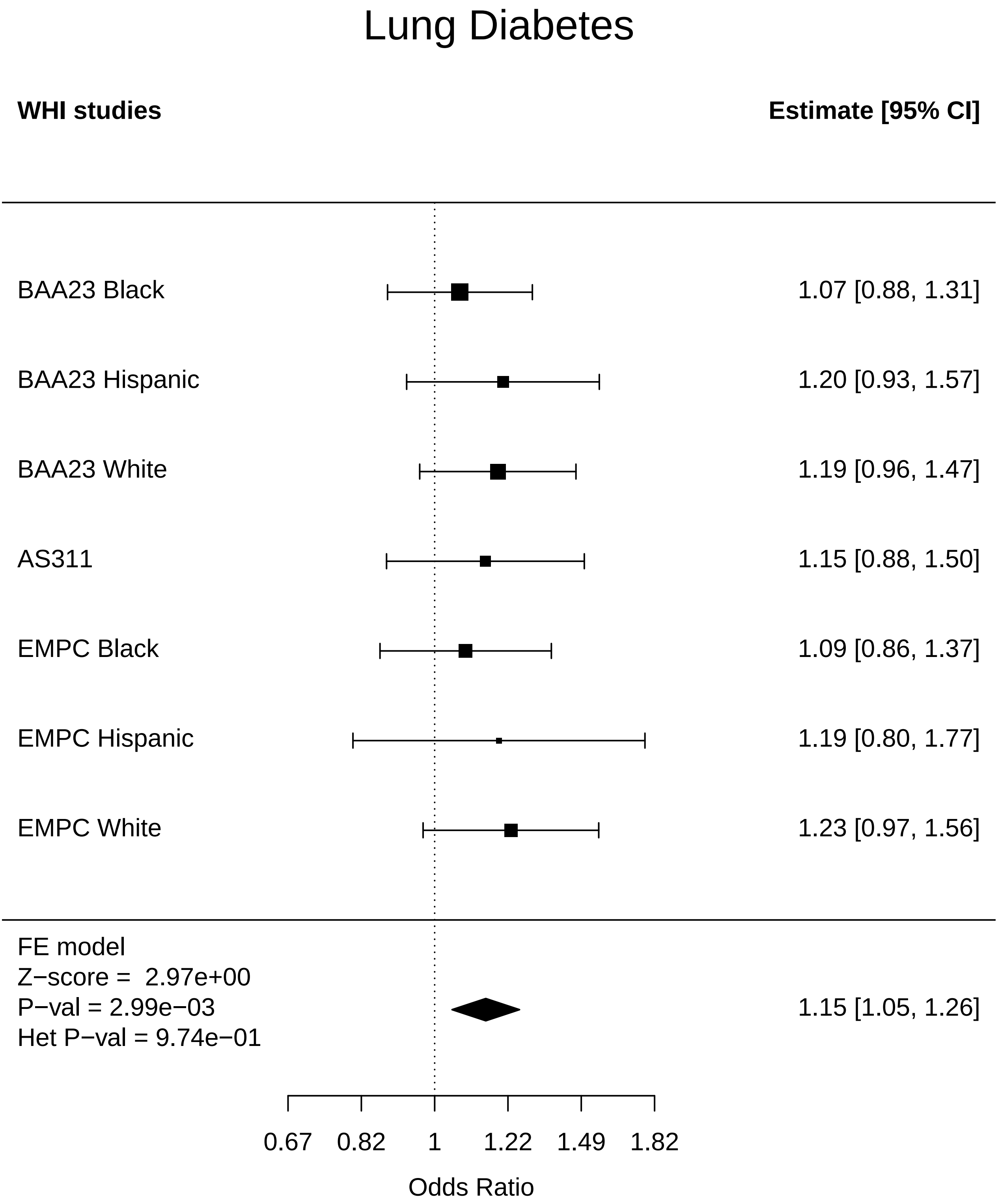

### Lung_Disease free.pdf

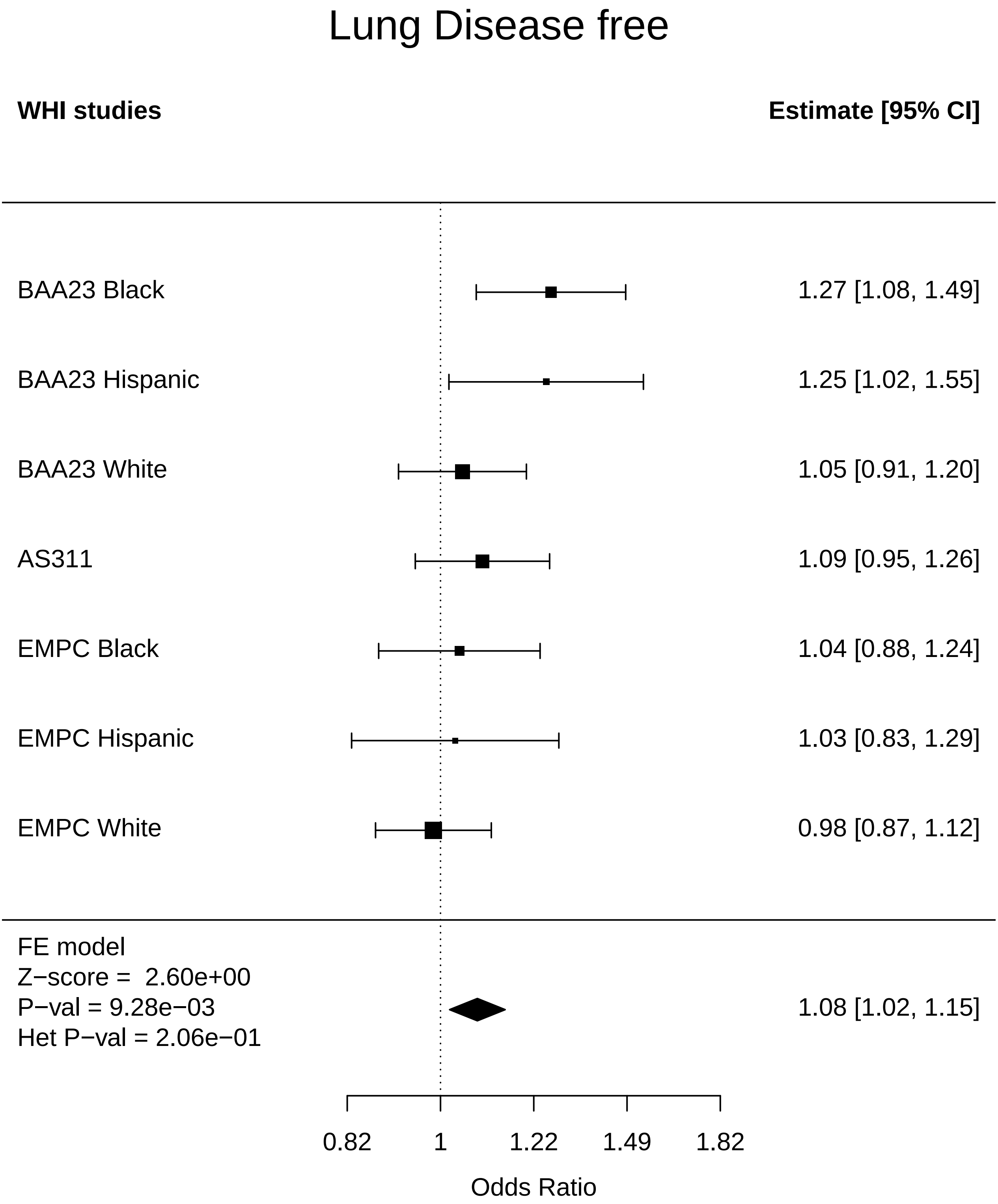

### Lung_Endometrial cancer.pdf

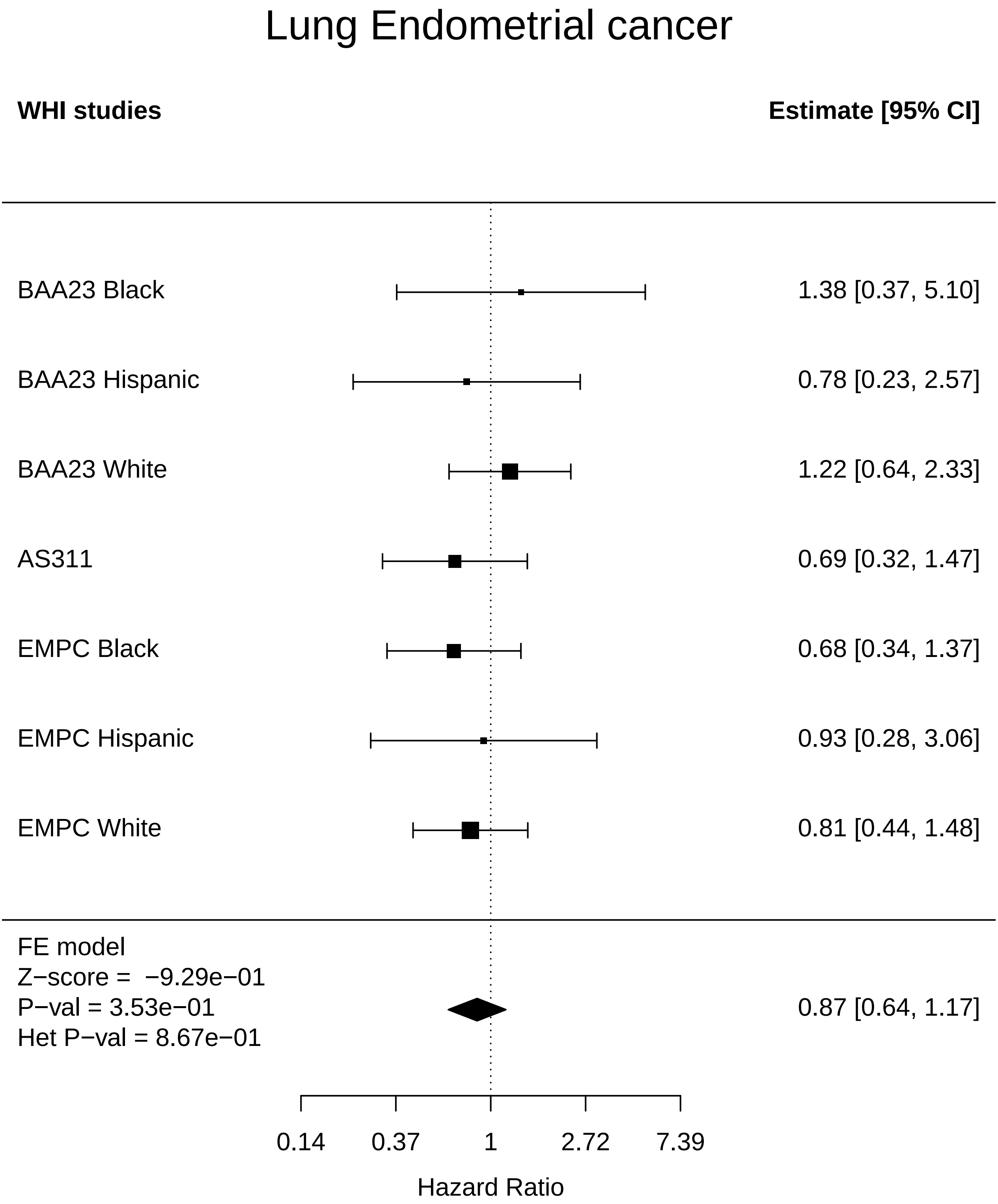

### Lung_Mortality.pdf

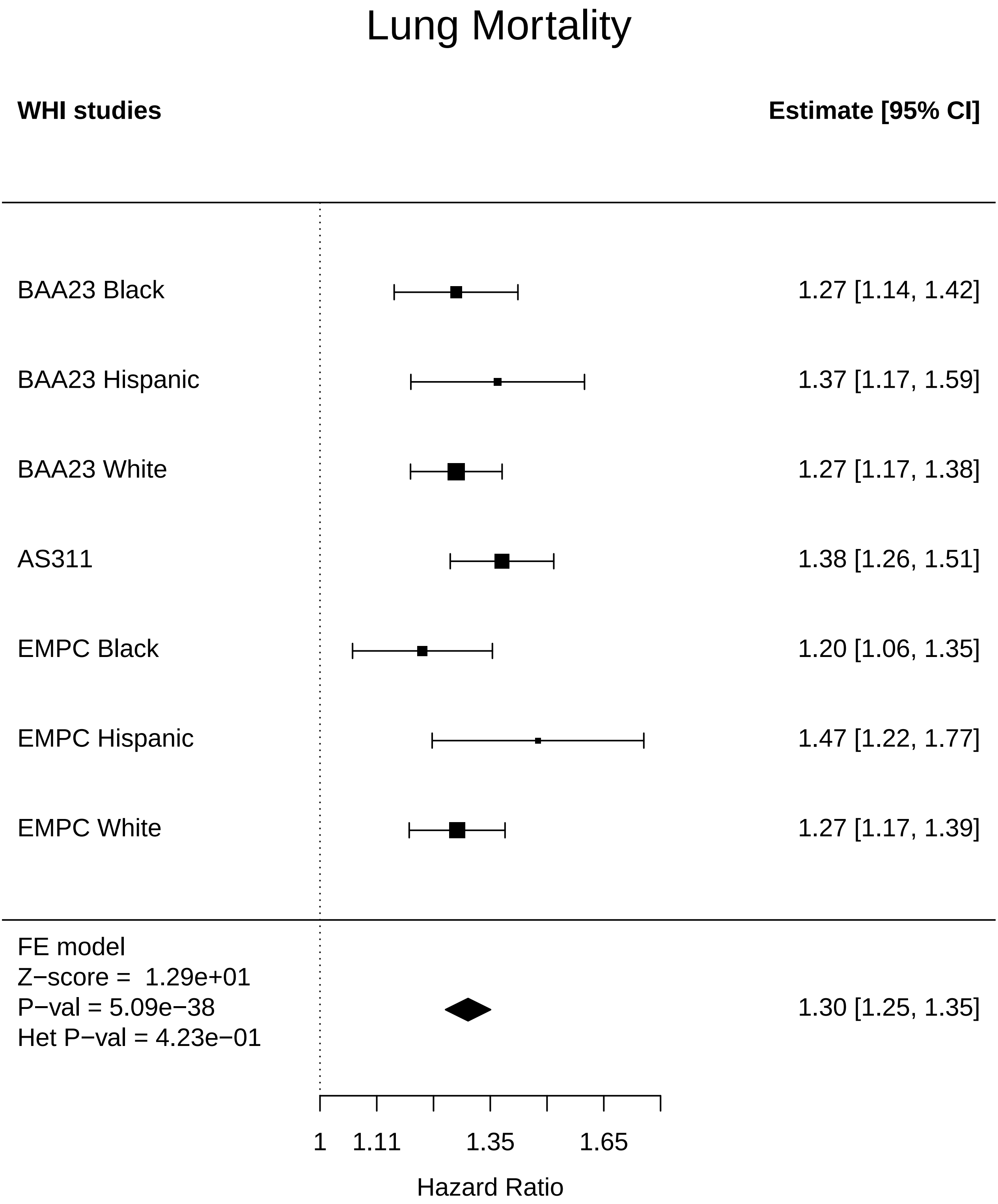

### Metabolic_Any Cancer.pdf

#### Metabolic Any Cancer

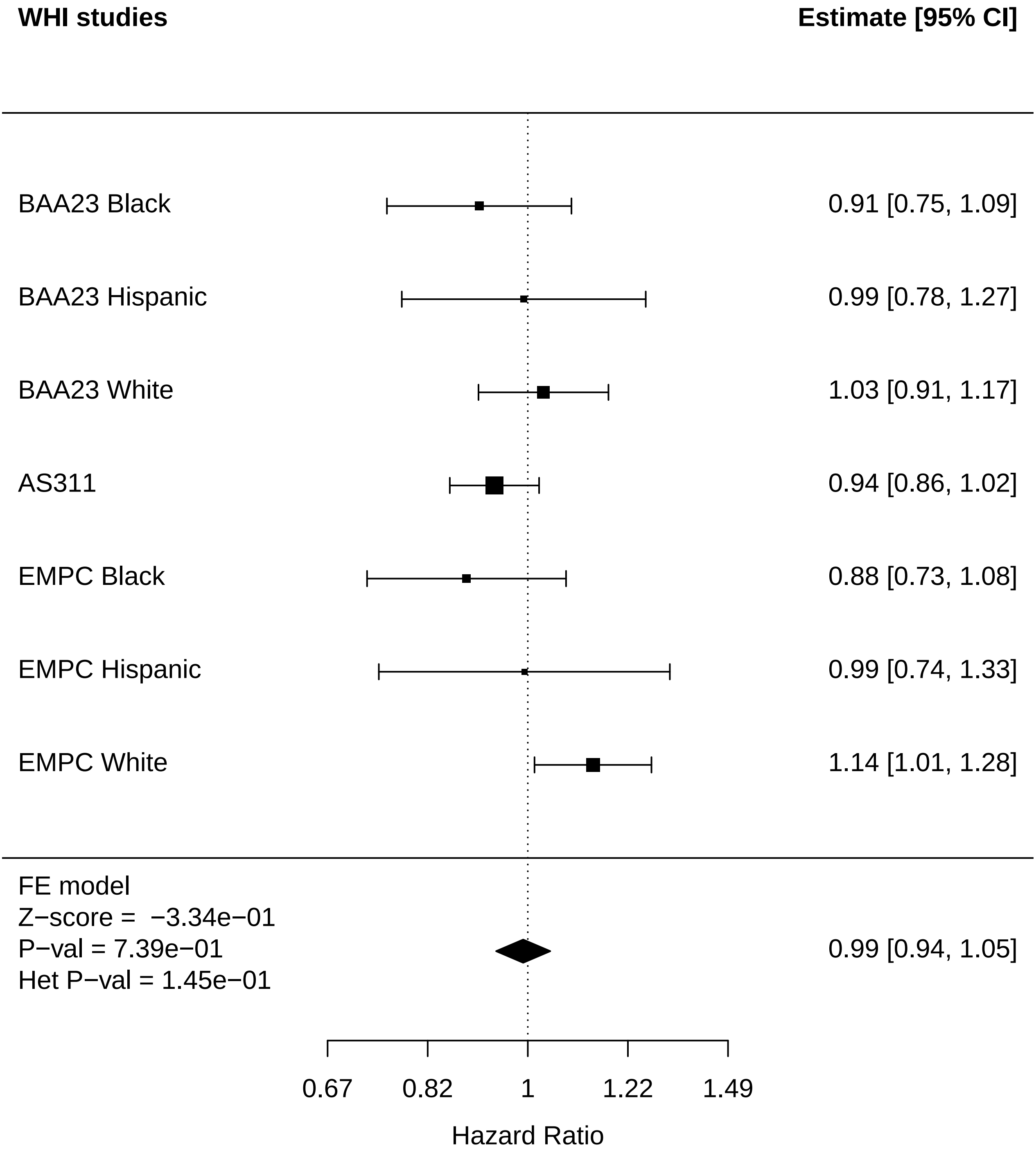

### Metabolic_Arthritis.pdf

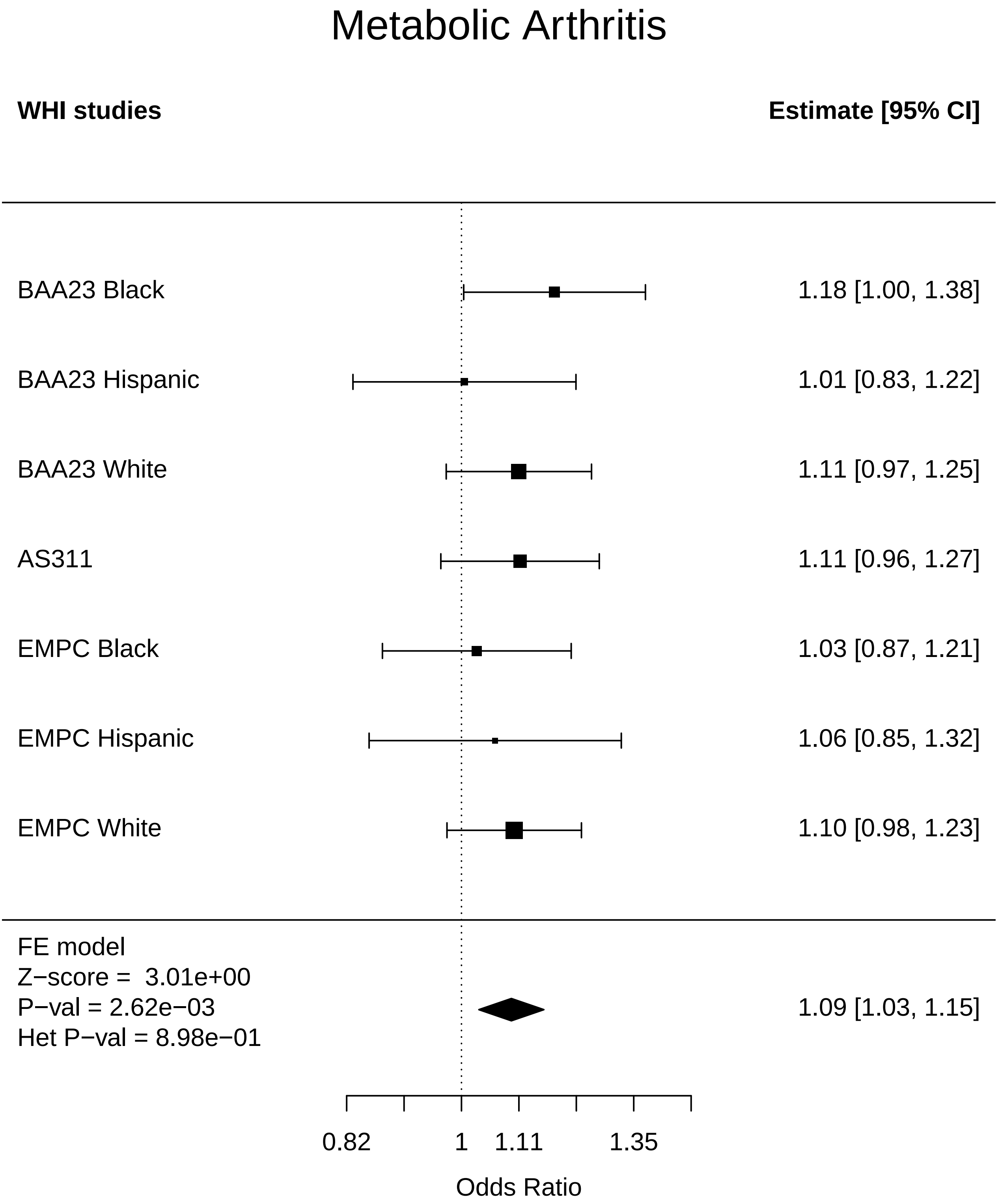

### Metabolic_Breast Cancer.pdf

#### Metabolic Breast Cancer

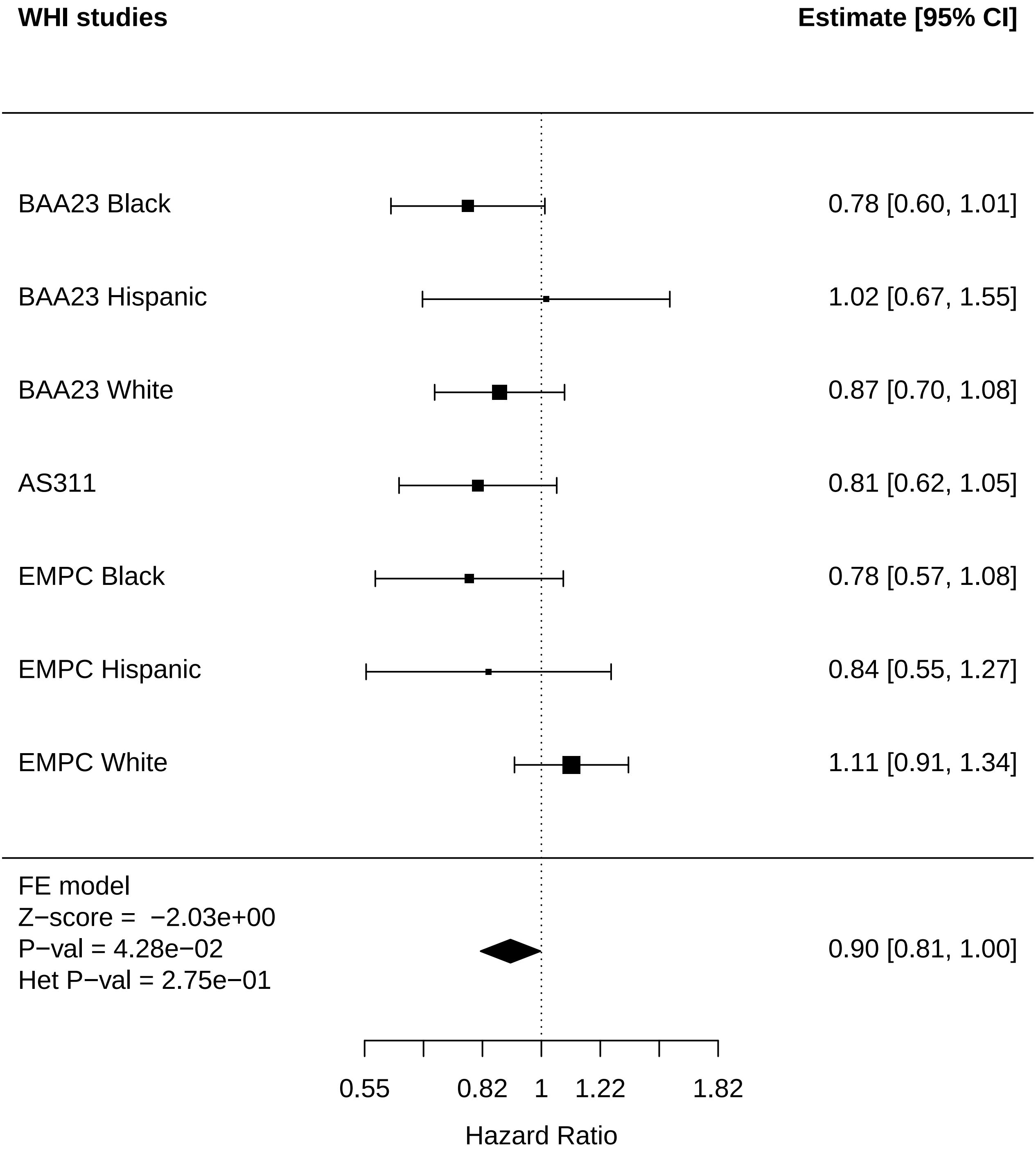

### Metabolic_CHD.pdf

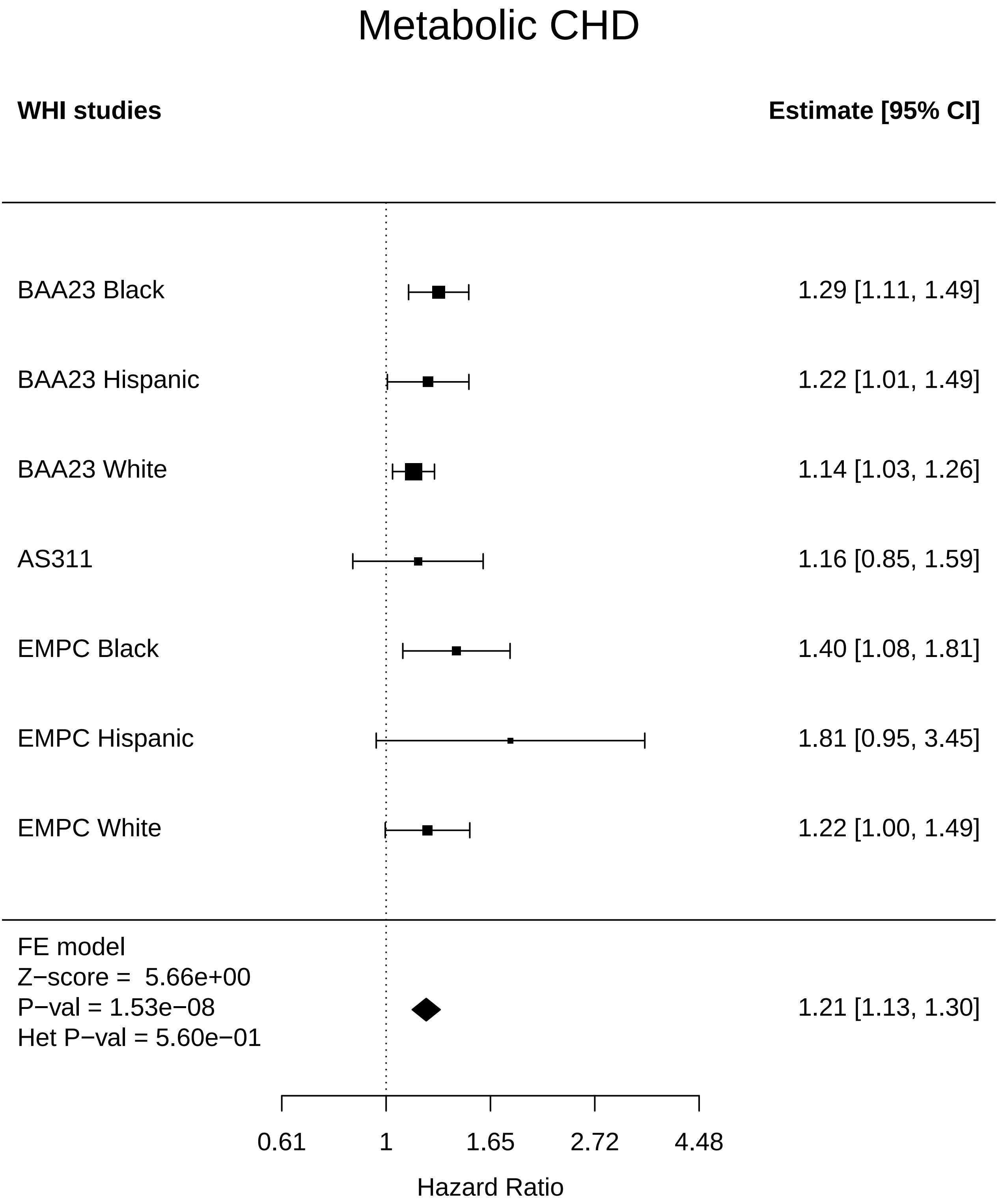

### Metabolic_Diabetes.pdf

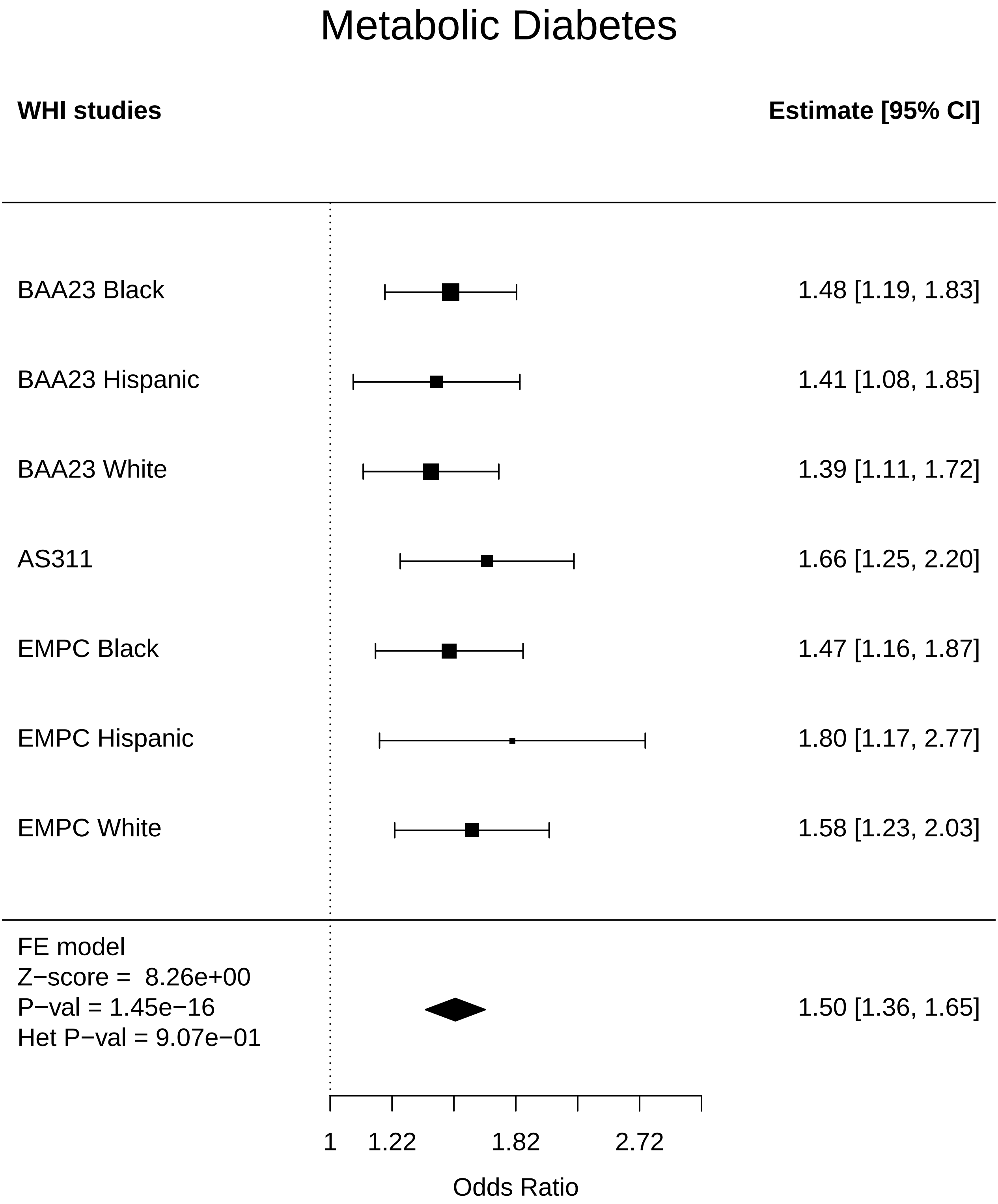

### Metabolic_Disease free.pdf

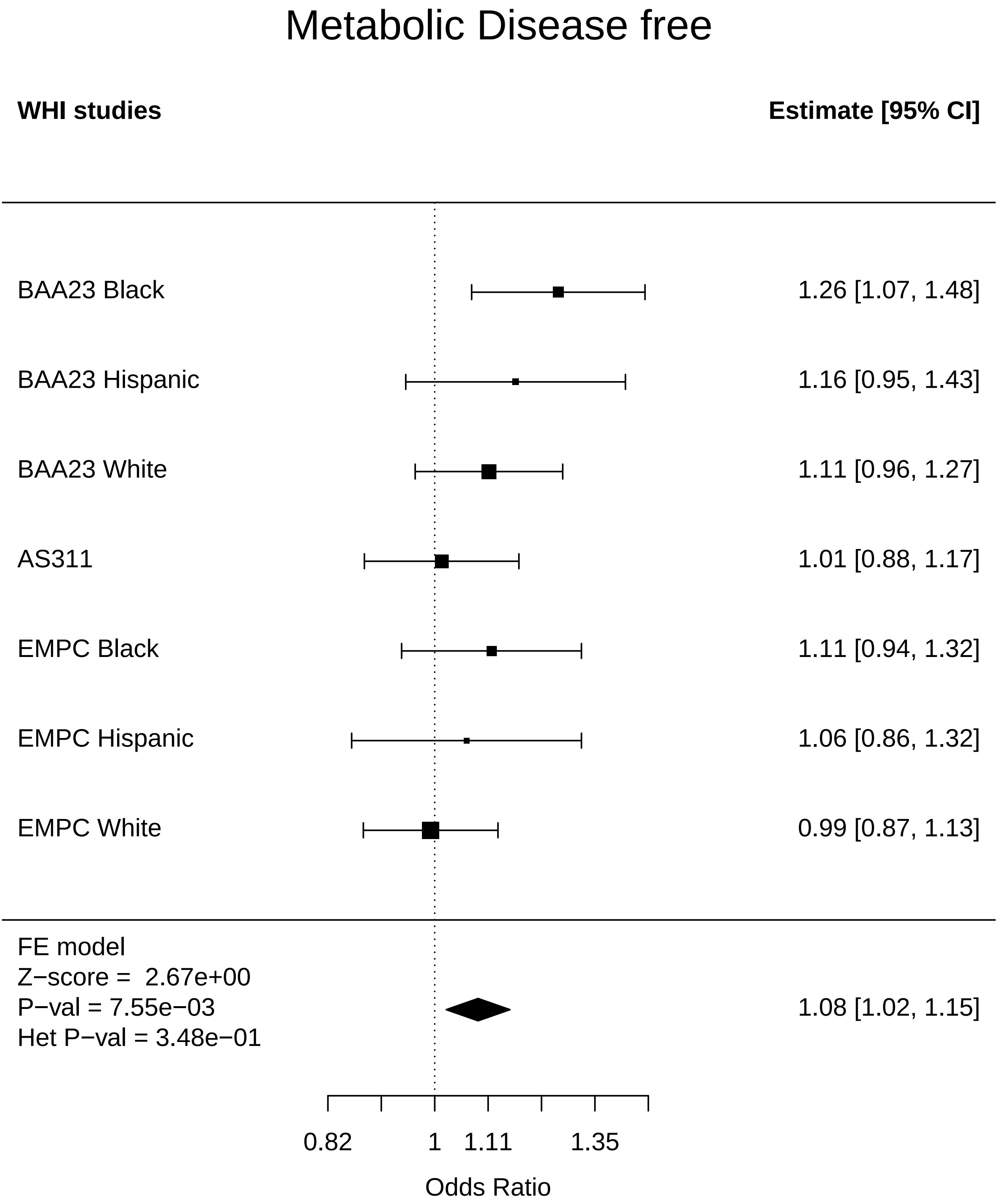

### Metabolic_Endometrial cancer.pdf

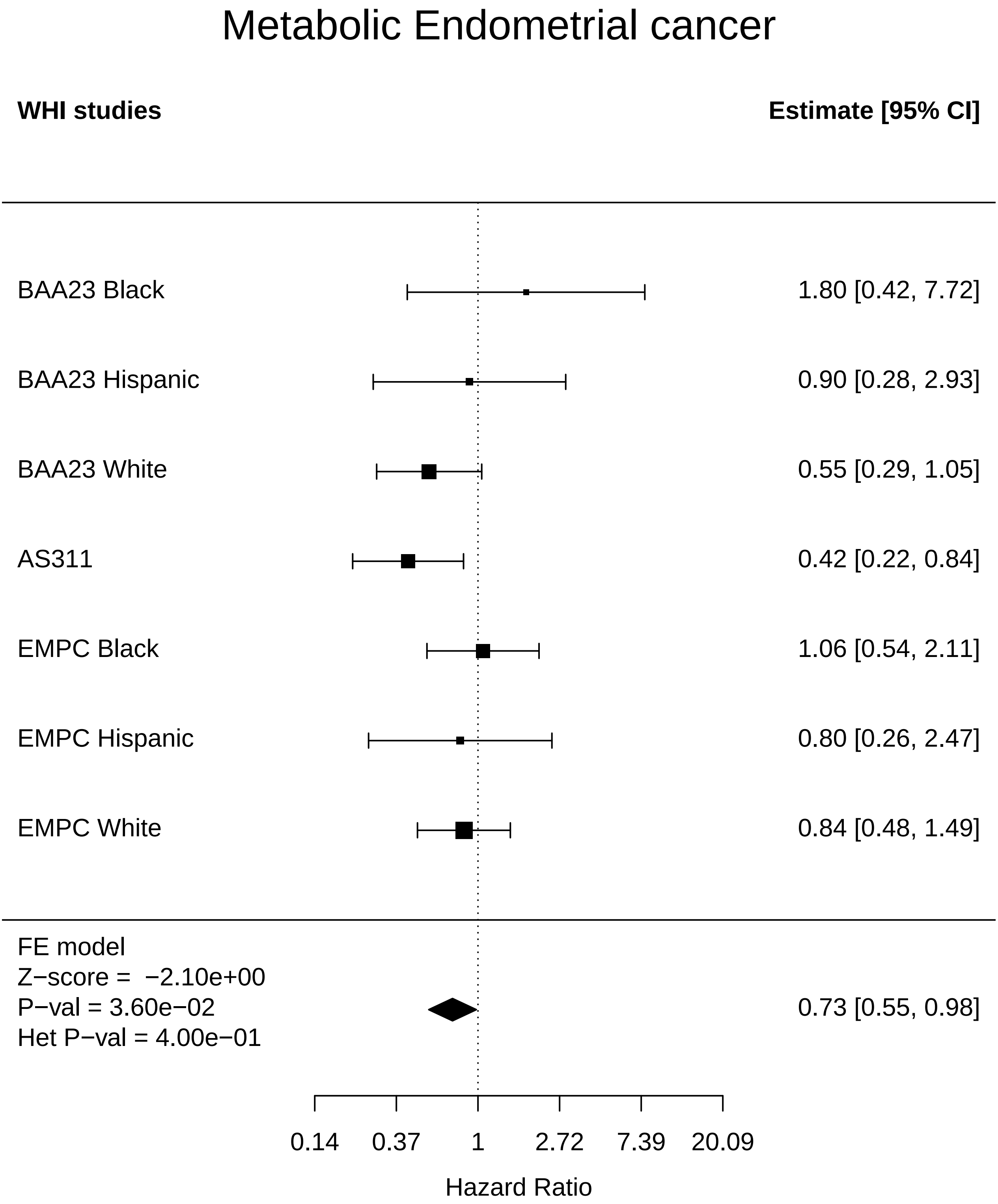

### Metabolic_Lung Cancer.pdf

#### Metabolic Lung Cancer

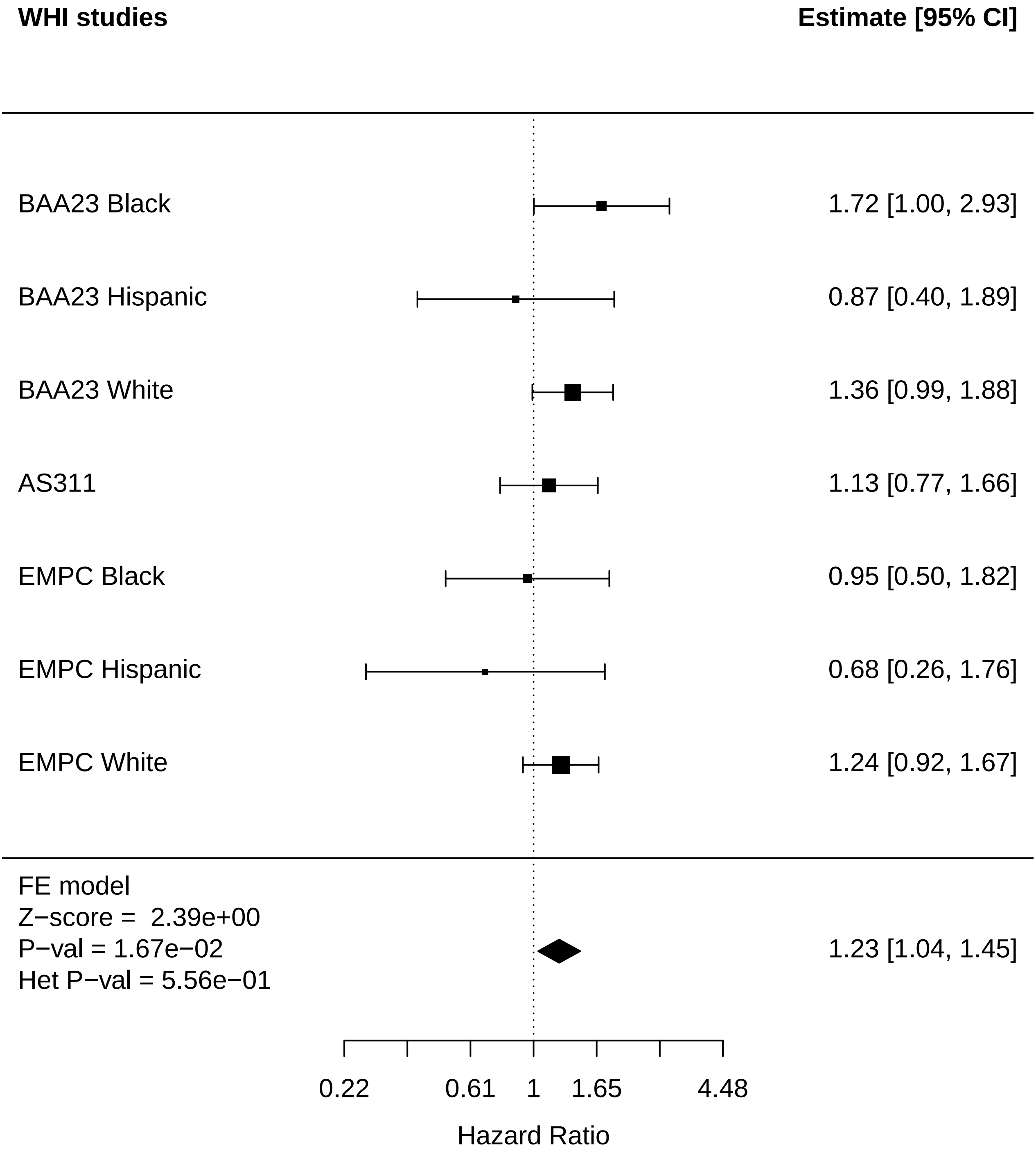

### Metabolic_MI.pdf

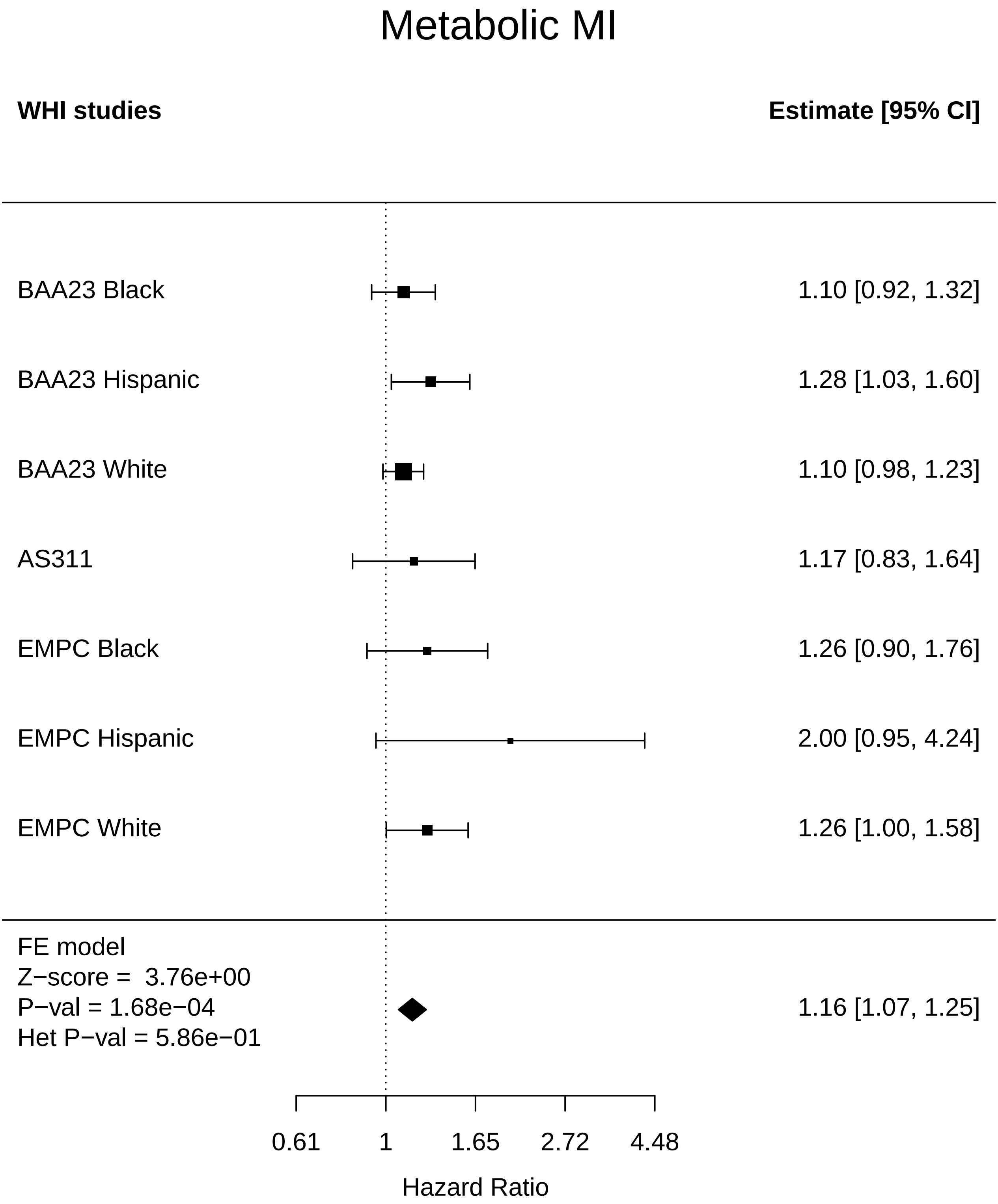

### Metabolic_Mortality.pdf

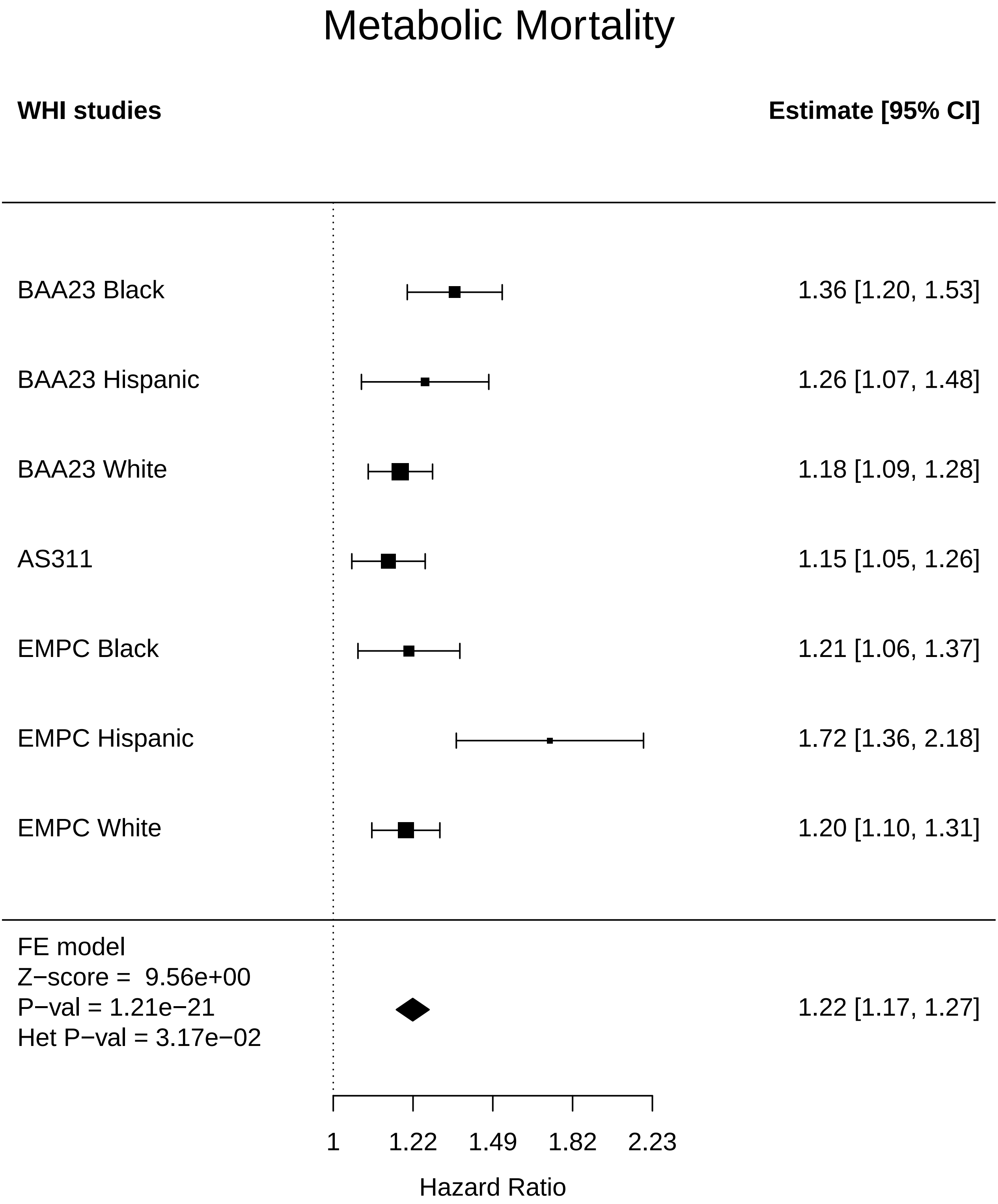

### Metabolic_Stroke.pdf

#### Metabolic Stroke

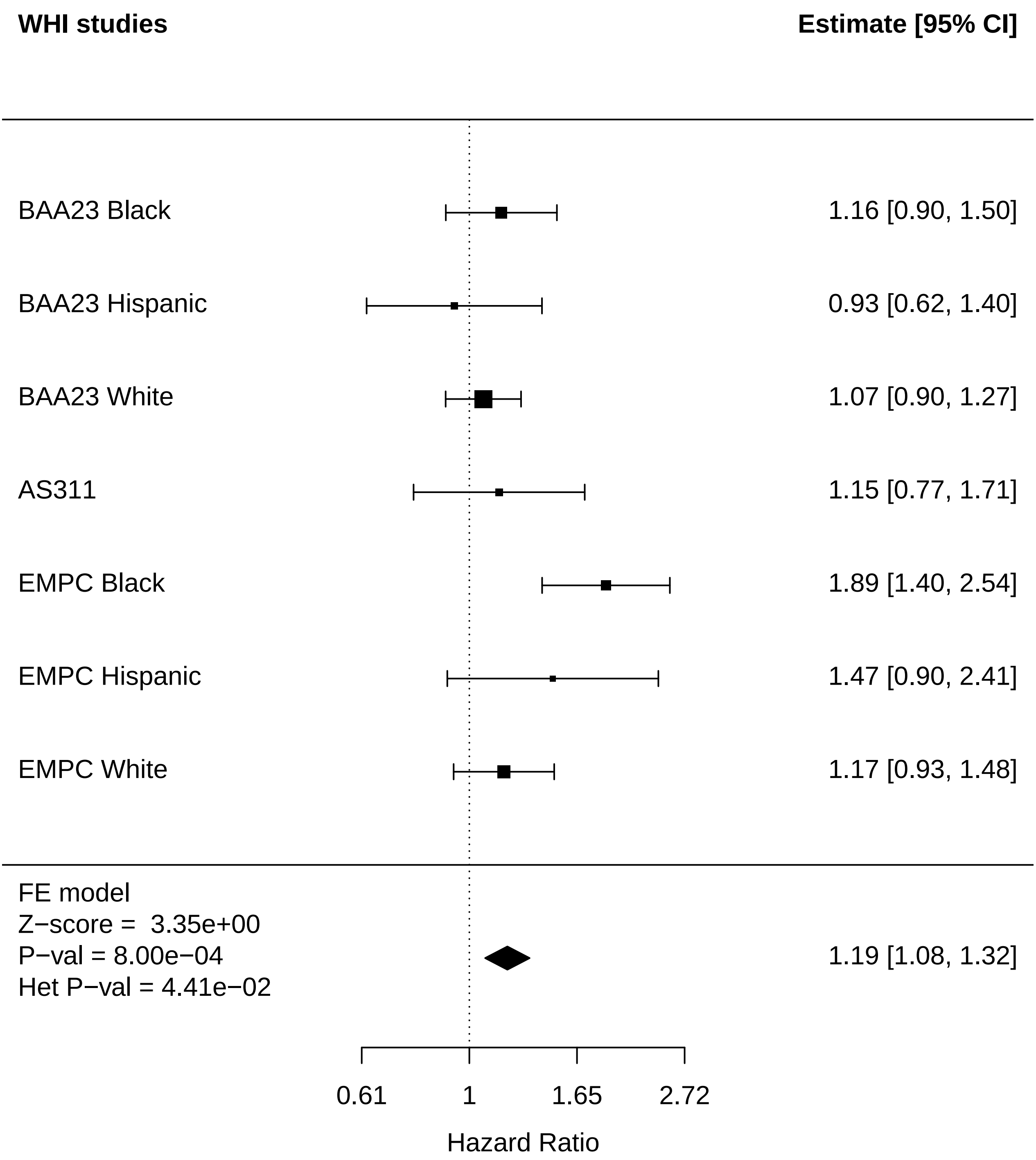

### Metabolic_Thyroid disease.pdf

#### Metabolic Thyroid disease

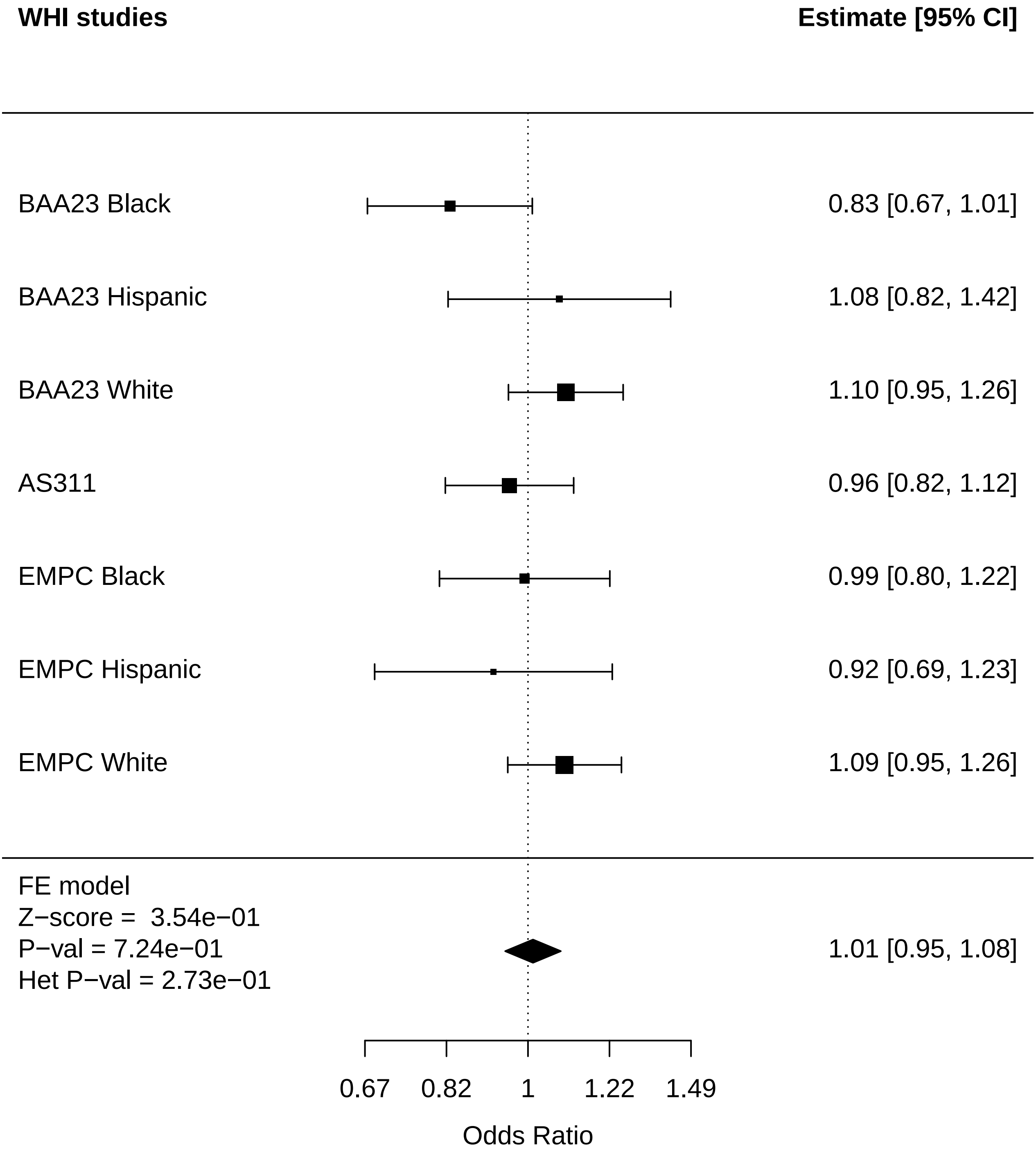

### MusculoSkeletal_Cataract.pdf

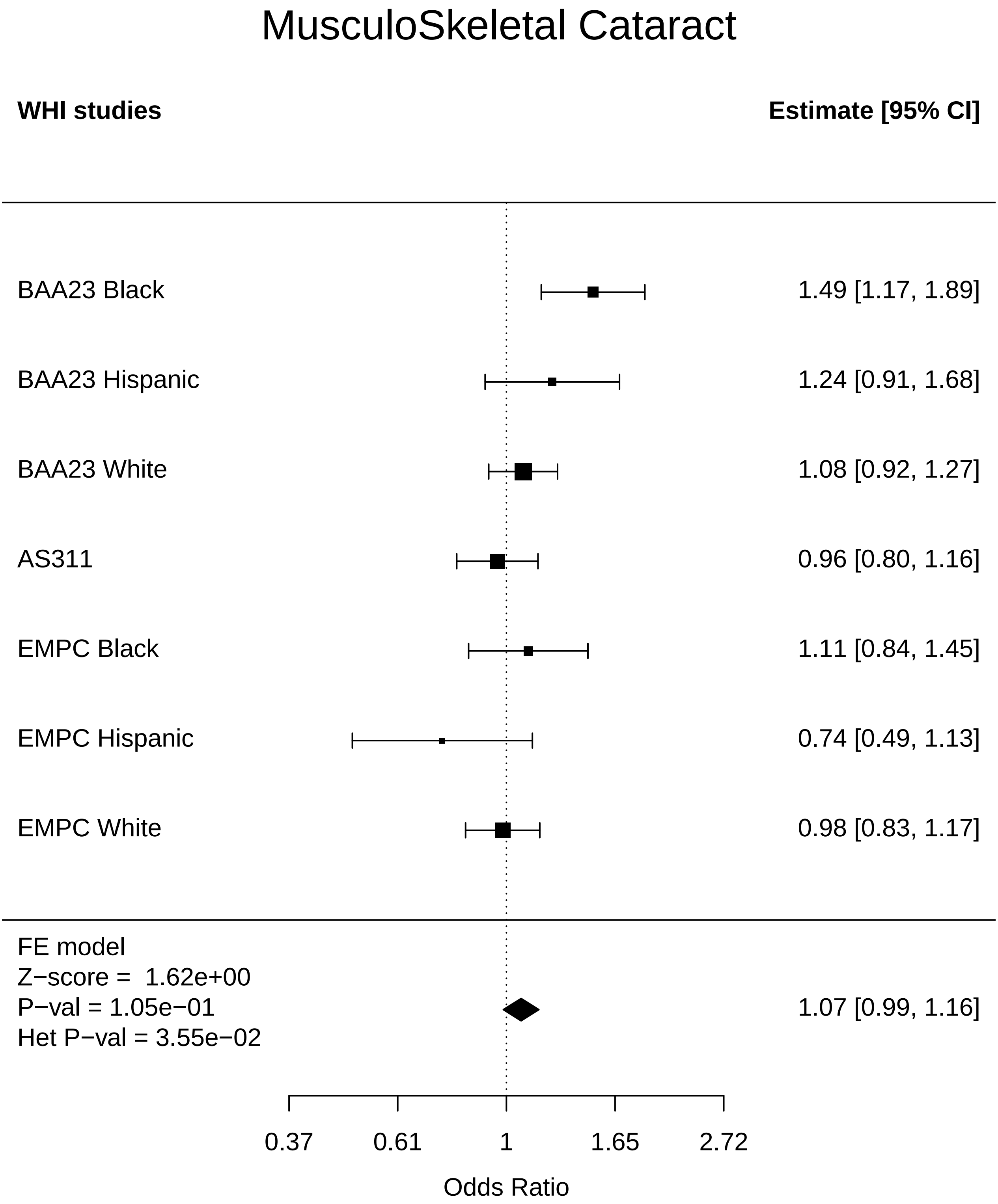

### MusculoSkeletal_CHD.pdf

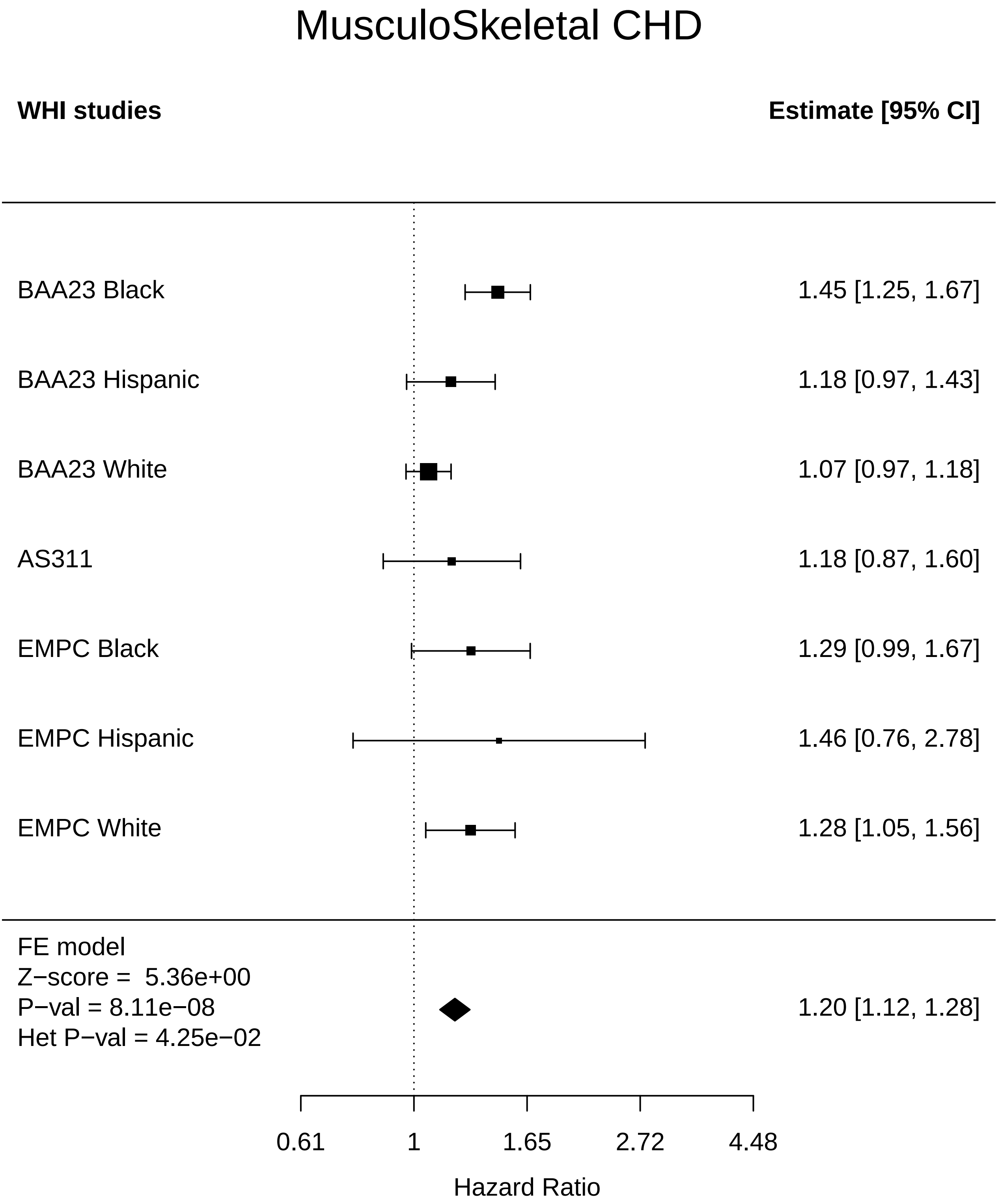

### MusculoSkeletal_Physical function.pdf

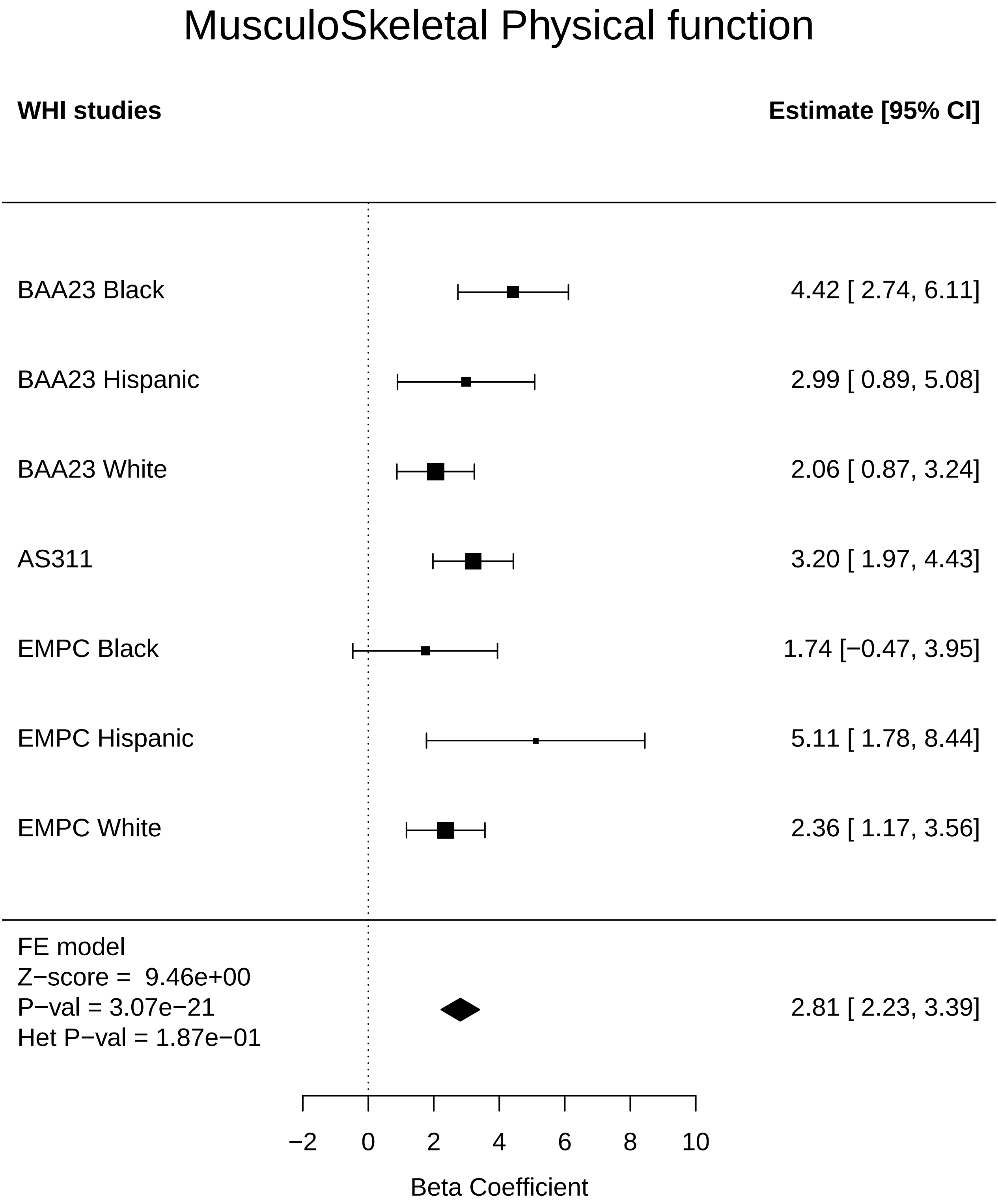
