## Supplementary figures and images for "Systems Age: A single blood methylation test to quantify aging heterogeneity across 11 physiological systems"

### Age_prediction_Any Cancer.pdf

#### Age_prediction Any Cancer

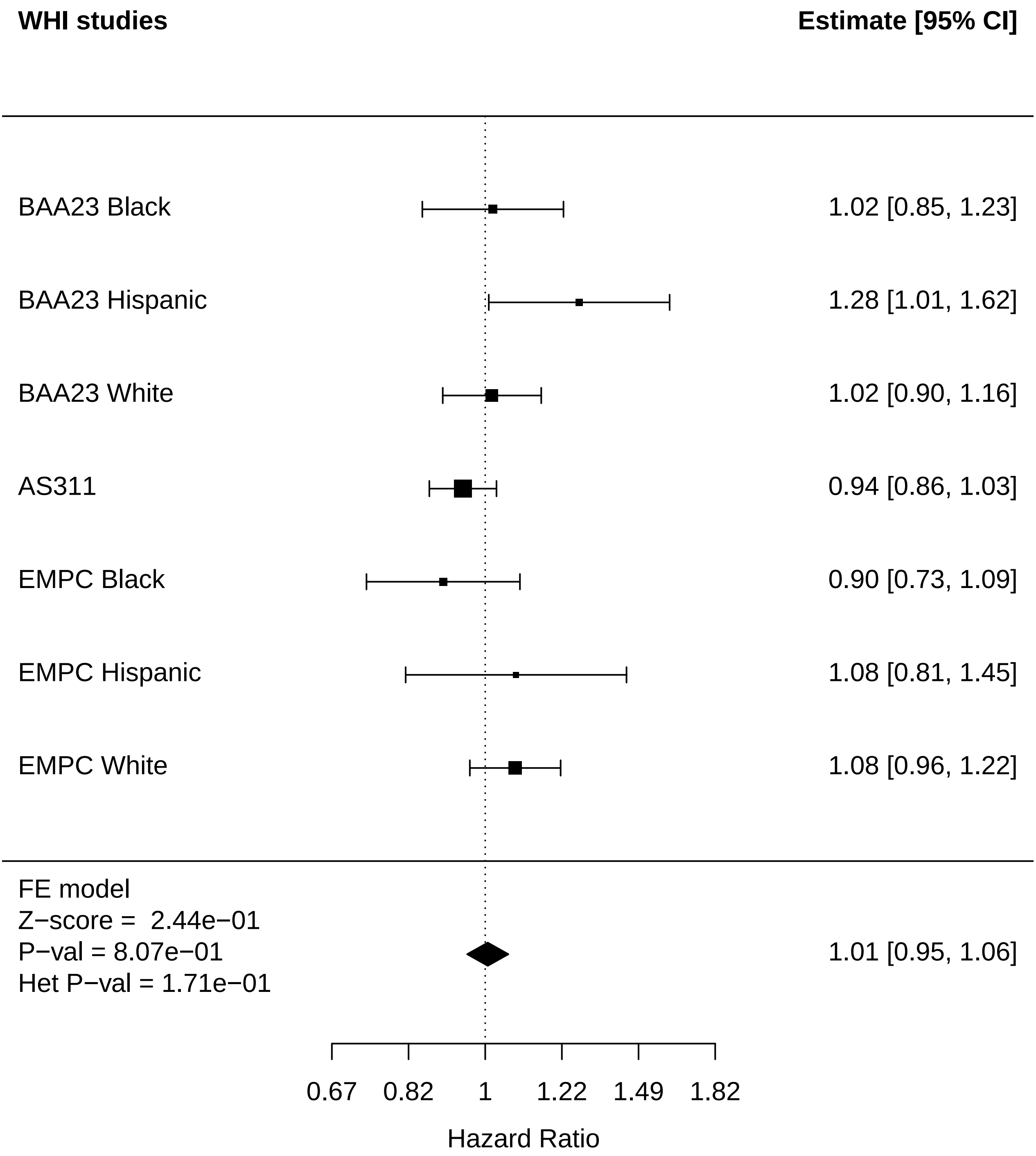

### Age_prediction_Arthritis.pdf

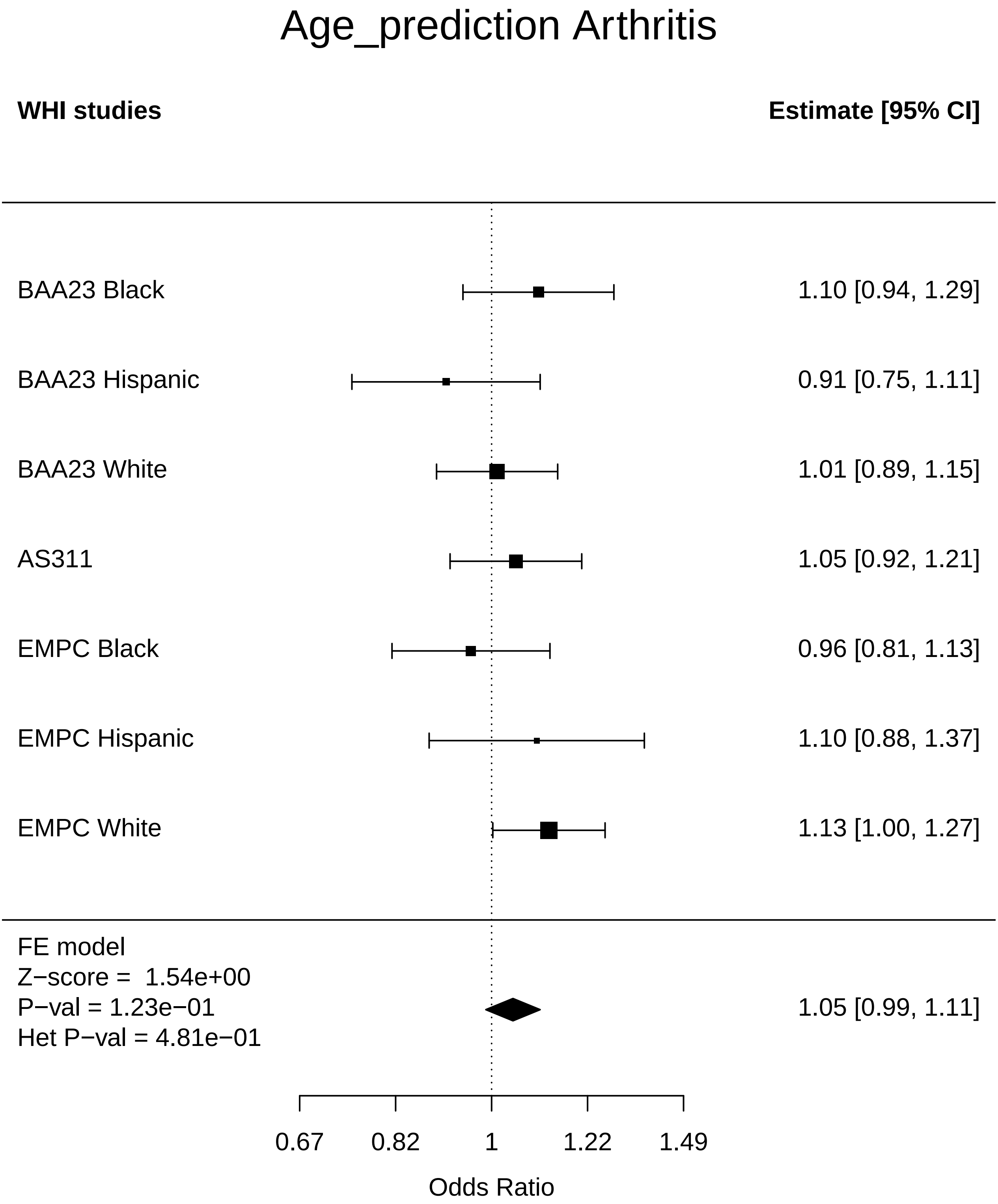

### Age_prediction_Breast Cancer.pdf

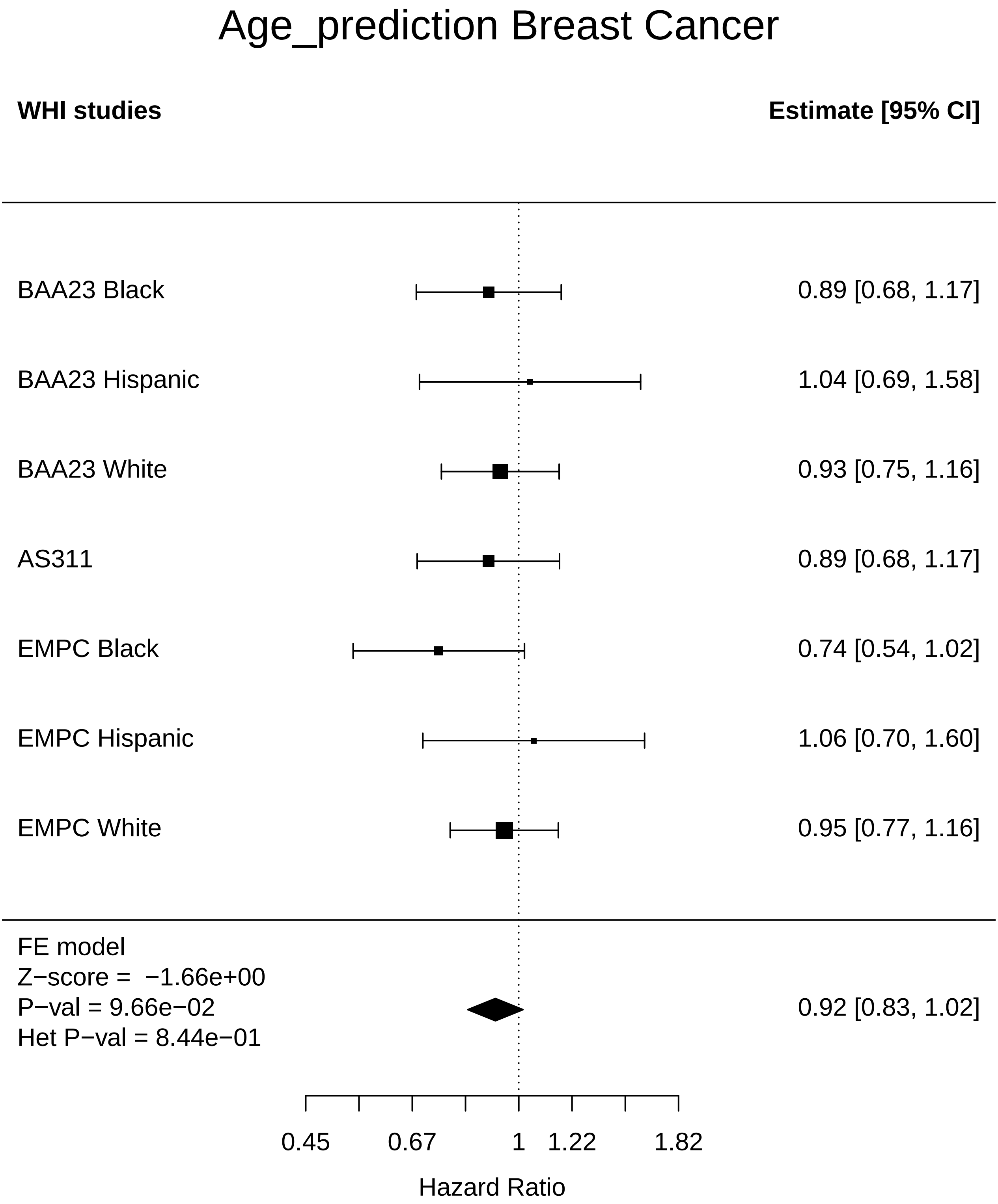

### Age_prediction_Cataract.pdf

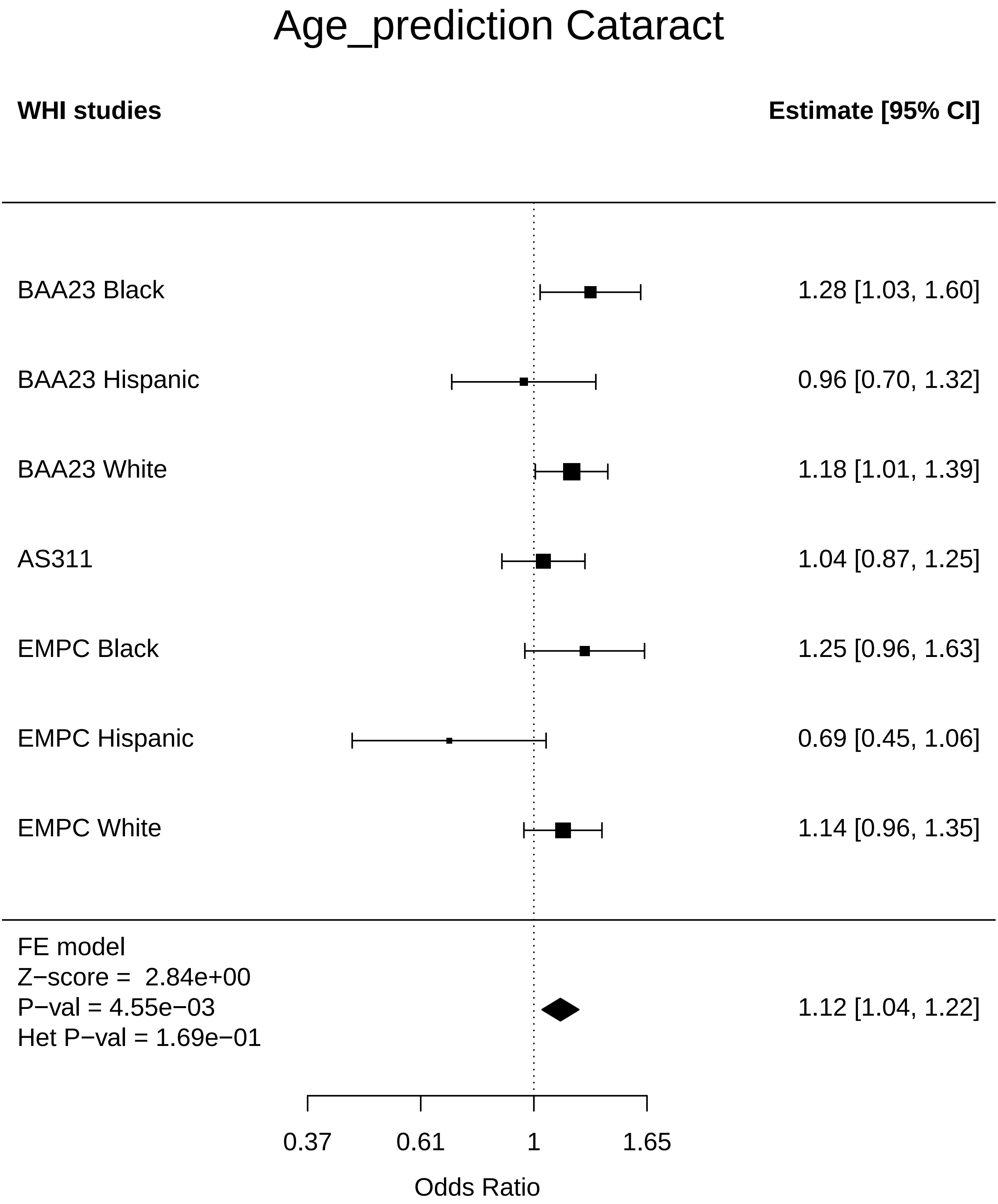

### Age_prediction_CHD.pdf

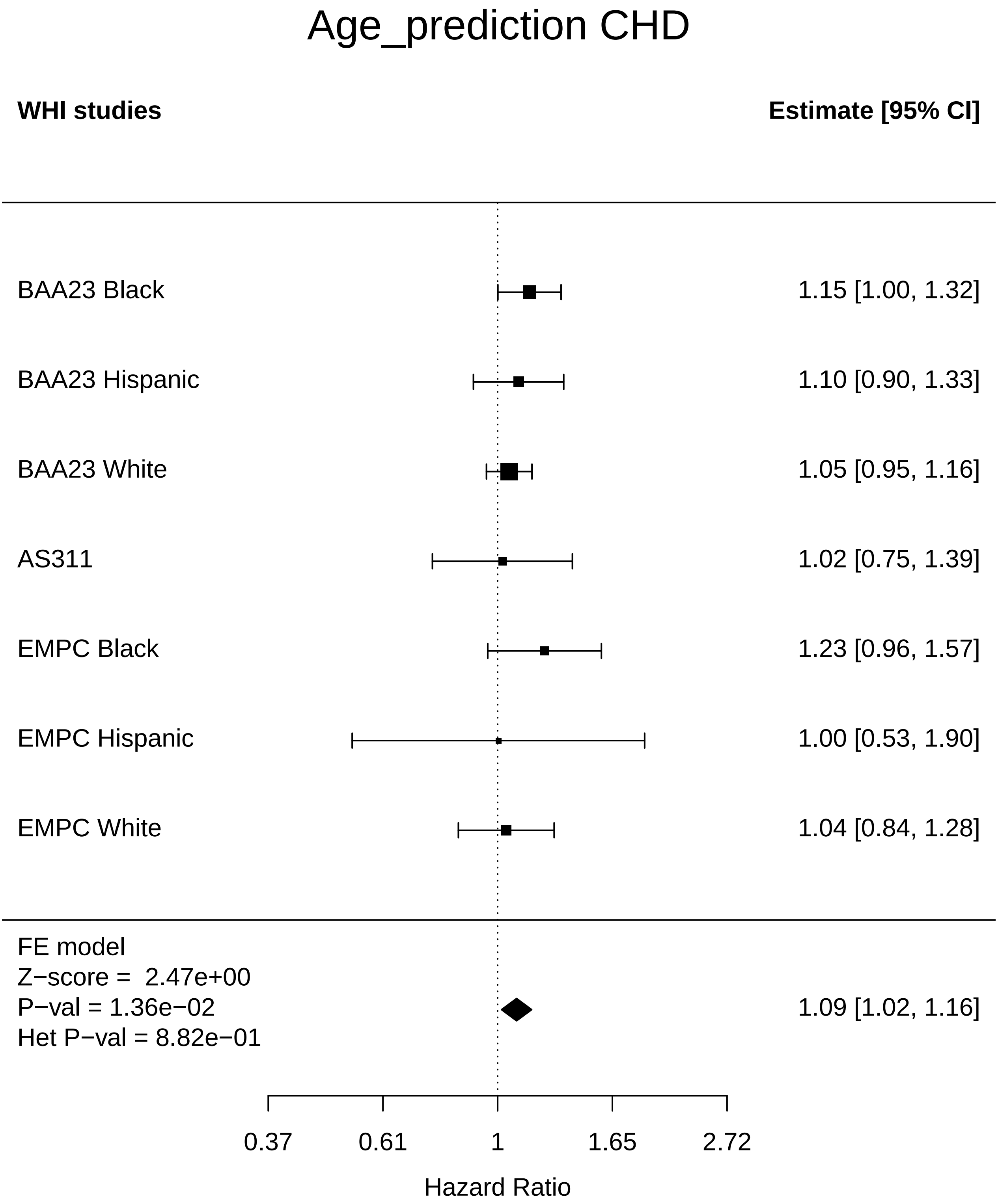

### Age_prediction_Cognitive function.pdf

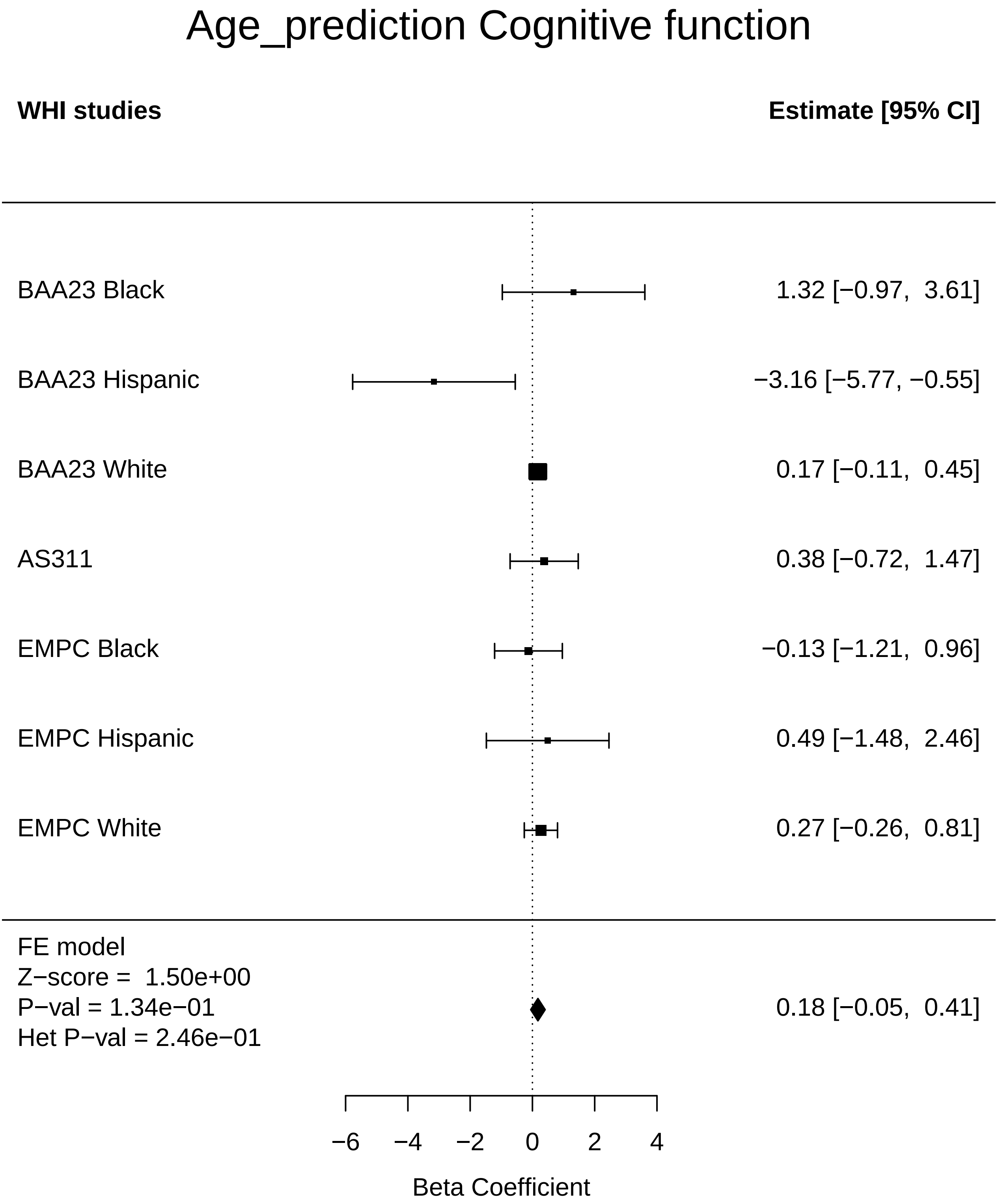

### Age_prediction_Colorectal cancer.pdf

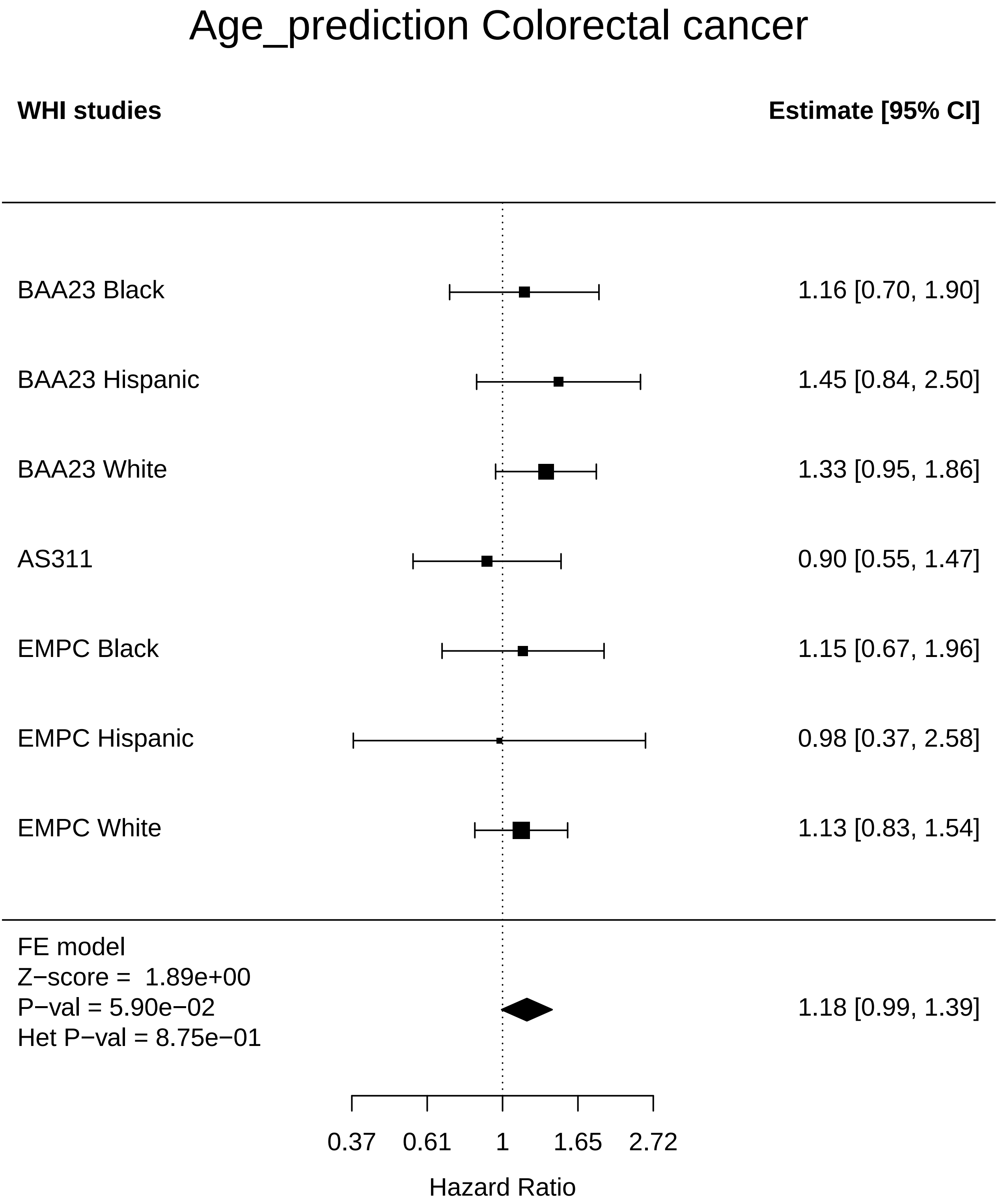

### Age_prediction_Diabetes.pdf

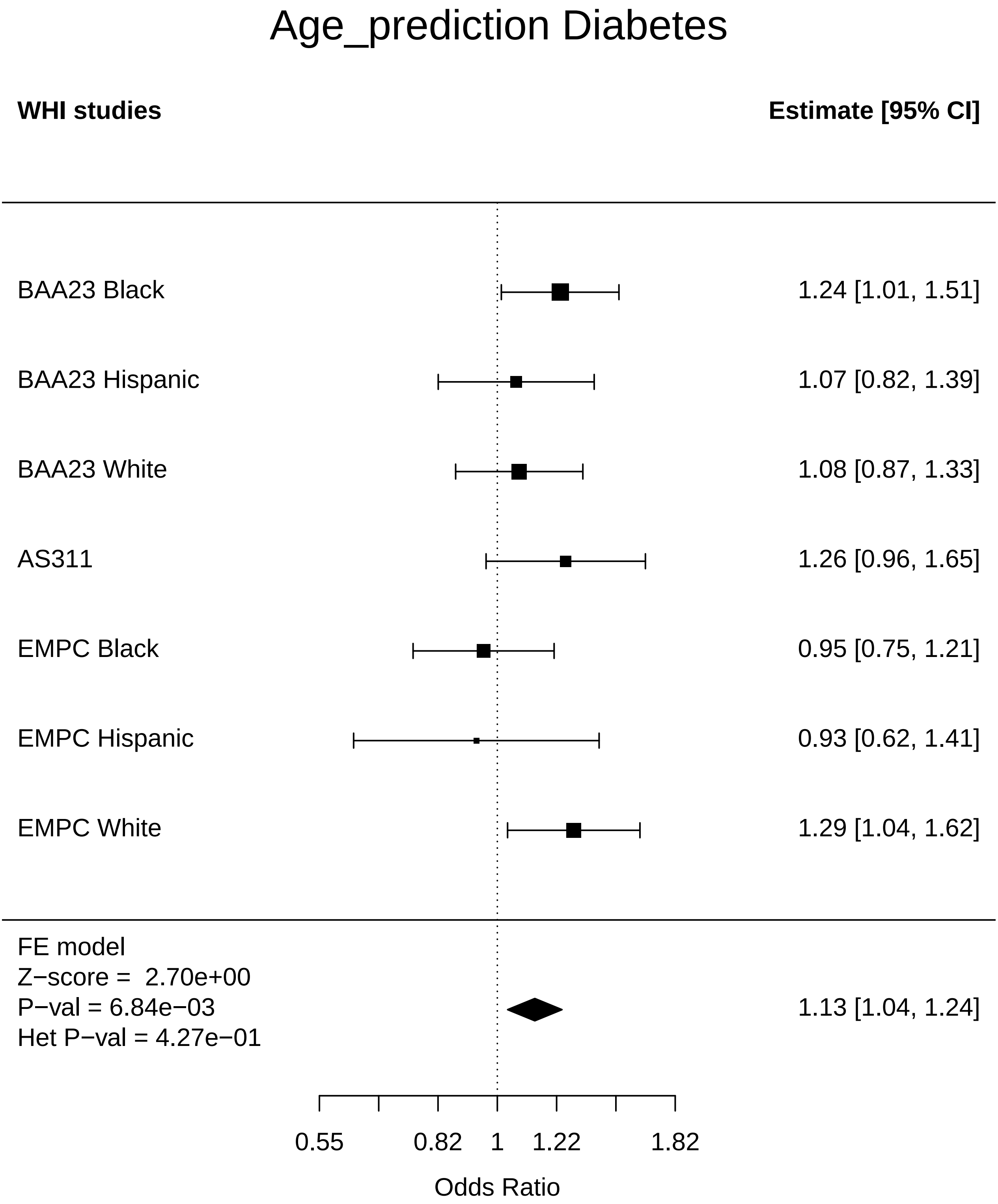

### Age_prediction_Endometrial cancer.pdf

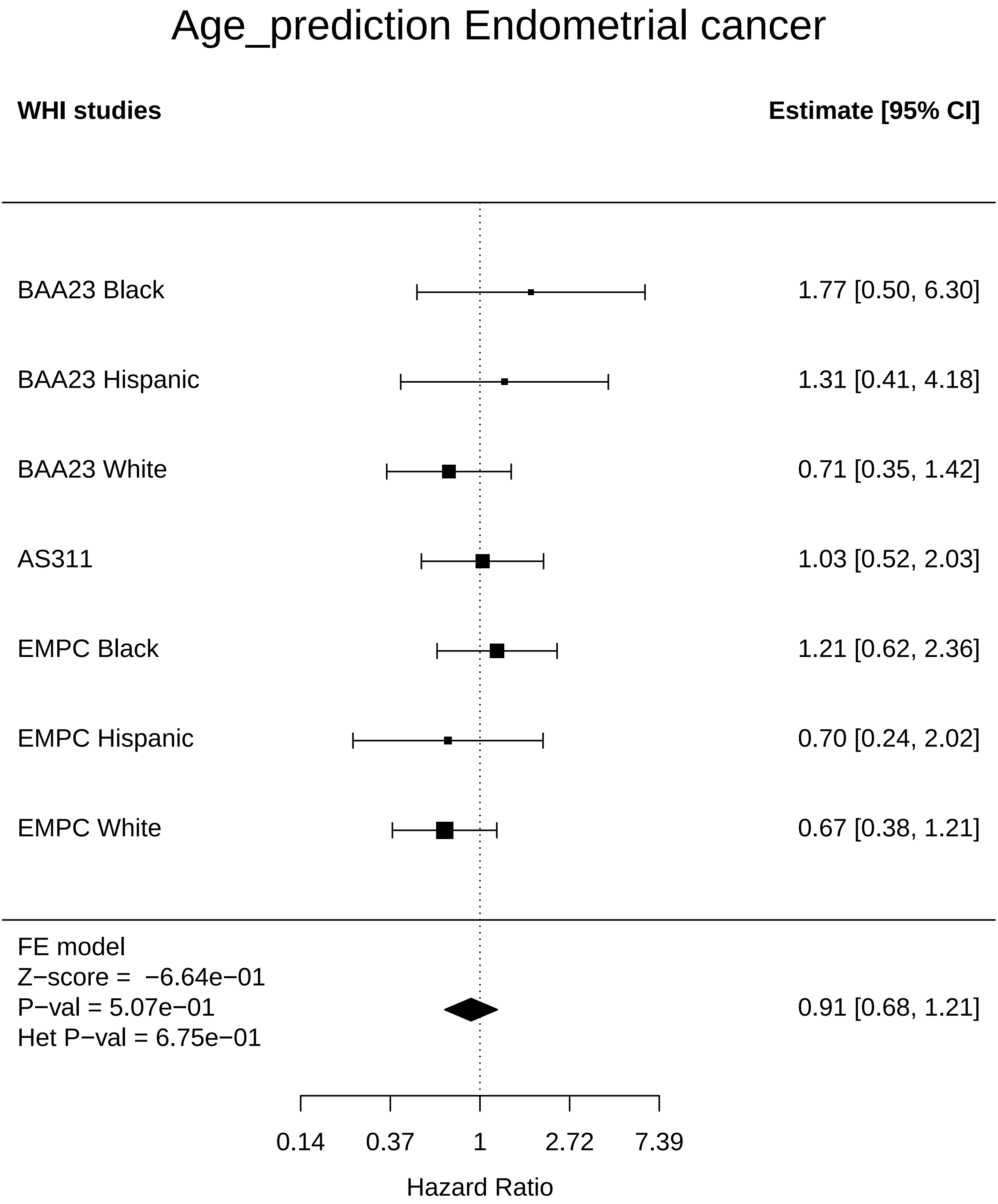

### Age_prediction_Leukemia.pdf

#### Age_prediction Leukemia

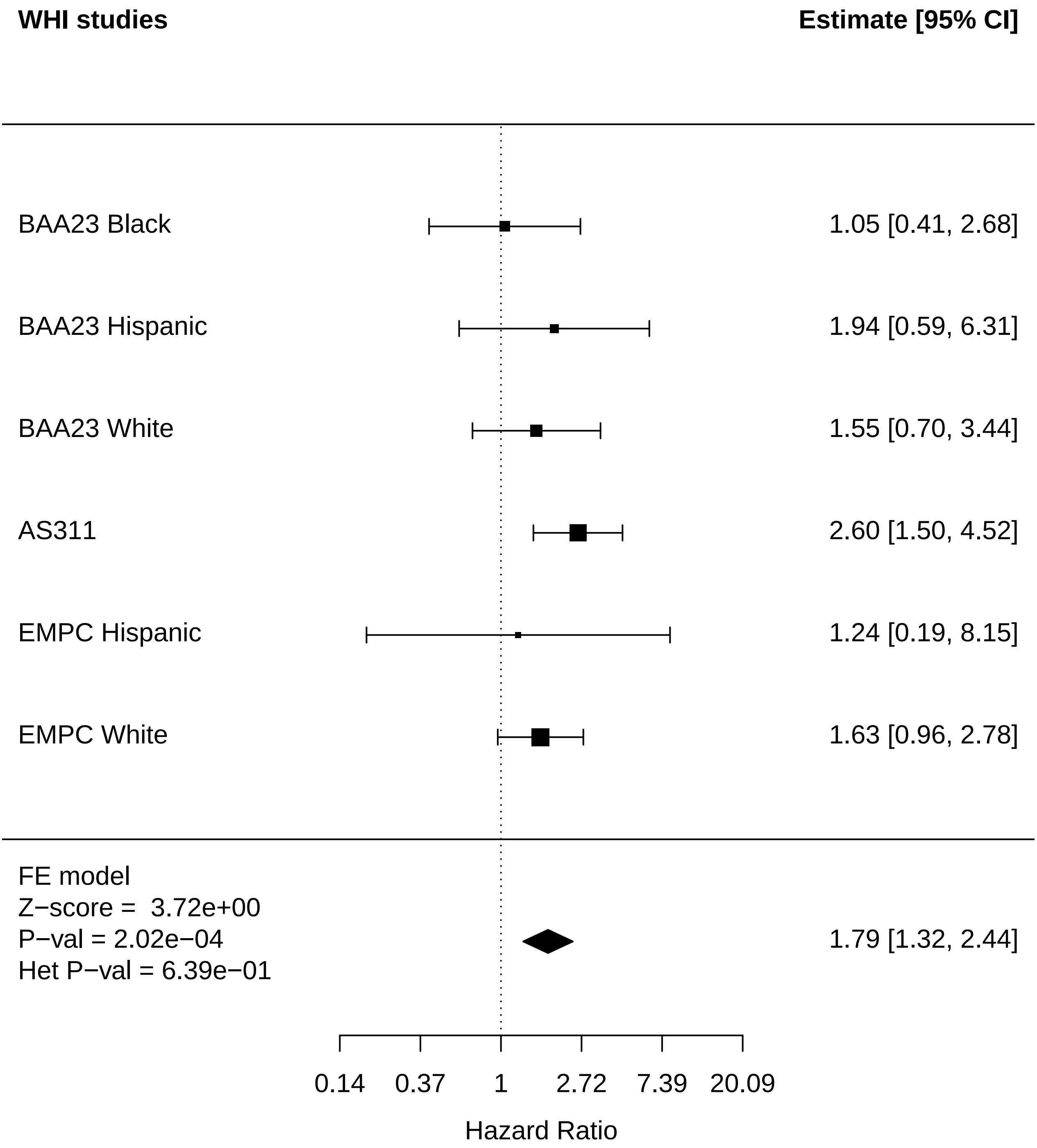

### Age_prediction_Lung Cancer.pdf

#### Age_prediction Lung Cancer

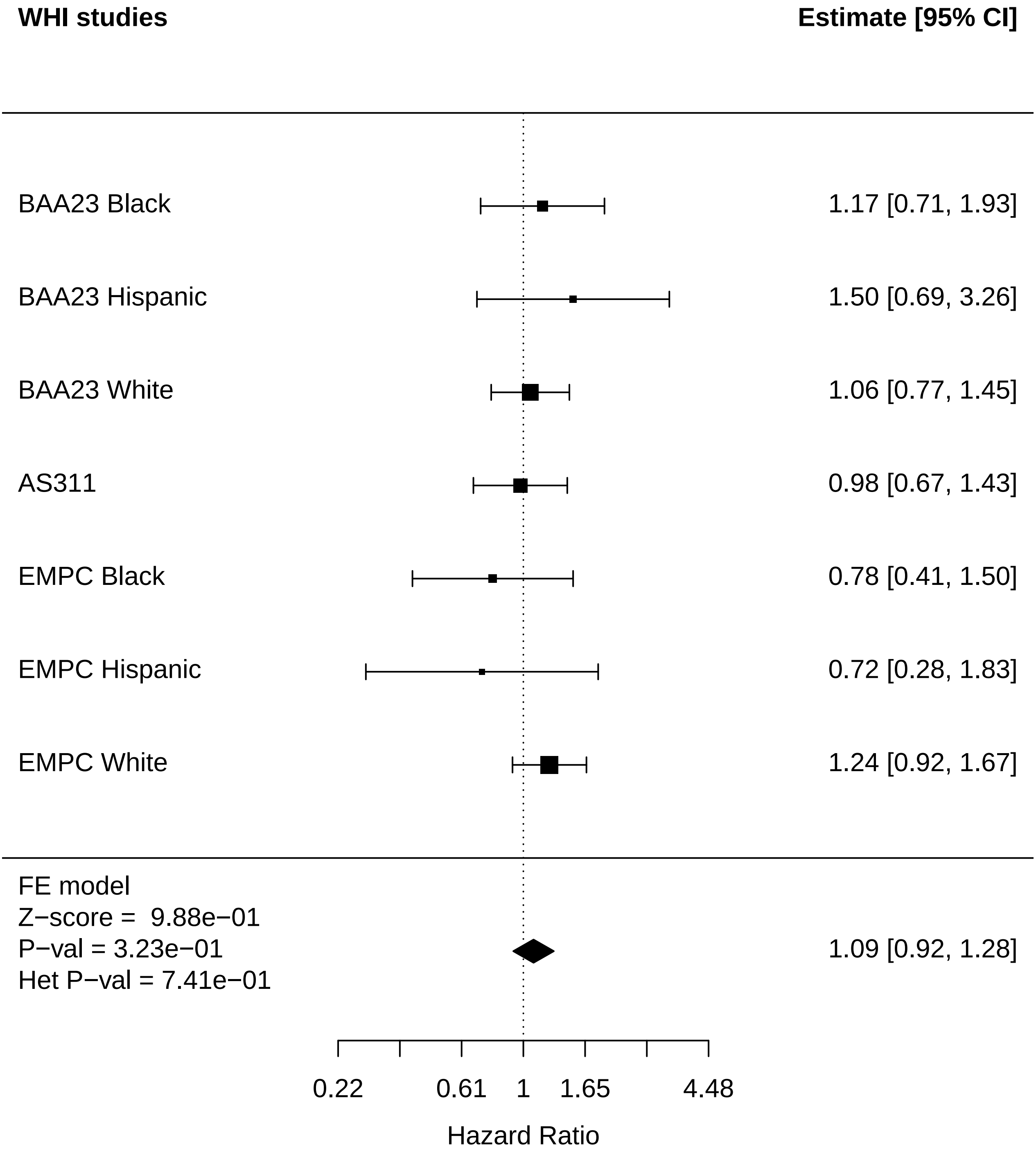

### Age_prediction_MI.pdf

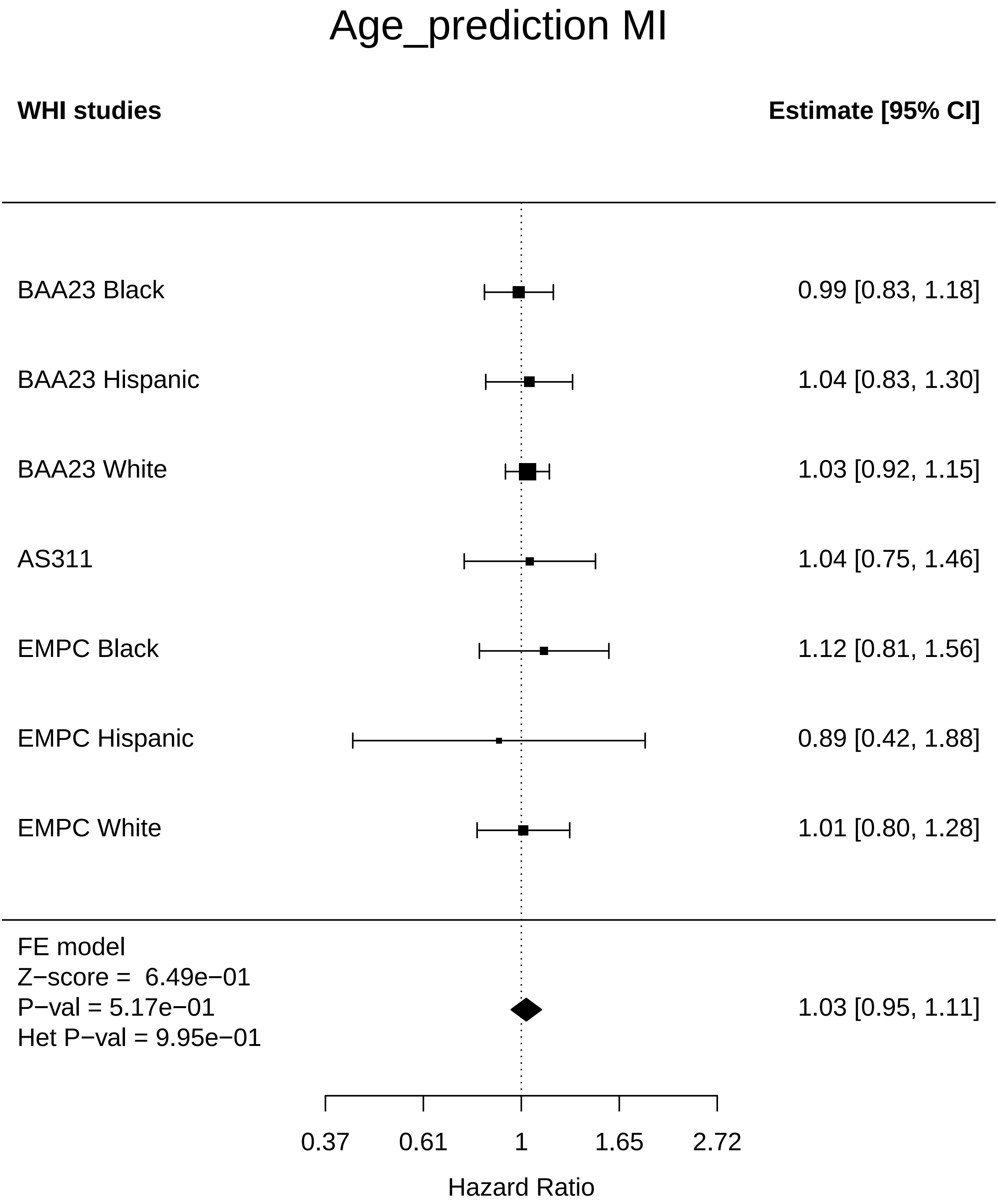

### Age_prediction_Mortality.pdf

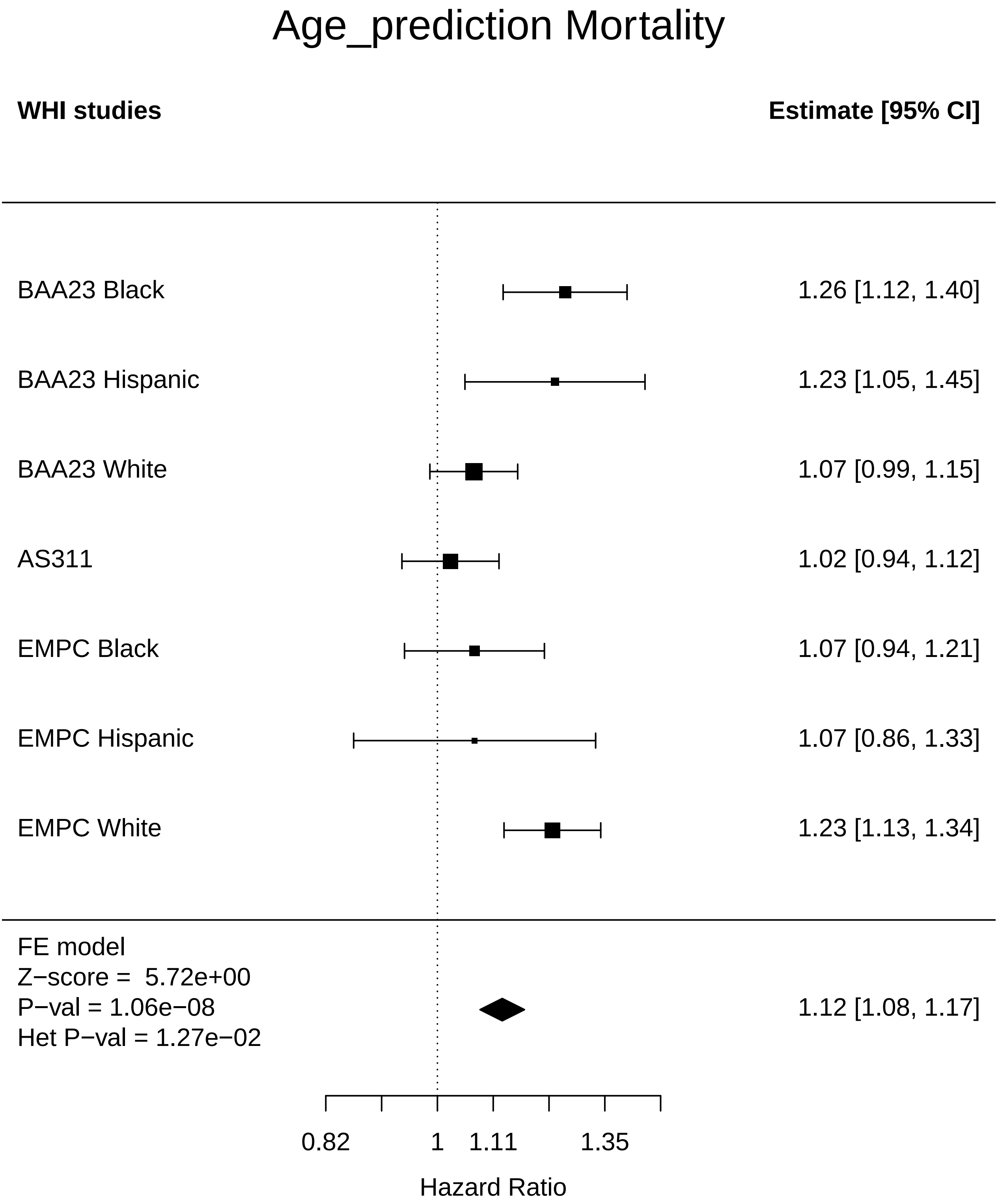

### Age_prediction_Physical function.pdf

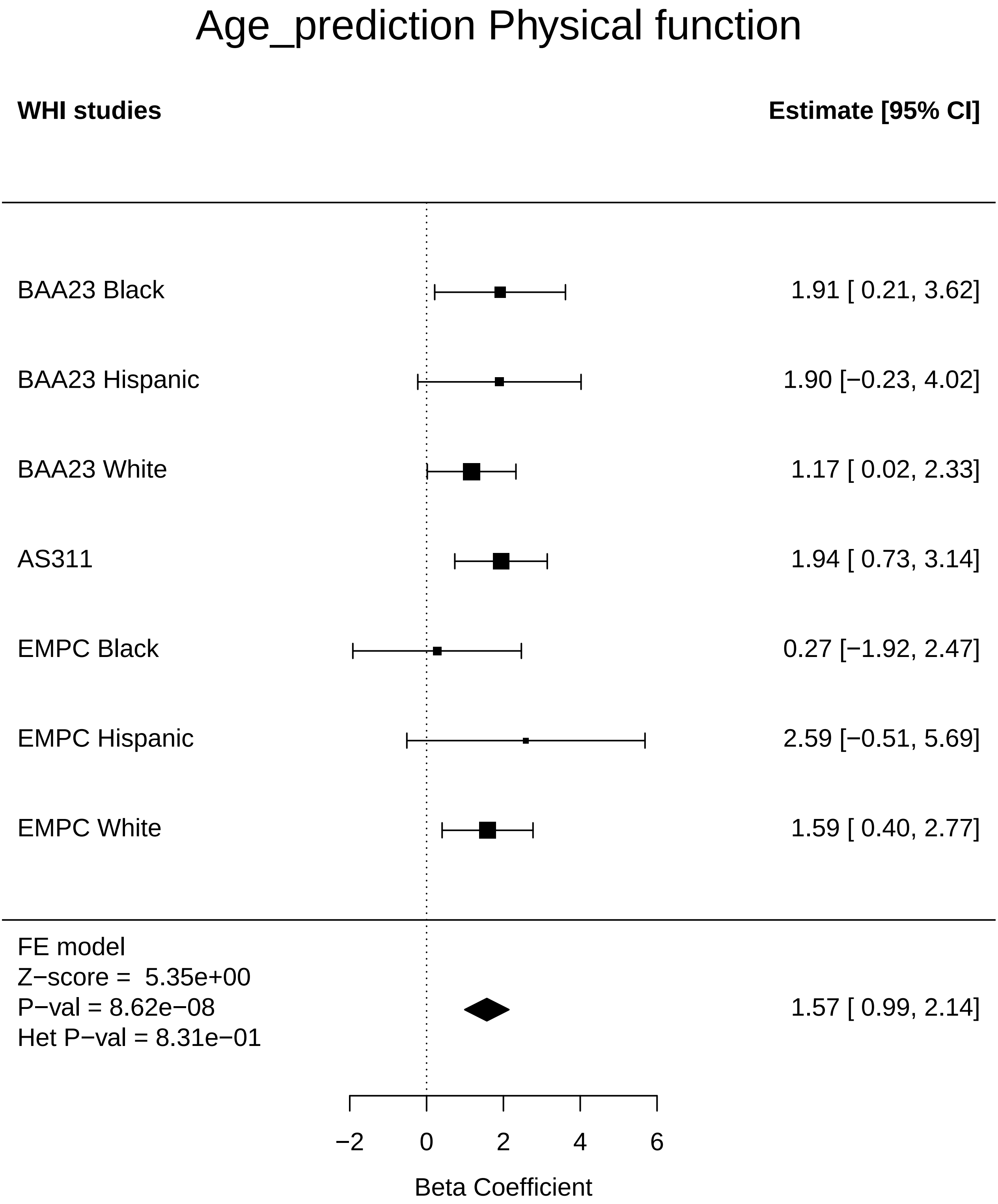

### Age_prediction_Stroke.pdf

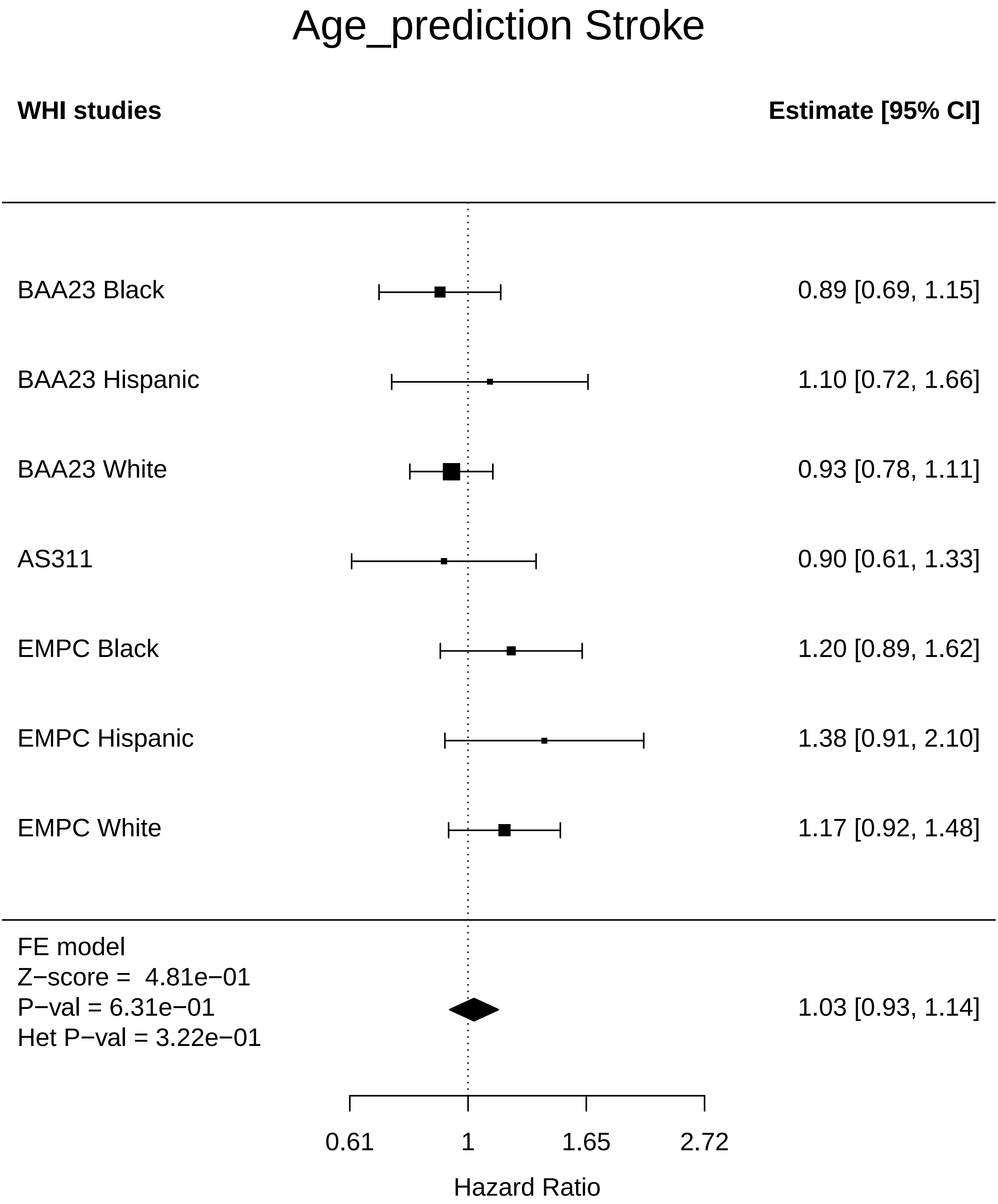

### Age_prediction_Thyroid disease.pdf

#### Age_prediction Thyroid disease

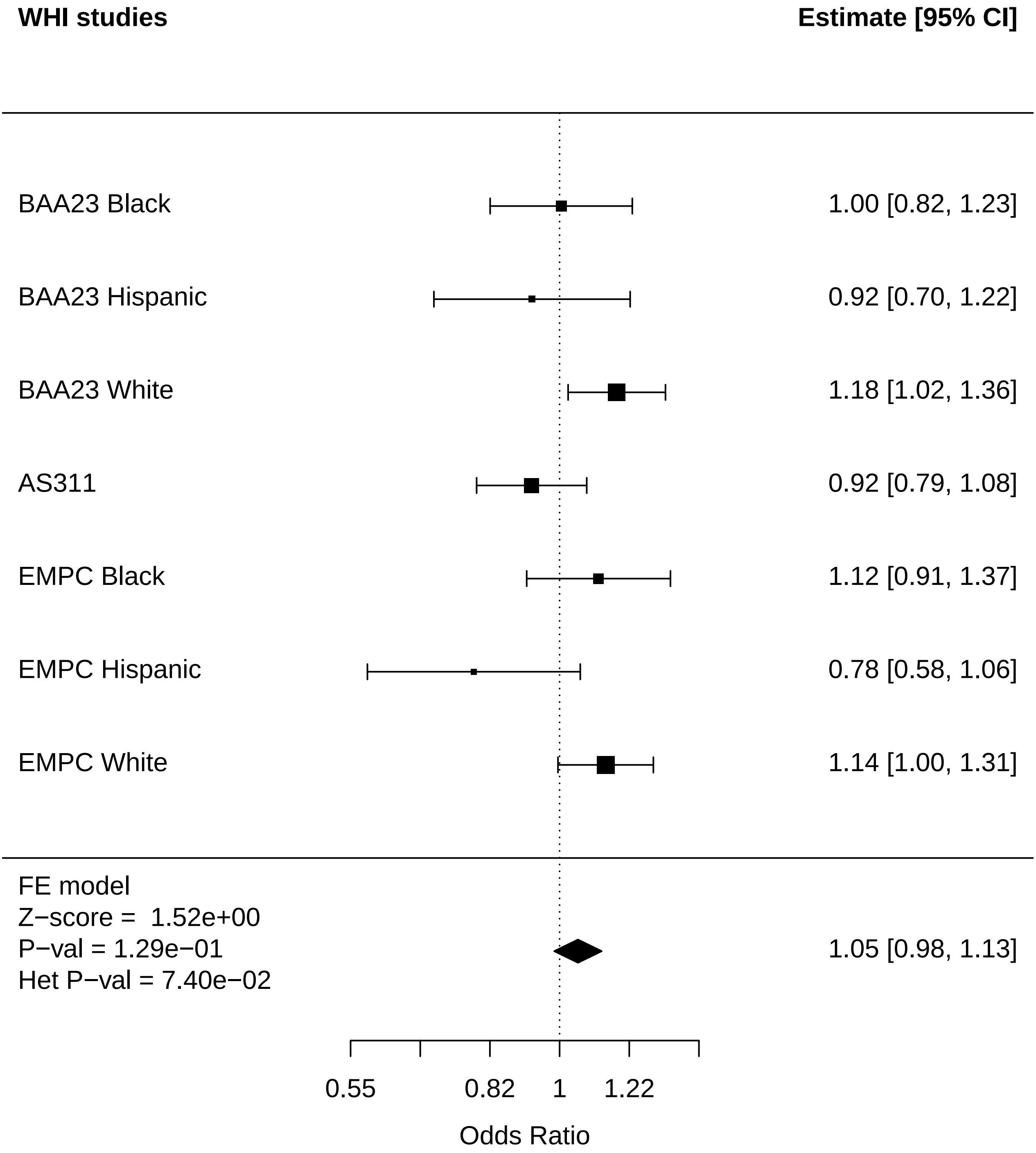

### Age_prediction_Total comorbidities.pdf

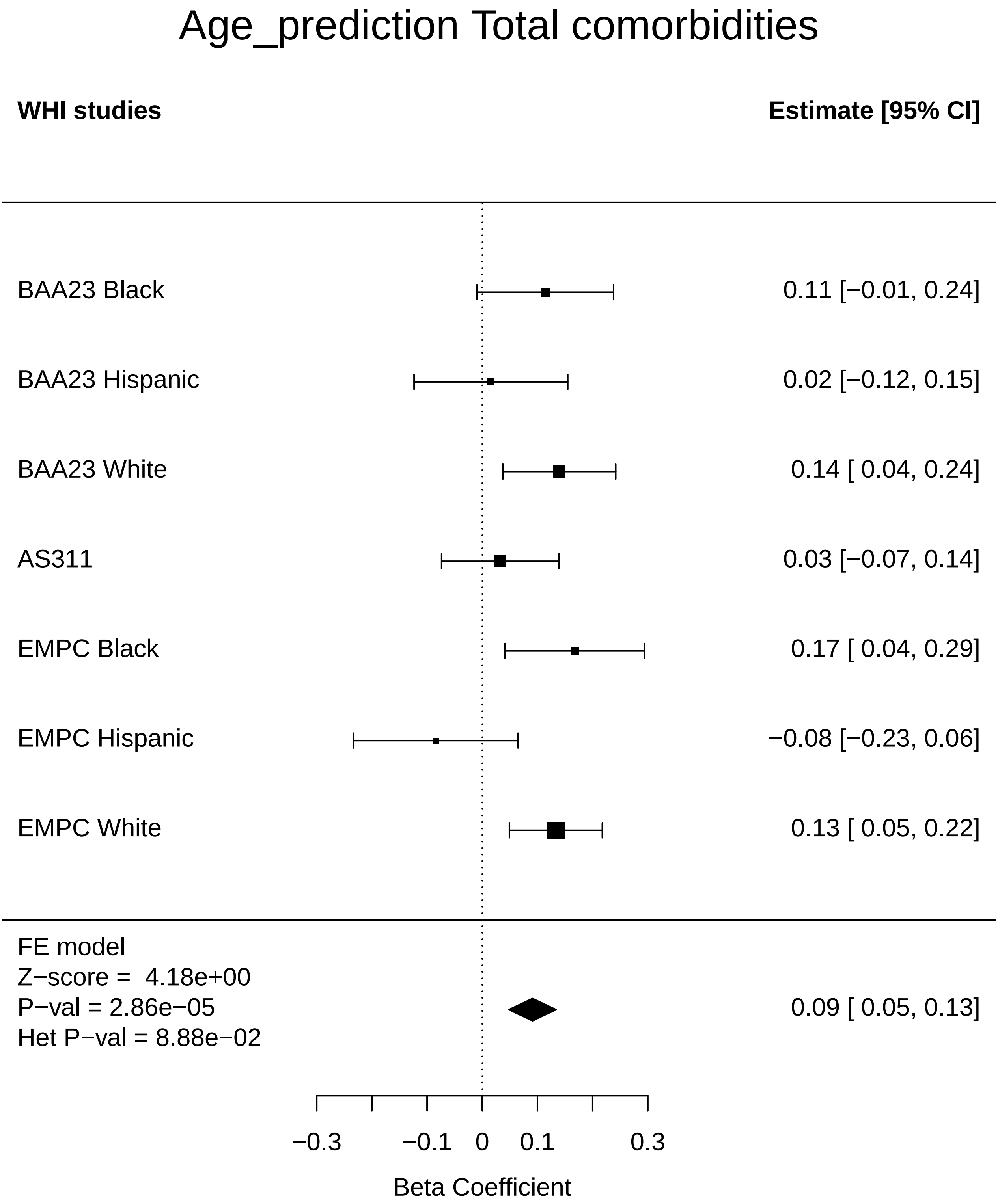

### DunedinPACE_Any Cancer.pdf

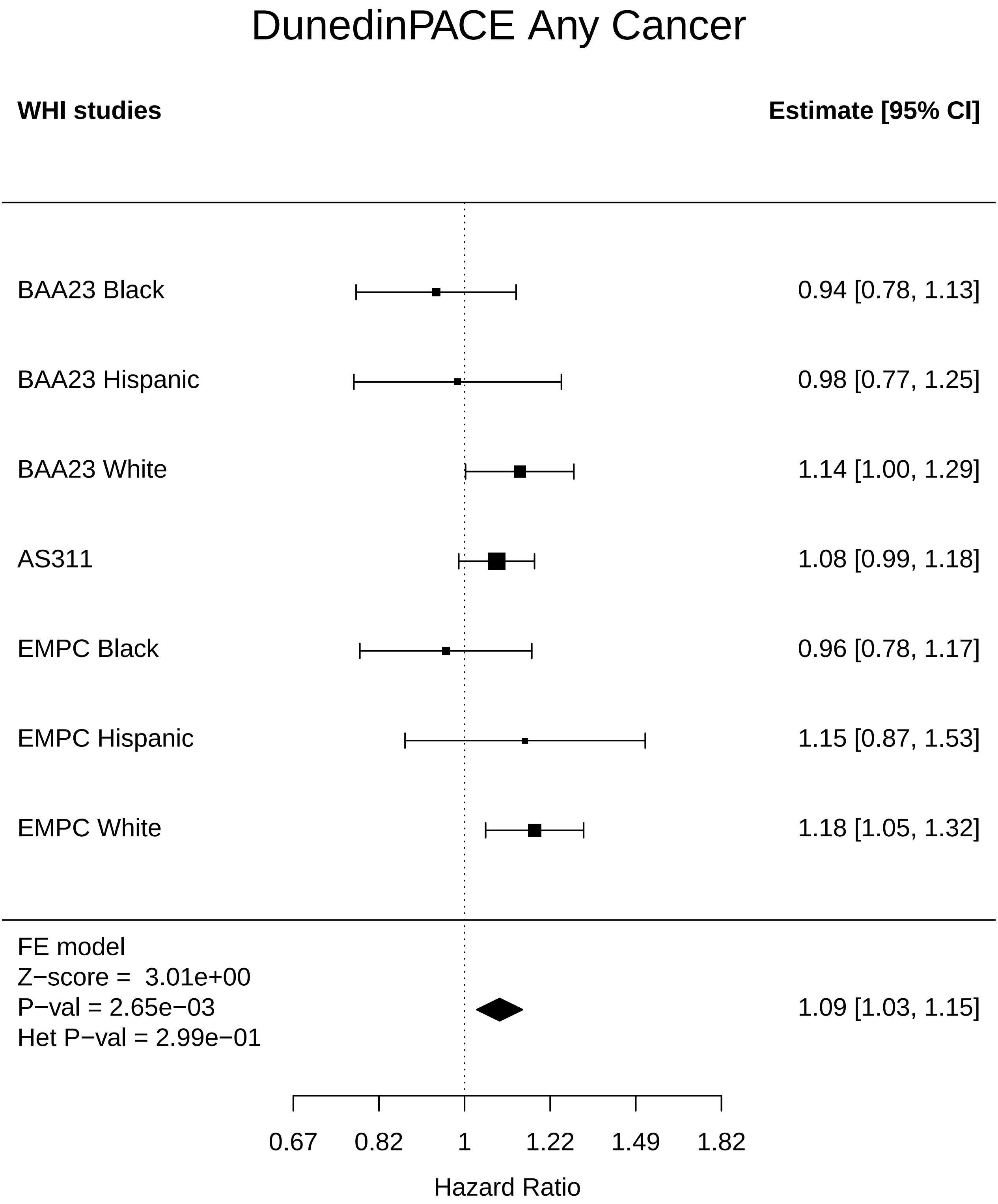

### DunedinPACE_Arthritis.pdf

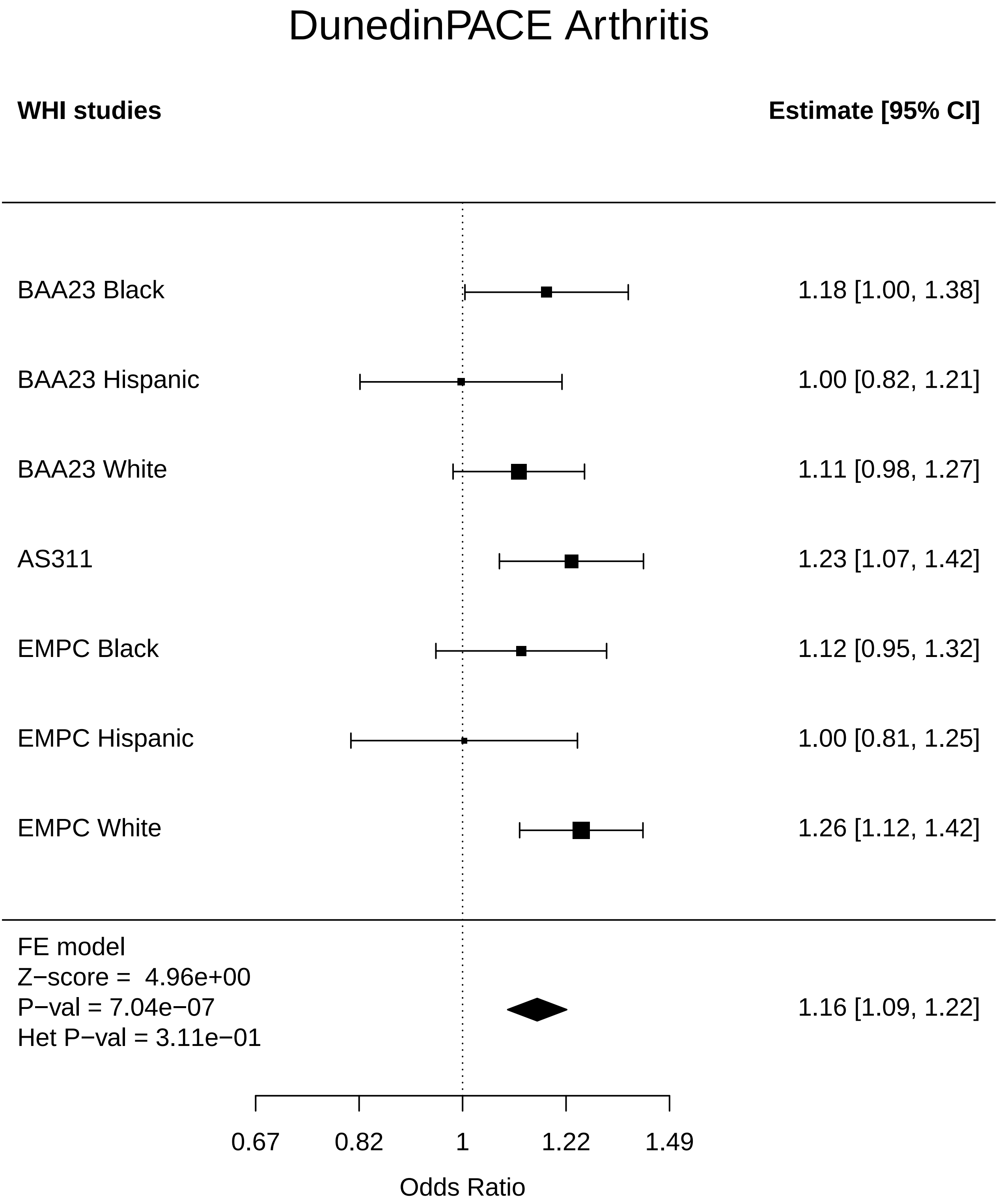

### DunedinPACE_Breast Cancer.pdf

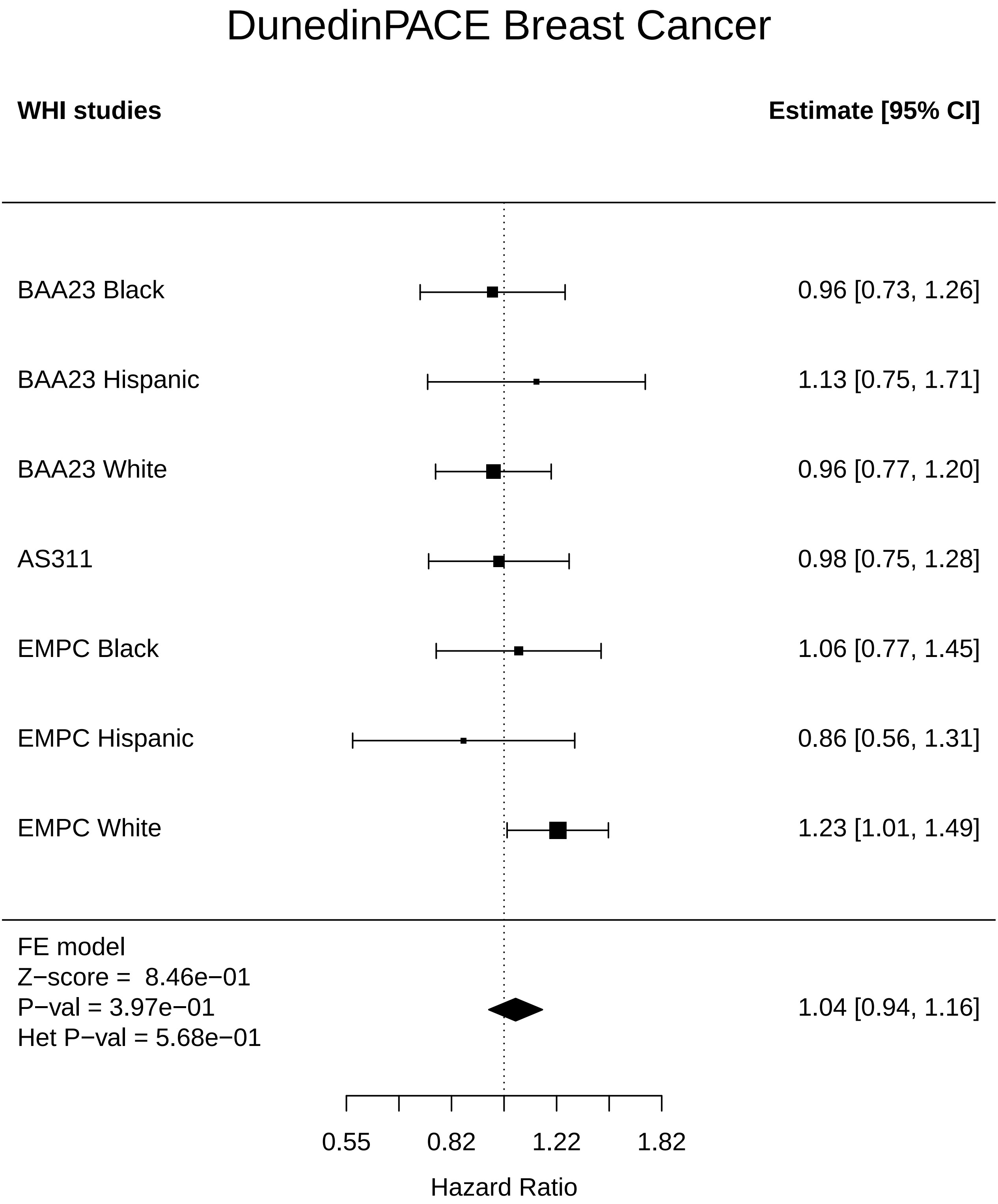

### DunedinPACE_Cataract.pdf

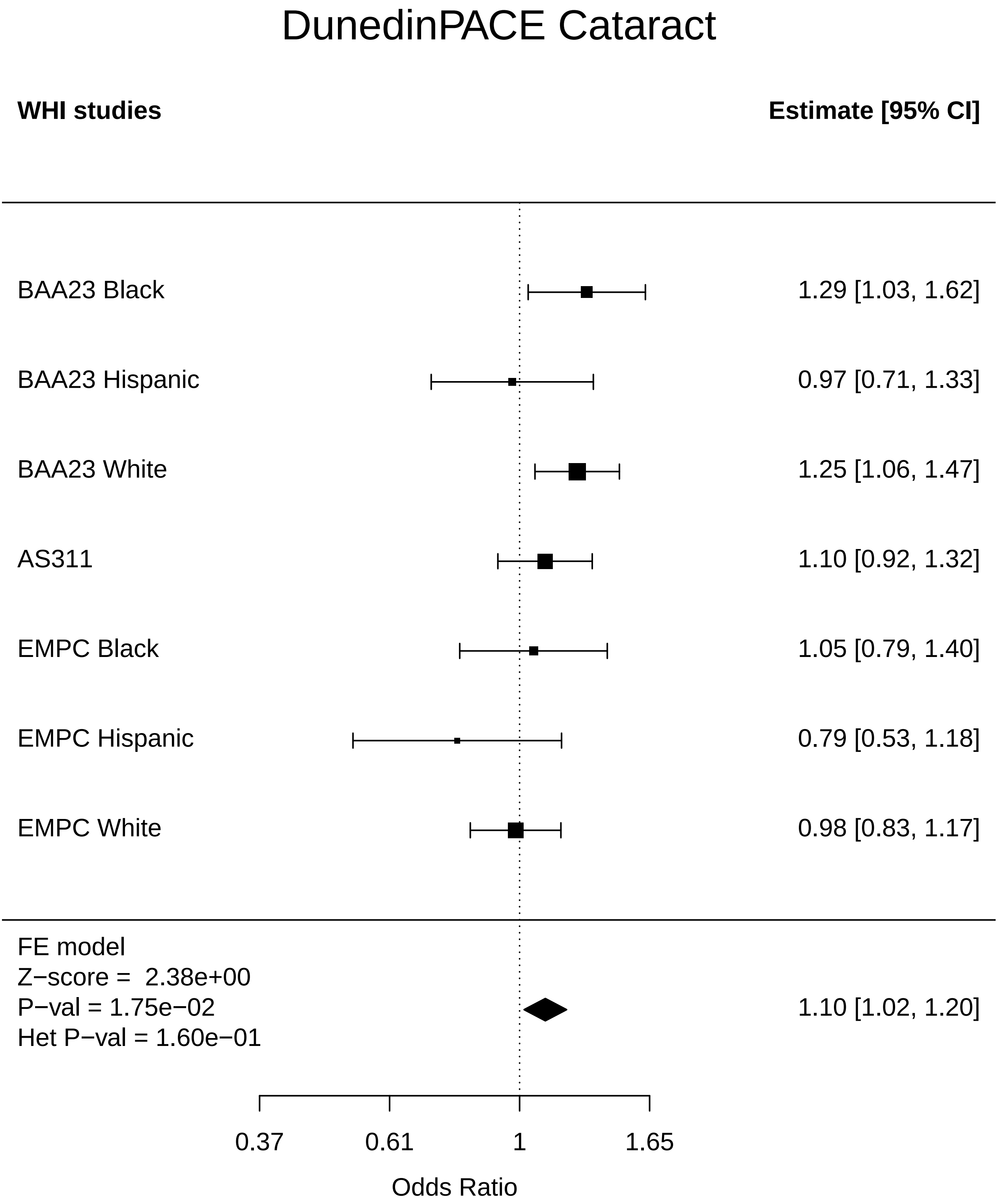

### DunedinPACE_CHD.pdf

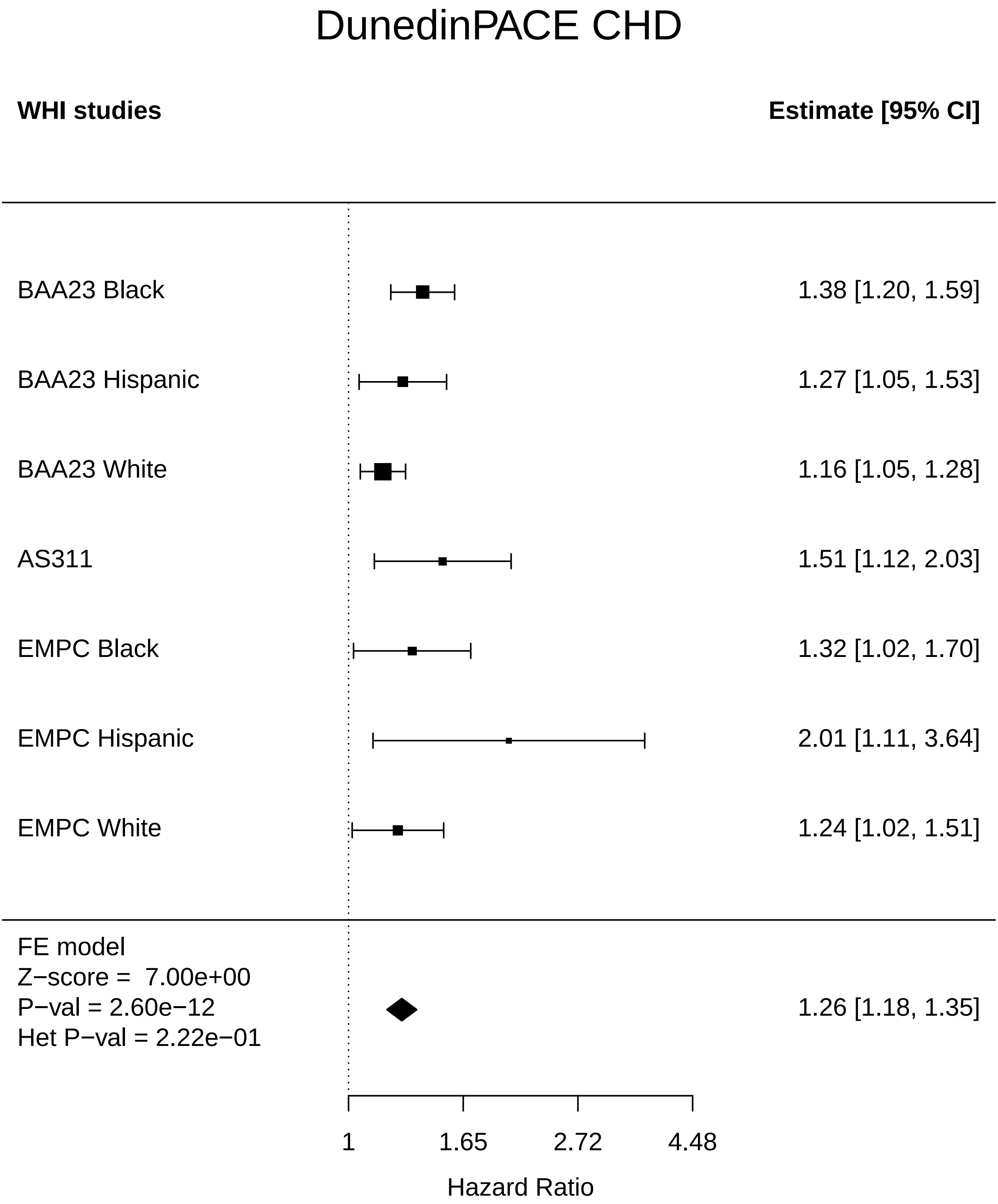

### DunedinPACE_Cognitive function.pdf

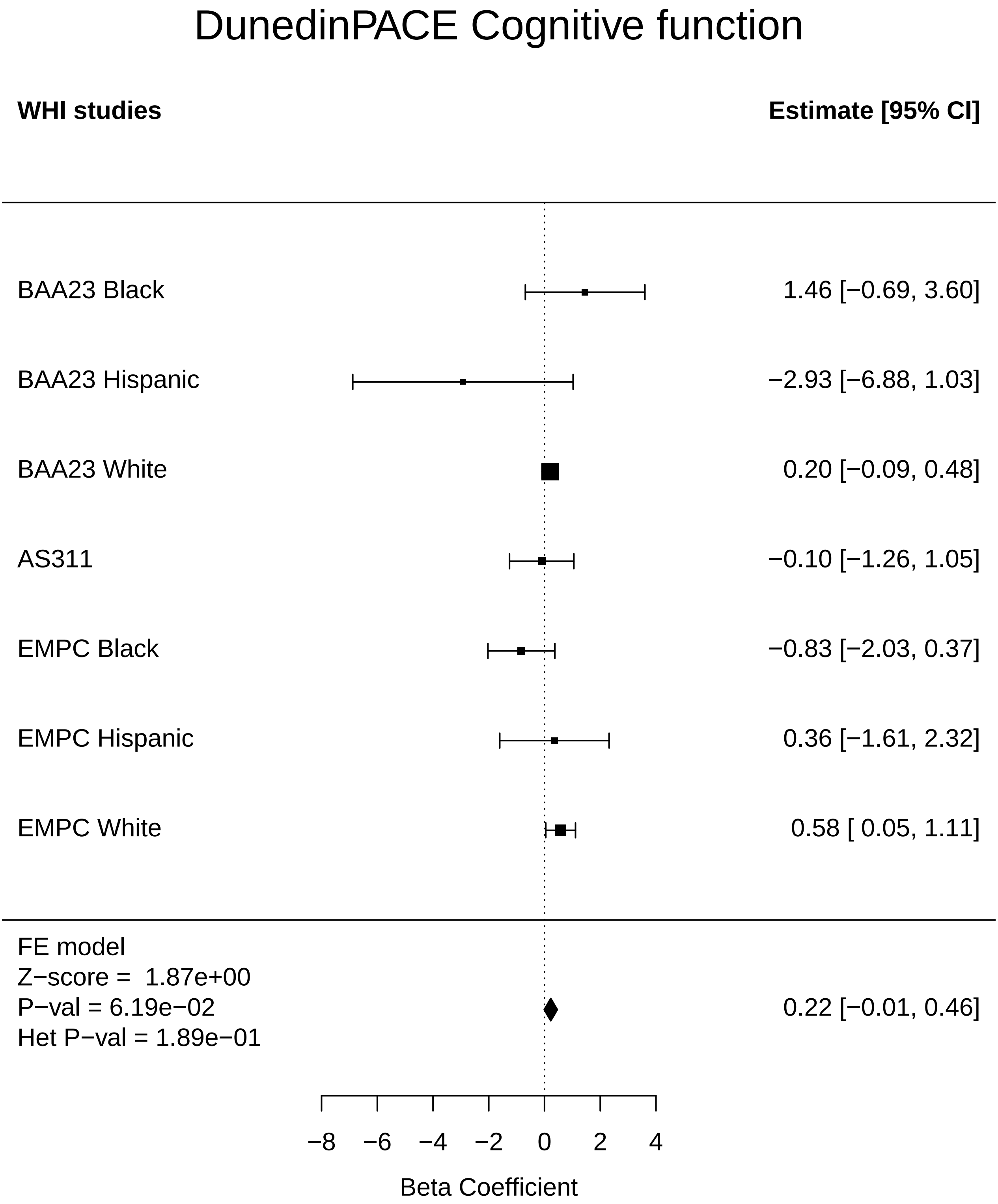

### DunedinPACE_Colorectal cancer.pdf

#### DunedinPACE Colorectal cancer

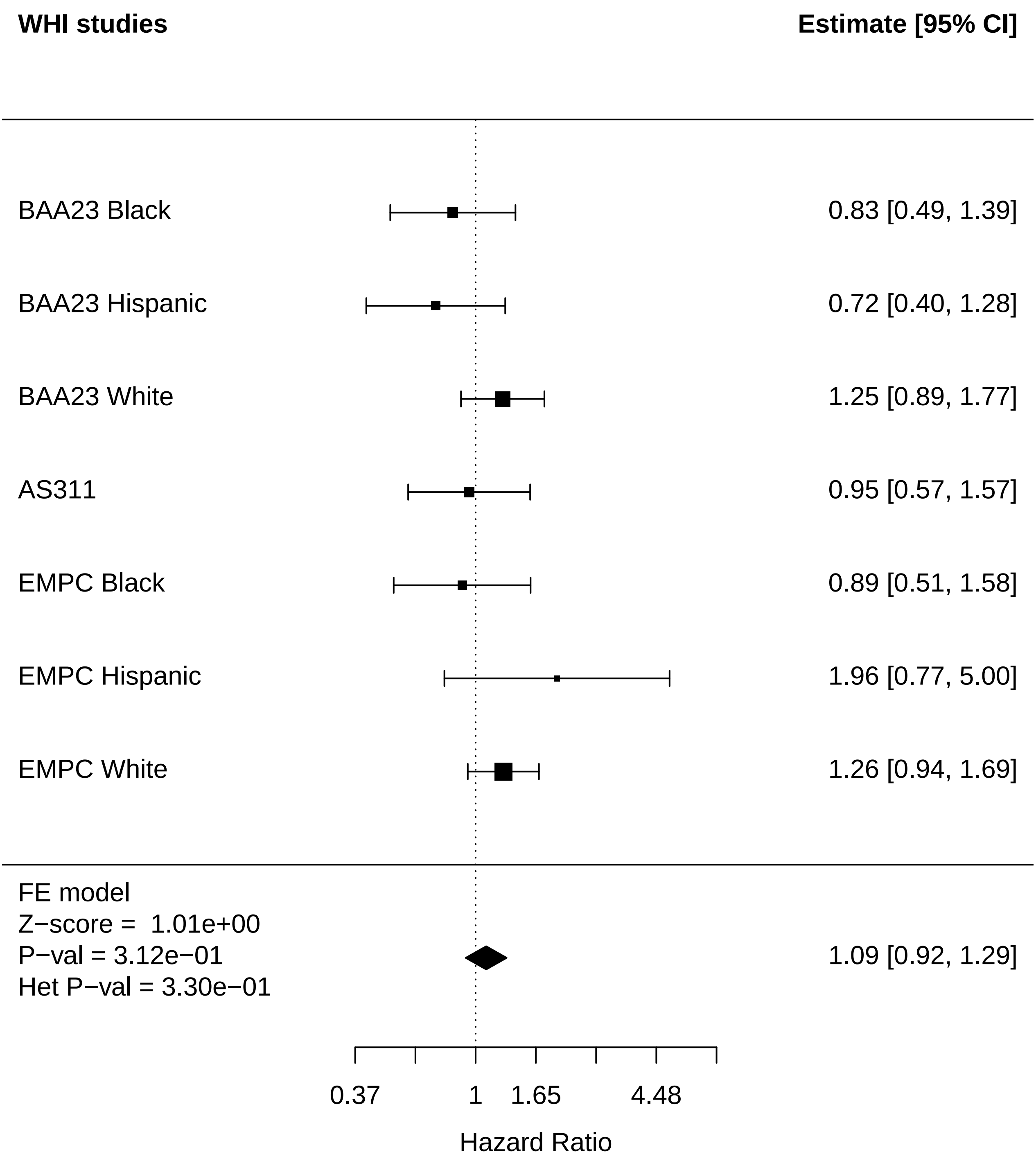

### DunedinPACE_Diabetes.pdf

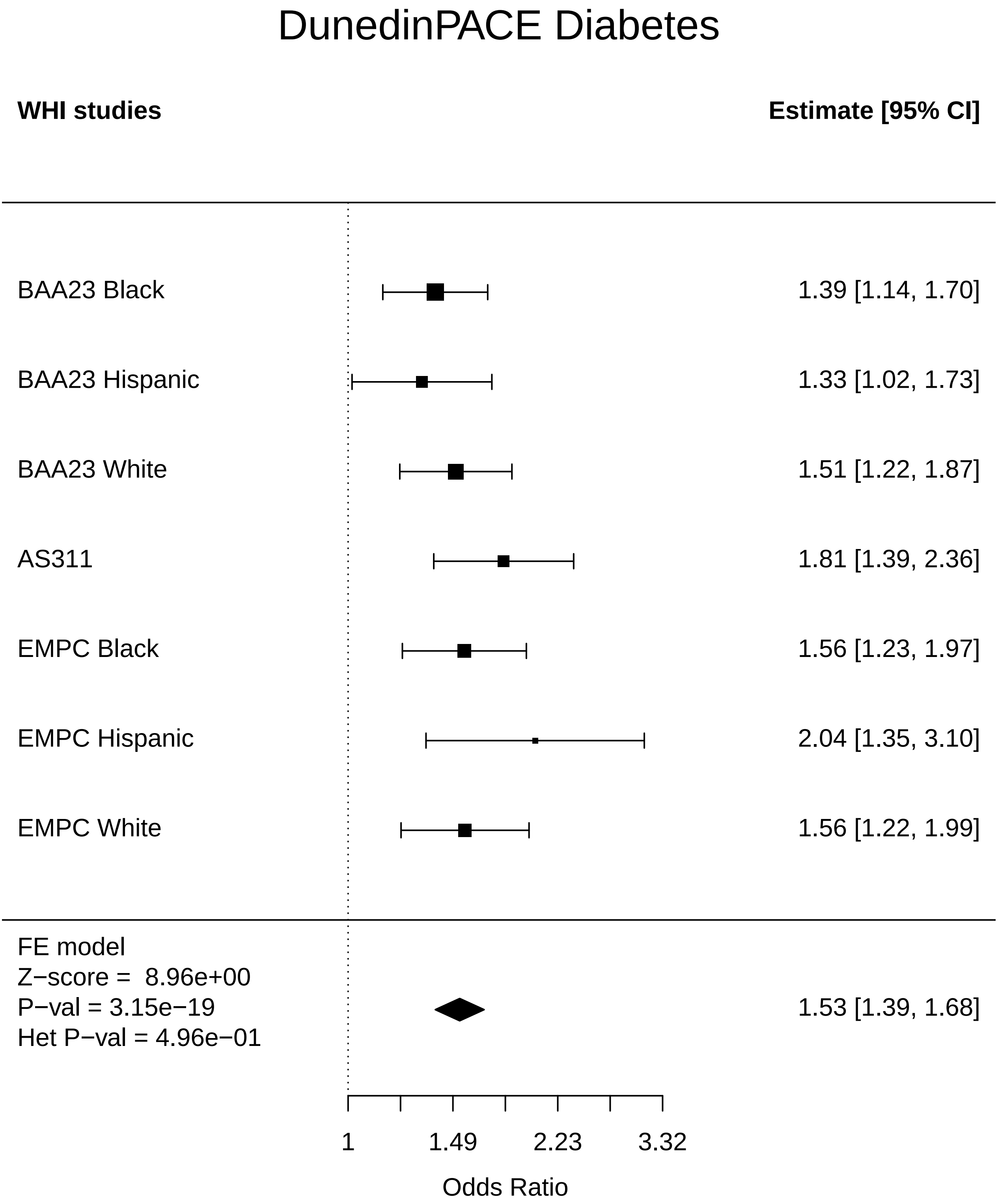

### DunedinPACE_Disease free.pdf

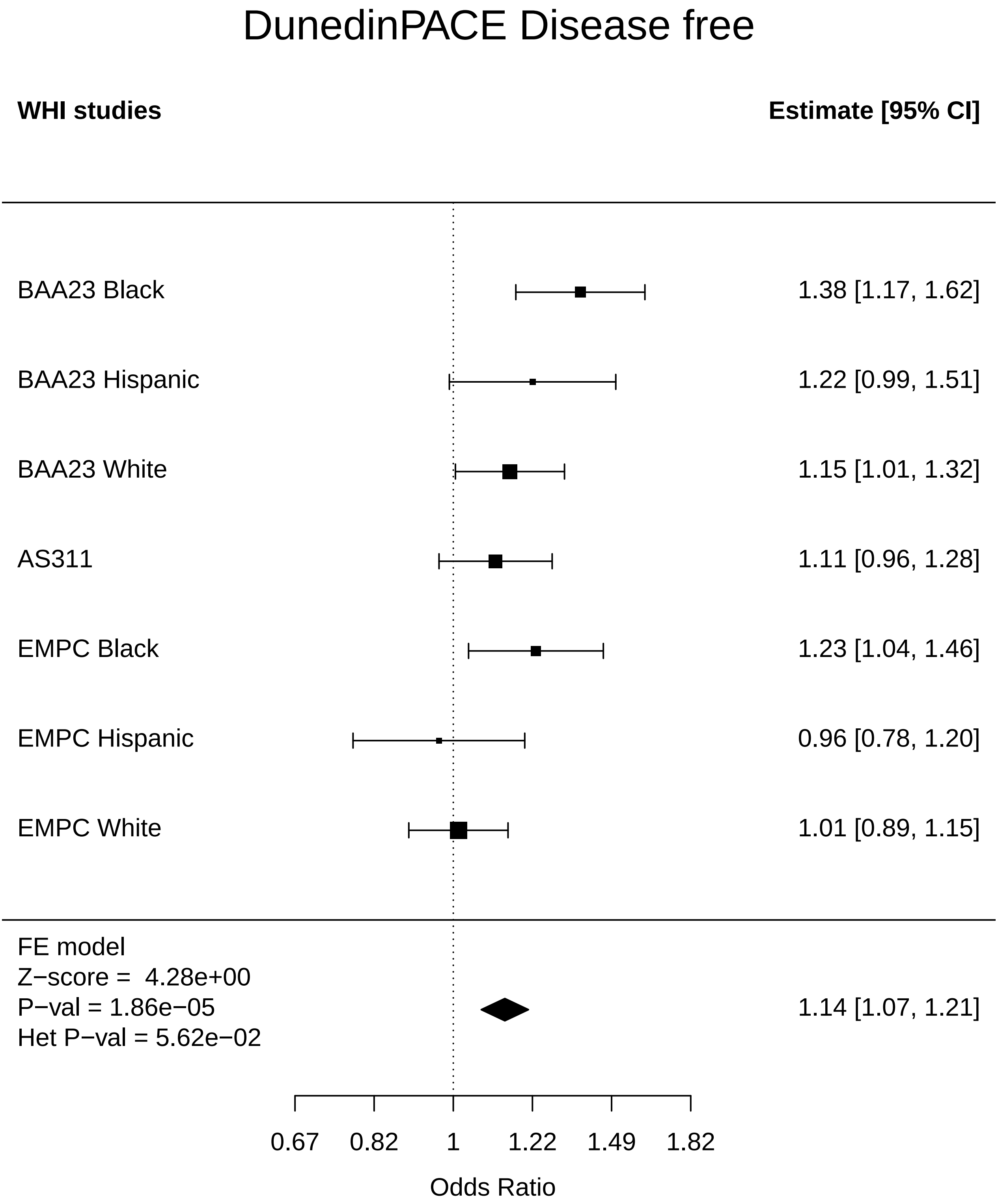

### DunedinPACE_Endometrial cancer.pdf

#### DunedinPACE Endometrial cancer

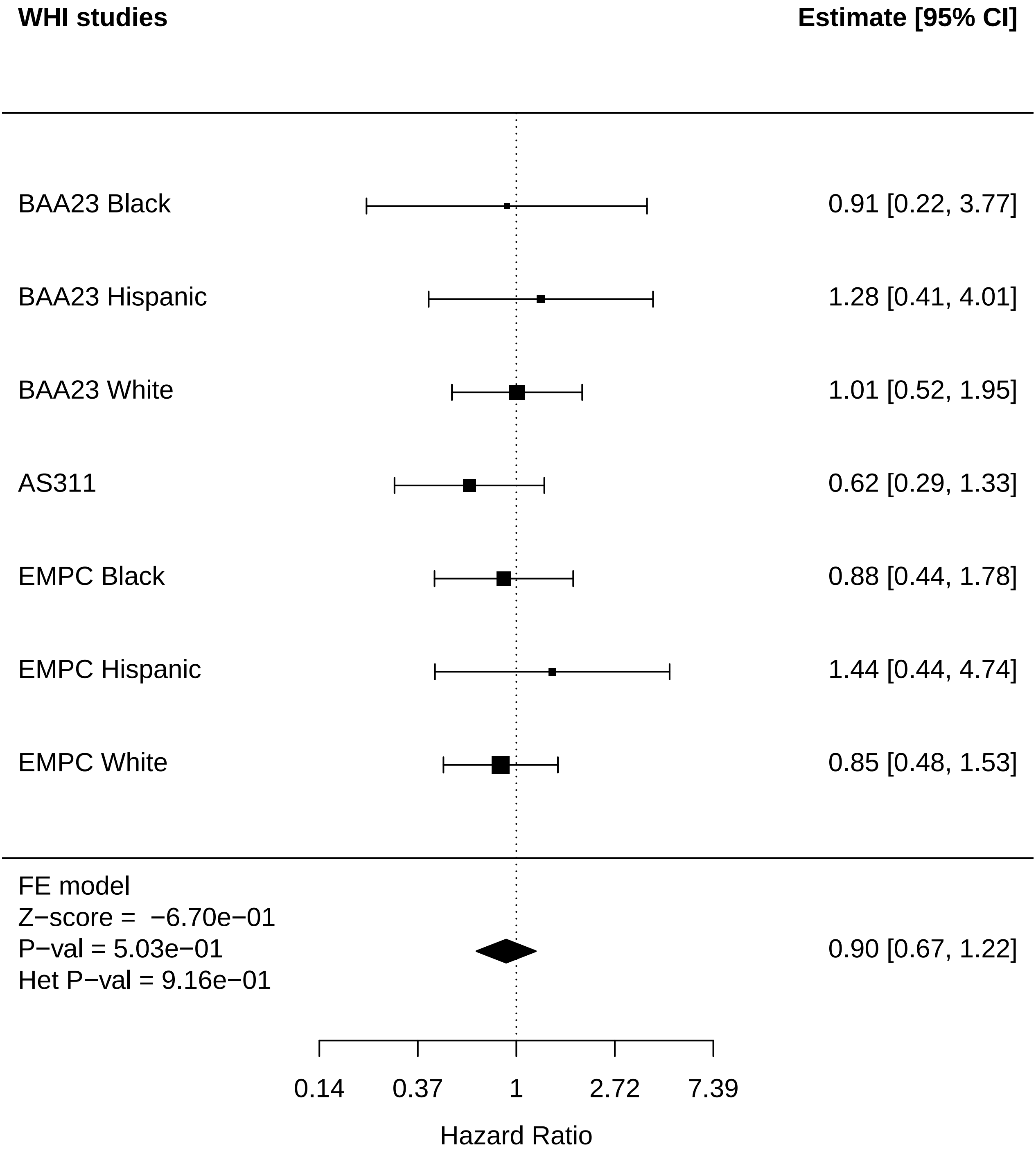

### DunedinPACE_Leukemia.pdf

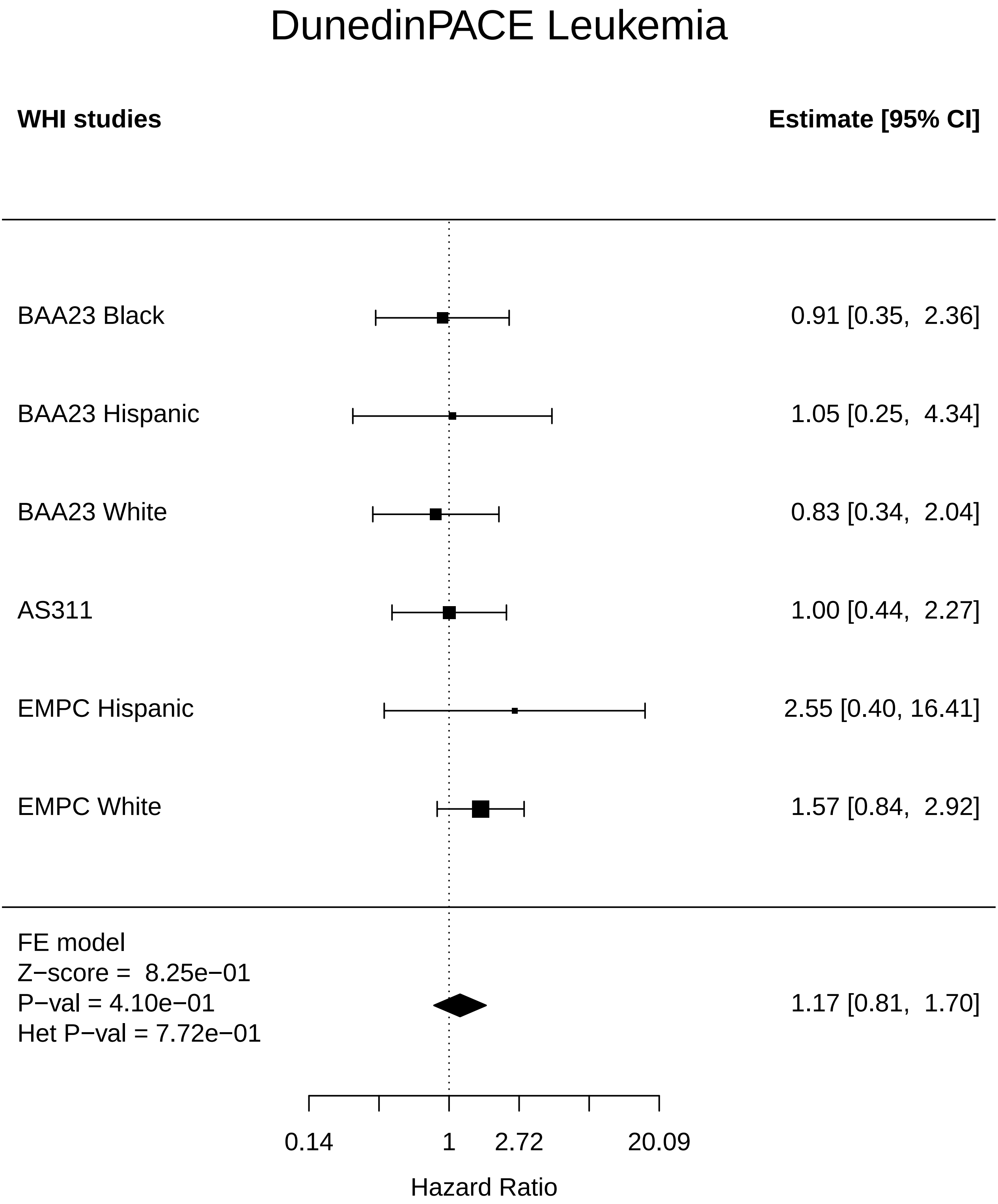

### DunedinPACE_Lung Cancer.pdf

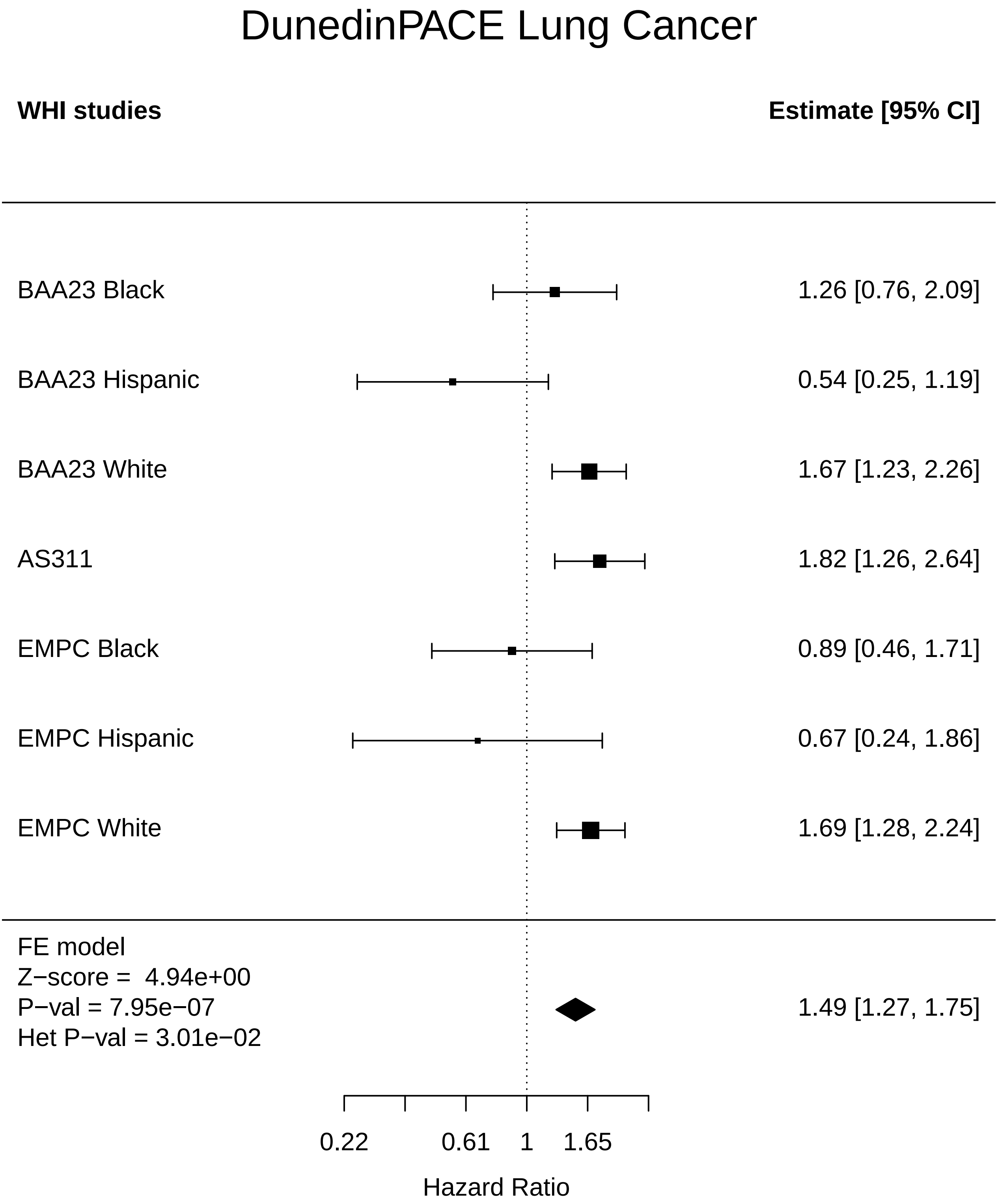

### DunedinPACE_MI.pdf

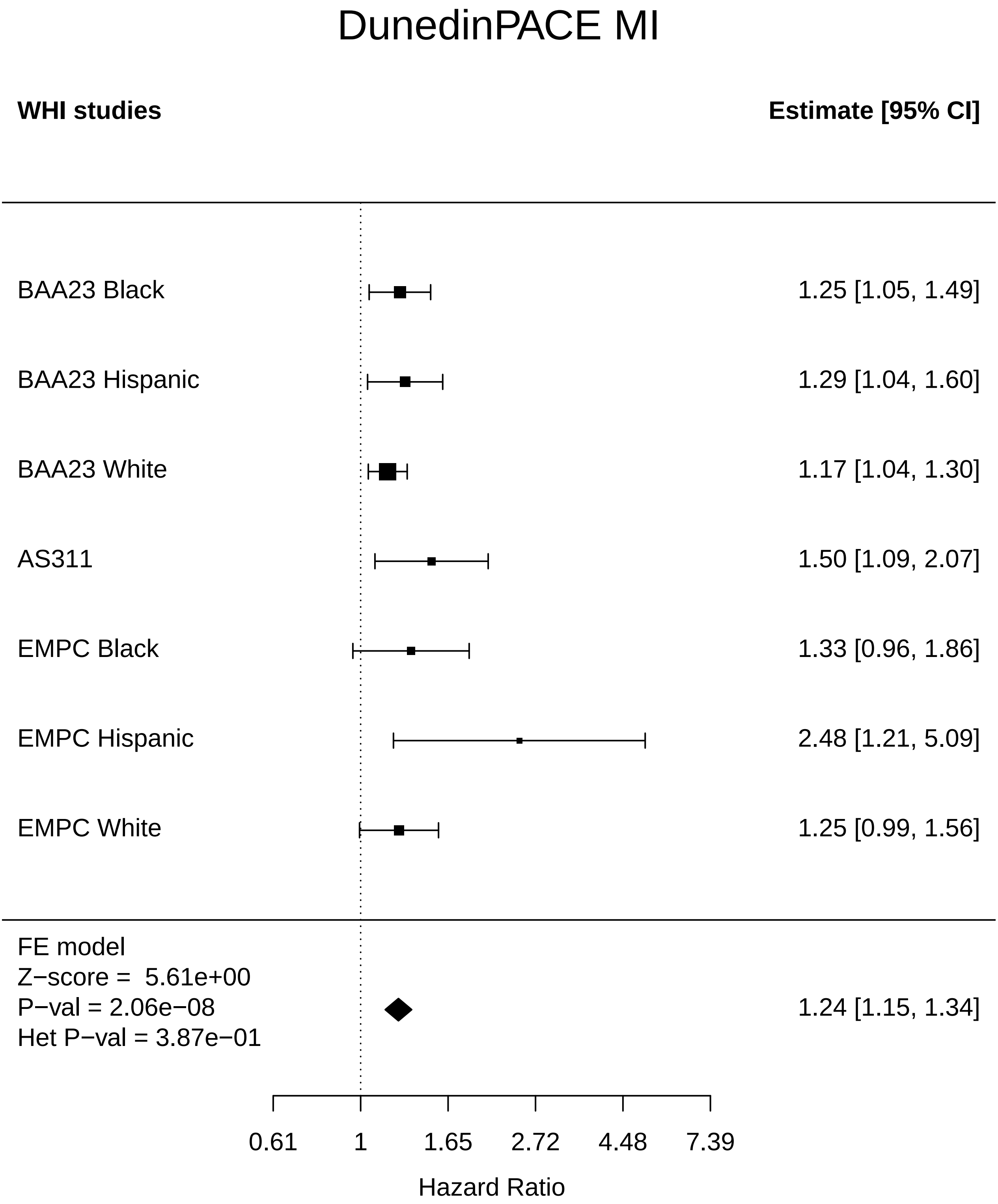

### DunedinPACE_Mortality.pdf

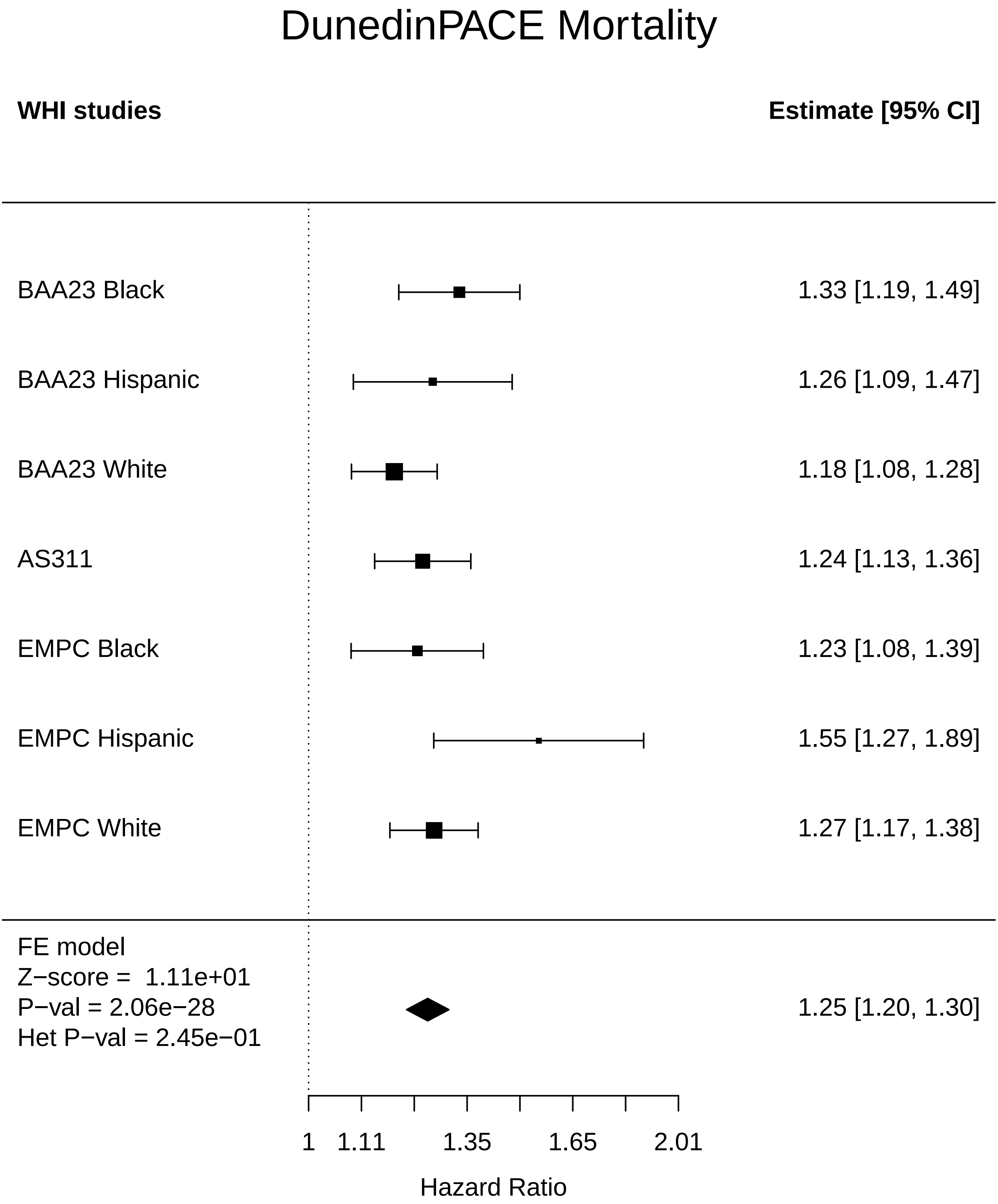

### DunedinPACE_Physical function.pdf

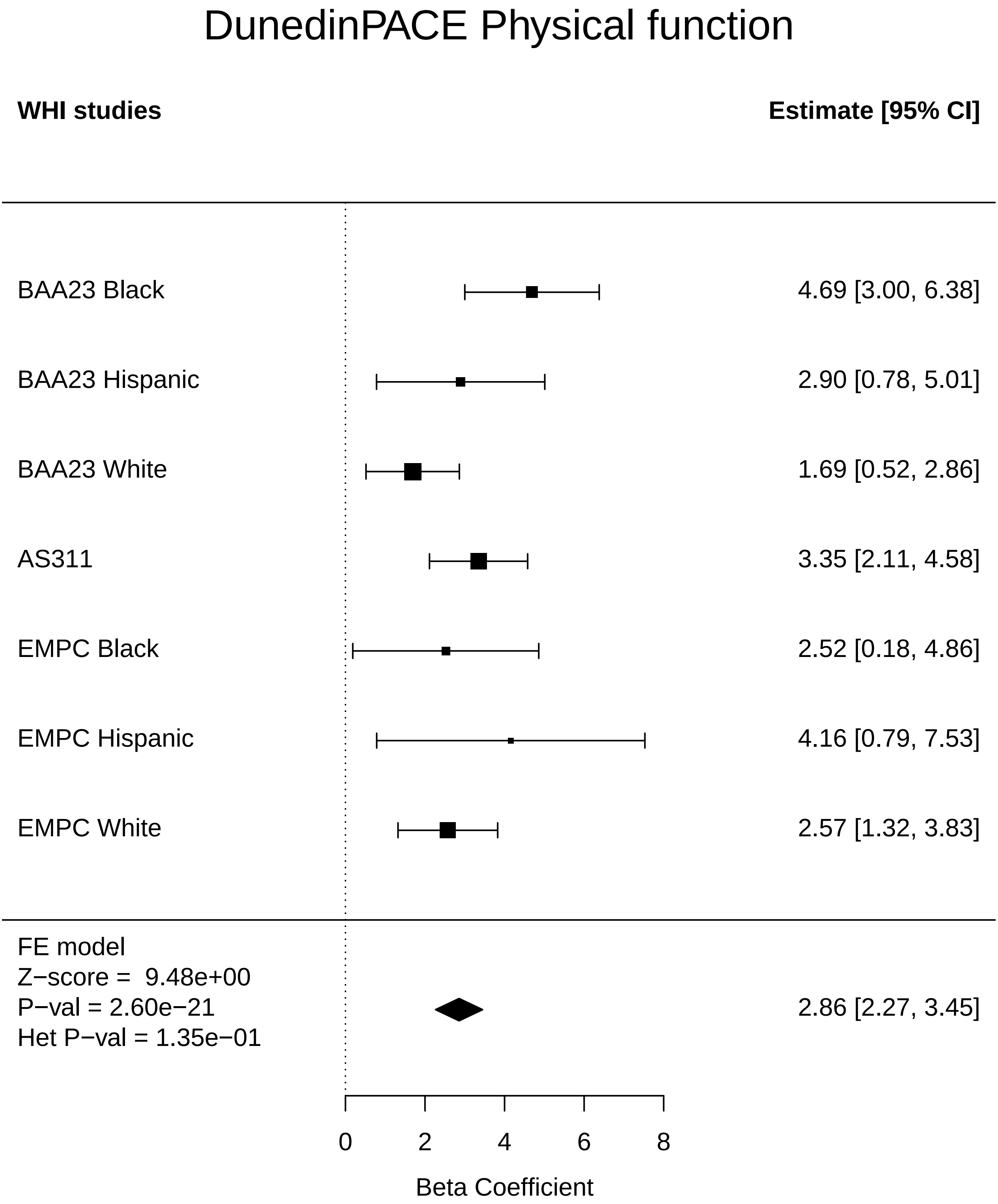

### DunedinPACE_Stroke.pdf

#### DunedinPACE Stroke

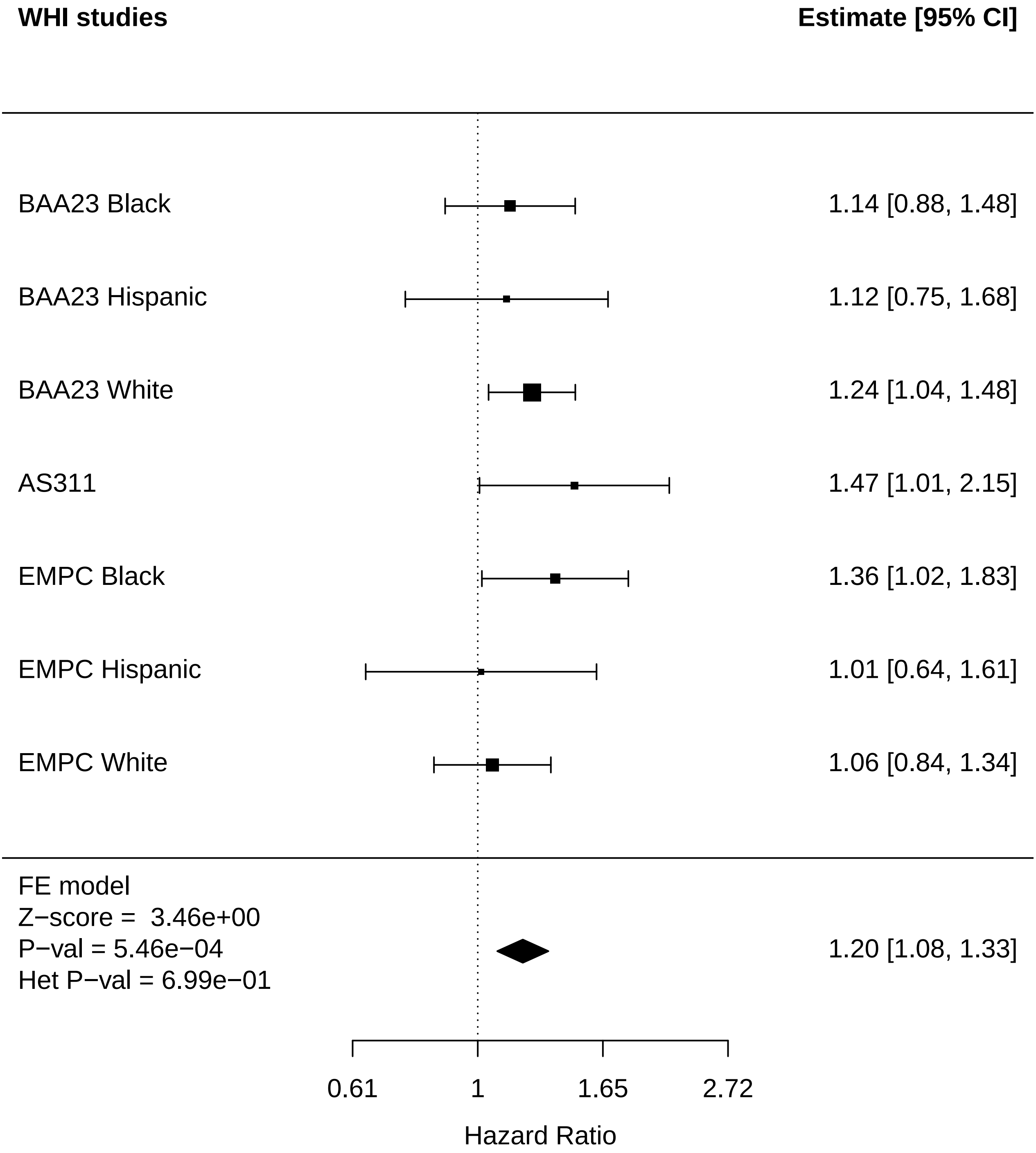

### DunedinPACE_Thyroid disease.pdf

#### DunedinPACE Thyroid disease

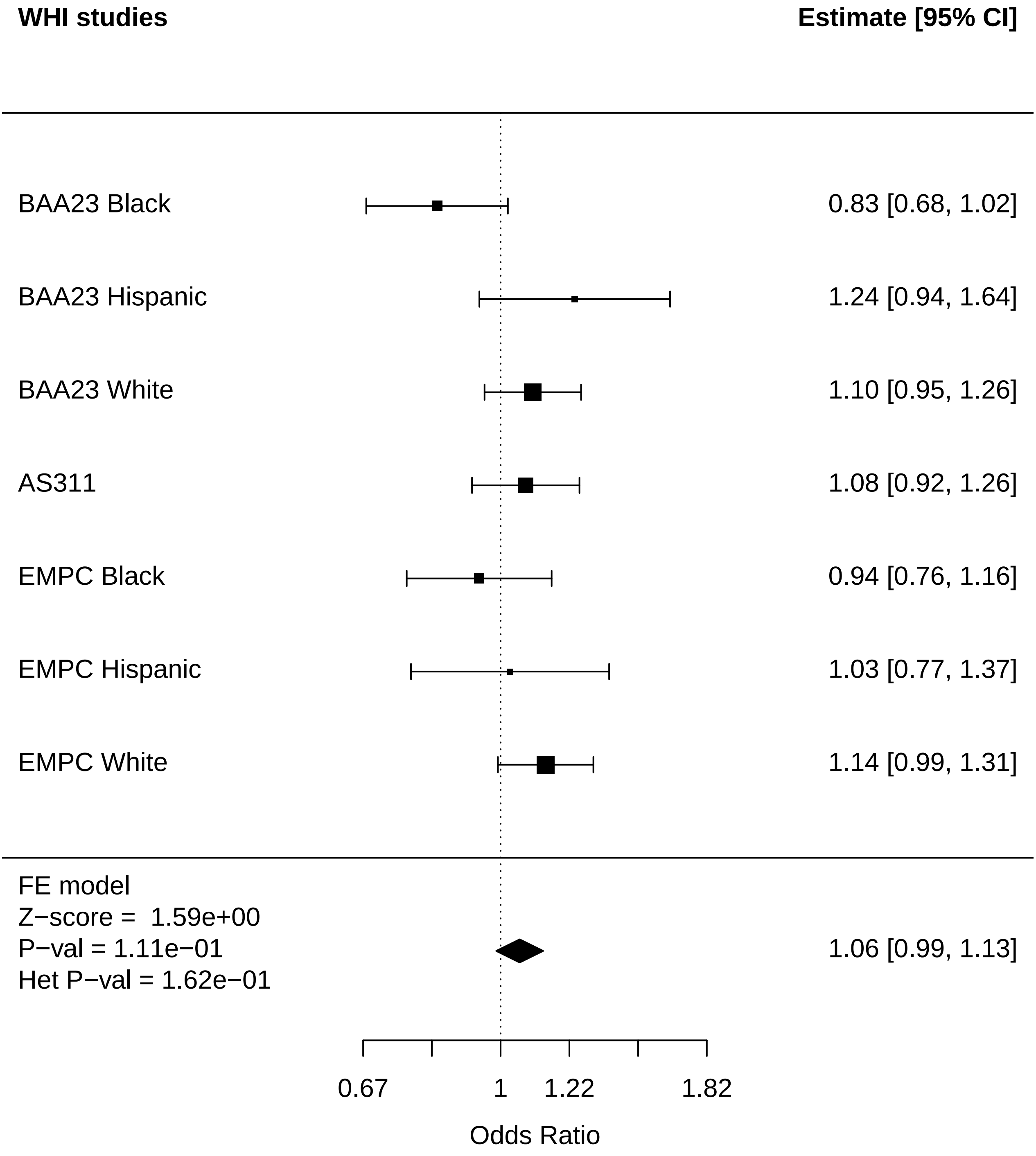

### DunedinPACE_Total comorbidities.pdf

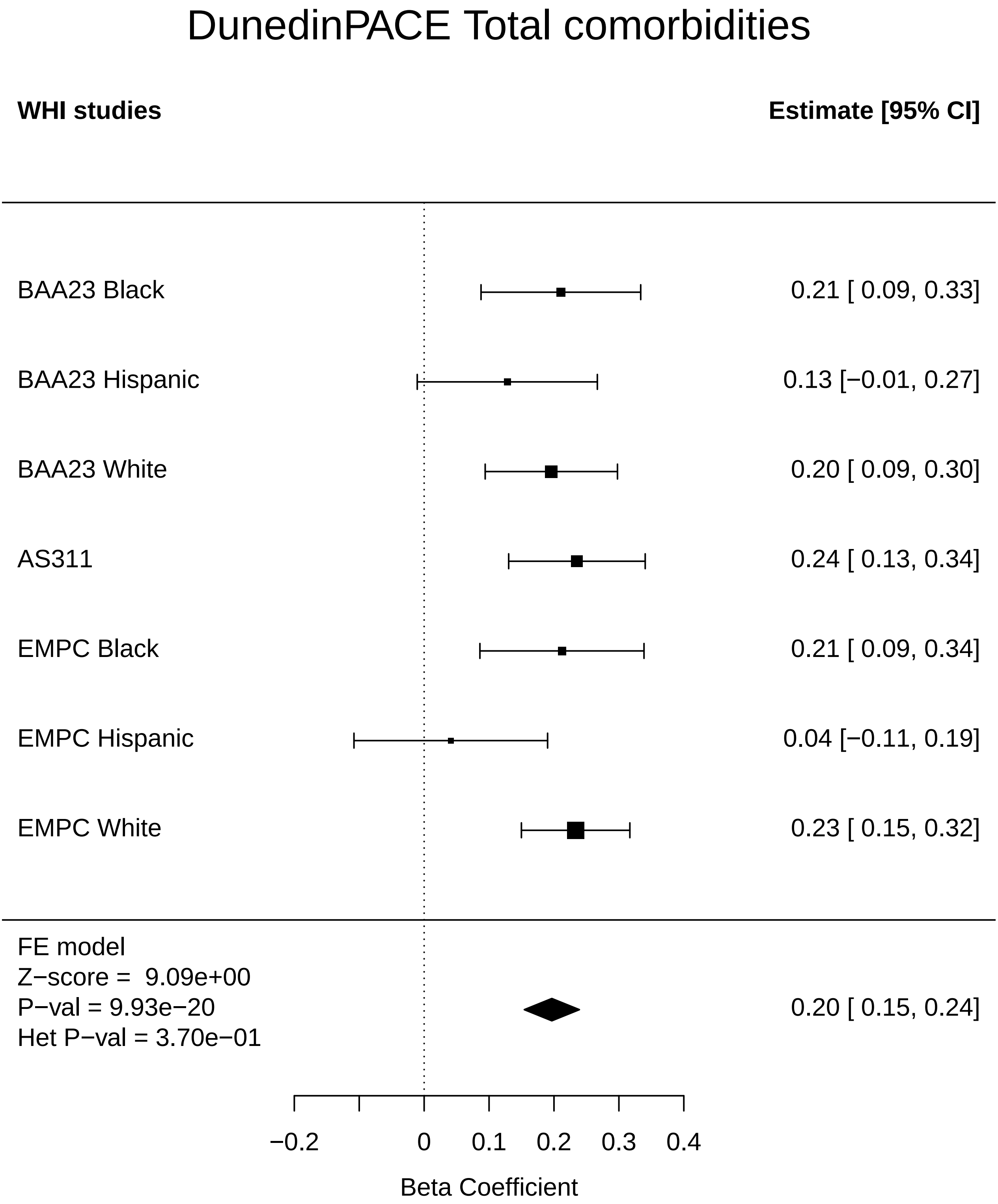

### Kidney_Any Cancer.pdf

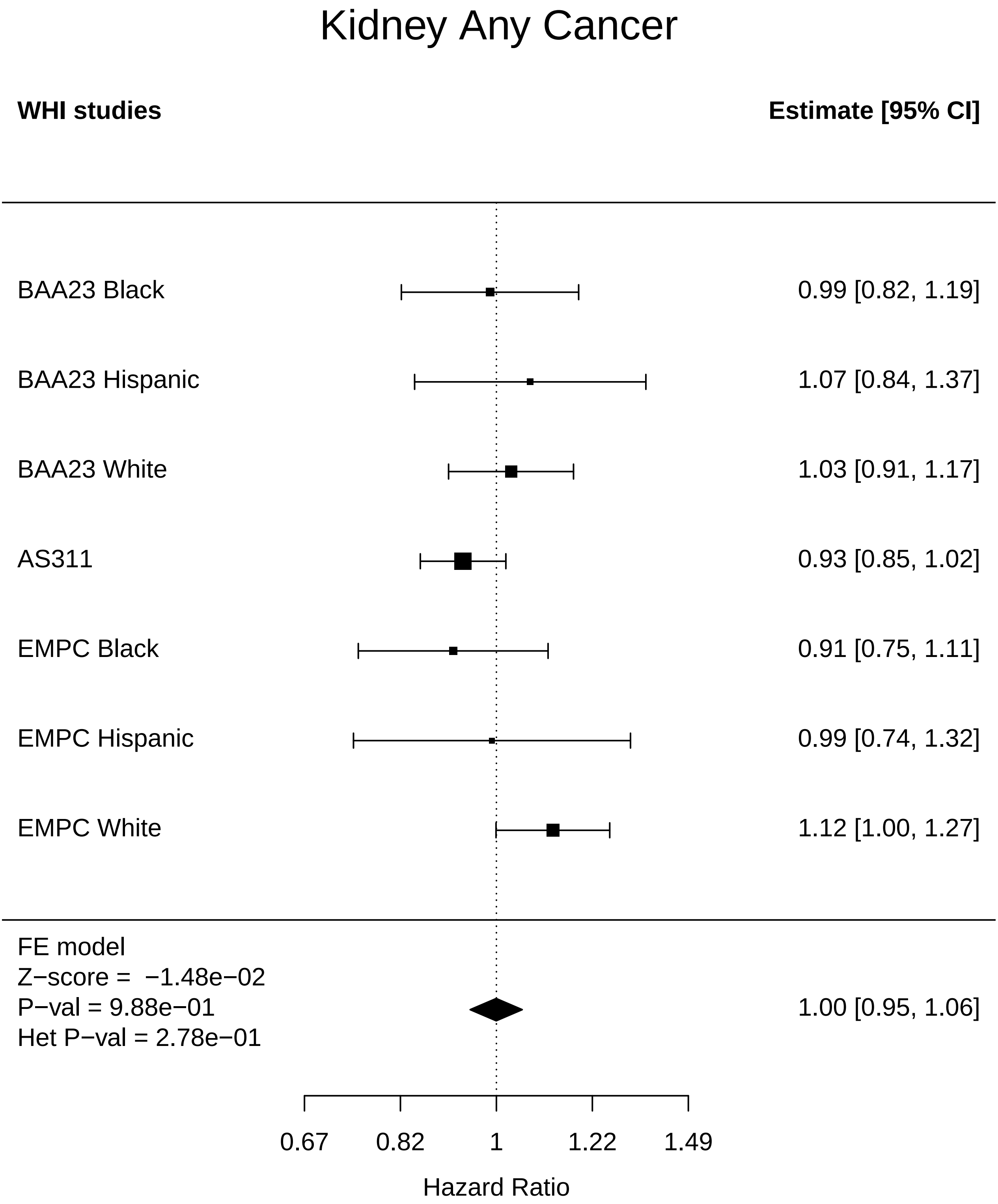

### Kidney_Breast Cancer.pdf

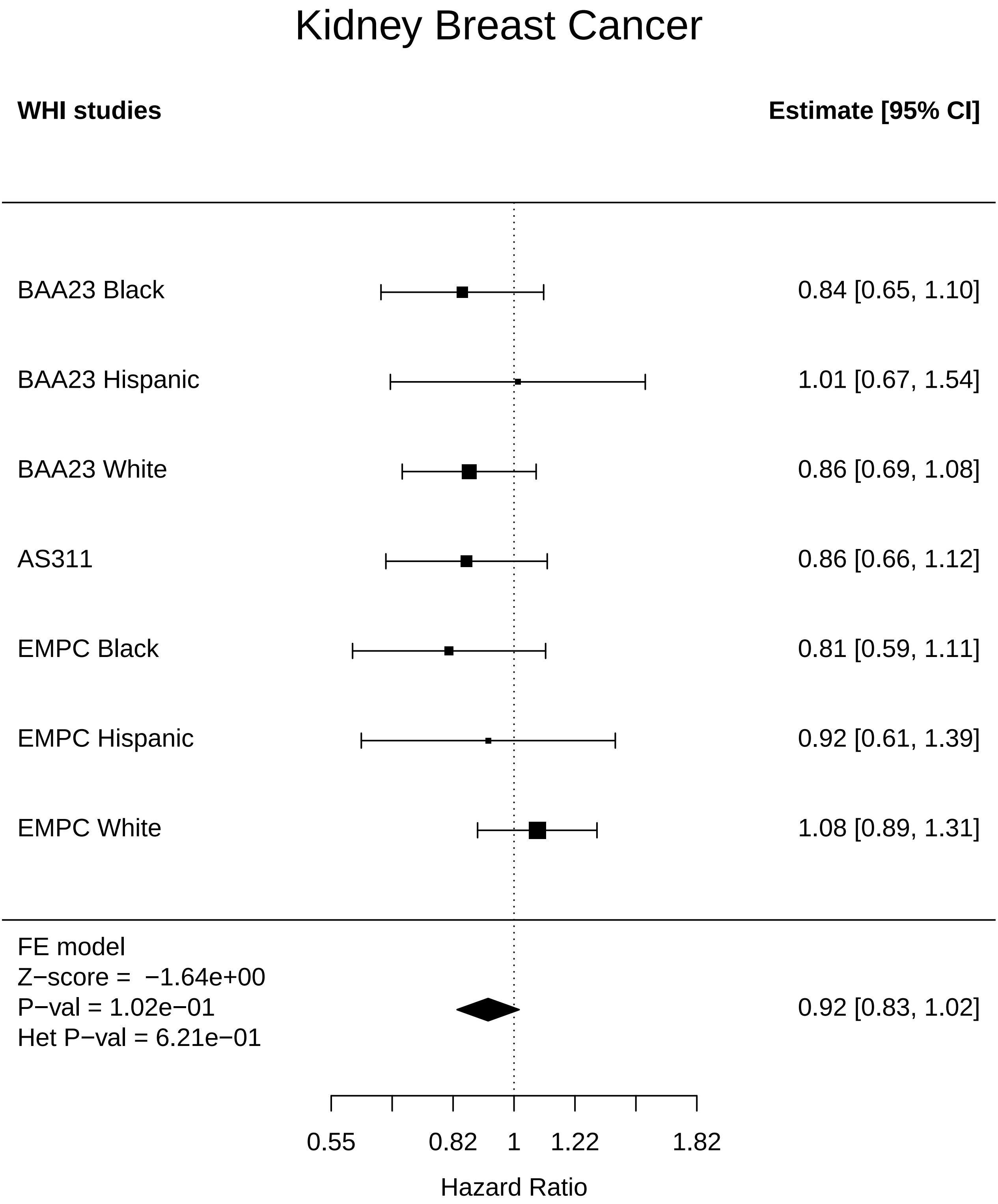

### Kidney_Cataract.pdf

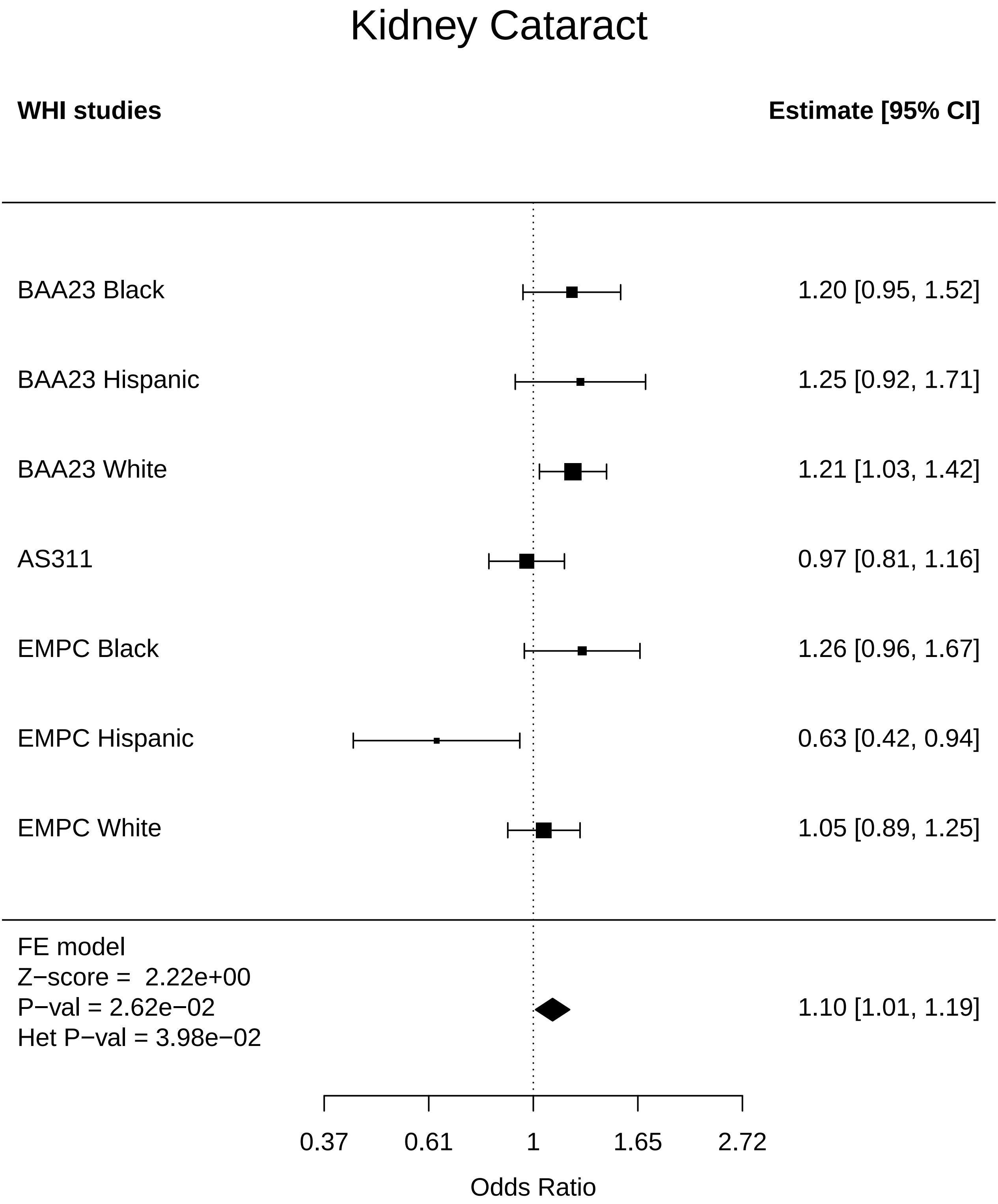

### Kidney_Colorectal cancer.pdf

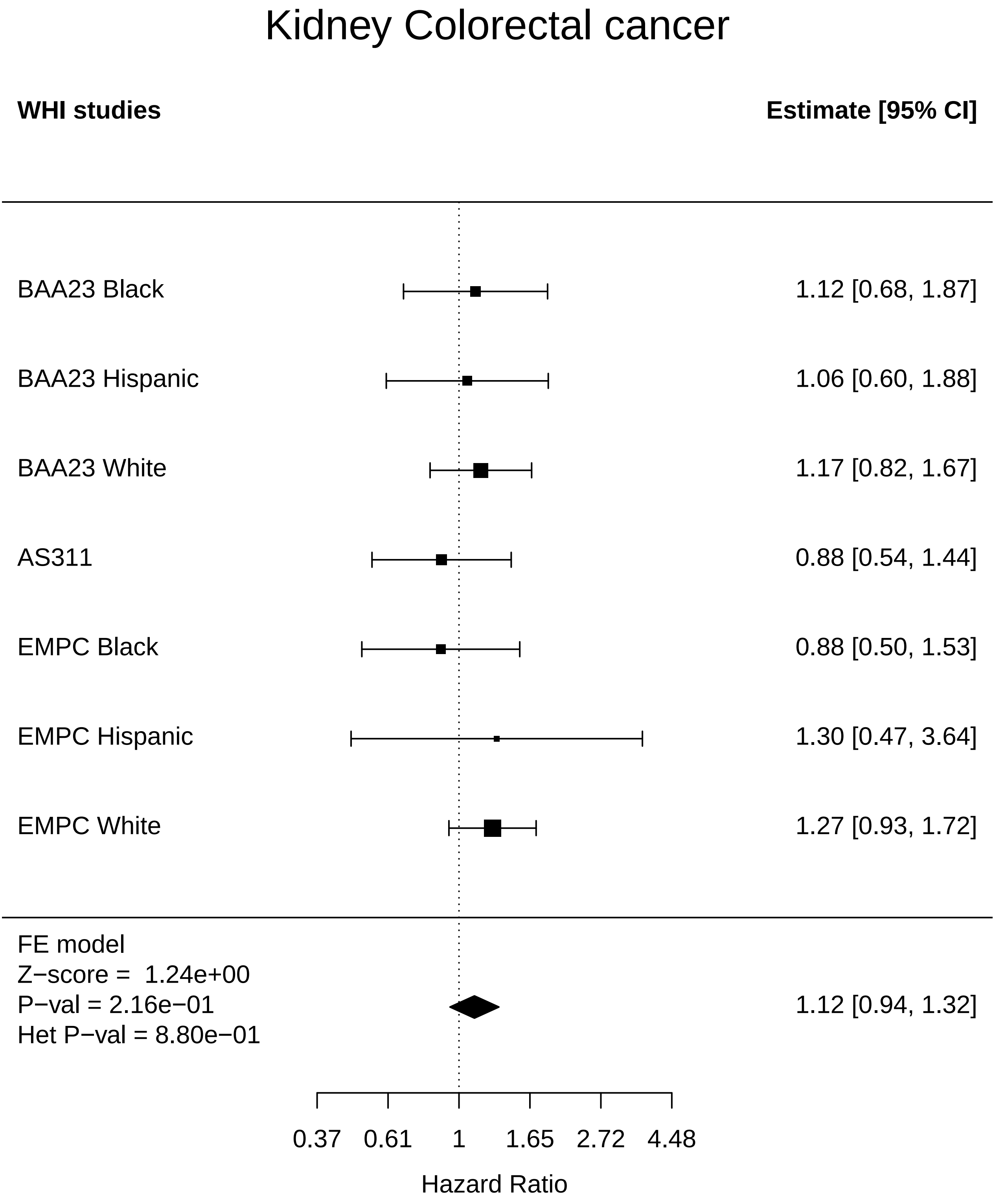

### Kidney_Leukemia.pdf

#### Kidney Leukemia

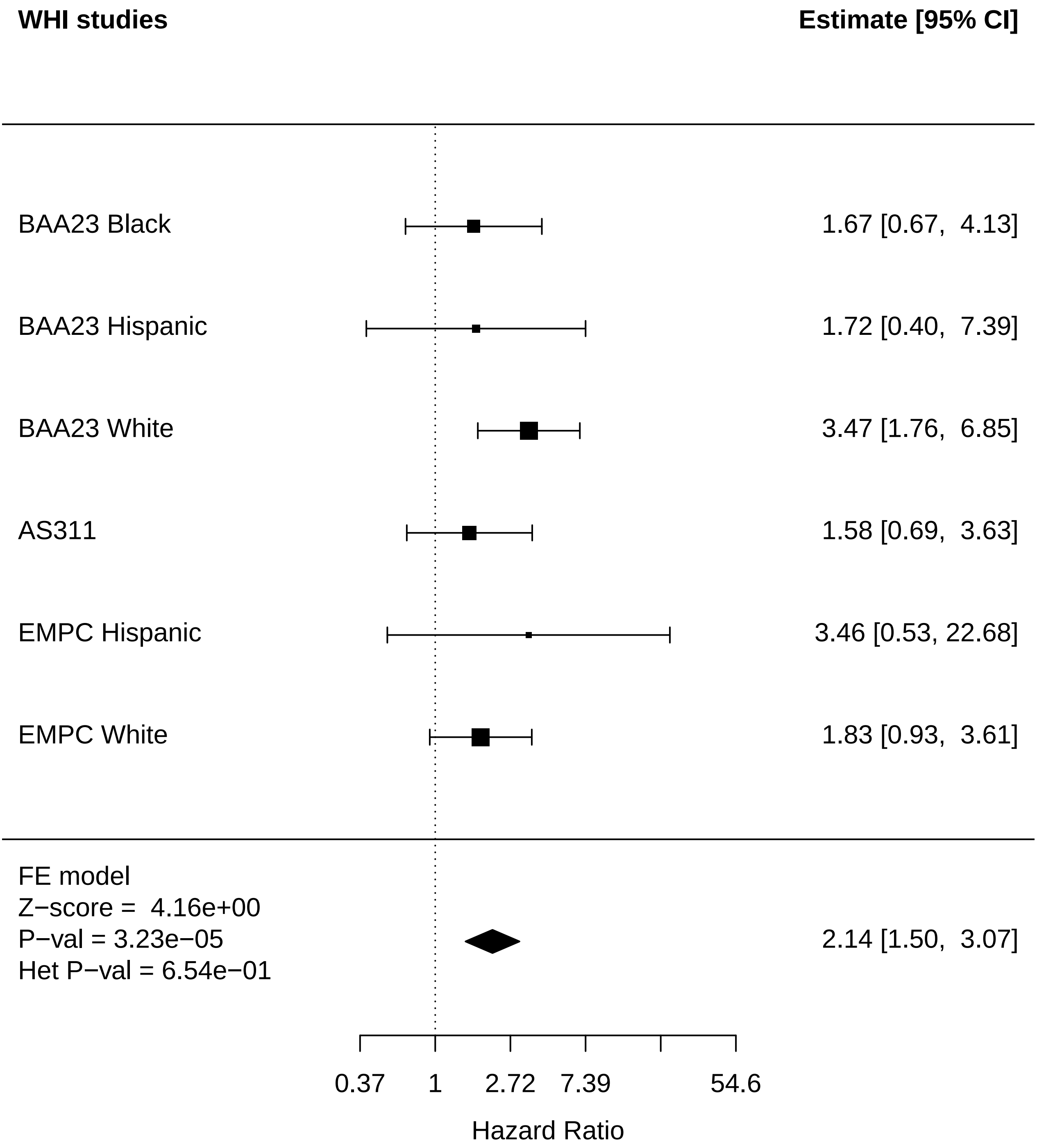

### Kidney_Mortality.pdf

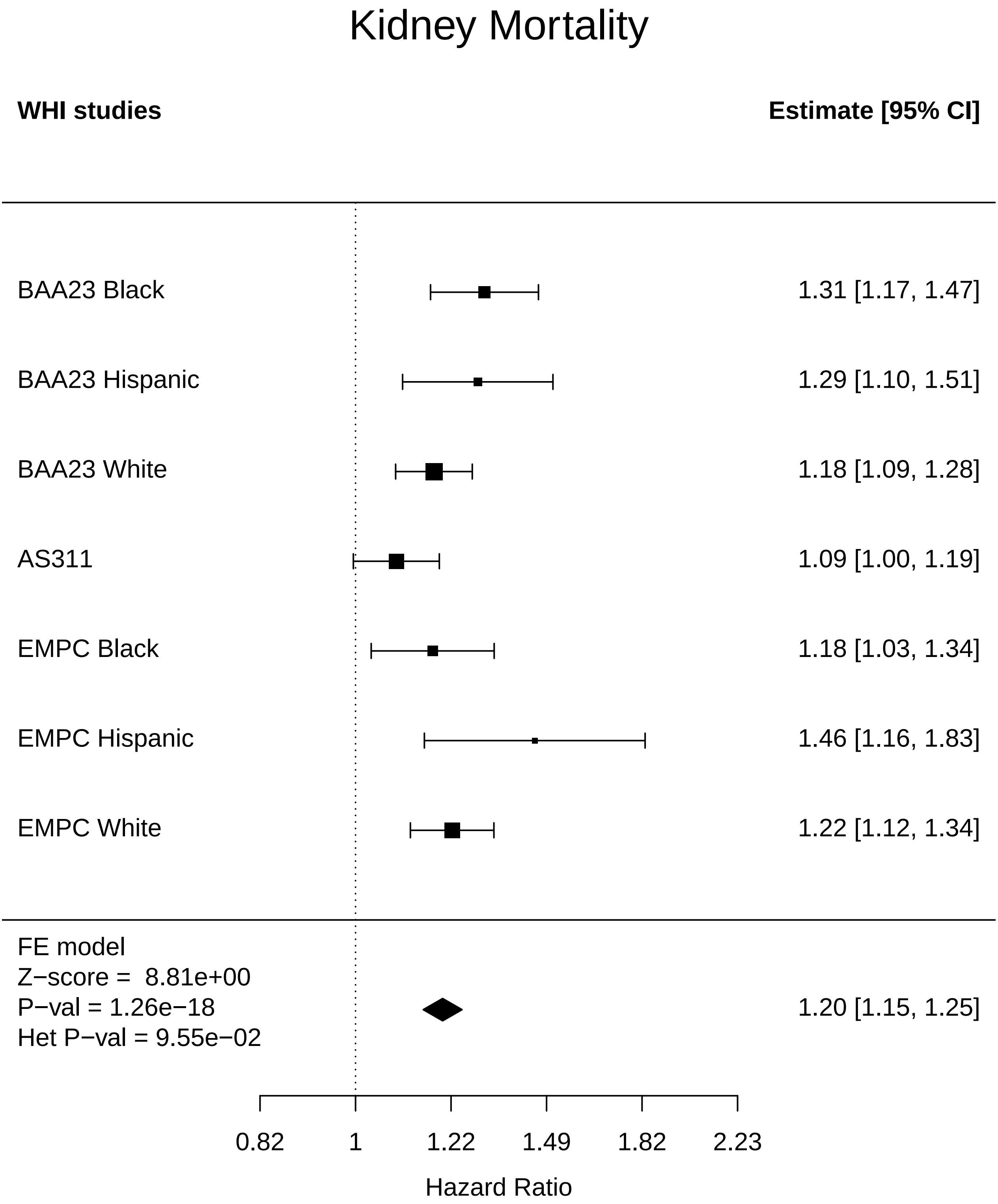

### Liver_Any Cancer.pdf

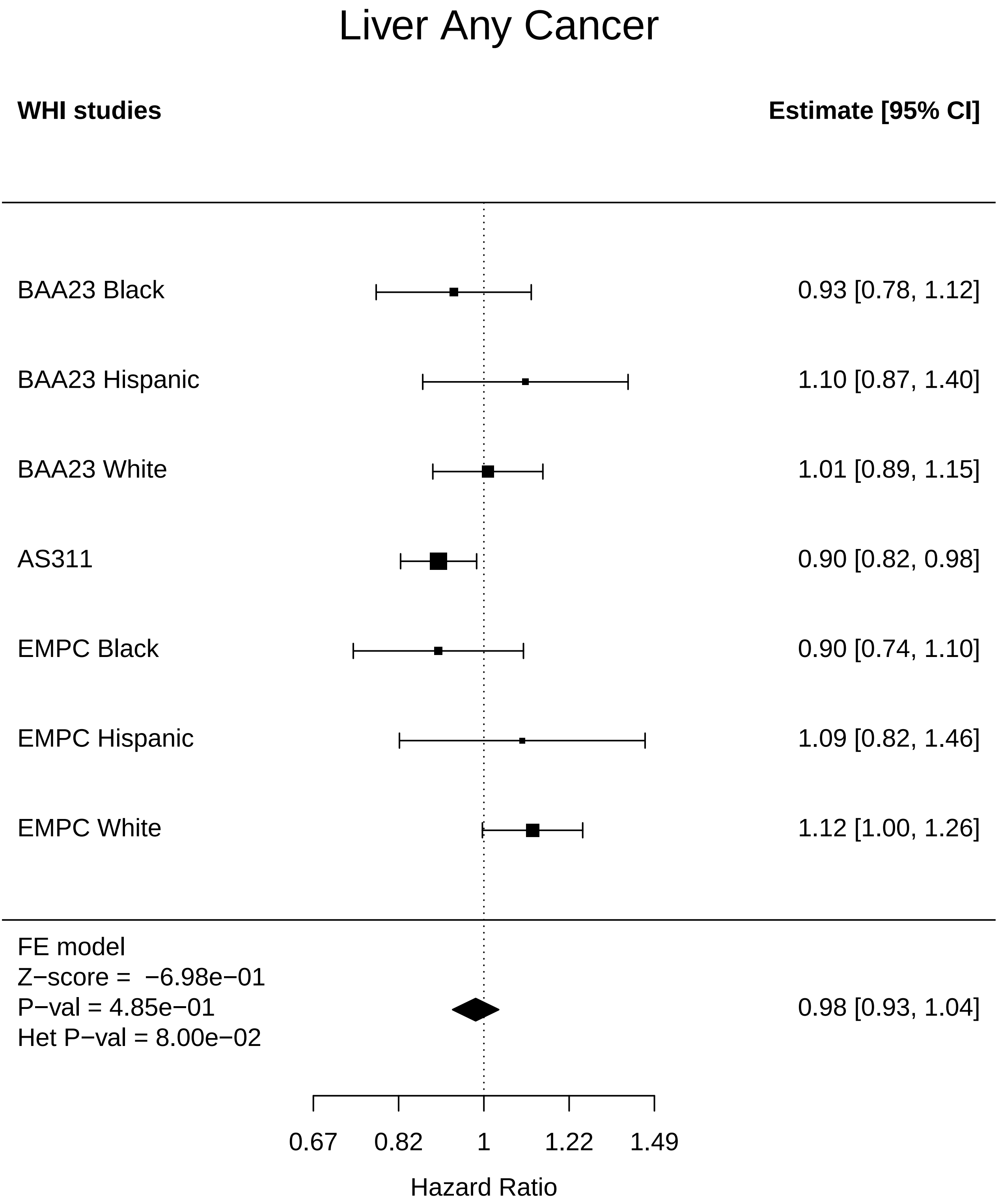

### Liver_Cataract.pdf

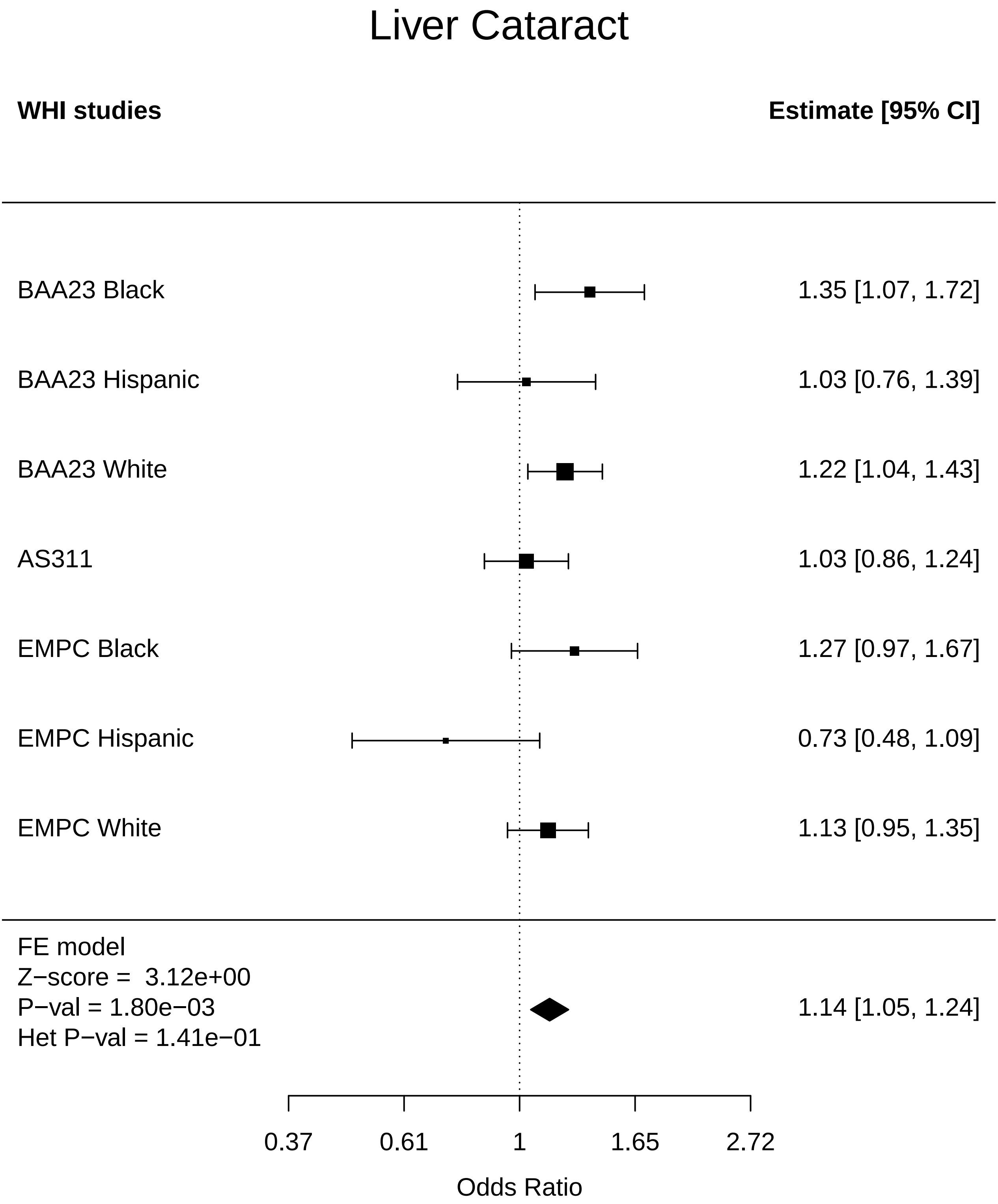

### Liver_CHD.pdf

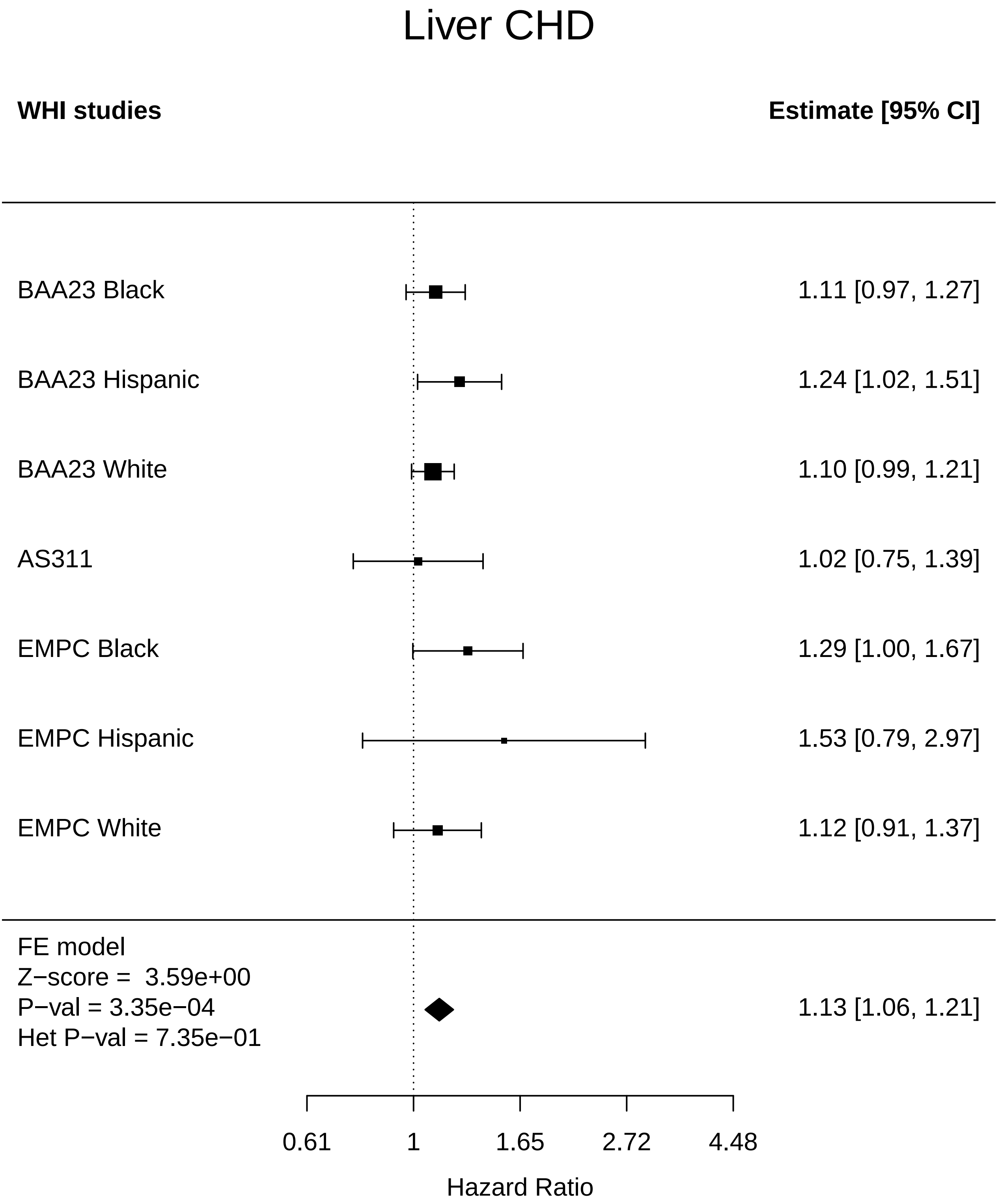

### Liver_Cognitive function.pdf

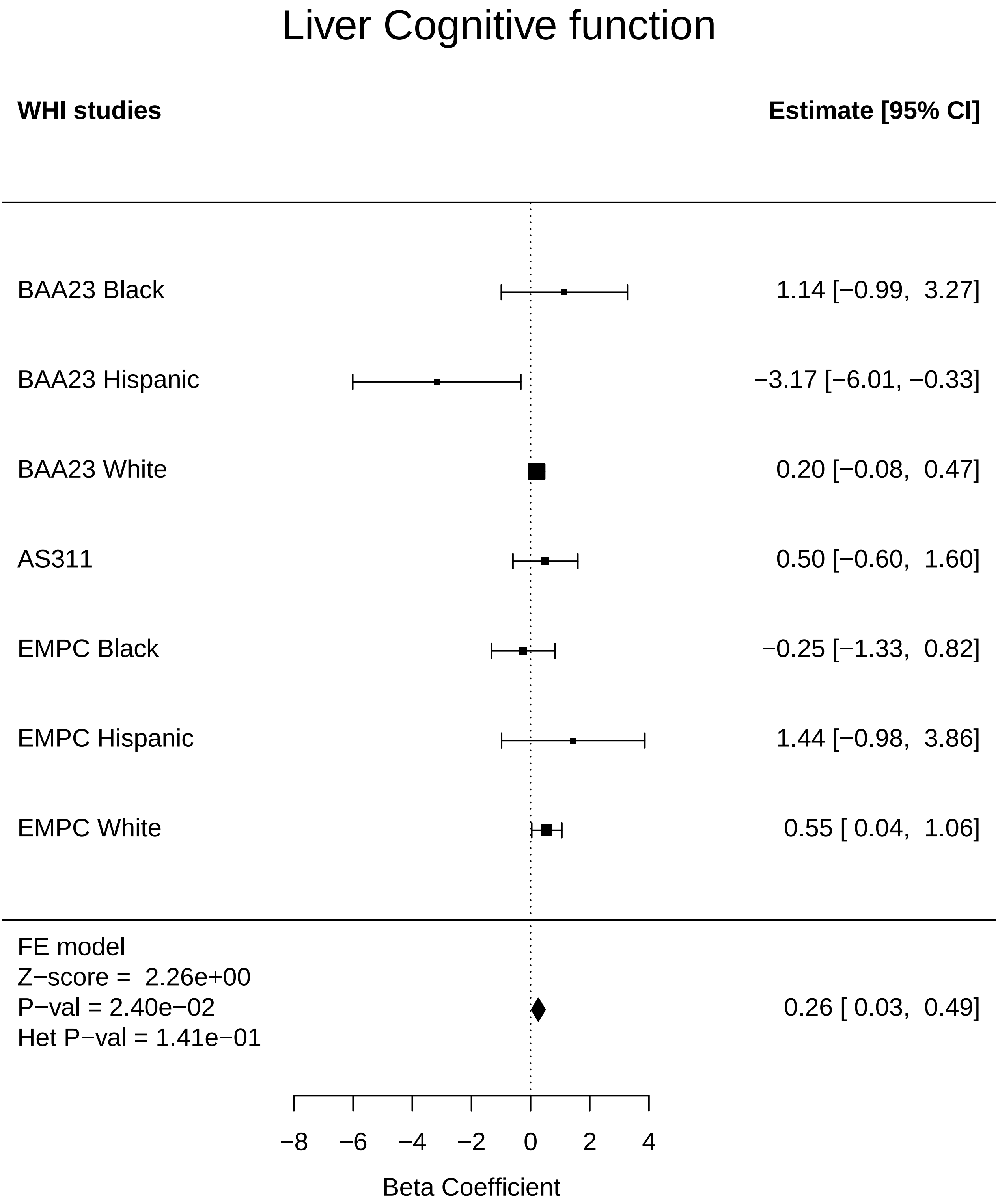

### Liver_Colorectal cancer.pdf

#### Liver Colorectal cancer

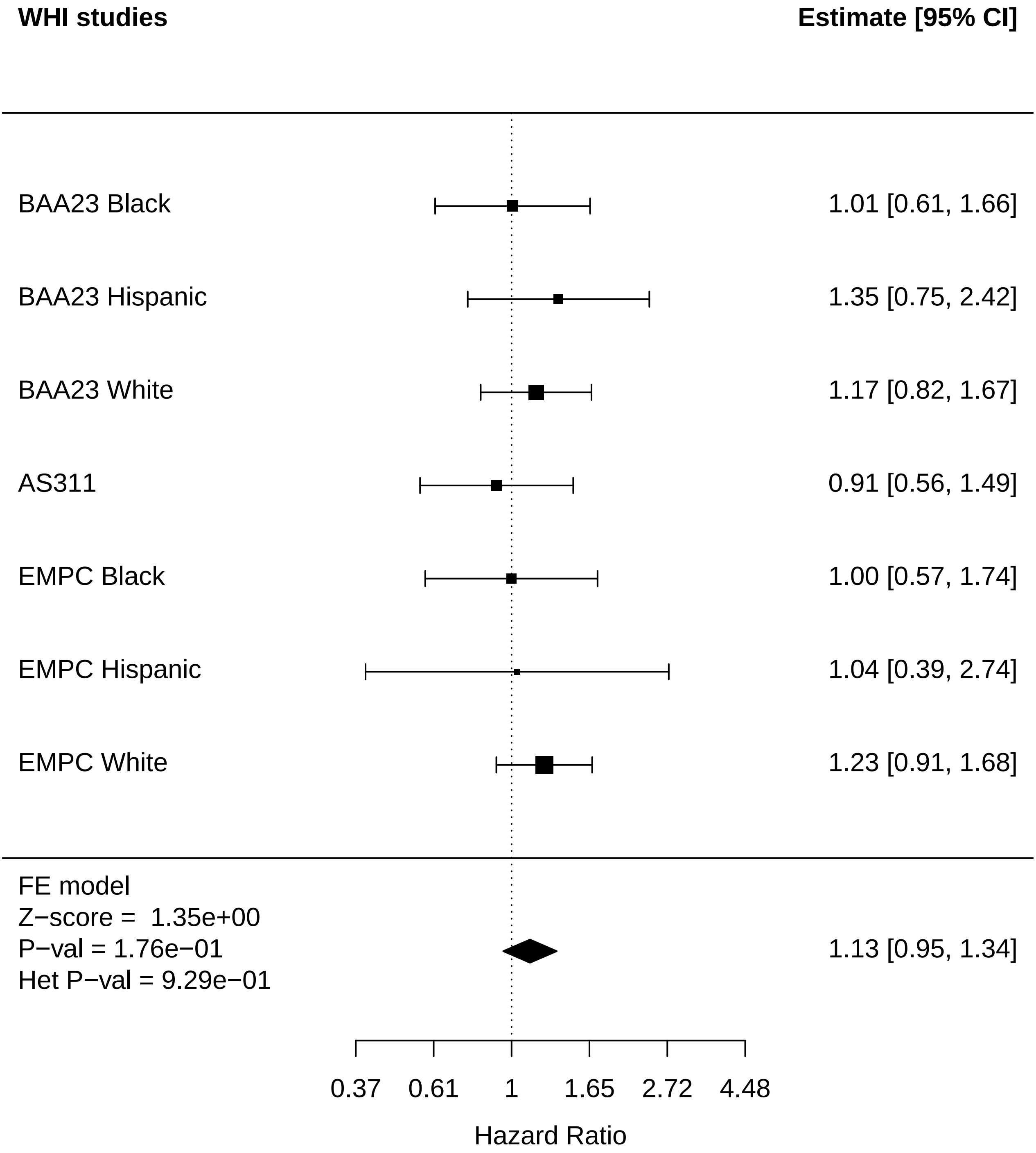

### Liver_Leukemia.pdf

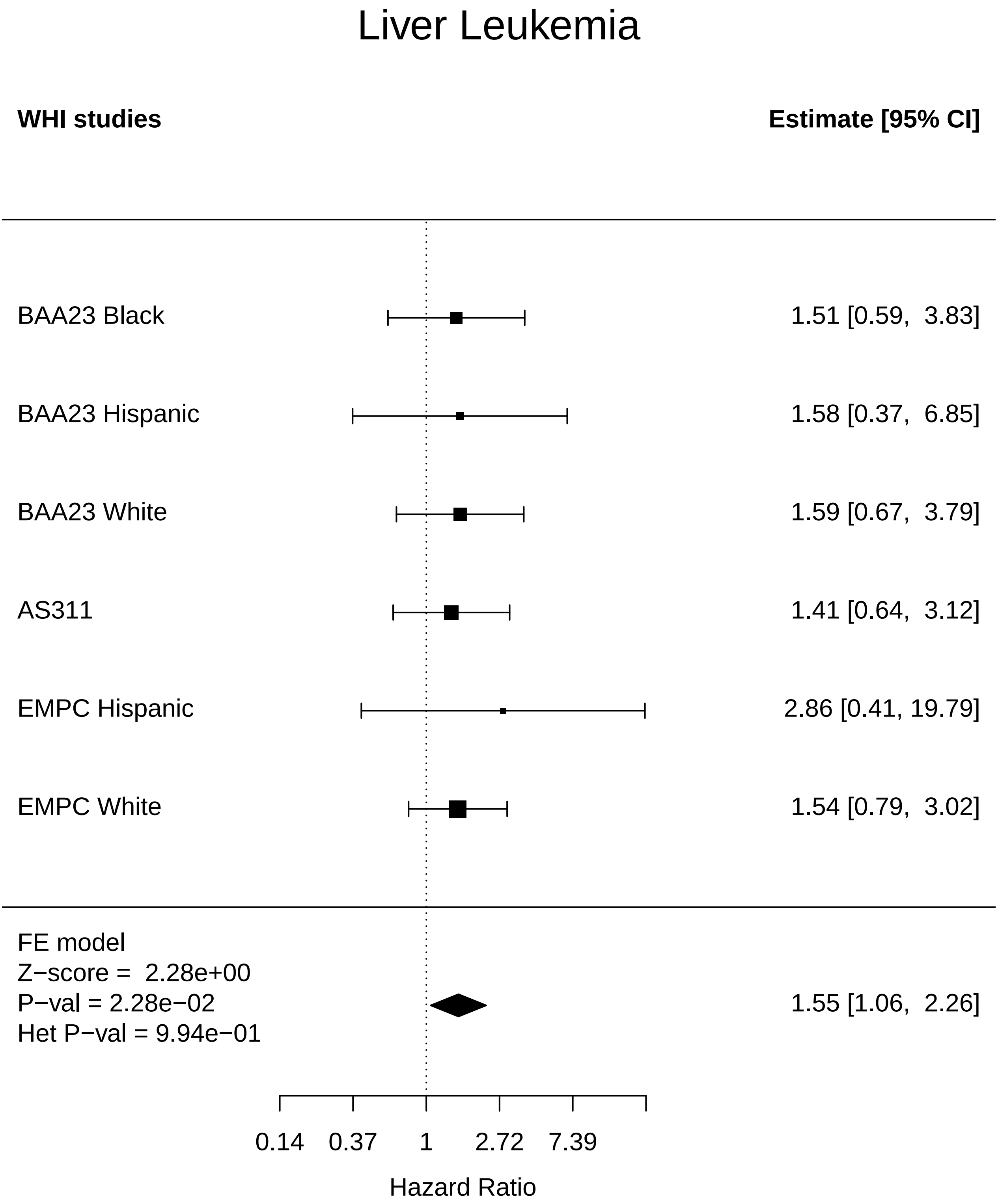

### Liver_Lung Cancer.pdf

#### Liver Lung Cancer

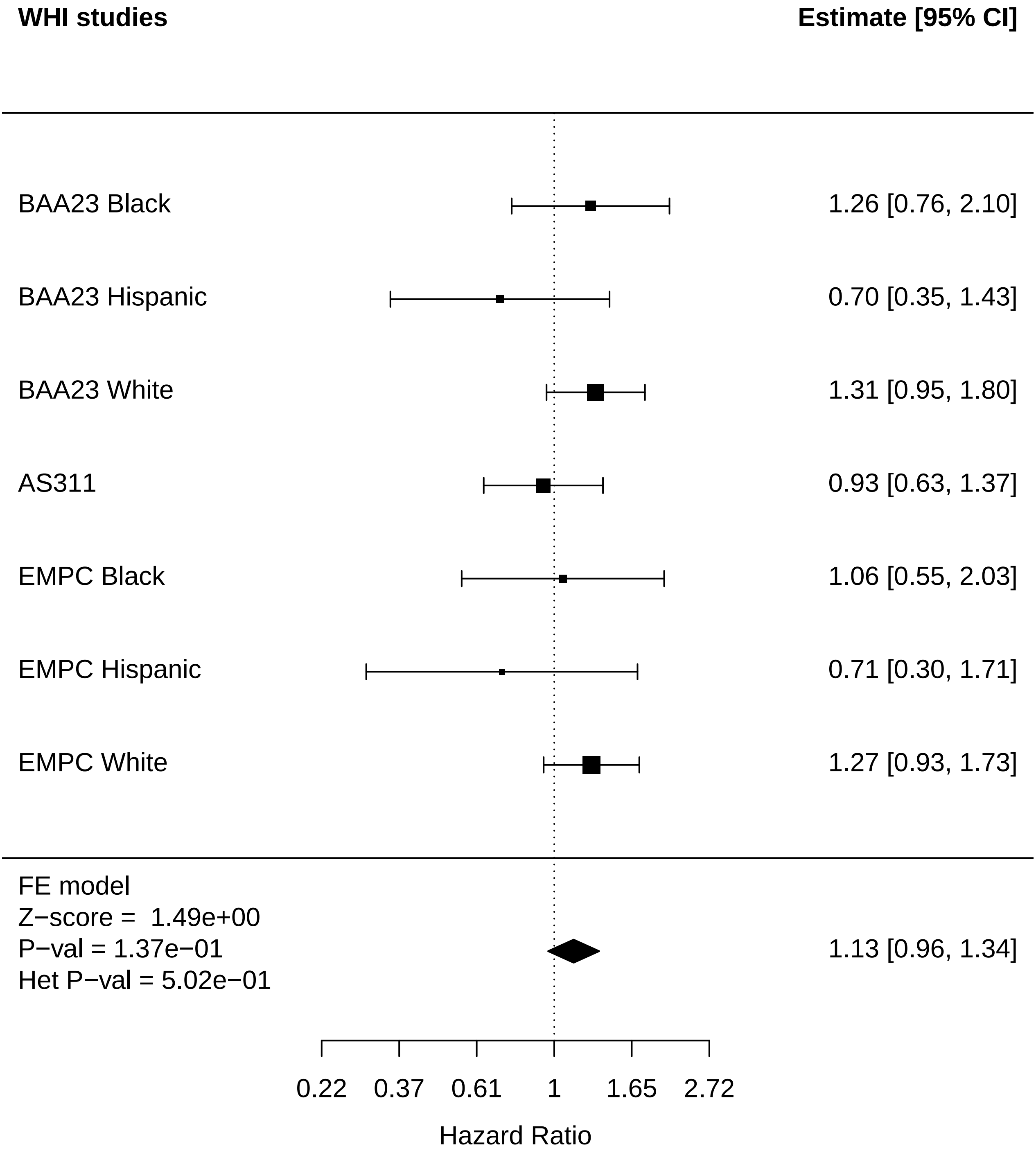

### Liver_Mortality.pdf

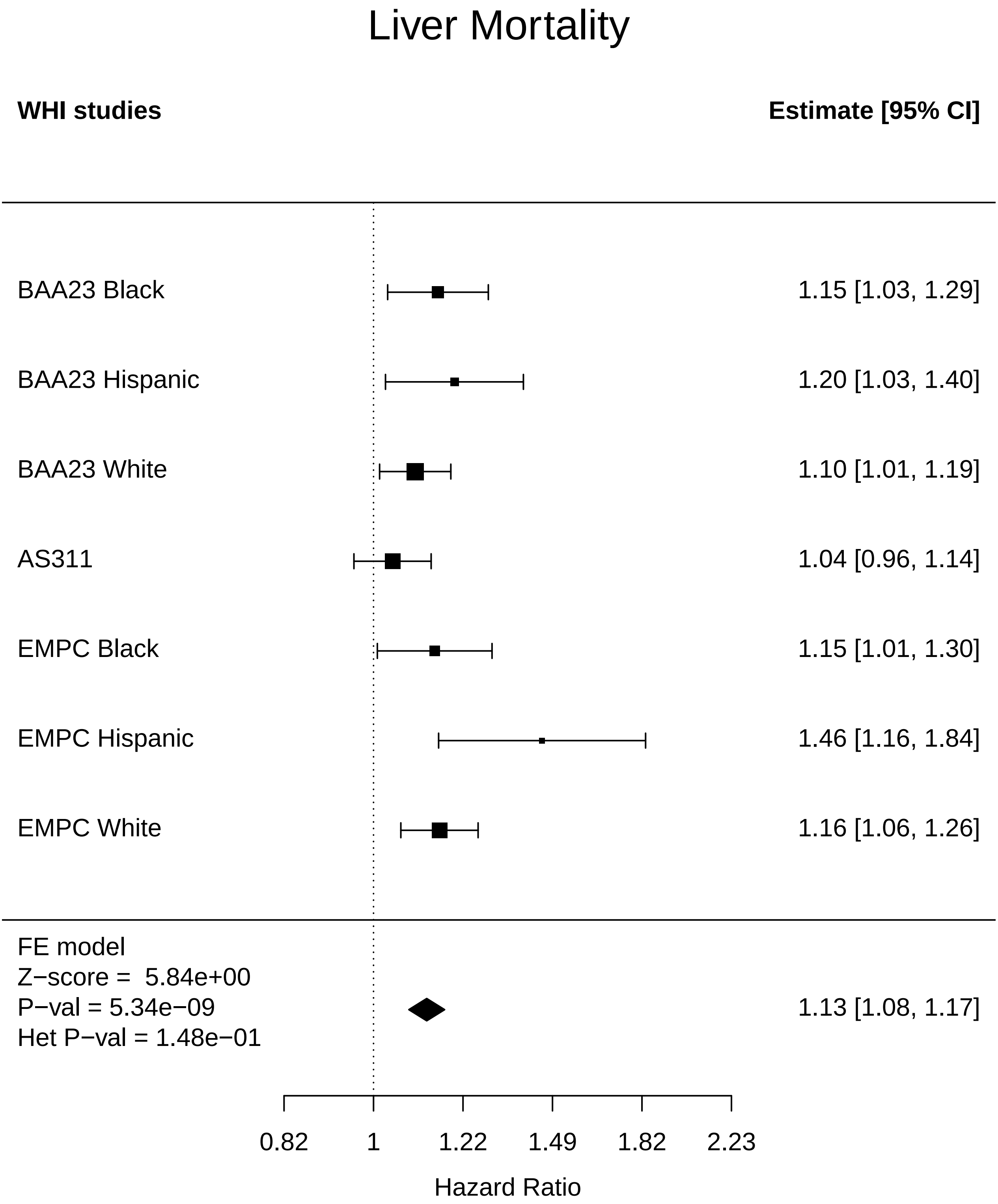

### Liver_Thyroid disease.pdf

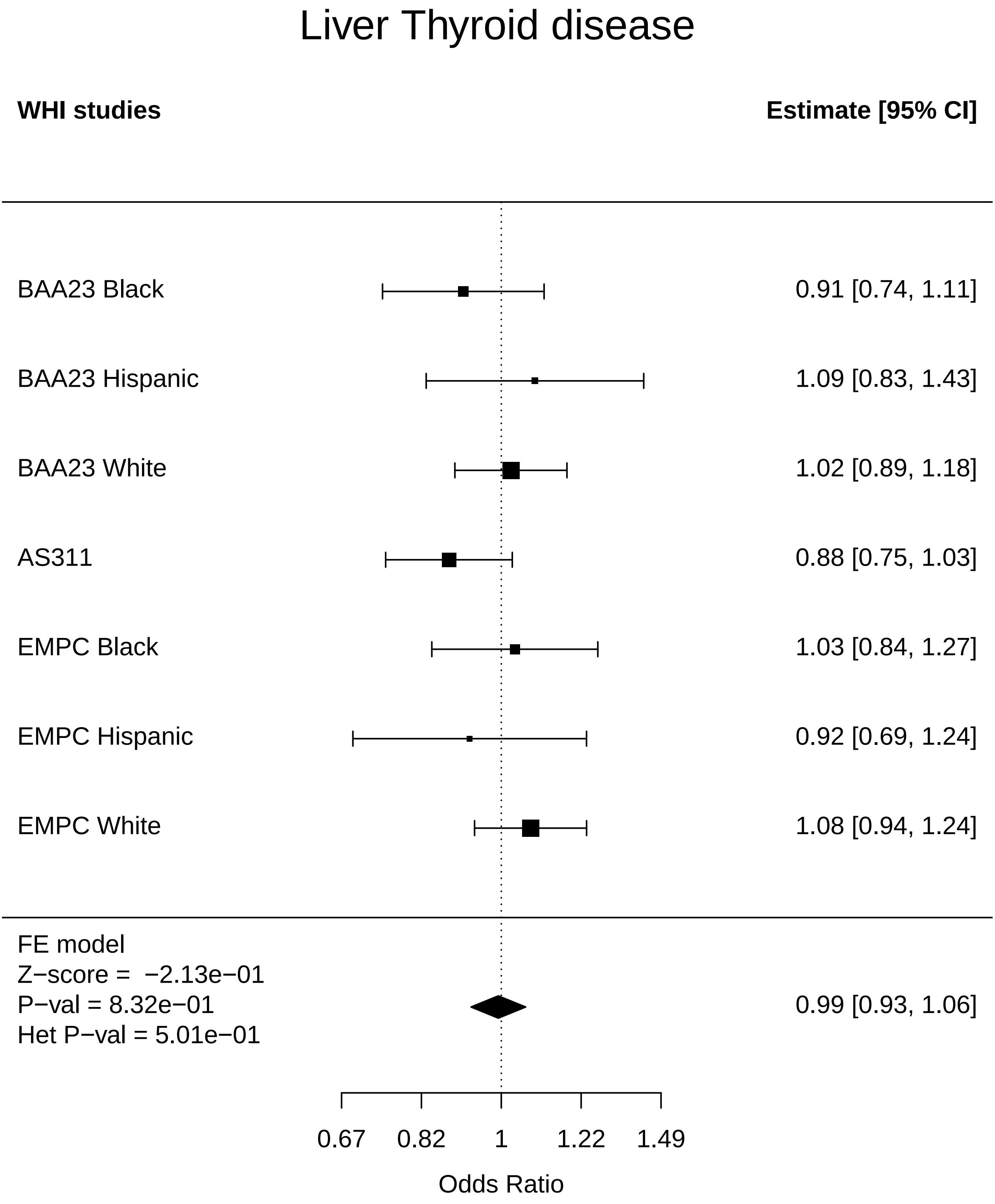

### Liver_Total comorbidities.pdf

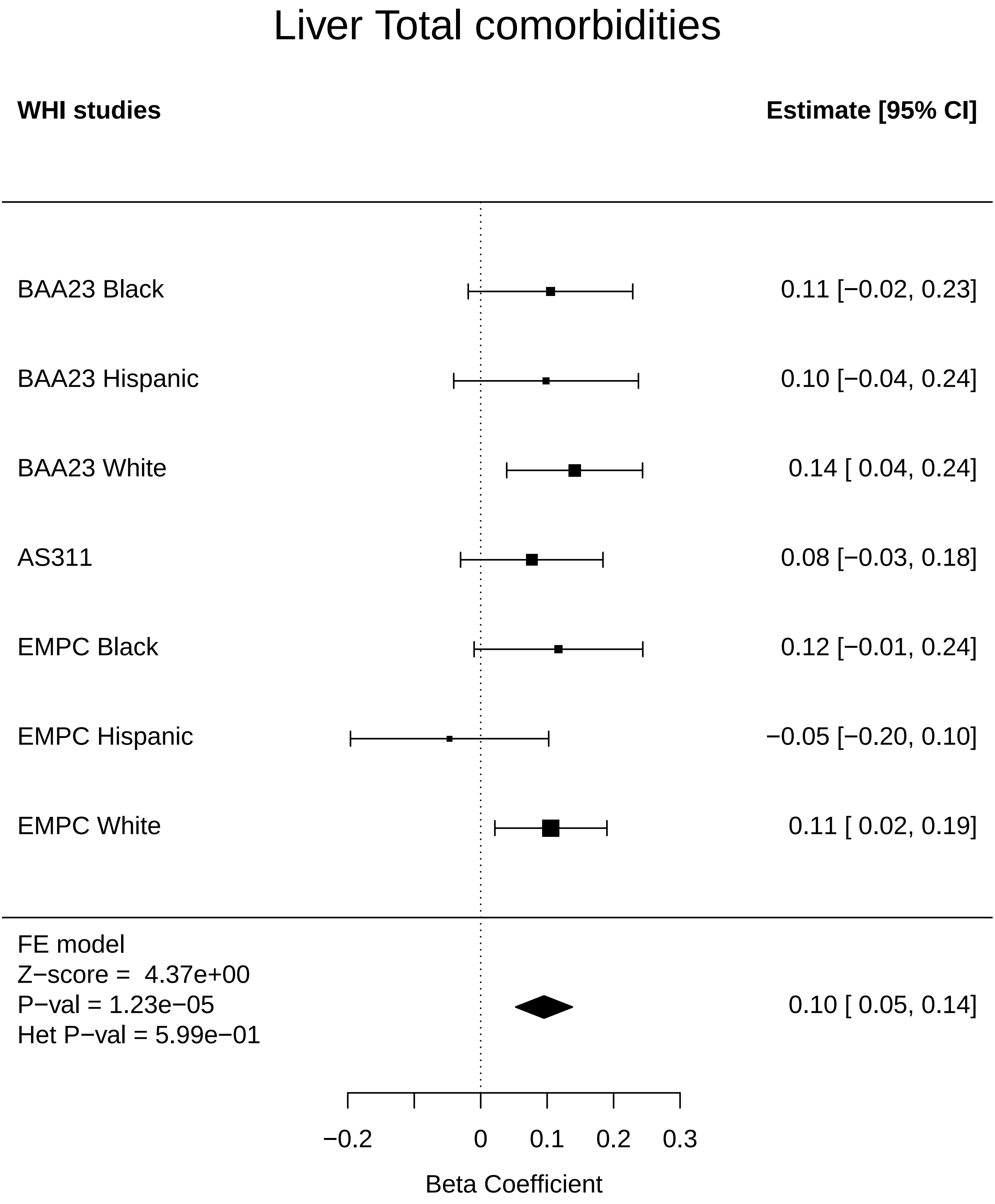

### Lung_Any Cancer.pdf

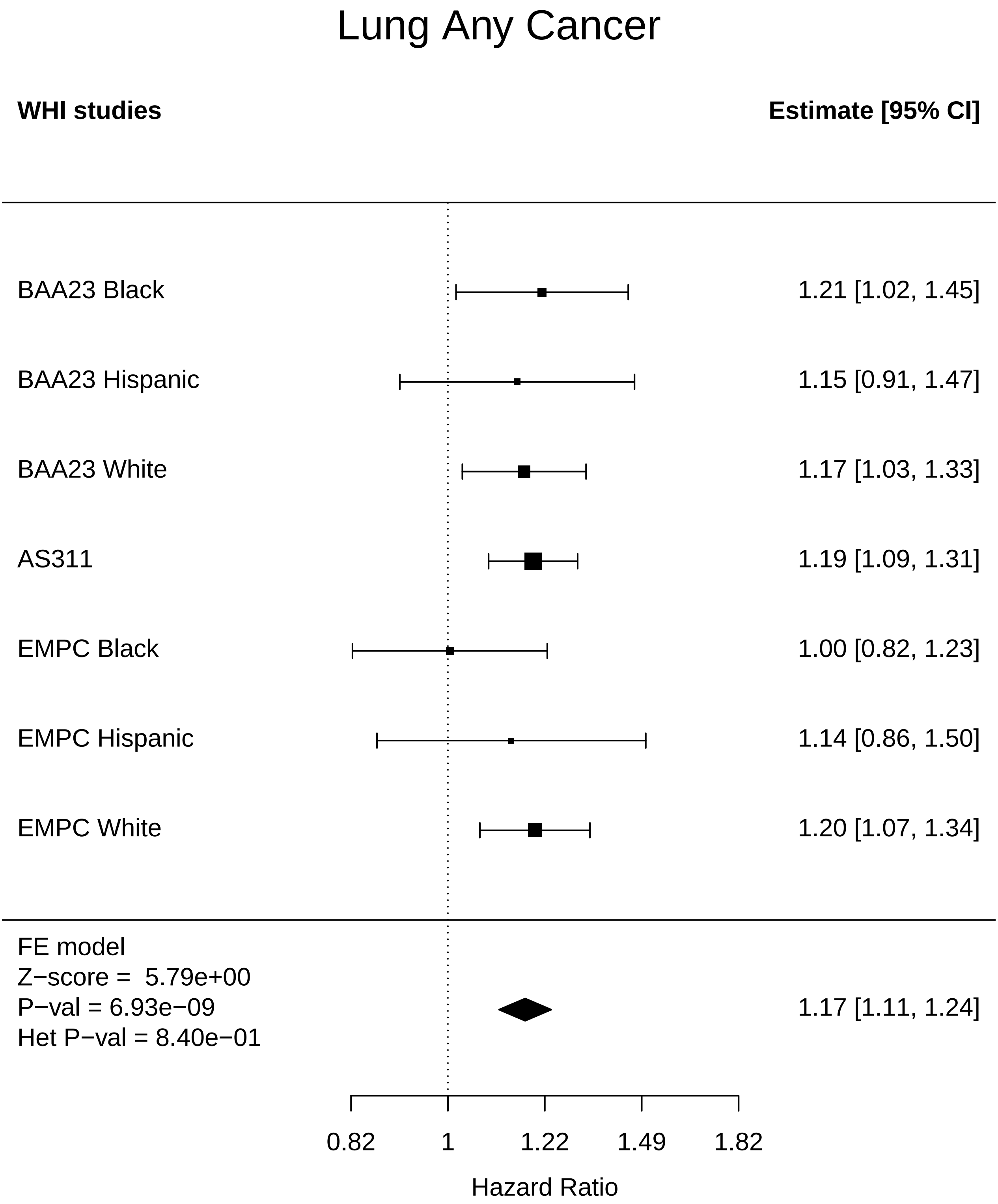

### Lung_Arthritis.pdf

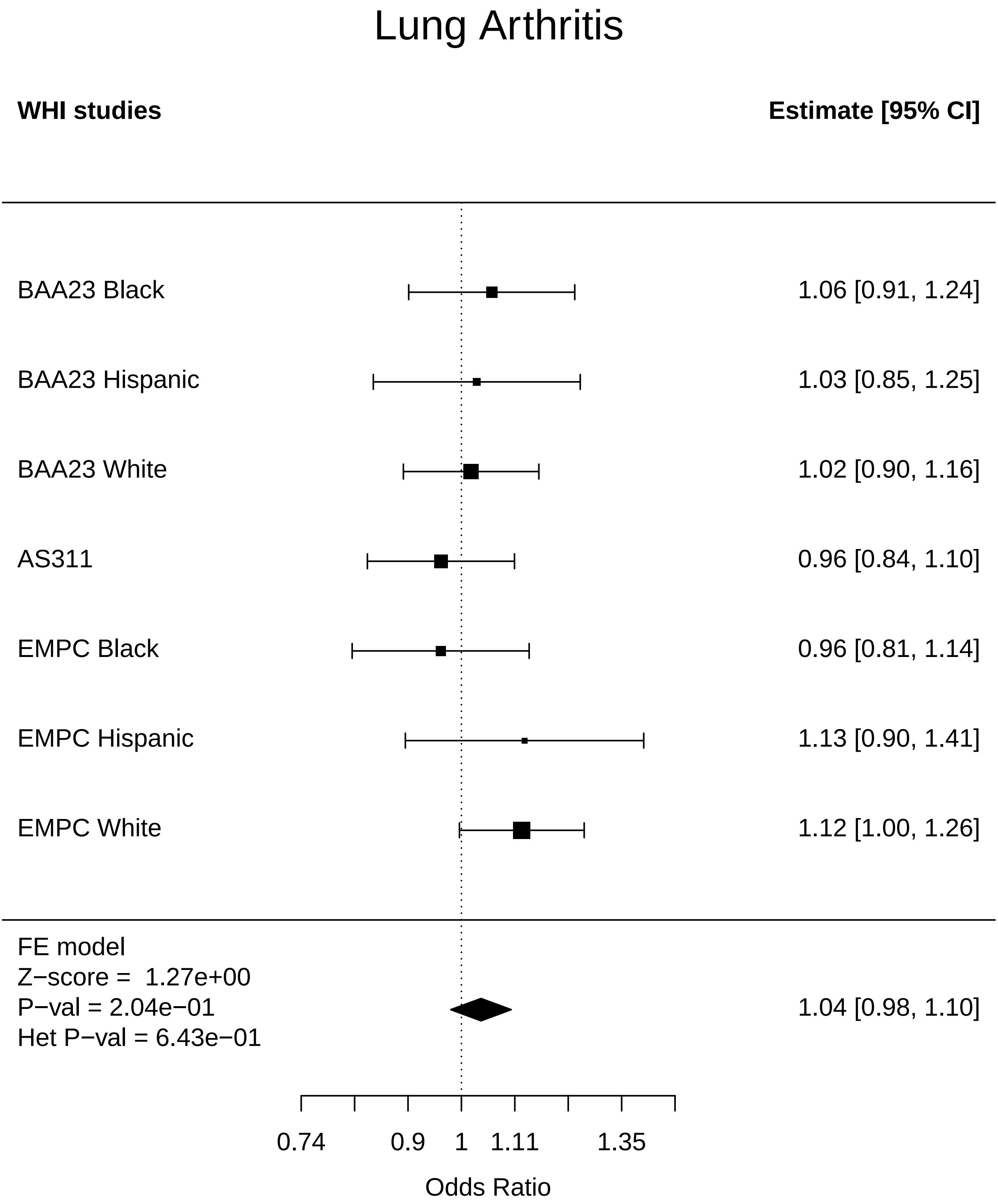

### Lung_Breast Cancer.pdf

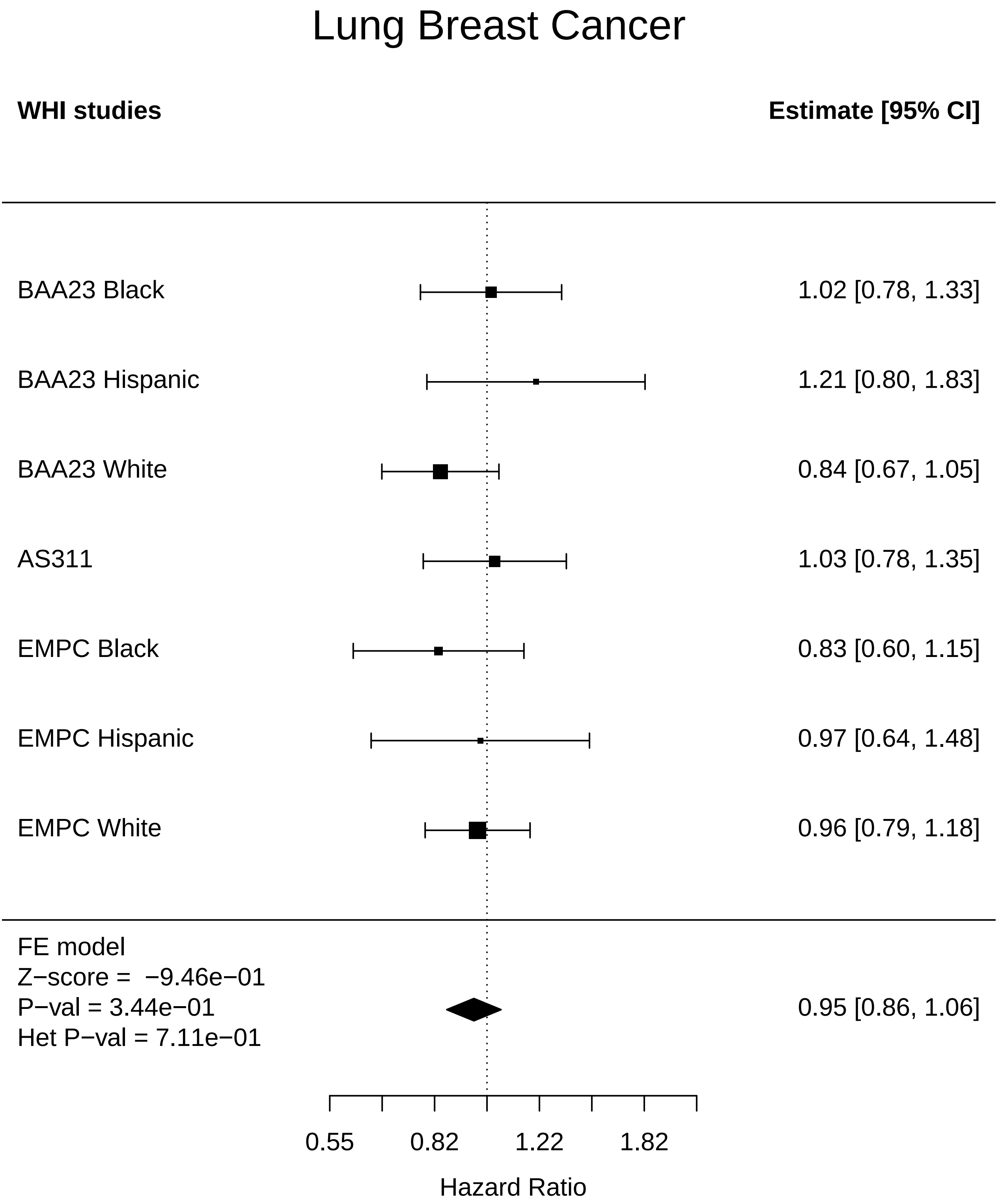

### Lung_Colorectal cancer.pdf

#### Lung Colorectal cancer

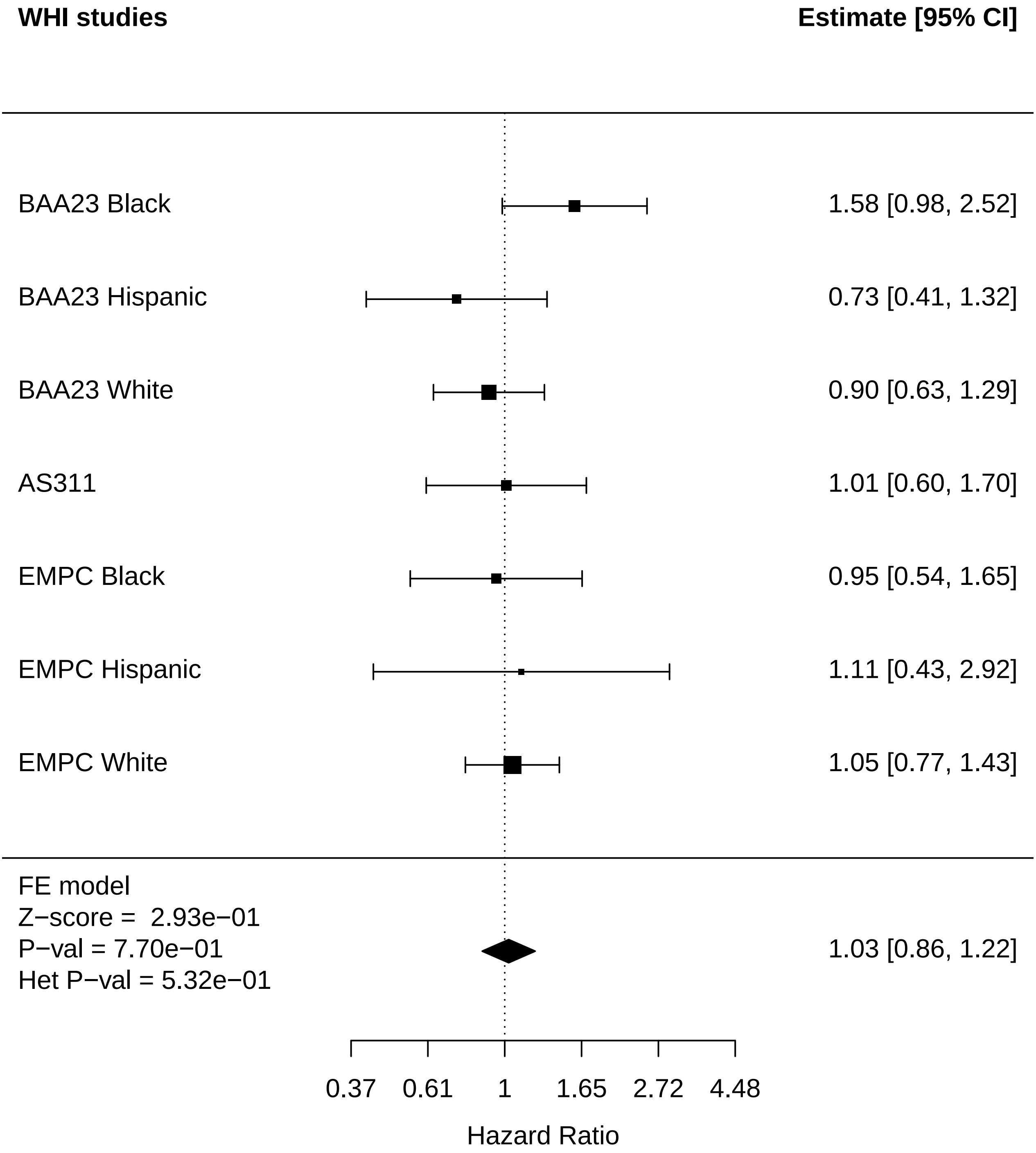

### Lung_Leukemia.pdf

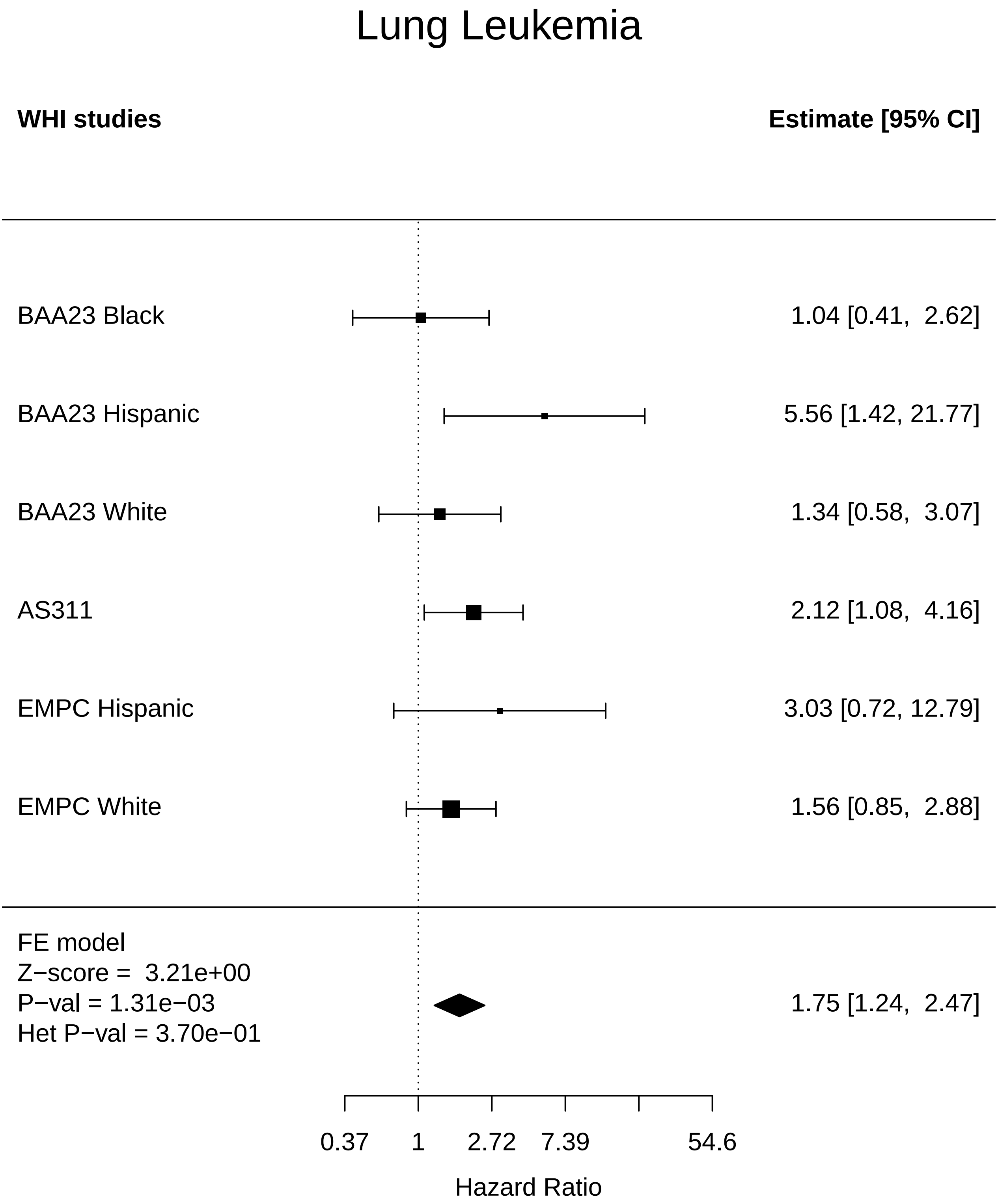

### Lung_Lung Cancer.pdf

#### Lung Lung Cancer

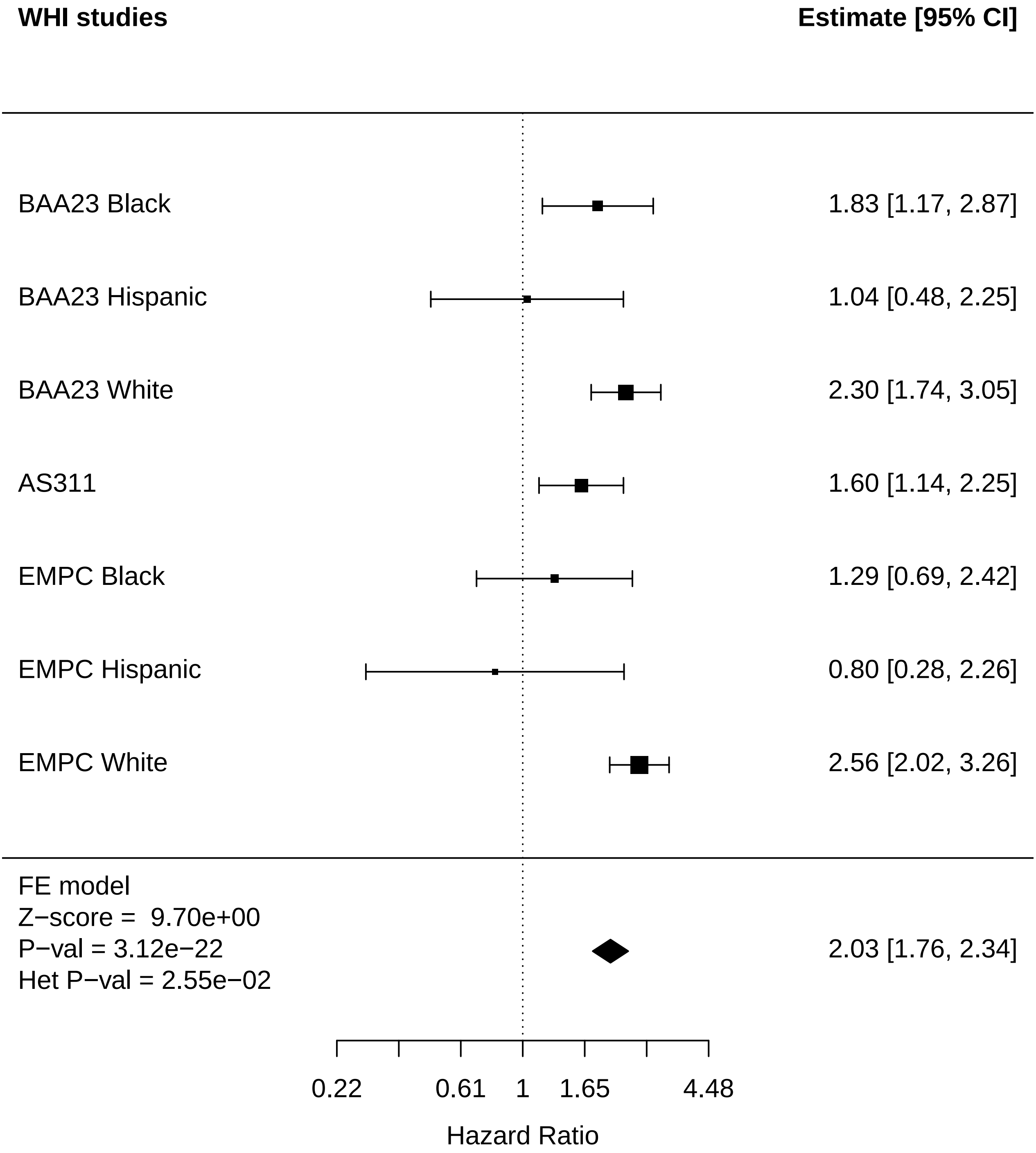

### Lung_MI.pdf

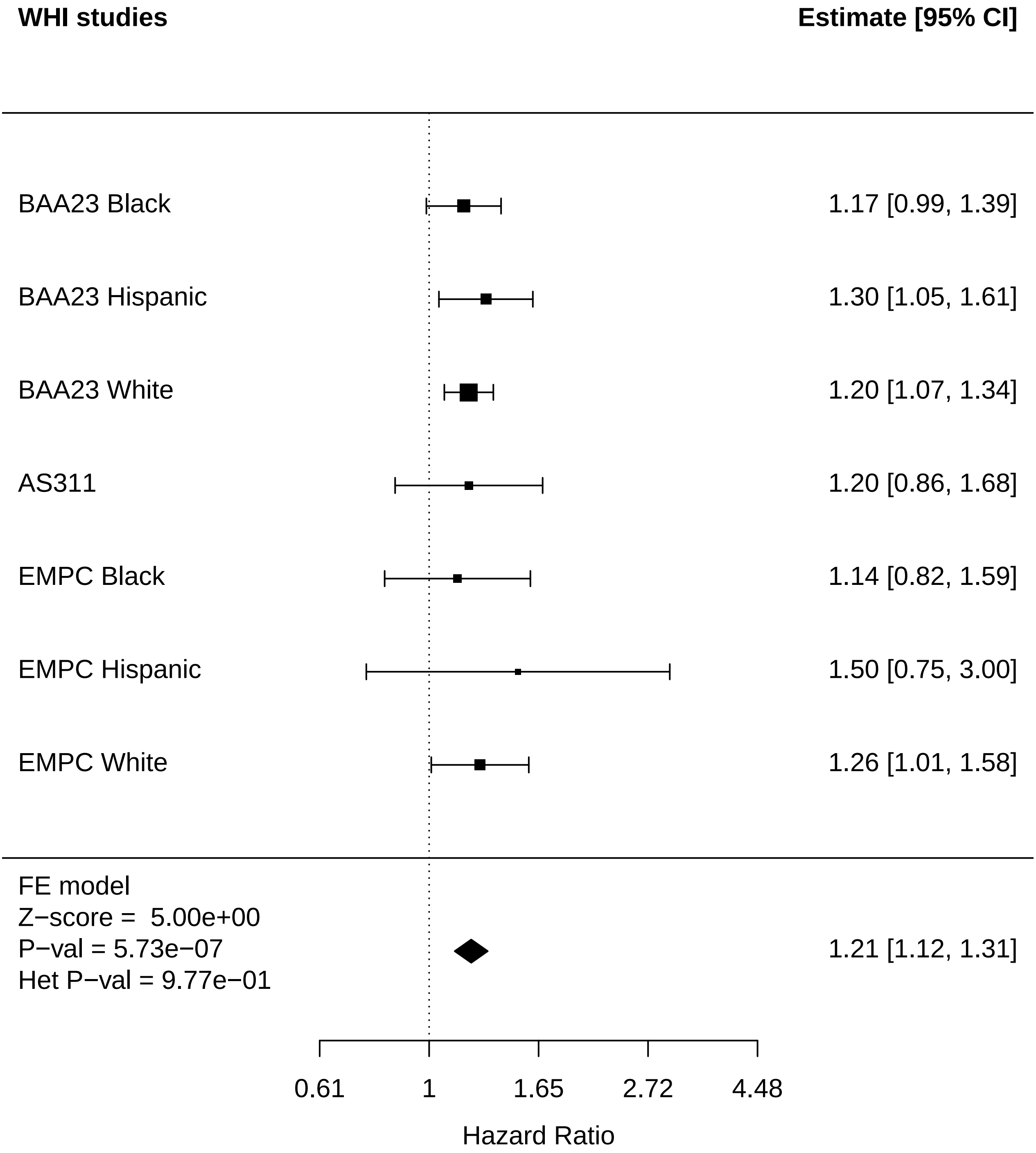

### Lung_Physical function.pdf

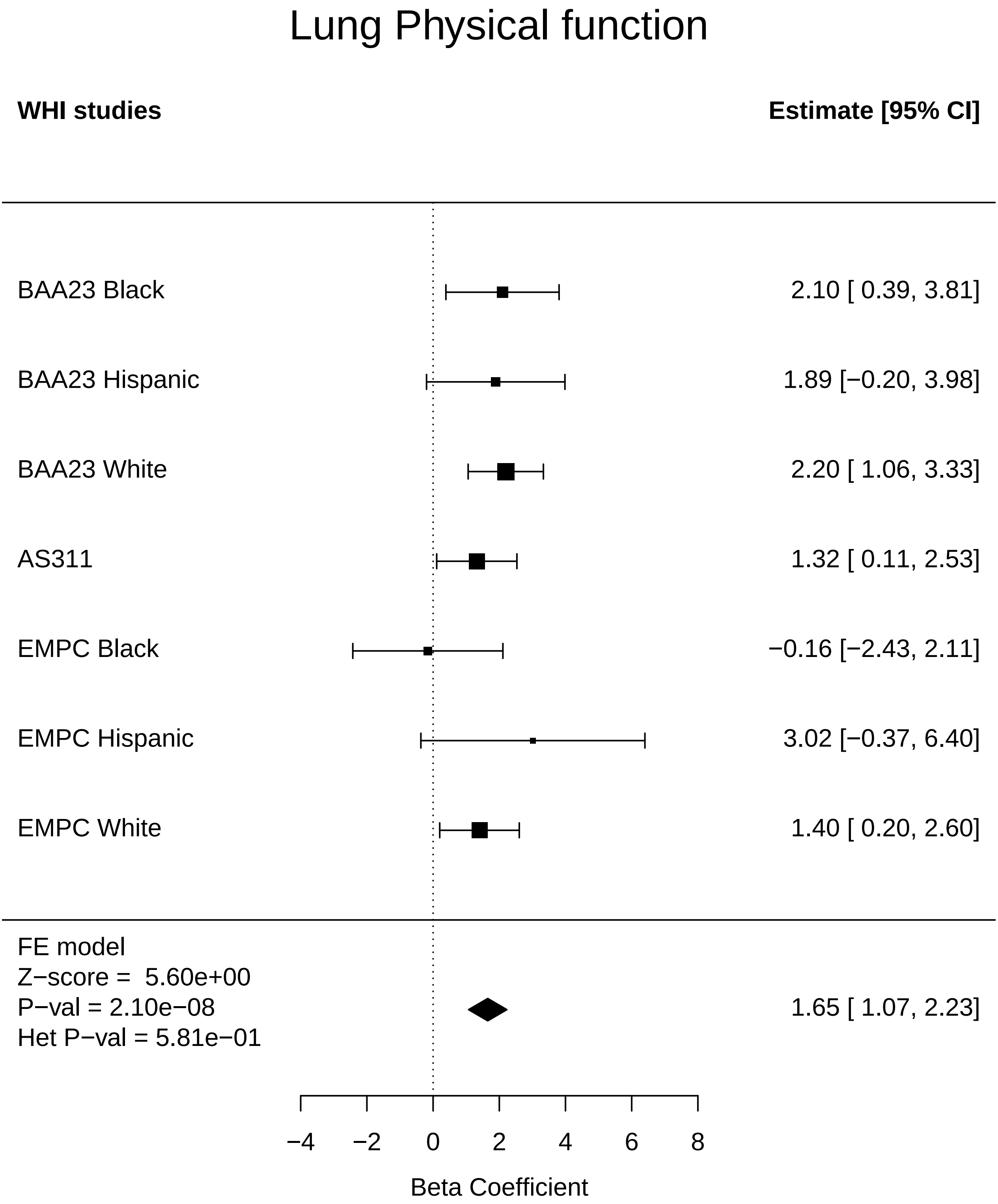

### Lung_Stroke.pdf

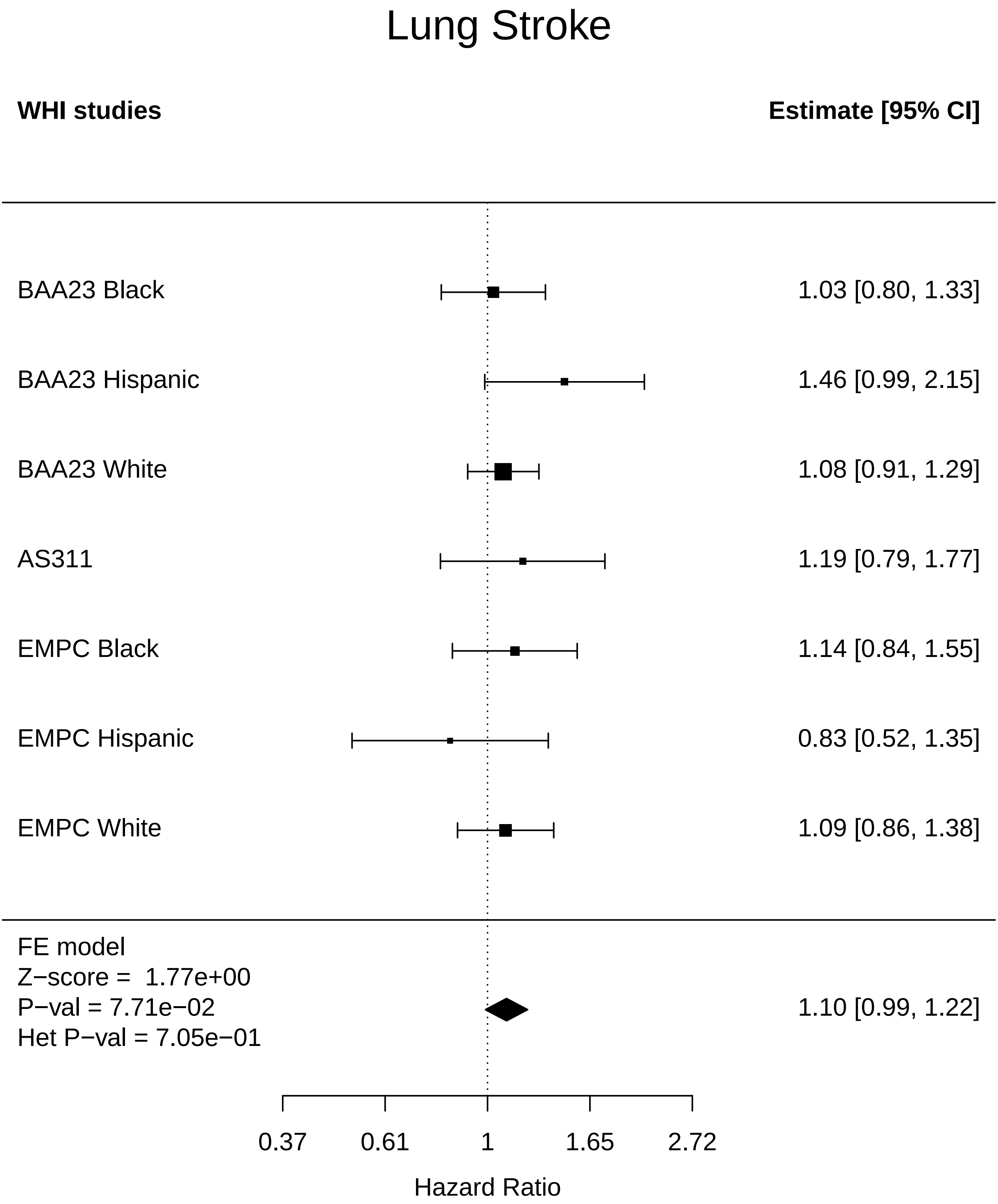

### Lung_Thyroid disease.pdf

#### Lung Thyroid disease

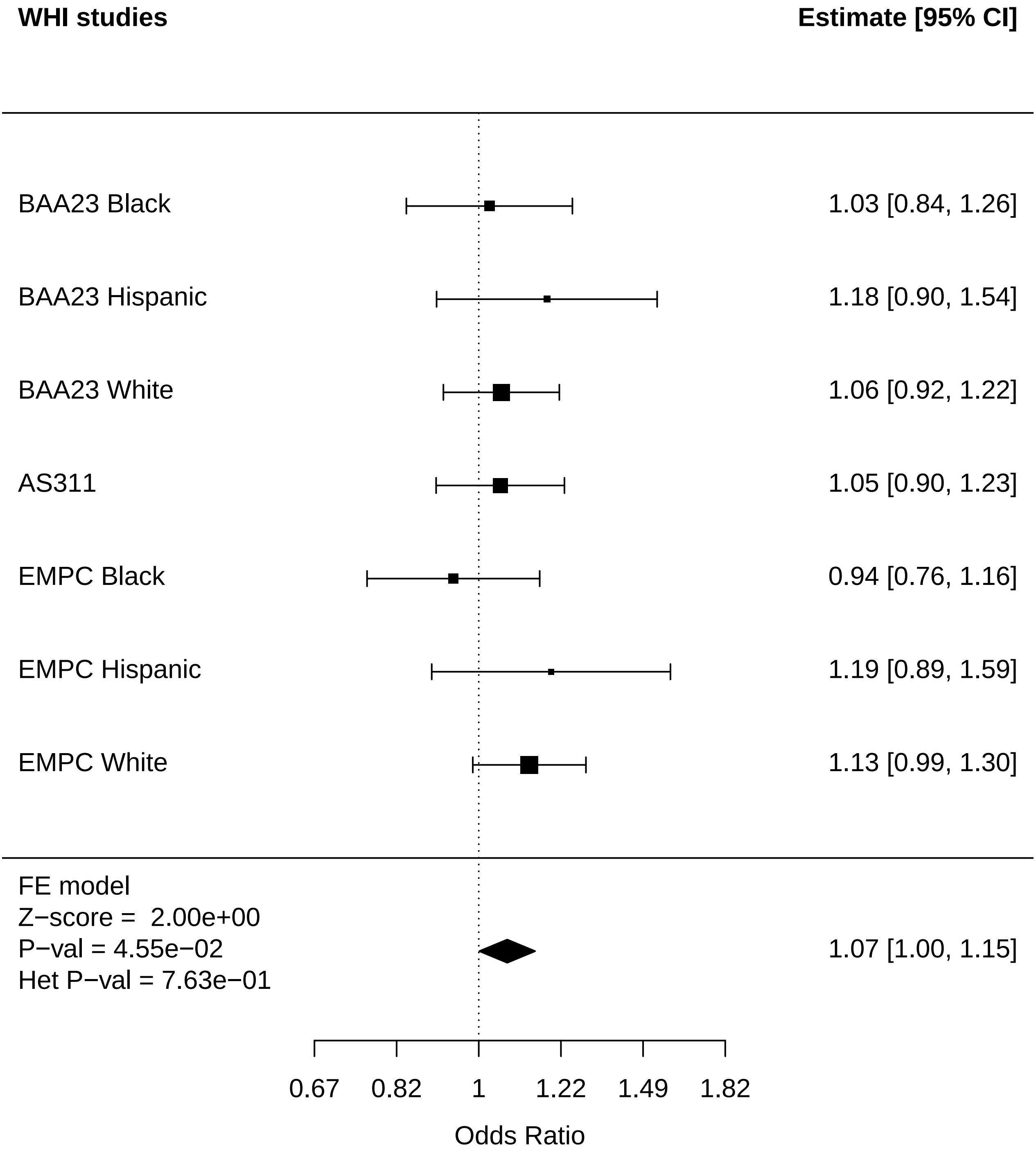

### Lung_Total comorbidities.pdf

#### Lung Total comorbidities

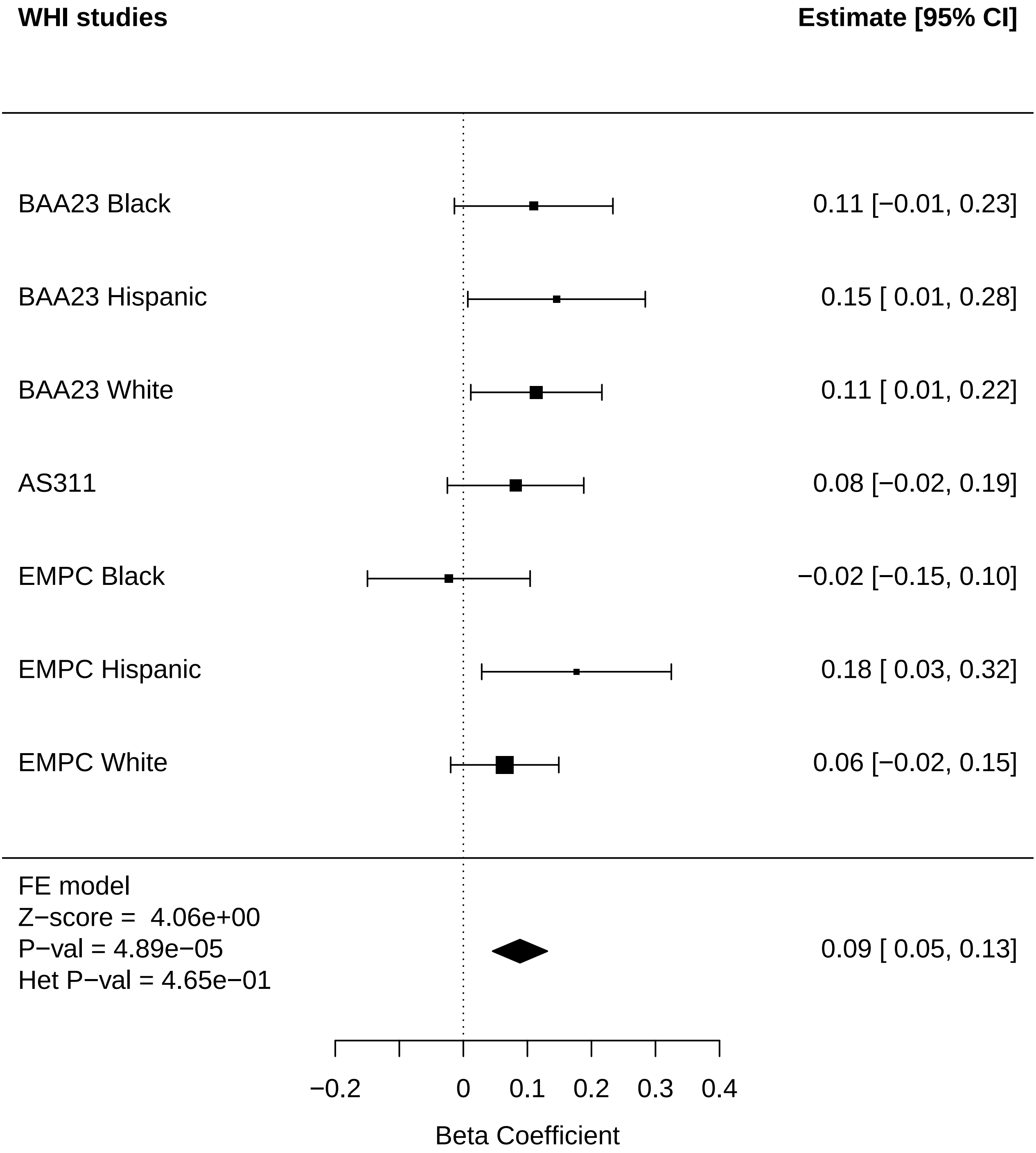

### Metabolic_Cataract.pdf

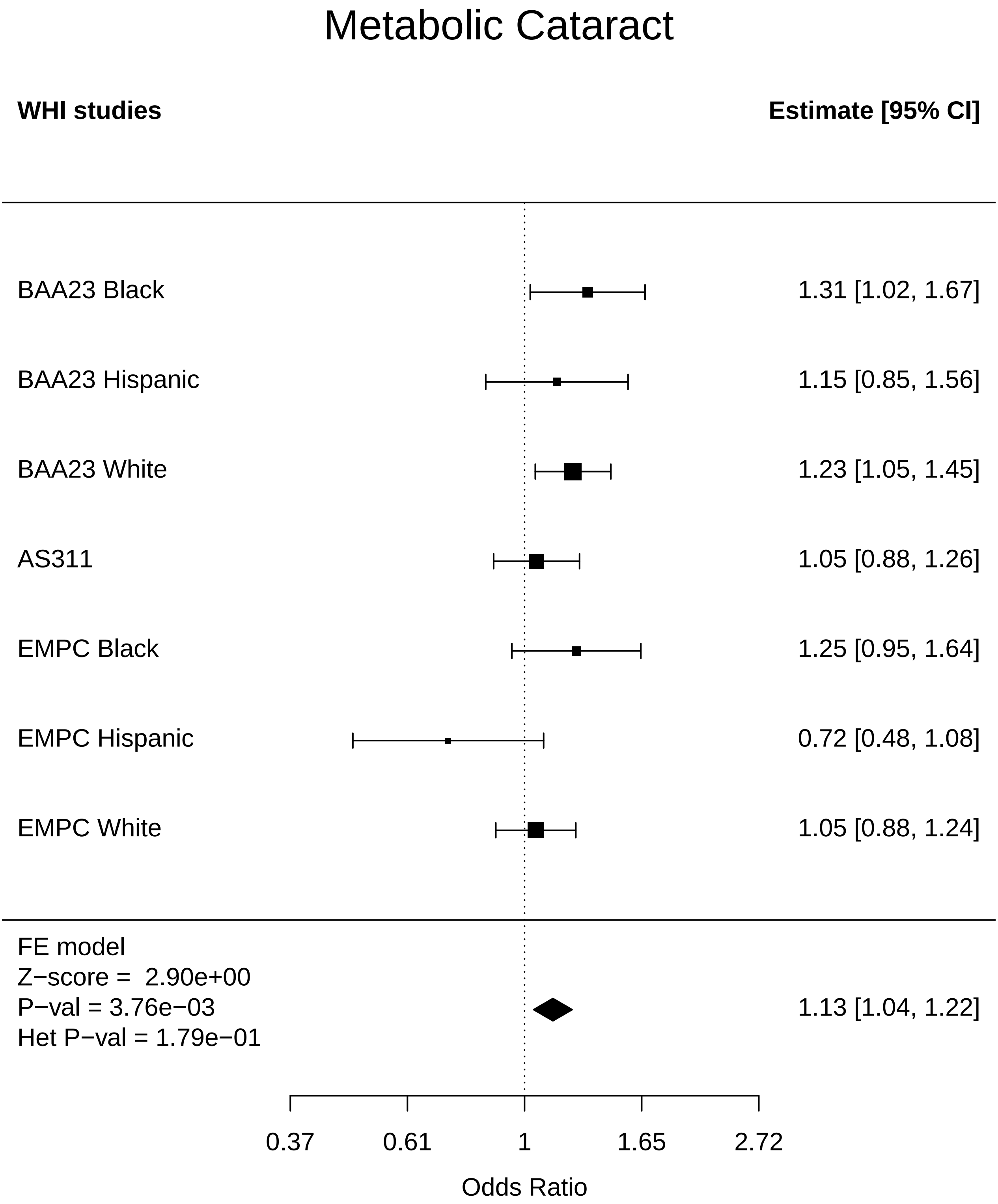

### Metabolic_Cognitive function.pdf

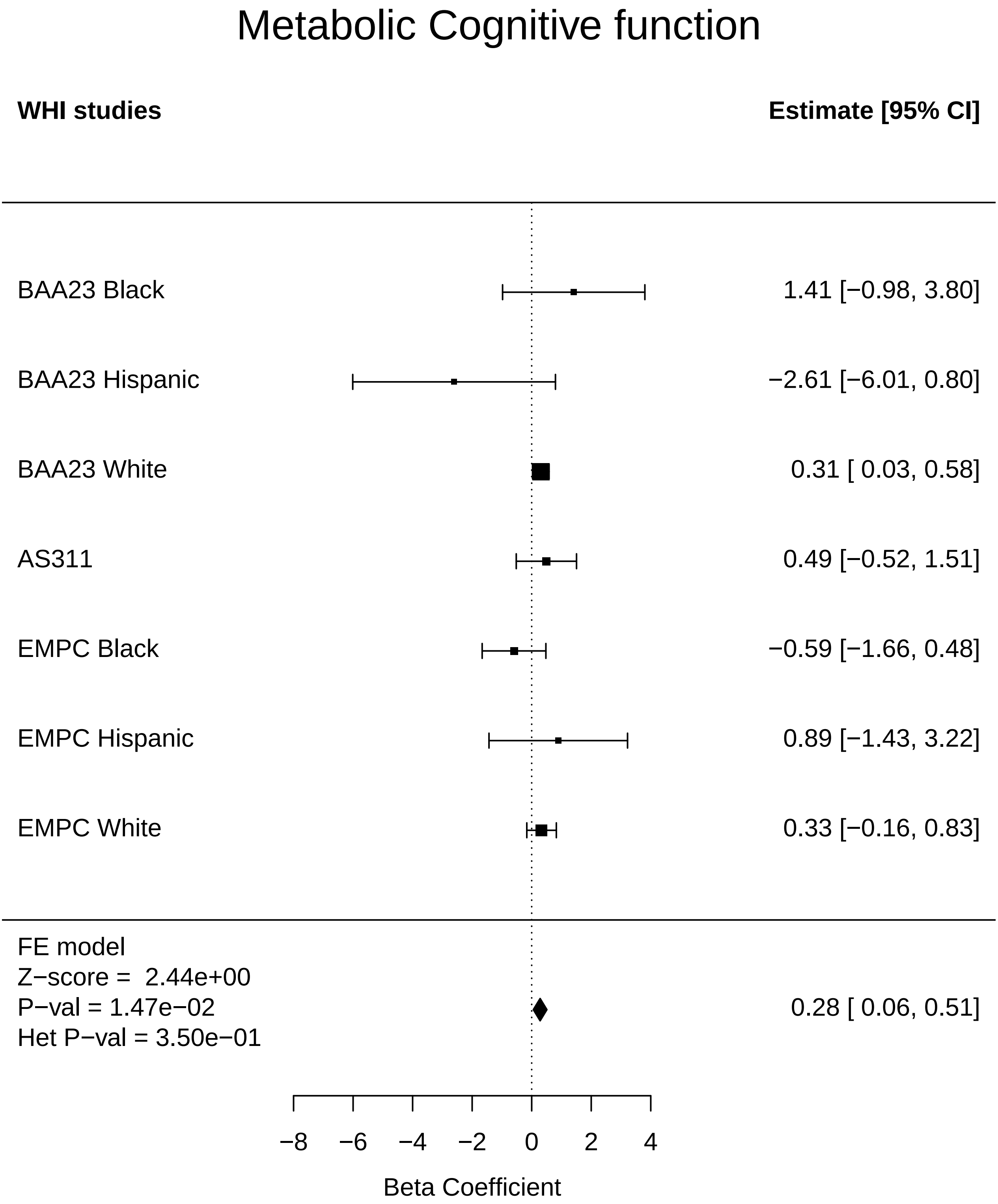

### Metabolic_Colorectal cancer.pdf

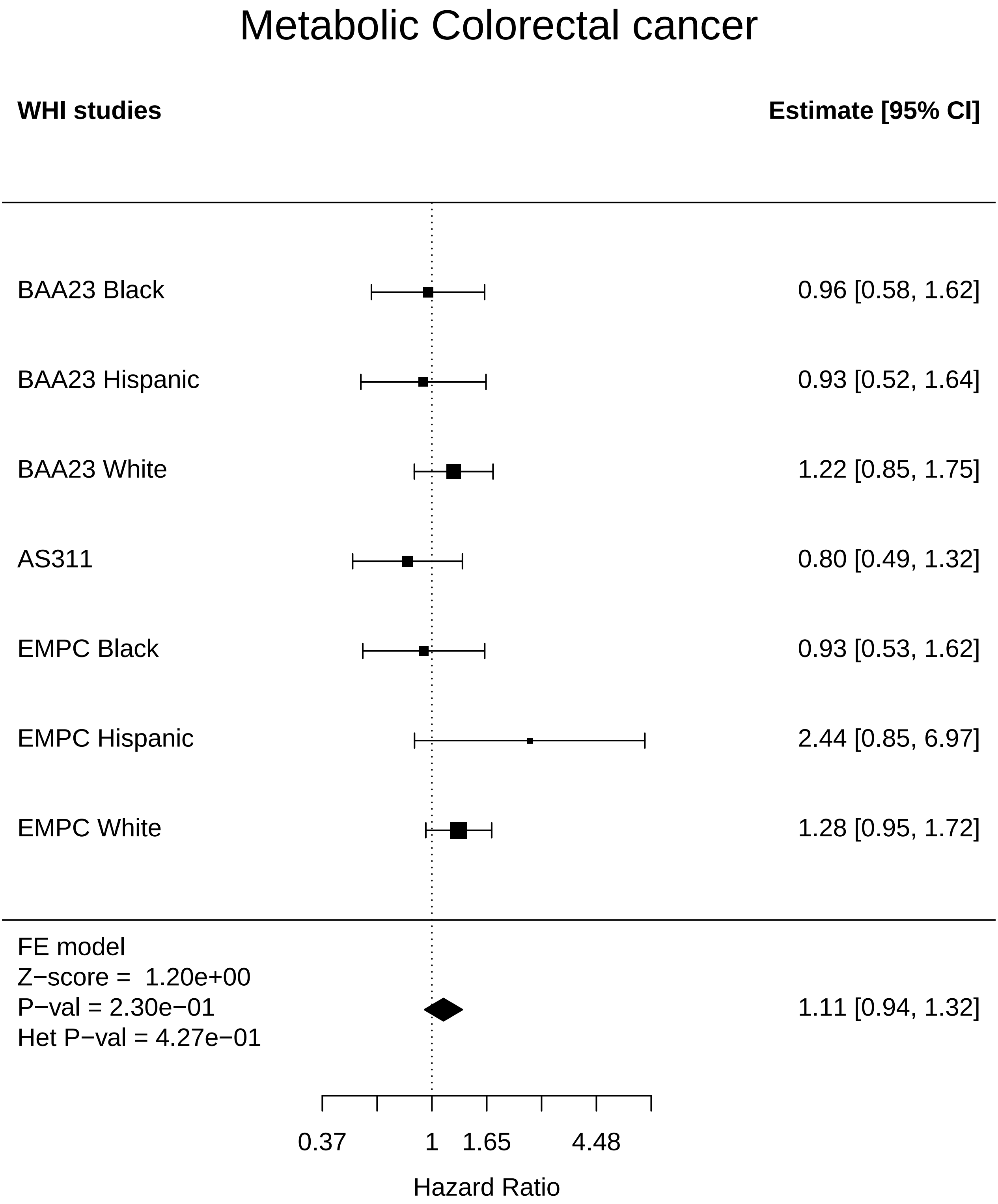

### Metabolic_Leukemia.pdf

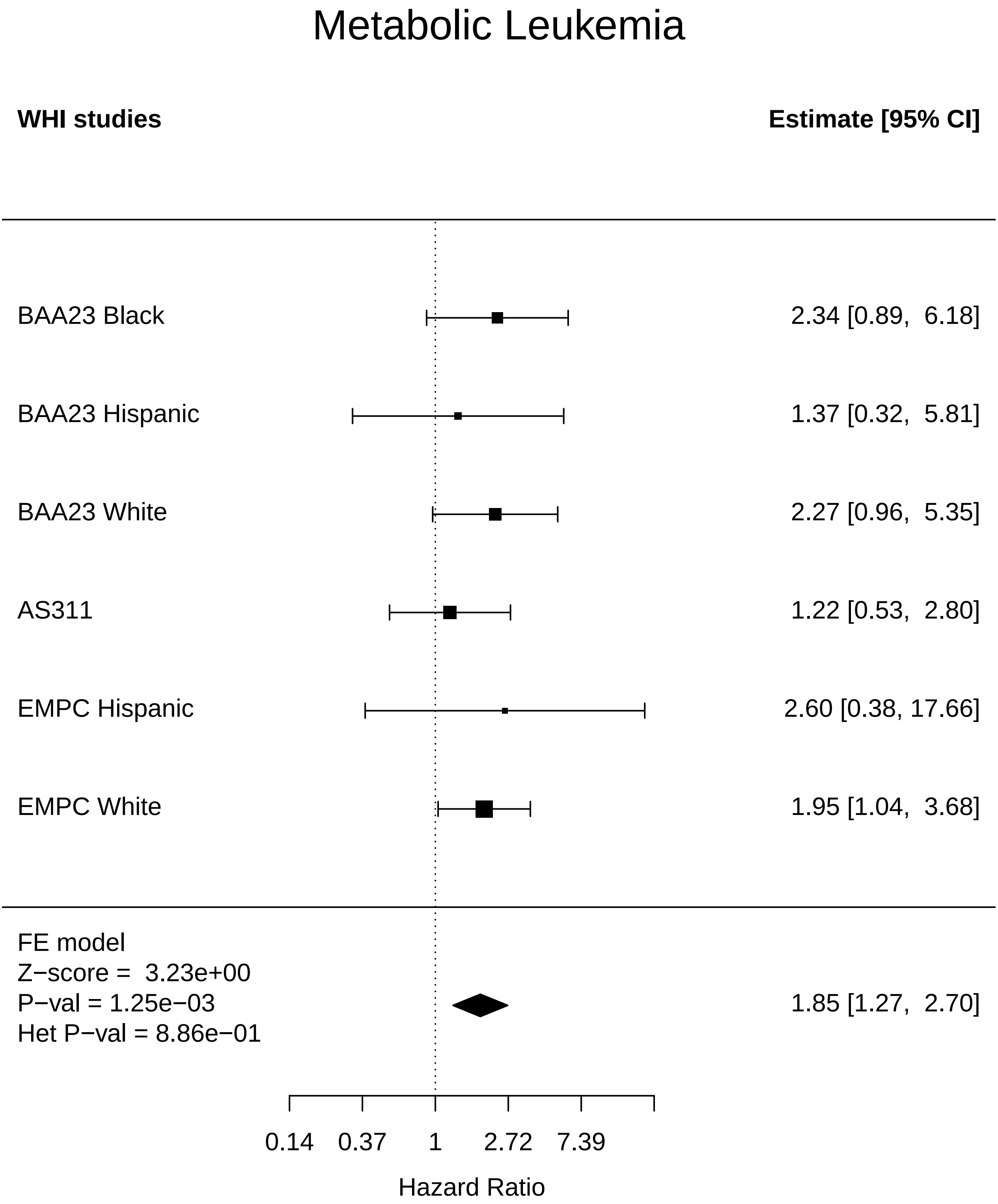

### Metabolic_Physical function.pdf

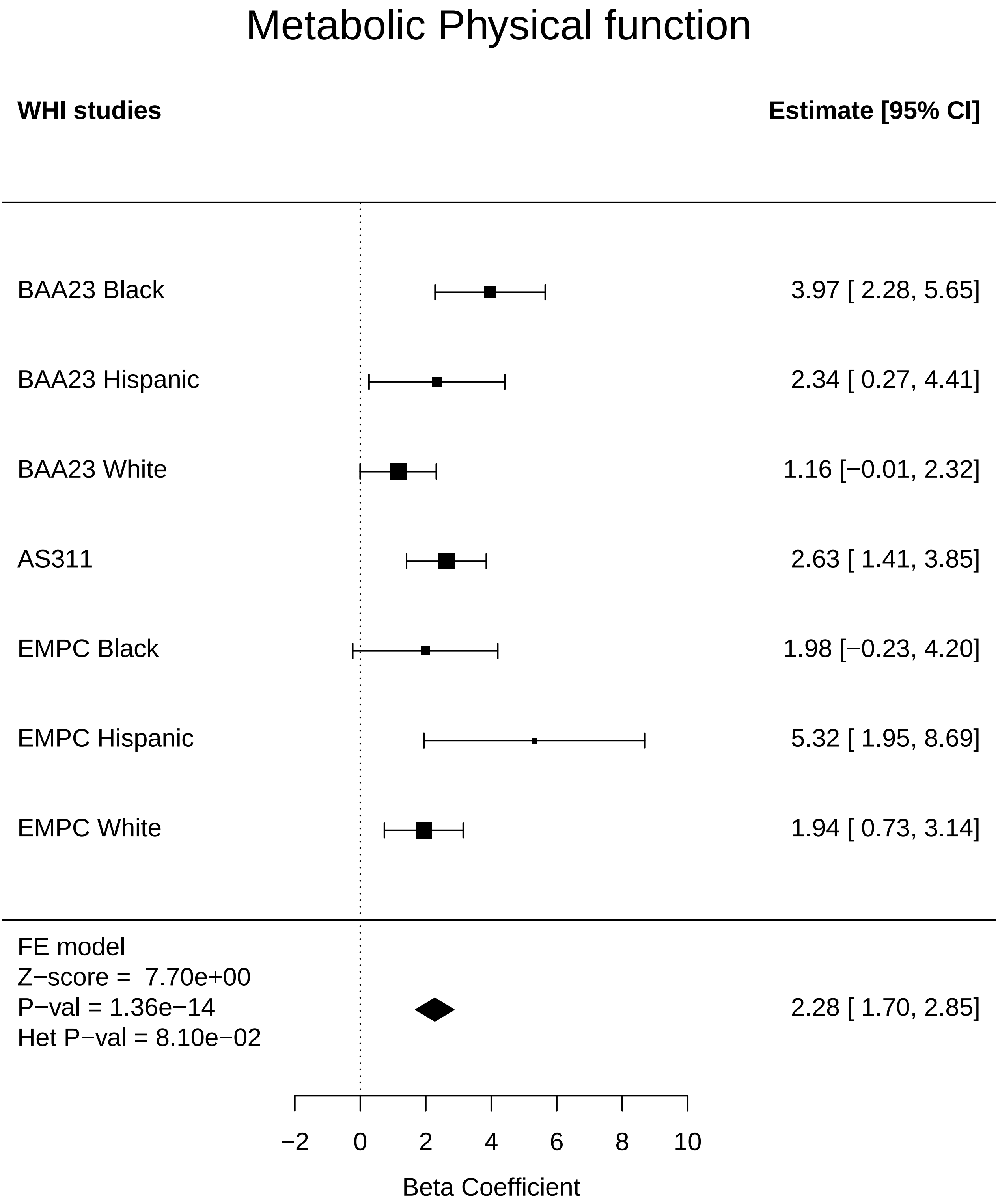

### Metabolic_Total comorbidities.pdf

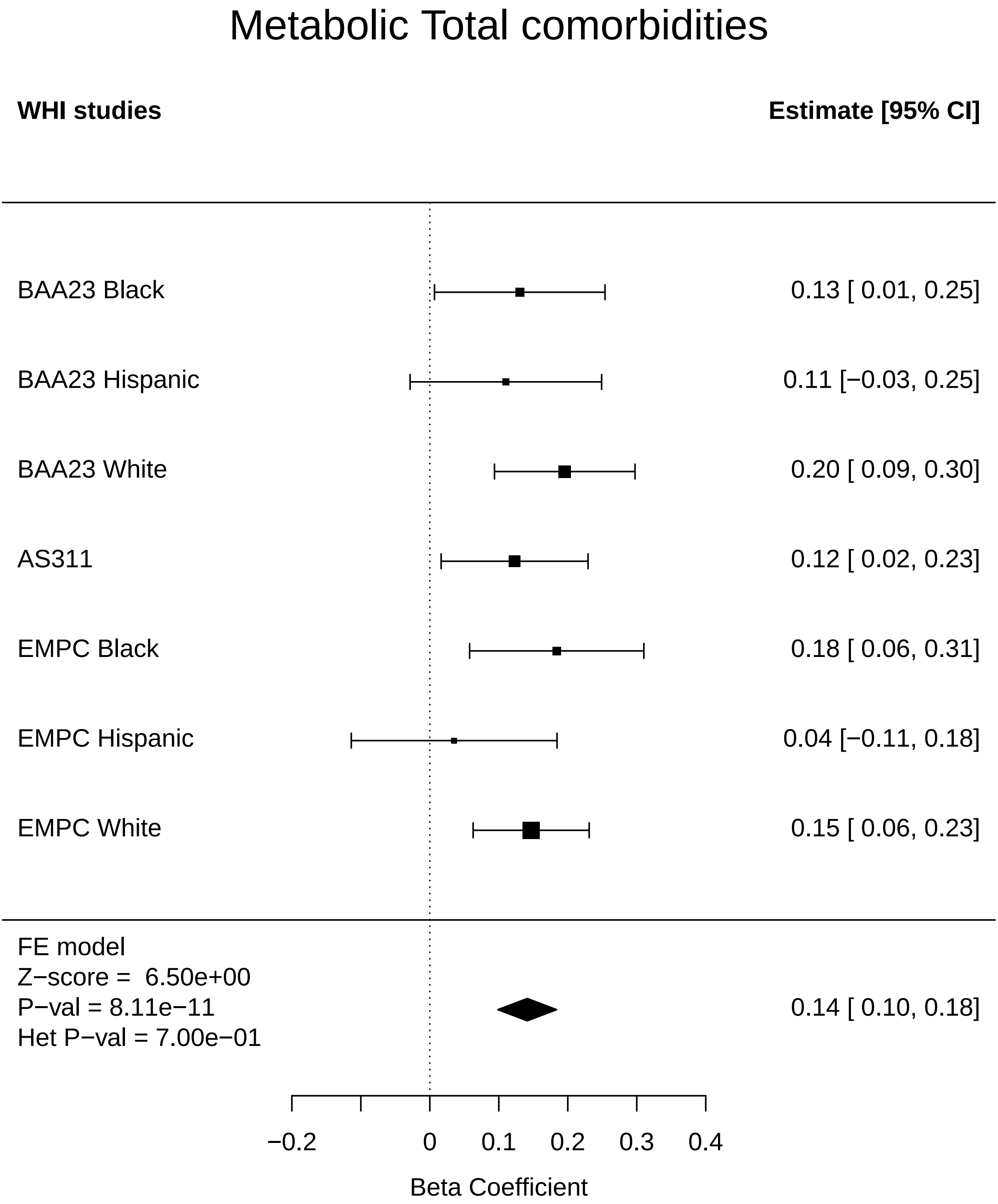

### MusculoSkeletal_Any Cancer.pdf

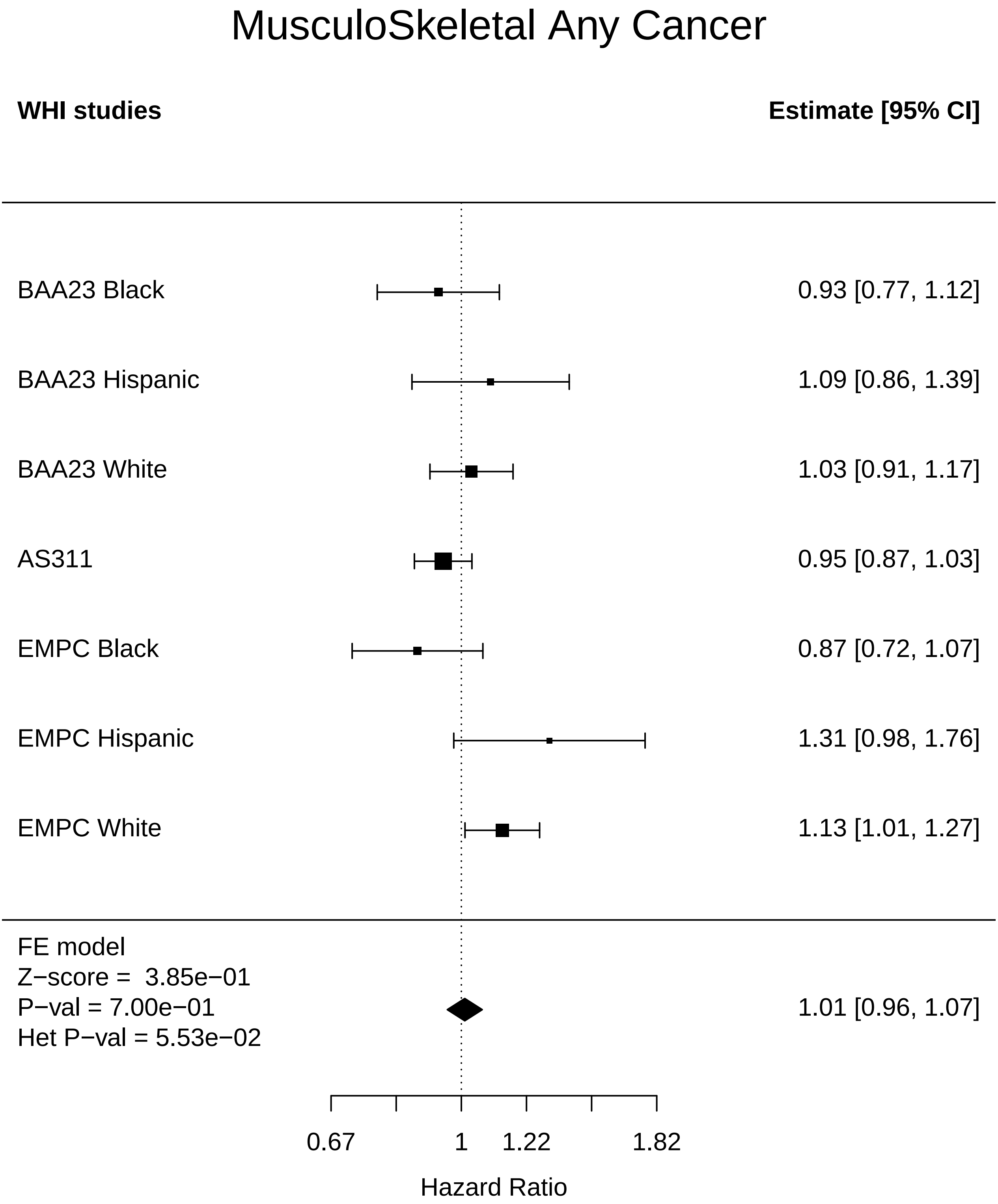

### MusculoSkeletal_Arthritis.pdf

#### MusculoSkeletal Arthritis

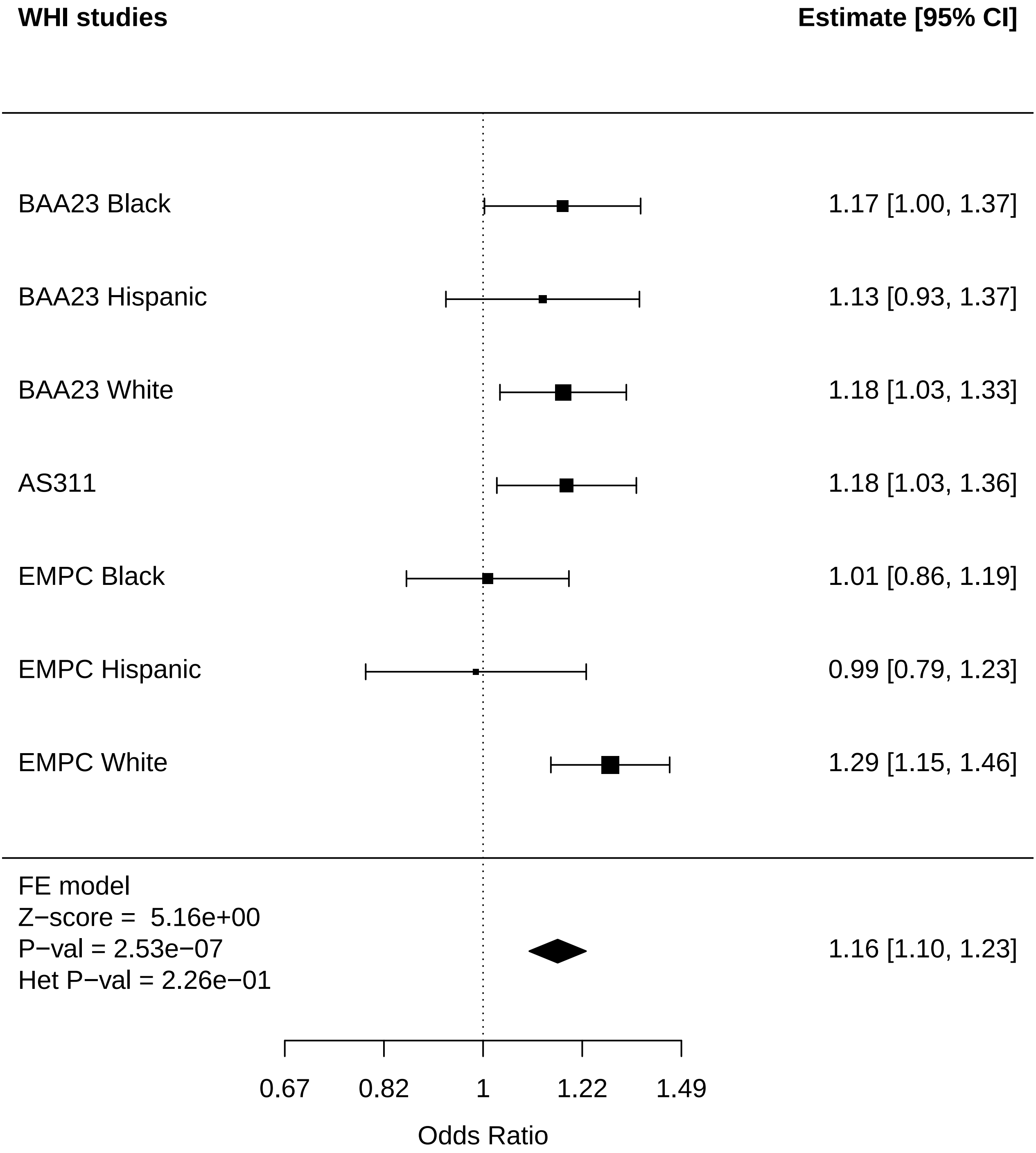

### MusculoSkeletal_Breast Cancer.pdf

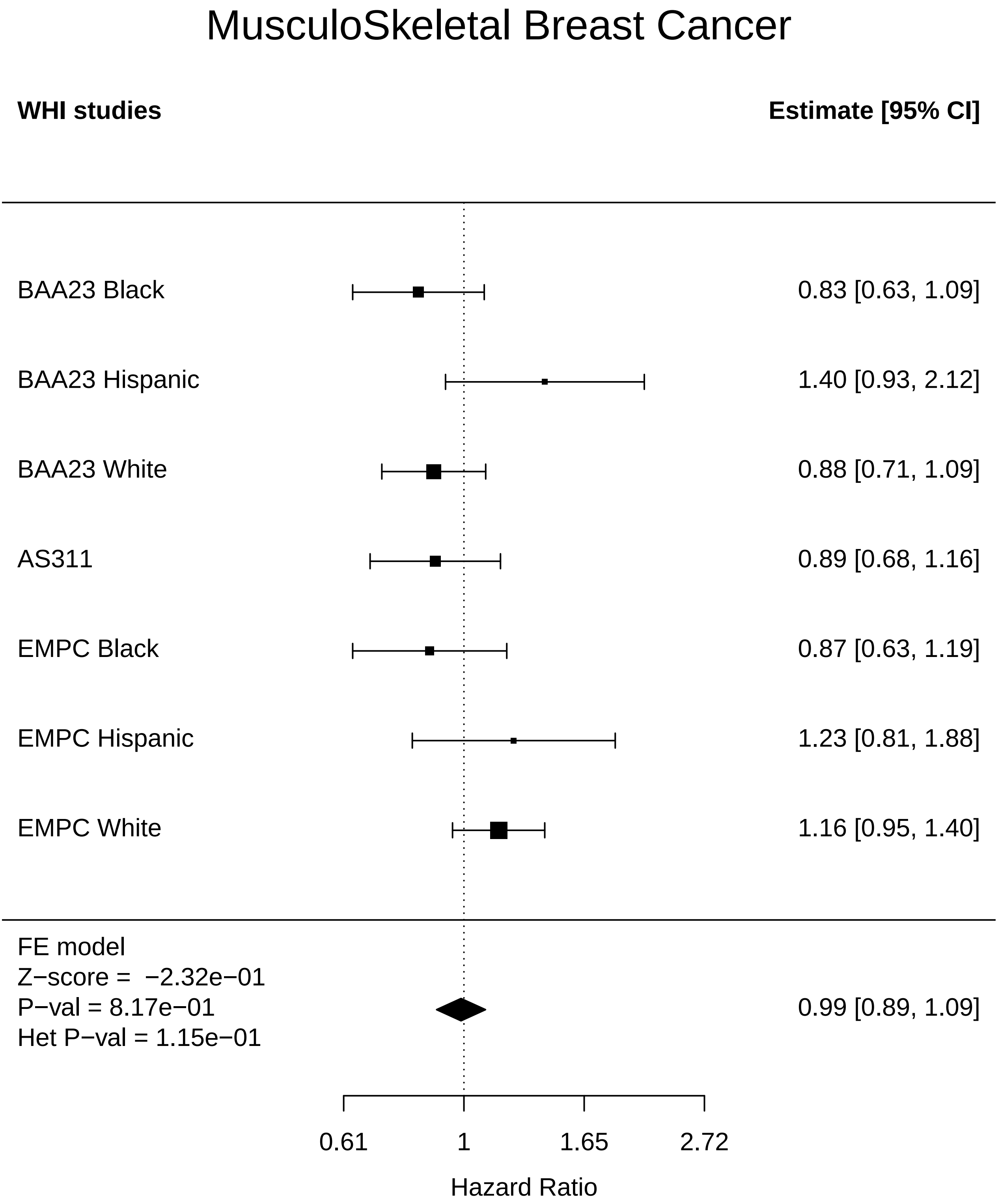

### MusculoSkeletal_Cognitive function.pdf

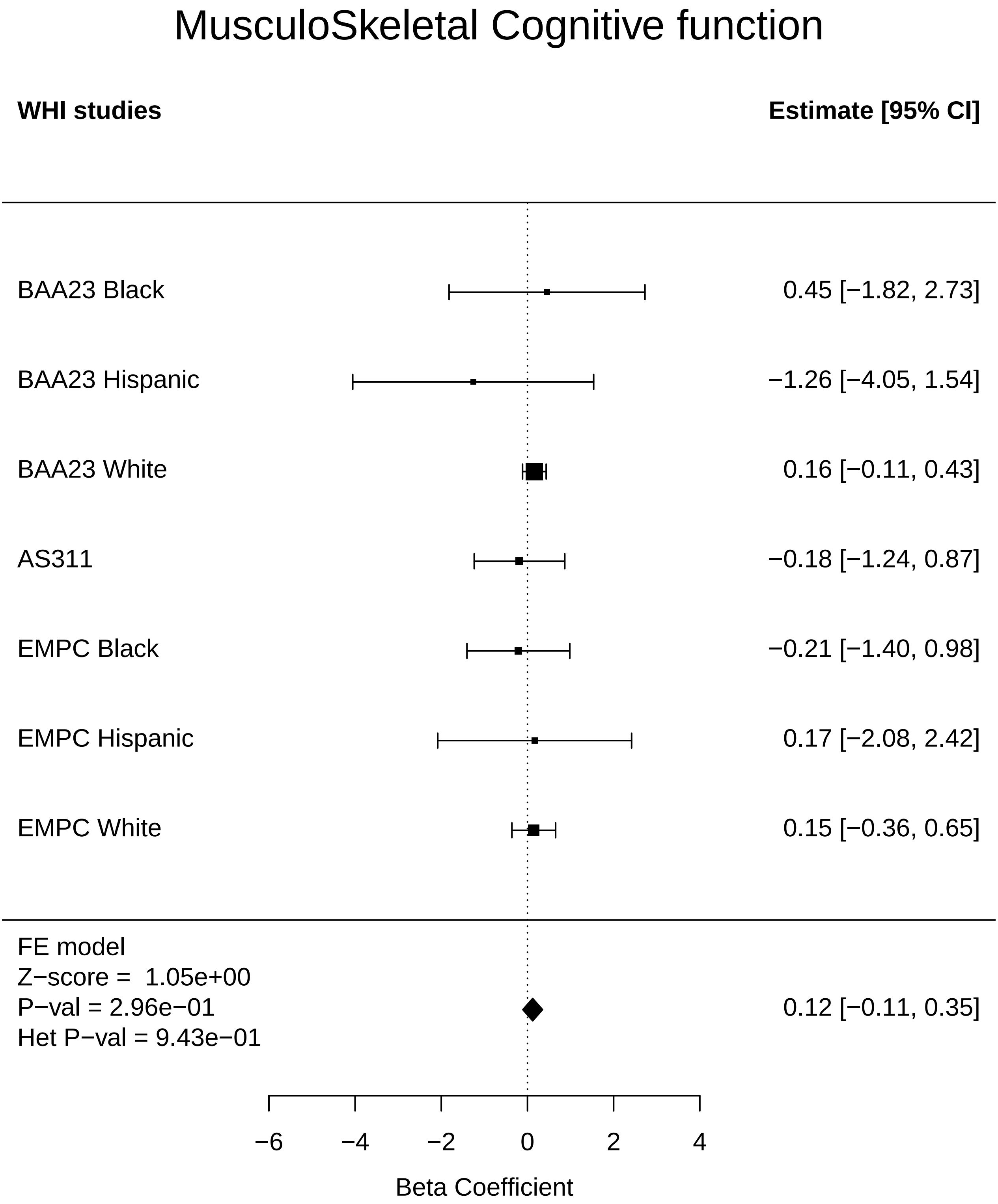

### MusculoSkeletal_Colorectal cancer.pdf

#### MusculoSkeletal Colorectal cancer

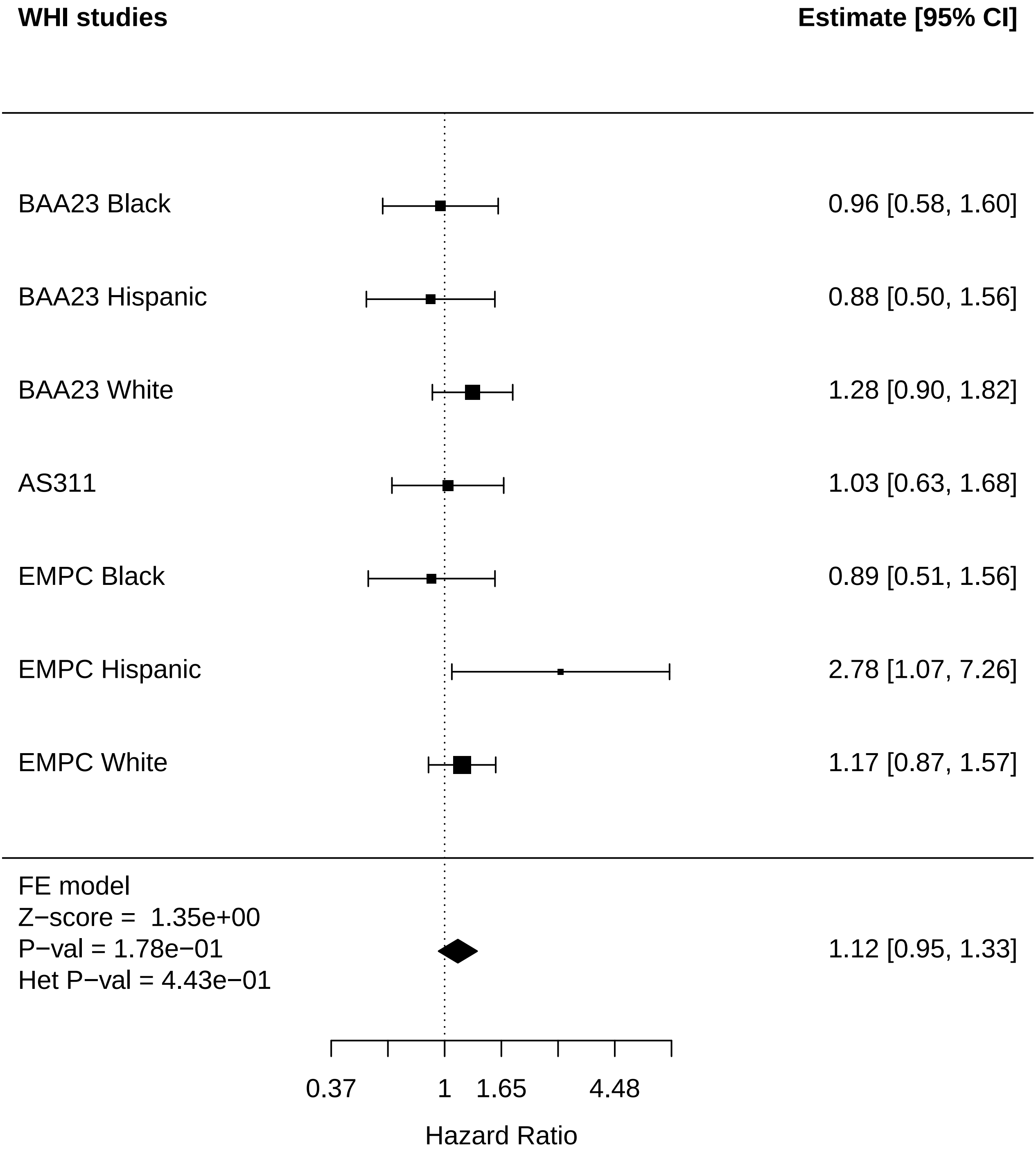

### MusculoSkeletal_Diabetes.pdf

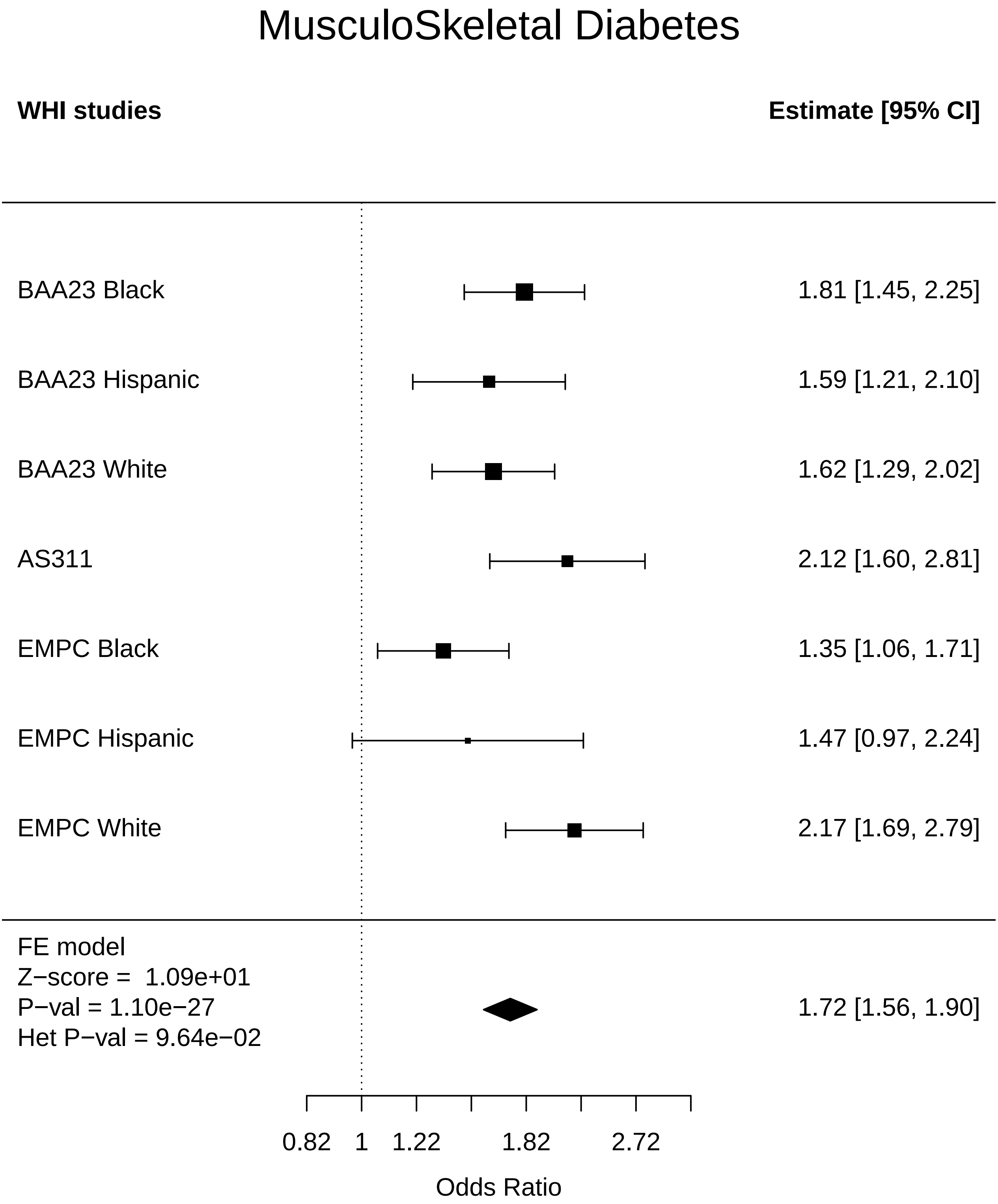

### MusculoSkeletal_Disease free.pdf

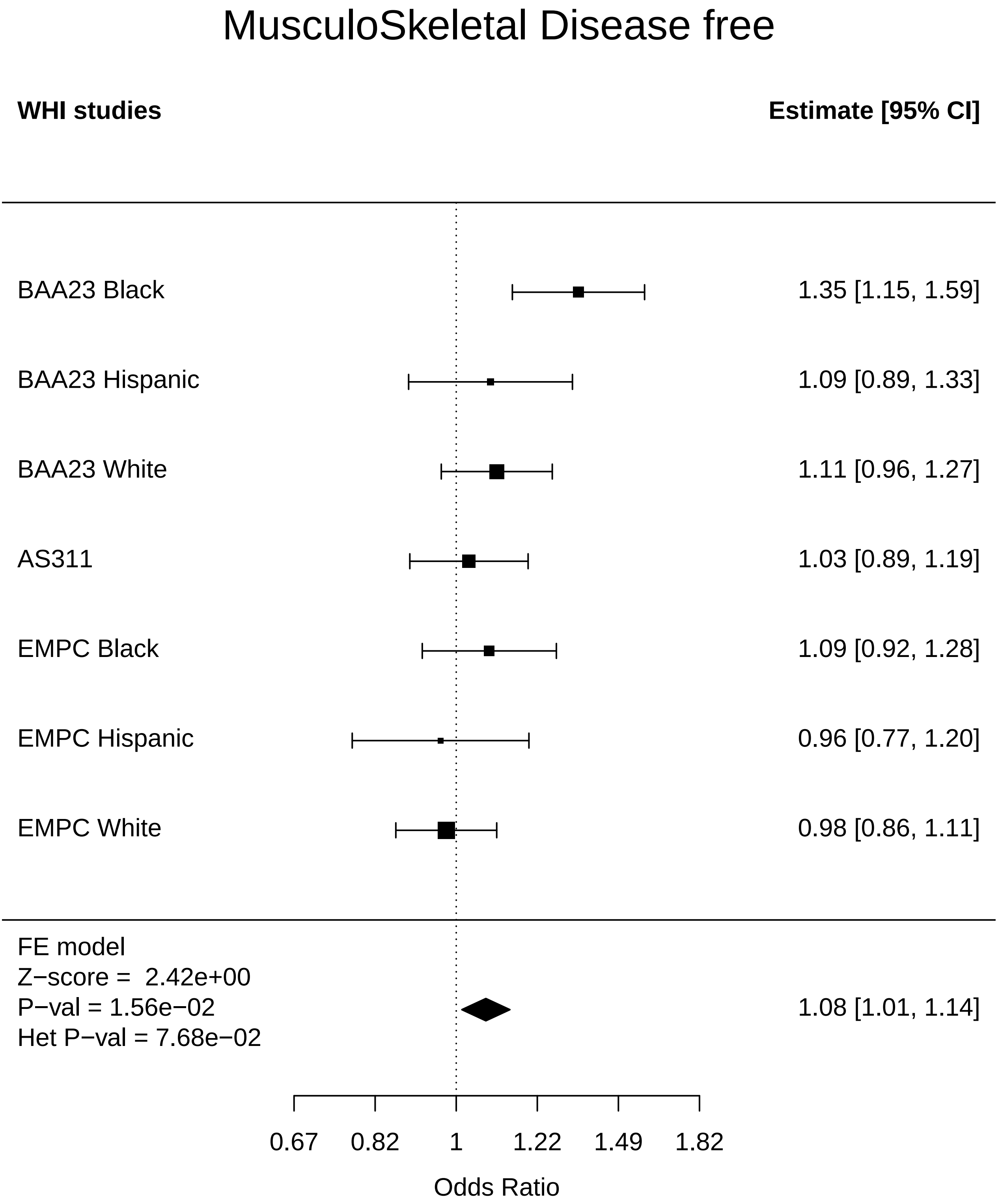

### MusculoSkeletal_Endometrial cancer.pdf

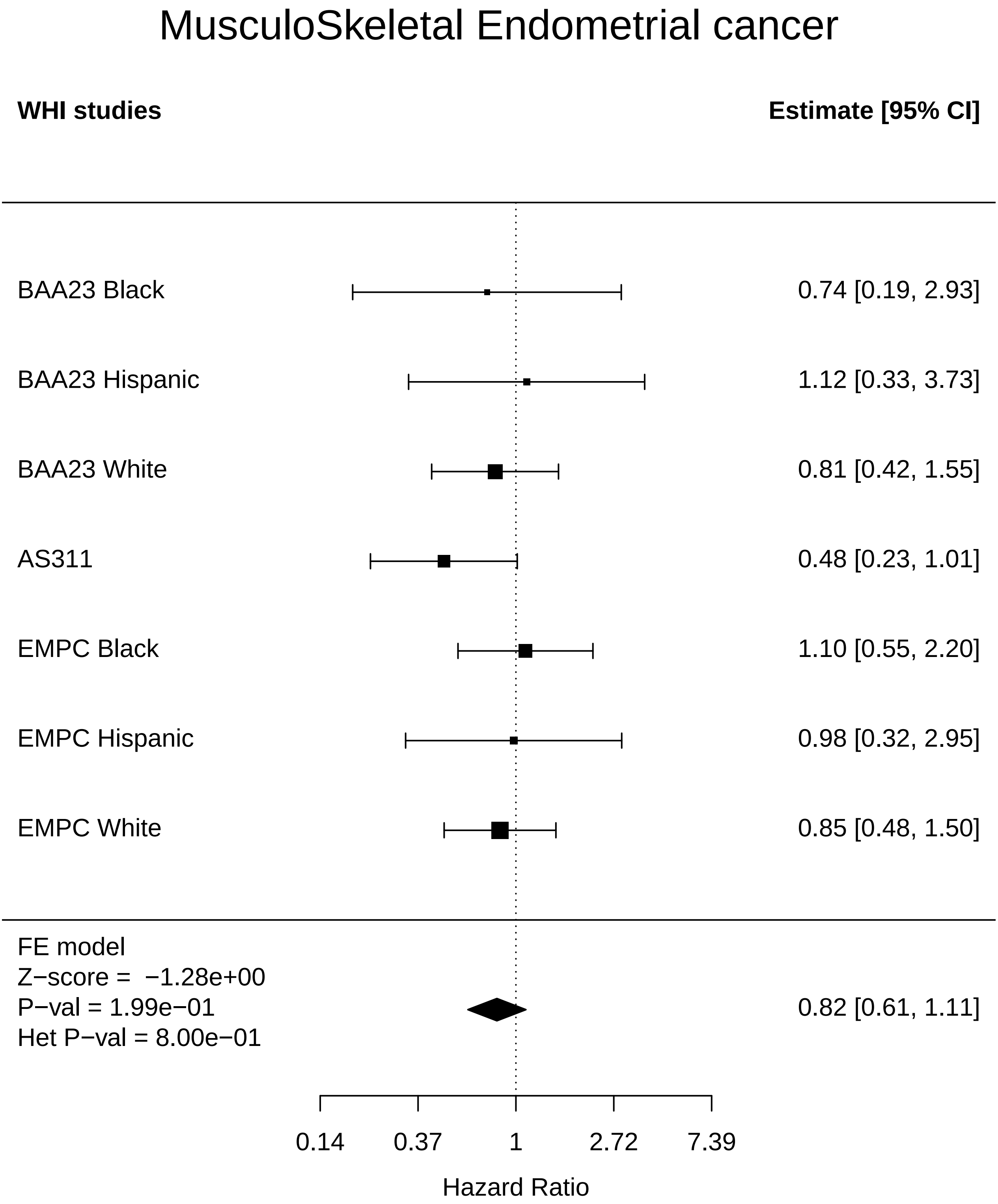

### MusculoSkeletal_Leukemia.pdf

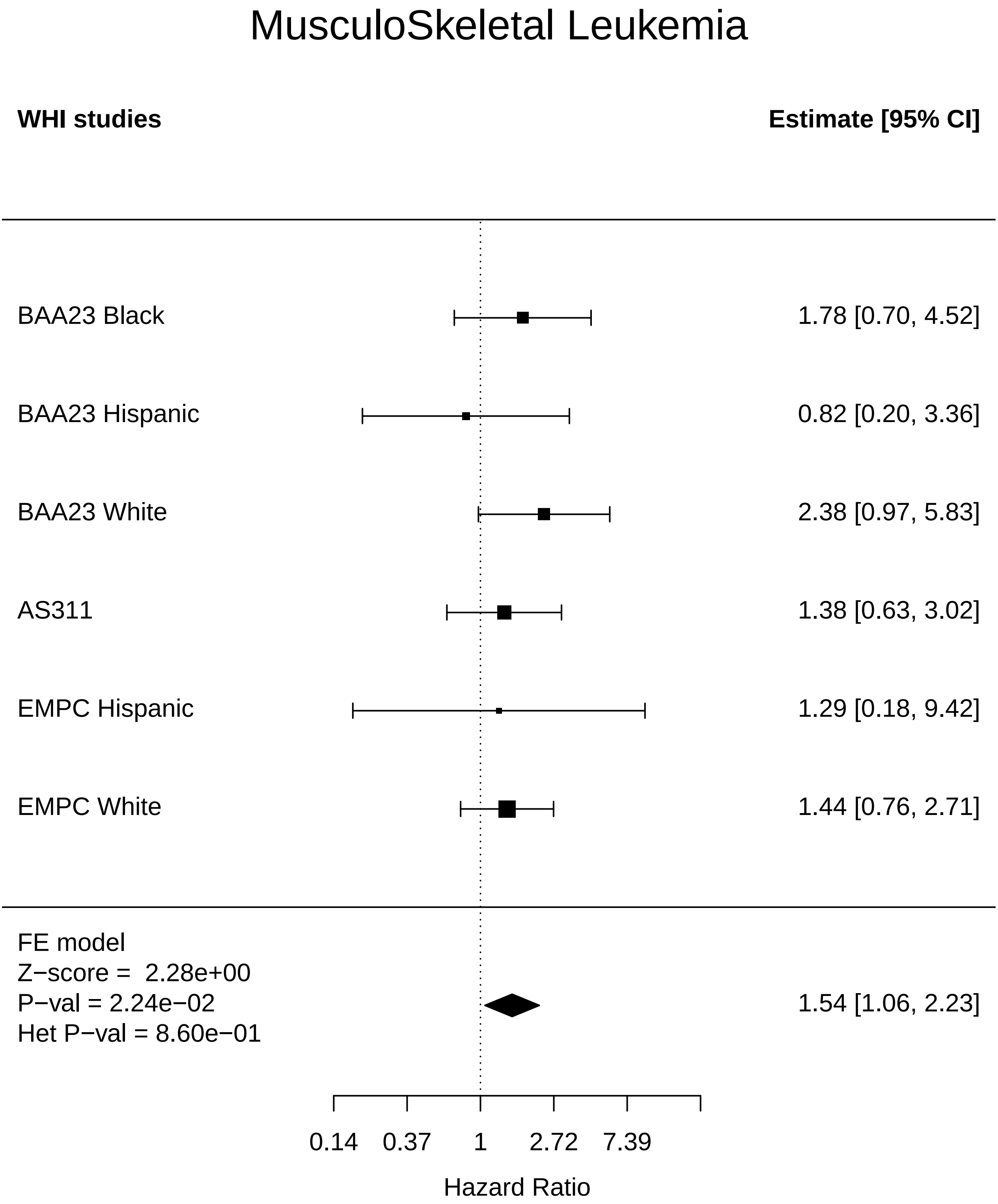

### MusculoSkeletal_Lung Cancer.pdf

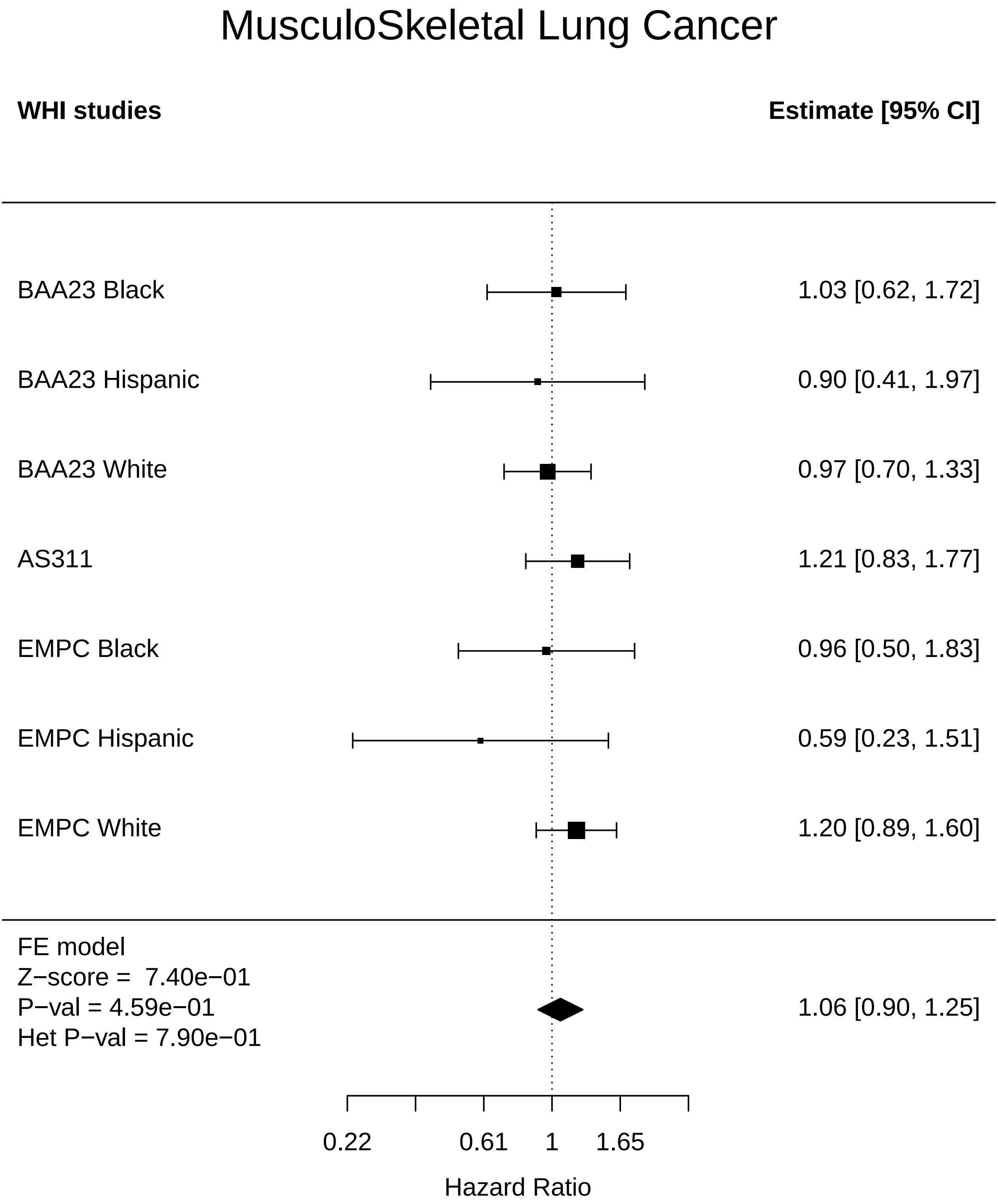

### MusculoSkeletal_MI.pdf

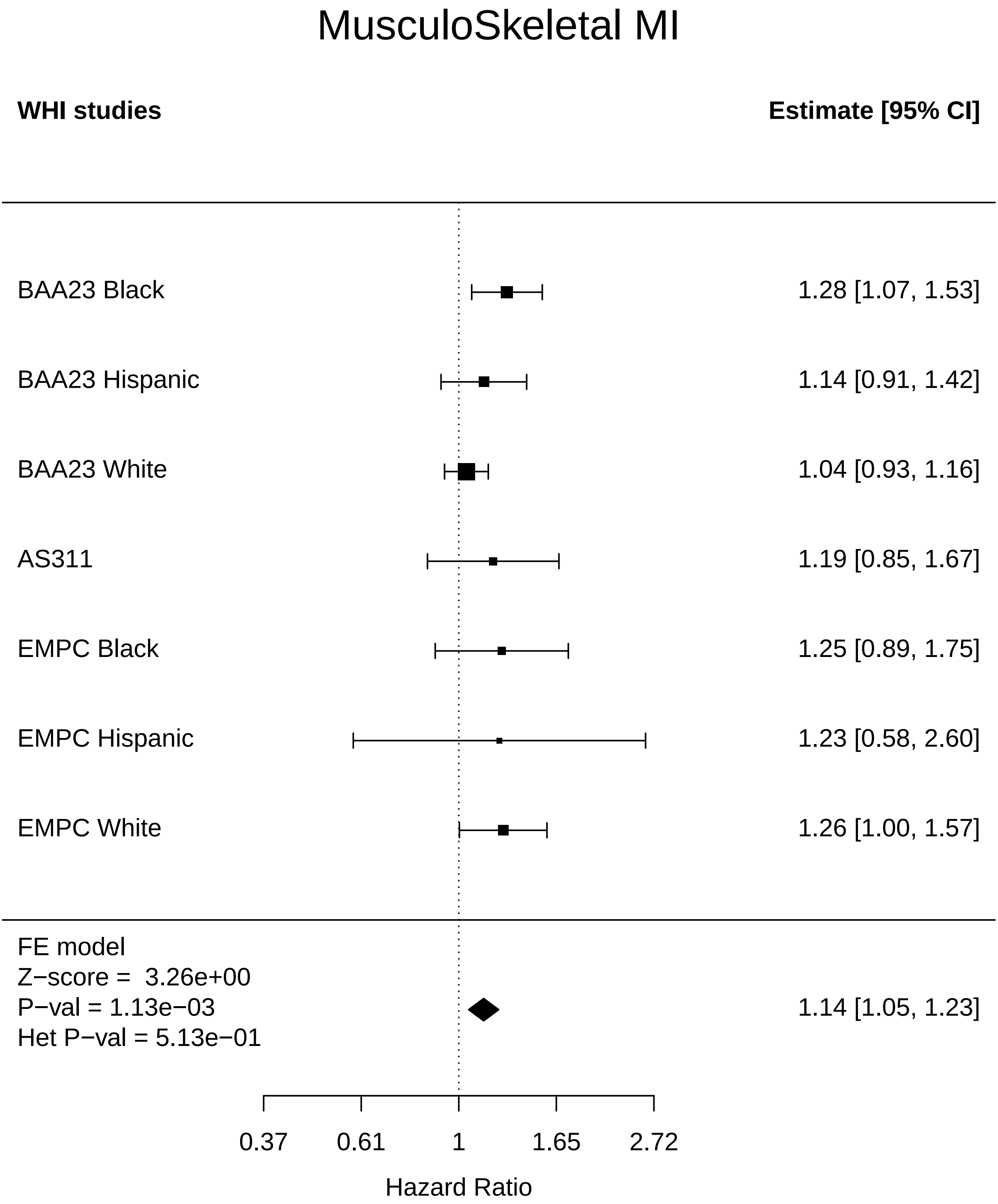

### MusculoSkeletal_Mortality.pdf

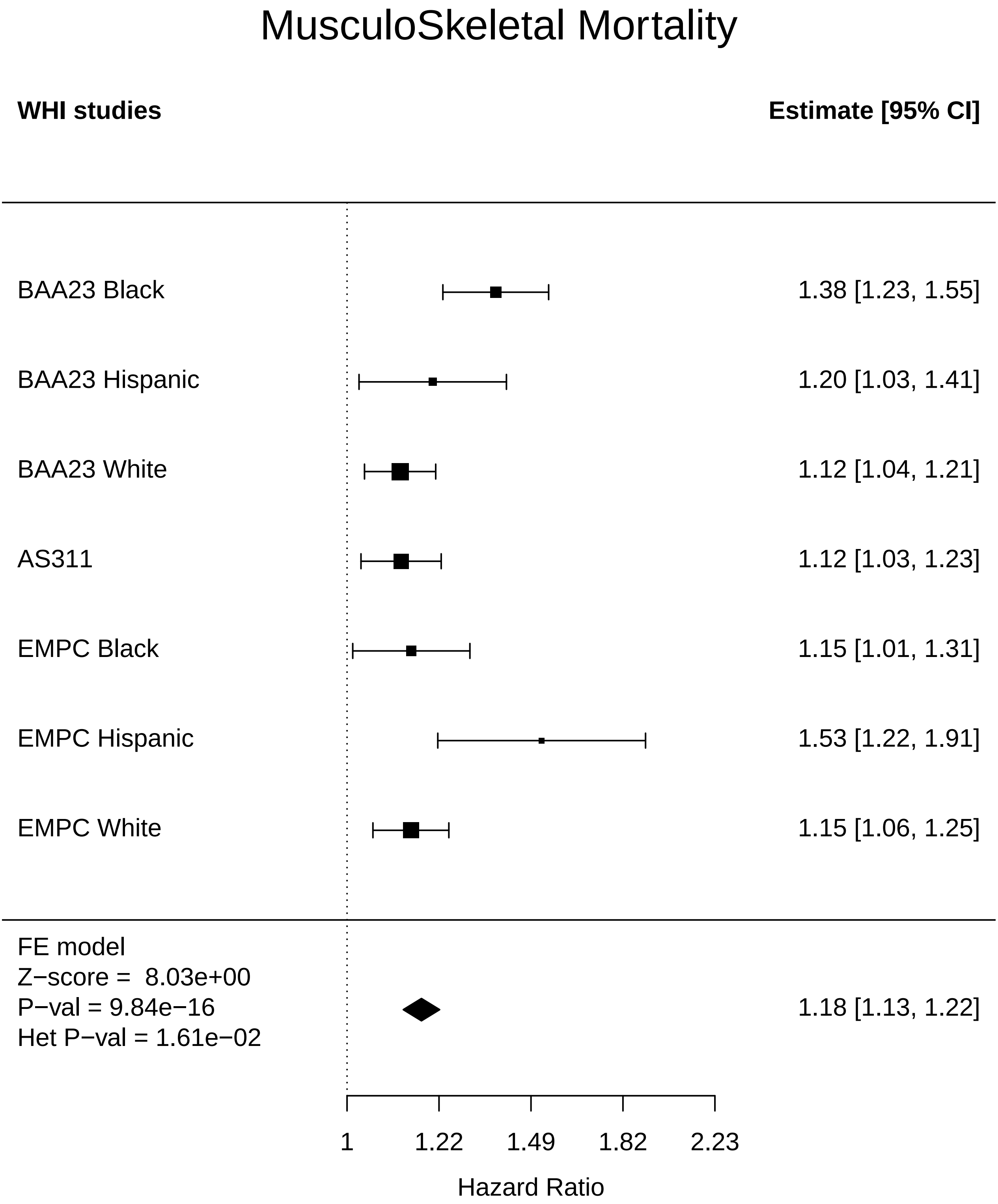

### MusculoSkeletal_Stroke.pdf

#### MusculoSkeletal Stroke

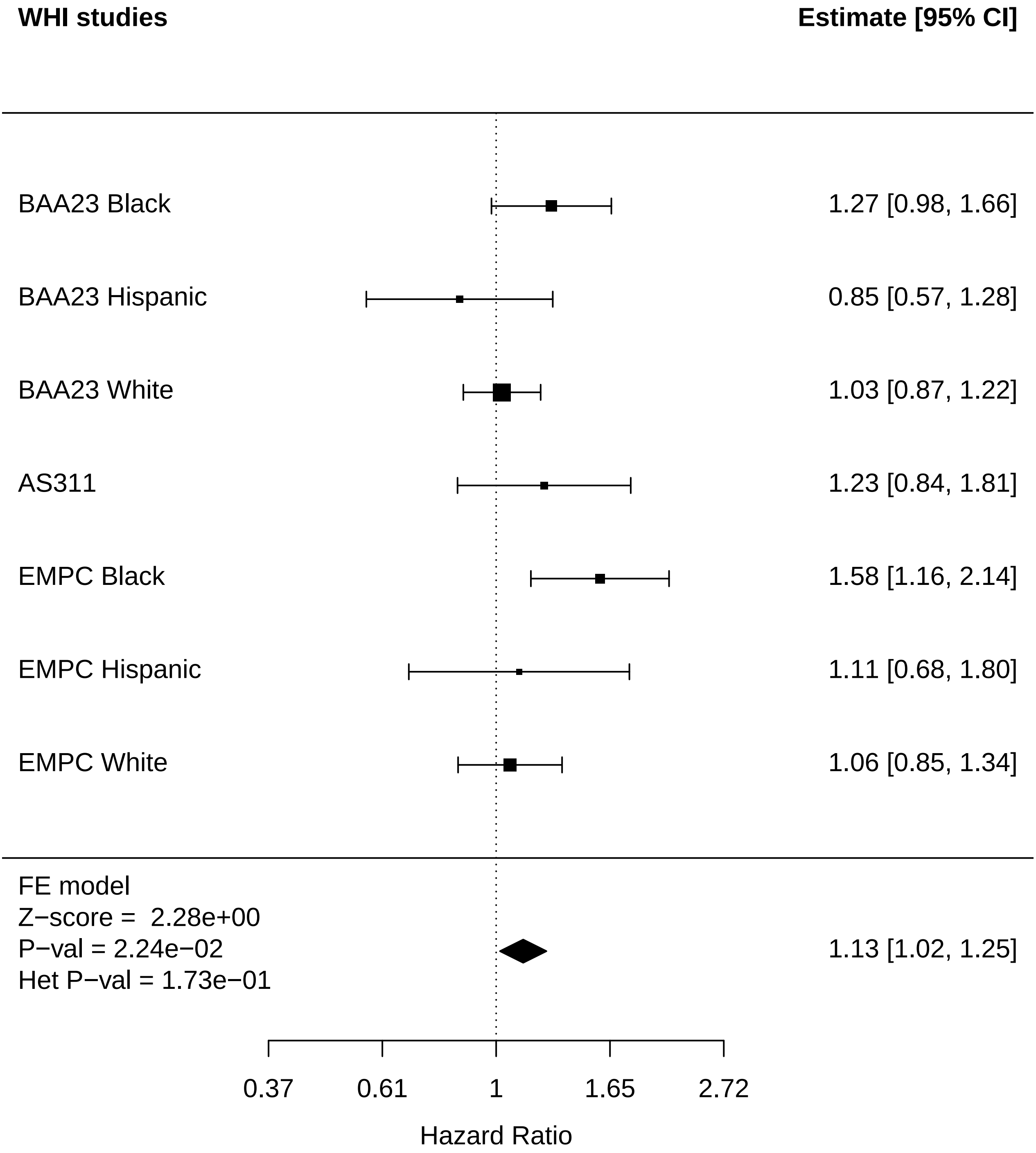

### MusculoSkeletal_Thyroid disease.pdf

#### MusculoSkeletal Thyroid disease

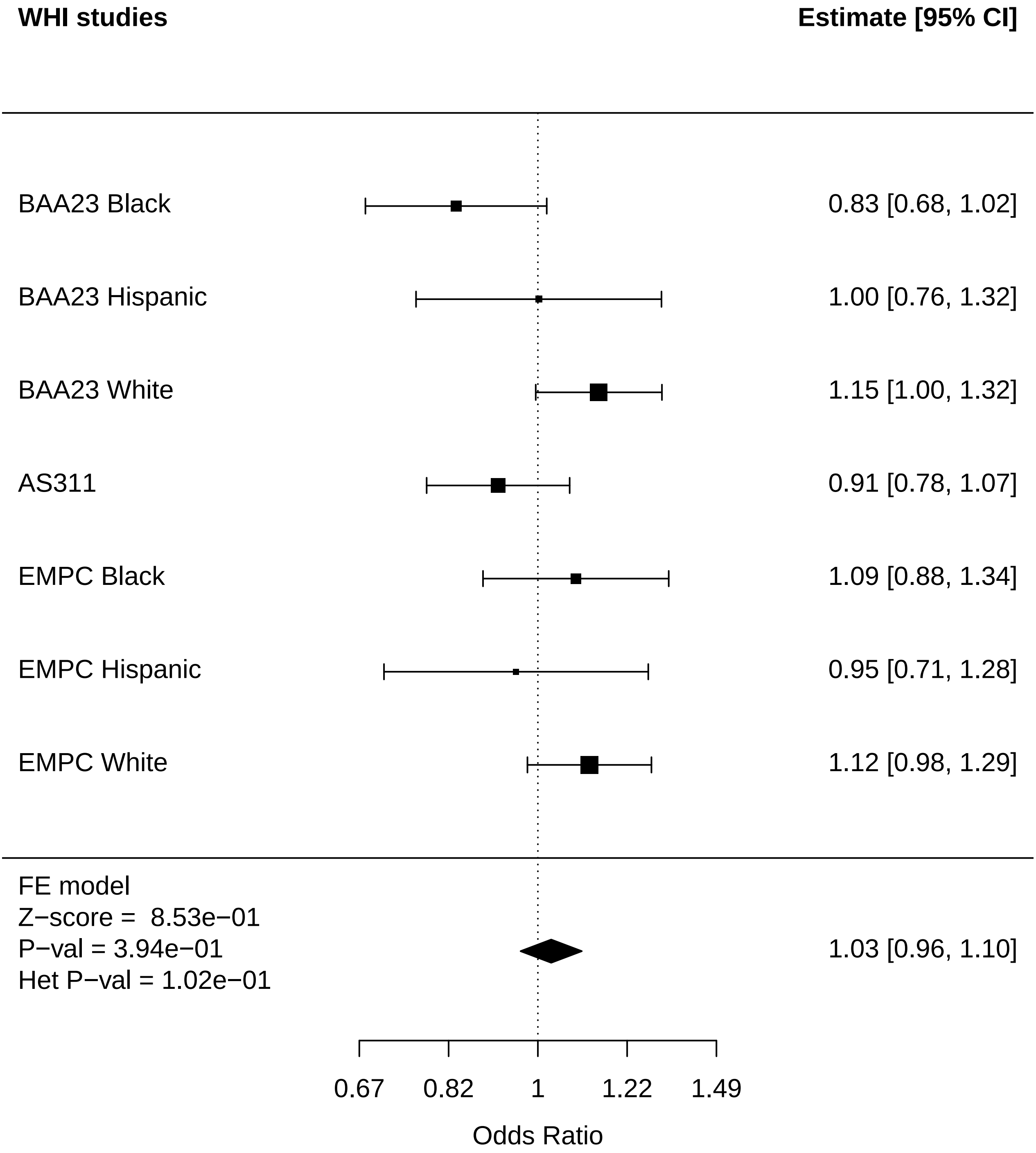

### MusculoSkeletal_Total comorbidities.pdf

#### MusculoSkeletal Total comorbidities

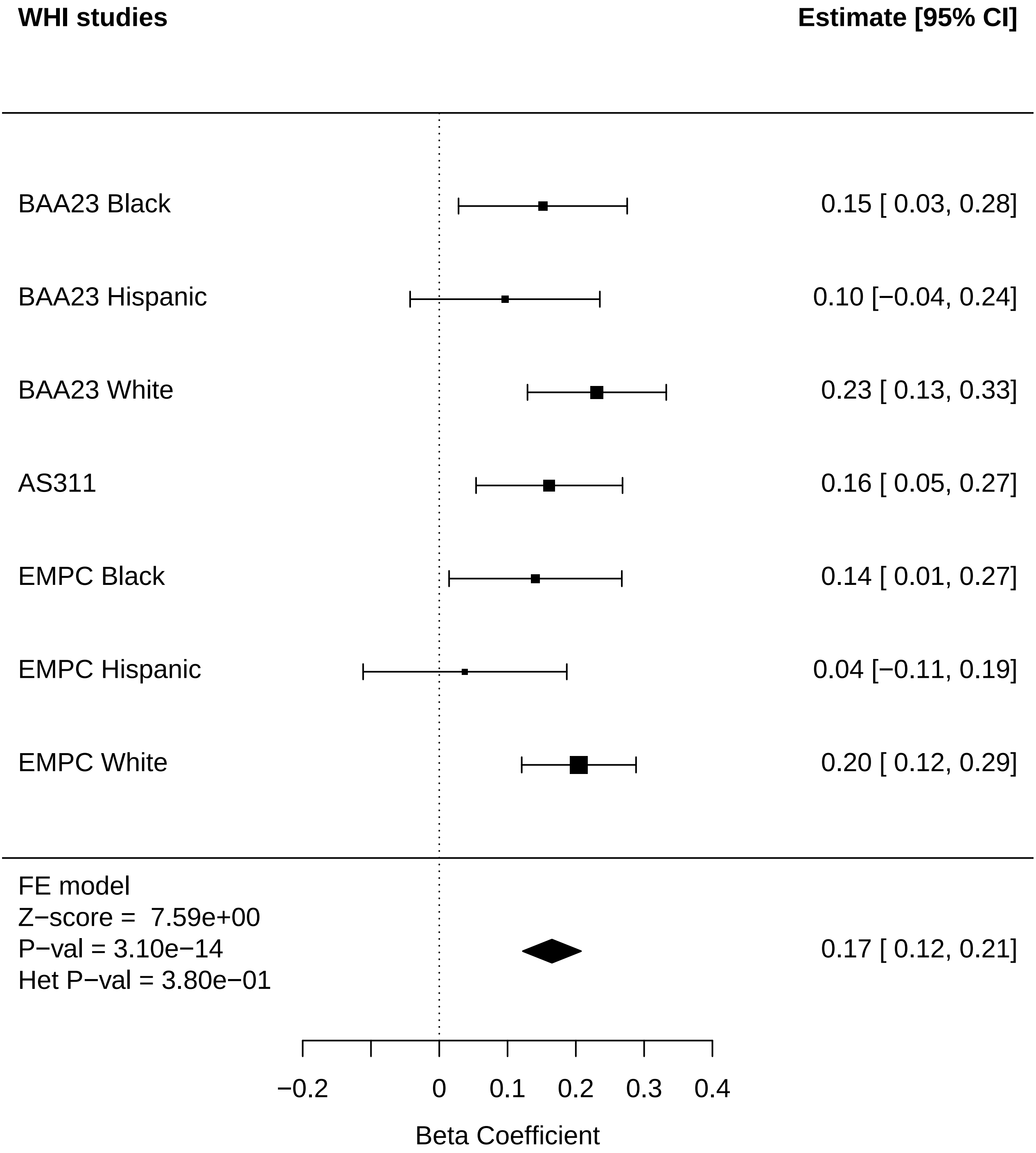

### Systems_Clock_Any Cancer.pdf

#### Systems_Clock Any Cancer

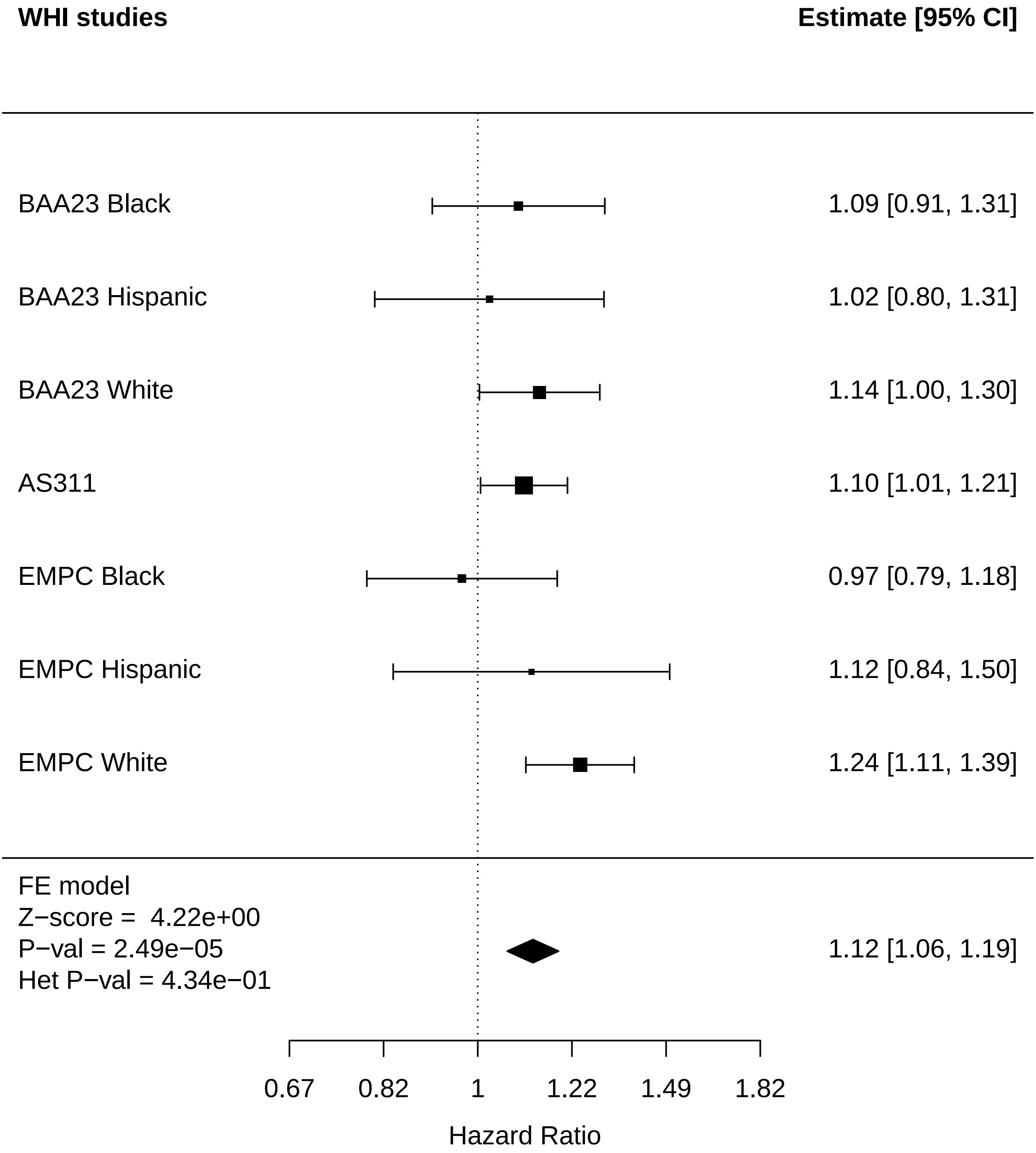

### Systems_Clock_Arthritis.pdf

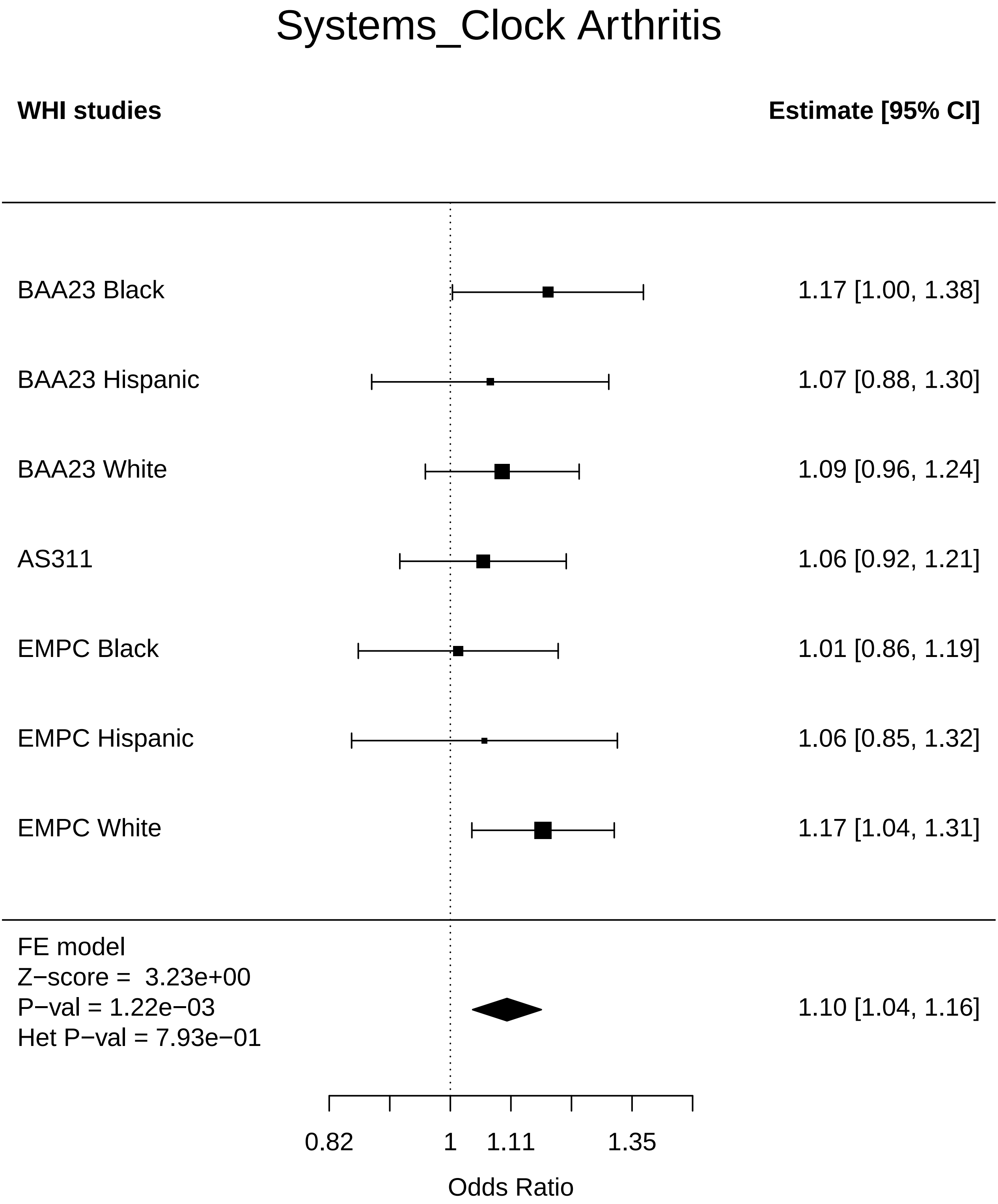

### Systems_Clock_Breast Cancer.pdf

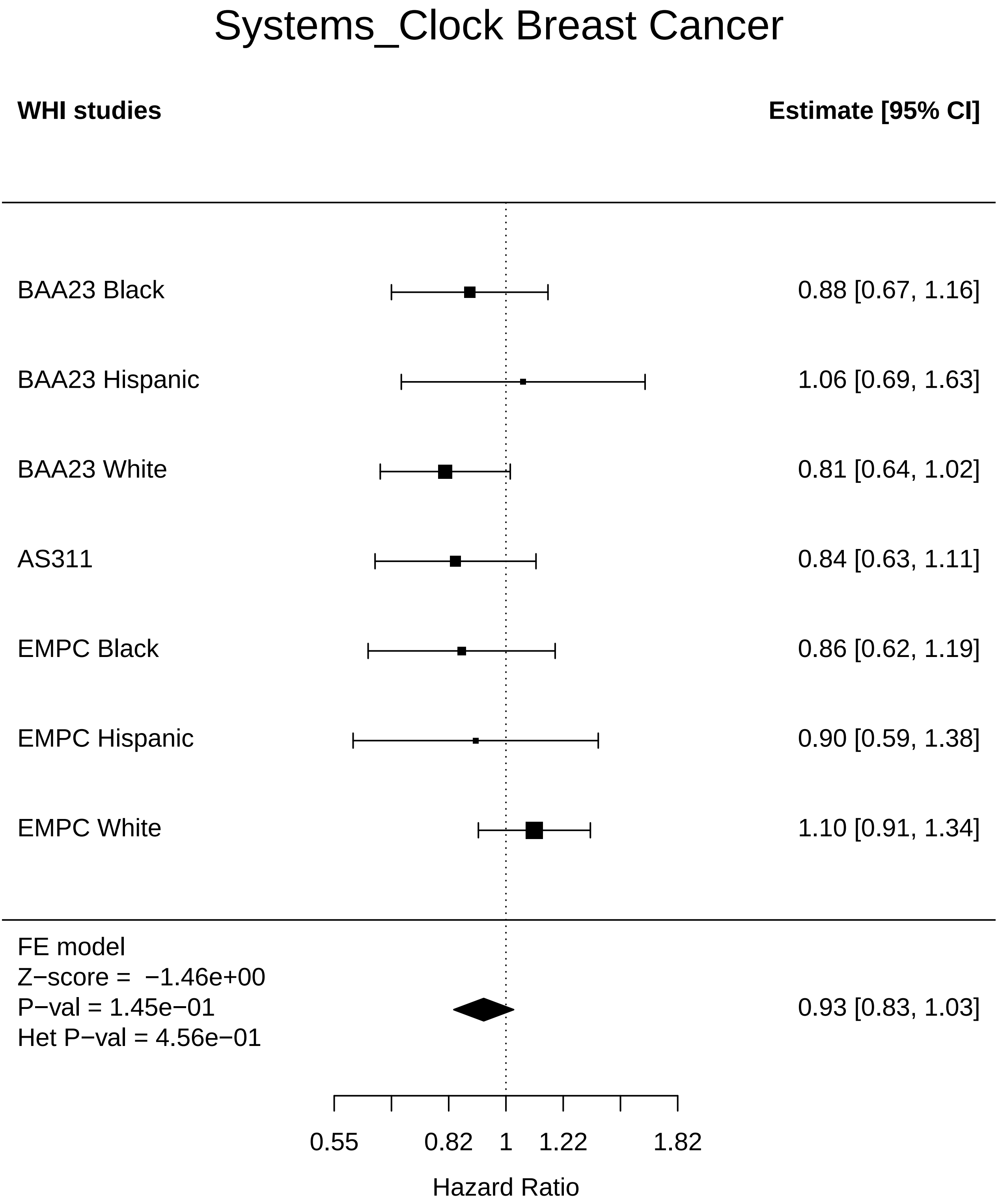

### Systems_Clock_Cataract.pdf

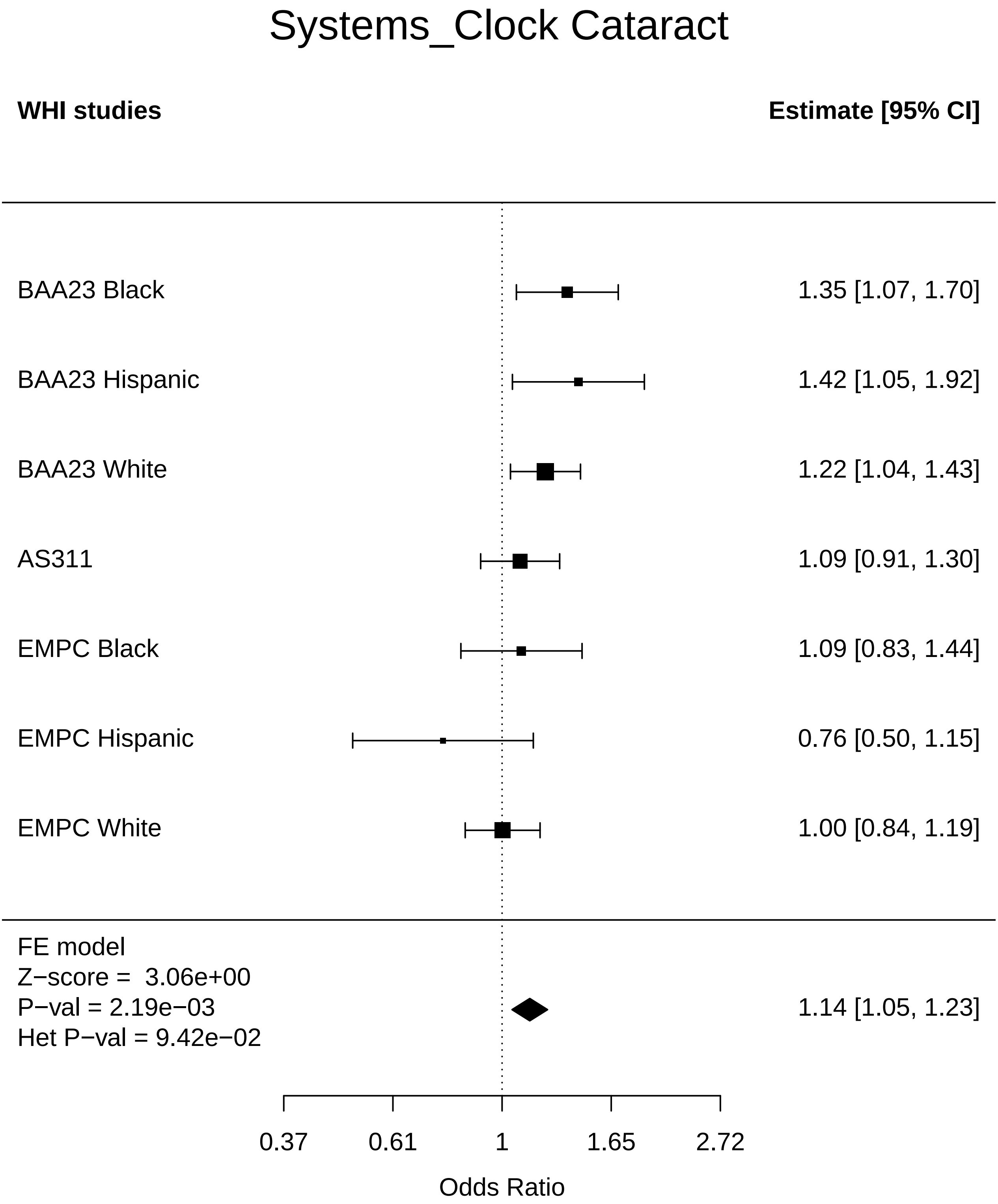

### Systems_Clock_CHD.pdf

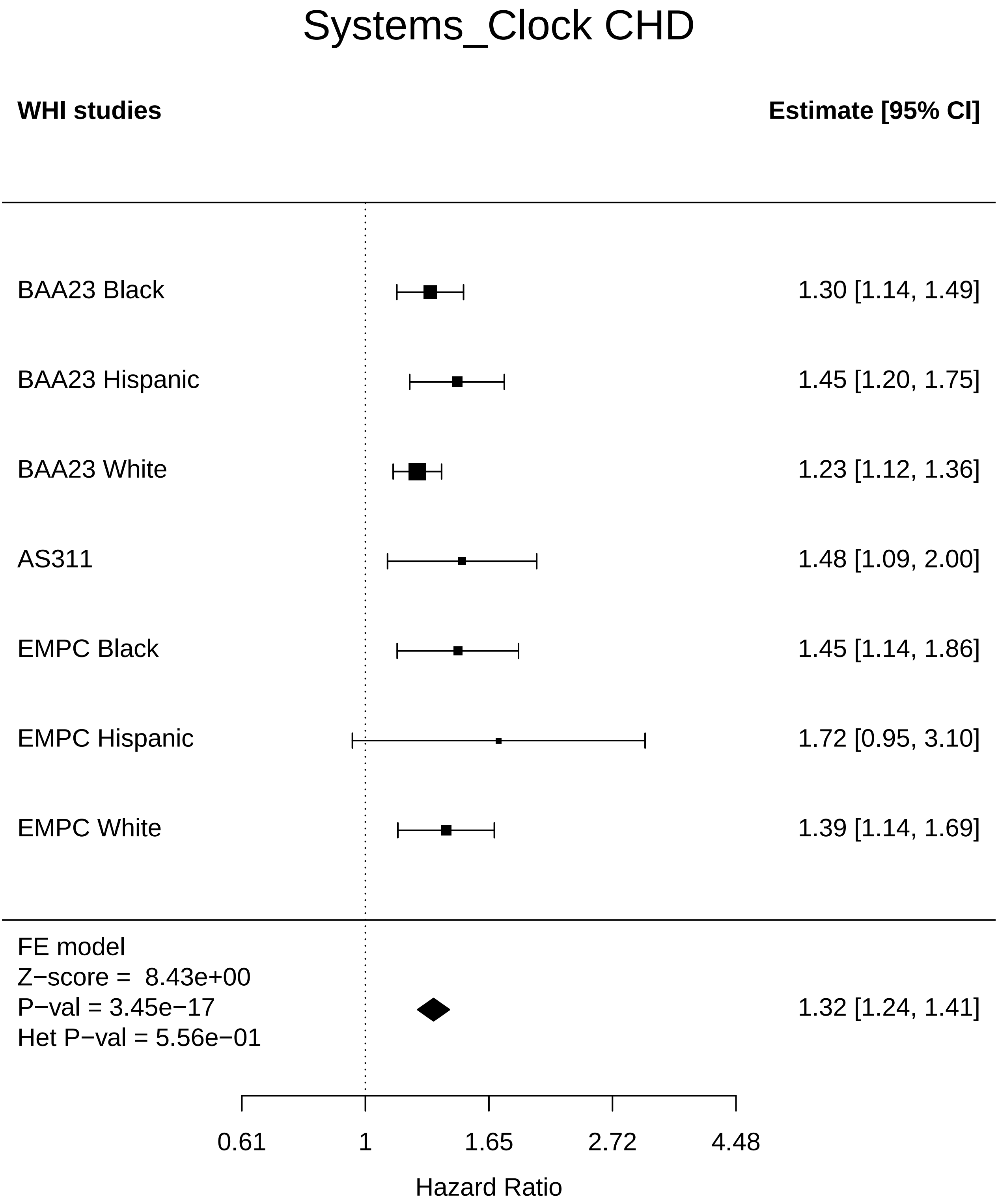

### Systems_Clock_Cognitive function.pdf

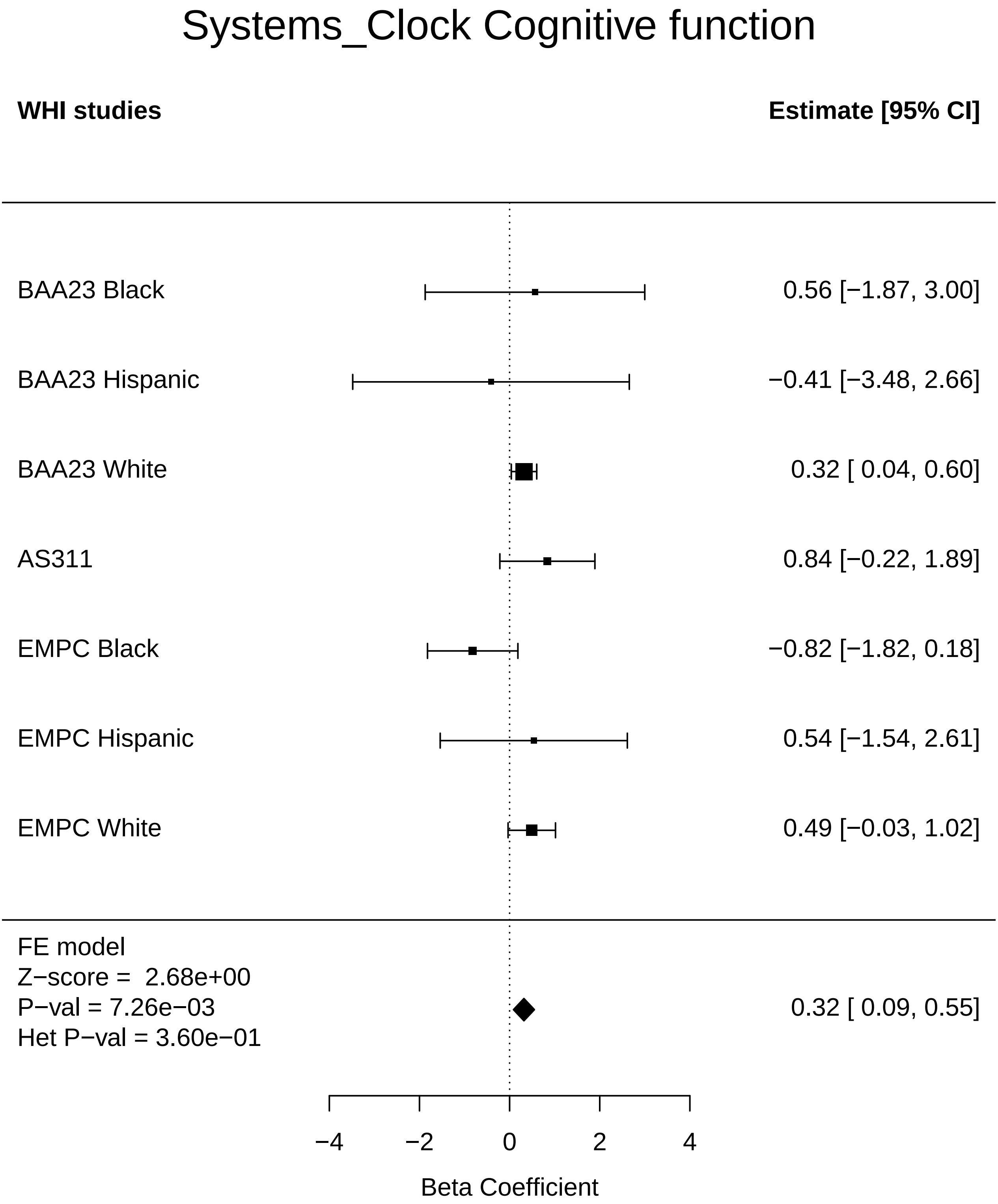

### Systems_Clock_Colorectal cancer.pdf

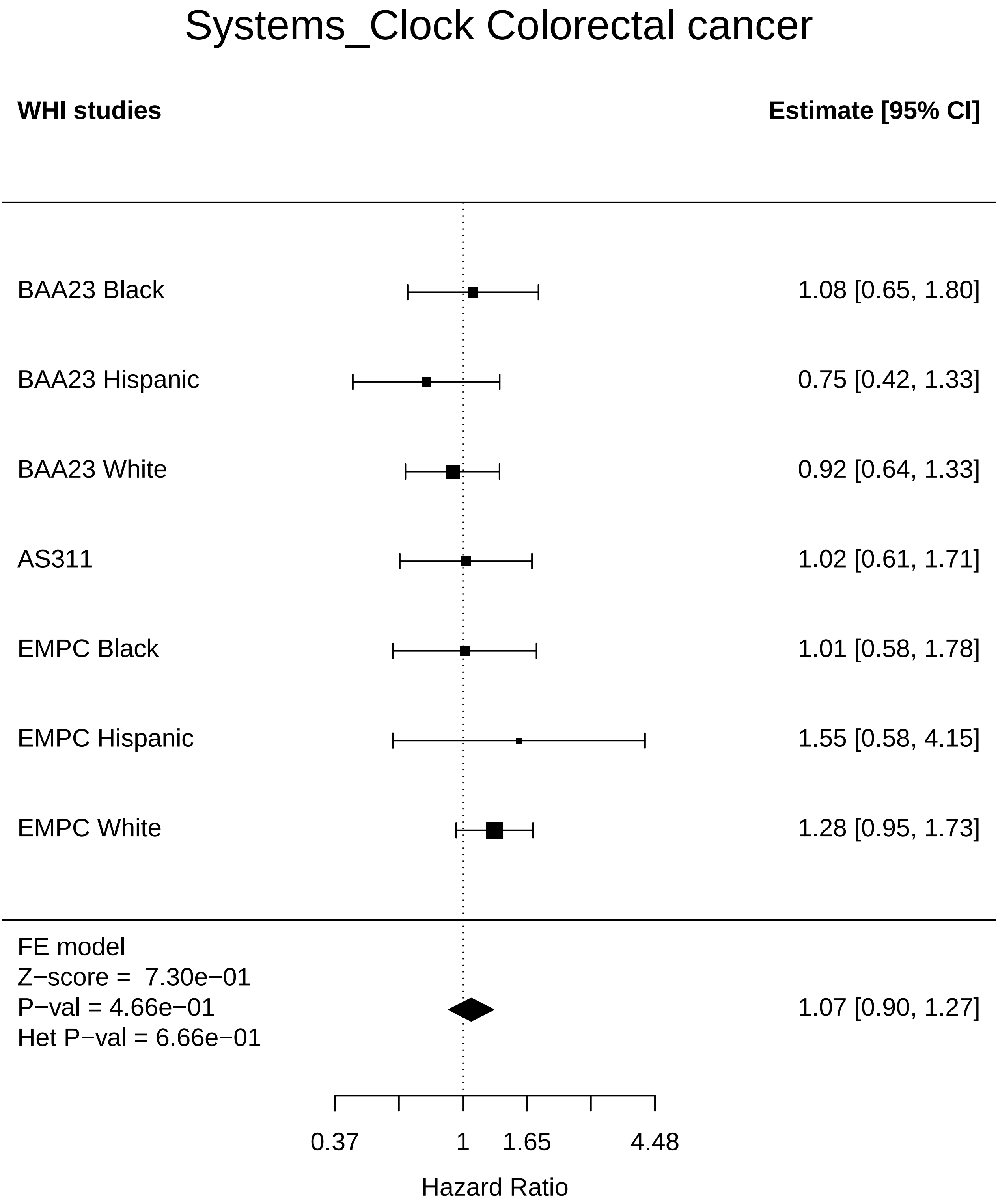

### Systems_Clock_Diabetes.pdf

#### Systems_Clock Diabetes

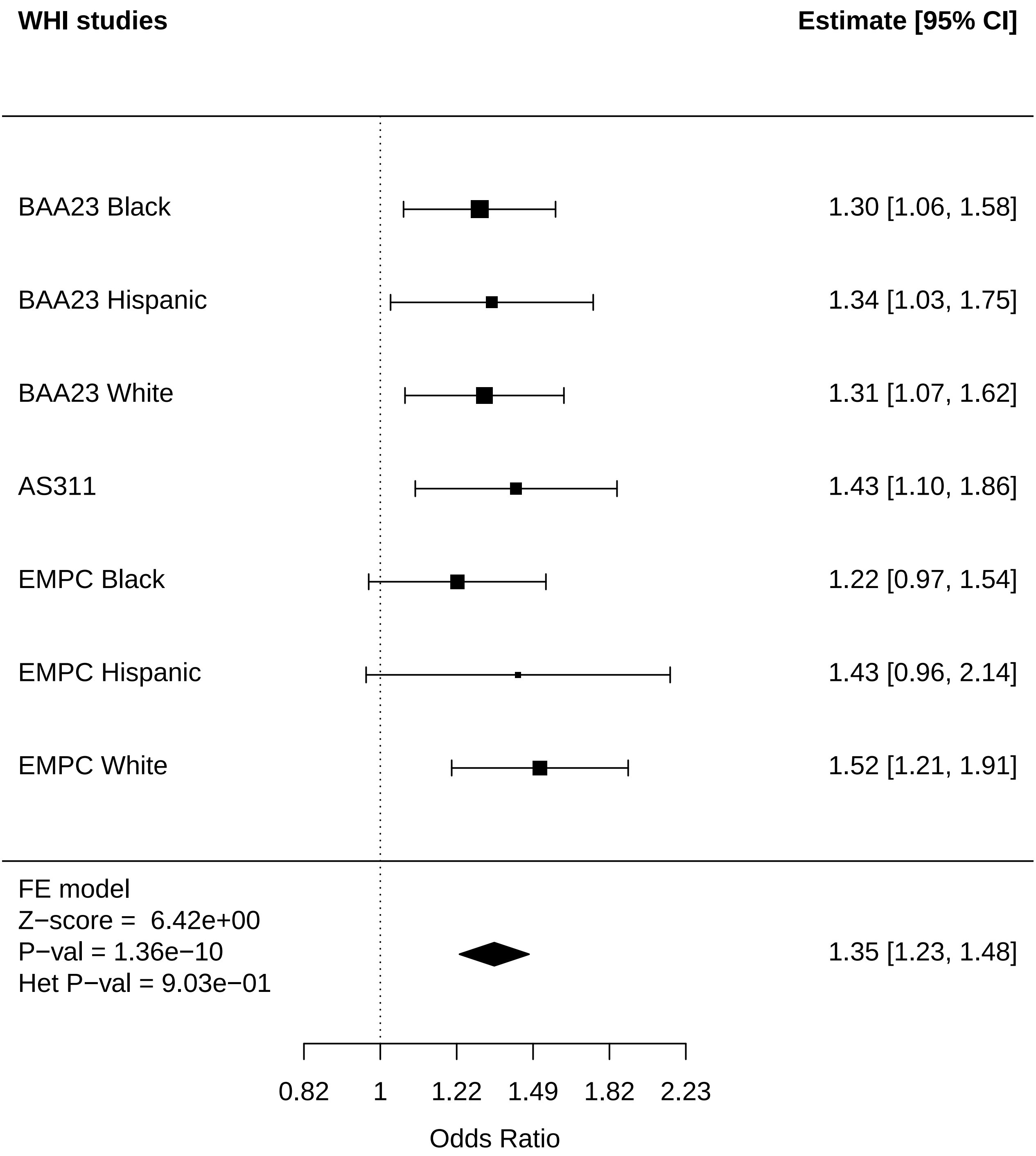

### Systems_Clock_Endometrial cancer.pdf

#### Systems_Clock Endometrial cancer

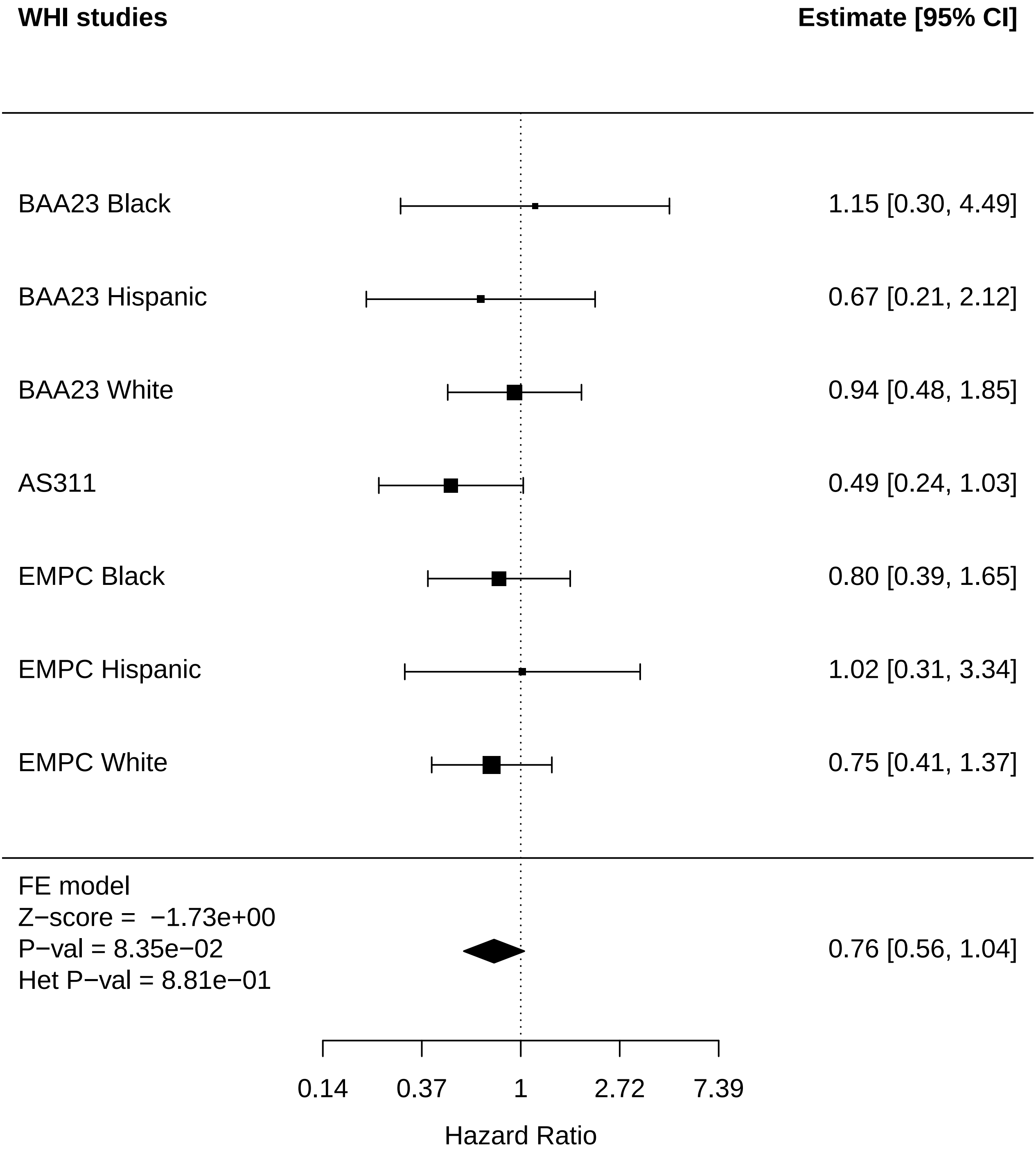

### Systems_Clock_Leukemia.pdf

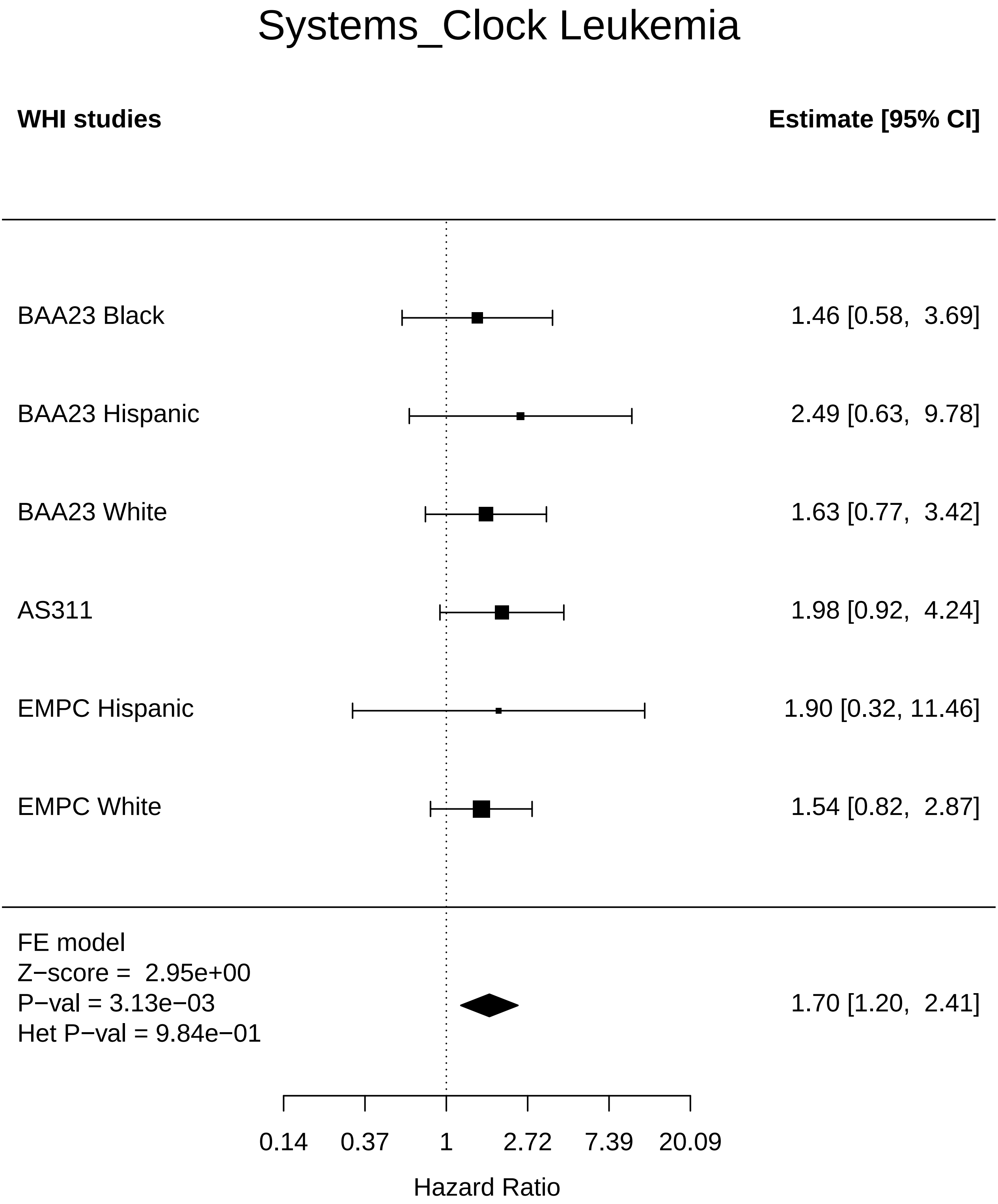

### Systems_Clock_Lung Cancer.pdf

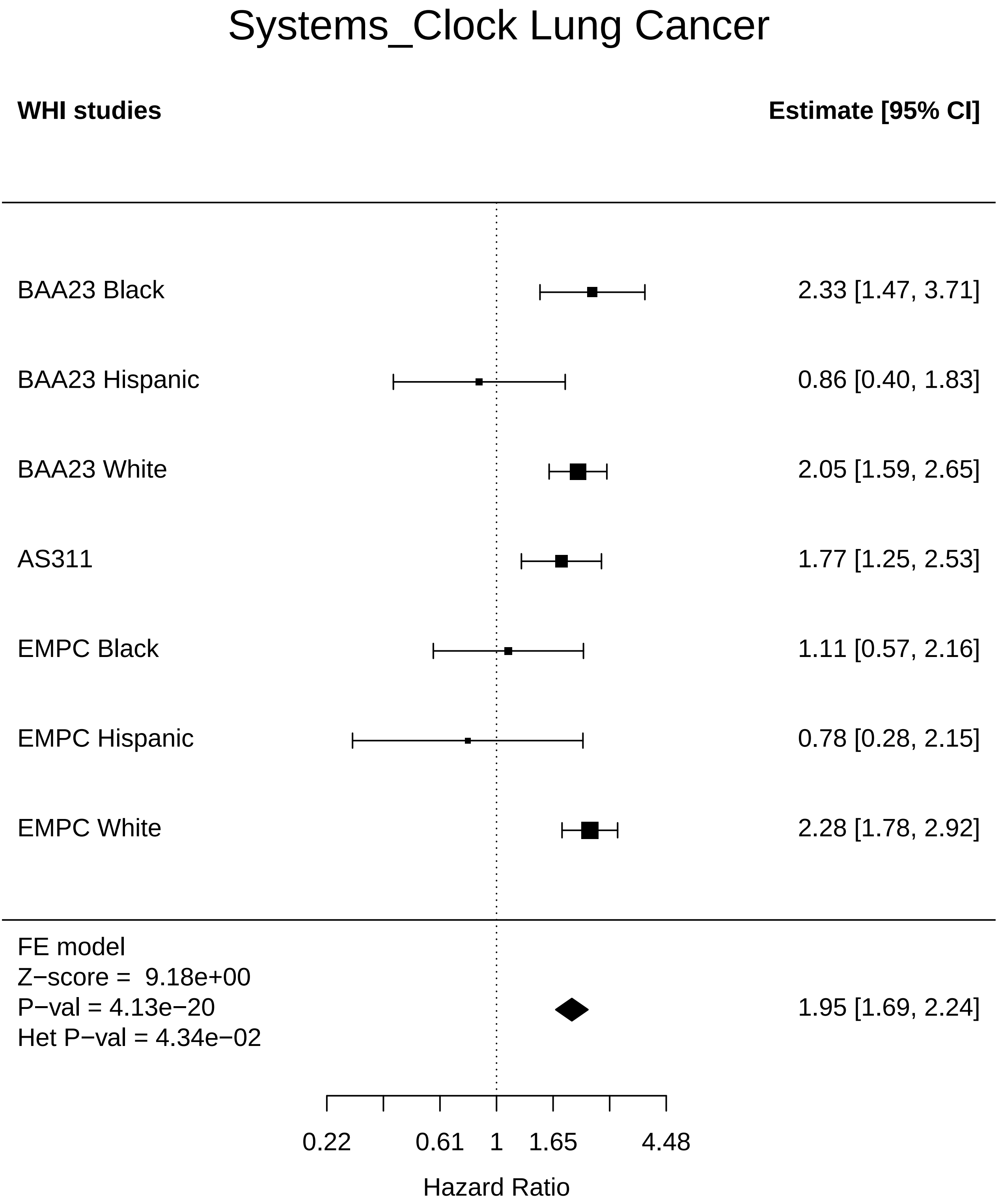

### Systems_Clock_MI.pdf

#### Systems_Clock MI

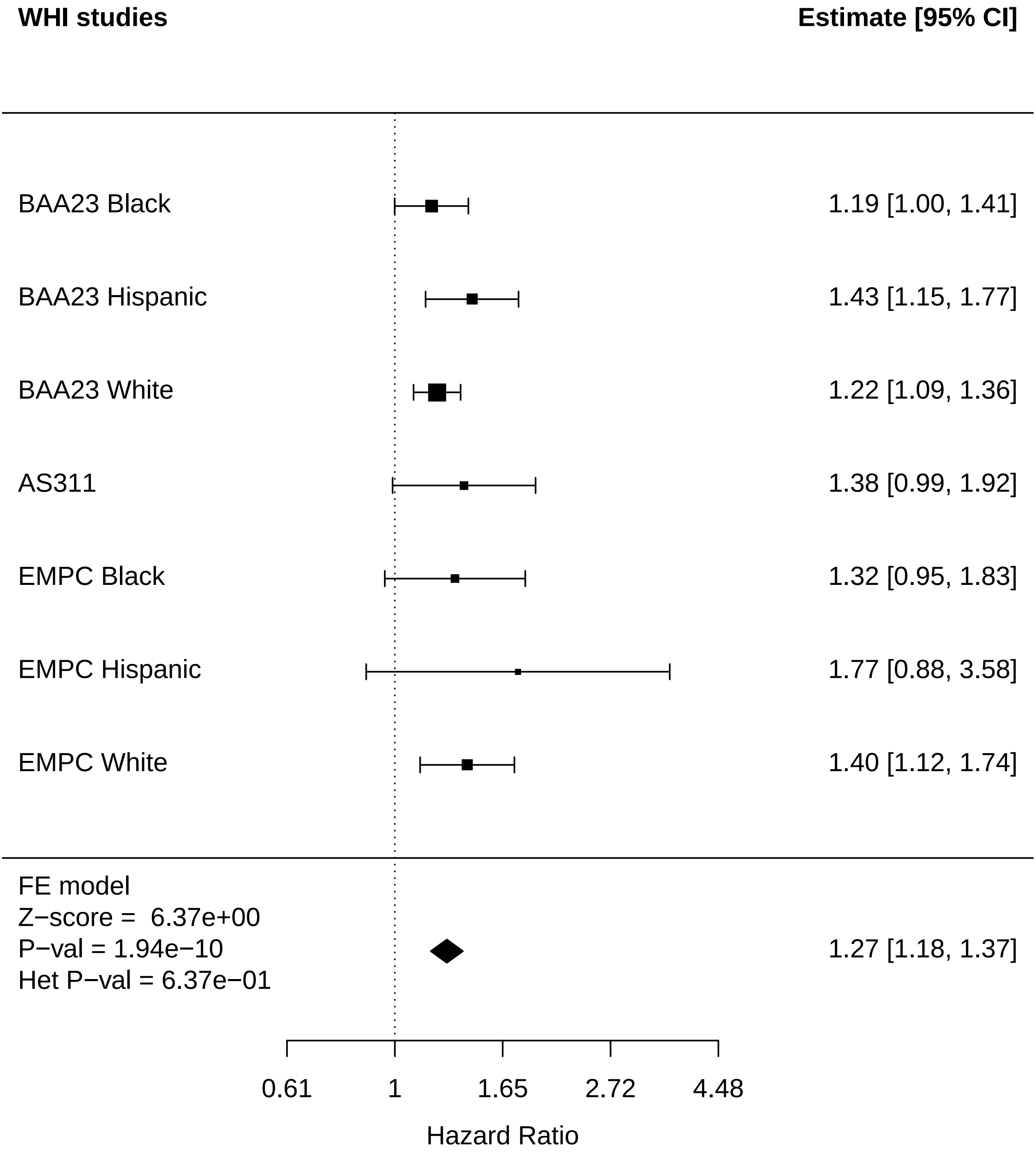

### Systems_Clock_Mortality.pdf

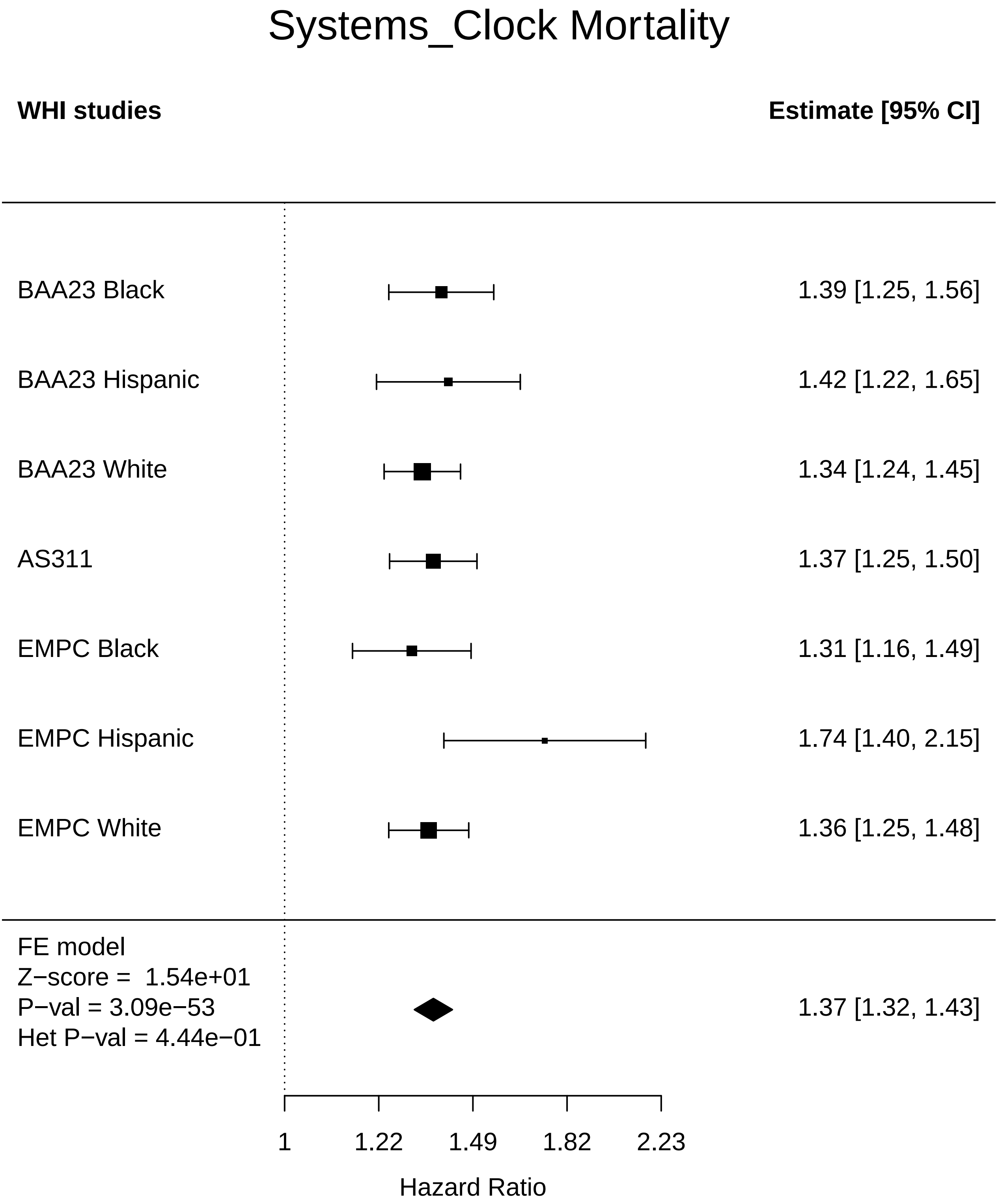

### Systems_Clock_Physical function.pdf

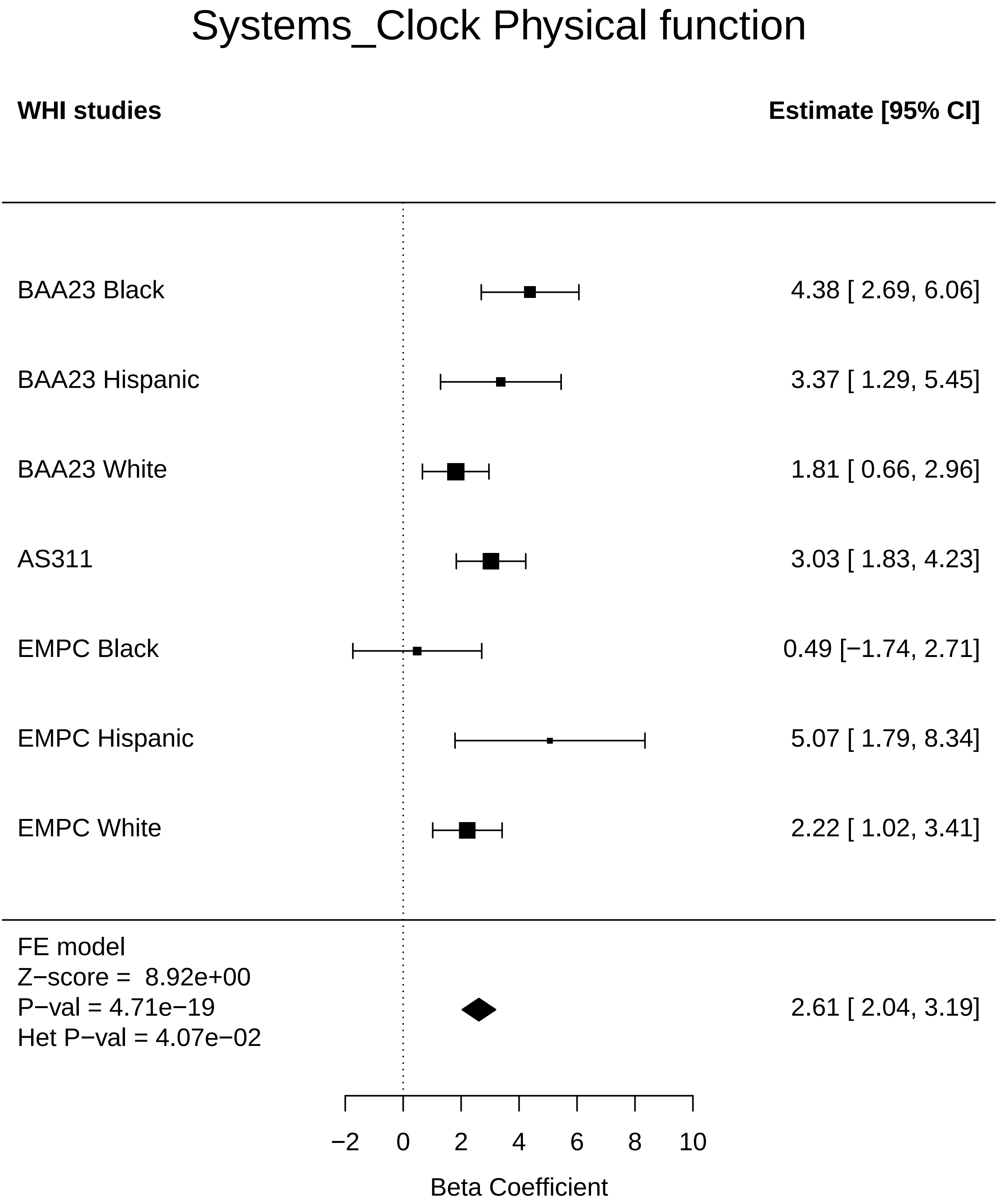

### Systems_Clock_Stroke.pdf

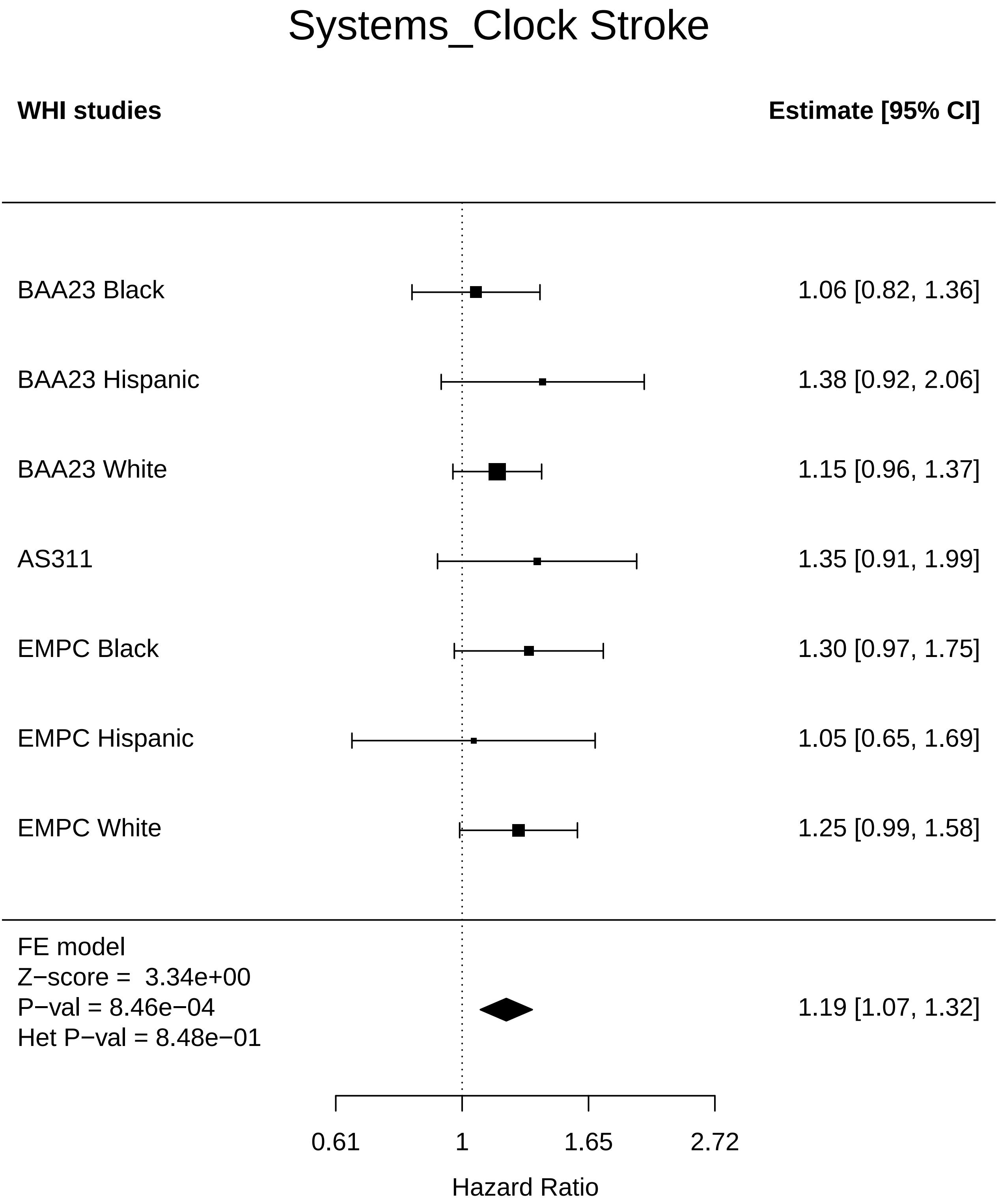

### Systems_Clock_Thyroid disease.pdf

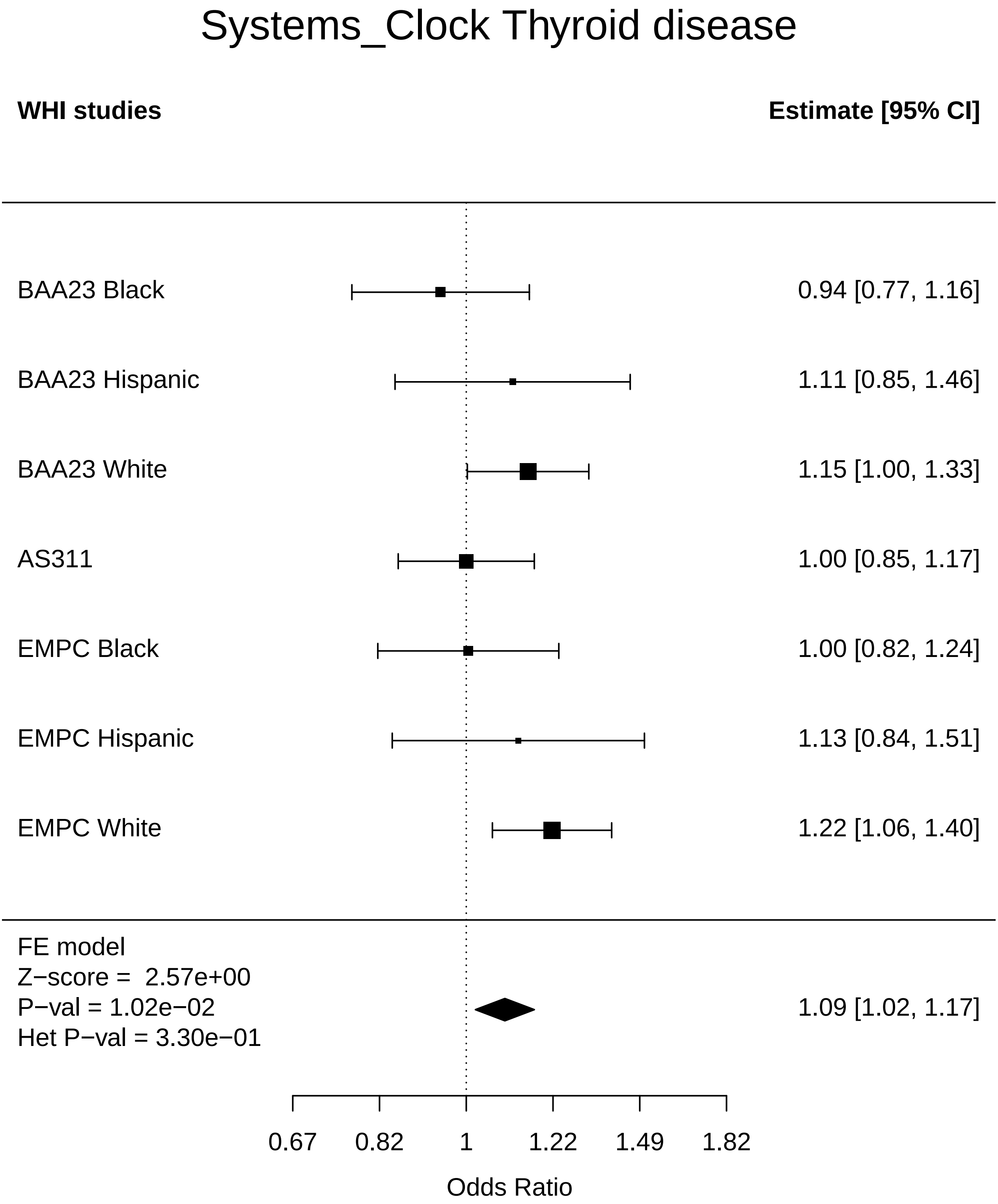

### Systems_Clock_Total comorbidities.pdf

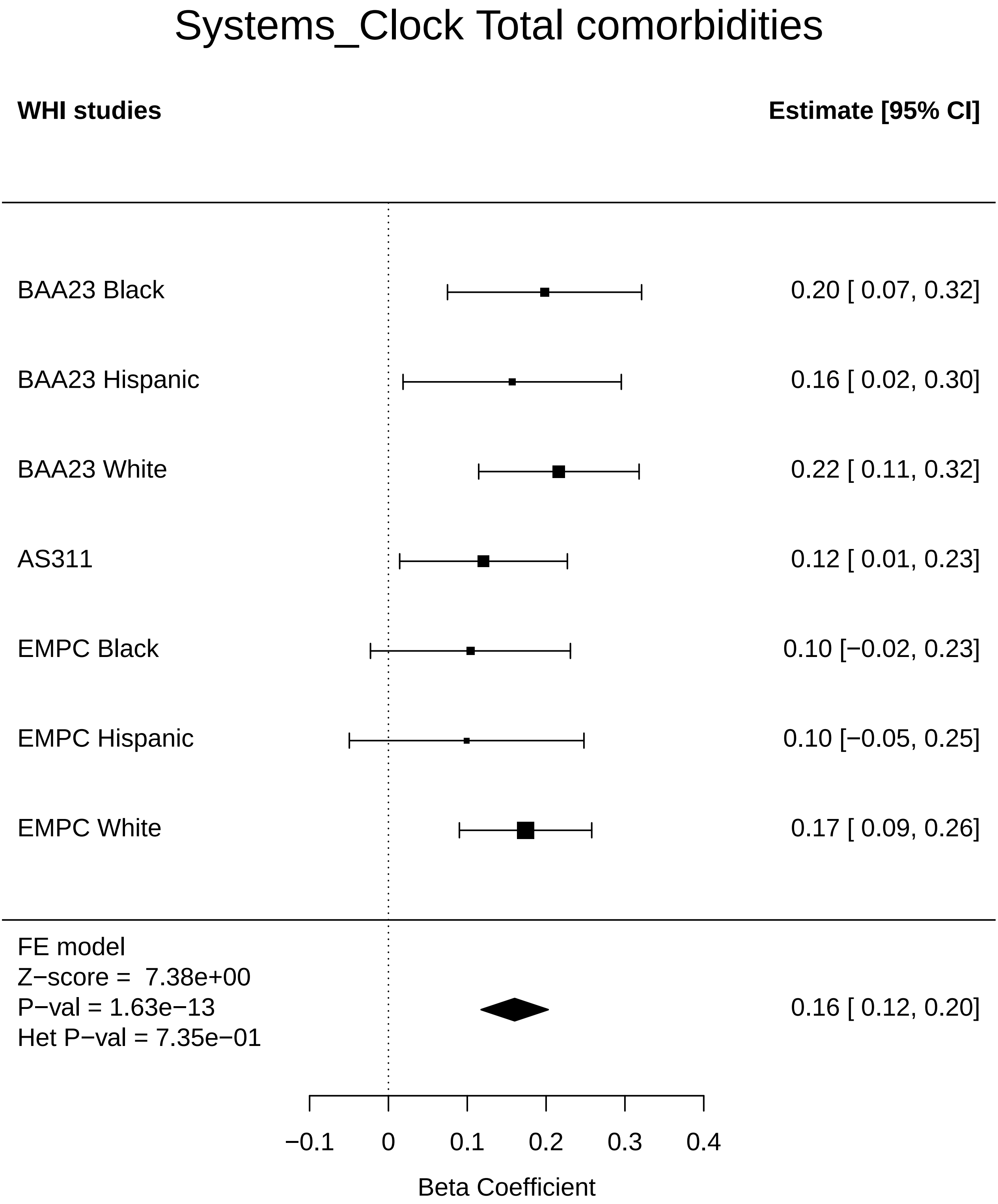
